# Postnatal persistence of sex-dependent renal developmentally programmed structural and molecular changes in nonhuman primates

**DOI:** 10.1101/2020.12.01.406355

**Authors:** Andrew C. Bishop, Kimberly D. Spradling-Reeves, Robert E. Shade, Kenneth J. Lange, Shifra Birnbaum, Kristin Favela, Edward J. Dick, Mark J. Nijland, Cun Li, Peter W. Nathanielsz, Laura A. Cox

## Abstract

**Background:** Poor nutrition during development programs kidney function. No studies on postnatal consequences of decreased perinatal nutrition exist in nonhuman primates (NHP) for translation to human renal disease. Our baboon model of moderate maternal nutrient restriction (MNR) produces intrauterine growth restricted (IUGR) and programs renal fetal phenotype. We hypothesized that the IUGR phenotype persists postnatally, influencing responses to a high-fat, high-carbohydrate, high-salt (HFCS) diet.

**Methods:** Pregnant baboons ate chow (Control; CON) or 70% of control intake (MNR) from 0.16 gestation through lactation. MNR offspring were IUGR at birth. At weaning, all offspring (CON and IUGR females and males, n=3/group) ate chow. At ~4.5 years of age, blood, urine, and kidney biopsies were collected before and after a 7-week HFCS diet challenge. Kidney function, unbiased kidney gene expression, and untargeted urine metabolomics were evaluated.

**Results:** IUGR female and male kidney transcriptome and urine metabolome differed from CON at 3.5 years, prior to HFCS. After the challenge, we observed sex-specific and fetal exposure-specific responses in urine creatinine, urine metabolites, and renal signaling pathways.

**Conclusions:** We previously showed mTOR signaling dysregulation in IUGR fetal kidneys. Before HFCS, gene expression analysis indicated that dysregulation persists postnatally in IUGR females. IUGR male offspring response to HFCS showed uncoordinated signaling pathway responses suggestive of proximal tubule injury. To our knowledge, this is the first study comparing CON and IUGR postnatal juvenile NHP and the impact of fetal and postnatal life caloric mismatch. Perinatal history needs to be taken into account when assessing renal disease risk.

## Introduction

Human epidemiological ^1–3^ and controlled maternal nutrient restriction (MNR) studies in rats, sheep and nonhuman primates (NHP) show that sub-optimal intrauterine nutrition leads to intrauterine growth restriction (IUGR), altering fetal development and predisposing to later-life chronic disease, including renal and heart disease and hypertension ^4–8^ and chronic kidney disease ^9, 10^. In 2013 nearly 35 million Americans experienced either food insecurity or hunger ^11^, including women of reproductive years. Developmental programming can be defined as *responses to challenges in critical developmental time windows that alter life course phenotype.* Poor maternal nutrition is a common programming challenge that leads to IUGR and offspring adverse life-course health ^12^.

Studies of mechanisms by which MNR during pregnancy negatively impacts offspring health have been mostly conducted in rodents ^13^ and are difficult to extrapolate to humans due to differences in rodent and primate renal developmental trajectories ^14^. Renal molecular and cellular effects of combined prenatal and postnatal stressors remain uninvestigated in NHP, the species closest to human for the study of renal programming.

We have developed a well-characterized baboon model of moderate MNR (70% of chow fed to controls, CON) during pregnancy, resulting in IUGR and fetal renal ^15–18^, cardiovascular ^19^, metabolic ^20–22^, hepatic ^23, 24^ and neural ^25^ phenotypes in female and male offspring ^18, 25, 26^. By 0.5 gestation (G), the fetal kidney showed decreased tubule length and down-regulation of genes directing kidney branching morphogenesis ^15^ and mTOR nutrient sensing ^27^. Renal programming effects persisted to 0.9 G ^17^.

We hypothesized that fetal kidney molecular and tubular changes persist postnatally and are accompanied by dysfunction in renal development. We quantified kidney molecular pathways and function in IUGR and age-matched CON baboons (4.5 years; human equivalent 13 years). We evaluated responses when IUGR kidneys are subjected to the second hit stress of postnatal dietary mismatch with a high-fat, high-carbohydrate, high-salt (HFCS) Western-style diet and high fructose drink. We monitored food and drink consumption, body weight, and urine output. We also collected blood, urine and kidney biopsies before and after the HFCS challenge to investigate molecular renal function with unbiased renal transcriptome and urine metabolome analyses. Our findings indicate that the impact of IUGR on female and male primate kidneys result in persistent renal dysfunction postnatally and response to HFCS differs by sex and fetal exposure.

## Methods

### Animal Selection and Management

All animal procedures were approved by the Institutional Animal Care and Use Committee at Texas Biomedical Research Institute and conducted in Association for Assessment and Accreditation of Laboratory Animal Care approved facilities at the Southwest National Primate Research Center. Details of housing, individual feeding cages, training, and environmental enrichment were previously published in ^28^.

Twelve non-pregnant female baboons *(Papio* species), 11.5 ± 0.51 (mean ± standard error of the mean [SEM]) years of age and of similar morphometric phenotype, were selected for study and group-housed in outdoor cages with a vasectomized male, thereby providing full social and physical activity. Animals were trained prior to pregnancy to feed in individual cages as described previously ^28^. Briefly, at feeding time all baboons passed along a chute and into individual feeding cages. Each baboon’s weight was obtained while crossing an electronic scale (GSE 665; GSE Scale Systems, Milwaukee, Wisconsin). Water was continuously available in the feeding cages via individual waterers (Lixit, Napa, California). Following the introduction of a fertile male and starting at 30 days of gestation (dG; equivalent to 0.16G), six females were randomly assigned to the CON group and fed Purina Monkey Diet 5038 (chow; Purina, St Louis, MO) *ad libitum.* The chow diet contained 12% energy from fat, 18% from protein, and 69% from carbohydrate, consisting of 0.29% glucose and 0.32% fructose. The remaining six females were assigned to the MNR group and fed 70% of the feed eaten by the CON females on a weight-adjusted basis from 0.16G through the rest of pregnancy and lactation ^15^. Water was available to all animals *ad libitum.*

CON and MNR mothers spontaneously delivered CON and IUGR offspring, respectively, at full term. The offspring were reared with their mothers in group housing until weaning at approximately nine months of age. Then they were moved to a juvenile cage of mixed males and females and maintained on the chow diet.

### HFCS Diet Challenge

At approximately 4.5 years of age (human equivalent of 13 years of age), 6 CON offspring (3 females and 3 males) and 6 age-matched IUGR offspring (3 females and 3 males) were challenged with a 7-week HFCS diet and given access to a high fructose drink in addition to water, which were all available to the animals throughout the study *ad libitum.* The HFCS diet contained 73% Purina Monkey Chow 5038 (a grain-based meal), 7% lard, 4% Crisco, 4% coconut oil, 10.5% flavored high fructose corn syrup, and1.5% water. Vitamins and mineral preparations were added to match the micronutrient composition of the chow diet. Palatability was enhanced using non-caloric artificial fruit flavors. Details have been published ^29^. The high fructose drink included water, high fructose corn syrup (2.83 Kcal/g, 76% sugar, 41.8% fructose, 34.2% dextrose, ISOSWEET 5500, Tate & Lyle, Decatur, IL), and artificial fruit flavoring ^29^.

During the HFCS challenge, all baboons from a single group cage were run once per week into individual feeding cages, passing over an electronic weighing scale. Urine was collected in a pan below each feeding cage for 3 hours. Food and drink were not available during this time to minimize contamination of urine. After the urine was collected during this 3-hour period, animals were given free access to the HFCS diet, high fructose drink, and water for 13.5 hours in the individual feeding cages. They were then returned to the group social area. The high fructose drink was provided in a Lixit waterer (Lixit, Napa, California) that was connected to a gauge to measure sugar drink consumption for each animal. Food consumption was recorded for each animal. Following the 13.5-hour consumption period, animals were returned to the group cage.

### Cardiovascular Telemetry

Due to budgetary constraints, telemetry was only measured in males. Based on other studies showing a greater impact of IUGR on male offspring ^30^, we predicted that IUGR would impact male offspring to a greater degree than IUGR female offspring and chose to use telemetry for IUGR and CON males. Two weeks prior to initiation of the HFCS challenge, the male offspring received a radio transmitter (Model PA-C40, Data Sciences International, St. Paul, MN) capable of measuring arterial blood pressure, electrocardiogram (ECG), and temperature. A 2-3 inch midline incision was made on the abdomen below the navel and above the pubis to open the peritoneum. A 1-2 inch incision was made on the interior aspect of the right hind limb above the femoral artery. The sterile body of the transmitter was placed inside the abdominal cavity and anchored via attachment holes on the implant to the ventral peritoneal wall during closure of the peritoneal incision. The pressure catheter exited adjacent to the peritoneal incision and was tunneled subcutaneously to the right hind limb incision. The femoral artery was dissected free and occluded at two points with suture material. A purse string of 3-O nonabsorbable suture material was placed on the surface of the artery and a small incision made in the center of the purse string. The catheter was introduced into the artery via the incision and the purse string used to seal the entry point. The catheter was then secured to surrounding connective tissue. ECG electrodes were also tunneled subcutaneously, one to the right shoulder and the other to the left groin. The skin incisions were closed in layers. Ketorolac was administered for pain management intramuscularly at the time of surgery (15-30mg) and SID for 2 additional days, and Cephalexin (25mg/kg) was administered twice daily for 7 days postoperatively via food.

Telemetry data were collected from the males weekly during the 13.5-hour period the animals were in the individual feeding cages. Recordings were obtained 1 week prior to initiation of the HFCS diet and during each week of the 7-week challenge. Therefore, a total of 8 weekly measurements of ~13.5 hours were made on each animal. One-minute recordings were averaged for 30-minute time periods for each collection.

### Morphometric Measurements

Morphometric measurements of each animal were collected before and after the HFCS challenge. Baboons were sedated with 10 mg/kg ketamine administered intramuscularly. To obtain the appropriate anthropometric measurements, hair was completely removed around the waist and hip circumference lines. Baboons were laid on a board with a flat surface, and body length (recumbent length) was measured using an anthropometer (cat. N101, Siber Hegner Ltd. Switzerland) from the crown of the head to the right tibia. The head was positioned firmly against the fixed board of anthropometer in the extended position. The right knee was extended, and the feet were flexed at right angles to the lower legs. Length was recorded to the nearest 0.1 centimeter. Waist circumference was measured by applying a tape measure mid-way between the lowest point of the ribs in the mid-axillary line (costal margin) (10th rib) and the iliac crest in the mid-axillary line. Hip circumference measurements were taken at the point of maximum circumference over the buttocks with a non-stretchable tape held in a horizontal plane, touching the skin, but not indenting the soft tissue. Waist depth (anterior-posterior abdominal distance) was measured with the anthropometer from the plane of the back to the anterior point of the abdomen at the level of navel. Body Mass Index (BMI) was calculated by dividing the weight in kilograms by the crown-rump length in cm^2^.

### Blood and Kidney Collections

Blood and kidney tissue samples were collected before and after the HFCS challenge at the same time as morphometric data were collected. Following an overnight fast, animals were premedicated with ketamine and then anesthetized with isoflurane (1.5% v/v, inhalation), and percutaneous venipuncture was performed on the femoral vein just caudal to the femoral triangle for blood sample collections. Percutaneous renal punch biopsies were also collected while the animals were anesthetized. The biopsy area was aseptically prepared, and the site of incision was locally infiltrated with lidocaine. The biopsies were then collected using ultrasound imaging to ensure consistent sampling among animals; kidney biopsies were frozen in liquid nitrogen and stored at −80°C until use. Specificity of kidney biopsy collections was determined based on gene expression specific for renal tubule segments ^31^.

### Urine Measures

Renal function was assessed using urine samples collected each week of the study while the animals were housed in individual feeding cages. Urine volume, urine creatinine excretion, and urinary sodium and potassium measurements were obtained for each sample. Urine volume was measured from each weekly overnight collection. Urine creatinine was measured using a Creatinine (urinary) Colorimetric Assay Kit (Item # 500701; Cayman Chemical Co. Ann Arbor, MI). Urinary sodium and potassium concentrations were measured by flame photometry (Corning model 450) and used to calculate sodium and potassium excretion, respectively, for the 3 hour urine collection period.

### Blood Measures

Serum creatinine was quantified with a Creatinine (serum) Assay Kit (Item # 700460, Cayman Chemical Co.), and cortisol concentration was measured by Immulite 1000 immunoassay kit (Item # 914038; Siemens Medical Solutions Diagnostics, Los Angeles, CA). Serum aldosterone excretion rates were measured by radioimmunoassay (Diagnostic Products, Los Angeles, CA) ^32^.

### RNA Isolation and Microarray Hybridization

Total RNA was isolated from an approximately 5 mg section of each frozen kidney biopsy using a Power Gen Homogenizer (Omni International, Wilmington, DE) and TRI Reagent™ Solution (Invitrogen™, Carlsbad, CA) as previously described in ^16^. RNA integrity was determined using an Agilent 2100 Bioanalyzer (Agilent Technologies Inc., Santa Clara, CA), and RNA concentration was determined by UV-Vis spectrophotometry using a Nanodrop™ 2000 (Thermo Fisher Scientific, Wilmington, DE). Complementary RNA (cRNA) was synthesized and biotinylated using the Illumina™ TotalPrep™-96 RNA Amplification Kit (Illumina Inc., San Diego, CA). cRNA samples were then hybridized to HumanHT-12 v3 Expression BeadChips (Illumina, Inc.).

### Microarray Data Analysis

Gene expression data were extracted and log2-transformed using GenomeStudio software (Illumina, Inc.) and subsequently analyzed using Partek®Genomics Suite (Partek®, St. Louis, MO). Signal intensities were all-median normalized, and differentially expressed genes were identified by Analysis of Variance (ANOVA; p < 0.05). Differentially expressed genes were overlaid onto canonical pathways and networks generated using differentially expressed genes in the dataset and the Ingenuity Pathway Analysis (IPA; QIAGEN Inc., https://www.qiagenbioinformatics.com/products/ingenuity-pathway-analysis) Knowledge Base. Right-tailed Fisher’s exact test was used to calculate a significant enrichment of differentially expressed genes in pathways, p< 0.01 ^33^. Networks were built using the IPA Knowledge Base, requiring direct connections between molecules based on experimental evidence.

We used an end-of-pathway gene expression approach to identify coordinated pathways. We hypothesized that a pathway may only be relevant to the kidney phenotype if gene expression profiles after the pathway were consistent with the overall pathway change. Therefore, pathways meeting this criterion, as well as those downstream of these pathways, were investigated. If there were no differentially expressed genes after a pathway, that pathway was not considered relevant to the phenotype ^16^.

We performed upstream regulatory network analysis to identify potential causal networks which integrate previously observed cause-effect relationships, leveraging experimental knowledge about the direction of effects to infer upstream regulatory molecules and potential mechanisms explaining observed gene expression changes. The analysis includes prediction of activation or inhibition of the network and consistency of molecules in the network with the predicted activation state. The calculated z-score makes predictions about regulation directions and infers the activation state of a putative regulator (i.e., activated or inhibited) and can be used to determine likely regulators based on statistical significance of the pattern match. The statistical models are provided in detail in Kramer et al., 2014 ^34^. Networks were considered significant for p < 0.05.

### Urine Processing and GC×GC-TOF MS Analysis

Urine metabolomics was performed on all study samples described above except for a single baseline sample for one IUGR male due to technical issues. Urine samples were processed following previously published methodology ^35^. Briefly, 10 mL urine aliquots were acidified to pH 2 with sulfuric acid. Metabolites were extracted with 2mL of methylene chloride followed by a 30-second vortex. Samples were centrifuged, and the organic phase was transferred to a new vial. 200 μL of the organic phase was transferred into sample vials containing sodium sulfate. For derivatization, 30μL of dry pyridine and 200 μL of N,O-bis(Trimethylsilyl)-trifluoroacetamide (BSTFA) was sequentially added to the sample vial followed by incubation at 60°C for 60 minutes.

Derivatized urine samples and reagent blank samples were injected onto a Pegasus 4D GC×GC-TOF MS system equipped with an Agilent 7890 gas chromatograph (Agilent Technologies) in line with a LECO Time-of-Flight mass spectrometer (LECO Corp, St Joseph, MI). The primary column was a 30 m x 0.25 mm i.d. x 0.25μm *d_f_* Rtx-1MS (Restek Corp., Belefonte, PA) and the secondary column a 2m x 0.18 mm i.d. x 0.18 μm *d_f_* Rxi-17M (Restek Corp.). Carrier gas was helium, and runs were performed in splitless mode. The initial temperature of the GC oven was 40°C, held for 1 minute, followed by ramping to 300°C at a rate of 5°C/minute, followed by a 5-minute hold. Secondary oven temperature offset was +5°C to the primary oven, and modulator temperature offset was set to +20°C of the secondary oven. The thermal modulation periods were set to 5 seconds, hot pulse time 0.6 seconds and cool time between stages 1.9 seconds. Column flow was set to 1.0 mL/minute, and the total run time was set to 34.80 minutes. The transfer line was set to 300°C and ion source 225°C. Acquisition rate was set to 100 spectra/second with a 500-second delay and mass range of 40-500 amu.

### Metabolomics Data Cleaning and Analysis

MS data were generated and processed using Chromatof software v. 4.72 (LECO Corp), which included baseline correction, peak deconvolution, peak calling and spectral library matching. Peak identification was assigned at a MS2 level consistent with spectra and fragmentation ions or accurate masses with proposed structure, as previously described ^36^. Only spectral match scores equal or greater than 70% (forward score) to the NIST2011 library were further considered. R-package, R2DGC ^37^, was used to align identified peaks by both retention times and unique mass. Alignments were performed both separately and together for baseline and challenge across all samples.

Each alignment was exported to Excel for further processing. Alignments were first filtered for metabolites present in at least 50% of the samples. Next the alignments were manually filtered for GC column and system contaminants. Lastly, the alignments were again manually filtered to remove compounds that were identified as background noise. The remaining compounds were normalized by the sum peak height of all annotated compounds remaining. Putative compounds were assessed based on present/absent between sex and treatment group. Normalized peak intensities that were present in all 3 samples of a group (e.g., all 3 males or all 3 females on HFCS diet, etc.) were compared by two-tailed t-test.

Identifiers from the Human Metabolome Database (HMDB) ^38^ and Kyoto Encyclopedia of Genes and Genomes (KEGG) ^39^ were inputted into MetaboloAnalyst 4.0 ^38^ to determine common pathways present in the urine metabolomics analysis.

### Statistical Analyses

Principal Component Analysis (PCA) and hierarchical clustering in Partek®Genomics Suite identified sex as the greatest source of variation in each dataset; therefore, therefore, we divided the data into female and male sets for statistical analysis. Weight-adjusted fructose consumption was calculated by dividing the 7-week average fructose consumption by the 7-week average body weight. In addition, the percent change for weight and BMI, comparing the end of challenge to baseline, were calculated by dividing the end of challenge value by the baseline value, multiplying by 100 and subtracting 100. Pairwise comparisons were performed using two-tailed t-tests. Data from each diet group were analyzed independently for pre- and post-challenge means and then evaluated using two-tailed t-test and two-way ANOVA with significance p < 0.05 and marginal significance 0.05 < p < 0.1.

## Results

### Kidney Biopsy Specificity

Transcriptome analysis of mouse kidney microdissected renal tubule segments provided 626 renal nepron-specific genes ^31^; 447 of these genes were expressed in our baboon renal biopsy samples (Supplemental Table 1).

### Sex Differences for Chow and HFCS

Kidney transcriptome analysis by PCA showed sex differences for both chow and HFCS where sex accounted for 53.0% of total variation for chow (Figure 1A) and 49.0% of total variation for HFCS (Figure 1B). Analysis of urine metabolome by PCA also revealed group clustering by sex, explaining 68.6% of total variation for chow (Figure 2A) and 71.1% of total variation for HFCS (Figure 2B).

**Figure 1:**
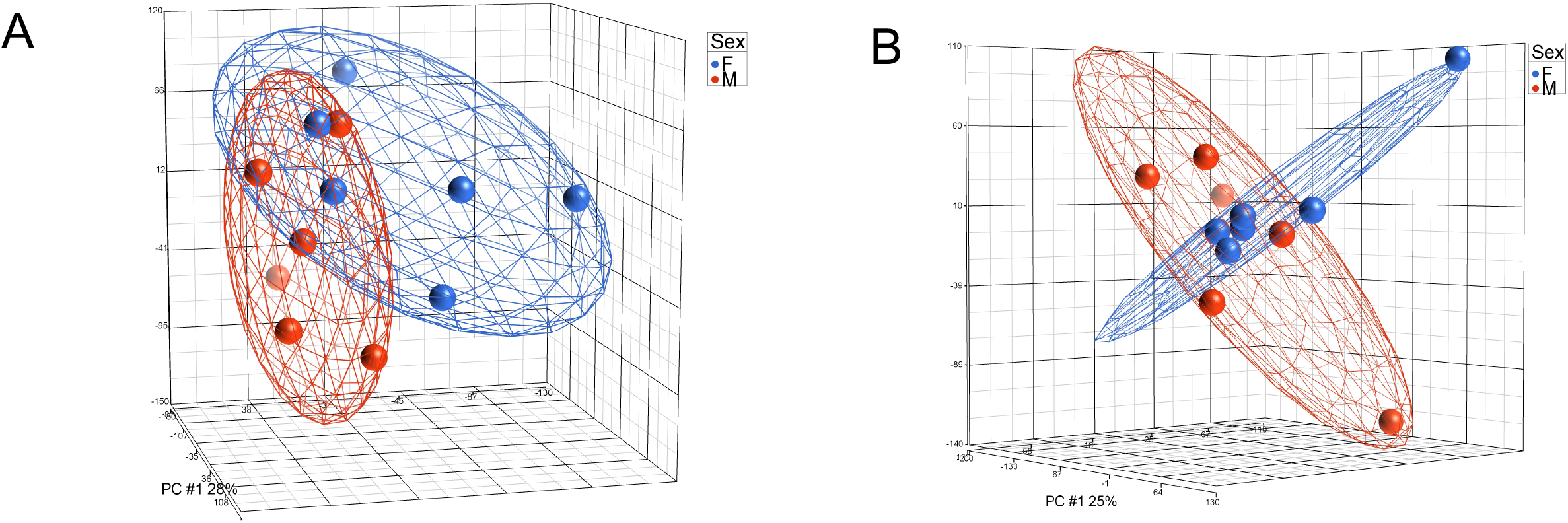
Principal component analysis (PCA) of sex differences within the renal transcriptome. PCA reveals sex differences in renal transcripts on both chow diet (A) and HFCS diet (B). Blue denotes females, red denotes males.

**Figure 2:**
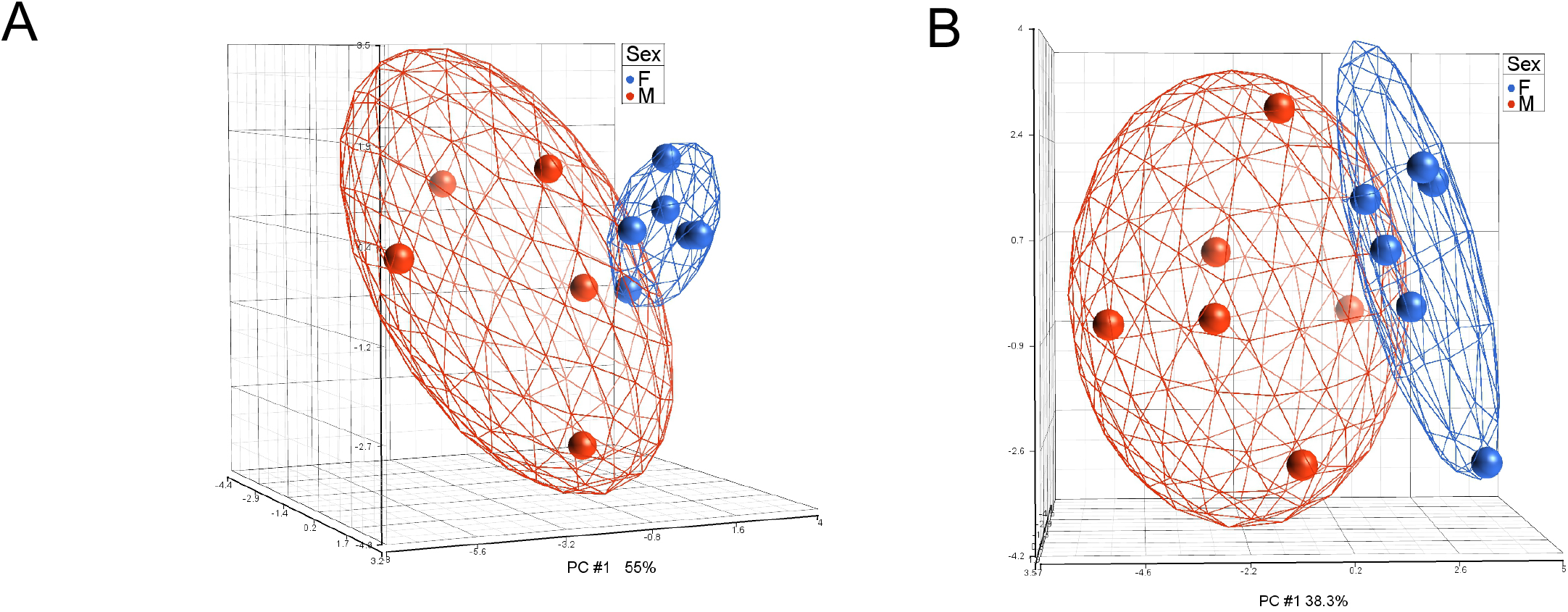
Principal component analysis (PCA) of sex differences within the urine metabolome. PCA reveals sex differences in urine metabolites signature on chow diet (A) and HFCS diet (B). Blue denotes females, red denotes males.

### IUGR vs CON Maintained on Chow

#### Morphometrics and Renal Function

At baseline, age-matched CON and IUGR female offspring were similar height, but IUGR weighed less and had lower BMI than CON (Table 1). Urine volume and urine creatinine were greater in IUGR than CON females (Table 2), while urine potassium (Table 2) and serum creatinine (Table 3) were less. Serum aldosterone and cortisol were similar between groups (Table 3). In males at baseline, age-matched CON and IUGR offspring were similar height and weight, but IUGR had lower BMI than CON (Table 1). Urine volume, urine potassium, serum creatinine, serum aldosterone, or serum cortisol were similar between IUGR and CON males (Tables 2 and 3); in addition, no differences were found for systolic, diastolic and mean arterial blood pressure, and heart rate (Table 4).

**Table 1.** Morphometrics

|  |  | Height (cm) |  |  | Weight (Kg) |  |  | Body Mass Index |  |  |
| --- | --- | --- | --- | --- | --- | --- | --- | --- | --- | --- |
|  |  | Mean | SEM | p-val | Mean | SEM | p-val | Mean | SEM | p-val |
| F CON | Chow | 87.8 | 3.35 |  | 12.9 | 0.61 |  | 16.70 | 0.49 |  |
| F IUGR |  | 87.0 | 3.21 | 0.87 | 9.9 | 0.95 | **0.058** | 13.02 | 0.44 | **0.005** |
| M CON |  | 90.1 | 0.46 |  | 15.0 | 0.85 |  | 18.45 | 1.16 |  |
| M IUGR |  | 89.9 | 6.15 | 0.98 | 12.0 | 1.86 | 0.22 | 14.68 | 0.63 | **0.046** |
| F CON | HFCS | 92.3 | 0.67 |  | 13.3 | 0.09 |  | 15.66 | 0.28 |  |
| F IUGR |  | 90.8 | 2.13 | 0.55 | 10.6 | 0.66 | **0.015** | 12.86 | 0.19 | **0.001** |
| M CON |  | 103.5 | 3.21 |  | 16.2 | 1.31 |  | 15.03 | 0.37 |  |
| M IUGR |  | 96.7 | 4.26 | 0.27 | 14.7 | 1.97 | 0.56 | 15.53 | 0.82 | 0.60 |
| F CON | Diff | 4.47 | 2.68 |  | 0.47 | 0.64 |  | -1.05 | 0.21 |  |
| F IUGR |  | 3.83 | 1.83 | 0.85 | 0.73 | 0.38 | 0.74 | -0.16 | 0.26 | **0.058** |
| M CON |  | 13.4 | 3.53 |  | 1.20 | 0.46 |  | -3.42 | 0.89 |  |
| M IUGR |  | 6.77 | 1.90 | 0.17 | 2.67 | 0.12 | **0.036** | 0.85 | 0.48 | **0.013** |

**Table 2.**
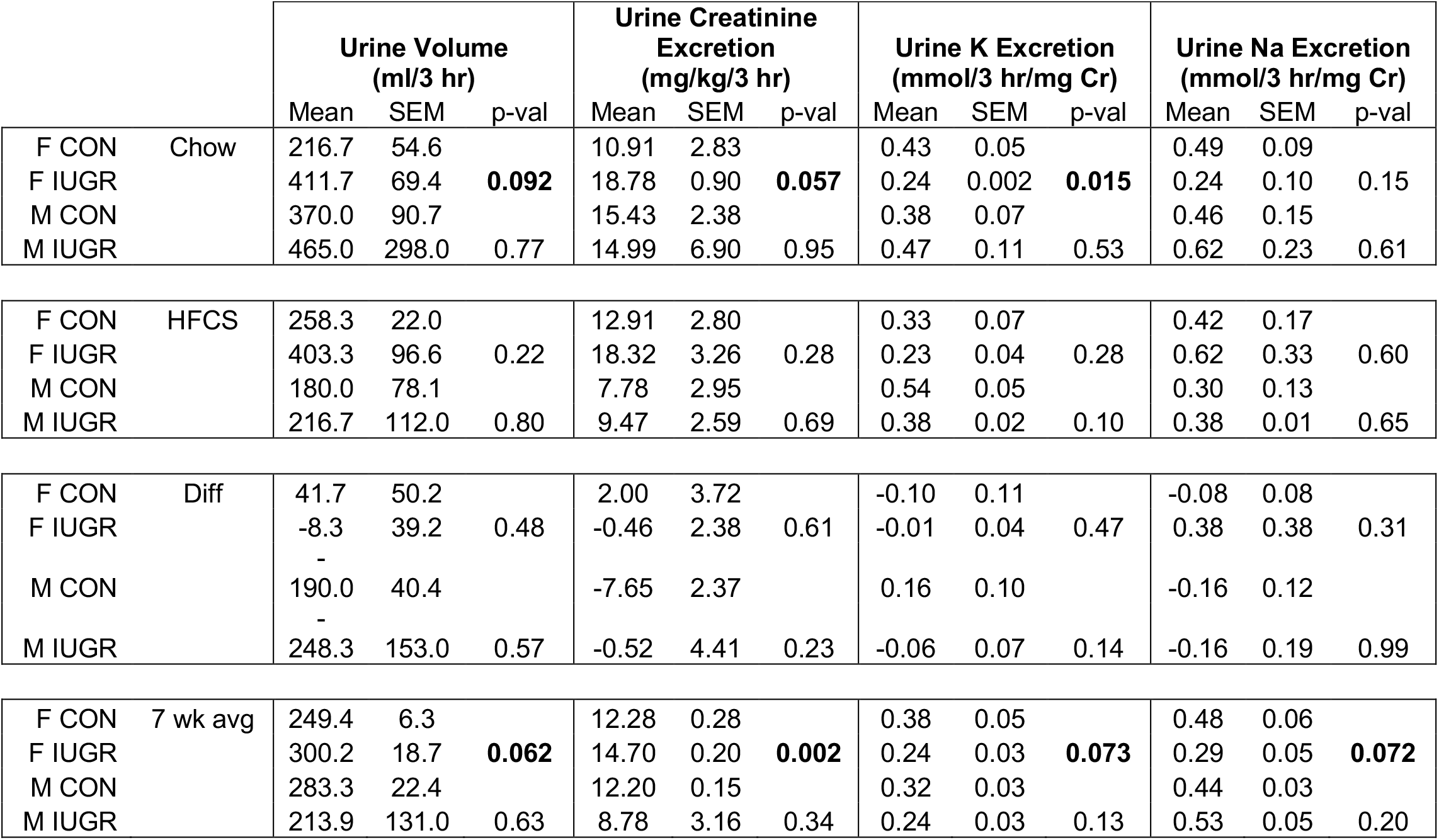
Urine Measures of Kidney Function

**Table 3.** Blood Measures of Kidney Function

|  |  | Serum creatinine (mg/ml) |  |  | Aldosterone (% bound) |  |  | Aldosterone (pg/ml) |  |  | Cortisol (ug/dL) |  |  |
| --- | --- | --- | --- | --- | --- | --- | --- | --- | --- | --- | --- | --- | --- |
|  |  | Mean | SEM | p-val | Mean | SEM | p-val | Mean | SEM | p-val | Mean | SEM | p-val |
| F CON | Chow | 1.19 | 0.08 |  | 0.20 | 0.03 |  | 194.23 | 39.28 |  | 37.17 | 3.92 |  |
| F IUGR |  | 0.88 | 0.11 | **0.084** | 0.30 | 0.10 | 0.40 | 176.87 | 126.58 | 0.90 | 33.60 | 1.35 | 0.44 |
| M CON |  | 0.94 | 0.06 |  | 0.48 | 0.04 |  | 33.06 | 8.19 |  | 43.03 | 3.67 |  |
| M IUGR |  | 1.09 | 0.16 | 0.44 | 0.33 | 0.09 | 0.21 | 118.82 | 70.10 | 0.29 | 52.67 | 9.63 | 0.40 |
| F CON | HFCS | 1.42 | 0.12 |  | 0.27 | 0.07 |  | 147.38 | 74.92 |  | 41.07 | 5.50 |  |
| F IUGR |  | 1.29 | 0.09 | 0.42 | 0.53 | 0.09 | 0.09 | 29.44 | 11.27 | 0.19 | 46.73 | 1.27 | 0.37 |
| M CON |  | 1.28 | 0.03 |  | 0.48 | 0.07 |  | 36.61 | 15.44 |  | 35.00 | 8.82 |  |
| M IUGR |  | 1.37 | 0.09 | 0.40 | 0.31 | 0.11 | 0.23 | 157.50 | 104.55 | 0.32 | 35.63 | 6.10 | 0.96 |
| F CON | Diff | 0.23 | 0.08 |  | 0.08 | 0.08 |  | -46.85 | 93.80 |  | 3.90 | 9.40 |  |
| F IUGR |  | 0.41 | 0.03 | **0.082** | 0.23 | 0.20 | 0.509 | -147.43 | 137.84 | 0.58 | 13.13 | 2.58 | 0.40 |
| M CON |  | 0.35 | 0.04 |  | 0.00 | 0.11 |  | 3.55 | 22.85 |  | -8.03 | 6.58 |  |
| M IUGR |  | 0.29 | 0.12 | 0.66 | -0.02 | 0.17 | 0.92 | 38.68 | 151.16 | 0.83 | -17.03 | 4.22 | 0.31 |

**Table 4.** Heart Rate and Blood Pressure

|  |  | Heart Rate (BPM) |  |  | Mean Blood Pressure (mg/Hg) |  |  | Systolic Blood Pressure (mg/Hg) |  |  | Diastolic Blood Pressure (mg/Hg) |  |  |
| --- | --- | --- | --- | --- | --- | --- | --- | --- | --- | --- | --- | --- | --- |
|  |  | Mean | SEM | p-val | Mean | SEM | p-val | Mean | SEM | p-val | Mean | SEM | p-val |
| M CON | Chow | 114.48 | 6.89 |  | 96.89 | 6.32 |  | 115.92 | 8.28 |  | 83.17 | 4.38 |  |
| M IUGR |  | 111.31 | 1.39 | 0.79 | 90.26 | 6.22 | 0.36 | 107.21 | 0.67 | 0.37 | 78.55 | 0.47 | 0.37 |
| M CON | HFCS | 105.34 | 7.43 |  | 88.41 | 5.60 |  | 113.75 | 3.37 |  | 70.89 | 5.96 |  |
| M IUGR |  | 108.47 | 2.05 | 0.79 | 81.26 | 7.67 | 0.27 | 102.99 | 0.60 | 0.11 | 66.87 | 0.47 | 0.56 |
| M CON | Diff | -9.14 | 1.72 |  | -8.49 | 4.91 |  | -2.17 | 6.27 |  | -12.28 | 5.90 |  |
| M IUGR |  | -2.84 | 3.19 | 0.21 | -9.00 | 1.87 | 0.92 | -4.22 | 0.75 | 0.80 | -11.68 | 0.15 | 0.93 |

#### Renal Transcriptome

In IUGR versus CON females, 687 genes were differentially expressed, 372 down and 315 up (Supplementary Table 1). Four pathways were enriched for differentially expressed genes and passed the end-of-pathway criteria - 3 down, including oxidative phosphorylation and mitochondrial function (Supplementary Table 2).

Causal network analysis in IUGR versus CON females showed 2 networks: RB1 regulatory network was inhibited and the HOXA10 network activated in IUGR versus CON females (Supplementary Table 3). Genes within this merged network were found in multiple signaling pathways including iron homeostasis, oxidative phosphorylation, glycolysis, and cell cycle control. The merged HOXA10 and RB1 inhibitory network shows the extent of overlap among genes (Figure 3).

**Figure 3:**
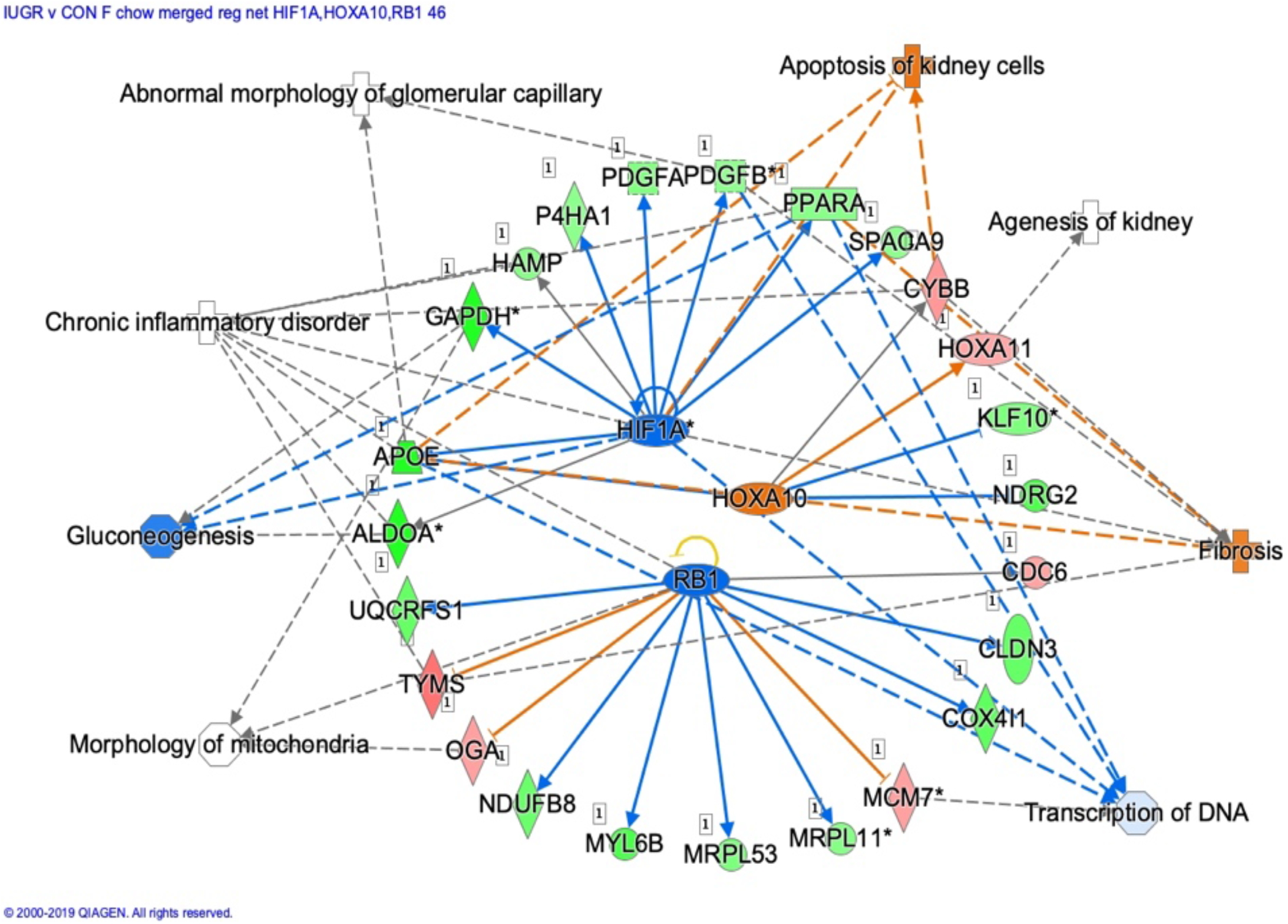
Renal transcriptome comparison of IUGR vs CON females maintained on chow diet. Merged HIF1A, HOXA10, and RB1 inhibitory regulatory networks in IUGR F vs CON F fed the chow diet. Red fill denotes up-regulated genes, green down-regulated, blue predicted down-regulated, orange predicted up-regulated, gray were not different, and white fill were not detected. Lines with arrows denote activation, lines ending in perpendicular line denote inhibition.

In IUGR versus CON males, 1613 genes were differentially expressed with 600 down and 1,013 up (Supplementary Table 1). Five pathways passed end-of-pathway criteria - 3 up-regulated pathways play roles in response to cell stress and DNA damage (Supplementary Table 2). Figure 4 shows activation of BRCA1 in DNA damage signaling in IUGR versus CON males.

**Figure 4:**
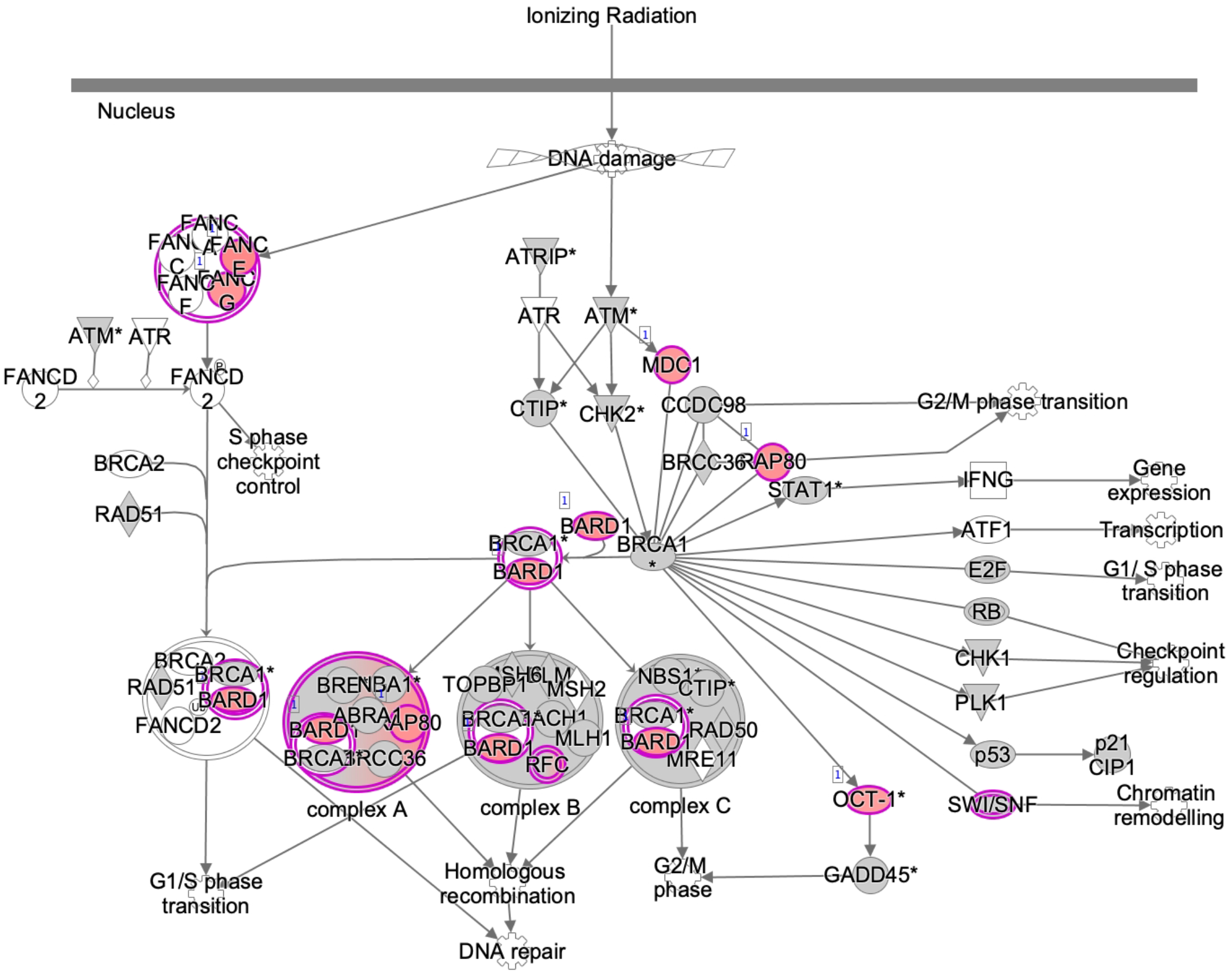
Renal transcriptome comparison of IUGR vs CON males maintained on chow diet. Role of BRCA1 in the DNA damage response signaling pathway in IUGR males compared with CON males fed the chow diet. Red fill denotes up-regulated genes in IUGR males compared with CON males, gray fill indicates genes that were not different and white fill were not detected. Lines with arrows denote activation.

#### Urine Metabolome in IUGR vs CON

Forty-seven metabolites were detected in females and males on chow diet (Supplementary Tables 4 and 5). Of these, 6 were different and 2 marginally different; females were lower than males (Table 5). Sixteen metabolites were present in females on chow but not on HFCS diet. Ten metabolites were present in both sexes, and one, lignoceric acid, was specific to CON females (Table 6, Supplementary Table 5).

**Table 5.**
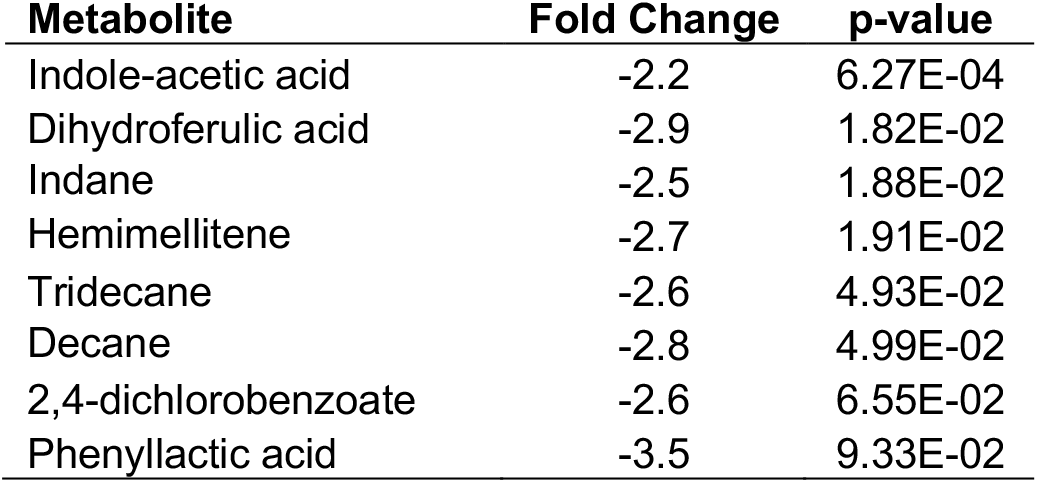
Females vs. Males Chow diet

**Table 6.**
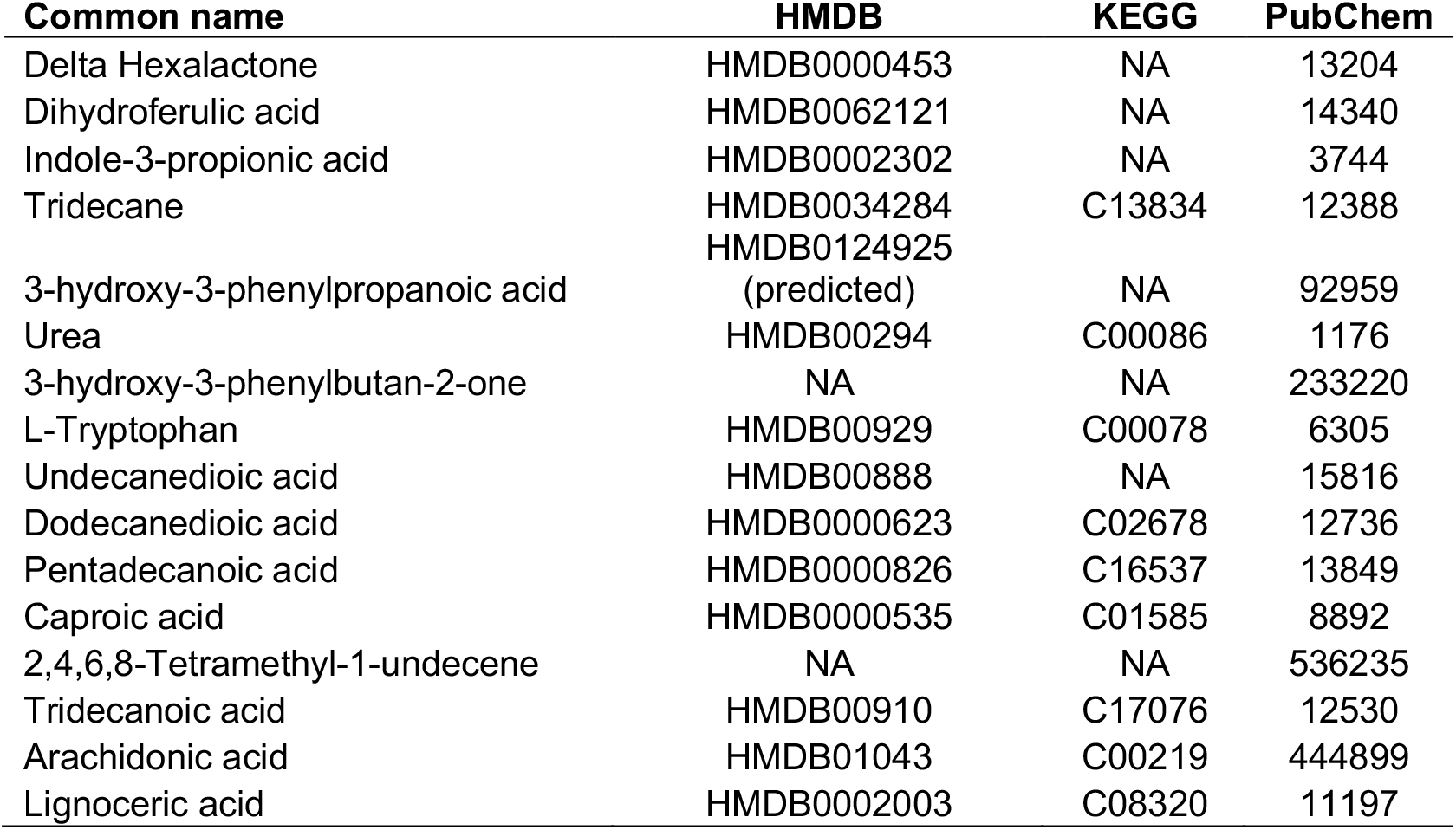
Urine Metabolites in Chow Diet Only

Thirty-six metabolites were detected in all female urine samples, while 15 were identified in all males (Supplementary Table 5). In females, abundances of 2 metabolites (2-ethyl-p-xylene and decane) were higher in IUGR than CON, and 3 metabolites (hemimellitene, tridecane, and indane) were marginally different, with higher abundances in IUGR versus CON (Table 7).

**Table 7.** IUGR vs. CON Females on Chow

| Metabolite | Fold Change | p-value |
| --- | --- | --- |
| 2-ethyl-p-xylene | 1.7 | 3.89E-02 |
| Decane | 1.6 | 4.44E-02 |
| Hemimellitene | 1.6 | 7.31E-02 |
| Tridecane | 2.3 | 7.66E-02 |
| Indane | 1.6 | 7.68E-02 |

### Response to the HFCS Diet Challenge

#### Food Consumption, Morphometrics & Renal Function

IUGR versus CON females were similar in food and fructose drink consumption during the HFCS challenge (not shown). Weight and BMI were less in IUGR than CON females after the challenge (Table 1). The 7-week average for urine volume, urine creatinine, and urine sodium was greater, while urine potassium was marginally less in IUGR versus CON females (Table 2). Serum measures of creatinine, aldosterone, and cortisol were similar in IUGR and CON females (Table 3). On average for the HFCS challenge urine potassium excretion was lower in IUGR than CON females (Table 2).

In IUGR versus CON males, food and fructose drink consumption were similar during the HFCS challenge. IUGR males’ weight and BMI after the challenge were similar to CON males. The change in BMI from after challenge to baseline showed an increase in IUGR and decrease in CON males (Table 1). The 7-week average for urine volume, urine creatinine, urine potassium, urine sodium, serum creatinine, aldosterone, and cortisol were similar between IUGR and CON (Tables 2–3), and showed a similar response to HFCS for systolic, diastolic and mean arterial blood pressure, and heart rate (Table 4).

#### Renal Transcriptome

##### CON Females HFCS vs Chow

Three-hundred-forty-nine genes were differentially expressed in CON females fed HFCS versus chow: 133 were down and 216 were up (Supplemental Table 6). Two pathways passed end-of-pathway criteria with one up and one down (Supplemental Table 2). Nine networks were inhibited and 8 activated (Figure 5, Supplemental Table 3). Genes in all 17 networks overlap with genes in the 3 canonical pathways. Genes in the networks have roles in multiple canonical pathways including prothrombin activation, fatty acid biosynthesis, Wnt/β-catenin signaling, cyclins and cell cycle regulation (Supplemental Table 3).

**Figure 5:**
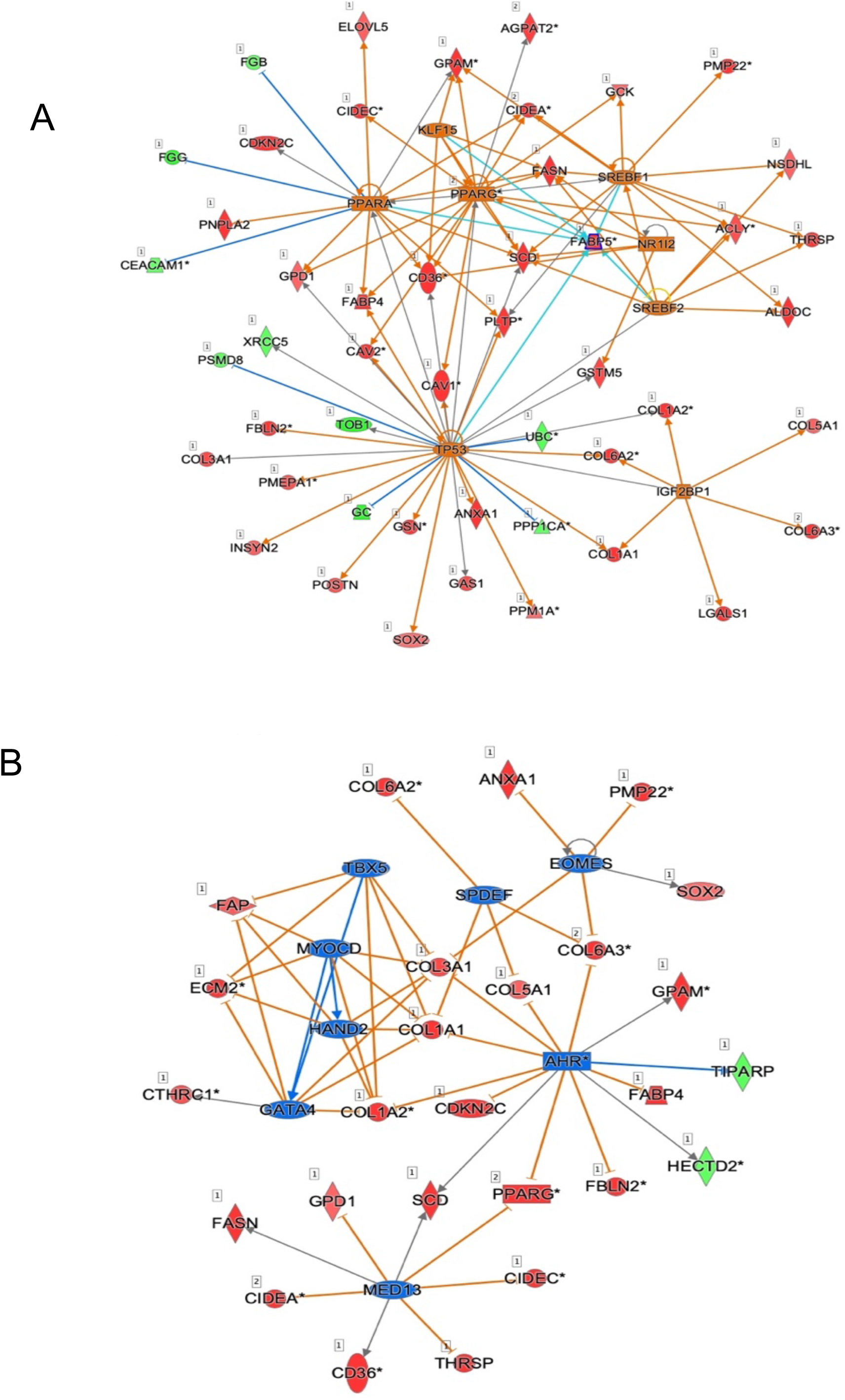
Renal transcriptome comparison of CON females fed the HFCS diet compared with chow diet. Activated (A) and inhibited (B) regulatory networks in CON females fed the HFCS diet versus chow diet. Red fill denotes up-regulated genes, green down-regulated, blue predicted down-regulated, orange predicted up-regulated, gray were not different, and white fill were not detected. Lines with arrows denote activation, lines ending in perpendicular line denote inhibition.

##### IUGR Females HFCS vs Chow

Seven-hundred-ninety-three genes were differentially expressed in IUGR females fed HFCS versus chow: 447 were down and 346 were up (Supplemental Table 6). Seven pathways passed end-of-pathway criteria - all down, including mTOR (Figure 6), eIF4 & p70S6K, and PI3K/AKT signaling. One network with regulator SMARCA4 was activated (Figure 7); genes in this network are included in canonical pathways glucocorticoid receptor, eNOS, oxidative stress response, calcium, and integrin signaling (Supplemental Table 3).

**Figure 6:**
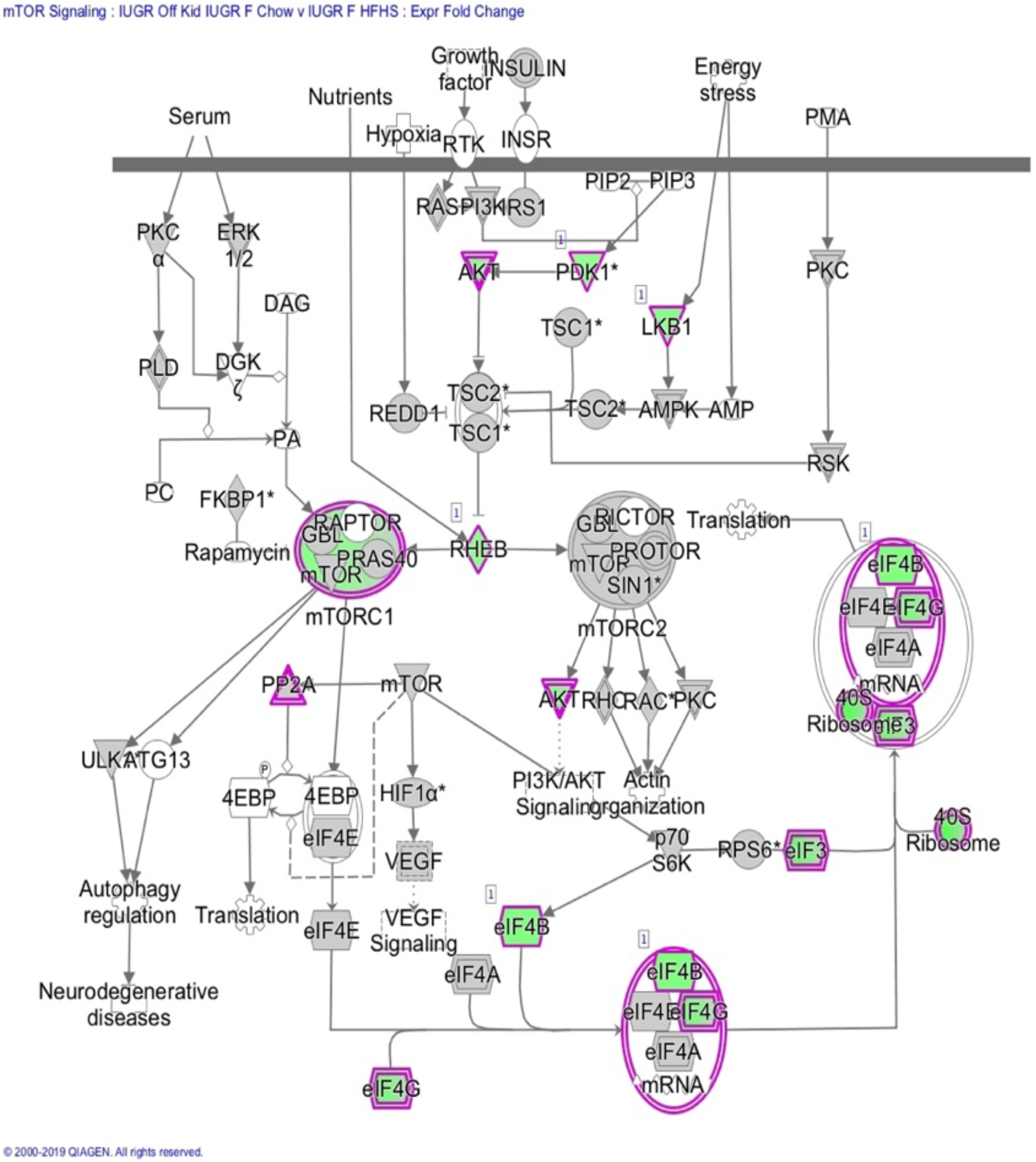
(A) Down regulation of mTOR signaling in kidneys of IUGR females fed the HFCS diet compared with chow diet. Green fill denotes down-regulated genes in IUGR females compared with CON females, gray fill indicates genes that were not different and white fill were not detected. Lines with arrows denote activation, lines ending in perpendicular line denote inhibition. (B) Activation of SMARCA4 regulatory network in renal transcriptome of IUGR females fed the HFCS diet compared with chow diet. Red fill denotes up-regulated genes, green down-regulated, and orange predicted up.

**Figure 7:**
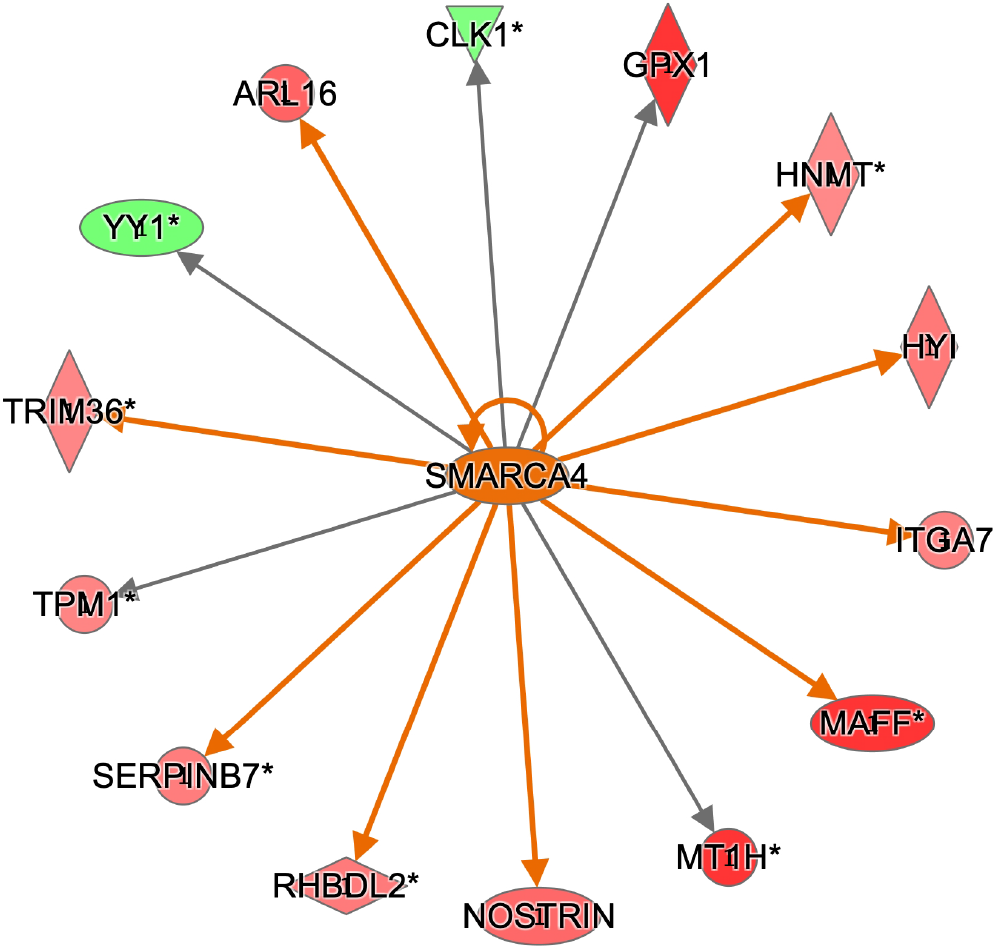
Renal transcriptome comparison of CON males fed the HFCS diet compared with chow diet. Merged CREB1, AR, and SRF activated regulatory networks in CON males fed the HFCS diet versus chow diet. Red fill denotes up-regulated genes, green down-regulated, and orange predicted up-regulated. Lines with arrows denote activation, lines ending in perpendicular line denote inhibition.

##### CON Males HFCS vs Chow

Nine-hundred-fifty-nine genes were differentially expressed in CON males fed HFCS compared with chow: 354 were down and 605 were up (Supplemental Table 6). Eight pathways passed end-of-pathway criteria - all were up, including cholesterol biosynthesis, and integrin and ILK signaling (Supplemental Table 2). Of 5 regulatory networks, 3 were activated by CREB1, AR, and SRF (Figure 8). Genes in each network overlapped with genes in 6 pathways. Genes in these networks have roles in corticotropin releasing hormone, cholesterol biosynthesis, glucocorticoid receptor, and integrin signaling (Supplemental Table 3).

**Figure 8:**
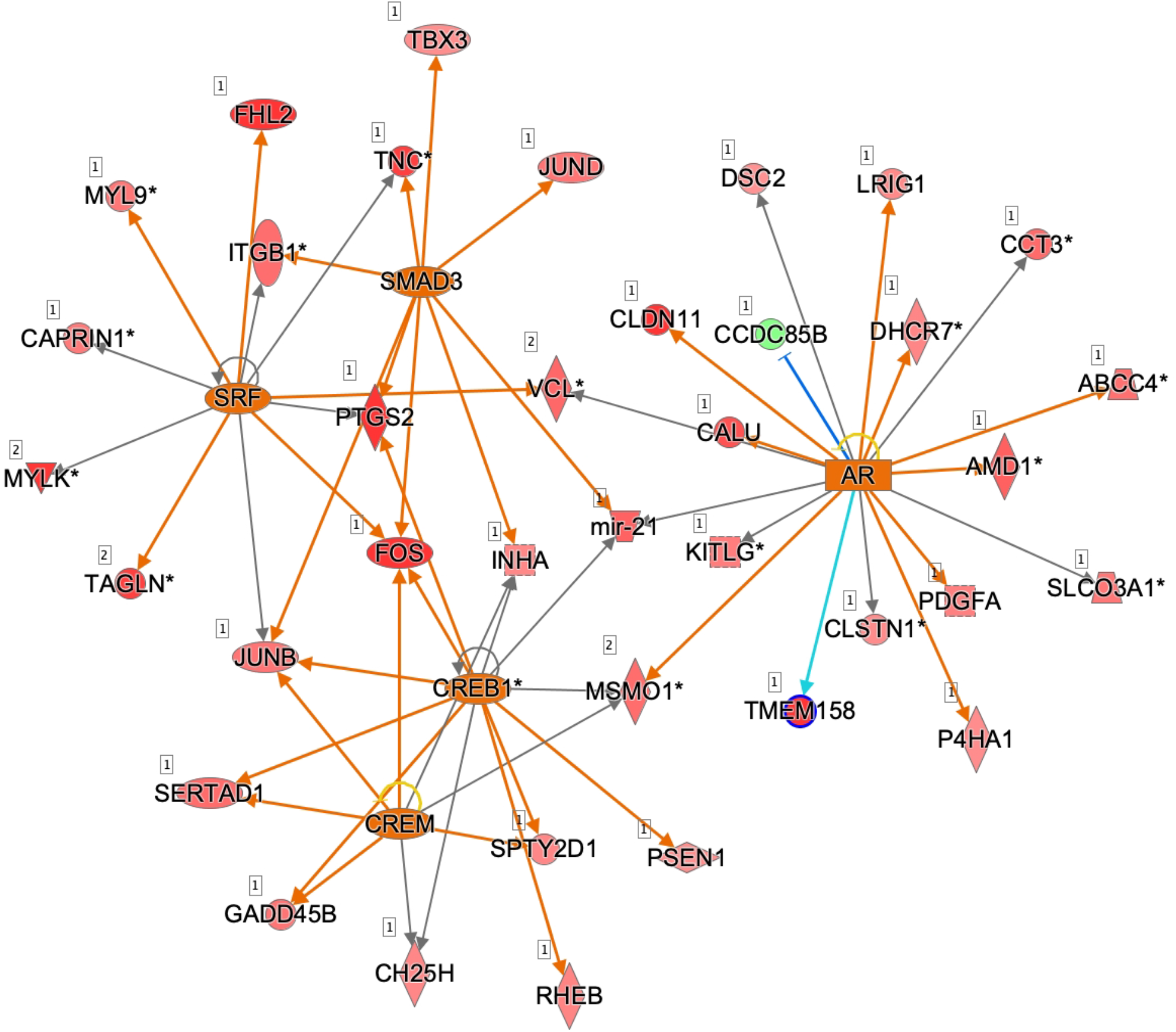
Renal transcriptome comparison of IUGR males fed the HFCS diet compared with chow diet. Merged PARP1 and POU4F1 inhibited regulatory networks in IUGR males fed the HFCS diet versus chow diet. Red fill denotes up-regulated genes, green down-regulated, and blue predicted down-regulated. Lines with arrows denote activation, lines ending in perpendicular line denote inhibition.

##### IUGR Males HFCS vs Chow

Twelve-hundred-nine genes were differentially expressed in IUGR males fed HFCS versus chow: 503 were down and 706 were up (Supplemental Table 6). Two pathways passed end-of-pathway criteria - mitochondrial function was up and assembly of RNA polymerase complex was down (Supplemental Table 2). Two networks with regulators PARP1 and POU4F1 were inhibited (Figure 9). Genes in these networks have roles in thrombin and glucocorticoid receptor signaling.

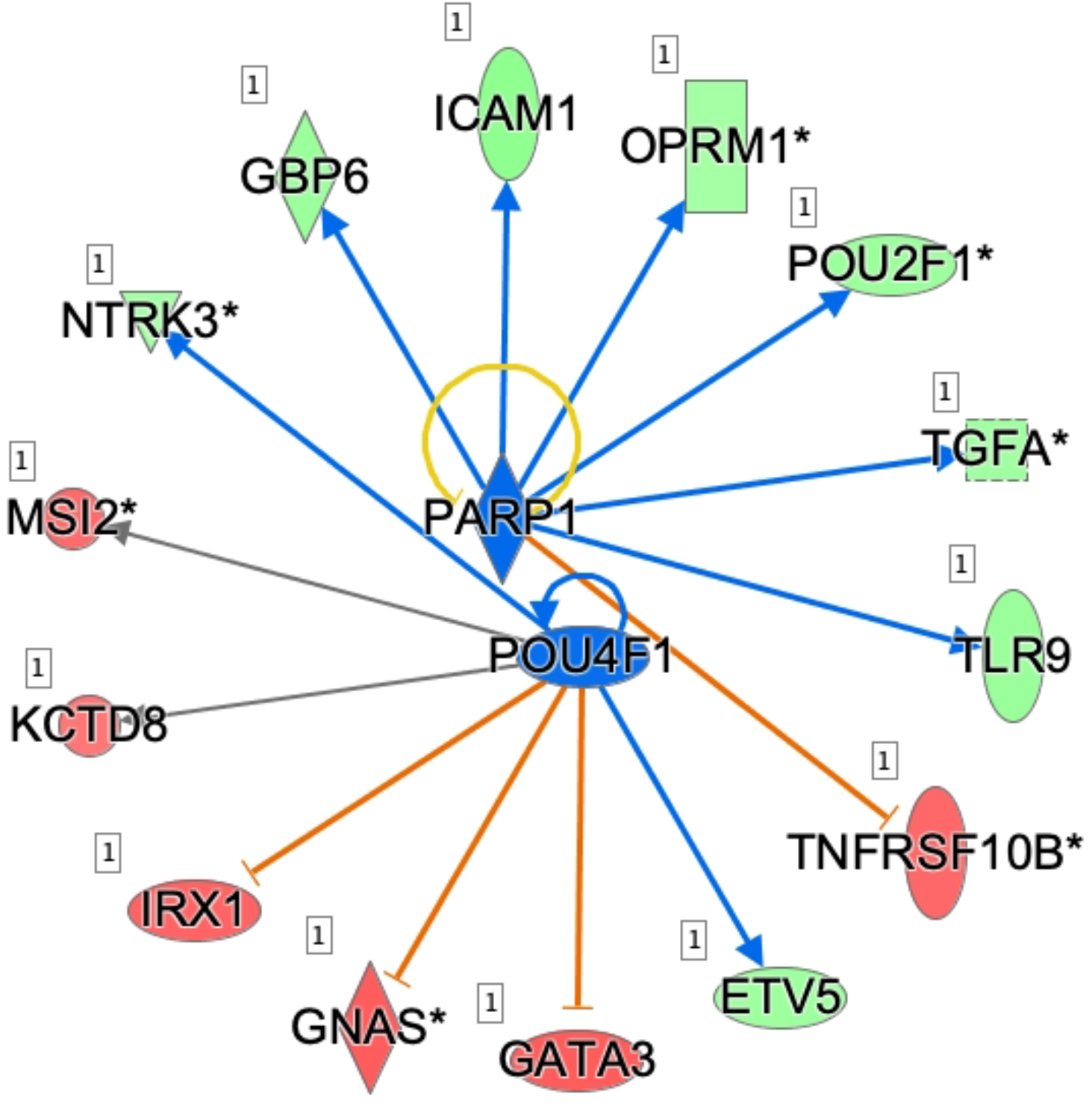

#### Urine Metabolome in IUGR vs CON Females and Males on HFCS

Thirty-eight metabolites were detected above background in urine from males and females on HFCS (Supplementary 5). Of these, seven metabolites were differentially expressed between females and males with 6 being less abundant in females (Table 8). PCA of these metabolites, revealed group clustering by sex (Figure 2B). Seven of the metabolites detected after the challenge were specific to HFCS (Table 9).

**Table 8.** Females vs. Males on HFCS Diet

| Metabolite | Fold Change | p-value |
| --- | --- | --- |
| Indole-acetic acid | -4.365 | 4.00E-03 |
| Azelaic acid | -3.036 | 8.07E-03 |
| Hippuric acid | -5.323 | 1.27E-02 |
| Prehnitene | -1.639 | 1.42E-02 |
| Hemimellitene | -1.550 | 2.82E-02 |
| Indane | -1.734 | 3.03E-02 |
| N,N-dimethyl-1-Pentadecanamine | 1.906 | 4.84E-02 |

**Table 9.** Diet-Specific Metabolites - Urine Metabolites in HFCS Diet Only

| Common name | HMDB ID | KEGG ID | PubChem ID |
| --- | --- | --- | --- |
| (S)3-hydroxyisobutyric acid (3-HIBA) | HMDB0000023 | C06001 | 440873 |
| Prehnitene | HMDB0059823 | NA | 10236 |
| 4-Hydroxyphenylpyruvic acid (4-HPPA) | HMDB0000707 | C01179 | 979 |
| 3-t-Pentylcyclopentanone | NA | NA | 551379 |
| 6-methyl-2-Hepten-4-one | NA | NA | 12502788 |
| 4-methylcatechol | HMDB0000873 | C06730 | 9958 |
| Mercaptoacetic acid | NA | C02086 | 1133 |

CON females fed HFCS versus chow showed 2 decreased metabolites - hemimellitene increased and azelaic acid (Table 10). Comparison of IUGR females fed HFCS versus chow revealed 6 marginally different metabolites. Stearic acid showed lower abundance, and all others higher abundance in HFCS versus chow (Table 11). PCA of metabolites present in both IUGR chow and HFCS female urine samples revealed increased separation of clusters versus CON females (Supplementary Figure 1).

**Table 10.**
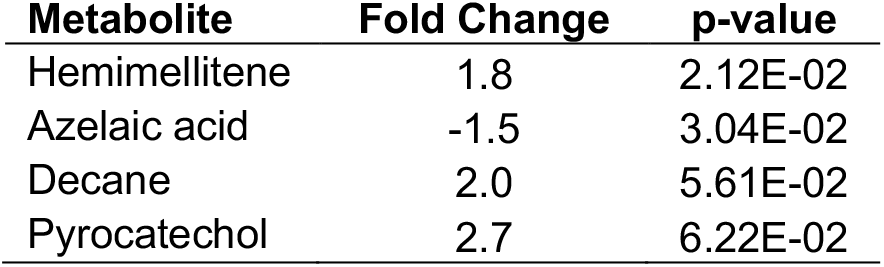
CON Females on HFCS vs. Chow Diet

**Table 11.** IUGR Females on HFCS vs. Chow Diet

| **Metabolite** | **Fold Change** | **p-value** |
| --- | --- | --- |
| Hemimellitene | 1.4 | 5.83E-02 |
| Stearic acid | -7.3 | 5.12E-02 |
| N,N-dimethyloctylamine | 5.1 | 5.72E-02 |
| Lauric acid | 2.7 | 6.23E-02 |
| Indane | 1.4 | 5.86E-02 |
| Hydrocinnamic acid | 1.6 | 8.64E-02 |

IUGR and CON male urine metabolite abundance was similar on chow. Following HFCS challenge, 4 metabolites were different in IUGR versus CON: N-mehtyl-N-octyl-1-octanamine and N,N-dimethyloctlamine decreased, and hippuric acid and lauric acid increased (Table 12).

**Table 12.** IUGR vs CON Males on HFCS Diet

| **Metabolite** | **Fold Change** | **p-value** |
| --- | --- | --- |
| N-methyl-N-octyl-1-Octanamine | 1.01 | 4.22E-03 |
| N,N-Dimethyloctylamine | 1.01 | 3.16E-02 |
| Hippuric acid | -1.03 | 3.26E-02 |
| Lauric acid | -1.00 | 4.90E-02 |

## Discussion

A poor fetal nutritional environment, due not only to food availability but also poor placental function, followed by a postnatal calorie-rich environment is common in developed countries. This prenatal/postnatal caloric mismatch predisposes IUGR offspring to early onset metabolic dysregulation leading to dyslipidemia and hypertension ^40^ as juveniles ^21 41^ and on cardiac function as young adults ^19^. Here we follow up two questions from our fetal kidney studies: 1) Do fetal MNR effects on kidney growth and function persist postnatally in IUGR offspring; 2) Does a second hit stressor of caloric mismatch between fetal and postnatal life reveal renal vulnerabilities undetected without the second hit.

### Comparison of CON and IUGR prior to the HFCS challenge

In postnatal IUGR females, measures of kidney function, gene expression and metabolite abundance differed from CON females. Signs of renal dysfunction were decreased urine potassium and serum creatinine, and increased urine creatinine in IUGR versus CON females. Urine metabolome analysis revealed elevated abundances of aromatic hydrocarbons in IUGR versus CON females, indicating changes in metabolic endproducts associated with mitochondrial dysfunction ^42^. Transcriptome analysis revealed down-regulation of oxidative phosphorylation and mitochondrial function and inhibition of causal networks regulated by RB1 and HOXA10 in IUGR versus CON females. These results indicate that juvenile IUGR female kidneys differ from CON at cellular and molecular levels. Decreased human urinary potassium excretion correlates with reduced renal proximal tubule function ^43, 44^. Aromatic hydrocarbons are associated with mitochondrial function in human chronic kidney disease ^42^. The presence of altered aromatic hydrocarbons may be related to differing inflammatory or oxidative states in CON and IUGR kidneys as well as responses to HFCS diet. Down-regulation of renal oxidative phosphorylation and mitochondrial function is associated with kidney injury ^45^. RB1 induces cellular senescence and hypertension ^46^. Functional annotations of genes in this network predict decreased glomerular growth and gluconeogenesis, and increased apoptosis in IUGR versus CON female kidneys. HOXA10 expression is specific for proximal tubules and involved in sodium overload response ^47, 48^; in rats HOXA10 activation correlates with early renal pathogenesis ^49^. Activation of the HOXA10 network is consistent with inhibition of oxidative phosphorylation in IUGR female kidneys versus CON. HIF1A expression, which is down-regulated, is specific to proximal tubules and involved development ^50^. In addition, regulators RB1 and HOXA10 are predicted to inhibit networks containing genes involved in mitochondrial function, cell cycle regulation and initiation of transcription. Together, these suggest that shorter proximal tubules ^15^ in females persist postnatally and that IUGR predisposes to reduced renal function.

In males on chow diet, transcriptional pathways related to cell stress and DNA damage were up-regulated in IUGR versus CON. More than 1,600 genes were differentially expressed in IUGR versus CON males (~2.5x more than IUGR versus CON females); however, only 5 pathways passed end-of-pathway criteria, with 3 related to DNA damage. No regulatory networks showed coordinated outcomes. These results show that IUGR differ from CON male kidneys. In addition, the large number of differentially expressed genes with lack of enrichment in coordinated pathways or networks suggest this difference is due to molecular dysregulation of the IUGR male kidney.

### Comparison of CON and IUGR response to HFCS

To address the question whether the second hit of a nutritional mismatch between fetal and postnatal life reveals renal vulnerabilities in juvenile primates, we compared IUGR and CON before and after the HFCS diet.

IUGR females weighed less with lower BMIs than CON after the 7-week challenge. The 7-week average urine volume, creatinine and sodium was greater with potassium nominally less in IUGR versus CON females, suggesting early preclinical dysfunction of female IUGR kidneys. In CON females, the transcriptome response to the HFCS challenge revealed coordinated expression of 2 canonical pathways and 17 regulatory networks. Of note is activation of SREBF1 and SREBF2 networks. Under conditions of ER stress, SREBF1 inhibits signaling pathways, such as NF-κB ^51^. SREBF2 increases renal cholesterol synthesis ^52^. Inclusion of genes in the top regulatory networks with roles in fatty acid biosynthesis and Wnt/β-catenin signaling (e.g., FASN, SOX2) suggest a coordinated transcriptional response to the HFCS diet. The significant overlap of genes in canonical pathways and regulatory networks provides additional support for CON females coordinately responding to the HFCS challenge.

In IUGR females, the transcriptome response to the HFCS diet revealed a coordinated response with multiple pathways down-regulated; down-regulation of mTOR and p706K indicates a persistence of decreased mTOR signaling demonstrated in IUGR versus CON in the 0.5G baboon fetal kidney ^15 16^. Our findings that mTOR signaling remains down in IUGR females with increased nutrient availability suggests that fetal MNR effects persist in these juveniles and lack of coordinated response to the HFCS may indicate inability to respond appropriately to the second hit. Additionally, up-regulation of genes within the SMARCA4 network are central to the DNA damage response (SMARCA4) ^53^, oxidative damage (GPX1) ^54^ and tissue remodeling (TPM1) ^55^, suggesting increased renal fibrosis due to HFCS diet.

The female IUGR urine metabolome responded differently to the HFCS challenge than the female CON urine metabolome. PCA of metabolites indicated greater metabolic differences in response to the HFCS diet in IUGR than CON females. These results are also consistent with our gene expression results showing differential diet response. Additional studies are needed to determine whether the IUGR metabolic profile is indicative of renal vulnerability.

We identified urine metabolites specific to chow or HFCS diets. Two metabolites, tryptophan and indole-3 propionic acid, associated with tryptophan metabolism, were detected on chow diet. Indole-3 propionic acid, a deamination product of tryptophan, has been associated with susceptibility to Type 2 Diabetes ^56^. Of the metabolites identified for HFCS diet but not chow, 3-hydroxyisobutyric acid (3-HIBA) and 4-hydroxyphenylpyruvic acid (4-HPPA) are associated with amino acid metabolism. 3-HIBA, detected in every urine sample following the HFCS diet, is a product of valine oxidation ^42^ and may have inhibitory effects on mitochondrial creatine kinase activity ^57^, important for energy homeostasis ^58^. 3-HIBA has been associated with diabetic kidney disease, and is lower in diabetics with kidney disease compared to diabetics without kidney disease ^42^.

In females, we observed increased presence of methylated metabolites (prehnitene, 6-methyl-heptenone and 4-methyl-catechol) in urine from the HFCS challenge, suggesting HFCS diet may alter overall methylation status of metabolites, which warrants future investigation of epimetabolites ^59^. In addition, the same enzymes methylate DNA, indicating the need to evaluate kidney DNA methylation’s role in epigenetic control.

The male transcriptional response to HFCS differed between IUGR and CON. CON males showed up-regulation of cholesterol synthesis and activation of a CREB1 regulated network; CREB1 activates proximal tubule sodium and fluid transport ^60^. The androgen receptor (AR) network is also of interest - androgens in blood and intrarenal synthesis ^61^ can bind to distal tubular AR cells, increasing epithelial sodium transport ^62^. SRF, one regulator in this network, is involved in maintenance of podocyte structure and function ^63^.

Unlike CON males, IUGR males showed no coordinated renal molecular response to HFCS. HFCS versus chow in IUGR males revealed 1209 differentially expressed genes, ~25% more than CON males’ response and ~50% more than IUGR females. In spite of the greater number of differentially expressed genes in IUGR males compared with all other groups, only 2 pathways passed end-of-pathway criteria - mitochondrial function and assembly of RNA polymerase complex, and only one network - PARP1, was identified. PARP inhibition is vasoprotective in models of renal failure ^64^.

The male CON and IUGR urine metabolome showed more variation than females. Although many of the same metabolites were detected in females and males, some metabolites were not detected in more than one male per group, limiting our ability to assess statistical differences between CON and IUGR animals on both diets.

Although we saw differences between IUGR and CON male response to HFCS at the molecular level, both gene expression and metabolite abundance, we did not find differences in measures of renal function, blood pressure, or heart rate. Few studies that have evaluated renal function in mammalian IUGR juvenile offspring demonstrated catch-up growth. Study of IUGR juvenile sheep found catch-up growth in response to an energy-dense diet and decreased nephron numbers ^65^. Importantly, in agreement with our blood pressure data from males, they found no evidence of hypertension. It is possible that clinical effects of IUGR on kidney function are not manifest during the juvenile period due to large metabolic demands of rapid growth. It is also possible that a 7-week challenge is too short to impact blood pressure.

This study showed that *in utero* renal effects of MNR persist in postnatal IUGR juvenile primates. Additionally, a second metabolic hit revealed sex-specific renal vulnerabilities beyond those programmed by IUGR alone and indicated reduced kidney function, altered urine metabolome and renal transcriptome activity, which are consistent with impaired nutrient sensing and consistent with findings of MNR effects on 0.5G primate kidneys. Exposure to the HFCS challenge resulted in renal transcriptome changes, suggesting a lack of coordinated molecular response to the challenge. Longer term study as animals age is required to determine if molecular changes observed in this study manifest as impairment of kidney function in IUGR offspring as adults.

## Author Contributions

LAC, MJN, RES, and PWN designed the study; ACB, KDSR, KJL, SB, ML, LAC, KF, EJD, and CL carried out the experiments; ACB, KDSR, LAC, RES, and MJN analyzed the data; ACB and LAC made the figures; ACB, PWN and LAC drafted and revised the paper; all authors approved the final version of the manuscript.

## Acknowledgments

This investigation was supported by P01 HD021350, P51 RR013986, and P51 OD011133. This investigation was conducted in facilities constructed with support from C06 RR14578, C06 RR15456, C06 RR013556, and C06 RR017515.

## Supplemental Material Table of Contents

Supplementary Table 1: IUGR vs CON Chow Diet Gene Expression

Supplementary Table 2: Pathways Enrichment Analysis

Supplementary Table 3: Causal Network Analysis

Supplementary Table 4. Normalized and transformed values of identified metabolites F=female, M=male, CON = Control, IUGR = Interuterine Growth Restricted, 0 = Chow Diet, 7 = HFCS diet.

Supplementary Table 5. Identifiers of metabolites including NIST Identifier, common name, HMDB ID, KEGG ID, PubChem ID, Class and Pathway. X = present in all animals, O = not present in all animals.

Supplementary Table 6: HFCS Diet vs Chow Diet Gene Expression

Supplemental Figure 1: Principal component analysis (PCA) of female urine metabolite signatures show altered response on HFCS diets compared to chow. Metabolite clustering show increase separation for B) IUGR female compared to A) CON females indicative of a greater response to the HFCS diet.

