## Supplemental Figure 1 and Tables 1-6 for "Postnatal persistence of sex-dependent renal developmentally programmed structural and molecular changes in nonhuman primates"

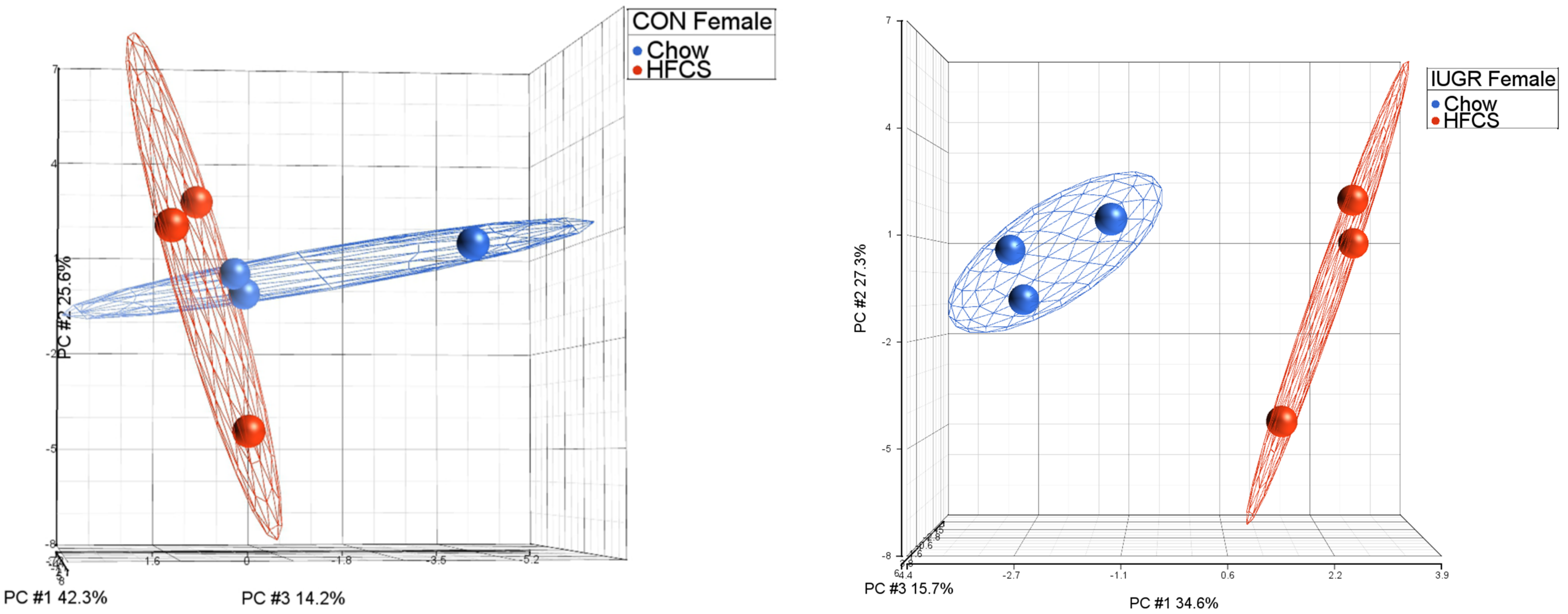

Supplementary Table 1: IUGR vs CON Chow Diet Gene Expression

| Gene Symbol | Gene Name | Accession No. | CON Chow F Mean | IUGR Chow F Mean | pval IUGR v CON F Chow | Ratio IUGR v CON F Chow | Nephron Segment-Specific REFSEQ ID | NEPHRON (MEAN RPKM) | COLLECTING DUCT (MEAN RPKM) |
|---|---|---|---|---|---|---|---|---|---|
| Xist | X inactive specific transcript (non-protein coding)(XIST) | NR_001564 | 599.6 | 836.2 | 0.0001 | 1.3 | | | |
| GTPBP6 | GTP binding protein 6 (putative)(GTPBP6) | NM_012227 | 244.8 | 183 | 0.0002 | 0.69 | | | |
| CSDE1 | cold shock domain containing E1(CSDE1) | NM_001007553 | 143 | 131.3 | 0.0003 | 0.75 | | | |
| CSDE1 | cold shock domain containing E1(CSDE1) | NM_001007553 | 187.8 | 156.9 | 0.0003 | 0.75 | | | |
| OSTM1 | osteopetrosis associated transmembrane protein 1(OSTM1) | NM_014028 | 91.4 | 131.6 | 0.0003 | 1.28 | | | |
| | | BQ186959 | 111.6 | 106.8 | 0.0004 | 0.88 | | | |
| | | DB336481 | 147.3 | 135.5 | 0.0004 | 0.82 | NM_017193 | 356.13 | 8.18 |
| UBC | ubiquitin C(UBC) | NM_021009 | 10602.5 | 9244.7 | 0.0005 | 0.68 | NM_017193 | 356.13 | 8.18 |

| | | | | | | | | | |
|---|---|---|---|---|---|---|---|---|---|
| UBC | ubiquitin C(UBC) | NM_021009 | 12020.7 | 11235.1 | 0.0005 | 0.68 | NM_017193 | 356.13 | 8.18 |
| ODC1 | ornithine decarboxylase 1(ODC1) | NM_002539 | 463.8 | 725.5 | 0.0006 | 1.42 | | | |
| PCDHA3 | protocadherin alpha 3(PCDHA3) | NM_018906 | 105.8 | 136.8 | 0.0006 | 1.15 | | | |
| SREBF1 | sterol regulatory element binding transcription factor 1(SREBF1) | NM_004176 | 150.6 | 118.2 | 0.0008 | 0.73 | | | |
| UAP1L1 | UDP-N-acetylglucosamine pyrophosphorylase 1 like 1(UAP1L1) | NM_207309 | 134.5 | 114.8 | 0.0008 | 0.78 | | | |
| DNAJB6 | DnaJ heat shock protein family (Hsp40) member B6(DNAJB6) | NM_005494 | 206 | 182.1 | 0.0011 | 0.79 | | | |
| UBE3C | ubiquitin protein ligase E3C(UBE3C) | NM_014671 | 160.3 | 131.2 | 0.0013 | 0.75 | | | |
| UBE3C | ubiquitin protein ligase E3C(UBE3C) | NM_014671 | 478.2 | 511.9 | 0.0013 | 0.75 | | | |
| | | XM_941684 | 297.7 | 538.9 | 0.0014 | 1.65 | | | |

| | | | | | | |
|---|---|---|---|---|---|---|
| PAH | phenylalanine hydroxylase(PAH) | NM_000277 | 126.8 | 100.9 | 0.0015 | 0.75 |
| ZNF467 | zinc finger protein 467(ZNF467) | NM_207336 | 264.2 | 193.3 | 0.0015 | 0.67 |
| CYB5D2 | cytochrome b5 domain containing 2(CYB5D2) | NM_144611 | 125.3 | 170.6 | 0.0016 | 1.18 |
| NDUFB11 | NADH:ubiquinone oxidoreductase subunit B11(NDUFB11) | NM_019056 | 1498.6 | 1222.1 | 0.0018 | 0.75 |
| APOE | apolipoprotein E(APOE) | NM_000041 | 2789.9 | 1250.2 | 0.0021 | 0.39 |
| LGMN | legumain(LGMN) | NM_001008530 | 135.2 | 143.9 | 0.0021 | 0.41 |
| LGMN | legumain(LGMN) | NM_001008530 | 203.6 | 186.1 | 0.0021 | 0.41 |
| LGMN | legumain(LGMN) | NM_001008530 | 1731.9 | 783.9 | 0.0021 | 0.41 |
| | | XM_941785 | 110.7 | 106.4 | 0.0022 | 0.88 |
| NUMA1 | nuclear mitotic apparatus protein 1(NUMA1) | NM_006185 | 123 | 158 | 0.0024 | 1.3 |
| NUMA1 | nuclear mitotic apparatus protein 1(NUMA1) | NM_006185 | 252 | 368.7 | 0.0024 | 1.3 |

| | | | | | | |
|---|---|---|---|---|---|---|
| MID1IP1 | MID1 interacting protein 1(MID1IP1) | NM_021242 | 116 | 115.3 | 0.0028 | 0.72 |
| MID1IP1 | MID1 interacting protein 1(MID1IP1) | NM_021242 | 199.5 | 158.2 | 0.0028 | 0.72 |
| SRSF12 | serine and arginine rich splicing factor 12(SRSF12) | NM_080743 | 97.8 | 136.2 | 0.0028 | 1.23 |
| | | XM_001719540 | 114.6 | 150.1 | 0.0029 | 1.14 |
| BAG3 | BCL2 associated athanogene 3(BAG3) | NM_004281 | 291.2 | 149.3 | 0.0031 | 0.47 |
| USP21 | ubiquitin specific peptidase 21(USP21) | NM_012475 | 133.7 | 142.8 | 0.0032 | 1.24 |
| USP21 | ubiquitin specific peptidase 21(USP21) | NM_012475 | 150 | 217.3 | 0.0032 | 1.24 |
| CASKIN2 | CASK interacting protein 2(CASKIN2) | NM_020753 | 116.9 | 106.8 | 0.0034 | 0.85 |
| | | XM_933893 | 108.3 | 149.8 | 0.0034 | 1.2 |
| TIPARP | TCDD inducible poly(ADP-ribose) polymerase(TIPARP) | NM_015508 | 396.4 | 319.1 | 0.0036 | 0.75 |

| | | | | | | |
|---|---|---|---|---|---|---|
| XYLT1 | xylosyltransferase 1(XYLT1) | NM_022166 | 95.4 | 125.5 | 0.0036 | 1.18 |
| | | BX111977 | 112.6 | 145.7 | 0.0036 | 1.14 |
| | | XM_934376 | 115.9 | 149.6 | 0.0037 | 1.13 |
| SBDSP1 | Shwachman-Bodian-Diamond syndrome pseudogene 1(SBDSP1) | NR_001588 | 336.4 | 258.4 | 0.0038 | 0.68 |
| SBDSP1 | Shwachman-Bodian-Diamond syndrome pseudogene 1(SBDSP1) | NR_001588 | 415.5 | 362.7 | 0.0038 | 0.68 |
| DCUN1D2 | defective in cullin neddylation 1 domain containing 2(DCUN1D2) | NM_001014283 | 205 | 190.3 | 0.0039 | 0.82 |
| ELMOD2 | ELMO domain containing 2(ELMOD2) | NM_153702 | 160.2 | 132.5 | 0.0041 | 0.77 |
| SCARA3 | scavenger receptor class A member 3(SCARA3) | NM_016240 | 97.4 | 131.1 | 0.0041 | 1.19 |
| OR6N2 | olfactory receptor family 6 subfamily N member 2(OR6N2) | NM_001005278 | 105.4 | 134 | 0.0044 | 1.14 |

| | | | | | | |
|---|---|---|---|---|---|---|
| | | XR_019029 | 129.5 | 175.5 | 0.0044 | 1.18 |
| ARHGEF37 | Rho guanine nucleotide exchange factor 37(ARHGEF37) | NM_001001669 | 124.1 | 173.1 | 0.0047 | 1.21 |
| CDCA7 | cell division cycle associated 7(CDCA7) | NM_031942 | 128.6 | 214.8 | 0.0049 | 1.45 |
| | | XR_039439 | 111.5 | 155.4 | 0.0049 | 1.21 |
| | | XR_039439 | 1038.8 | 1435 | 0.0049 | 1.21 |
| DPCD | deleted in primary ciliary dyskinesia homolog (mouse)(DPCD) | NM_015448 | 281.5 | 229.7 | 0.005 | 0.72 |
| SEL1L3 | SEL1L family member 3(SEL1L3) | NM_015187 | 113.5 | 162.3 | 0.0053 | 1.23 |
| | | XR_038700 | 102.2 | 149.8 | 0.0053 | 1.26 |
| | | XR_018838 | 192.5 | 163.3 | 0.0056 | 0.77 |
| MRPL53 | mitochondrial ribosomal protein L53(MRPL53) | NM_053050 | 231.1 | 205 | 0.0057 | 0.77 |

| | | | | | | | | | |
|---|---|---|---|---|---|---|---|---|---|
| KCNJ14 | potassium voltage-gated channel subfamily J member 14(KCNJ14) | NM_170720 | 120.1 | 162.7 | 0.0059 | 1.18 | | | |
| KMT2C | lysine methyltransferase 2C(KMT2C) | NM_170606 | 127.7 | 200.7 | 0.0059 | 1.34 | | | |
| KMT2C | lysine methyltransferase 2C(KMT2C) | NM_170606 | 138.7 | 155.8 | 0.0059 | 1.34 | | | |
| TRAPPC2 | trafficking protein particle complex 2(TRAPPC2) | NM_001011658 | 108.3 | 140.7 | 0.006 | 1.14 | | | |
| c1orf216 | chromosome 1 open reading frame 216(C1orf216) | NM_152374 | 123.1 | 162 | 0.0061 | 1.15 | | | |
| HAUS4 | HAUS augmin like complex subunit 4(HAUS4) | NM_017815 | 164.4 | 138.2 | 0.0062 | 0.76 | NM_053502 | 4.02 | 18.01 |
| SCAPER | S-phase cyclin A associated protein in the ER(SCAPER) | NM_020843 | 200.8 | 181.1 | 0.0063 | 0.78 | NM_053502 | 4.02 | 18.01 |

| | | | | | | | | | |
|---|---|---|---|---|---|---|---|---|---|
| c3p1 | complement component 3 precursor pseudogene(C3P1) | NM_001013640 | 125.7 | 168.7 | 0.0066 | 1.17 | NM_053502 | 4.02 | 18.01 |
| | | AK129699 | 141.4 | 332 | 0.0069 | 2.08 | | | |
| PEX2 | peroxisomal biogenesis factor 2(PEX2) | NM_000318 | 242.1 | 194.7 | 0.007 | 0.71 | | | |
| | | XM_928619 | 183.9 | 175 | 0.007 | 0.86 | | | |
| MIR939 | microRNA 939(MIR939) | NR_030635 | 137.4 | 183.9 | 0.0071 | 1.16 | | | |
| KLF10 | Kruppel like factor 10(KLF10) | NM_005655 | 154.5 | 164.7 | 0.0072 | 0.78 | | | |
| KLF10 | Kruppel like factor 10(KLF10) | NM_005655 | 192.8 | 164.2 | 0.0072 | 0.78 | NM_001009670 | 33.30 | 2.04 |
| PTK2 | protein tyrosine kinase 2(PTK2) | NM_153831 | 119 | 119.9 | 0.0072 | 0.9 | | | |
| TMEM184C | transmembrane protein 184C(TMEM184C) | NM_018241 | 110.5 | 106.7 | 0.0072 | 0.89 | | | |
| TMEM184C | transmembrane protein 184C(TMEM184C) | NM_018241 | 331.1 | 360.5 | 0.0072 | 0.89 | | | |
| RNF146 | ring finger protein 146(RNF146) | NM_030963 | 117.2 | 104.6 | 0.0073 | 0.82 | | | |
| RNF146 | ring finger protein 146(RNF146) | NM_030963 | 152.3 | 156.8 | 0.0073 | 0.82 | | | |

| | | | | | | |
|---|---|---|---|---|---|---|
| FAM174A | family with sequence similarity 174 member A(FAM174A) | NM_198507 | 283.1 | 227.5 | 0.0076 | 0.71 |
| SLC41A3 | solute carrier family 41 member 3(SLC41A3) | NM_001008485 | 612.9 | 375.5 | 0.0076 | 0.58 |
| TBC1D8B | TBC1 domain family member 8B(TBC1D8B) | NM_017752 | 113.6 | 121.8 | 0.0076 | 0.81 |
| TBC1D8B | TBC1 domain family member 8B(TBC1D8B) | NM_017752 | 176.8 | 157.7 | 0.0076 | 0.81 |
| RPL17P39 | ribosomal protein L17 pseudogene 39(RPL17P39) | NM_001093733 | 519.6 | 880.1 | 0.0078 | 1.57 |
| RRAGB | Ras related GTP binding B(RRAGB) | NM_006064 | 98.7 | 152.4 | 0.0079 | 1.36 |
| RRAGB | Ras related GTP binding B(RRAGB) | NM_006064 | 126.7 | 129.2 | 0.0079 | 1.36 |
| CNEP1R1 | CTD nuclear envelope phosphatase 1 regulatory subunit 1(CNEP1R1) | NM_153261 | 111.5 | 147.8 | 0.008 | 1.18 |
| ARHGAP1 | Rho GTPase activating protein 1(ARHGAP1) | NM_004308 | 1017.5 | 907.2 | 0.0081 | 0.83 |

| | | | | | | |
|---|---|---|---|---|---|---|
| ATP6V0D1 | ATPase H+ transporting V0 subunit d1(ATP6V0D1) | NM_004691 | 430.6 | 338.6 | 0.0083 | 0.73 |
| GAPDH | glyceraldehyde-3-phosphate dehydrogenase(GAPDH) | NM_002046 | 142.9 | 141.5 | 0.0085 | 0.43 |
| GAPDH | glyceraldehyde-3-phosphate dehydrogenase(GAPDH) | NM_002046 | 1842.1 | 1026.5 | 0.0085 | 0.43 |
| GAPDH | glyceraldehyde-3-phosphate dehydrogenase(GAPDH) | NM_002046 | 3640.2 | 1836.1 | 0.0085 | 0.43 |
| TYMS | thymidylate synthetase(TYMS) | NM_001071 | 113.5 | 232.8 | 0.0086 | 1.72 |
| FNTA | farnesyltransferase, CAAX box, alpha(FNTA) | NM_001018677 | 302.3 | 399.5 | 0.0089 | 1.21 |
| RPLP2 | ribosomal protein lateral stalk subunit P2(RPLP2) | NM_001004 | 10890.7 | 9676.9 | 0.009 | 0.66 |

| | | | | | | |
|---|---|---|---|---|---|---|
| HACE1 | HECT domain and ankyrin repeat containing E3 ubiquitin protein ligase 1(HACE1) | NM_020771 | 127.4 | 120.2 | 0.0091 | 0.86 |
| EIF1 | eukaryotic translation initiation factor 1(EIF1) | NM_005801 | 1591.4 | 202.6 | 0.0093 | 0.12 |
| PTS | 6-pyruvoyltetrahydropterin synthase(PTS) | NM_000317 | 316.4 | 243.3 | 0.0093 | 0.67 |
| TFDP1 | transcription factor Dp-1(TFDP1) | NM_007111 | 196.2 | 339.1 | 0.0093 | 1.52 |
| TFDP1 | transcription factor Dp-1(TFDP1) | NM_007111 | 492.5 | 636.7 | 0.0093 | 1.52 |
| | | XM_001718145 | 121.7 | 114.7 | 0.0094 | 0.86 |
| | | XM_927228 | 109.4 | 137.4 | 0.0095 | 1.11 |
| HNRNPH2 | heterogeneous nuclear ribonucleoprotein H2 (H')(HNRNPH2) | NM_019597 | 293.3 | 236.1 | 0.0096 | 0.74 |
| | | XM_938755 | 1571.1 | 2625.7 | 0.0097 | 1.49 |

| | | | | | | |
|---|---|---|---|---|---|---|
| AP2M1 | adaptor related protein complex 2 mu 1 subunit(AP2M1) | NM_004068 | 126.6 | 119.3 | 0.0098 | 0.86 |
| AP2M1 | adaptor related protein complex 2 mu 1 subunit(AP2M1) | NM_004068 | 271 | 288.9 | 0.0098 | 0.86 |
| LOC105377348 | uncharacterized LOC105377348(LOC105377348) | AK093356 | 153.6 | 138.2 | 0.0099 | 0.82 |
| STRAP | serine/threonine kinase receptor associated protein(STRAP) | NM_007178 | 186.5 | 321.8 | 0.01 | 1.5 |
| ANXA7 | annexin A7(ANXA7) | NM_004034 | 145 | 191.6 | 0.0101 | 2.02 |
| ANXA7 | annexin A7(ANXA7) | NM_004034 | 345.7 | 830.6 | 0.0101 | 2.02 |
| GHSR | growth hormone secretagogue receptor(GHSR) | NM_004122 | 103.9 | 133.9 | 0.0101 | 1.13 |
| PSMD5 | proteasome 26S subunit, non-ATPase 5(PSMD5) | NM_005047 | 163.1 | 234.6 | 0.0101 | 1.25 |
| | | XM_001721968 | 107.3 | 141.8 | 0.0101 | 1.15 |

| | | | | | | |
|---|---|---|---|---|---|---|
| SDR42E1 | short chain dehydrogenase/reductase family 42E, member 1(SDR42E1) | NM_145168 | 119.6 | 119.1 | 0.0105 | 0.9 |
| ZBTB40 | zinc finger and BTB domain containing 40(ZBTB40) | NM_001083621 | 148 | 234.5 | 0.0106 | 1.36 |
| | | XM_001722060 | 469.6 | 1000.1 | 0.0108 | 1.96 |
| | | XM_939687 | 1087.7 | 1810 | 0.0108 | 1.56 |
| USE1 | unconventional SNARE in the ER 1(USE1) | NM_018467 | 148.9 | 131.6 | 0.011 | 0.81 |
| | | XM_944813 | 121.9 | 121.3 | 0.011 | 0.9 |
| | | XR_040895 | 136.4 | 182.2 | 0.011 | 1.16 |
| | | XR_017960 | 91.5 | 123 | 0.0111 | 1.22 |
| ACD | adrenocortical dysplasia homolog(ACD) | NM_022914 | 137.6 | 185.5 | 0.0113 | 1.17 |
| ISG20L2 | interferon stimulated exonuclease gene 20 like 2(ISG20L2) | NM_030980 | 144.8 | 199.4 | 0.0115 | 1.2 |

| | | | | | | |
|---|---|---|---|---|---|---|
| RSRC2 | arginine and serine rich coiled-coil 2(RSRC2) | NM_023012 | 121.6 | 144.5 | 0.0115 | 1.29 |
| RSRC2 | arginine and serine rich coiled-coil 2(RSRC2) | NM_023012 | 656.4 | 918.9 | 0.0115 | 1.29 |
| C11orf74 | chromosome 11 open reading frame 74(C11orf74) | NM_138787 | 216.8 | 185.6 | 0.0116 | 0.76 |
| PUM2 | pumilio RNA binding family member 2(PUM2) | NM_015317 | 458.7 | 667.5 | 0.0118 | 1.33 |
| ANKRA2 | ankyrin repeat family A member 2(ANKRA2) | NM_023039 | 116.8 | 150 | 0.012 | 1.14 |
| ANKRA2 | ankyrin repeat family A member 2(ANKRA2) | NM_023039 | 177.1 | 229.6 | 0.012 | 1.14 |
| fxyd2 | FXYD domain containing ion transport regulator 2(FXYD2) | NM_001680 | 128.4 | 146.9 | 0.012 | 0.46 |
| fxyd2 | FXYD domain containing ion transport regulator 2(FXYD2) | NM_001680 | 12803.3 | 6710.8 | 0.012 | 0.46 |

| | | | | | | | | | |
|---|---|---|---|---|---|---|---|---|---|
| OTUB1 | OTU deubiquitinase, ubiquitin aldehyde binding 1(OTUB1) | NR_003089 | 148.1 | 139.2 | 0.012 | 0.84 | | | |
| MRAP2 | melanocortin 2 receptor accessory protein 2(MRAP2) | NM_138409 | 141.4 | 116.5 | 0.0122 | 0.76 | | | |
| MRAP2 | melanocortin 2 receptor accessory protein 2(MRAP2) | NM_138409 | 161.3 | 174.9 | 0.0122 | 0.76 | | | |
| RPS6 | ribosomal protein S6(RPS6) | NM_001010 | 2996.1 | 6286.2 | 0.0122 | 1.6 | | | |
| RPS6 | ribosomal protein S6(RPS6) | NM_001010 | 6172 | 8646.3 | 0.0122 | 1.6 | | | |
| TRIM36 | tripartite motif containing 36(TRIM36) | NM_001017397 | 103.5 | 133 | 0.0122 | 1.14 | | | |
| PNMA1 | paraneoplastic Ma antigen 1(PNMA1) | NM_006029 | 322.8 | 245.5 | 0.0123 | 0.7 | | | |
| oxtr | oxytocin receptor(OXTR) | NM_000916 | 118.9 | 112.8 | 0.0124 | 0.86 | | | |
| PWWP2B | PWWP domain containing 2B(PWWP2B) | NM_138499 | 116.1 | 107 | 0.0128 | 0.84 | NM_001107943 | 0.63 | 112.77 |
| PWWP2B | PWWP domain containing 2B(PWWP2B) | NM_138499 | 143 | 161.6 | 0.0128 | 0.84 | | | |

| | | | | | | |
|---|---|---|---|---|---|---|
| FAM127B | family with sequence similarity 127 member B(FAM127B) | NM_001078172 | 157.9 | 178.2 | 0.0129 | 0.71 |
| FAM127B | family with sequence similarity 127 member B(FAM127B) | NM_001078172 | 299.4 | 236.3 | 0.0129 | 0.71 |
| | | XM_001726456 | 115.4 | 155 | 0.0129 | 1.16 |
| GNG10 | G protein subunit gamma 10(GNG10) | NM_001017998 | 897.8 | 460.7 | 0.0131 | 0.53 |
| SLC25A4 | solute carrier family 25 member 4(SLC25A4) | NM_001151 | 1239.8 | 632.8 | 0.0131 | 0.43 |
| SLC25A4 | solute carrier family 25 member 4(SLC25A4) | NM_001151 | 2794.8 | 1377.6 | 0.0131 | 0.43 |
| GPATCH8 | G-patch domain containing 8(GPATCH8) | NM_001002909 | 120.3 | 158.8 | 0.0132 | 1.17 |
| STX8 | syntaxin 8(STX8) | NM_004853 | 462.3 | 367.9 | 0.0133 | 0.71 |
| | | XR_015954 | 126 | 195.4 | 0.0134 | 1.36 |
| MINK1 | misshapen like kinase 1(MINK1) | NM_153827 | 118.2 | 113.2 | 0.0135 | 0.87 |

| | | | | | | | | | |
|---|---|---|---|---|---|---|---|---|---|
| MINK1 | misshapen like kinase 1(MINK1) | NM_153827 | 118.7 | 138.1 | 0.0135 | 0.87 | NM_134372 | 22.51 | 0.03 |
| GPR34 | G protein-coupled receptor 34(GPR34) | NM_001033513 | 123.6 | 167.1 | 0.0136 | 1.17 | | | |
| MZB1 | marginal zone B and B1 cell specific protein(MZB1) | NM_016459 | 92.9 | 112 | 0.0136 | 2.51 | | | |
| MZB1 | marginal zone B and B1 cell specific protein(MZB1) | NM_016459 | 179.2 | 514.5 | 0.0136 | 2.51 | | | |
| PGA3 | pepsinogen 3, group I (pepsinogen A)(PGA3) | NM_001079807 | 95.5 | 110.5 | 0.0137 | 1.17 | | | |
| PGA3 | pepsinogen 3, group I (pepsinogen A)(PGA3) | NM_001079807 | 110.8 | 120.2 | 0.0137 | 1.17 | | | |
| PGA3 | pepsinogen 3, group I (pepsinogen A)(PGA3) | NM_001079807 | 111.8 | 150.6 | 0.0137 | 1.17 | | | |
| | | XR_017249 | 103.6 | 146.2 | 0.0137 | 1.25 | | | |

| | | | | | | |
|---|---|---|---|---|---|---|
| CEACAM1 | carcinoembryonic antigen related cell adhesion molecule 1(CEACAM1) | NM_001712 | 120 | 110 | 0.0141 | 0.83 |
| GPRC5C | G protein-coupled receptor class C group 5 member C(GPRC5C) | NM_018653 | 427.4 | 290.7 | 0.0141 | 0.61 |
| LINC01278 | long intergenic non-protein coding RNA 1278(LINC01278) | AK130294 | 290.9 | 239.4 | 0.0141 | 0.73 |
| BUB3 | BUB3, mitotic checkpoint protein(BUB3) | NM_004725 | 182.1 | 251.4 | 0.0142 | 1.73 |
| BUB3 | BUB3, mitotic checkpoint protein(BUB3) | NM_004725 | 261.4 | 499.8 | 0.0142 | 1.73 |
| LIG3 | DNA ligase 3(LIG3) | NM_013975 | 160.3 | 207.6 | 0.0143 | 1.12 |
| FAM96B | family with sequence similarity 96 member B(FAM96B) | NM_016062 | 415.4 | 361.1 | 0.0144 | 0.79 |
| RPL26P30 | ribosomal protein L26 pseudogene 30(RPL26P30) | NM_001093761 | 366.6 | 296 | 0.0145 | 0.73 |

| | | | | | | | | | |
|---|---|---|---|---|---|---|---|---|---|
| FAM127A | family with sequence similarity 127 member A(FAM127A) | NM_001078171 | 363 | 277.5 | 0.0146 | 0.7 | | | |
| GLS2 | glutaminase 2(GLS2) | NM_013267 | 122 | 118.8 | 0.0146 | 0.89 | | | |
| AKT3 | AKT serine/threonine kinase 3(AKT3) | NM_005465 | 91.9 | 117.9 | 0.0147 | 1.18 | | | |
| OPRM1 | opioid receptor mu 1(OPRM1) | NM_001008505 | 116.7 | 144.5 | 0.0147 | 1.09 | | | |
| OPRM1 | opioid receptor mu 1(OPRM1) | NM_001008505 | 177.5 | 216.2 | 0.0147 | 1.09 | | | |
| RPL19P9 | ribosomal protein L19 pseudogene 9(RPL19P9) | NM_001080544 | 429.2 | 660.6 | 0.0147 | 1.38 | | | |
| RPL19P9 | ribosomal protein L19 pseudogene 9(RPL19P9) | NM_001080544 | 4171.5 | 6597.3 | 0.0147 | 1.38 | | | |
| | | XM_001726945 | 106.8 | 105.8 | 0.0147 | 0.9 | NM_001031645 | 1.93 | 10.04 |
| RAB11FIP1 | RAB11 family interacting protein 1(RAB11FIP1) | NM_001002814 | 459.8 | 697.6 | 0.0148 | 1.39 | | | |
| E2F4 | E2F transcription factor 4(E2F4) | NM_001950 | 289.3 | 407.9 | 0.0149 | 1.27 | | | |
| | | XR_037945 | 99.3 | 139.2 | 0.0149 | 1.23 | | | |

| | | | | | | |
|---|---|---|---|---|---|---|
| PHOSPHO2 | phosphatase, orphan 2(PHOSPHO2) | NM_001008489 | 299 | 208 | 0.015 | 0.65 |
| ADGRE5 | adhesion G protein-coupled receptor E5(ADGRE5) | NM_078481 | 103.5 | 130.9 | 0.0155 | 1.12 |
| ADGRE5 | adhesion G protein-coupled receptor E5(ADGRE5) | NM_078481 | 107.1 | 124.5 | 0.0155 | 1.12 |
| | | XM_001133677 | 94.8 | 106.4 | 0.0155 | 1.13 |
| | | XM_001133677 | 106.8 | 138.9 | 0.0155 | 1.13 |
| | | BM987531 | 112.8 | 142.2 | 0.0156 | 1.11 |
| FAM96A | family with sequence similarity 96 member A(FAM96A) | NM_032231 | 460 | 369.9 | 0.0157 | 0.75 |
| PLCD4 | phospholipase C delta 4(PLCD4) | NM_032726 | 242 | 179.8 | 0.0158 | 0.65 |
| ASB8 | ankyrin repeat and SOCS box containing 8(ASB8) | NM_024095 | 118.5 | 120 | 0.0159 | 0.91 |
| ASB8 | ankyrin repeat and SOCS box containing 8(ASB8) | NM_024095 | 121.6 | 122.4 | 0.0159 | 0.91 |

| | | | | | | | | | |
|---|---|---|---|---|---|---|---|---|---|
| P2RY2 | purinergic receptor P2Y2(P2RY2) | NM_002564 | 185.8 | 141.2 | 0.0159 | 0.67 | | | |
| | | XM_001723699 | 112.9 | 142.5 | 0.0159 | 1.12 | NM_144748 | 66.99 | 0.14 |
| | | XR_038699 | 1022.6 | 1448.6 | 0.0159 | 1.31 | | | |
| ATP6V0E1 | ATPase H+ transporting V0 subunit e1(ATP6V0E1) | NM_003945 | 970.5 | 897.5 | 0.0162 | 0.85 | | | |
| ATP6V0E1 | ATPase H+ transporting V0 subunit e1(ATP6V0E1) | NM_003945 | 1668.7 | 1759 | 0.0162 | 0.85 | | | |
| GDI2 | GDP dissociation inhibitor 2(GDI2) | NM_001494 | 146.7 | 216 | 0.0163 | 1.3 | | | |
| MRPL20 | mitochondrial ribosomal protein L20(MRPL20) | NM_017971 | 482 | 407.1 | 0.0164 | 0.77 | | | |
| ZNF622 | zinc finger protein 622(ZNF622) | NM_033414 | 482 | 463.4 | 0.0164 | 0.87 | | | |
| | | XM_001714310 | 100.6 | 126 | 0.0164 | 1.12 | | | |
| STT3A | STT3A, catalytic subunit of the oligosaccharyltransferase complex(STT3A) | NM_152713 | 482.8 | 723.5 | 0.0165 | 1.38 | NM_001107793 | 77.29 | 6.85 |
| | | XR_019270 | 119.8 | 157.2 | 0.0165 | 1.13 | NM_001107793 | 77.29 | 6.85 |

| | | | | | | | | | |
|---|---|---|---|---|---|---|---|---|---|
| | | XR_039539 | 113.8 | 109.9 | 0.0165 | 0.88 | NM_001107793 | 77.29 | 6.85 |
| HOMER2 | homer scaffolding protein 2(HOMER2) | NM_199332 | 212.4 | 171.6 | 0.0169 | 0.73 | | | |
| BTF3 | basic transcription factor 3(BTF3) | NM_001207 | 1296.2 | 614.3 | 0.0172 | 0.49 | | | |
| SCARNA20 | small Cajal body-specific RNA 20(SCARNA20) | NR_002999 | 104.2 | 131.7 | 0.0172 | 1.13 | | | |
| SNORA1 | small nucleolar RNA, H/ACA box 1(SNORA1) | NR_003026 | 100.9 | 128.2 | 0.0173 | 1.14 | | | |
| PSMB11 | proteasome subunit beta 11(PSMB11) | NM_001099780 | 108.5 | 138.9 | 0.0178 | 1.13 | | | |
| STEAP2 | STEAP2 metalloreductase(STEAP2) | NM_152999 | 127.7 | 115.4 | 0.0178 | 0.72 | | | |
| STEAP2 | STEAP2 metalloreductase(STEAP2) | NM_152999 | 201.8 | 157.9 | 0.0178 | 0.72 | | | |
| GLMP | glycosylated lysosomal membrane protein(GLMP) | NM_144580 | 845.9 | 613.6 | 0.0179 | 0.66 | | | |
| CYBB | cytochrome b-245 beta chain(CYBB) | NM_000397 | 106.7 | 152.9 | 0.0183 | 1.26 | | | |
| RPL14 | ribosomal protein L14(RPL14) | NM_001034996 | 864.7 | 1435.9 | 0.0184 | 1.51 | | | |

| | | | | | | |
|---|---|---|---|---|---|---|
| TIGD6 | tigger transposable element derived 6(TIGD6) | NM_030953 | 121.5 | 122.8 | 0.0184 | 0.91 |
| MRPL43 | mitochondrial ribosomal protein L43(MRPL43) | NM_032112 | 496.5 | 372.2 | 0.0186 | 0.68 |
| MRPL43 | mitochondrial ribosomal protein L43(MRPL43) | NM_032112 | 935.3 | 791.2 | 0.0186 | 0.68 |
| OXA1L | OXA1L, mitochondrial inner membrane protein(OXA1L) | NM_005015 | 738 | 478.1 | 0.0186 | 0.59 |
| | | AI911349 | 136.7 | 183.5 | 0.0186 | 1.16 |
| | | XR_037977 | 780.1 | 1280.1 | 0.0186 | 1.53 |
| ATG4B | autophagy related 4B cysteine peptidase(ATG4B) | NM_178326 | 173.9 | 164.2 | 0.0187 | 0.83 |
| HPX | hemopexin(HPX) | NM_000613 | 145.1 | 116.2 | 0.0187 | 0.72 |
| RAPGEF1 | Rap guanine nucleotide exchange factor 1(RAPGEF1) | NM_005312 | 136.4 | 177.8 | 0.0188 | 1.14 |
| DEFB103A | defensin beta 103A(DEFB103A) | NM_001081551 | 104 | 136.2 | 0.019 | 1.15 |

| | | | | | | |
|---|---|---|---|---|---|---|
| DOLK | dolichol kinase(DOLK) | NM_014908 | 218.9 | 191 | 0.019 | 0.8 |
| RAD51AP1 | RAD51 associated protein 1(RAD51AP1) | NM_006479 | 99.3 | 126.4 | 0.019 | 1.15 |
| RTN4IP1 | reticulon 4 interacting protein 1(RTN4IP1) | NM_032730 | 199.8 | 190.4 | 0.019 | 0.84 |
| METTL4 | methyltransferase like 4(METTL4) | NM_022840 | 160.5 | 227.8 | 0.0191 | 1.21 |
| SERPINB1 | serpin family B member 1(SERPINB1) | NM_030666 | 99.8 | 130.8 | 0.0192 | 1.16 |
| | | XM_934839 | 104 | 138.5 | 0.0192 | 1.17 |
| FGD1 | FYVE, RhoGEF and PH domain containing 1(FGD1) | NM_004463 | 118.3 | 115.7 | 0.0193 | 0.88 |
| BCR | BCR, RhoGEF and GTPase activating protein(BCR) | NM_004327 | 137 | 157.7 | 0.0196 | 0.74 |
| BCR | BCR, RhoGEF and GTPase activating protein(BCR) | NM_004327 | 215.7 | 175.7 | 0.0196 | 0.74 |

| | | | | | | | | | |
|---|---|---|---|---|---|---|---|---|---|
| LINC00544 | long intergenic non-protein coding RNA 544(LINC00544) | NM_001099778 | 116 | 160.1 | 0.0196 | 1.2 | | | |
| | | XM_934553 | 100 | 124.9 | 0.0197 | 1.11 | | | |
| MYH9 | myosin heavy chain 9(MYH9) | NM_002473 | 806.9 | 2308.3 | 0.02 | 2.53 | NM_001009603 | 28.32 | 0.90 |
| MYH9 | myosin heavy chain 9(MYH9) | NM_002473 | 2233.6 | 3165.4 | 0.02 | 2.53 | | | |
| | | XM_925919 | 112 | 108.5 | 0.02 | 0.88 | | | |
| KCNJ13 | potassium voltage-gated channel subfamily J member 13(KCNJ13) | NM_002242 | 107.3 | 104.4 | 0.0201 | 0.89 | | | |
| STXBP6 | syntaxin binding protein 6(STXBP6) | NM_014178 | 151.4 | 138.1 | 0.0201 | 0.82 | | | |
| | | XR_038898 | 160.3 | 250 | 0.0204 | 1.42 | | | |
| ZNF684 | zinc finger protein 684(ZNF684) | NM_152373 | 97.3 | 127.8 | 0.0207 | 1.17 | | | |
| Tas2r43 | taste 2 receptor member 43(TAS2R43) | NM_176884 | 113 | 110.3 | 0.0208 | 0.89 | | | |
| TRAPPC13 | trafficking protein particle complex 13(TRAPPC13) | NM_001093756 | 201.4 | 162.3 | 0.0209 | 0.72 | | | |

| | | | | | | |
|---|---|---|---|---|---|---|
| | | BQ185080 | 112.1 | 145.6 | 0.0209 | 1.13 |
| | | CR748464 | 101.1 | 127.6 | 0.0209 | 1.12 |
| | | XR_017653 | 1485 | 2142.3 | 0.021 | 1.28 |
| ZFAND6 | zinc finger AN1-type containing 6(ZFAND6) | NM_019006 | 257.2 | 254.4 | 0.0212 | 0.75 |
| ZFAND6 | zinc finger AN1-type containing 6(ZFAND6) | NM_019006 | 509.7 | 405.2 | 0.0212 | 0.75 |
| | | XR_018412 | 106.9 | 132.9 | 0.0213 | 1.11 |
| HERPUD2 | HERPUD family member 2(HERPUD2) | NM_022373 | 109.7 | 139.4 | 0.0214 | 1.13 |
| HERPUD2 | HERPUD family member 2(HERPUD2) | NM_022373 | 120.7 | 132.2 | 0.0214 | 1.13 |
| CMC1 | C-X9-C motif containing 1(CMC1) | NM_182523 | 274.2 | 215.2 | 0.0217 | 0.69 |
| CLEC11A | C-type lectin domain family 11 member A(CLEC11A) | NM_002975 | 176 | 156.4 | 0.0218 | 0.78 |
| TPI1P2 | triosephosphate isomerase 1 pseudogene 2(TPI1P2) | NR_002187 | 143.8 | 133.8 | 0.0218 | 0.84 |

| | | | | | | |
|---|---|---|---|---|---|---|
| UQCC1 | ubiquinol-cytochrome c reductase complex assembly factor 1(UQCC1) | NM_199487 | 217.1 | 198.8 | 0.022 | 0.79 |
| | | DB299679 | 123.7 | 117.3 | 0.0223 | 0.86 |
| ADCK1 | aarF domain containing kinase 1(ADCK1) | NM_020421 | 153.3 | 150.7 | 0.0224 | 0.87 |
| SH3TC1 | SH3 domain and tetratricopeptide repeats 1(SH3TC1) | NM_018986 | 539.6 | 360.7 | 0.0224 | 0.62 |
| ANGEL1 | angel homolog 1(ANGEL1) | NM_015305 | 166.5 | 276 | 0.0226 | 1.45 |
| MACROD1 | MACRO domain containing 1(MACROD1) | NM_014067 | 552.7 | 291.7 | 0.0226 | 0.5 |
| TMEM185A | transmembrane protein 185A(TMEM185A) | NM_032508 | 144.5 | 162.4 | 0.0227 | 0.83 |
| TMEM185A | transmembrane protein 185A(TMEM185A) | NM_032508 | 200.1 | 184.3 | 0.0227 | 0.83 |
| PSMF1 | proteasome inhibitor subunit 1(PSMF1) | NM_178579 | 377.6 | 336.1 | 0.0228 | 0.79 |

| | | | | | | |
|---|---|---|---|---|---|---|
| | | XM_495909 | 105.7 | 134.6 | 0.0228 | 1.14 |
| csnk1g3 | casein kinase 1 gamma 3(CSNK1G3) | NM_001031812 | 142.5 | 190.8 | 0.0229 | 1.17 |
| ZNF385C | zinc finger protein 385C(ZNF385C) | NM_001013624 | 146.1 | 209.7 | 0.0229 | 1.23 |
| | | XM_942071 | 137.4 | 190.7 | 0.0229 | 1.21 |
| NOTUM | NOTUM, palmitoleoyl-protein carboxylesterase(NOTUM) | NM_178493 | 113.6 | 115 | 0.023 | 0.91 |
| RPAIN | RPA interacting protein(RPAIN) | NM_001033002 | 326.6 | 295.8 | 0.023 | 0.8 |
| CHRNA6 | cholinergic receptor nicotinic alpha 6 subunit(CHRNA6) | NM_004198 | 114.7 | 107.6 | 0.0232 | 0.86 |
| GORASP1 | golgi reassembly stacking protein 1(GORASP1) | NM_031899 | 169.5 | 129.7 | 0.0236 | 0.71 |
| | | XM_001725669 | 640.4 | 415.7 | 0.0237 | 0.61 |
| EPB41L1 | erythrocyte membrane protein band 4.1 like 1(EPB41L1) | NM_012156 | 326.1 | 273.9 | 0.024 | 0.74 |

| | | | | | | |
|---|---|---|---|---|---|---|
| | | XM_926231 | 156.3 | 244.2 | 0.024 | 1.38 |
| ARCN1 | archain 1(ARCN1) | NM_001655 | 305 | 433.4 | 0.0241 | 1.29 |
| ARRDC4 | arrestin domain containing 4(ARRDC4) | NM_183376 | 90.1 | 120.7 | 0.0241 | 1.21 |
| | | XR_016385 | 258.3 | 386.7 | 0.0241 | 1.35 |
| RPL32 | ribosomal protein L32(RPL32) | NM_001007074 | 102.7 | 118.6 | 0.0242 | 1.59 |
| RPL32 | ribosomal protein L32(RPL32) | NM_001007074 | 1495.3 | 2663.2 | 0.0242 | 1.59 |
| SPACA9 | sperm acrosome associated 9(SPACA9) | NM_018956 | 157.3 | 159.6 | 0.0242 | 0.89 |
| BBC3 | BCL2 binding component 3(BBC3) | NM_014417 | 180.4 | 179.1 | 0.0245 | 0.89 |
| LYPD6 | LY6/PLAUR domain containing 6(LYPD6) | NM_194317 | 98.1 | 121.9 | 0.0245 | 1.13 |
| | | AW665973 | 268.9 | 505.5 | 0.0246 | 1.58 |
| ADAT1 | adenosine deaminase, tRNA specific 1(ADAT1) | NM_012091 | 116.9 | 118.7 | 0.0249 | 0.92 |
| CTSF | cathepsin F(CTSF) | NM_003793 | 559.1 | 489.3 | 0.0249 | 0.81 |

| | | | | | | |
|---|---|---|---|---|---|---|
| PTPN3 | protein tyrosine phosphatase, non-receptor type 3(PTPN3) | NM_002829 | 338.6 | 222.4 | 0.0251 | 0.6 |
| PDGFA | platelet derived growth factor subunit A(PDGFA) | NM_033023 | 167.2 | 149.9 | 0.0252 | 0.8 |
| | | XR_016832 | 153.2 | 224.8 | 0.0252 | 1.28 |
| NDUFA11 | NADH:ubiquinone oxidoreductase subunit A11(NDUFA11) | NM_175614 | 1766.9 | 1305.1 | 0.0254 | 0.67 |
| PLA1A | phospholipase A1 member A(PLA1A) | NM_015900 | 349.7 | 213.4 | 0.0254 | 0.57 |
| TMEM189-UBE2V1 | TMEM189-UBE2V1 readthrough(TMEM189-UBE2V1) | NM_003349 | 236.2 | 171.9 | 0.0254 | 0.69 |
| MRPL34 | mitochondrial ribosomal protein L34(MRPL34) | NM_023937 | 399.5 | 384.9 | 0.0256 | 0.87 |
| PSMC1 | proteasome 26S subunit, ATPase 1(PSMC1) | NM_002802 | 1659.8 | 1339.7 | 0.0256 | 0.73 |
| PSMC1 | proteasome 26S subunit, ATPase 1(PSMC1) | NM_002802 | 1673.9 | 1645.6 | 0.0256 | 0.73 |

| | | | | | | |
|---|---|---|---|---|---|---|
| SIRPG | signal regulatory protein gamma(SIRPG) | NM_001039508 | 102.8 | 157.2 | 0.0256 | 1.33 |
| CHCHD6 | coiled-coil-helix-coiled-coil-helix domain containing 6(CHCHD6) | NM_032343 | 302.2 | 248.4 | 0.0259 | 0.73 |
| fgl1 | fibrinogen like 1(FGL1) | NM_201553 | 167.9 | 119.9 | 0.0259 | 0.65 |
| | | XM_944079 | 122.9 | 164.4 | 0.026 | 1.16 |
| Cox4I1 | cytochrome c oxidase subunit 4I1(COX4I1) | NM_001861 | 10161.5 | 7591.4 | 0.0261 | 0.63 |
| FAM131B | family with sequence similarity 131 member B(FAM131B) | NM_014690 | 107.2 | 134.9 | 0.0261 | 1.11 |
| HSPBP1 | HSPA (Hsp70) binding protein 1(HSPBP1) | NM_012267 | 110.4 | 114.8 | 0.0261 | 0.87 |
| HSPBP1 | HSPA (Hsp70) binding protein 1(HSPBP1) | NM_012267 | 121.7 | 115.7 | 0.0261 | 0.87 |
| TCEA3 | transcription elongation factor A3(TCEA3) | NM_003196 | 704.6 | 419.6 | 0.0261 | 0.53 |

| | | | | | | |
|---|---|---|---|---|---|---|
| RTFDC1 | replication termination factor 2 domain containing 1(RTFDC1) | NM_016407 | 474.5 | 421.8 | 0.0262 | 0.79 |
| TRAF4 | TNF receptor associated factor 4(TRAF4) | NM_004295 | 117.9 | 112.6 | 0.0262 | 0.87 |
| ALDOA | aldolase, fructose-bisphosphate A(ALDOA) | NM_184043 | 4490.9 | 1955.6 | 0.0263 | 0.36 |
| | | XM_945148 | 123.5 | 162.6 | 0.0263 | 1.16 |
| | | XR_019429 | 141.7 | 241.6 | 0.0264 | 1.51 |
| GLUD1 | glutamate dehydrogenase 1(GLUD1) | NM_005271 | 111.9 | 110.9 | 0.0265 | 0.9 |
| PIGL | phosphatidylinositol glycan anchor biosynthesis class L(PIGL) | NM_004278 | 104.4 | 128.1 | 0.0265 | 1.1 |
| ANKS1B | ankyrin repeat and sterile alpha motif domain containing 1B(ANKS1B) | NM_020140 | 112.3 | 111 | 0.0266 | 0.91 |
| slc35e2b | solute carrier family 35 member E2B(SLC35E2B) | NM_001110781 | 118.7 | 155 | 0.0267 | 1.15 |
| | | AK002088 | 91.1 | 116.5 | 0.0267 | 1.15 |

| | | | | | | | | | |
|---|---|---|---|---|---|---|---|---|---|
| MSMO1 | methylsterol monooxygenase 1(MSMO1) | NM_001017369 | 108.3 | 108.6 | 0.0268 | 0.91 | | | |
| MSMO1 | methylsterol monooxygenase 1(MSMO1) | NM_001017369 | 177.3 | 230.5 | 0.0268 | 0.91 | | | |
| | | NR_003252 | 153.5 | 152.5 | 0.0268 | 0.88 | | | |
| | | XM_926590 | 99.3 | 128.8 | 0.0268 | 1.17 | | | |
| GC | GC, vitamin D binding protein(GC) | NM_000583 | 172 | 110.3 | 0.027 | 0.58 | | | |
| ESRRAP2 | estrogen-related receptor alpha pseudogene 2(ESRRAP2) | NR_000033 | 430.1 | 343.1 | 0.0271 | 0.71 | | | |
| SEH1L | SEH1 like nucleoporin(SEH1L) | NM_001013437 | 149 | 209.3 | 0.0271 | 1.25 | | | |
| SEH1L | SEH1 like nucleoporin(SEH1L) | NM_001013437 | 245.1 | 273.8 | 0.0271 | 1.25 | NM_144744 | 0.11 | 40.01 |
| | | XM_001727022 | 145.8 | 132.9 | 0.0271 | 0.83 | | | |
| GORAB | golgin, RAB6 interacting(GORAB) | NM_152281 | 114.4 | 140.1 | 0.0272 | 1.09 | | | |
| CORO2B | coronin 2B(CORO2B) | NM_006091 | 102.2 | 114.1 | 0.0273 | 0.83 | | | |

| | | | | | | |
|---|---|---|---|---|---|---|
| CORO2B | coronin 2B(CORO2B) | NM_006091 | 124 | 113.4 | 0.0273 | 0.83 |
| TMEM206 | transmembrane protein 206(TMEM206) | NM_018252 | 105.7 | 98.3 | 0.0274 | 0.86 |
| | | XM_940962 | 170 | 155.8 | 0.0274 | 0.82 |
| CCDC106 | coiled-coil domain containing 106(CCDC106) | NM_013301 | 349 | 317.4 | 0.0276 | 0.81 |
| | | XM_001720172 | 494.9 | 722 | 0.0276 | 1.33 |
| JADE1 | jade family PHD finger 1(JADE1) | NM_024900 | 215 | 199 | 0.0277 | 0.82 |
| ZRANB1 | zinc finger RANBP2-type containing 1(ZRANB1) | NM_017580 | 156 | 154.2 | 0.0277 | 0.87 |
| DNAJC27 | DnaJ heat shock protein family (Hsp40) member C27(DNAJC27) | NM_016544 | 297.1 | 260.3 | 0.0278 | 0.76 |
| | | XM_001714755 | 167.8 | 136.1 | 0.0278 | 0.73 |
| ICAM4 | intercellular adhesion molecule 4 (Landsteiner-Wiener blood group)(ICAM4) | NM_022377 | 112.3 | 107.3 | 0.028 | 0.88 |

| | | | | | | |
|---|---|---|---|---|---|---|
| | | XR_038602 | 1134.2 | 1738.1 | 0.0282 | 1.42 |
| HOXD3 | homeobox D3(HOXD3) | NM_006898 | 203.3 | 175.7 | 0.0285 | 0.79 |
| SERPINC1 | serpin family C member 1(SERPINC1) | NM_000488 | 329.7 | 175.3 | 0.0285 | 0.46 |
| LIN7B | lin-7 homolog B, crumbs cell polarity complex component(LIN7B) | NM_022165 | 140.9 | 131.4 | 0.0287 | 0.82 |
| HOXA11 | homeobox A11(HOXA11) | NM_005523 | 109 | 131.5 | 0.0288 | 1.08 |
| PDGFB | platelet derived growth factor subunit B(PDGFB) | NM_033016 | 126.5 | 123.4 | 0.0288 | 0.88 |
| CYB561A3 | cytochrome b561 family member A3(CYB561A3) | NM_153611 | 201.9 | 405.9 | 0.0289 | 1.77 |
| KREMEN2 | kringle containing transmembrane protein 2(KREMEN2) | NM_024507 | 113.8 | 111.5 | 0.0291 | 0.89 |
| PAXIP1 | PAX interacting protein 1(PAXIP1) | NM_007349 | 171.7 | 237.5 | 0.0292 | 1.21 |
| TMEM67 | transmembrane protein 67(TMEM67) | NM_153704 | 116.7 | 111.4 | 0.0292 | 0.88 |

| | | | | | | |
|---|---|---|---|---|---|---|
| CADM4 | cell adhesion molecule 4(CADM4) | NM_145296 | 330.3 | 222.6 | 0.0293 | 0.63 |
| | | XM_944183 | 114.3 | 138.9 | 0.0294 | 1.08 |
| IGSF9B | immunoglobulin superfamily member 9B(IGSF9B) | NM_014987 | 111.1 | 141.8 | 0.0295 | 1.13 |
| PLAA | phospholipase A2 activating protein(PLAA) | NM_001031689 | 91.5 | 118.1 | 0.0295 | 1.16 |
| PLAA | phospholipase A2 activating protein(PLAA) | NM_001031689 | 114.9 | 126.3 | 0.0295 | 1.16 |
| SNX11 | sorting nexin 11(SNX11) | NM_152244 | 376 | 508 | 0.0295 | 1.21 |
| PLBD2 | phospholipase B domain containing 2(PLBD2) | NM_173542 | 224 | 210.6 | 0.0296 | 0.84 |
| TUBA3FP | tubulin alpha 3f pseudogene(TUBA3FP) | NR_003608 | 120.7 | 122.8 | 0.0296 | 0.91 |
| CENPA | centromere protein A(CENPA) | NM_001042426 | 98.7 | 122.8 | 0.0297 | 1.14 |
| CENPA | centromere protein A(CENPA) | NM_001042426 | 104.5 | 135.1 | 0.0297 | 1.14 |

| | | | | | | |
|---|---|---|---|---|---|---|
| | | XR_019267 | 129.8 | 178.9 | 0.0297 | 1.19 |
| SLC26A10 | solute carrier family 26 member 10(SLC26A10) | NM_133489 | 110.1 | 99.9 | 0.0298 | 0.84 |
| SLC26A10 | solute carrier family 26 member 10(SLC26A10) | NM_133489 | 125.5 | 144.5 | 0.0298 | 0.84 |
| SRP14 | signal recognition particle 14(SRP14) | NM_003134 | 257.6 | 405.8 | 0.03 | 0.72 |
| SRP14 | signal recognition particle 14(SRP14) | NM_003134 | 4633.3 | 3832.9 | 0.03 | 0.72 |
| hamp | hepcidin antimicrobial peptide(HAMP) | NM_021175 | 136.1 | 129.4 | 0.0301 | 0.84 |
| KIF16B | kinesin family member 16B(KIF16B) | NM_024704 | 111.6 | 111.5 | 0.0302 | 0.81 |
| KIF16B | kinesin family member 16B(KIF16B) | NM_024704 | 136.6 | 123.9 | 0.0302 | 0.81 |
| MAN1C1 | mannosidase alpha class 1C member 1(MAN1C1) | NM_020379 | 589.2 | 269 | 0.0302 | 0.41 |
| DDX19A | DEAD-box helicase 19A(DDX19A) | NM_018332 | 119 | 127.9 | 0.0303 | 1.17 |

| | | | | | | |
|---|---|---|---|---|---|---|
| DDX19A | DEAD-box helicase 19A(DDX19A) | NM_018332 | 261.7 | 346.5 | 0.0303 | 1.17 |
| SSTR3 | somatostatin receptor 3(SSTR3) | NM_001051 | 107.6 | 105.6 | 0.0303 | 0.9 |
| ATP5J | ATP synthase, H+ transporting, mitochondrial Fo complex subunit F6(ATP5J) | NM_001003701 | 4092.1 | 3324.4 | 0.0304 | 0.69 |
| | | AA658833 | 108 | 104.5 | 0.0304 | 0.89 |
| | | BM684558 | 101.2 | 128.4 | 0.0304 | 1.12 |
| ASAH1 | N-acylsphingosine amidohydrolase 1(ASAH1) | NM_177924 | 122.3 | 125.4 | 0.0307 | 0.92 |
| PSMB5 | proteasome subunit beta 5(PSMB5) | NM_002797 | 1255.4 | 1083.9 | 0.0307 | 0.78 |
| RRAGD | Ras related GTP binding D(RRAGD) | NM_021244 | 219.1 | 161.2 | 0.0307 | 0.68 |
| SARM1 | sterile alpha and TIR motif containing 1(SARM1) | NM_015077 | 106.1 | 131.7 | 0.0308 | 1.1 |
| | | XR_038738 | 230.6 | 359.9 | 0.0308 | 1.41 |

| | | | | | | |
|---|---|---|---|---|---|---|
| DDAH2 | dimethylarginine dimethylaminohydrolase 2(DDAH2) | NM_013974 | 244.6 | 204.9 | 0.031 | 0.76 |
| DTD2 | D-tyrosyl-tRNA deacylase 2 (putative)(DTD2) | NM_080664 | 119.2 | 112 | 0.031 | 0.84 |
| NME6 | NME/NM23 nucleoside diphosphate kinase 6(NME6) | NM_005793 | 114.3 | 147.2 | 0.031 | 1.13 |
| NME6 | NME/NM23 nucleoside diphosphate kinase 6(NME6) | NM_005793 | 139.1 | 168.6 | 0.031 | 1.13 |
| | | XR_015351 | 120 | 119.9 | 0.0312 | 0.89 |
| NMRK2 | nicotinamide riboside kinase 2(NMRK2) | NM_170678 | 104 | 92.5 | 0.0313 | 0.83 |
| NMRK2 | nicotinamide riboside kinase 2(NMRK2) | NM_170678 | 118.8 | 123.2 | 0.0313 | 0.83 |
| PDE6H | phosphodiesterase 6H(PDE6H) | NM_006205 | 82.4 | 113.1 | 0.0313 | 1.2 |
| MED31 | mediator complex subunit 31(MED31) | NM_016060 | 441.8 | 405.1 | 0.0314 | 0.83 |
| | | XM_935907 | 416.3 | 658.5 | 0.0314 | 1.48 |

| | | | | | | |
|---|---|---|---|---|---|---|
| CCDC85C | coiled-coil domain containing 85C(CCDC85C) | AK123840 | 368.8 | 309.1 | 0.0316 | 0.73 |
| PRMT6 | protein arginine methyltransferase 6(PRMT6) | NM_018137 | 177.1 | 161.2 | 0.0316 | 0.81 |
| PRMT6 | protein arginine methyltransferase 6(PRMT6) | NM_018137 | 212.6 | 203.4 | 0.0316 | 0.81 |
| ARSD | arylsulfatase D(ARSD) | NM_001669 | 970.1 | 728.8 | 0.0318 | 0.68 |
| | | XR_017498 | 151.5 | 232.3 | 0.0318 | 1.35 |
| | | XR_017498 | 458.5 | 519.2 | 0.0318 | 1.35 |
| EFHD1 | EF-hand domain family member D1(EFHD1) | NM_025202 | 600 | 427.1 | 0.0319 | 0.66 |
| N4BP1 | NEDD4 binding protein 1(N4BP1) | NM_153029 | 142.1 | 198.4 | 0.0319 | 1.2 |
| UNC5C | unc-5 netrin receptor C(UNC5C) | NM_003728 | 110.8 | 111.9 | 0.0319 | 0.91 |
| AUH | AU RNA binding methylglutaconyl-CoA hydratase(AUH) | NM_001698 | 505.3 | 369.1 | 0.032 | 0.67 |
| CFDP1 | craniofacial development protein 1(CFDP1) | NM_006324 | 889.4 | 731.3 | 0.032 | 0.77 |

| | | | | | | |
|---|---|---|---|---|---|---|
| TIMM23 | translocase of inner mitochondrial membrane 23(TIMM23) | NM_006327 | 154.6 | 176 | 0.032 | 0.72 |
| TIMM23 | translocase of inner mitochondrial membrane 23(TIMM23) | NM_006327 | 419.4 | 365.1 | 0.032 | 0.72 |
| TIMM23 | translocase of inner mitochondrial membrane 23(TIMM23) | NM_006327 | 932.1 | 729.9 | 0.032 | 0.72 |
| TIMM23 | translocase of inner mitochondrial membrane 23(TIMM23) | NM_006327 | 1595 | 1344.1 | 0.032 | 0.72 |
| FAM26D | family with sequence similarity 26 member D(FAM26D) | NM_153036 | 101.6 | 118.4 | 0.0325 | 1.09 |
| FAM26D | family with sequence similarity 26 member D(FAM26D) | NM_153036 | 112.3 | 139.5 | 0.0325 | 1.09 |

| | | | | | | |
|---|---|---|---|---|---|---|
| OR4D6 | olfactory receptor family 4 subfamily D member 6(OR4D6) | NM_001004708 | 126.5 | 161.3 | 0.0325 | 1.12 |
| coa1 | cytochrome c oxidase assembly factor 1 homolog(COA1) | NM_018224 | 193.7 | 177.5 | 0.0326 | 0.8 |
| FBXL6 | F-box and leucine rich repeat protein 6(FBXL6) | NM_012162 | 147.7 | 191.4 | 0.0326 | 1.13 |
| | | XM_001723545 | 108.1 | 135.1 | 0.0326 | 1.1 |
| NDRG2 | NDRG family member 2(NDRG2) | NM_201539 | 700 | 431.8 | 0.0327 | 0.57 |
| RBM10 | RNA binding motif protein 10(RBM10) | NM_152856 | 345.5 | 461.1 | 0.0329 | 1.24 |
| ttll5 | tubulin tyrosine ligase like 5(TTLL5) | NM_015072 | 179.7 | 252.8 | 0.0329 | 1.23 |
| PPIP5K1 | diphosphoinositol pentakisphosphate kinase 1(PPIP5K1) | NM_001024463 | 200.4 | 194 | 0.033 | 0.85 |
| nr0b1 | nuclear receptor subfamily 0 group B member 1(NR0B1) | NM_000475 | 108.5 | 96.6 | 0.0331 | 0.83 |

| | | | | | | |
|---|---|---|---|---|---|---|
| | | XR_016424 | 2184.7 | 3530.7 | 0.0331 | 1.33 |
| TMEM163 | transmembrane protein 163(TMEM163) | NM_030923 | 138.5 | 191.9 | 0.0333 | 1.21 |
| ERI1 | exoribonuclease 1(ERI1) | NM_153332 | 159.9 | 208.2 | 0.0334 | 1.14 |
| BTF3 | basic transcription factor 3(BTF3) | NM_001037637 | 2422.2 | 1588.4 | 0.0335 | 0.61 |
| TCERG1 | transcription elongation regulator 1(TCERG1) | NM_006706 | 113.3 | 127.8 | 0.0336 | 1.21 |
| TCERG1 | transcription elongation regulator 1(TCERG1) | NM_006706 | 181.5 | 250.2 | 0.0336 | 1.21 |
| TPPP3 | tubulin polymerization promoting protein family member 3(TPPP3) | NM_015964 | 116 | 112.2 | 0.0336 | 0.89 |
| | | XM_942669 | 679.3 | 1229.2 | 0.0338 | 1.73 |
| | | XR_016813 | 114.1 | 155.6 | 0.0338 | 1.2 |
| CDC123 | cell division cycle 123(CDC123) | NM_006023 | 169.5 | 259.4 | 0.0339 | 1.37 |
| | | AI024182 | 128.9 | 126.5 | 0.034 | 0.88 |

| | | | | | | | | | |
|---|---|---|---|---|---|---|---|---|---|
| MSI2 | musashi RNA binding protein 2(MSI2) | NM_138962 | 111.3 | 117.6 | 0.0341 | 0.84 | | | |
| MSI2 | musashi RNA binding protein 2(MSI2) | NM_138962 | 188.6 | 182.9 | 0.0341 | 0.84 | | | |
| | | XR_017366 | 106.1 | 139 | 0.0342 | 1.14 | | | |
| GIN1 | gypsy retrotransposon integrase 1(GIN1) | NM_017676 | 119.6 | 123.6 | 0.0343 | 0.83 | | | |
| GIN1 | gypsy retrotransposon integrase 1(GIN1) | NM_017676 | 146.4 | 132.4 | 0.0343 | 0.83 | | | |
| | | XM_926898 | 134.4 | 172.2 | 0.0345 | 1.11 | NM_017196 | 14.73 | 0.69 |
| | | XR_016036 | 194.9 | 302.4 | 0.0347 | 1.37 | NM_017196 | 14.73 | 0.69 |
| | | XR_015255 | 105.7 | 147.3 | 0.0349 | 1.23 | NM_017196 | 14.73 | 0.69 |
| | | XR_015611 | 139.9 | 213.1 | 0.0349 | 1.34 | | | |
| APOO | apolipoprotein O(APOO) | NM_024122 | 648.6 | 459.8 | 0.035 | 0.65 | | | |
| Fam129c | family with sequence similarity 129 member C(FAM129C) | NM_173544 | 111.1 | 141.3 | 0.0351 | 1.12 | | | |

| | | | | | | | | | |
|---|---|---|---|---|---|---|---|---|---|
| FNDC5 | fibronectin type III domain containing 5(FNDC5) | NM_153756 | 233.1 | 143.2 | 0.0351 | 0.55 | NM_031356 | 5.43 | 24.53 |
| PFKM | phosphofructokinase, muscle(PFKM) | NM_000289 | 1167.2 | 418.7 | 0.0351 | 0.32 | NM_031356 | 5.43 | 24.53 |
| | | CR605357 | 135.1 | 133 | 0.0351 | 0.88 | | | |
| CA10 | carbonic anhydrase 10(CA10) | NM_020178 | 125.5 | 118.9 | 0.0354 | 0.86 | | | |
| DPPA2 | developmental pluripotency associated 2(DPPA2) | NM_138815 | 114.9 | 155.5 | 0.0354 | 1.17 | | | |
| SYNGR1 | synaptogyrin 1(SYNGR1) | NM_004711 | 135.2 | 123 | 0.0354 | 0.83 | | | |
| DRC3 | dynein regulatory complex subunit 3(DRC3) | NM_031294 | 127.7 | 113.6 | 0.0355 | 0.81 | | | |
| NDUFA1 | NADH:ubiquinone oxidoreductase subunit A1(NDUFA1) | NM_004541 | 1004.1 | 723.3 | 0.0355 | 0.65 | | | |
| TSEN34 | tRNA splicing endonuclease subunit 34(TSEN34) | NM_001077446 | 123.2 | 139.1 | 0.0355 | 0.78 | | | |

| | | | | | | | | | |
|---|---|---|---|---|---|---|---|---|---|
| TSEN34 | tRNA splicing endonuclease subunit 34(TSEN34) | NM_001077446 | 1078.5 | 938.7 | 0.0355 | 0.78 | | | |
| | | XM_927955 | 4868.5 | 3764.5 | 0.0356 | 0.67 | | | |
| | | AW364673 | 134.6 | 122.6 | 0.0357 | 0.83 | | | |
| | | XR_017072 | 119.2 | 177.6 | 0.0357 | 1.28 | | | |
| CLIP4 | CAP-Gly domain containing linker protein family member 4(CLIP4) | NM_024692 | 249.4 | 222.4 | 0.0359 | 0.79 | | | |
| SAYSD1 | SAYSVFN motif domain containing 1(SAYSD1) | NM_018322 | 94.6 | 119.8 | 0.0359 | 1.13 | | | |
| ANKRD9 | ankyrin repeat domain 9(ANKRD9) | NM_152326 | 107.3 | 115.4 | 0.036 | 0.67 | | | |
| ANKRD9 | ankyrin repeat domain 9(ANKRD9) | NM_152326 | 317 | 230 | 0.036 | 0.67 | NM_017135 | 41.45 | 2.88 |
| STYX | serine/threonine/tyrosine interacting protein(STYX) | NM_145251 | 109.1 | 110.1 | 0.0364 | 0.91 | NM_017135 | 41.45 | 2.88 |
| | | XM_209824 | 111 | 135.6 | 0.0365 | 1.08 | NM_017135 | 41.45 | 2.88 |

| | | | | | | | | | |
|---|---|---|---|---|---|---|---|---|---|
| PTPN20 | protein tyrosine phosphatase, non-receptor type 20(PTPN20) | NM_015605 | 101.9 | 128.9 | 0.0366 | 1.11 | NM_017135 | 41.45 | 2.88 |
| NTRK3 | neurotrophic receptor tyrosine kinase 3(NTRK3) | NM_002530 | 102.8 | 123.3 | 0.0367 | 1.08 | | | |
| TNFAIP1 | TNF alpha induced protein 1(TNFAIP1) | NM_021137 | 171.2 | 263.4 | 0.0367 | 1.35 | | | |
| | | XM_934518 | 126.6 | 171.7 | 0.0367 | 1.17 | | | |
| BYSL | bystin like(BYSL) | NM_004053 | 114.8 | 154.9 | 0.0369 | 1.17 | | | |
| CCDC177 | coiled-coil domain containing 177(CCDC177) | NM_020181 | 100.9 | 131.6 | 0.0369 | 1.14 | | | |
| USP38 | ubiquitin specific peptidase 38(USP38) | NM_032557 | 126 | 115 | 0.0369 | 0.83 | | | |
| | | NR_031743 | 108.4 | 138.9 | 0.0369 | 1.13 | NM_001033653 | 43.46 | 1.56 |
| TM2D1 | TM2 domain containing 1(TM2D1) | NM_032027 | 155.4 | 193.9 | 0.0374 | 0.81 | | | |
| TM2D1 | TM2 domain containing 1(TM2D1) | NM_032027 | 202 | 178.1 | 0.0374 | 0.81 | | | |

| | | | | | | |
|---|---|---|---|---|---|---|
| Cox7a1 | cytochrome c oxidase subunit 7A1(COX7A1) | NM_001864 | 710.5 | 487.2 | 0.0375 | 0.62 |
| | | XM_372926 | 2409.2 | 4504.4 | 0.0375 | 1.49 |
| MGEA5 | meningioma expressed antigen 5 (hyaluronidase)(MGEA5) | NM_012215 | 109.8 | 143.7 | 0.0376 | 1.14 |
| PIAS2 | protein inhibitor of activated STAT 2(PIAS2) | NM_004671 | 115.3 | 126.3 | 0.0376 | 0.87 |
| PIAS2 | protein inhibitor of activated STAT 2(PIAS2) | NM_004671 | 160.8 | 157.4 | 0.0376 | 0.87 |
| TUBBP6 | tubulin beta class I pseudogene 6(TUBBP6) | NR_003598 | 110.7 | 143.4 | 0.0376 | 1.13 |
| | | XM_928996 | 116.4 | 146.6 | 0.0376 | 1.1 |
| CLASP1 | cytoplasmic linker associated protein 1(CLASP1) | NM_015282 | 422.7 | 357.9 | 0.0378 | 0.75 |
| RPL36 | ribosomal protein L36(RPL36) | NM_033643 | 1083.6 | 1757.7 | 0.038 | 1.41 |
| TNFRSF19 | TNF receptor superfamily member 19(TNFRSF19) | NM_018647 | 117.1 | 111.3 | 0.038 | 0.85 |

| | | | | | | | | | |
|---|---|---|---|---|---|---|---|---|---|
| | | XM_374276 | 109.5 | 105 | 0.038 | 0.88 | | | |
| | | XR_018261 | 655.9 | 1141.9 | 0.038 | 1.54 | | | |
| PPTC7 | PTC7 protein phosphatase homolog(PPTC7) | NM_139283 | 282 | 240 | 0.0381 | 0.75 | | | |
| ACAP2 | ArfGAP with coiled-coil, ankyrin repeat and PH domains 2(ACAP2) | NM_012287 | 118.4 | 164.3 | 0.0382 | 1.34 | | | |
| ACAP2 | ArfGAP with coiled-coil, ankyrin repeat and PH domains 2(ACAP2) | NM_012287 | 126.2 | 170.7 | 0.0382 | 1.34 | | | |
| ACAP2 | ArfGAP with coiled-coil, ankyrin repeat and PH domains 2(ACAP2) | NM_012287 | 157.8 | 242.1 | 0.0382 | 1.34 | | | |
| | | XM_377426 | 115.4 | 206.1 | 0.0382 | 1.56 | | | |
| STMN1 | stathmin 1(STMN1) | NM_203401 | 102.2 | 147.4 | 0.0384 | 1.26 | | | |
| | | XM_929427 | 141.3 | 190.2 | 0.0384 | 1.16 | | | |
| KEAP1 | kelch like ECH associated protein 1(KEAP1) | NM_012289 | 249.4 | 238.4 | 0.0387 | 0.8 | NM_134407 | 50.47 | 11.16 |

| | | | | | | |
|---|---|---|---|---|---|---|
| KEAP1 | kelch like ECH associated protein 1(KEAP1) | NM_012289 | 329.6 | 296.7 | 0.0387 | 0.8 |
| | | XR_038871 | 541.3 | 964.7 | 0.0389 | 1.62 |
| YIF1B | Yip1 interacting factor homolog B, membrane trafficking protein(YIF1B) | NM_033557 | 124 | 125.9 | 0.039 | 0.91 |
| ZFP82 | ZFP82 zinc finger protein(ZFP82) | NM_133466 | 125.4 | 162 | 0.039 | 1.12 |
| | | AW780096 | 100.2 | 119 | 0.039 | 1.08 |
| | | XR_016632 | 716 | 1233 | 0.039 | 1.62 |
| | | XR_038356 | 129 | 195.1 | 0.0391 | 1.28 |
| NDUFB8 | NADH:ubiquinone oxidoreductase subunit B8(NDUFB8) | NM_005004 | 5604.2 | 4470.4 | 0.0396 | 0.68 |
| NCBP1 | nuclear cap binding protein subunit 1(NCBP1) | NM_002486 | 292.3 | 382.8 | 0.0397 | 1.21 |
| PDS5A | PDS5 cohesin associated factor A(PDS5A) | NM_015200 | 104.4 | 132.4 | 0.0398 | 1.14 |
| | | XR_039693 | 147.7 | 224.7 | 0.0398 | 1.34 |

| | | | | | | | | | |
|---|---|---|---|---|---|---|---|---|---|
| CDC6 | cell division cycle 6(CDC6) | NM_001254 | 106.1 | 136.6 | 0.0399 | 1.13 | | | |
| GATB | glutamyl-tRNA amidotransferase subunit B(GATB) | NM_004564 | 200.2 | 184.2 | 0.0401 | 0.82 | | | |
| MBTPS1 | membrane bound transcription factor peptidase, site 1(MBTPS1) | NM_003791 | 195.7 | 210.8 | 0.0402 | 1.19 | | | |
| MBTPS1 | membrane bound transcription factor peptidase, site 1(MBTPS1) | NM_003791 | 197.7 | 264.4 | 0.0402 | 1.19 | | | |
| UBR3 | ubiquitin protein ligase E3 component n-recognin 3 (putative)(UBR3) | NM_172070 | 199.2 | 199.1 | 0.0403 | 0.85 | | | |
| UBR3 | ubiquitin protein ligase E3 component n-recognin 3 (putative)(UBR3) | NM_172070 | 220.9 | 211.2 | 0.0403 | 0.85 | NM_001033706 | 2.42 | 10.86 |
| BRD1 | bromodomain containing 1(BRD1) | NM_014577 | 143.5 | 190.5 | 0.0404 | 1.16 | | | |

| | | | | | | | | | |
|---|---|---|---|---|---|---|---|---|---|
| TRPM4 | transient receptor potential cation channel subfamily M member 4(TRPM4) | NM_017636 | 366.4 | 255.3 | 0.0404 | 0.61 | | | |
| CDK12 | cyclin dependent kinase 12(CDK12) | NM_016507 | 217.3 | 498.7 | 0.0405 | 2.02 | | | |
| ZDHHC17 | zinc finger DHHC-type containing 17(ZDHHC17) | NM_015336 | 95.9 | 121.1 | 0.0405 | 1.13 | | | |
| | | NM_001080831 | 145.7 | 206.3 | 0.0405 | 1.23 | | | |
| TRAF3IP2 | TRAF3 interacting protein 2(TRAF3IP2) | NM_147686 | 392.8 | 312.2 | 0.0406 | 0.71 | NM_153300 | 0.14 | 61.17 |
| SIGLEC17P | sialic acid binding Ig like lectin 17, pseudogene(SIGLEC17P) | NR_002804 | 128.8 | 166.5 | 0.0407 | 1.12 | | | |
| TMEM208 | transmembrane protein 208(TMEM208) | NM_014187 | 250.3 | 237.5 | 0.0407 | 0.85 | NM_032416 | 108.51 | 1.87 |
| TMEM208 | transmembrane protein 208(TMEM208) | NM_014187 | 299.6 | 302 | 0.0407 | 0.85 | | | |
| Znf763 | zinc finger protein 763(ZNF763) | NM_001012753 | 106.5 | 151.7 | 0.0407 | 1.23 | | | |

| | | | | | | |
|---|---|---|---|---|---|---|
| EI24 | EI24, autophagy associated transmembrane protein(EI24) | NM_004879 | 206.8 | 173.7 | 0.0408 | 0.78 |
| CCNDBP1 | cyclin D1 binding protein 1(CCNDBP1) | NM_012142 | 163.9 | 152.5 | 0.0409 | 0.83 |
| josd2 | Josephin domain containing 2(JOSD2) | NM_138334 | 120.2 | 117.2 | 0.0409 | 0.88 |
| NDRG4 | NDRG family member 4(NDRG4) | NM_020465 | 117.7 | 128.3 | 0.041 | 0.9 |
| NDRG4 | NDRG family member 4(NDRG4) | NM_020465 | 119.4 | 120.7 | 0.041 | 0.9 |
| SAFB2 | scaffold attachment factor B2(SAFB2) | NM_014649 | 124.3 | 136.7 | 0.041 | 1.25 |
| SAFB2 | scaffold attachment factor B2(SAFB2) | NM_014649 | 653.9 | 894.6 | 0.041 | 1.25 |
| FGF18 | fibroblast growth factor 18(FGF18) | NM_003862 | 119.7 | 120.1 | 0.0412 | 0.91 |
| APIP | APAF1 interacting protein(APIP) | NM_015957 | 198.8 | 200.5 | 0.0413 | 0.76 |
| APIP | APAF1 interacting protein(APIP) | NM_015957 | 350.3 | 301.3 | 0.0413 | 0.76 |

| | | | | | | | | | |
|---|---|---|---|---|---|---|---|---|---|
| CCT6P1 | chaperonin containing TCP1 subunit 6 pseudogene 1(CCT6P1) | NR_003110 | 508.4 | 644.7 | 0.0413 | 1.16 | NM_001191088 | 10.07 | 0.03 |
| | | XM_938599 | 503.5 | 827.3 | 0.0413 | 1.52 | NM_001191088 | 10.07 | 0.03 |
| ESD | esterase D(ESD) | NM_001984 | 247.8 | 204.4 | 0.0414 | 0.72 | | | |
| ESD | esterase D(ESD) | NM_001984 | 1970.5 | 1884.5 | 0.0414 | 0.72 | | | |
| MRPS33 | mitochondrial ribosomal protein S33(MRPS33) | NM_016071 | 126.7 | 116.9 | 0.0414 | 0.84 | | | |
| MRPS33 | mitochondrial ribosomal protein S33(MRPS33) | NM_016071 | 318.9 | 316.3 | 0.0414 | 0.84 | NM_012496 | 1095.39 | 8.76 |
| MXD1 | MAX dimerization protein 1(MXD1) | NM_002357 | 118.3 | 150.4 | 0.0414 | 1.1 | | | |
| SKP2 | S-phase kinase associated protein 2(SKP2) | NM_032637 | 103.7 | 135.7 | 0.0414 | 1.16 | | | |
| CTLA4 | cytotoxic T-lymphocyte associated protein 4(CTLA4) | NM_005214 | 108 | 156.4 | 0.0415 | 1.27 | | | |
| LMOD3 | leiomodin 3(LMOD3) | NM_198271 | 658.6 | 1079.3 | 0.0415 | 1.53 | | | |

| | | | | | | |
|---|---|---|---|---|---|---|
| P4HA1 | prolyl 4-hydroxylase subunit alpha 1(P4HA1) | NM_000917 | 125.8 | 126.1 | 0.0415 | 0.9 |
| PON3 | paraoxonase 3(PON3) | NM_000940 | 111.4 | 105.6 | 0.0415 | 0.87 |
| RANBP3 | RAN binding protein 3(RANBP3) | NM_003624 | 103.6 | 130.3 | 0.0415 | 1.11 |
| RANBP3 | RAN binding protein 3(RANBP3) | NM_003624 | 105.5 | 132.3 | 0.0415 | 1.11 |
| MTFR1L | mitochondrial fission regulator 1 like(MTFR1L) | NM_019557 | 193.3 | 171.3 | 0.0416 | 0.8 |
| | | CN479898 | 136.7 | 129.1 | 0.0416 | 0.86 |
| | | XM_001719755 | 116.7 | 150.2 | 0.0416 | 1.13 |
| DCAF6 | DDB1 and CUL4 associated factor 6(DCAF6) | NM_0010177977 | 156.4 | 135.3 | 0.0418 | 0.79 |
| DCAF6 | DDB1 and CUL4 associated factor 6(DCAF6) | NM_0010177977 | 1084.1 | 931.7 | 0.0418 | 0.79 |
| TPR | translocated promoter region, nuclear basket protein(TPR) | NM_003292 | 357.6 | 500.3 | 0.042 | 1.25 |
| | | XM_940014 | 117.3 | 114.8 | 0.0422 | 0.89 |

| | | | | | | |
|---|---|---|---|---|---|---|
| MCM7 | minichromosome maintenance complex component 7(MCM7) | NM_005916 | 107.5 | 138.9 | 0.0423 | 1.14 |
| MYL6B | myosin light chain 6B(MYL6B) | NM_002475 | 1326.3 | 797.2 | 0.0423 | 0.57 |
| PPARA | peroxisome proliferator activated receptor alpha(PPARA) | NM_001001928 | 143 | 136.8 | 0.0423 | 0.85 |
| MICU1 | mitochondrial calcium uptake 1(MICU1) | NM_006077 | 273.1 | 227 | 0.0424 | 0.72 |
| ORC1 | origin recognition complex subunit 1(ORC1) | NM_004153 | 126.5 | 163.1 | 0.0424 | 1.13 |
| GLRX5 | glutaredoxin 5(GLRX5) | NM_016417 | 3240.7 | 2238.3 | 0.0425 | 0.62 |
| PIH1D1 | PIH1 domain containing 1(PIH1D1) | NM_017916 | 236.8 | 216.1 | 0.0425 | 0.8 |
| SDHA | succinate dehydrogenase complex flavoprotein subunit A(SDHA) | NM_004168 | 722 | 712.9 | 0.0425 | 0.51 |

| | | | | | | |
|---|---|---|---|---|---|---|
| SDHA | succinate dehydrogenase complex flavoprotein subunit A(SDHA) | NM_004168 | 744.3 | 418 | 0.0425 | 0.51 |
| | | XR_037304 | 109.2 | 101.7 | 0.0425 | 0.86 |
| KDELR1 | KDEL endoplasmic reticulum protein retention receptor 1(KDELR1) | NM_006801 | 130.8 | 170.7 | 0.0427 | 1.13 |
| PJA1 | praja ring finger ubiquitin ligase 1(PJA1) | NM_001032396 | 205.6 | 177.3 | 0.0428 | 0.78 |
| CHMP4B | charged multivesicular body protein 4B(CHMP4B) | NM_176812 | 465.5 | 415.2 | 0.0429 | 0.8 |
| FGA | fibrinogen alpha chain(FGA) | NM_000508 | 113.1 | 106.4 | 0.0429 | 0.87 |
| MAPKBP1 | mitogen-activated protein kinase binding protein 1(MAPKBP1) | NM_014994 | 107.3 | 134 | 0.043 | 1.11 |
| polr2m | RNA polymerase II subunit M(POLR2M) | NM_0010188102 | 106.2 | 116.4 | 0.043 | 0.82 |
| polr2m | RNA polymerase II subunit M(POLR2M) | NM_0010188102 | 127.1 | 112.7 | 0.043 | 0.82 |

| | | | | | | | | | |
|---|---|---|---|---|---|---|---|---|---|
| TIMM9 | translocase of inner mitochondrial membrane 9(TIMM9) | NM_012460 | 271.6 | 254.4 | 0.043 | 0.84 | | | |
| UNQ6494 | uncharacterized LOC100129066(UNQ6494) | NR_024280 | 107 | 133.4 | 0.043 | 1.11 | | | |
| | | XM_001719035 | 364.6 | 522.4 | 0.043 | 1.36 | | | |
| | | XR_039704 | 101.7 | 132.1 | 0.043 | 1.15 | | | |
| CFAP69 | cilia and flagella associated protein 69(CFAP69) | NM_001039706 | 171.7 | 153.9 | 0.0431 | 0.82 | NM_013059 | 21.02 | 0.11 |
| PALM | paralemmin(PALM) | NM_002579 | 107.5 | 113.4 | 0.0431 | 0.61 | | | |
| PALM | paralemmin(PALM) | NM_002579 | 244 | 218.2 | 0.0431 | 0.61 | | | |
| PALM | paralemmin(PALM) | NM_002579 | 1139.7 | 756.9 | 0.0431 | 0.61 | | | |
| TMSB4X | thymosin beta 4, X-linked(TMSB4X) | NM_021109 | 139.5 | 266.2 | 0.0432 | 1.73 | | | |
| TMSB4X | thymosin beta 4, X-linked(TMSB4X) | NM_021109 | 16760.5 | 17031.2 | 0.0432 | 1.73 | | | |
| | | AA193588 | 125.4 | 163.4 | 0.0433 | 1.13 | | | |
| | | XM_933938 | 99.3 | 127.8 | 0.0434 | 1.14 | | | |

| | | | | | | | | | |
|---|---|---|---|---|---|---|---|---|---|
| KIAA0586 | KIAA0586(KIAA0586) | NM_014749 | 118.7 | 117.6 | 0.0435 | 0.89 | | | |
| PARP16 | poly(ADP-ribose) polymerase family member 16(PARP16) | NM_017851 | 104.2 | 135.5 | 0.0435 | 1.16 | | | |
| PEX7 | peroxisomal biogenesis factor 7(PEX7) | NM_000288 | 278.2 | 256.5 | 0.0436 | 0.83 | | | |
| FAM122A | family with sequence similarity 122A(FAM122A) | NM_138333 | 105.7 | 129.6 | 0.0437 | 1.09 | | | |
| pnkd | paroxysmal nonkinesigenic dyskinesia(PNKD) | NM_015488 | 245.6 | 174.8 | 0.0438 | 0.65 | NM_012816 | 19.78 | 2.28 |
| RPL32P18 | ribosomal protein L32 pseudogene 18(RPL32P18) | NM_001090027 | 1617.9 | 2874.7 | 0.0438 | 1.48 | NM_012816 | 19.78 | 2.28 |
| ZC3H18-AS1 | ZC3H18 antisense RNA 1 (head to head)(ZC3H18-AS1) | NM_0010011682 | 103.9 | 127.5 | 0.0438 | 1.1 | | | |
| | | XM_927868 | 845.1 | 653.3 | 0.0438 | 0.72 | | | |
| CDK18 | cyclin dependent kinase 18(CDK18) | NM_212503 | 151.1 | 146.7 | 0.0439 | 0.85 | | | |

| | | | | | | |
|---|---|---|---|---|---|---|
| FREM2 | FRAS1 related extracellular matrix protein 2(FREM2) | NM_207361 | 117.9 | 104.5 | 0.044 | 0.81 |
| MRPL11 | mitochondrial ribosomal protein L11(MRPL11) | NM_016050 | 541.9 | 488.5 | 0.0441 | 0.83 |
| | | AI183825 | 117.1 | 117.5 | 0.0441 | 0.9 |
| ANAPC15 | anaphase promoting complex subunit 15(ANAPC15) | NM_014042 | 187.5 | 160.1 | 0.0442 | 0.75 |
| APOA2 | apolipoprotein A2(APOA2) | NM_001643 | 157.9 | 104.6 | 0.0442 | 0.6 |
| KIAA1551 | KIAA1551(KIAA1551) | NM_018169 | 96.6 | 136.8 | 0.0442 | 1.24 |
| KIAA1551 | KIAA1551(KIAA1551) | NM_018169 | 100.5 | 119.9 | 0.0442 | 1.24 |
| PPIL6 | peptidylprolyl isomerase like 6(PPIL6) | NM_173672 | 112.6 | 103.3 | 0.0442 | 0.84 |
| | | XR_018020 | 106.7 | 134.1 | 0.0442 | 1.11 |
| ADAM8 | ADAM metallopeptidase domain 8(ADAM8) | NM_001109 | 92.3 | 121.7 | 0.0443 | 1.16 |
| GOT2 | glutamic-oxaloacetic transaminase 2(GOT2) | NM_002080 | 1233.7 | 865.2 | 0.0443 | 0.65 |

| | | | | | | | | | |
|---|---|---|---|---|---|---|---|---|---|
| RFX5 | regulatory factor X5(RFX5) | NM_001025603 | 106.6 | 141.5 | 0.0443 | 1.17 | | | |
| RPUSD3 | RNA pseudouridylate synthase domain containing 3(RPUSD3) | NM_173659 | 260 | 249.8 | 0.0443 | 0.84 | | | |
| HAX1 | HCLS1 associated protein X-1(HAX1) | NM_006118 | 138.7 | 135.6 | 0.0444 | 0.88 | | | |
| HAX1 | HCLS1 associated protein X-1(HAX1) | NM_006118 | 334.7 | 354.8 | 0.0444 | 0.88 | | | |
| ACTR3 | ARP3 actin related protein 3 homolog(ACTR3) | NM_005721 | 162.5 | 310.8 | 0.0445 | 1.71 | NM_031544 | 4.79 | 38.59 |
| | | XM_001713657 | 730 | 488.3 | 0.0445 | 0.62 | NM_031544 | 4.79 | 38.59 |
| | | XM_371878 | 91.6 | 116.4 | 0.0445 | 1.14 | | | |
| FAM45A | family with sequence similarity 45 member A(FAM45A) | NM_207009 | 117.1 | 110.3 | 0.0446 | 0.84 | | | |

| | | | | | | |
|---|---|---|---|---|---|---|
| HAR1B | highly accelerated region 1B (non-protein coding)(HAR1B) | NR_003245 | 111.9 | 141 | 0.0447 | 1.11 |
| PRKRIP1 | PRKR interacting protein 1 (IL11 inducible)(PRKRIP1) | NM_024653 | 123.9 | 157 | 0.0447 | 1.11 |
| USP33 | ubiquitin specific peptidase 33(USP33) | NM_201624 | 104.6 | 101 | 0.0447 | 0.89 |
| RRP1B | ribosomal RNA processing 1B(RRP1B) | NM_015056 | 109.3 | 104.6 | 0.0448 | 0.87 |
| | | XM_001724539 | 453.1 | 729.2 | 0.0448 | 1.51 |
| | | XR_016197 | 81.4 | 120.8 | 0.0448 | 1.29 |
| CDC45 | cell division cycle 45(CDC45) | NM_003504 | 86.1 | 134.6 | 0.0449 | 1.35 |
| FGG | fibrinogen gamma chain(FGG) | NM_000509 | 188.8 | 101.8 | 0.0449 | 0.48 |
| GPR89B | G protein-coupled receptor 89B(GPR89B) | NM_016334 | 150.5 | 138.8 | 0.045 | 0.85 |
| | | AW014327 | 106.7 | 136.9 | 0.045 | 1.12 |
| | | XM_928029 | 152.5 | 153.4 | 0.045 | 0.89 |

| | | | | | | |
|---|---|---|---|---|---|---|
| CCDC28B | coiled-coil domain containing 28B(CCDC28B) | NM_024296 | 119.7 | 125.2 | 0.0451 | 0.88 |
| CCDC28B | coiled-coil domain containing 28B(CCDC28B) | NM_024296 | 140.9 | 139.9 | 0.0451 | 0.88 |
| CD8A | CD8a molecule(CD8A) | NM_001768 | 115.5 | 169.7 | 0.0451 | 1.29 |
| | | XR_016732 | 2595.4 | 4667.1 | 0.0451 | 1.4 |
| | | XM_0017144765 | 114.2 | 111.8 | 0.0453 | 0.9 |
| MPRIP | myosin phosphatase Rho interacting protein(MPRIP) | NM_201274 | 195 | 166.4 | 0.0454 | 0.78 |
| Uqcrfs1 | ubiquinol-cytochrome c reductase, Rieske iron-sulfur polypeptide 1(UQCRFS1) | NM_006003 | 4410.4 | 3323 | 0.0455 | 0.66 |
| COPA | coatomer protein complex subunit alpha(COPA) | NM_004371 | 233.5 | 325.7 | 0.0456 | 1.25 |
| PPP1CA | protein phosphatase 1 catalytic subunit alpha(PPP1CA) | NM_002708 | 121.5 | 125.2 | 0.0456 | 0.92 |

| | | | | | | |
|---|---|---|---|---|---|---|
| | | AW028158 | 101 | 131.2 | 0.0457 | 1.15 |
| | | XM_001128975 | 160.9 | 212.8 | 0.0457 | 1.15 |
| SLC26A8 | solute carrier family 26 member 8(SLC26A8) | NM_052961 | 126.3 | 167.9 | 0.0458 | 1.16 |
| | | XM_940853 | 111.7 | 110.7 | 0.0458 | 0.91 |
| CEBPZ | CCAAT/enhancer binding protein zeta(CEBPZ) | NM_005760 | 446.1 | 598.6 | 0.0461 | 1.23 |
| | | XM_001719341 | 1141.8 | 2040.2 | 0.0462 | 1.61 |
| | | XR_039643 | 131.7 | 193.8 | 0.0462 | 1.3 |
| | | XM_001126676 | 127.7 | 125 | 0.0463 | 0.88 |
| | | XR_040489 | 121.9 | 125.3 | 0.0463 | 0.91 |
| ANTXR1 | anthrax toxin receptor 1(ANTXR1) | NM_032208 | 167.3 | 277.1 | 0.0464 | 1.51 |
| | | XM_927511 | 107.3 | 128.4 | 0.0464 | 1.07 |
| | | XM_001717905 | 120 | 150.6 | 0.0465 | 1.1 |
| DPH2 | DPH2 homolog(DPH2) | NM_001384 | 169.1 | 165.1 | 0.0466 | 0.87 |
| DUSP13 | dual specificity phosphatase 13(DUSP13) | NM_016364 | 141.4 | 115.6 | 0.0466 | 0.73 |

| | | | | | | |
|---|---|---|---|---|---|---|
| DUSP13 | dual specificity phosphatase 13(DUSP13) | NM_016364 | 190.8 | 210.7 | 0.0466 | 0.73 |
| HSD11B2 | hydroxysteroid 11-beta dehydrogenase 2(HSD11B2) | NM_000196 | 458.6 | 245.1 | 0.0466 | 0.51 |
| TRABD | TraB domain containing(TRABD) | NM_025204 | 624 | 944 | 0.0467 | 1.38 |
| WDR33 | WD repeat domain 33(WDR33) | NM_001006623 | 110.6 | 135.3 | 0.0467 | 1.08 |
| WDR33 | WD repeat domain 33(WDR33) | NM_001006623 | 131.2 | 163.2 | 0.0467 | 1.08 |
| ABCD3 | ATP binding cassette subfamily D member 3(ABCD3) | NM_002858 | 148.9 | 126.4 | 0.047 | 0.77 |
| SLC6A10P | solute carrier family 6 member 10, pseudogene(SLC6A10P) | NR_003083 | 192.7 | 160 | 0.047 | 0.76 |
| | | XM_941195 | 1831 | 2947.7 | 0.047 | 1.42 |

| | | | | | | |
|---|---|---|---|---|---|---|
| FAM210 A | family with sequence similarity 210 member A(FAM210A) | NM_15235 2 | 189.1 | 155.6 | 0.0471 | 0.77 |
| KIF13B | kinesin family member 13B(KIF13B) | NM_01525 4 | 794.4 | 649.9 | 0.0472 | 0.75 |
| MIR125 3 | microRNA 1253(MIR1253) | NR_03165 4 | 143.3 | 191.4 | 0.0472 | 1.17 |
| STXBP5 | syntaxin binding protein 5(STXBP5) | NM_13924 4 | 152.3 | 211.1 | 0.0473 | 1.23 |
| KLHDC 3 | kelch domain containing 3(KLHDC3) | NM_05716 1 | 957.9 | 896.4 | 0.0474 | 0.87 |
| TTR | transthyretin(TTR) | NM_00037 1 | 205.9 | 141 | 0.0474 | 0.59 |
| | | XR_01774 7 | 109.6 | 110.3 | 0.0474 | 0.92 |
| | | XR_01774 7 | 122.2 | 127.7 | 0.0474 | 0.92 |
| | | XR_03781 2 | 171.6 | 232.6 | 0.0474 | 1.2 |
| | | XR_03842 9 | 99.6 | 124.8 | 0.0474 | 1.13 |
| CLDN3 | claudin 3(CLDN3) | NM_00130 6 | 214.1 | 151.1 | 0.0476 | 0.67 |

| | | | | | | |
|---|---|---|---|---|---|---|
| OR2T12 | olfactory receptor family 2 subfamily T member 12(OR2T12) | NM_001004692 | 167.6 | 232 | 0.0476 | 1.2 |
| IFITM4P | interferon induced transmembrane protein 4 pseudogene(IFITM4P) | NR_001590 | 128.1 | 167.1 | 0.0477 | 1.13 |
| GPD1L | glycerol-3-phosphate dehydrogenase 1-like(GPD1L) | NM_015141 | 758.3 | 659.4 | 0.0479 | 0.79 |
| | | XR_037672 | 131.4 | 130.6 | 0.0479 | 0.9 |
| GGT6 | gamma-glutamyltransferase 6(GGT6) | NM_153338 | 146.5 | 125.8 | 0.048 | 0.77 |
| LINC01420 | long intergenic non-protein coding RNA 1420(LINC01420) | NR_015367 | 1185.8 | 1017.2 | 0.048 | 0.8 |
| | | XR_037282 | 174.7 | 263.9 | 0.048 | 1.34 |
| RIPK4 | receptor interacting serine/threonine kinase 4(RIPK4) | NM_020639 | 129.6 | 128.8 | 0.0481 | 0.89 |

| | | | | | | |
|---|---|---|---|---|---|---|
| RPS3A | ribosomal protein S3A(RPS3A) | NM_001006 | 5938.6 | 8852.1 | 0.0482 | 1.18 |
| RPS3A | ribosomal protein S3A(RPS3A) | NM_001006 | 6427 | 9708 | 0.0482 | 1.18 |
| RPS3A | ribosomal protein S3A(RPS3A) | NM_001006 | 6930.3 | 9567.1 | 0.0482 | 1.18 |
| MDM2 | MDM2 proto-oncogene(MDM2) | NM_002392 | 146.5 | 194.2 | 0.0483 | 1.14 |
| HSBP1 | heat shock factor binding protein 1(HSBP1) | NM_001537 | 182.4 | 411.4 | 0.0485 | 1.9 |
| | | XM_930111 | 407.7 | 571.9 | 0.0485 | 1.31 |
| | | XM_940214 | 409 | 599.6 | 0.0485 | 1.37 |
| AKT3 | AKT serine/threonine kinase 3(AKT3) | NM_181690 | 117 | 155.4 | 0.0488 | 1.15 |
| RPS21 | ribosomal protein S21(RPS21) | NM_001024 | 129.8 | 165.5 | 0.0488 | 1.11 |
| | | XR_041286 | 1127.7 | 1954 | 0.0488 | 1.51 |
| | | XR_041286 | 3769.8 | 5828 | 0.0488 | 1.51 |
| SMIM13 | small integral membrane protein 13(SMIM13) | AJ420516 | 123 | 114.8 | 0.0489 | 0.86 |

| | | | | | | |
|---|---|---|---|---|---|---|
| ECEL1 | endothelin converting enzyme like 1(ECEL1) | NM_004826 | 117.2 | 149.2 | 0.0491 | 1.12 |
| | | XM_0011255941 | 270.5 | 659.8 | 0.0491 | 2.22 |
| | | XM_0011255941 | 414.5 | 851 | 0.0491 | 2.22 |
| | | XM_929608 | 103.9 | 100 | 0.0491 | 0.88 |
| FAM13B | family with sequence similarity 13 member B(FAM13B) | NM_016603 | 98.9 | 123.7 | 0.0492 | 1.13 |
| HMBS | hydroxymethylbilane synthase(HMBS) | NM_000190 | 99.5 | 110.7 | 0.0492 | 0.81 |
| HMBS | hydroxymethylbilane synthase(HMBS) | NM_000190 | 149.5 | 135.3 | 0.0492 | 0.81 |
| | | BQ007862 | 105.5 | 129.8 | 0.0492 | 1.09 |
| CECR5 | cat eye syndrome chromosome region, candidate 5(CECR5) | NM_017829 | 137.7 | 135.1 | 0.0494 | 0.88 |
| LYPLAL1 | lysophospholipase like 1(LYPLAL1) | NM_138794 | 125.1 | 132.7 | 0.0494 | 0.72 |

| | | | | | | |
|---|---|---|---|---|---|---|
| LYPLAL1 | lysophospholipase like 1(LYPLAL1) | NM_138794 | 401.4 | 315 | 0.0494 | 0.72 |
| | | XM_932583 | 105.1 | 134.6 | 0.0494 | 1.14 |
| EML1 | echinoderm microtubule associated protein like 1(EML1) | NM_004434 | 145.4 | 142.5 | 0.0495 | 0.79 |
| EML1 | echinoderm microtubule associated protein like 1(EML1) | NM_004434 | 210.2 | 187 | 0.0495 | 0.79 |
| ENTPD4 | ectonucleoside triphosphate diphosphohydrolase 4(ENTPD4) | NM_004901 | 150.7 | 196.6 | 0.0495 | 1.13 |
| GADD45A | growth arrest and DNA damage inducible alpha(GADD45A) | NM_001924 | 1030.8 | 588.5 | 0.0495 | 0.51 |
| GADD45A | growth arrest and DNA damage inducible alpha(GADD45A) | NM_001924 | 1193 | 939.9 | 0.0495 | 0.51 |
| ALB | albumin(ALB) | NM_000477 | 1072.9 | 368.6 | 0.0497 | 0.31 |
| ALB | albumin(ALB) | NM_000477 | 1474.5 | 558.4 | 0.0497 | 0.31 |

| | | | | | | | | | |
|---|---|---|---|---|---|---|---|---|---|
| ATP5D | ATP synthase, H+ transporting, mitochondrial F1 complex, delta subunit(ATP5D) | NM_001687 | 244.3 | 225.6 | 0.0497 | 0.76 | | | |
| ATP5D | ATP synthase, H+ transporting, mitochondrial F1 complex, delta subunit(ATP5D) | NM_001687 | 910.5 | 742.3 | 0.0497 | 0.76 | NM_031012 | 19.39 | 0.85 |
| MIR1185-1 | microRNA 1185-1(MIR1185-1) | NR_031575 | 138.9 | 141.9 | 0.0497 | 0.91 | | | |
| FGA | fibrinogen alpha chain(FGA) | NM_021871 | 187.3 | 127.9 | 0.0498 | 0.6 | | | |
| ZC3H3 | zinc finger CCCH-type containing 3(ZC3H3) | NM_015117 | 184.4 | 242.7 | 0.0499 | 1.18 | NM_012904 | 10.16 | 55.22 |
| PHF1 | PHD finger protein 1(PHF1) | NM_024165 | 110.7 | 129.5 | 0.05 | 1.13 | | | |
| PHF1 | PHD finger protein 1(PHF1) | NM_024165 | 308.9 | 381.8 | 0.05 | 1.13 | | | |

| Gene Symbol | Gene Name | Accession No. | CON Chow M Mean | IUGR Chow M Mean | pval IUGR v CON M Chow | Ratio IUGR v CON M Chow | REFSEQ ID | NEPHRON (MEAN RPKM) | COLLECTING DUCT (MEAN RPKM) |
|---|---|---|---|---|---|---|---|---|---|
| GTPBP6 | GTP binding protein 6 (putative)(GTPBP6) | NM_012227 | 159.9 | 197 | 0.0000 | 1.41 | | | |

| | | | | | | | | | |
|---|---|---|---|---|---|---|---|---|---|
| PDGFA | platelet derived growth factor subunit A(PDGFA) | NM_033023 | 123.1 | 163.8 | 0.0000 | 1.55 | | | |
| | | AW780096 | 106.2 | 112.4 | 0.0000 | 1.16 | | | |
| | | XM_927511 | 115.2 | 92.1 | 0.0000 | 0.85 | | | |
| KDF1 | keratinocyte differentiation factor 1(KDF1) | NM_152365 | 139 | 102.2 | 0.0000 | 0.8 | | | |
| csnk1e | casein kinase 1 epsilon(CSNK1E) | NM_152221 | 127.6 | 189.3 | 0.0000 | 1.68 | | | |
| csnk1e | casein kinase 1 epsilon(CSNK1E) | NM_152221 | 142.1 | 149.5 | 0.0000 | 1.68 | NM_017193 | 356.13 | 8.18 |
| RNA5S9 | RNA, 5S ribosomal 9(RNA5S9) | NR_023371 | 156.7 | 218.1 | 0.0000 | 1.57 | NM_017193 | 356.13 | 8.18 |
| | | XR_039453 | 110.5 | 109.9 | 0.0000 | 1.19 | NM_017193 | 356.13 | 8.18 |
| | | XR_039453 | 196.8 | 203.3 | 0.0000 | 1.19 | | | |
| | | XM_931775 | 134.4 | 100.5 | 0.0000 | 0.81 | | | |
| UPF3A | UPF3 regulator of nonsense transcripts homolog A (yeast)(UPF3A) | NM_023011 | 231 | 347.5 | 0.0000 | 1.68 | | | |

| | | | | | | |
|---|---|---|---|---|---|---|
| MTHFD2L | methylenetetrahydrofolate dehydrogenase (NADP+ dependent) 2-like(MTHFD2L) | NM_001004346 | 95.7 | 108 | 0.0000 | 1.23 |
| ZNF33B | zinc finger protein 33B(ZNF33B) | NM_006955 | 127.8 | 148.8 | 0.0000 | 1.35 |
| | | XM_942469 | 114.8 | 150.9 | 0.0000 | 1.5 |
| | | XM_942469 | 115.9 | 105 | 0.0000 | 1.5 |
| FAM133B | family with sequence similarity 133 member B(FAM133B) | NM_152789 | 273.3 | 368.5 | 0.0000 | 1.5 |
| CAPG | capping actin protein, gelsolin like(CAPG) | NM_001747 | 97.6 | 113.5 | 0.0000 | 1.28 |
| MERTK | MER proto-oncogene, tyrosine kinase(MERTK) | NM_006343 | 113.3 | 127.8 | 0.0000 | 1.25 |
| MERTK | MER proto-oncogene, tyrosine kinase(MERTK) | NM_006343 | 181.3 | 237.3 | 0.0000 | 1.25 |
| TCL1B | T-cell leukemia/lymphoma 1B(TCL1B) | NM_004918 | 114 | 138.6 | 0.0000 | 1.39 |

| | | | | | | |
|---|---|---|---|---|---|---|
| ARID4B | AT-rich interaction domain 4B(ARID4B) | NM_016374 | 126.6 | 127.9 | 0.0000 | 2.14 |
| ARID4B | AT-rich interaction domain 4B(ARID4B) | NM_016374 | 141.3 | 210.3 | 0.0000 | 2.14 |
| ARID4B | AT-rich interaction domain 4B(ARID4B) | NM_016374 | 226.6 | 430.2 | 0.0000 | 2.14 |
| THOC2 | THO complex 2(THOC2) | NM_020449 | 245.8 | 402.9 | 0.0000 | 1.83 |
| TDGP1 | thymine-DNA glycosylase pseudogene 1(TDGP1) | NR_024382 | 94.7 | 112.1 | 0.0000 | 1.29 |
| ZBTB43 | zinc finger and BTB domain containing 43(ZBTB43) | NM_014007 | 116.3 | 139.1 | 0.0000 | 1.34 |
| cep350 | centrosomal protein 350(CEP350) | NM_014810 | 155.4 | 199.8 | 0.0000 | 1.45 |
| KDM5A | lysine demethylase 5A(KDM5A) | NM_005056 | 144.1 | 241 | 0.0000 | 1.89 |

| | | | | | | |
|---|---|---|---|---|---|---|
| NUFIP2 | NUFIP2, FMR1 interacting protein 2(NUFIP2) | NM_020772 | 111.9 | 158.4 | 0.0000 | 1.83 |
| NUFIP2 | NUFIP2, FMR1 interacting protein 2(NUFIP2) | NM_020772 | 211.3 | 341.6 | 0.0000 | 1.83 |
| | | XM_933083 | 143.5 | 101.3 | 0.0000 | 0.77 |
| EXOC1 | exocyst complex component 1(EXOC1) | NM_001024924 | 111.3 | 135 | 0.0000 | 1.38 |
| TAOK1 | TAO kinase 1(TAOK1) | NM_020791 | 96.6 | 114 | 0.0000 | 1.29 |
| TDG | thymine DNA glycosylase(TDG) | NM_003211 | 133 | 190.1 | 0.0000 | 1.6 |
| TDG | thymine DNA glycosylase(TDG) | NM_003211 | 280.2 | 326.3 | 0.0000 | 1.6 |
| | | BF107605 | 111.8 | 119.9 | 0.0000 | 1.21 |
| ATXN3 | ataxin 3(ATXN3) | NM_004993 | 105.1 | 116.8 | 0.0000 | 1.24 |
| NUMBL | NUMB like, endocytic adaptor protein(NUMBL) | NM_004756 | 94.2 | 106 | 0.0000 | 1.24 |
| RAD21 | RAD21 cohesin complex component(RAD21) | NM_006265 | 294.9 | 498 | 0.0000 | 1.89 |

| | | | | | | |
|---|---|---|---|---|---|---|
| RAD21 | RAD21 cohesin complex component(RAD21) | NM_006265 | 311.8 | 296.7 | 0.0000 | 1.89 |
| RRBP1 | ribosome binding protein 1(RRBP1) | NM_001042576 | 130.7 | 165.2 | 0.0000 | 1.44 |
| RRBP1 | ribosome binding protein 1(RRBP1) | NM_001042576 | 358.3 | 314.1 | 0.0000 | 1.44 |
| VRK3 | vaccinia related kinase 3(VRK3) | NM_001025778 | 106.9 | 96.8 | 0.0000 | 1.24 |
| VRK3 | vaccinia related kinase 3(VRK3) | NM_001025778 | 167.3 | 180.3 | 0.0000 | 1.24 |
| WWOX | WW domain containing oxidoreductase(WWOX) | NM_130844 | 115.6 | 157.5 | 0.0000 | 1.53 |
| PCBP2 | poly(rC) binding protein 2(PCBP2) | NM_001098620 | 692.1 | 1075.5 | 0.0000 | 1.75 |
| | | NM_033531 | 98.3 | 152 | 0.0000 | 1.72 |
| ANKRD17 | ankyrin repeat domain 17(ANKRD17) | NM_032217 | 98.6 | 126 | 0.0000 | 1.43 |
| RRAGB | Ras related GTP binding B(RRAGB) | NM_006064 | 124.9 | 110.8 | 0.0010 | 0.66 |
| RRAGB | Ras related GTP binding B(RRAGB) | NM_006064 | 146.7 | 91.4 | 0.0010 | 0.66 |

| | | | | | | |
|---|---|---|---|---|---|---|
| ADAT1 | adenosine deaminase, tRNA specific 1(ADAT1) | NM_012091 | 107.3 | 112.3 | 0.0010 | 1.15 |
| TUBA3FP | tubulin alpha 3f pseudogene(TUBA3FP) | NR_003608 | 118.3 | 95.8 | 0.0010 | 0.86 |
| ASAH1 | N-acylsphingosine amidohydrolase 1(ASAH1) | NM_177924 | 120.5 | 96.2 | 0.0010 | 0.86 |
| FRYL | FRY like transcription coactivator(FRYL) | NM_015030 | 101.3 | 99 | 0.0010 | 1.41 |
| FRYL | FRY like transcription coactivator(FRYL) | NM_015030 | 115.5 | 105.1 | 0.0010 | 1.41 |
| FRYL | FRY like transcription coactivator(FRYL) | NM_015030 | 187.9 | 237.9 | 0.0010 | 1.41 |
| | | XM_001131286 | 127.8 | 99.7 | 0.0010 | 0.83 |
| RSPRY1 | ring finger and SPRY domain containing 1(RSPRY1) | NM_133368 | 218.9 | 294.2 | 0.0010 | 1.5 |
| | | AW274669 | 124.3 | 94.1 | 0.0010 | 0.82 |

| | | | | | | | | | |
|---|---|---|---|---|---|---|---|---|---|
| SUZ12 | SUZ12 polycomb repressive complex 2 subunit(SUZ12) | NM_015355 | 297.5 | 493.9 | 0.0010 | 1.88 | | | |
| | | XR_039881 | 94.7 | 115.4 | 0.0010 | 1.33 | | | |
| PHF7 | PHD finger protein 7(PHF7) | NM_016483 | 116.6 | 96.1 | 0.0010 | 0.88 | NM_053502 | 4.02 | 18.01 |
| NPAT | nuclear protein, coactivator of histone transcription(NPAT) | NM_002519 | 92.4 | 103 | 0.0010 | 1.21 | NM_053502 | 4.02 | 18.01 |
| C1orf64 | chromosome 1 open reading frame 64(C1orf64) | NM_178840 | 117.9 | 89.7 | 0.0010 | 0.81 | NM_053502 | 4.02 | 18.01 |
| FCGR1B | Fc fragment of IgG receptor Ib(FCGR1B) | NM_001017986 | 93.1 | 105.4 | 0.0010 | 1.23 | | | |
| ABHD2 | abhydrolase domain containing 2(ABHD2) | NM_152924 | 116.5 | 120.2 | 0.0010 | 1.23 | | | |
| ABHD2 | abhydrolase domain containing 2(ABHD2) | NM_152924 | 116.9 | 126.8 | 0.0010 | 1.23 | | | |

| | | | | | | | | | |
|---|---|---|---|---|---|---|---|---|---|
| whsc1l1 | Wolf-Hirschhorn syndrome candidate 1-like 1(WHSC1L1) | NM_023034 | 96.6 | 112.4 | 0.0010 | 1.27 | | | |
| SLC38A3 | solute carrier family 38 member 3(SLC38A3) | NM_006841 | 210.4 | 114.2 | 0.0010 | 0.59 | | | |
| CREB1 | cAMP responsive element binding protein 1(CREB1) | AK126342 | 178 | 297.7 | 0.0010 | 1.89 | NM_001009670 | 33.30 | 2.04 |
| | | NM_001001977 | 1360.1 | 813.4 | 0.0010 | 0.69 | | | |
| RPRD2 | regulation of nuclear pre-mRNA domain containing 2(RPRD2) | NM_015203 | 103.1 | 111.3 | 0.0010 | 1.18 | | | |
| RPRD2 | regulation of nuclear pre-mRNA domain containing 2(RPRD2) | NM_015203 | 194.2 | 177.9 | 0.0010 | 1.18 | | | |
| | | XR_040782 | 111.2 | 92.2 | 0.0010 | 0.89 | | | |
| UPF2 | UPF2 regulator of nonsense transcripts homolog (yeast)(UPF2) | NM_015542 | 267.8 | 500.7 | 0.0010 | 2.13 | | | |
| ZNF451 | zinc finger protein 451(ZNF451) | NM_001031623 | 101.2 | 104.1 | 0.0010 | 1.37 | | | |

| | | | | | | |
|---|---|---|---|---|---|---|
| ZNF451 | zinc finger protein 451(ZNF451) | NM_001031623 | 105.3 | 127.4 | 0.0010 | 1.37 |
| CWC22 | CWC22 spliceosome associated protein homolog(CWC22) | NM_020943 | 94.6 | 108.8 | 0.0010 | 1.26 |
| PCBP2 | poly(rC) binding protein 2(PCBP2) | NM_005016 | 1645.9 | 2165.2 | 0.0010 | 1.4 |
| UBE2D2 | ubiquitin conjugating enzyme E2 D2(UBE2D2) | NM_181838 | 93.5 | 103.9 | 0.0010 | 1.21 |
| PSMD9 | proteasome 26S subunit, non-ATPase 9(PSMD9) | NM_002813 | 141.1 | 103.9 | 0.0010 | 0.79 |
| C21orf58 | chromosome 21 open reading frame 58(C21orf58) | NM_058180 | 1075.2 | 1541.9 | 0.0010 | 1.64 |
| HOOK3 | hook microtubule tethering protein 3(HOOK3) | NM_032410 | 96.2 | 111.3 | 0.0010 | 1.26 |
| POGZ | pogo transposable element with ZNF domain(POGZ) | NM_015100 | 112.2 | 139.6 | 0.0010 | 1.4 |

| | | | | | | |
|---|---|---|---|---|---|---|
| LSM8 | LSM8 homolog, U6 small nuclear RNA associated(LSM8) | NM_016200 | 98.6 | 124.7 | 0.0010 | 1.4 |
| C1orf115 | chromosome 1 open reading frame 115(C1orf115) | NM_024709 | 249.6 | 110.1 | 0.0010 | 0.47 |
| RBCK1 | RANBP2-type and C3HC4-type zinc finger containing 1(RBCK1) | NM_006462 | 153 | 152.4 | 0.0010 | 1.13 |
| MPHOSPH10 | M-phase phosphoprotein 10(MPHOSPH10) | NM_005791 | 364.7 | 409.1 | 0.0010 | 1.27 |
| ZNF689 | zinc finger protein 689(ZNF689) | NM_138447 | 146.5 | 170.6 | 0.0010 | 1.34 |
| CARS | cysteinyl-tRNA synthetase(CARS) | NM_001014438 | 146.1 | 224.4 | 0.0010 | 1.75 |
| ATXN3 | ataxin 3(ATXN3) | NM_030660 | 126.2 | 167.2 | 0.0010 | 1.49 |
| GMCL1 | germ cell-less, spermatogenesis associated 1(GMCL1) | NM_178439 | 99 | 95.1 | 0.0010 | 1.44 |

| | | | | | | |
|---|---|---|---|---|---|---|
| GMCL1 | germ cell-less, spermatogenesis associated 1(GMCL1) | NM_178439 | 123 | 160.1 | 0.0010 | 1.44 |
| VPS13C | vacuolar protein sorting 13 homolog C(VPS13C) | NM_018080 | 125 | 149.5 | 0.0010 | 1.36 |
| RBBP6 | RB binding protein 6, ubiquitin ligase(RBBP6) | NM_032626 | 144.7 | 204.2 | 0.0010 | 1.59 |
| | | XM_294209 | 92.7 | 103 | 0.0010 | 1.19 |
| VCP | valosin containing protein(VCP) | NM_007126 | 504.6 | 324.9 | 0.0010 | 0.73 |
| | | XM_001725606 | 104.8 | 113.7 | 0.0010 | 1.2 |
| RGPD5 | RANBP2-like and GRIP domain containing 5(RGPD5) | NM_005054 | 85.4 | 101.5 | 0.0010 | 1.28 |
| ELF2 | E74 like ETS transcription factor 2(ELF2) | NM_006874 | 94.4 | 122.1 | 0.0010 | 1.45 |
| ELF2 | E74 like ETS transcription factor 2(ELF2) | NM_006874 | 146 | 149.3 | 0.0010 | 1.45 |
| bbx | BBX, HMG-box containing(BBX) | NM_020235 | 236.9 | 373.6 | 0.0010 | 1.74 |

| | | | | | | |
|---|---|---|---|---|---|---|
| RUFY3 | RUN and FYVE domain containing 3(RUFY3) | NM_014961 | 118.1 | 152.9 | 0.0010 | 1.43 |
| | | XM_001714086 | 94.7 | 117.5 | 0.0010 | 1.37 |
| | | XR_019071 | 332.4 | 502.8 | 0.0010 | 1.72 |
| FOXO3 | forkhead box O3(FOXO3) | NM_201559 | 197.9 | 362.9 | 0.0010 | 2.01 |
| FOXO3 | forkhead box O3(FOXO3) | NM_201559 | 273 | 307.7 | 0.0010 | 2.01 |
| TIA1 | TIA1 cytotoxic granule associated RNA binding protein(TIA1) | NM_022037 | 165.9 | 239 | 0.0010 | 1.6 |
| UBC | ubiquitin C(UBC) | NM_021009 | 7920.3 | 5031 | 0.0020 | 0.66 |
| UBC | ubiquitin C(UBC) | NM_021009 | 9534 | 7321.4 | 0.0020 | 0.66 |
| XRCC5 | X-ray repair cross complementing 5(XRCC5) | NM_021141 | 116.8 | 129.5 | 0.0020 | 1.27 |
| | | CD520639 | 103.8 | 109.2 | 0.0020 | 1.15 |
| | | DA877611 | 109.7 | 89.1 | 0.0020 | 0.86 |
| CREBRF | CREB3 regulatory factor(CREBRF) | NM_153607 | 195.6 | 260.6 | 0.0020 | 1.45 |

| | | | | | | |
|---|---|---|---|---|---|---|
| CREBRF | CREB3 regulatory factor(CREBRF) | NM_153607 | 206.9 | 332 | 0.0020 | 1.45 |
| BICD2 | BICD cargo adaptor 2(BICD2) | NM_001003800 | 159.2 | 209.9 | 0.0020 | 1.5 |
| MAD1L1 | MAD1 mitotic arrest deficient like 1(MAD1L1) | NM_001013836 | 98.6 | 109 | 0.0020 | 1.2 |
| MAD1L1 | MAD1 mitotic arrest deficient like 1(MAD1L1) | NM_001013836 | 109.5 | 97.4 | 0.0020 | 1.2 |
| OR6C74 | olfactory receptor family 6 subfamily C member 74(OR6C74) | NM_001005490 | 117.6 | 92.6 | 0.0020 | 0.83 |
| ZNF669 | zinc finger protein 669(ZNF669) | NM_024804 | 229.4 | 271.1 | 0.0020 | 1.3 |
| ZNF669 | zinc finger protein 669(ZNF669) | NM_024804 | 490.1 | 450 | 0.0020 | 1.3 |
| LRRC29 | leucine rich repeat containing 29(LRRC29) | NM_001004055 | 215.6 | 135.7 | 0.0020 | 0.71 |
| CDHR2 | cadherin related family member 2(CDHR2) | NM_017675 | 140.3 | 106.7 | 0.0020 | 0.83 |
| | | XM_926303 | 117.5 | 95 | 0.0020 | 0.88 |

| | | | | | | |
|---|---|---|---|---|---|---|
| PCYT2 | phosphate cytidylyltransferase 2, ethanolamine(PCYT2) | NM_002861 | 161.6 | 104.6 | 0.0020 | 0.71 |
| | | XM_942896 | 174.8 | 122.8 | 0.0020 | 0.79 |
| CLK2P1 | CDC like kinase 2, pseudogene 1(CLK2P1) | NR_002711 | 117.3 | 141.4 | 0.0020 | 1.35 |
| C2orf78 | chromosome 2 open reading frame 78(C2orf78) | NM_001080474 | 116.5 | 93.7 | 0.0020 | 0.84 |
| PNLDC1 | PARN like, ribonuclease domain containing 1(PNLDC1) | NM_173516 | 97.6 | 105.4 | 0.0020 | 1.19 |
| AMMECR1L | AMMECR1 like(AMMECR1L) | NM_031445 | 134.5 | 164.7 | 0.0020 | 1.38 |
| AGO2 | argonaute 2, RISC catalytic component(AGO2) | NM_012154 | 221.4 | 378.9 | 0.0020 | 1.91 |
| MYLK | myosin light chain kinase(MYLK) | NM_053026 | 142.6 | 161.7 | 0.0020 | 1.29 |
| | | XM_001726093 | 161.4 | 114.8 | 0.0020 | 0.79 |

| | | | | | | | | | |
|---|---|---|---|---|---|---|---|---|---|
| SMG1P2 | SMG1P2, nonsense mediated mRNA decay associated PI3K related kinase pseudogene 2(SMG1P2) | NR_002473 | 118.5 | 129.8 | 0.0020 | 1.22 | | | |
| DENND1B | DENN domain containing 1B(DENND1B) | NM_019049 | 175.9 | 218.2 | 0.0020 | 1.46 | | | |
| MSL3 | male-specific lethal 3 homolog (Drosophila)(MSL3) | NM_078629 | 177.7 | 374.5 | 0.0020 | 2.39 | NM_001107943 | 0.63 | 112.77 |
| MSL3 | male-specific lethal 3 homolog (Drosophila)(MSL3) | NM_078629 | 179.7 | 311 | 0.0020 | 2.39 | | | |
| | | XM_933942 | 144.2 | 111.9 | 0.0020 | 0.85 | | | |
| CCNDBP1 | cyclin D1 binding protein 1(CCNDBP1) | NM_037370 | 391.9 | 472.1 | 0.0020 | 1.39 | | | |
| Snora2b | small nucleolar RNA, H/ACA box 2B(SNORA2B) | NR_002951 | 114.4 | 136 | 0.0020 | 1.36 | | | |
| GSR | glutathione-disulfide reductase(GSR) | NM_000637 | 93.1 | 104.7 | 0.0020 | 1.22 | | | |

| | | | | | | | | | |
|---|---|---|---|---|---|---|---|---|---|
| GSR | glutathione-disulfide reductase(GSR) | NM_000637 | 127.3 | 104.6 | 0.0020 | 1.22 | | | |
| PLEKHG7 | pleckstrin homology and RhoGEF domain containing G7(PLEKHG7) | NM_001004330 | 98.6 | 108.1 | 0.0020 | 1.2 | | | |
| LYPD4 | LY6/PLAUR domain containing 4(LYPD4) | NM_173506 | 135.8 | 106.4 | 0.0020 | 0.86 | | | |
| VSIG8 | V-set and immunoglobulin domain containing 8(VSIG8) | NM_001013661 | 86 | 100.1 | 0.0020 | 1.26 | | | |
| KIF5A | kinesin family member 5A(KIF5A) | NM_004984 | 146.4 | 111.7 | 0.0020 | 0.84 | | | |
| RABGAP1 | RAB GTPase activating protein 1(RABGAP1) | NM_012197 | 309 | 385.1 | 0.0020 | 1.38 | | | |
| PMS1 | PMS1 homolog 1, mismatch repair system component(PMS1) | NM_000534 | 98.7 | 107 | 0.0020 | 1.18 | NM_134372 | 22.51 | 0.03 |
| | | CD242195 | 119.9 | 94.6 | 0.0020 | 0.84 | | | |

| | | | | | | |
|---|---|---|---|---|---|---|
| ZRANB2 | zinc finger RANBP2-type containing 2(ZRANB2) | NM_005455 | 117.1 | 118.4 | 0.0020 | 1.37 |
| ZRANB2 | zinc finger RANBP2-type containing 2(ZRANB2) | NM_005455 | 155.9 | 186.5 | 0.0020 | 1.37 |
| | | XR_038027 | 89.7 | 98.5 | 0.0020 | 1.19 |
| | | XR_038027 | 113.2 | 100.4 | 0.0020 | 1.19 |
| ASXL2 | additional sex combs like 2, transcriptional regulator(ASXL2) | NM_018263 | 130.2 | 206.1 | 0.0020 | 1.82 |
| TTC14 | tetratricopeptide repeat domain 14(TTC14) | NM_001042601 | 252.9 | 364.7 | 0.0020 | 1.62 |
| STAU1 | staufen double-stranded RNA binding protein 1(STAU1) | NM_017453 | 95.8 | 112.9 | 0.0020 | 1.31 |
| | | BM715751 | 123.2 | 96.7 | 0.0030 | 0.86 |
| CCDC9 | coiled-coil domain containing 9(CCDC9) | NM_015603 | 106.3 | 111.7 | 0.0030 | 1.16 |

| | | | | | | |
|---|---|---|---|---|---|---|
| RMND5B | required for meiotic nuclear division 5 homolog B(RMND5B) | NM_022762 | 120.5 | 133 | 0.0030 | 1.27 |
| RRS1 | ribosome biogenesis regulator homolog(RRS1) | NM_015169 | 119.4 | 131.8 | 0.0030 | 1.23 |
| | | XM_0011126824 | 150.6 | 111.8 | 0.0030 | 0.82 |
| MTERF1 | mitochondrial transcription termination factor 1(MTERF1) | NM_006980 | 156.3 | 192.2 | 0.0030 | 1.41 |
| TOX2 | TOX high mobility group box family member 2(TOX2) | NM_001098796 | 95.6 | 102.3 | 0.0030 | 1.17 |
| WRN | Werner syndrome RecQ like helicase(WRN) | NM_000553 | 119.9 | 136.4 | 0.0030 | 1.29 |
| ARMCX6 | armadillo repeat containing, X-linked 6(ARMCX6) | NM_019007 | 99 | 97.4 | 0.0030 | 1.3 |
| ARMCX6 | armadillo repeat containing, X-linked 6(ARMCX6) | NM_019007 | 115.1 | 132.9 | 0.0030 | 1.3 |

| | | | | | | | | | |
|---|---|---|---|---|---|---|---|---|---|
| FAM98C | family with sequence similarity 98 member C(FAM98C) | NM_174905 | 153.5 | 177 | 0.0030 | 1.32 | | | |
| ZSCAN9 | zinc finger and SCAN domain containing 9(ZSCAN9) | NM_006299 | 93.8 | 113.4 | 0.0030 | 1.32 | | | |
| SIRT5 | sirtuin 5(SIRT5) | NM_031244 | 101.3 | 117.7 | 0.0030 | 1.28 | | | |
| SIRT5 | sirtuin 5(SIRT5) | NM_031244 | 107.1 | 102.1 | 0.0030 | 1.28 | | | |
| MTF2 | metal response element binding transcription factor 2(MTF2) | NM_007358 | 177.6 | 318 | 0.0030 | 2.02 | NM_001031645 | 1.93 | 10.04 |
| aste1 | asteroid homolog 1 (Drosophila)(ASTE1) | NM_014065 | 90.3 | 102.8 | 0.0030 | 1.22 | | | |
| ADGRF5 | adhesion G protein-coupled receptor F5(ADGRF5) | NM_015234 | 1392.1 | 1820.8 | 0.0030 | 1.45 | | | |
| | | AK095804 | 139.3 | 101.1 | 0.0030 | 0.78 | | | |
| TADA2A | transcriptional adaptor 2A(TADA2A) | NM_001488 | 101.5 | 107.8 | 0.0030 | 1.17 | | | |
| | | XM_930777 | 109.7 | 126.2 | 0.0030 | 1.29 | | | |

| | | | | | | | | | |
|---|---|---|---|---|---|---|---|---|---|
| UGCG | UDP-glucose ceramide glucosyltransferase(UGCG) | NM_003358 | 265.4 | 517.2 | 0.0030 | 2.22 | | | |
| C11orf84 | chromosome 11 open reading frame 84(C11orf84) | NM_138471 | 99.7 | 107.1 | 0.0030 | 1.18 | | | |
| | | AA651832 | 100.5 | 106 | 0.0030 | 1.17 | | | |
| RIMKLB | ribosomal modification protein rimK like family member B(RIMKLB) | NM_020734 | 106.4 | 116.8 | 0.0030 | 1.21 | | | |
| | | XR_038607 | 111.1 | 129.7 | 0.0030 | 1.32 | | | |
| PSRC1 | proline and serine rich coiled-coil 1(PSRC1) | NM_001032290 | 137.2 | 107.7 | 0.0030 | 0.86 | | | |
| LILRA4 | leukocyte immunoglobulin like receptor A4(LILRA4) | NM_012276 | 93.9 | 107 | 0.0030 | 1.24 | | | |
| | | XM_930350 | 108.5 | 121.4 | 0.0030 | 1.27 | | | |
| | | XM_928281 | 108.4 | 118.5 | 0.0030 | 1.2 | | | |
| | | XM_928619 | 177.5 | 132.5 | 0.0040 | 0.85 | NM_144748 | 66.99 | 0.14 |

| | | | | | | | | | |
|---|---|---|---|---|---|---|---|---|---|
| TRIM36 | tripartite motif containing 36(TRIM36) | NM_001017397 | 109.7 | 87.1 | 0.0040 | 0.84 | | | |
| IFITM4P | interferon induced transmembrane protein 4 pseudogene(IFITM4P) | NR_001590 | 143.1 | 107.8 | 0.0040 | 0.82 | | | |
| LOC102724002 | uncharacterized LOC102724002(LOC102724002) | AK025166 | 167 | 190.7 | 0.0040 | 1.31 | | | |
| UBE2E3 | ubiquitin conjugating enzyme E2 E3(UBE2E3) | NM_006357 | 127.3 | 102.7 | 0.0040 | 0.87 | | | |
| UBE2E3 | ubiquitin conjugating enzyme E2 E3(UBE2E3) | NM_006357 | 157.4 | 139.3 | 0.0040 | 0.87 | | | |
| UBE2E3 | ubiquitin conjugating enzyme E2 E3(UBE2E3) | NM_006357 | 431 | 339.9 | 0.0040 | 0.87 | | | |
| | | XR_015946 | 132.6 | 140.9 | 0.0040 | 1.21 | | | |
| | | XR_015946 | 228 | 225.6 | 0.0040 | 1.21 | NM_001107793 | 77.29 | 6.85 |
| LOC643923 | uncharacterized LOC643923(LOC643923) | NM_001039772 | 94.1 | 107 | 0.0040 | 1.25 | NM_001107793 | 77.29 | 6.85 |

| | | | | | | | | | |
|---|---|---|---|---|---|---|---|---|---|
| CHRNA5 | cholinergic receptor nicotinic alpha 5 subunit(CHRNA5) | NM_000745 | 897.2 | 1337 | 0.0040 | 1.64 | NM_001107793 | 77.29 | 6.85 |
| LRRK2 | leucine rich repeat kinase 2(LRRK2) | NM_198578 | 99.3 | 104.1 | 0.0040 | 1.14 | | | |
| DCAF16 | DDB1 and CUL4 associated factor 16(DCAF16) | NM_017741 | 99.8 | 106.4 | 0.0040 | 1.15 | | | |
| ZNF317 | zinc finger protein 317(ZNF317) | NM_020933 | 127.1 | 143.5 | 0.0040 | 1.29 | | | |
| DLGAP4 | DLG associated protein 4(DLGAP4) | NM_183006 | 117.8 | 96.9 | 0.0040 | 0.89 | | | |
| RPN2 | ribophorin II(RPN2) | NM_002951 | 285.7 | 330.9 | 0.0040 | 1.29 | | | |
| | | XR_037397 | 93.1 | 104.6 | 0.0040 | 1.23 | | | |
| | | XM_372319 | 144.1 | 108.4 | 0.0040 | 0.82 | | | |
| TTLL3 | tubulin tyrosine ligase like 3(TTLL3) | NM_015644 | 126.8 | 102.6 | 0.0040 | 0.87 | | | |
| TTLL3 | tubulin tyrosine ligase like 3(TTLL3) | NM_015644 | 160.9 | 152.3 | 0.0040 | 0.87 | | | |
| NEK8 | NIMA related kinase 8(NEK8) | NM_178170 | 125.8 | 136.7 | 0.0040 | 1.25 | | | |

| | | | | | | |
|---|---|---|---|---|---|---|
| | | XM_927957 | 130.6 | 98.5 | 0.0040 | 0.81 |
| C9orf85 | chromosome 9 open reading frame 85(C9orf85) | NM_182505 | 116.8 | 133.4 | 0.0040 | 1.27 |
| TXLNGY | taxilin gamma pseudogene, Y-linked(TXLNGY) | NM_0010058 52 | 88.2 | 106.8 | 0.0040 | 1.32 |
| ZNF148 | zinc finger protein 148(ZNF148) | NM_021964 | 585.7 | 881.1 | 0.0040 | 1.69 |
| RASAL2 | RAS protein activator like 2(RASAL2) | NM_170692 | 92.3 | 104.5 | 0.0040 | 1.22 |
| M6PR | mannose-6-phosphate receptor, cation dependent(M6PR) | NM_002355 | 675 | 1018 | 0.0040 | 1.64 |
| WDFY3 | WD repeat and FYVE domain containing 3(WDFY3) | NM_178583 | 113.6 | 92.1 | 0.0050 | 0.85 |
| SCARNA18 | small Cajal body-specific RNA 18(SCARNA18) | NR_003139 | 102.7 | 117.9 | 0.0050 | 1.29 |
| PIP5K1B | phosphatidylinositol-4-phosphate 5-kinase type 1 beta(PIP5K1B) | NM_003558 | 99.6 | 114.7 | 0.0050 | 1.27 |

| | | | | | | |
|---|---|---|---|---|---|---|
| | | XM_001720423 | 149.9 | 108 | 0.0050 | 0.81 |
| SCML1 | sex comb on midleg-like 1 (Drosophila)(SCML1) | NM_001037540 | 123.8 | 127.6 | 0.0050 | 1.21 |
| SCML1 | sex comb on midleg-like 1 (Drosophila)(SCML1) | NM_001037540 | 128.5 | 137.4 | 0.0050 | 1.21 |
| | | XR_017689 | 128.7 | 105.4 | 0.0050 | 0.89 |
| | | XR_038018 | 343.3 | 198.7 | 0.0050 | 0.66 |
| TRIM52 | tripartite motif containing 52(TRIM52) | NM_032765 | 109.9 | 143.1 | 0.0050 | 1.46 |
| | | XR_015455 | 131.6 | 112.3 | 0.0050 | 1.36 |
| | | XR_015455 | 134.9 | 157 | 0.0050 | 1.36 |
| MAP3K7 | mitogen-activated protein kinase kinase kinase 7(MAP3K7) | NM_145331 | 159.4 | 212.6 | 0.0050 | 1.49 |
| | | AB074162 | 112.2 | 132.5 | 0.0050 | 1.33 |
| HJURP | Holliday junction recognition protein(HJURP) | NM_018410 | 93.4 | 103.8 | 0.0050 | 1.21 |

| | | | | | | | | | |
|---|---|---|---|---|---|---|---|---|---|
| HNRNPA3P1 | heterogeneous nuclear ribonucleoprotein A3 pseudogene 1(HNRNPA3P1) | NR_002726 | 114.9 | 124 | 0.0050 | 1.21 | | | |
| BAGE5 | B melanoma antigen family member 5(BAGE5) | NM_182484 | 110.4 | 112.5 | 0.0050 | 1.13 | NM_001009603 | 28.32 | 0.90 |
| | | XR_037956 | 121.3 | 93.8 | 0.0050 | 0.84 | | | |
| SRP19 | signal recognition particle 19(SRP19) | NM_003135 | 216.5 | 275.1 | 0.0050 | 1.4 | | | |
| CHD2 | chromodomain helicase DNA binding protein 2(CHD2) | NM_001042572 | 88.6 | 103 | 0.0050 | 1.27 | | | |
| | | AW297854 | 122.1 | 97.9 | 0.0050 | 0.86 | | | |
| BCLAF1 | BCL2 associated transcription factor 1(BCLAF1) | NM_001077440 | 107.8 | 132.4 | 0.0050 | 1.36 | | | |
| BCLAF1 | BCL2 associated transcription factor 1(BCLAF1) | NM_001077440 | 556.2 | 636.2 | 0.0050 | 1.36 | | | |
| CBLL1 | Cbl proto-oncogene like 1(CBLL1) | NM_024814 | 114 | 91.5 | 0.0050 | 0.86 | | | |
| CBLL1 | Cbl proto-oncogene like 1(CBLL1) | NM_024814 | 141.2 | 115.2 | 0.0050 | 0.86 | | | |

| | | | | | | |
|---|---|---|---|---|---|---|
| | | XM_927142 | 102.6 | 112.5 | 0.0050 | 1.2 |
| FBXO11 | F-box protein 11(FBXO11) | NM_025133 | 351.7 | 374.5 | 0.0050 | 1.31 |
| FBXO11 | F-box protein 11(FBXO11) | NM_025133 | 514.1 | 596.4 | 0.0050 | 1.31 |
| CDK11A | cyclin dependent kinase 11A(CDK11A) | NM_024011 | 112.4 | 131.7 | 0.0050 | 1.29 |
| KAZN | kazrin, periplakin interacting protein(KAZN) | NM_001018000 | 128.2 | 152.1 | 0.0050 | 1.37 |
| KAZN | kazrin, periplakin interacting protein(KAZN) | NM_001018000 | 212.7 | 253.4 | 0.0050 | 1.37 |
| SF3B1 | splicing factor 3b subunit 1(SF3B1) | NM_012433 | 114.4 | 148.3 | 0.0050 | 1.43 |
| SF3B1 | splicing factor 3b subunit 1(SF3B1) | NM_012433 | 542.8 | 636.6 | 0.0050 | 1.43 |
| FAM122A | family with sequence similarity 122A(FAM122A) | NM_138333 | 118.4 | 96.9 | 0.0060 | 0.89 |
| CPNE1 | copine 1(CPNE1) | NM_003915 | 96.3 | 103.7 | 0.0060 | 1.17 |
| | | XM_929565 | 217.9 | 143.8 | 0.0060 | 0.76 |

| | | | | | | |
|---|---|---|---|---|---|---|
| PITPNC 1 | phosphatidylinositol transfer protein, cytoplasmic 1(PITPNC1) | NM_181671 | 142.3 | 159.1 | 0.0060 | 1.27 |
| | | BQ027703 | 100.5 | 105.4 | 0.0060 | 1.14 |
| | | BM668469 | 112.9 | 119.6 | 0.0060 | 1.19 |
| mapk13 | mitogen-activated protein kinase 13(MAPK13) | NM_002754 | 106.1 | 114.2 | 0.0060 | 1.19 |
| SCLT1 | sodium channel and clathrin linker 1(SCLT1) | NM_144643 | 88.3 | 102 | 0.0060 | 1.24 |
| CRCP | CGRP receptor component(CRCP) | NM_014478 | 115.9 | 112.3 | 0.0060 | 1.46 |
| CRCP | CGRP receptor component(CRCP) | NM_014478 | 2824.8 | 3977.5 | 0.0060 | 1.46 |
| RBBP5 | RB binding protein 5, histone lysine methyltransferase complex subunit(RBBP5) | NM_005057 | 114.2 | 114 | 0.0060 | 1.11 |
| | | XM_934567 | 123.1 | 126.1 | 0.0060 | 1.16 |
| | | NM_199345 | 148.2 | 185.7 | 0.0060 | 1.42 |
| | | BX105338 | 133 | 194.5 | 0.0060 | 1.65 |

| | | | | | | |
|---|---|---|---|---|---|---|
| FAM50A | family with sequence similarity 50 member A(FAM50A) | NM_004699 | 164.6 | 179.7 | 0.0060 | 1.26 |
| | | DB038929 | 143.4 | 110.9 | 0.0060 | 0.84 |
| | | XM_001134215 | 124.1 | 116 | 0.0060 | 1.44 |
| | | XM_001134215 | 216.9 | 275.7 | 0.0060 | 1.44 |
| S100A13 | S100 calcium binding protein A13(S100A13) | NM_001024211 | 122.2 | 128.7 | 0.0060 | 0.77 |
| S100A13 | S100 calcium binding protein A13(S100A13) | NM_001024211 | 170.6 | 116.5 | 0.0060 | 0.77 |
| MTIF2 | mitochondrial translational initiation factor 2(MTIF2) | NM_001005369 | 553.3 | 724.5 | 0.0060 | 1.44 |
| MEIS3P1 | Meis homeobox 3 pseudogene 1(MEIS3P1) | NR_002211 | 212.4 | 240.6 | 0.0060 | 1.27 |
| | | XM_001128983 | 102.6 | 111 | 0.0060 | 1.19 |
| | | XM_001713654 | 6200.5 | 5439.2 | 0.0060 | 0.85 |
| ARHGEF7 | Rho guanine nucleotide exchange factor 7(ARHGEF7) | NM_145735 | 86.5 | 108.6 | 0.0060 | 1.37 |

| | | | | | | |
|---|---|---|---|---|---|---|
| GCC1 | GRIP and coiled-coil domain containing 1(GCC1) | NM_024523 | 193.4 | 216.1 | 0.0060 | 1.28 |
| | | XM_946257 | 101.3 | 107.6 | 0.0060 | 1.17 |
| INTS2 | integrator complex subunit 2(INTS2) | NM_020748 | 154.7 | 184.2 | 0.0060 | 1.36 |
| UGGT1 | UDP-glucose glycoprotein glucosyltransferase 1(UGGT1) | NM_020120 | 119.2 | 151.8 | 0.0060 | 1.44 |
| | | AI905886 | 121.5 | 95.1 | 0.0060 | 0.83 |
| | | AY475219 | 89.2 | 104.7 | 0.0060 | 1.27 |
| | | XM_933437 | 144.3 | 100.8 | 0.0060 | 0.75 |
| PEX11B | peroxisomal biogenesis factor 11 beta(PEX11B) | NM_003846 | 506.9 | 289.2 | 0.0060 | 0.65 |
| | | XM_001131379 | 105.3 | 110.6 | 0.0060 | 1.15 |
| RBM6 | RNA binding motif protein 6(RBM6) | NM_005777 | 118.2 | 123.5 | 0.0060 | 1.18 |
| NRDE2 | NRDE-2, necessary for RNA interference, domain containing(NRDE2) | NM_017970 | 152.5 | 192.5 | 0.0060 | 1.44 |

| | | | | | | |
|---|---|---|---|---|---|---|
| | | XM_932354 | 142.7 | 107 | 0.0060 | 0.83 |
| | | XM_001126418 | 114.3 | 117.8 | 0.0060 | 1.14 |
| C1orf101 | chromosome 1 open reading frame 101(C1orf101) | NM_173807 | 138.7 | 108 | 0.0060 | 0.85 |
| MTMR14 | myotubularin related protein 14(MTMR14) | NM_001077525 | 133.8 | 139.1 | 0.0060 | 1.25 |
| MTMR14 | myotubularin related protein 14(MTMR14) | NM_001077525 | 169.9 | 186.3 | 0.0060 | 1.25 |
| DDIT4 | DNA damage inducible transcript 4(DDIT4) | NM_019058 | 131.9 | 102.7 | 0.0060 | 0.86 |
| MS4A6A | membrane spanning 4-domains A6A(MS4A6A) | NM_022349 | 98.3 | 104.3 | 0.0060 | 1.16 |
| BAGE3 | B melanoma antigen family member 3(BAGE3) | NM_182481 | 87.8 | 97.7 | 0.0060 | 1.19 |
| RNF220 | ring finger protein 220(RNF220) | NM_018150 | 208 | 238.3 | 0.0060 | 1.3 |

| | | | | | | |
|---|---|---|---|---|---|---|
| PGA3 | pepsinogen 3, group I (pepsinogen A)(PGA3) | NM_001079807 | 90.1 | 94.4 | 0.0070 | 0.84 |
| PGA3 | pepsinogen 3, group I (pepsinogen A)(PGA3) | NM_001079807 | 113.2 | 99.3 | 0.0070 | 0.84 |
| PGA3 | pepsinogen 3, group I (pepsinogen A)(PGA3) | NM_001079807 | 130.2 | 101.7 | 0.0070 | 0.84 |
| | | AI911349 | 142.2 | 108.9 | 0.0070 | 0.82 |
| SULT1A1 | sulfotransferase family 1A member 1(SULT1A1) | NM_177530 | 2668.3 | 3612.2 | 0.0070 | 1.4 |
| | | XM_498969 | 203.5 | 118.8 | 0.0070 | 0.65 |
| HCFC2 | host cell factor C2(HCFC2) | NM_013320 | 110.7 | 116.6 | 0.0070 | 1.16 |
| TMEM263 | transmembrane protein 263(TMEM263) | NM_152261 | 93 | 101.2 | 0.0070 | 1.19 |
| | | XR_019471 | 956.6 | 1383.3 | 0.0070 | 1.57 |
| WDR37 | WD repeat domain 37(WDR37) | NM_014023 | 121.7 | 139.7 | 0.0070 | 1.29 |

| | | | | | | |
|---|---|---|---|---|---|---|
| WDR37 | WD repeat domain 37(WDR37) | NM_014023 | 150.9 | 176.7 | 0.0070 | 1.29 |
| AKAP10 | A-kinase anchoring protein 10(AKAP10) | NM_007202 | 121.3 | 135.3 | 0.0070 | 1.26 |
| | | XM_001716137 | 99.1 | 104.8 | 0.0070 | 1.15 |
| MACF1 | microtubule-actin crosslinking factor 1(MACF1) | NM_012090 | 101.9 | 113.5 | 0.0070 | 1.23 |
| MACF1 | microtubule-actin crosslinking factor 1(MACF1) | NM_012090 | 486.3 | 494.9 | 0.0070 | 1.23 |
| GSK3B | glycogen synthase kinase 3 beta(GSK3B) | NM_002093 | 234.2 | 260.7 | 0.0070 | 1.25 |
| APH1A | aph-1 homolog A, gamma-secretase subunit(APH1A) | NM_001077628 | 213.3 | 181 | 0.0070 | 0.77 |
| APH1A | aph-1 homolog A, gamma-secretase subunit(APH1A) | NM_001077628 | 275.7 | 186.2 | 0.0070 | 0.77 |
| NACC2 | NACC family member 2(NACC2) | AK094914 | 443.2 | 275.7 | 0.0070 | 0.69 |
| MGA | MGA, MAX dimerization protein(MGA) | NM_001080541 | 102.8 | 104.7 | 0.0070 | 0.85 |

| | | | | | | | | | |
|---|---|---|---|---|---|---|---|---|---|
| MGA | MGA, MAX dimerization protein(MGA) | NM_001080541 | 155.6 | 117.3 | 0.0070 | 0.85 | | | |
| PPFIBP1 | PPFIA binding protein 1(PPFIBP1) | NM_003622 | 99.8 | 114.9 | 0.0070 | 1.25 | | | |
| PPFIBP1 | PPFIA binding protein 1(PPFIBP1) | NM_003622 | 111 | 103 | 0.0070 | 1.25 | | | |
| PPFIBP1 | PPFIA binding protein 1(PPFIBP1) | NM_003622 | 168.4 | 124 | 0.0070 | 1.25 | | | |
| ZNF654 | zinc finger protein 654(ZNF654) | NM_018293 | 89.3 | 100 | 0.0070 | 1.21 | NM_144744 | 0.11 | 40.01 |
| ZNF654 | zinc finger protein 654(ZNF654) | NM_018293 | 102 | 94.7 | 0.0070 | 1.21 | | | |
| ZNF654 | zinc finger protein 654(ZNF654) | NM_018293 | 102.6 | 101.7 | 0.0070 | 1.21 | | | |
| ZNF654 | zinc finger protein 654(ZNF654) | NM_018293 | 102.6 | 105.3 | 0.0070 | 1.21 | | | |
| | | BQ447196 | 99.1 | 104.7 | 0.0070 | 1.15 | | | |
| | | AA430240 | 92.9 | 100.6 | 0.0070 | 1.16 | | | |
| OR2G2 | olfactory receptor family 2 subfamily G member 2(OR2G2) | NM_001001915 | 148.6 | 101 | 0.0070 | 0.74 | | | |
| | | XM_001723369 | 125.5 | 102.3 | 0.0070 | 0.88 | | | |
| ZMYND11 | zinc finger MYND-type containing 11(ZMYND11) | NM_006624 | 93.7 | 91.5 | 0.0070 | 1.3 | | | |

| | | | | | | |
|---|---|---|---|---|---|---|
| ZMYND 11 | zinc finger MYND-type containing 11(ZMYND11) | NM_00662 4 | 1043.1 | 1211.7 | 0.0070 | 1.3 |
| | | XR_03935 7 | 101 | 114.2 | 0.0070 | 1.24 |
| | | AK024852 | 109.8 | 145 | 0.0070 | 1.5 |
| FXYD5 | FXYD domain containing ion transport regulator 5(FXYD5) | NM_14477 9 | 103.4 | 180.9 | 0.0070 | 1.14 |
| FXYD5 | FXYD domain containing ion transport regulator 5(FXYD5) | NM_14477 9 | 109.4 | 113 | 0.0070 | 1.14 |
| DUX4L1 | double homeobox 4 like 1(DUX4L1) | NM_03317 8 | 113.3 | 97.3 | 0.0070 | 0.93 |
| EIF5A | eukaryotic translation initiation factor 5A(EIF5A) | NM_00197 0 | 412.5 | 457.4 | 0.0070 | 1.26 |
| sorbs1 | sorbin and SH3 domain containing 1(SORBS1) | NM_00103 4954 | 101.8 | 108.2 | 0.0070 | 1.16 |
| sorbs1 | sorbin and SH3 domain containing 1(SORBS1) | NM_00103 4954 | 260.9 | 275.7 | 0.0070 | 1.16 |

| | | | | | | |
|---|---|---|---|---|---|---|
| SELENOP | selenoprotein P(SELENOP) | NM_005410 | 116.2 | 85.6 | 0.0070 | 0.79 |
| CCL25 | C-C motif chemokine ligand 25(CCL25) | NM_005624 | 152.6 | 112.7 | 0.0070 | 0.82 |
| STX16 | syntaxin 16(STX16) | NM_001001433 | 137.2 | 167 | 0.0070 | 1.39 |
| STX16 | syntaxin 16(STX16) | NM_001001433 | 288.6 | 381.4 | 0.0070 | 1.39 |
| | | XR_039439 | 110.9 | 86.2 | 0.0080 | 0.81 |
| | | XR_039439 | 649.4 | 557.2 | 0.0080 | 0.81 |
| EXTL2 | exostosin like glycosyltransferase 2(EXTL2) | NM_001033025 | 148.9 | 170.7 | 0.0080 | 1.29 |
| ALX3 | ALX homeobox 3(ALX3) | NM_006492 | 121.1 | 106.9 | 0.0080 | 0.85 |
| ALX3 | ALX homeobox 3(ALX3) | NM_006492 | 132.8 | 105.6 | 0.0080 | 0.85 |
| MPPE1 | metallophosphoesterase 1(MPPE1) | NM_023075 | 161 | 194.9 | 0.0080 | 1.38 |
| MCM3AP | minichromosome maintenance complex component 3 associated protein(MCM3AP) | NM_003906 | 101.4 | 107.7 | 0.0080 | 1.18 |

| | | | | | | |
|---|---|---|---|---|---|---|
| OR2B11 | olfactory receptor family 2 subfamily B member 11(OR2B11) | NM_001004492 | 131.4 | 101.4 | 0.0080 | 0.83 |
| OCIAD2 | OCIA domain containing 2(OCIAD2) | NM_152398 | 147.5 | 101.9 | 0.0080 | 0.74 |
| spop | speckle type BTB/POZ protein(SPOP) | NM_001007230 | 556.6 | 727.7 | 0.0080 | 1.48 |
| ADGRG2 | adhesion G protein-coupled receptor G2(ADGRG2) | NM_001079859 | 96.3 | 103.4 | 0.0080 | 1.15 |
| CYP11A1 | cytochrome P450 family 11 subfamily A member 1(CYP11A1) | NM_000781 | 145.6 | 113.6 | 0.0080 | 0.86 |
| | | XM_937648 | 134.9 | 106.7 | 0.0080 | 0.85 |
| ZC2HC1A | zinc finger C2HC-type containing 1A(ZC2HC1A) | NM_016010 | 101.2 | 106.2 | 0.0080 | 1.38 |
| ZC2HC1A | zinc finger C2HC-type containing 1A(ZC2HC1A) | NM_016010 | 126.1 | 152 | 0.0080 | 1.38 |
| TLR2 | toll like receptor 2(TLR2) | NM_003264 | 98.5 | 127.1 | 0.0080 | 1.45 |
| tcf20 | transcription factor 20(TCF20) | NM_181492 | 107.8 | 101 | 0.0080 | 1.38 |

| | | | | | | |
|---|---|---|---|---|---|---|
| tcf20 | transcription factor 20(TCF20) | NM_181492 | 242.8 | 304.4 | 0.0080 | 1.38 |
| | | XM_926130 | 110.9 | 141.1 | 0.0080 | 1.44 |
| USP33 | ubiquitin specific peptidase 33(USP33) | NM_015017 | 102.4 | 117.1 | 0.0080 | 1.27 |
| ZNF565 | zinc finger protein 565(ZNF565) | NM_152477 | 116.1 | 127.7 | 0.0080 | 1.23 |
| ZNF565 | zinc finger protein 565(ZNF565) | NM_152477 | 121 | 127.7 | 0.0080 | 1.23 |
| POU2F1 | POU class 2 homeobox 1(POU2F1) | NM_002697 | 98.3 | 114.2 | 0.0080 | 1.27 |
| hist1h4c | histone cluster 1 H4 family member c(HIST1H4C) | NM_003542 | 397.6 | 1481.1 | 0.0080 | 3.75 |
| CNNM3 | cyclin and CBS domain divalent metal cation transport mediator 3(CNNM3) | NM_017623 | 108.2 | 93.2 | 0.0080 | 1.2 |
| CNNM3 | cyclin and CBS domain divalent metal cation transport mediator 3(CNNM3) | NM_017623 | 110.7 | 121.7 | 0.0080 | 1.2 |

| | | | | | | |
|---|---|---|---|---|---|---|
| FBXW2 | F-box and WD repeat domain containing 2(FBXW2) | NM_012164 | 115.3 | 135.4 | 0.0080 | 1.34 |
| PHF14 | PHD finger protein 14(PHF14) | NM_014660 | 120.2 | 100.5 | 0.0080 | 0.91 |
| FOXJ2 | forkhead box J2(FOXJ2) | NM_018416 | 431.6 | 548 | 0.0080 | 1.47 |
| or8g1 | olfactory receptor family 8 subfamily G member 1 (gene/pseudogene)(OR8G1) | NM_001002905 | 111.3 | 90.1 | 0.0080 | 0.85 |
| SKP2 | S-phase kinase associated protein 2(SKP2) | NM_032637 | 104.4 | 113.8 | 0.0090 | 1.21 |
| | | XR_017962 | 118.5 | 98.1 | 0.0090 | 0.84 |
| | | XR_017962 | 129.4 | 99.9 | 0.0090 | 0.84 |
| | | XR_039678 | 126.8 | 92.9 | 0.0090 | 0.79 |
| MAGEB1 | MAGE family member B1(MAGEB1) | NM_002363 | 131.2 | 101.1 | 0.0090 | 0.84 |
| FMC1 | formation of mitochondrial complex V assembly factor 1 homolog(FMC1) | NM_197964 | 268.8 | 199.6 | 0.0090 | 1.43 |

| | | | | | | |
|---|---|---|---|---|---|---|
| FMC1 | formation of mitochondrial complex V assembly factor 1 homolog(FMC1) | NM_197964 | 1082.9 | 1361.3 | 0.0090 | 1.43 |
| FMC1 | formation of mitochondrial complex V assembly factor 1 homolog(FMC1) | NM_197964 | 1369.2 | 870 | 0.0090 | 1.43 |
| ERGIC1 | endoplasmic reticulum-golgi intermediate compartment 1(ERGIC1) | NM_001031711 | 171.4 | 196.6 | 0.0090 | 1.31 |
| LMBRD2 | LMBR1 domain containing 2(LMBRD2) | NM_001007527 | 112.2 | 121.7 | 0.0090 | 1.19 |
| BPIFA3 | BPI fold containing family A member 3(BPIFA3) | NM_178466 | 92.5 | 98.6 | 0.0090 | 1.14 |
| HLA-DQA2 | major histocompatibility complex, class II, DQ alpha 2(HLA-DQA2) | NM_020056 | 147.6 | 114.8 | 0.0090 | 0.86 |
| ZBED5 | zinc finger BED-type containing 5(ZBED5) | NM_021211 | 286.7 | 329.4 | 0.0090 | 1.55 |

| | | | | | | |
|---|---|---|---|---|---|---|
| ZBED5 | zinc finger BED-type containing 5(ZBED5) | NM_021211 | 496.9 | 685.8 | 0.0090 | 1.55 |
| CLPTM1L | CLPTM1 like(CLPTM1L) | NM_030782 | 222.8 | 269.7 | 0.0090 | 1.42 |
| SMOC1 | SPARC related modular calcium binding 1(SMOC1) | NM_001034852 | 114.4 | 101.6 | 0.0090 | 0.81 |
| SMOC1 | SPARC related modular calcium binding 1(SMOC1) | NM_001034852 | 118.7 | 90.2 | 0.0090 | 0.81 |
| RPS26 | ribosomal protein S26(RPS26) | NM_001029 | 89.2 | 93 | 0.0090 | 1.11 |
| RPS26 | ribosomal protein S26(RPS26) | NM_001029 | 1360.8 | 1438.4 | 0.0090 | 1.11 |
| FDPS | farnesyl diphosphate synthase(FDPS) | NM_002004 | 142.8 | 160.3 | 0.0090 | 1.25 |
| | | NM_020231 | 221.5 | 330.2 | 0.0090 | 1.68 |
| ZZEF1 | zinc finger ZZ-type and EF-hand domain containing 1(ZZEF1) | NM_015113 | 210.3 | 255.3 | 0.0090 | 1.39 |
| KLHL9 | kelch like family member 9(KLHL9) | NM_018847 | 404.9 | 500.1 | 0.0090 | 1.39 |

| | | | | | | |
|---|---|---|---|---|---|---|
| C1orf137 | chromosome 1 open reading frame 137(C1orf137) | NM_001013643 | 142.2 | 106.9 | 0.0090 | 0.83 |
| UGDH | UDP-glucose 6-dehydrogenase(UGDH) | NM_003359 | 122.4 | 92.2 | 0.0090 | 0.81 |
| BDP1 | B double prime 1, subunit of RNA polymerase III transcription initiation factor IIIB(BDP1) | NM_018429 | 118.1 | 120.3 | 0.0090 | 1.14 |
| GABBR1 | gamma-aminobutyric acid type B receptor subunit 1(GABBR1) | NM_021905 | 100.3 | 104.8 | 0.0090 | 1.14 |
| GABBR1 | gamma-aminobutyric acid type B receptor subunit 1(GABBR1) | NM_021905 | 105.7 | 108.6 | 0.0090 | 1.14 |
| | | XR_038461 | 100 | 108.1 | 0.0090 | 1.17 |
| ercc1 | ERCC excision repair 1, endonuclease non-catalytic subunit(ERCC1) | NM_001983 | 108 | 100.1 | 0.0090 | 1.33 |

| | | | | | | |
|---|---|---|---|---|---|---|
| ercc1 | ERCC excision repair 1, endonuclease non-catalytic subunit(ERCC1) | NM_001983 | 165.3 | 196.6 | 0.0090 | 1.33 |
| ercc1 | ERCC excision repair 1, endonuclease non-catalytic subunit(ERCC1) | NM_001983 | 177.3 | 179.5 | 0.0090 | 1.33 |
| UBE2D3 | ubiquitin conjugating enzyme E2 D3(UBE2D3) | NM_181890 | 1309.4 | 1623.6 | 0.0090 | 1.36 |
| UBE2D3 | ubiquitin conjugating enzyme E2 D3(UBE2D3) | NM_181890 | 1649.2 | 2032.9 | 0.0090 | 1.36 |
| WDR20 | WD repeat domain 20(WDR20) | NM_144574 | 124.1 | 97.5 | 0.0090 | 0.82 |
| GRIA3 | glutamate ionotropic receptor AMPA type subunit 3(GRIA3) | NM_181894 | 94.1 | 101 | 0.0090 | 1.16 |
| | | AW051978 | 132.7 | 106.3 | 0.0090 | 0.87 |
| TMEM184C | transmembrane protein 184C(TMEM184C) | NM_018241 | 99.6 | 103.1 | 0.0100 | 1.11 |

| | | | | | | |
|---|---|---|---|---|---|---|
| TMEM184C | transmembrane protein 184C(TMEM184C) | NM_018241 | 289.7 | 256.8 | 0.0100 | 1.11 |
| KEAP1 | kelch like ECH associated protein 1(KEAP1) | NM_012289 | 211.3 | 176.5 | 0.0100 | 0.75 |
| KEAP1 | kelch like ECH associated protein 1(KEAP1) | NM_012289 | 311.4 | 210.9 | 0.0100 | 0.75 |
| POMGNT2 | protein O-linked mannose N-acetylglucosaminyltransferase 2 (beta 1,4-)(POMGNT2) | NM_032806 | 234.8 | 141.8 | 0.0100 | 0.68 |
| | | XM_929444 | 151.3 | 114.8 | 0.0100 | 0.85 |
| RBM3 | RNA binding motif (RNP1, RRM) protein 3(RBM3) | NM_001017430 | 105.3 | 112.6 | 0.0100 | 1.46 |
| RBM3 | RNA binding motif (RNP1, RRM) protein 3(RBM3) | NM_001017430 | 167.5 | 208.4 | 0.0100 | 1.46 |
| BEND6 | BEN domain containing 6(BEND6) | NM_152731 | 93.3 | 101.8 | 0.0100 | 1.19 |

| | | | | | | | | | |
|---|---|---|---|---|---|---|---|---|---|
| BEND6 | BEN domain containing 6(BEND6) | NM_152731 | 98.5 | 99 | 0.0100 | 1.19 | | | |
| | | XM_0017144670 | 131.9 | 144.9 | 0.0100 | 1.25 | | | |
| | | XM_0011262202 | 146.5 | 194.1 | 0.0100 | 1.51 | | | |
| | | XM_945223 | 118 | 95.1 | 0.0100 | 0.87 | | | |
| trak1 | trafficking kinesin protein 1(TRAK1) | NM_001042646 | 106.4 | 114.6 | 0.0100 | 1.18 | | | |
| trak1 | trafficking kinesin protein 1(TRAK1) | NM_001042646 | 160.3 | 131.9 | 0.0100 | 1.18 | | | |
| TCEAL9 | transcription elongation factor A like 9(TCEAL9) | NM_001006612 | 142.1 | 167.3 | 0.0100 | 1.39 | | | |
| | | XM_939730 | 126 | 98.3 | 0.0100 | 0.85 | | | |
| SNHG3 | small nucleolar RNA host gene 3(SNHG3) | NR_002909 | 120.3 | 96.7 | 0.0100 | 0.87 | NM_017196 | 14.73 | 0.69 |
| fsd1l | fibronectin type III and SPRY domain containing 1 like(FSD1L) | NM_207647 | 102.8 | 111.3 | 0.0100 | 1.18 | NM_017196 | 14.73 | 0.69 |
| fsd1l | fibronectin type III and SPRY domain containing 1 like(FSD1L) | NM_207647 | 126 | 108.7 | 0.0100 | 1.18 | NM_017196 | 14.73 | 0.69 |

| | | | | | | | | | |
|---|---|---|---|---|---|---|---|---|---|
| CGGBP 1 | CGG triplet repeat binding protein 1(CGGBP1) | NM_00366 3 | 599.8 | 867.6 | 0.0100 | 1.58 | | | |
| | | BX106221 | 124.5 | 102 | 0.0100 | 0.9 | | | |
| | | XM_94591 8 | 141 | 107.1 | 0.0100 | 0.83 | | | |
| NUTF2 | nuclear transport factor 2(NUTF2) | NM_00579 6 | 117.6 | 137.9 | 0.0100 | 1.31 | NM_0313 56 | 5.43 | 24.53 |
| | | XM_00171 4845 | 110.4 | 92.4 | 0.0100 | 0.87 | NM_0313 56 | 5.43 | 24.53 |
| TARBP2 | TARBP2, RISC loading complex RNA binding subunit(TARBP2) | NM_13432 3 | 100 | 107.1 | 0.0100 | 1.17 | | | |
| FBXO2 | F-box protein 2(FBXO2) | NM_01216 8 | 216.3 | 144 | 0.0100 | 0.75 | | | |
| CYB561 | cytochrome b561(CYB561) | NM_00101 7917 | 111.9 | 114.3 | 0.0100 | 1.14 | | | |
| IFT88 | intraflagellar transport 88(IFT88) | NM_17560 5 | 90.9 | 102.1 | 0.0100 | 1.21 | | | |
| CARF | calcium responsive transcription factor(CARF) | NM_02474 4 | 103.2 | 106 | 0.0100 | 1.12 | | | |
| CLIC5 | chloride intracellular channel 5(CLIC5) | NM_01692 9 | 196.4 | 347.5 | 0.0100 | 2 | | | |
| | | AA705831 | 119.7 | 100.3 | 0.0100 | 0.89 | | | |

| | | | | | | | | | |
|---|---|---|---|---|---|---|---|---|---|
| | | XR_017601 | 136.4 | 109.8 | 0.0100 | 0.89 | | | |
| | | CN283269 | 135 | 110.1 | 0.0100 | 0.9 | | | |
| | | XM_942071 | 156.1 | 111.3 | 0.0110 | 0.79 | | | |
| | | XM_001726941 | 118.7 | 96.5 | 0.0110 | 0.88 | | | |
| EFNA4 | ephrin A4(EFNA4) | NM_182689 | 99.5 | 102.7 | 0.0110 | 1.13 | | | |
| APOPT1 | apoptogenic 1, mitochondrial(APOPT1) | NM_032374 | 1681.1 | 2076.8 | 0.0110 | 1.31 | | | |
| | | AI082160 | 151 | 116.3 | 0.0110 | 0.85 | | | |
| | | XM_001132307 | 119.6 | 95.6 | 0.0110 | 0.85 | NM_017135 | 41.45 | 2.88 |
| | | XM_001713723 | 18151.9 | 14953.9 | 0.0110 | 0.81 | NM_017135 | 41.45 | 2.88 |
| RERE | arginine-glutamic acid dipeptide repeats(RERE) | NM_012102 | 293.4 | 307.3 | 0.0110 | 1.18 | NM_017135 | 41.45 | 2.88 |
| ADPRHL1 | ADP-ribosylhydrolase like 1(ADPRHL1) | NM_138430 | 145.2 | 117.4 | 0.0110 | 0.89 | NM_017135 | 41.45 | 2.88 |
| VEZT | vezatin, adherens junctions transmembrane protein(VEZT) | NM_017599 | 194.2 | 251 | 0.0110 | 1.44 | | | |
| BARD1 | BRCA1 associated RING domain 1(BARD1) | NM_000465 | 98.6 | 130.8 | 0.0110 | 1.47 | | | |

| | | | | | | | | | |
|---|---|---|---|---|---|---|---|---|---|
| MYO1C | myosin IC(MYO1C) | NM_033375 | 146.8 | 113.7 | 0.0110 | 0.86 | | | |
| FBXW5 | F-box and WD repeat domain containing 5(FBXW5) | NM_018998 | 95 | 101 | 0.0110 | 1.15 | | | |
| AUP1 | ancient ubiquitous protein 1(AUP1) | NM_181575 | 227.6 | 268.1 | 0.0110 | 1.31 | | | |
| mri1 | methylthioribose-1-phosphate isomerase 1(MRI1) | NM_032285 | 122.7 | 138.6 | 0.0110 | 1.28 | | | |
| | | BX108993 | 114.4 | 91.9 | 0.0110 | 0.84 | NM_001033653 | 43.46 | 1.56 |
| | | BE465760 | 192.4 | 102 | 0.0110 | 0.54 | | | |
| ZNF341 | zinc finger protein 341(ZNF341) | NM_032819 | 94.6 | 106.8 | 0.0110 | 1.24 | | | |
| RYBP | RING1 and YY1 binding protein(RYBP) | NM_012234 | 340.3 | 521.8 | 0.0110 | 1.73 | | | |
| | | NR_004402 | 108.4 | 114.4 | 0.0110 | 1.16 | | | |
| LOC105376064 | uncharacterized LOC105376064(LOC105376064) | AW205071 | 201.8 | 129.1 | 0.0110 | 0.73 | | | |
| WDFY2 | WD repeat and FYVE domain containing 2(WDFY2) | NM_052950 | 121.4 | 135.6 | 0.0110 | 1.25 | | | |

| | | | | | | |
|---|---|---|---|---|---|---|
| NDUFB11 | NADH:ubiquinone oxidoreductase subunit B11(NDUFB11) | NM_019056 | 1103.5 | 755.4 | 0.0120 | 0.78 |
| NEUROG3 | neurogenin 3(NEUROG3) | NM_020999 | 174.3 | 121.5 | 0.0120 | 0.78 |
| TNFSF14 | tumor necrosis factor superfamily member 14(TNFSF14) | NM_003807 | 89 | 96.3 | 0.0120 | 1.15 |
| SCYL1 | SCY1 like pseudokinase 1(SCYL1) | NM_020680 | 172.9 | 156 | 0.0120 | 1.22 |
| SCYL1 | SCY1 like pseudokinase 1(SCYL1) | NM_020680 | 174.8 | 182.8 | 0.0120 | 1.22 |
| KCNG2 | potassium voltage-gated channel modifier subfamily G member 2(KCNG2) | NM_012283 | 95.5 | 100.2 | 0.0120 | 1.14 |
| CNOT6L | CCR4-NOT transcription complex subunit 6 like(CNOT6L) | NM_144571 | 131.2 | 163.4 | 0.0120 | 1.37 |
| CSF1 | colony stimulating factor 1(CSF1) | NM_172212 | 113.9 | 93.2 | 0.0120 | 0.87 |
| | | XR_041749 | 223.6 | 118 | 0.0120 | 0.6 |

| | | | | | | | | | |
|---|---|---|---|---|---|---|---|---|---|
| | | BX102609 | 122.5 | 97.7 | 0.0120 | 0.87 | | | |
| ASS1 | argininosuccinate synthase 1(ASS1) | NM_000050 | 127.4 | 94 | 0.0120 | 0.81 | | | |
| ASS1 | argininosuccinate synthase 1(ASS1) | NM_000050 | 1505.7 | 823.6 | 0.0120 | 0.81 | | | |
| PPARD | peroxisome proliferator activated receptor delta(PPARD) | NM_006238 | 94.5 | 98 | 0.0120 | 0.84 | | | |
| PPARD | peroxisome proliferator activated receptor delta(PPARD) | NM_006238 | 129.7 | 102.3 | 0.0120 | 0.84 | | | |
| PTPN9 | protein tyrosine phosphatase, non-receptor type 9(PTPN9) | NM_002833 | 104 | 108 | 0.0120 | 1.13 | | | |
| CEP164 | centrosomal protein 164(CEP164) | NM_014956 | 133.7 | 141.7 | 0.0120 | 1.22 | NM_134407 | 50.47 | 11.16 |
| NIN | ninein(NIN) | NM_020921 | 118.1 | 136.9 | 0.0120 | 1.31 | | | |
| F2R | coagulation factor II thrombin receptor(F2R) | NM_001992 | 16746 | 18218.7 | 0.0120 | 1.21 | | | |
| ACADVL | acyl-CoA dehydrogenase, very long chain(ACADVL) | NM_001033859 | 174.1 | 121.1 | 0.0120 | 0.79 | | | |

| | | | | | | |
|---|---|---|---|---|---|---|
| RCN3 | reticulocalbin 3(RCN3) | NM_020650 | 151.3 | 117.6 | 0.0120 | 0.86 |
| RPUSD4 | RNA pseudouridylate synthase domain containing 4(RPUSD4) | NM_032795 | 117.1 | 123.5 | 0.0120 | 1.19 |
| | | NR_024266 | 118.5 | 126.6 | 0.0120 | 1.2 |
| | | NM_138926 | 157.8 | 211.4 | 0.0120 | 1.48 |
| PCIF1 | PDX1 C-terminal inhibiting factor 1(PCIF1) | NM_022104 | 113.3 | 120.8 | 0.0120 | 1.21 |
| ST5 | suppression of tumorigenicity 5(ST5) | NM_139157 | 92.1 | 100.5 | 0.0120 | 1.17 |
| ST5 | suppression of tumorigenicity 5(ST5) | NM_139157 | 120.2 | 119.8 | 0.0120 | 1.17 |
| NRXN2 | neurexin 2(NRXN2) | NM_138734 | 127.5 | 128.6 | 0.0120 | 0.85 |
| NRXN2 | neurexin 2(NRXN2) | NM_138734 | 159.3 | 121.3 | 0.0120 | 0.85 |
| HMGXB4 | HMG-box containing 4(HMGXB4) | NM_005487 | 107.8 | 99.7 | 0.0120 | 1.17 |
| HMGXB4 | HMG-box containing 4(HMGXB4) | NM_005487 | 113.4 | 120.2 | 0.0120 | 1.17 |

| | | | | | | | | | |
|---|---|---|---|---|---|---|---|---|---|
| | | XM_942086 | 98.5 | 107.6 | 0.0120 | 1.18 | | | |
| TIA1 | TIA1 cytotoxic granule associated RNA binding protein(TIA1) | NM_022173 | 267.7 | 432.7 | 0.0120 | 1.82 | | | |
| CDK5RAP2 | CDK5 regulatory subunit associated protein 2(CDK5RAP2) | NM_001011649 | 122.1 | 150.1 | 0.0120 | 1.39 | NM_001033706 | 2.42 | 10.86 |
| TCOF1 | treacle ribosome biogenesis factor 1(TCOF1) | NM_000356 | 145.9 | 109.5 | 0.0120 | 0.83 | | | |
| RNU6-1 | RNA, U6 small nuclear 1(RNU6-1) | NR_004394 | 1021.3 | 1283.5 | 0.0120 | 1.39 | | | |
| SNX5 | sorting nexin 5(SNX5) | NM_014426 | 165.7 | 204.2 | 0.0120 | 1.41 | | | |
| LRP4 | LDL receptor related protein 4(LRP4) | NM_002334 | 98.8 | 114.9 | 0.0120 | 1.28 | | | |
| SLTM | SAFB like transcription modulator(SLTM) | NM_024755 | 110.5 | 132.4 | 0.0120 | 1.32 | | | |
| SLTM | SAFB like transcription modulator(SLTM) | NM_024755 | 334.1 | 342.5 | 0.0120 | 1.32 | NM_153300 | 0.14 | 61.17 |

| | | | | | | | | | |
|---|---|---|---|---|---|---|---|---|---|
| SLC6A10P | solute carrier family 6 member 10, pseudogene(SLC6A10P) | NR_003083 | 184.2 | 120.9 | 0.0130 | 0.7 | | | |
| DCAF13 | DDB1 and CUL4 associated factor 13(DCAF13) | NM_015420 | 157.5 | 169.6 | 0.0130 | 1.23 | NM_032416 | 108.51 | 1.87 |
| FAM63A | family with sequence similarity 63 member A(FAM63A) | NM_018379 | 119.6 | 110 | 0.0130 | 1.41 | | | |
| FAM63A | family with sequence similarity 63 member A(FAM63A) | NM_018379 | 539 | 672.1 | 0.0130 | 1.41 | | | |
| | | XM_001133026 | 591.5 | 817.8 | 0.0130 | 1.59 | | | |
| | | XM_001134374 | 115.9 | 94.1 | 0.0130 | 0.86 | | | |
| | | AI887987 | 94.6 | 100.2 | 0.0130 | 1.13 | | | |
| rab41 | RAB41, member RAS oncogene family(RAB41) | NM_001032726 | 159.4 | 118.4 | 0.0130 | 0.83 | | | |
| CT45A6 | cancer/testis antigen family 45 member A6(CT45A6) | NM_001017438 | 119.5 | 95.1 | 0.0130 | 0.87 | | | |

| | | | | | | | | | |
|---|---|---|---|---|---|---|---|---|---|
| | | XM_939521 | 103.3 | 105.4 | 0.0130 | 1.11 | | | |
| RNF150 | ring finger protein 150(RNF150) | NM_020724 | 104.9 | 151.1 | 0.0130 | 1.63 | | | |
| KAZN | kazrin, periplakin interacting protein(KAZN) | NM_001017999 | 114.1 | 116.3 | 0.0130 | 1.12 | | | |
| ACKR3 | atypical chemokine receptor 3(ACKR3) | NM_001047841 | 152.2 | 225.8 | 0.0130 | 1.68 | | | |
| BAIAP2 | BAI1 associated protein 2(BAIAP2) | NM_006340 | 123.1 | 97 | 0.0130 | 0.86 | | | |
| BAIAP2 | BAI1 associated protein 2(BAIAP2) | NM_006340 | 183 | 133.2 | 0.0130 | 0.86 | NM_001191088 | 10.07 | 0.03 |
| | | XM_001131851 | 121.6 | 131.7 | 0.0130 | 1.21 | NM_001191088 | 10.07 | 0.03 |
| ZAP70 | zeta chain of T cell receptor associated protein kinase 70(ZAP70) | NM_001079 | 93.7 | 140 | 0.0130 | 0.83 | | | |
| ZAP70 | zeta chain of T cell receptor associated protein kinase 70(ZAP70) | NM_001079 | 157.2 | 117.9 | 0.0130 | 0.83 | | | |
| | | XR_038597 | 334.9 | 402.6 | 0.0130 | 1.39 | | | |

| | | | | | | | | | |
|---|---|---|---|---|---|---|---|---|---|
| GABBR 1 | gamma-aminobutyric acid type B receptor subunit 1(GABBR1) | NM_02190 3 | 130.2 | 105.2 | 0.0130 | 0.88 | NM_0124 96 | 1095.39 | 8.76 |
| | | XM_94251 6 | 110.1 | 91.9 | 0.0130 | 0.89 | | | |
| ZNF787 | zinc finger protein 787(ZNF787) | NM_00100 2836 | 160.4 | 166.5 | 0.0130 | 1.17 | | | |
| | | XR_01635 3 | 118.7 | 91.6 | 0.0130 | 0.83 | | | |
| SRSF4 | serine and arginine rich splicing factor 4(SRSF4) | NM_00562 6 | 813.3 | 1029.5 | 0.0130 | 1.4 | | | |
| CSNK2 A1 | casein kinase 2 alpha 1(CSNK2A1) | NM_00189 5 | 133.1 | 136.4 | 0.0130 | 0.78 | | | |
| CSNK2 A1 | casein kinase 2 alpha 1(CSNK2A1) | NM_00189 5 | 270 | 185.3 | 0.0130 | 0.78 | | | |
| | | XM_00172 3239 | 13523.8 | 15446.4 | 0.0130 | 1.22 | | | |
| | | XR_01753 2 | 190.2 | 214.7 | 0.0130 | 1.28 | | | |
| | | XM_93554 0 | 97.3 | 107 | 0.0130 | 1.18 | | | |

| | | | | | | |
|---|---|---|---|---|---|---|
| KCNH2 | potassium voltage-gated channel subfamily H member 2(KCNH2) | NM_172056 | 108.7 | 111.7 | 0.0130 | 1.14 |
| | | BX107471 | 114.9 | 92.6 | 0.0130 | 0.86 |
| FAM208A | family with sequence similarity 208 member A(FAM208A) | NM_015224 | 337 | 175.6 | 0.0140 | 0.61 |
| LOC729609 | uncharacterized LOC729609(LOC729609) | NR_024440 | 115.3 | 93.9 | 0.0140 | 0.88 |
| RNF5 | ring finger protein 5(RNF5) | NM_006913 | 537.7 | 347.2 | 0.0140 | 0.74 |
| HS6ST2-AS1 | HS6ST2 antisense RNA 1(HS6ST2-AS1) | BQ001311 | 115.6 | 95.2 | 0.0140 | 0.88 |
| THOP1 | thimet oligopeptidase 1(THOP1) | NM_003249 | 121.3 | 144.9 | 0.0140 | 1.34 |
| | | XM_942288 | 119.2 | 139 | 0.0140 | 1.31 |
| | | BQ188349 | 98.3 | 103.9 | 0.0140 | 1.14 |
| OPN5 | opsin 5(OPN5) | NM_001030051 | 174.7 | 120.6 | 0.0140 | 0.77 |
| TLR9 | toll like receptor 9(TLR9) | NM_017442 | 88.6 | 98.7 | 0.0140 | 1.19 |

| | | | | | | |
|---|---|---|---|---|---|---|
| REXO4 | REX4 homolog, 3'-5' exonuclease(REXO4) | NM_020385 | 190.5 | 216.7 | 0.0140 | 1.31 |
| TXNDC11 | thioredoxin domain containing 11(TXNDC11) | NM_015914 | 101.5 | 112.8 | 0.0140 | 1.22 |
| NGFR | nerve growth factor receptor(NGFR) | NM_002507 | 94.8 | 100.5 | 0.0140 | 1.13 |
| TAF1C | TATA-box binding protein associated factor, RNA polymerase I subunit C(TAF1C) | NM_139353 | 143.4 | 156 | 0.0140 | 1.24 |
| | | XR_015650 | 118.6 | 129.5 | 0.0140 | 1.23 |
| MCOLN1 | mucolipin 1(MCOLN1) | NM_020533 | 103.6 | 141.9 | 0.0140 | 1.54 |
| KHDRBS3 | KH RNA binding domain containing, signal transduction associated 3(KHDRBS3) | NM_006558 | 123.7 | 131.3 | 0.0140 | 1.5 |

| | | | | | | |
|---|---|---|---|---|---|---|
| KHDRBS3 | KH RNA binding domain containing, signal transduction associated 3(KHDRBS3) | NM_006558 | 208.7 | 278 | 0.0140 | 1.5 |
| | | AW044615 | 120.4 | 98.3 | 0.0140 | 0.88 |
| HUS1 | HUS1 checkpoint clamp component(HUS1) | NM_004507 | 120.9 | 97.1 | 0.0140 | 0.85 |
| PSMD6-AS2 | PSMD6 antisense RNA 2(PSMD6-AS2) | AK023371 | 107.5 | 109.9 | 0.0140 | 1.11 |
| MRPL39 | mitochondrial ribosomal protein L39(MRPL39) | NM_080794 | 106.8 | 121.7 | 0.0140 | 1.29 |
| KRTAP12-4 | keratin associated protein 12-4(KRTAP12-4) | NM_198698 | 122.6 | 97 | 0.0140 | 0.85 |
| RBM26 | RNA binding motif protein 26(RBM26) | NM_022118 | 144.5 | 170.5 | 0.0140 | 1.34 |
| ATAD2B | ATPase family, AAA domain containing 2B(ATAD2B) | NM_017552 | 110.4 | 114.3 | 0.0140 | 1.16 |
| EXOSC10 | exosome component 10(EXOSC10) | NM_001001998 | 117.1 | 124.9 | 0.0140 | 1.2 |

| | | | | | | | | | |
|---|---|---|---|---|---|---|---|---|---|
| EXOSC10 | exosome component 10(EXOSC10) | NM_001001998 | 418.7 | 407.8 | 0.0140 | 1.2 | NM_013059 | 21.02 | 0.11 |
| SBDSP1 | Shwachman-Bodian-Diamond syndrome pseudogene 1(SBDSP1) | NR_001588 | 279.4 | 179.6 | 0.0150 | 0.72 | | | |
| SBDSP1 | Shwachman-Bodian-Diamond syndrome pseudogene 1(SBDSP1) | NR_001588 | 299 | 214 | 0.0150 | 0.72 | | | |
| | | BQ185080 | 121.5 | 98.8 | 0.0150 | 0.87 | | | |
| SEH1L | SEH1 like nucleoporin(SEH1L) | NM_001013437 | 137.3 | 156.3 | 0.0150 | 1.29 | | | |
| SEH1L | SEH1 like nucleoporin(SEH1L) | NM_001013437 | 221.8 | 217.1 | 0.0150 | 1.29 | | | |
| WDR35 | WD repeat domain 35(WDR35) | NM_001006657 | 257.6 | 166 | 0.0150 | 0.73 | | | |
| CCNI | cyclin I(CCNI) | NM_006835 | 3100.4 | 3698.5 | 0.0150 | 1.27 | | | |
| | | XM_927542 | 155.8 | 116.5 | 0.0150 | 0.83 | | | |
| | | XM_001721042 | 100.4 | 92.1 | 0.0150 | 0.85 | | | |
| | | XM_001721042 | 181.6 | 135.5 | 0.0150 | 0.85 | | | |

| | | | | | | | | | |
|---|---|---|---|---|---|---|---|---|---|
| | | XM_001725241 | 99.2 | 102.1 | 0.0150 | 1.12 | | | |
| MAGEL2 | MAGE family member L2(MAGEL2) | NM_019066 | 176.2 | 124.1 | 0.0150 | 0.79 | NM_012816 | 19.78 | 2.28 |
| | | XR_001434 | 123.5 | 102.6 | 0.0150 | 0.89 | NM_012816 | 19.78 | 2.28 |
| DNAH6 | dynein axonemal heavy chain 6(DNAH6) | NM_173645 | 166 | 185.3 | 0.0150 | 1.27 | | | |
| RAD51C | RAD51 paralog C(RAD51C) | NM_002876 | 103.2 | 122.6 | 0.0150 | 1.32 | | | |
| CSNK1A1 | casein kinase 1 alpha 1(CSNK1A1) | NM_001892 | 195.1 | 256 | 0.0150 | 1.5 | | | |
| SF3B1 | splicing factor 3b subunit 1(SF3B1) | NM_001005526 | 96.4 | 105.3 | 0.0150 | 1.18 | | | |
| CSTF3 | cleavage stimulation factor subunit 3(CSTF3) | NM_001033506 | 112.2 | 124.4 | 0.0150 | 1.25 | | | |
| PLA2G6 | phospholipase A2 group VI(PLA2G6) | NM_001004426 | 182.4 | 129 | 0.0150 | 0.8 | | | |
| ZNF462 | zinc finger protein 462(ZNF462) | NM_021224 | 130.2 | 175.5 | 0.0150 | 1.5 | | | |
| ZCCHC6 | zinc finger CCHC-type containing 6(ZCCHC6) | NM_024617 | 142.7 | 332.1 | 0.0150 | 2.69 | | | |
| | | XM_936462 | 100.1 | 104.5 | 0.0150 | 1.13 | | | |

| | | | | | | | | | |
|---|---|---|---|---|---|---|---|---|---|
| CLCN6 | chloride voltage-gated channel 6(CLCN6) | NM_001286 | 152.7 | 151.3 | 0.0150 | 1.13 | | | |
| ANKRD12 | ankyrin repeat domain 12(ANKRD12) | NM_015208 | 122.6 | 140.8 | 0.0150 | 1.3 | | | |
| RORA | RAR related orphan receptor A(RORA) | AK055969 | 117.3 | 134.3 | 0.0150 | 1.27 | | | |
| PBXIP1 | PBX homeobox interacting protein 1(PBXIP1) | NM_020524 | 161.6 | 122.6 | 0.0150 | 0.85 | | | |
| nfic | nuclear factor I C(NFIC) | NM_005597 | 114.1 | 105.2 | 0.0150 | 0.83 | | | |
| nfic | nuclear factor I C(NFIC) | NM_005597 | 115.2 | 89.5 | 0.0150 | 0.83 | | | |
| DPT | dermatopontin(DPT) | NM_001937 | 670.8 | 373.6 | 0.0150 | 0.64 | | | |
| LOC642947 | uncharacterized LOC642947(LOC642947) | NM_001039895 | 11130.4 | 12980.5 | 0.0150 | 1.28 | | | |
| FRA10AC1 | FRA10A associated CGG repeat 1(FRA10AC1) | NM_203440 | 100.4 | 106 | 0.0150 | 1.14 | | | |
| NXN | nucleoredoxin(NXN) | NM_022463 | 136.1 | 105.1 | 0.0150 | 0.84 | NM_031544 | 4.79 | 38.59 |

| | | | | | | | | | |
|---|---|---|---|---|---|---|---|---|---|
| SREK1 | splicing regulatory glutamic acid and lysine rich protein 1(SREK1) | NM_139168 | 95.7 | 102.7 | 0.0150 | 1.17 | NM_031544 | 4.79 | 38.59 |
| | | XM_001720144 | 97.4 | 102.9 | 0.0150 | 1.14 | | | |
| KTI12 | KTI12 chromatin associated homolog(KTI12) | NM_138417 | 138.8 | 107.1 | 0.0150 | 0.83 | | | |
| | | XM_944214 | 113.1 | 93.7 | 0.0150 | 0.89 | | | |
| WDR19 | WD repeat domain 19(WDR19) | NM_025132 | 146.6 | 167.2 | 0.0150 | 1.28 | | | |
| PAPOLA | poly(A) polymerase alpha(PAPOLA) | NM_032632 | 661.5 | 930.2 | 0.0160 | 1.54 | | | |
| RAB2B | RAB2B, member RAS oncogene family(RAB2B) | NM_032846 | 292.2 | 348.1 | 0.0160 | 1.33 | | | |
| ITGA4 | integrin subunit alpha 4(ITGA4) | NM_000885 | 89.1 | 98.8 | 0.0160 | 1.2 | | | |
| | | XM_928404 | 176.5 | 130.9 | 0.0160 | 0.84 | | | |
| FLJ41327 | FLJ41327 protein(FLJ41327) | NM_207485 | 98.4 | 101.4 | 0.0160 | 1.12 | | | |
| ANKRD16 | ankyrin repeat domain 16(ANKRD16) | NM_0010099941 | 113.3 | 118.9 | 0.0160 | 1.2 | | | |

| | | | | | | |
|---|---|---|---|---|---|---|
| U2SURP | U2 snRNP associated SURP domain containing(U2SURP) | NM_001080415 | 243.1 | 272.7 | 0.0160 | 1.28 |
| PFKFB2 | 6-phosphofructo-2-kinase/fructose-2,6-biphosphatase 2(PFKFB2) | NM_001018053 | 124.3 | 100.1 | 0.0160 | 0.87 |
| PFKFB2 | 6-phosphofructo-2-kinase/fructose-2,6-biphosphatase 2(PFKFB2) | NM_001018053 | 126.5 | 114.8 | 0.0160 | 0.87 |
| | | XM_942545 | 96.5 | 103.1 | 0.0160 | 1.18 |
| DZIP1 | DAZ interacting zinc finger protein 1(DZIP1) | NM_198968 | 124.2 | 171.1 | 0.0160 | 1.54 |
| TLE3 | transducin like enhancer of split 3(TLE3) | NM_005078 | 96.4 | 104.6 | 0.0160 | 1.18 |
| CYP2A13 | cytochrome P450 family 2 subfamily A member 13(CYP2A13) | NM_000766 | 89 | 99.6 | 0.0160 | 1.2 |
| CYP2A13 | cytochrome P450 family 2 subfamily A member 13(CYP2A13) | NM_000766 | 111.3 | 100.3 | 0.0160 | 1.2 |

| | | | | | | |
|---|---|---|---|---|---|---|
| GRIN2A | glutamate ionotropic receptor NMDA type subunit 2A(GRIN2A) | NM_000833 | 111.9 | 93.5 | 0.0160 | 0.88 |
| SETX | senataxin(SETX) | NM_015046 | 104.1 | 123.3 | 0.0160 | 1.28 |
| | | AW975165 | 101.1 | 107.5 | 0.0160 | 1.14 |
| | | XM_0011134195 | 121.6 | 99.8 | 0.0160 | 0.89 |
| | | XM_927996 | 114.5 | 95 | 0.0160 | 0.88 |
| CAPRIN2 | caprin family member 2(CAPRIN2) | NM_001002259 | 235.2 | 252.4 | 0.0160 | 1.23 |
| | | XM_001725231 | 115.2 | 92.7 | 0.0160 | 0.85 |
| EPS8L3 | EPS8 like 3(EPS8L3) | NM_024526 | 112 | 94.3 | 0.0160 | 0.9 |
| LRRC15 | leucine rich repeat containing 15(LRRC15) | NM_130830 | 105.1 | 110.5 | 0.0160 | 1.16 |
| | | XM_001721880 | 107.4 | 101.2 | 0.0160 | 1.15 |
| | | XM_001721880 | 175.1 | 177.7 | 0.0160 | 1.15 |
| SCAF4 | SR-related CTD associated factor 4(SCAF4) | NM_020706 | 183.1 | 219.9 | 0.0160 | 1.34 |
| | | XM_001715244 | 115.4 | 99 | 0.0160 | 0.91 |

| | | | | | | |
|---|---|---|---|---|---|---|
| UBE2Q2 | ubiquitin conjugating enzyme E2 Q2(UBE2Q2) | NM_173469 | 105.9 | 114.7 | 0.0160 | 1.19 |
| | | XM_942049 | 126.3 | 98.4 | 0.0160 | 0.85 |
| NCOR1P1 | nuclear receptor corepressor 1 pseudogene 1(NCOR1P1) | NM_001039379 | 222.7 | 270.1 | 0.0160 | 1.35 |
| BCL2 | BCL2, apoptosis regulator(BCL2) | NM_000633 | 106.3 | 116.7 | 0.0170 | 1.23 |
| BCL2 | BCL2, apoptosis regulator(BCL2) | NM_000633 | 280.2 | 304.5 | 0.0170 | 1.23 |
| GNB3 | G protein subunit beta 3(GNB3) | NM_002075 | 142.1 | 105.4 | 0.0170 | 0.82 |
| ALKBH5 | alkB homolog 5, RNA demethylase(ALKBH5) | NM_017758 | 2465.6 | 1848.7 | 0.0170 | 0.83 |
| CTGF | connective tissue growth factor(CTGF) | NM_001901 | 128 | 92.9 | 0.0170 | 0.79 |
| TSSK1B | testis specific serine kinase 1B(TSSK1B) | NM_032028 | 102 | 107.2 | 0.0170 | 1.15 |
| | | BM671800 | 141.5 | 110.5 | 0.0170 | 0.85 |
| HIC2 | HIC ZBTB transcriptional repressor 2(HIC2) | NM_015094 | 145.2 | 158.9 | 0.0170 | 1.24 |

| | | | | | | |
|---|---|---|---|---|---|---|
| TNRC6A | trinucleotide repeat containing 6A(TNRC6A) | NM_014494 | 124.2 | 128.5 | 0.0170 | 1.15 |
| TNRC6A | trinucleotide repeat containing 6A(TNRC6A) | NM_014494 | 138.8 | 134.2 | 0.0170 | 1.15 |
| TNRC6A | trinucleotide repeat containing 6A(TNRC6A) | NM_014494 | 366.9 | 310.9 | 0.0170 | 1.15 |
| CALCOCO1 | calcium binding and coiled-coil domain 1(CALCOCO1) | NM_020898 | 209.2 | 148.7 | 0.0170 | 0.81 |
| EXOSC9 | exosome component 9(EXOSC9) | NM_001034194 | 170 | 169 | 0.0170 | 1.35 |
| EXOSC9 | exosome component 9(EXOSC9) | NM_001034194 | 170.9 | 205.6 | 0.0170 | 1.35 |
| PTGIR | prostaglandin I2 (prostacyclin) receptor (IP)(PTGIR) | NM_000960 | 131.9 | 101.3 | 0.0170 | 0.84 |
| | | XM_942713 | 95.4 | 101.9 | 0.0170 | 1.17 |
| SNORA12 | small nucleolar RNA, H/ACA box 12(SNORA12) | NR_002954 | 99.6 | 110.3 | 0.0170 | 1.19 |
| TBC1D8 | TBC1 domain family member 8(TBC1D8) | NM_007063 | 129.2 | 139.8 | 0.0170 | 1.23 |

| | | | | | | |
|---|---|---|---|---|---|---|
| NCOA5 | nuclear receptor coactivator 5(NCOA5) | NM_020967 | 117.3 | 122.9 | 0.0170 | 1.17 |
| MCM9 | minichromosome maintenance 9 homologous recombination repair factor(MCM9) | NM_153255 | 108.4 | 127.1 | 0.0170 | 1.27 |
| NDC1 | NDC1 transmembrane nucleoporin(NDC1) | NM_018087 | 96 | 105.5 | 0.0170 | 1.2 |
| CDK7 | cyclin dependent kinase 7(CDK7) | NM_001799 | 180.2 | 223.1 | 0.0170 | 1.4 |
| FA2H | fatty acid 2-hydroxylase(FA2H) | NM_024306 | 180 | 119.9 | 0.0170 | 0.75 |
| MXRA7 | matrix remodeling associated 7(MXRA7) | NM_001008528 | 106.5 | 111.3 | 0.0170 | 1.17 |
| VPS39 | VPS39, HOPS complex subunit(VPS39) | NM_015289 | 111.5 | 112.4 | 0.0170 | 1.12 |
| WNK1 | WNK lysine deficient protein kinase 1(WNK1) | NM_018979 | 154.4 | 142 | 0.0170 | 1.73 |
| WNK1 | WNK lysine deficient protein kinase 1(WNK1) | NM_018979 | 196.9 | 293.3 | 0.0170 | 1.73 |

| | | | | | | |
|---|---|---|---|---|---|---|
| EXOSC3 | exosome component 3(EXOSC3) | NM_016042 | 141.8 | 168.3 | 0.0170 | 1.34 |
| CNRIP1 | cannabinoid receptor interacting protein 1(CNRIP1) | NM_001111101 | 96.3 | 102.1 | 0.0170 | 1.15 |
| SGSH | N-sulfoglucosamine sulfohydrolase(SGSH) | NM_000199 | 364.2 | 245.8 | 0.0170 | 0.78 |
| HAX1 | HCLS1 associated protein X-1(HAX1) | NM_006118 | 120.3 | 122.7 | 0.0180 | 1.16 |
| HAX1 | HCLS1 associated protein X-1(HAX1) | NM_006118 | 310.8 | 288.5 | 0.0180 | 1.16 |
| CD200R1 | CD200 receptor 1(CD200R1) | NM_138806 | 95.7 | 102.4 | 0.0180 | 1.16 |
| COQ10A | coenzyme Q10A(COQ10A) | NM_144576 | 331 | 176.8 | 0.0180 | 0.6 |
| ZMYM5 | zinc finger MYM-type containing 5(ZMYM5) | NM_001039650 | 113.5 | 118.2 | 0.0180 | 1.17 |
| MBP | myelin basic protein(MBP) | NM_001025090 | 189.1 | 252.9 | 0.0180 | 1.48 |
| NAG18 | NAG18 mRNA(NAG18) | NR_002568 | 19695.2 | 16538.4 | 0.0180 | 0.81 |

| | | | | | | |
|---|---|---|---|---|---|---|
| RFC4 | replication factor C subunit 4(RFC4) | NM_181573 | 95.2 | 113.1 | 0.0180 | 1.31 |
| FAM160B2 | family with sequence similarity 160 member B2(FAM160B2) | NM_022749 | 114.1 | 107.6 | 0.0180 | 1.15 |
| FAM160B2 | family with sequence similarity 160 member B2(FAM160B2) | NM_022749 | 124.6 | 128.4 | 0.0180 | 1.15 |
| FAM131A | family with sequence similarity 131 member A(FAM131A) | NM_144635 | 220.5 | 155.3 | 0.0180 | 0.78 |
| MRPL42 | mitochondrial ribosomal protein L42(MRPL42) | NM_172177 | 100.1 | 107 | 0.0180 | 1.16 |
| SLC45A4 | solute carrier family 45 member 4(SLC45A4) | NM_001080431 | 104.9 | 86.9 | 0.0180 | 0.85 |
| CES3 | carboxylesterase 3(CES3) | NM_024922 | 120.8 | 98.3 | 0.0180 | 0.86 |
| LINC00854 | long intergenic non-protein coding RNA 854(LINC00854) | BC073918 | 122.7 | 97.5 | 0.0180 | 0.85 |

| | | | | | | |
|---|---|---|---|---|---|---|
| GPX5 | glutathione peroxidase 5(GPX5) | NM_001509 | 124.2 | 91.8 | 0.0180 | 0.76 |
| GPX5 | glutathione peroxidase 5(GPX5) | NM_001509 | 135.3 | 95 | 0.0180 | 0.76 |
| MCMDC2 | minichromosome maintenance domain containing 2(MCMDC2) | NM_173518 | 109.1 | 103.5 | 0.0180 | 1.28 |
| MCMDC2 | minichromosome maintenance domain containing 2(MCMDC2) | NM_173518 | 3616.2 | 4452.2 | 0.0180 | 1.28 |
| SELENOK | selenoprotein K(SELENOK) | NM_021237 | 119.7 | 97.3 | 0.0180 | 0.88 |
| | | NM_006674 | 130.6 | 133.4 | 0.0180 | 1.17 |
| KCTD12 | potassium channel tetramerization domain containing 12(KCTD12) | NM_138444 | 191.9 | 360.6 | 0.0180 | 2.1 |
| | | XR_038277 | 110.6 | 113.4 | 0.0180 | 1.14 |
| | | XM_0017233218 | 110.5 | 92.1 | 0.0180 | 0.88 |

| | | | | | | | | | |
|---|---|---|---|---|---|---|---|---|---|
| NAA15 | N(alpha)-acetyltransferase 15, NatA auxiliary subunit(NAA15) | NM_057175 | 102.9 | 104.5 | 0.0180 | 1.1 | | | |
| | | XR_015544 | 218.2 | 151.3 | 0.0180 | 0.79 | | | |
| | | XM_001722663 | 148.1 | 113.9 | 0.0180 | 0.86 | | | |
| RND3 | Rho family GTPase 3(RND3) | NM_005168 | 107.3 | 121.3 | 0.0180 | 1.27 | | | |
| AGPAT3 | 1-acylglycerol-3-phosphate O-acyltransferase 3(AGPAT3) | NM_020132 | 153.7 | 118.1 | 0.0180 | 0.86 | | | |
| SNORD33 | small nucleolar RNA, C/D box 33(SNORD33) | NR_000020 | 113.1 | 123 | 0.0180 | 1.18 | | | |
| ANKRD26 | ankyrin repeat domain 26(ANKRD26) | NM_014915 | 131.5 | 146.1 | 0.0180 | 1.22 | | | |
| KCTD17 | potassium channel tetramerization domain containing 17(KCTD17) | NM_024681 | 103.4 | 112 | 0.0180 | 1.19 | NM_031012 | 19.39 | 0.85 |
| PPT1 | palmitoyl-protein thioesterase 1(PPT1) | NM_000310 | 1353.3 | 1657.5 | 0.0180 | 1.32 | | | |

| | | | | | | | | |
|---|---|---|---|---|---|---|---|---|
| RPLP2 | ribosomal protein lateral stalk subunit P2(RPLP2) | NM_001004 | 7146.7 | 9850.2 | 0.0190 | 1.43 | | |
| | | XM_934553 | 112.5 | 93.5 | 0.0190 | 0.9 | NM_012904 | 10.16 | 55.22 |
| | | XM_937753 | 156.6 | 119.7 | 0.0190 | 0.85 | | |
| FUT4 | fucosyltransferase 4(FUT4) | NM_002033 | 89 | 97.5 | 0.0190 | 1.18 | | |
| Snora21 | small nucleolar RNA, H/ACA box 21(SNORA21) | NR_002576 | 106.7 | 117.3 | 0.0190 | 1.23 | | |
| MOCS1 | molybdenum cofactor synthesis 1(MOCS1) | NM_005943 | 221.3 | 150.3 | 0.0190 | 0.76 | | |
| TOP1P2 | topoisomerase (DNA) I pseudogene 2(TOP1P2) | NR_001283 | 153.7 | 184.9 | 0.0190 | 1.36 | | |
| YWHAB | tyrosine 3-monooxygenase/tryptophan 5-monooxygenase activation protein beta(YWHAB) | NM_139323 | 1349.8 | 1627.9 | 0.0190 | 1.31 | | |
| EIF4G3 | eukaryotic translation initiation factor 4 gamma 3(EIF4G3) | NM_003760 | 200.8 | 256.1 | 0.0190 | 1.45 | | |

| | | | | | | | | | |
|---|---|---|---|---|---|---|---|---|---|
| CEACAM4 | carcinoembryonic antigen related cell adhesion molecule 4(CEACAM4) | NM_001817 | 155.2 | 117.1 | 0.0190 | 0.84 | | | |
| | | XM_001133189 | 106.8 | 88.6 | 0.0190 | 0.87 | | | |
| | | XM_001133189 | 234 | 189.4 | 0.0190 | 0.87 | | | |
| KLHL23 | kelch like family member 23(KLHL23) | NM_144711 | 94.7 | 103.4 | 0.0190 | 1.18 | NM_012823 | 17.52 | 0.04 |
| AOC2 | amine oxidase, copper containing 2(AOC2) | NM_001158 | 200.7 | 128.4 | 0.0190 | 0.73 | | | |
| TSPEAR | thrombospondin type laminin G domain and EAR repeats(TSPEAR) | NM_144991 | 122.2 | 100.2 | 0.0190 | 0.88 | | | |
| ADNP | activity dependent neuroprotector homeobox(ADNP) | NM_015339 | 115.9 | 124.2 | 0.0190 | 1.19 | | | |
| | | XM_929248 | 156.7 | 116.6 | 0.0190 | 0.83 | | | |
| URI1 | URI1, prefoldin like chaperone(URI1) | NM_134447 | 154.4 | 158.4 | 0.0190 | 1.45 | | | |

| | | | | | | |
|---|---|---|---|---|---|---|
| URI1 | URI1, prefoldin like chaperone(URI1) | NM_134447 | 476.1 | 600.9 | 0.0190 | 1.45 |
| DAZAP1 | DAZ associated protein 1(DAZAP1) | NM_170711 | 154.8 | 168.1 | 0.0190 | 1.26 |
| ZNF740 | zinc finger protein 740(ZNF740) | NM_001004304 | 165.4 | 126 | 0.0190 | 0.86 |
| CEP131 | centrosomal protein 131(CEP131) | NM_014984 | 100.2 | 112.7 | 0.0190 | 1.25 |
| TMEM9 | transmembrane protein 9(TMEM9) | NM_016456 | 229.1 | 151.5 | 0.0190 | 0.74 |
| ATXN10 | ataxin 10(ATXN10) | NM_013236 | 174.8 | 128.1 | 0.0190 | 0.83 |
| | | XR_016385 | 204.3 | 102.5 | 0.0200 | 0.56 |
| MRPL34 | mitochondrial ribosomal protein L34(MRPL34) | NM_023937 | 324.5 | 240.3 | 0.0200 | 0.84 |
| ZNF22 | zinc finger protein 22(ZNF22) | NM_006963 | 246.9 | 364.2 | 0.0200 | 1.63 |
| ZNF22 | zinc finger protein 22(ZNF22) | NM_006963 | 353 | 311.8 | 0.0200 | 1.63 |
| NSUN5 | NOP2/Sun RNA methyltransferase family member 5(NSUN5) | NM_018044 | 92.7 | 99.7 | 0.0200 | 1.14 |

| | | | | | | |
|---|---|---|---|---|---|---|
| NSUN5 | NOP2/Sun RNA methyltransferase family member 5(NSUN5) | NM_018044 | 136.7 | 130.9 | 0.0200 | 1.14 |
| SLC17A5 | solute carrier family 17 member 5(SLC17A5) | NM_012434 | 100.7 | 103.8 | 0.0200 | 1.12 |
| USP16 | ubiquitin specific peptidase 16(USP16) | NM_006447 | 292.9 | 275.6 | 0.0200 | 1.29 |
| USP16 | ubiquitin specific peptidase 16(USP16) | NM_006447 | 337.4 | 394.9 | 0.0200 | 1.29 |
| | | XM_929287 | 89.9 | 99.1 | 0.0200 | 1.16 |
| DUSP18 | dual specificity phosphatase 18(DUSP18) | NM_152511 | 103.2 | 111 | 0.0200 | 1.18 |
| ZNF302 | zinc finger protein 302(ZNF302) | NM_018443 | 136.8 | 155.1 | 0.0200 | 1.3 |
| KRTAP10-1 | keratin associated protein 10-1(KRTAP10-1) | NM_198691 | 106.4 | 87.3 | 0.0200 | 0.87 |
| FAT1 | FAT atypical cadherin 1(FAT1) | NM_005245 | 259.4 | 183.4 | 0.0200 | 0.64 |
| FAT1 | FAT atypical cadherin 1(FAT1) | NM_005245 | 291.5 | 168.2 | 0.0200 | 0.64 |
| TMOD3 | tropomodulin 3(TMOD3) | NM_014547 | 89.4 | 99 | 0.0200 | 1.18 |

| | | | | | | |
|---|---|---|---|---|---|---|
| TMOD3 | tropomodulin 3(TMOD3) | NM_014547 | 161.7 | 183.8 | 0.0200 | 1.18 |
| SERPINA11 | serpin family A member 11(SERPINA11) | NM_001080451 | 123.9 | 97.9 | 0.0200 | 0.85 |
| DLD | dihydrolipoamide dehydrogenase(DLD) | NM_000108 | 554.4 | 309.5 | 0.0200 | 0.62 |
| | | XR_038983 | 121.7 | 97.3 | 0.0200 | 0.86 |
| ZNF426 | zinc finger protein 426(ZNF426) | NM_024106 | 103.6 | 111.5 | 0.0200 | 1.17 |
| USF1 | upstream transcription factor 1(USF1) | NM_007122 | 114.7 | 163.8 | 0.0200 | 1.6 |
| NALCN | sodium leak channel, non-selective(NALCN) | NM_052867 | 102.1 | 102.5 | 0.0200 | 1.1 |
| FZD1 | frizzled class receptor 1(FZD1) | NM_003505 | 117.6 | 119.8 | 0.0200 | 1.11 |
| FZD1 | frizzled class receptor 1(FZD1) | NM_003505 | 251.1 | 149.7 | 0.0200 | 1.11 |
| SELENOH | selenoprotein H(SELENOH) | NM_170746 | 101.1 | 109.7 | 0.0200 | 1.19 |
| ACKR3 | atypical chemokine receptor 3(ACKR3) | NM_020311 | 289.6 | 535.3 | 0.0200 | 2.12 |

| | | | | | | |
|---|---|---|---|---|---|---|
| DHX15 | DEAH-box helicase 15(DHX15) | NM_001358 | 230 | 241.5 | 0.0200 | 1.56 |
| DHX15 | DEAH-box helicase 15(DHX15) | NM_001358 | 752.7 | 1044.7 | 0.0200 | 1.56 |
| | | XM_940223 | 135 | 90.8 | 0.0200 | 0.75 |
| BAIAP2L2 | BAI1 associated protein 2 like 2(BAIAP2L2) | NM_025045 | 306.1 | 152.5 | 0.0200 | 0.56 |
| CERS4 | ceramide synthase 4(CERS4) | NM_024552 | 124.5 | 100.5 | 0.0200 | 0.85 |
| CASKIN2 | CASK interacting protein 2(CASKIN2) | NM_020753 | 105.8 | 107.8 | 0.0210 | 1.12 |
| Snord96a | small nucleolar RNA, C/D box 96A(SNORD96A) | NR_002592 | 99.3 | 114.7 | 0.0210 | 1.27 |
| CHMP2A | charged multivesicular body protein 2A(CHMP2A) | NM_198426 | 182.8 | 124.1 | 0.0210 | 0.76 |
| | | AV652851 | 115.8 | 150.5 | 0.0210 | 1.42 |
| | | XM_001715037 | 124 | 142.4 | 0.0210 | 1.29 |

| | | | | | | | | | |
|---|---|---|---|---|---|---|---|---|---|
| MMAB | methylmalonic aciduria (cobalamin deficiency) cblB type(MMAB) | NM_052845 | 152.9 | 117.8 | 0.0210 | 0.86 | | | |
| MMAB | methylmalonic aciduria (cobalamin deficiency) cblB type(MMAB) | NM_052845 | 221.6 | 166 | 0.0210 | 0.86 | | | |
| FBRS | fibrosin(FBRS) | NM_001105079 | 188.8 | 217.5 | 0.0210 | 1.33 | | | |
| PTGER4 | prostaglandin E receptor 4(PTGER4) | NM_000958 | 442.3 | 547.7 | 0.0210 | 1.41 | NM_017170 | 13.37 | 0.05 |
| HMGB1P1 | high mobility group box 1 pseudogene 1(HMGB1P1) | NM_001008735 | 215.7 | 378.6 | 0.0210 | 2.01 | | | |
| HMGB1P1 | high mobility group box 1 pseudogene 1(HMGB1P1) | NM_001008735 | 264.9 | 352.8 | 0.0210 | 2.01 | | | |
| MRPS35 | mitochondrial ribosomal protein S35(MRPS35) | NM_021821 | 214.1 | 143.6 | 0.0210 | 0.74 | | | |
| RNVU1-18 | RNA, variant U1 small nuclear 18(RNVU1-18) | NR_004400 | 170 | 208.3 | 0.0210 | 1.38 | | | |

| | | | | | | | | | |
|---|---|---|---|---|---|---|---|---|---|
| ING3 | inhibitor of growth family member 3(ING3) | NM_198267 | 108.3 | 122.6 | 0.0210 | 1.24 | | | |
| RNU1-3 | RNA, U1 small nuclear 3(RNU1-3) | NR_004408 | 157.3 | 196.5 | 0.0210 | 1.4 | | | |
| C9ORF24 | chromosome 9 open reading frame 24(C9orf24) | NM_147169 | 131.7 | 105.7 | 0.0210 | 0.87 | | | |
| KCNH6 | potassium voltage-gated channel subfamily H member 6(KCNH6) | NM_030779 | 12897.2 | 14787 | 0.0210 | 1.29 | NM_001106492 | 2.61 | 10.85 |
| | | XM_942932 | 125.1 | 98.5 | 0.0210 | 0.85 | NM_001106492 | 2.61 | 10.85 |
| | | XM_927702 | 112.2 | 93 | 0.0210 | 0.88 | | | |
| | | NM_001013840 | 122.3 | 123.3 | 0.0210 | 1.14 | | | |
| | | NM_001013840 | 130.5 | 108.1 | 0.0210 | 1.14 | | | |
| FLII | FLII, actin remodeling protein(FLII) | NM_002018 | 165.1 | 179.2 | 0.0210 | 1.24 | | | |
| SYNJ1 | synaptojanin 1(SYNJ1) | NM_203446 | 199.5 | 234.2 | 0.0210 | 1.33 | | | |

| | | | | | | |
|---|---|---|---|---|---|---|
| b4galnt4 | beta-1,4-N-acetyl-galactosaminyltransferase 4(B4GALNT4) | NM_178537 | 89.4 | 99.2 | 0.0210 | 1.19 |
| TRMT10A | tRNA methyltransferase 10A(TRMT10A) | NM_152292 | 95 | 101 | 0.0210 | 1.15 |
| GAPT | GRB2 binding adaptor protein, transmembrane(GAPT) | NM_152687 | 112.4 | 107.4 | 0.0210 | 0.79 |
| GAPT | GRB2 binding adaptor protein, transmembrane(GAPT) | NM_152687 | 154.8 | 109.6 | 0.0210 | 0.79 |
| OR52E5 | olfactory receptor family 52 subfamily E member 5(OR52E5) | NM_001005166 | 106.9 | 85.8 | 0.0210 | 0.84 |
| ACY3 | aminoacylase 3(ACY3) | NM_080658 | 147.2 | 108.7 | 0.0210 | 0.81 |
| SLC35E1 | solute carrier family 35 member E1(SLC35E1) | NM_024881 | 109.2 | 105.9 | 0.0210 | 1.07 |
| SLC35E1 | solute carrier family 35 member E1(SLC35E1) | NM_024881 | 1302.4 | 1529.4 | 0.0210 | 1.07 |
| | | AI349750 | 87 | 96.5 | 0.0210 | 1.18 |

| | | | | | | | | | |
|---|---|---|---|---|---|---|---|---|---|
| CTDSP1 | CTD small phosphatase 1(CTDSP1) | NM_021198 | 113.3 | 90.5 | 0.0210 | 0.83 | | | |
| CTDSP1 | CTD small phosphatase 1(CTDSP1) | NM_021198 | 310.6 | 287.9 | 0.0210 | 0.83 | | | |
| CEP85L | centrosomal protein 85 like(CEP85L) | NM_001042475 | 107.7 | 125 | 0.0210 | 1.29 | | | |
| PYM1 | PYM homolog 1, exon junction complex associated factor(PYM1) | NM_032345 | 107.5 | 121.7 | 0.0210 | 1.26 | | | |
| BLMH | bleomycin hydrolase(BLMH) | NM_000386 | 127.3 | 147.7 | 0.0210 | 1.32 | NM_001270681 | 26.79 | 0.09 |
| SAMD1 | sterile alpha motif domain containing 1(SAMD1) | NM_138352 | 109 | 118.2 | 0.0210 | 1.2 | | | |
| | | DB305811 | 125.1 | 103 | 0.0210 | 0.88 | | | |
| RAD54L2 | RAD54-like 2 (S. cerevisiae)(RAD54L2) | NM_015106 | 116.7 | 128 | 0.0210 | 1.24 | | | |
| | | XM_001717507 | 150.8 | 111.1 | 0.0210 | 0.82 | | | |
| rn7sk | RNA, 7SK small nuclear(RN7SK) | NR_001445 | 279 | 410.6 | 0.0210 | 1.64 | | | |
| KDM2B | lysine demethylase 2B(KDM2B) | NM_001005366 | 279.2 | 159.7 | 0.0210 | 0.66 | | | |

| | | | | | | | | | |
|---|---|---|---|---|---|---|---|---|---|
| ACTR5 | ARP5 actin-related protein 5 homolog(ACTR5) | NM_024855 | 125.2 | 130.7 | 0.0220 | 1.19 | NM_019373 | 263.23 | 1.01 |
| | | AA744250 | 147 | 153.4 | 0.0220 | 1.2 | | | |
| FAM156A | family with sequence similarity 156 member A(FAM156A) | NM_014138 | 278.8 | 299.8 | 0.0220 | 1.23 | | | |
| NKTR | natural killer cell triggering receptor(NKTR) | NM_005385 | 168.2 | 216.3 | 0.0220 | 1.44 | | | |
| MITD1 | microtubule interacting and trafficking domain containing 1(MITD1) | NM_138798 | 146.5 | 173.7 | 0.0220 | 1.34 | | | |
| GRK4 | G protein-coupled receptor kinase 4(GRK4) | NM_001004056 | 119.5 | 164.7 | 0.0220 | 1.54 | | | |
| ANKRD6 | ankyrin repeat domain 6(ANKRD6) | NM_014942 | 102.3 | 104.5 | 0.0220 | 1.12 | | | |
| STX11 | syntaxin 11(STX11) | NM_003764 | 104.9 | 159.5 | 0.0220 | 1.71 | | | |
| CCDC130 | coiled-coil domain containing 130(CCDC130) | NM_030818 | 407.7 | 487.2 | 0.0220 | 1.38 | | | |

| | | | | | | | | | |
|---|---|---|---|---|---|---|---|---|---|
| BRD3 | bromodomain containing 3(BRD3) | NM_007371 | 227.9 | 287.9 | 0.0220 | 1.46 | | | |
| RBMS1 | RNA binding motif single stranded interacting protein 1(RBMS1) | NM_002897 | 134.1 | 162 | 0.0220 | 1.66 | | | |
| RBMS1 | RNA binding motif single stranded interacting protein 1(RBMS1) | NM_002897 | 293.1 | 431.7 | 0.0220 | 1.66 | | | |
| RBMS1 | RNA binding motif single stranded interacting protein 1(RBMS1) | NM_002897 | 337.9 | 419.5 | 0.0220 | 1.66 | NM_012778 | 744.78 | 0.71 |
| ZMYND12 | zinc finger MYND-type containing 12(ZMYND12) | NM_032257 | 198.3 | 130.4 | 0.0220 | 0.73 | NM_012778 | 744.78 | 0.71 |
| | | BX101207 | 106.6 | 112.6 | 0.0220 | 1.18 | | | |
| TTC14 | tetratricopeptide repeat domain 14(TTC14) | NM_133462 | 297 | 398.9 | 0.0220 | 1.52 | | | |
| AKAP13 | A-kinase anchoring protein 13(AKAP13) | NM_006738 | 144.6 | 168.4 | 0.0220 | 1.32 | NM_012909 | 6.73 | 5427.66 |

| | | | | | | | | | |
|---|---|---|---|---|---|---|---|---|---|
| GNAQ | G protein subunit alpha q(GNAQ) | NM_002072 | 128.9 | 101.4 | 0.0220 | 0.86 | NM_022181 | 0.40 | 160.77 |
| | | XM_001724189 | 130.5 | 148.3 | 0.0220 | 1.26 | NM_019157 | 14.48 | 0.05 |
| SOX17 | SRY-box 17(SOX17) | NM_022454 | 98.6 | 119.6 | 0.0220 | 1.34 | | | |
| SNORD16 | small nucleolar RNA, C/D box 16(SNORD16) | NR_002440 | 112.6 | 132.5 | 0.0220 | 1.3 | | | |
| LRRCC1 | leucine rich repeat and coiled-coil centrosomal protein 1(LRRCC1) | NM_033402 | 122.6 | 101.1 | 0.0220 | 1.3 | | | |
| LRRCC1 | leucine rich repeat and coiled-coil centrosomal protein 1(LRRCC1) | NM_033402 | 127.3 | 153.5 | 0.0220 | 1.3 | | | |
| RNU4-1 | RNA, U4 small nuclear 1(RNU4-1) | NR_003925 | 111.3 | 120.7 | 0.0220 | 1.21 | | | |
| slc26a6 | solute carrier family 26 member 6(SLC26A6) | NM_001040454 | 107.9 | 88.9 | 0.0220 | 0.87 | | | |
| tmem187 | transmembrane protein 187(TMEM187) | NM_003492 | 102.1 | 113 | 0.0220 | 1.2 | | | |

| | | | | | | |
|---|---|---|---|---|---|---|
| FEN1 | flap structure-specific endonuclease 1(FEN1) | NM_004111 | 99.3 | 109.8 | 0.0220 | 1.21 |
| FEN1 | flap structure-specific endonuclease 1(FEN1) | NM_004111 | 131.7 | 154.5 | 0.0220 | 1.21 |
| YY1 | YY1 transcription factor(YY1) | NM_003403 | 276.4 | 282.3 | 0.0220 | 1.58 |
| YY1 | YY1 transcription factor(YY1) | NM_003403 | 728.1 | 1066.2 | 0.0220 | 1.58 |
| SNRPB2 | small nuclear ribonucleoprotein polypeptide B2(SNRPB2) | NM_003092 | 216.6 | 153.9 | 0.0230 | 0.81 |
| LIG3 | DNA ligase 3(LIG3) | NM_002311 | 98.5 | 106 | 0.0230 | 1.16 |
| | | XM_937260 | 104.3 | 109 | 0.0230 | 1.15 |
| NPIPB15 | nuclear pore complex interacting protein family member B15(NPIPB15) | NM_001018059 | 118.4 | 98.7 | 0.0230 | 0.89 |

| | | | | | | | | | |
|---|---|---|---|---|---|---|---|---|---|
| YWHAE | tyrosine 3-monooxygenase/tryptophan 5-monooxygenase activation protein epsilon(YWHAE) | NM_006761 | 109 | 115.5 | 0.0230 | 1.18 | | | |
| B3GAT1 | beta-1,3-glucuronyltransferase 1(B3GAT1) | NM_054025 | 171.9 | 123.6 | 0.0230 | 0.82 | | | |
| PIWIL4 | piwi like RNA-mediated gene silencing 4(PIWIL4) | NM_152431 | 87.6 | 97.7 | 0.0230 | 1.2 | | | |
| STX18 | syntaxin 18(STX18) | NM_016930 | 113.5 | 96.4 | 0.0230 | 0.9 | | | |
| | | BF509118 | 115.1 | 123.4 | 0.0230 | 1.2 | | | |
| C3orf62 | chromosome 3 open reading frame 62(C3orf62) | NM_198562 | 105.3 | 110.3 | 0.0230 | 1.15 | | | |
| NDUFAF7 | NADH:ubiquinone oxidoreductase complex assembly factor 7(NDUFAF7) | NM_144736 | 122.7 | 114.2 | 0.0230 | 1.36 | | | |
| NDUFAF7 | NADH:ubiquinone oxidoreductase complex assembly factor 7(NDUFAF7) | NM_144736 | 296.5 | 349.9 | 0.0230 | 1.36 | NM_019168 | 8.26 | 61.49 |

| | | | | | | |
|---|---|---|---|---|---|---|
| ZNF263 | zinc finger protein 263(ZNF263) | NM_005741 | 160.8 | 181.7 | 0.0230 | 1.29 |
| | | XR_040873 | 110.8 | 93.1 | 0.0230 | 0.88 |
| RNF115 | ring finger protein 115(RNF115) | NM_014455 | 126.8 | 145.2 | 0.0230 | 1.55 |
| RNF115 | ring finger protein 115(RNF115) | NM_014455 | 462.1 | 652.9 | 0.0230 | 1.55 |
| RUFY3 | RUN and FYVE domain containing 3(RUFY3) | NM_001037442 | 104 | 106.6 | 0.0230 | 1.13 |
| RUFY3 | RUN and FYVE domain containing 3(RUFY3) | NM_001037442 | 349.3 | 389.7 | 0.0230 | 1.13 |
| PICALM | phosphatidylinositol binding clathrin assembly protein(PICALM) | NM_007166 | 121.6 | 97.9 | 0.0230 | 0.84 |
| PICALM | phosphatidylinositol binding clathrin assembly protein(PICALM) | NM_007166 | 866.6 | 648.3 | 0.0230 | 0.84 |
| | | BX111977 | 120.5 | 101.4 | 0.0240 | 0.91 |
| MBTPS1 | membrane bound transcription factor peptidase, site 1(MBTPS1) | NM_003791 | 166.4 | 156.6 | 0.0240 | 1.21 |

| | | | | | | |
|---|---|---|---|---|---|---|
| MBTPS1 | membrane bound transcription factor peptidase, site 1(MBTPS1) | NM_003791 | 182.9 | 195.8 | 0.0240 | 1.21 |
| C11orf57 | chromosome 11 open reading frame 57(C11orf57) | NM_018195 | 191.7 | 142.5 | 0.0240 | 0.84 |
| ARF1 | ADP ribosylation factor 1(ARF1) | NM_001024227 | 1348.5 | 902.1 | 0.0240 | 0.76 |
| | | XM_939026 | 127.1 | 102.8 | 0.0240 | 0.88 |
| VPS37D | VPS37D, ESCRT-I subunit(VPS37D) | NM_001077621 | 567.1 | 324.4 | 0.0240 | 0.65 |
| GPATCH4 | G-patch domain containing 4(GPATCH4) | NM_182679 | 97.6 | 98 | 0.0240 | 0.89 |
| GPATCH4 | G-patch domain containing 4(GPATCH4) | NM_182679 | 117.7 | 97.8 | 0.0240 | 0.89 |
| ZNF366 | zinc finger protein 366(ZNF366) | NM_152625 | 114.4 | 122.4 | 0.0240 | 1.19 |
| SPDYE8P | speedy/RINGO cell cycle regulator family member E8, pseudogene(SPDYE8P) | NR_003664 | 157.4 | 106.3 | 0.0240 | 0.75 |

| | | | | | | |
|---|---|---|---|---|---|---|
| SPDYE8P | speedy/RINGO cell cycle regulator family member E8, pseudogene(SPDYE8P) | NR_003664 | 11806.9 | 13009 | 0.0240 | 0.75 |
| SCP2 | sterol carrier protein 2(SCP2) | NM_001007098 | 286 | 178.5 | 0.0240 | 0.73 |
| nthl1 | nth like DNA glycosylase 1(NTHL1) | NM_002528 | 119.6 | 131.8 | 0.0240 | 1.24 |
| ZBTB8A | zinc finger and BTB domain containing 8A(ZBTB8A) | NM_001040441 | 102.8 | 114.1 | 0.0240 | 1.21 |
| MATR3 | matrin 3(MATR3) | NM_199189 | 97.7 | 91.5 | 0.0240 | 1.15 |
| MATR3 | matrin 3(MATR3) | NM_199189 | 100.8 | 107.2 | 0.0240 | 1.15 |
| MATR3 | matrin 3(MATR3) | NM_199189 | 1659.9 | 2099.5 | 0.0240 | 1.15 |
| BRPF1 | bromodomain and PHD finger containing 1(BRPF1) | NM_004634 | 432.9 | 664.6 | 0.0240 | 1.72 |
| TLDC1 | TBC/LysM-associated domain containing 1(TLDC1) | NM_020947 | 99.8 | 102.9 | 0.0240 | 1.12 |

| | | | | | | |
|---|---|---|---|---|---|---|
| SMARCC2 | SWI/SNF related, matrix associated, actin dependent regulator of chromatin subfamily c member 2(SMARCC2) | NM_139067 | 248.7 | 301.3 | 0.0240 | 1.35 |
| | | XR_018904 | 123.1 | 133.3 | 0.0240 | 1.22 |
| ZNF676 | zinc finger protein 676(ZNF676) | NM_001001411 | 102.6 | 106.4 | 0.0240 | 1.14 |
| | | XM_0017244301 | 101.8 | 105.2 | 0.0240 | 1.13 |
| GLMN | glomulin, FKBP associated protein(GLMN) | NM_053274 | 222.4 | 274.3 | 0.0240 | 1.4 |
| | | XM_0017244735 | 153.2 | 167.9 | 0.0240 | 1.25 |
| DDX19B | DEAD-box helicase 19B(DDX19B) | NM_0010144449 | 124 | 132.7 | 0.0240 | 1.19 |
| | | XM_927306 | 111.3 | 93.2 | 0.0240 | 0.86 |
| PRR19 | proline rich 19(PRR19) | NM_199285 | 131.5 | 107.1 | 0.0240 | 0.88 |

| | | | | | | |
|---|---|---|---|---|---|---|
| CMTM3 | CKLF like MARVEL transmembrane domain containing 3(CMTM3) | NM_181553 | 186.3 | 132 | 0.0240 | 0.81 |
| UBE2O | ubiquitin conjugating enzyme E2 O(UBE2O) | NM_022066 | 133.9 | 143.6 | 0.0240 | 1.19 |
| | | XM_939186 | 159 | 190.9 | 0.0240 | 1.38 |
| ANAPC11 | anaphase promoting complex subunit 11(ANAPC11) | NM_001002246 | 106.2 | 112.6 | 0.0240 | 1.17 |
| ANAPC11 | anaphase promoting complex subunit 11(ANAPC11) | NM_001002246 | 414 | 382.8 | 0.0240 | 1.17 |
| AGXT | alanine-glyoxylate aminotransferase(AGXT) | NM_000030 | 1763.1 | 657.1 | 0.0240 | 0.42 |
| PHF20L1 | PHD finger protein 20-like 1(PHF20L1) | NM_198513 | 97.4 | 107.4 | 0.0240 | 1.19 |
| NSA2 | NSA2, ribosome biogenesis homolog(NSA2) | NM_014886 | 1103.8 | 1275 | 0.0240 | 1.26 |

| | | | | | | | | | |
|---|---|---|---|---|---|---|---|---|---|
| NSA2 | NSA2, ribosome biogenesis homolog(NSA2) | NM_014886 | 1434.7 | 1488 | 0.0240 | 1.26 | | | |
| RAP2B | RAP2B, member of RAS oncogene family(RAP2B) | NM_002886 | 227.5 | 151 | 0.0250 | 0.75 | | | |
| or1l4 | olfactory receptor family 1 subfamily L member 4(OR1L4) | NM_001005235 | 103.7 | 95.1 | 0.0250 | 0.88 | | | |
| or1l4 | olfactory receptor family 1 subfamily L member 4(OR1L4) | NM_001005235 | 144.4 | 115.3 | 0.0250 | 0.88 | NM_001106061 | 2.22 | 12.22 |
| NOC3L | NOC3 like DNA replication regulator(NOC3L) | NM_022451 | 160.2 | 214.1 | 0.0250 | 1.53 | | | |
| HEATR6 | HEAT repeat containing 6(HEATR6) | NM_022070 | 154.9 | 206.3 | 0.0250 | 1.47 | | | |
| EP400 | E1A binding protein p400(EP400) | NM_015409 | 131.6 | 145.5 | 0.0250 | 1.24 | | | |
| KIAA0430 | KIAA0430(KIAA0430) | NM_014647 | 369.1 | 401.5 | 0.0250 | 1.25 | | | |
| SNORA14B | small nucleolar RNA, H/ACA box 14B(SNORA14B) | NR_002956 | 106.5 | 108.6 | 0.0250 | 1.11 | | | |

| | | | | | | |
|---|---|---|---|---|---|---|
| NUB1 | negative regulator of ubiquitin like proteins 1(NUB1) | NM_016118 | 128.1 | 103 | 0.0250 | 0.88 |
| NUB1 | negative regulator of ubiquitin like proteins 1(NUB1) | NM_016118 | 364.7 | 343.6 | 0.0250 | 0.88 |
| | | XM_926147 | 99 | 105.4 | 0.0250 | 1.15 |
| ATOH8 | atonal bHLH transcription factor 8(ATOH8) | NM_032827 | 99.4 | 117.3 | 0.0250 | 1.32 |
| mthfd2 | methylenetetrahydrofolate dehydrogenase (NADP+ dependent) 2, methenyltetrahydrofolate cyclohydrolase(MTHFD2) | NM_006636 | 89 | 98 | 0.0250 | 1.19 |
| CEPT1 | choline/ethanolamine phosphotransferase 1(CEPT1) | NM_006090 | 332.9 | 524.5 | 0.0250 | 1.81 |
| TRIM9 | tripartite motif containing 9(TRIM9) | NM_015163 | 122.3 | 98.1 | 0.0250 | 0.87 |
| | | XR_039516 | 112.6 | 92.6 | 0.0250 | 0.88 |

| | | | | | | |
|---|---|---|---|---|---|---|
| | | XR_039516 | 184.9 | 188.3 | 0.0250 | 0.88 |
| USP19 | ubiquitin specific peptidase 19(USP19) | NM_006677 | 107.6 | 113.3 | 0.0250 | 1.16 |
| PLAGL1 | PLAG1 like zinc finger 1(PLAGL1) | NM_001080952 | 113.4 | 119.2 | 0.0250 | 1.18 |
| SFSWAP | splicing factor SWAP homolog(SFSWAP) | NM_004592 | 143.9 | 155.8 | 0.0250 | 1.22 |
| NUDT6 | nudix hydrolase 6(NUDT6) | NM_007083 | 115.7 | 102.6 | 0.0250 | 0.79 |
| NUDT6 | nudix hydrolase 6(NUDT6) | NM_007083 | 132.2 | 110.8 | 0.0250 | 0.79 |
| NUDT6 | nudix hydrolase 6(NUDT6) | NM_007083 | 172 | 122.8 | 0.0250 | 0.79 |
| GJC2 | gap junction protein gamma 2(GJC2) | NM_020435 | 189.5 | 133.9 | 0.0250 | 0.79 |
| | | XR_017492 | 128.7 | 162.5 | 0.0250 | 1.46 |
| | | XR_017492 | 153.4 | 141.9 | 0.0250 | 1.46 |
| | | XR_017492 | 1141.3 | 1826.9 | 0.0250 | 1.46 |
| UBAP2L | ubiquitin associated protein 2 like(UBAP2L) | NM_014847 | 224 | 284.6 | 0.0250 | 1.42 |

| | | | | | | |
|---|---|---|---|---|---|---|
| TSC1 | tuberous sclerosis 1(TSC1) | NM_000368 | 127.4 | 116.9 | 0.0250 | 1.19 |
| TSC1 | tuberous sclerosis 1(TSC1) | NM_000368 | 177.3 | 183.4 | 0.0250 | 1.19 |
| TSC1 | tuberous sclerosis 1(TSC1) | NM_000368 | 444.8 | 471.5 | 0.0250 | 1.19 |
| AP2A2 | adaptor related protein complex 2 alpha 2 subunit(AP2A2) | NM_012305 | 128.7 | 100.4 | 0.0250 | 0.85 |
| PIH1D1 | PIH1 domain containing 1(PIH1D1) | NM_017916 | 186.1 | 209.3 | 0.0260 | 1.27 |
| FAM32A | family with sequence similarity 32 member A(FAM32A) | NM_014077 | 132.7 | 153.7 | 0.0260 | 1.31 |
| SNORA70C | small nucleolar RNA, H/ACA box 70C(SNORA70C) | NR_003708 | 127 | 129.6 | 0.0260 | 1.13 |
| | | XR_037303 | 112.5 | 92.1 | 0.0260 | 0.86 |
| P2RY6 | pyrimidinergic receptor P2Y6(P2RY6) | NM_176796 | 93.7 | 109.9 | 0.0260 | 1.29 |
| HPN | hepsin(HPN) | NM_002151 | 329.3 | 231.5 | 0.0260 | 0.53 |
| HPN | hepsin(HPN) | NM_002151 | 739.4 | 351.3 | 0.0260 | 0.53 |

| | | | | | | |
|---|---|---|---|---|---|---|
| CASP2 | caspase 2(CASP2) | NM_032983 | 106.4 | 120.7 | 0.0260 | 1.26 |
| PPP1R12B | protein phosphatase 1 regulatory subunit 12B(PPP1R12B) | NM_032103 | 94 | 98.1 | 0.0260 | 1.12 |
| PPP1R12B | protein phosphatase 1 regulatory subunit 12B(PPP1R12B) | NM_032103 | 101.4 | 103.6 | 0.0260 | 1.12 |
| RAB20 | RAB20, member RAS oncogene family(RAB20) | NM_017817 | 102.9 | 105.4 | 0.0260 | 1.13 |
| UQCR11 | ubiquinol-cytochrome c reductase, complex III subunit XI(UQCR11) | NM_006830 | 126.6 | 102.6 | 0.0260 | 0.88 |
| pik3r6 | phosphoinositide-3-kinase regulatory subunit 6(PIK3R6) | NM_001010855 | 148.7 | 110.2 | 0.0260 | 0.83 |
| RAD52 | RAD52 homolog, DNA repair protein(RAD52) | NM_134424 | 108.8 | 92.2 | 0.0260 | 0.88 |

| | | | | | | | | | |
|---|---|---|---|---|---|---|---|---|---|
| SC5D | sterol-C5-desaturase(SC5D) | NM_006918 | 532.6 | 324.3 | 0.0260 | 0.69 | | | |
| | | XM_938671 | 117.4 | 122.6 | 0.0260 | 1.16 | | | |
| | | XM_936333 | 109.8 | 113.2 | 0.0260 | 1.14 | | | |
| RAE1 | ribonucleic acid export 1(RAE1) | NM_003610 | 128.1 | 100.3 | 0.0260 | 0.86 | | | |
| RAE1 | ribonucleic acid export 1(RAE1) | NM_003610 | 281.3 | 269.9 | 0.0260 | 0.86 | | | |
| | | XM_001720371 | 96.9 | 103.9 | 0.0260 | 1.17 | | | |
| | | XM_001715936 | 103.4 | 109.1 | 0.0260 | 1.15 | | | |
| STT3B | STT3B, catalytic subunit of the oligosaccharyltransferase complex(STT3B) | NM_178862 | 163.6 | 188.3 | 0.0260 | 1.28 | | | |
| | | XM_938093 | 97.3 | 110 | 0.0260 | 1.23 | | | |
| | | XM_938093 | 150.6 | 186 | 0.0260 | 1.23 | | | |
| | | XM_933275 | 119.4 | 98.7 | 0.0260 | 0.9 | | | |
| SMYD5 | SMYD family member 5(SMYD5) | NM_006062 | 194.1 | 143.4 | 0.0260 | 0.84 | | | |
| | | XM_931329 | 116.3 | 95.5 | 0.0260 | 0.87 | NM_001014273 | 3.32 | 15.01 |

| | | | | | | | | | |
|---|---|---|---|---|---|---|---|---|---|
| CCNY | cyclin Y(CCNY) | NM_145012 | 202.3 | 272.5 | 0.0260 | 1.45 | NM_001014273 | 3.32 | 15.01 |
| NICN1 | nicolin 1(NICN1) | NM_032316 | 103.3 | 107 | 0.0260 | 1.13 | NM_001014273 | 3.32 | 15.01 |
| LCOR | ligand dependent nuclear receptor corepressor(LCOR) | NM_032440 | 125.2 | 111.6 | 0.0260 | 1.28 | | | |
| LCOR | ligand dependent nuclear receptor corepressor(LCOR) | NM_032440 | 139.9 | 164.1 | 0.0260 | 1.28 | | | |
| aire | autoimmune regulator(AIRE) | NM_000658 | 101.8 | 106.7 | 0.0260 | 1.16 | | | |
| SLC13A2 | solute carrier family 13 member 2(SLC13A2) | NM_003984 | 281.3 | 166.8 | 0.0260 | 0.68 | | | |
| SCNM1 | sodium channel modifier 1(SCNM1) | NM_024041 | 113 | 110.4 | 0.0260 | 1.18 | | | |
| SCNM1 | sodium channel modifier 1(SCNM1) | NM_024041 | 138.3 | 145.4 | 0.0260 | 1.18 | | | |
| ZNF786 | zinc finger protein 786(ZNF786) | NM_152411 | 966.8 | 1160.6 | 0.0270 | 1.39 | | | |
| NFASC | neurofascin(NFASC) | NM_001005387 | 114.6 | 98.2 | 0.0270 | 0.92 | | | |
| | | XM_001132115 | 157.4 | 100.9 | 0.0270 | 0.71 | | | |

| | | | | | | |
|---|---|---|---|---|---|---|
| TARBP1 | TAR (HIV-1) RNA binding protein 1(TARBP1) | NM_005646 | 113.6 | 122.8 | 0.0270 | 1.22 |
| | | XR_017629 | 115.4 | 126.5 | 0.0270 | 1.23 |
| CPNE6 | copine 6(CPNE6) | NM_006032 | 260.4 | 119.5 | 0.0270 | 0.53 |
| MIR488 | microRNA 488(MIR488) | NR_030163 | 117.2 | 99.1 | 0.0270 | 0.91 |
| METTL15 | methyltransferase like 15(METTL15) | NM_152636 | 107.7 | 111.2 | 0.0270 | 1.13 |
| DDX52 | DExD-box helicase 52(DDX52) | NM_007010 | 294.2 | 333.2 | 0.0270 | 1.27 |
| TANGO2 | transport and golgi organization 2 homolog(TANGO2) | NM_152906 | 113.2 | 138.2 | 0.0270 | 1.37 |
| | | XM_001725354 | 119.8 | 95.8 | 0.0270 | 0.87 |
| MNT | MAX network transcriptional repressor(MNT) | NM_020310 | 152 | 167.2 | 0.0270 | 1.23 |
| ZNF274 | zinc finger protein 274(ZNF274) | NM_016324 | 126.2 | 99.8 | 0.0270 | 0.86 |
| GON4L | gon-4 like(GON4L) | NM_001037533 | 277.2 | 349.3 | 0.0270 | 1.49 |
| | | XR_038286 | 110.5 | 87.6 | 0.0270 | 0.83 |

| | | | | | | |
|---|---|---|---|---|---|---|
| CHCHD1 | coiled-coil-helix-coiled-coil-helix domain containing 1(CHCHD1) | NM_203298 | 111.9 | 115.8 | 0.0270 | 1.15 |
| TUBB3 | tubulin beta 3 class III(TUBB3) | NM_006086 | 144.5 | 111.1 | 0.0270 | 0.85 |
| RNASEH1 | ribonuclease H1(RNASEH1) | NM_002936 | 114.7 | 123.8 | 0.0270 | 1.22 |
| | | NM_183041 | 112.9 | 94.4 | 0.0270 | 0.89 |
| | | XM_498608 | 115.6 | 95.9 | 0.0270 | 0.9 |
| TSPAN31 | tetraspanin 31(TSPAN31) | NM_005981 | 261.6 | 280.4 | 0.0270 | 1.22 |
| TRAF7 | TNF receptor associated factor 7(TRAF7) | NM_032271 | 132.5 | 127.3 | 0.0270 | 0.85 |
| TRAF7 | TNF receptor associated factor 7(TRAF7) | NM_032271 | 141.6 | 109 | 0.0270 | 0.85 |
| | | XR_016218 | 303.5 | 114.4 | 0.0280 | 0.43 |
| | | XM_944104 | 172.2 | 99.2 | 0.0280 | 0.65 |
| | | XM_944104 | 2909.6 | 2692.8 | 0.0280 | 0.65 |
| ODF3L2 | outer dense fiber of sperm tails 3 like 2(ODF3L2) | NM_182577 | 137.8 | 111.1 | 0.0280 | 0.89 |

| | | | | | | |
|---|---|---|---|---|---|---|
| ACYP2 | acylphosphatase 2(ACYP2) | NM_138448 | 183 | 134.3 | 0.0280 | 0.83 |
| ACYP2 | acylphosphatase 2(ACYP2) | NM_138448 | 255.9 | 167.5 | 0.0280 | 0.83 |
| ZNF304 | zinc finger protein 304(ZNF304) | NM_020657 | 105.8 | 107.2 | 0.0280 | 1.12 |
| SERF1B | small EDRK-rich factor 1B(SERF1B) | NM_022978 | 111.8 | 88 | 0.0280 | 0.84 |
| ARID4A | AT-rich interaction domain 4A(ARID4A) | NM_023001 | 101.7 | 106.8 | 0.0280 | 1.15 |
| plxdc2 | plexin domain containing 2(PLXDC2) | NM_032812 | 118.9 | 137.3 | 0.0280 | 1.28 |
| RALGDS | ral guanine nucleotide dissociation stimulator(RALGDS) | NM_006266 | 252.8 | 476.7 | 0.0280 | 2.11 |
| LILRB5 | leukocyte immunoglobulin like receptor B5(LILRB5) | NM_006840 | 108.6 | 111.5 | 0.0280 | 1.1 |
| | | XM_932143 | 190.8 | 215.8 | 0.0280 | 1.27 |
| | | XR_015609 | 111.5 | 95.8 | 0.0280 | 0.92 |
| | | XM_001718130 | 127.1 | 100.9 | 0.0280 | 0.87 |

| | | | | | | |
|---|---|---|---|---|---|---|
| f8a1 | coagulation factor VIII-associated 1(F8A1) | NM_012151 | 233.4 | 159.3 | 0.0280 | 0.78 |
| RARRES1 | retinoic acid receptor responder 1(RARRES1) | NM_206963 | 92.6 | 104.2 | 0.0280 | 1.23 |
| HOXC13 | homeobox C13(HOXC13) | NM_017410 | 116.4 | 92.7 | 0.0280 | 0.84 |
| MOSPD2 | motile sperm domain containing 2(MOSPD2) | NM_152581 | 122.9 | 101.6 | 0.0280 | 1.25 |
| MOSPD2 | motile sperm domain containing 2(MOSPD2) | NM_152581 | 151.1 | 169.8 | 0.0280 | 1.25 |
| UBE2Z | ubiquitin conjugating enzyme E2 Z(UBE2Z) | NM_023079 | 116.7 | 129 | 0.0280 | 1.24 |
| AJAP1 | adherens junctions associated protein 1(AJAP1) | NM_001042478 | 103.6 | 117.8 | 0.0280 | 1.26 |
| | | XR_037881 | 112.8 | 93.1 | 0.0280 | 0.86 |
| | | XR_037881 | 835.7 | 882.2 | 0.0280 | 0.86 |

| | | | | | | |
|---|---|---|---|---|---|---|
| CDKN2AIPNL | CDKN2A interacting protein N-terminal like(CDKN2AIPNL) | NM_080656 | 9284.9 | 11373 | 0.0280 | 1.34 |
| | | XM_927976 | 84.7 | 96.2 | 0.0280 | 1.2 |
| SLC35D1 | solute carrier family 35 member D1(SLC35D1) | NM_015139 | 113 | 96.2 | 0.0280 | 0.91 |
| ALS2CL | ALS2 C-terminal like(ALS2CL) | NM_182775 | 122.6 | 124.5 | 0.0290 | 1.14 |
| CRCP | CGRP receptor component(CRCP) | NM_001040648 | 119.4 | 122.8 | 0.0290 | 1.16 |
| SLCO1A2 | solute carrier organic anion transporter family member 1A2(SLCO1A2) | NM_134431 | 325.3 | 184 | 0.0290 | 0.66 |
| FLT4 | fms related tyrosine kinase 4(FLT4) | NM_002020 | 124.4 | 100.7 | 0.0290 | 0.89 |
| | | XM_944907 | 341.4 | 451.1 | 0.0290 | 1.53 |
| SPIN4 | spindlin family member 4(SPIN4) | NM_001012968 | 105.1 | 111 | 0.0290 | 1.17 |

| | | | | | | |
|---|---|---|---|---|---|---|
| Snord89 | small nucleolar RNA, C/D box 89(SNORD89) | NR_003070 | 381.2 | 655.7 | 0.0290 | 1.88 |
| | | AL110176 | 107.3 | 108.5 | 0.0290 | 1.11 |
| | | XM_001715878 | 125.7 | 101.3 | 0.0290 | 0.88 |
| H1FOO | H1 histone family member O, oocyte specific(H1FOO) | NM_153833 | 111.7 | 95.5 | 0.0290 | 0.9 |
| PSMC3IP | PSMC3 interacting protein(PSMC3IP) | NM_013290 | 118.1 | 96.7 | 0.0290 | 0.87 |
| ARSB | arylsulfatase B(ARSB) | NM_000046 | 124.2 | 102.4 | 0.0290 | 0.9 |
| ARSB | arylsulfatase B(ARSB) | NM_000046 | 149.8 | 126.1 | 0.0290 | 0.9 |
| ARSG | arylsulfatase G(ARSG) | NM_014960 | 195.9 | 145.7 | 0.0290 | 0.84 |
| SNAPIN | SNAP associated protein(SNAPIN) | NM_012437 | 112.2 | 118.8 | 0.0290 | 1.16 |
| HSPA9 | heat shock protein family A (Hsp70) member 9(HSPA9) | NM_004134 | 758.7 | 422.6 | 0.0290 | 0.63 |
| RNVU1-7 | RNA, variant U1 small nuclear 7(RNVU1-7) | NR_004426 | 156.7 | 187.9 | 0.0290 | 1.35 |

| | | | | | | |
|---|---|---|---|---|---|---|
| LCAT | lecithin-cholesterol acyltransferase(LCAT) | NM_000229 | 250 | 312.9 | 0.0290 | 1.44 |
| UBXN7 | UBX domain protein 7(UBXN7) | NM_015562 | 91.5 | 100.5 | 0.0290 | 1.19 |
| DNAAF5 | dynein axonemal assembly factor 5(DNAAF5) | NM_017802 | 88.2 | 107.2 | 0.0290 | 1.33 |
| FAM63B | family with sequence similarity 63 member B(FAM63B) | NM_001040450 | 115.2 | 139.8 | 0.0290 | 1.34 |
| | | XM_930835 | 202.8 | 145.5 | 0.0290 | 0.82 |
| NR6A1 | nuclear receptor subfamily 6 group A member 1(NR6A1) | NM_001489 | 109.4 | 93.9 | 0.0290 | 0.9 |
| SNX22 | sorting nexin 22(SNX22) | NM_024798 | 133.3 | 101.7 | 0.0290 | 0.82 |
| STRN3 | striatin 3(STRN3) | NM_014574 | 311.9 | 385.9 | 0.0290 | 1.39 |
| | | XM_925901 | 99.2 | 102.1 | 0.0290 | 1.11 |
| ENY2 | ENY2, transcription and export complex 2 subunit(ENY2) | NM_020189 | 972.6 | 641.6 | 0.0290 | 0.73 |

| | | | | | | | | | |
|---|---|---|---|---|---|---|---|---|---|
| OR2B2 | olfactory receptor family 2 subfamily B member 2(OR2B2) | NM_033057 | 129.6 | 101.4 | 0.0290 | 0.82 | | | |
| CACTIN | cactin, spliceosome C complex subunit(CACTIN) | NM_001080543 | 114.2 | 113.2 | 0.0300 | 1.1 | | | |
| | | XR_017883 | 222.1 | 139.3 | 0.0300 | 0.72 | | | |
| MED1 | mediator complex subunit 1(MED1) | NM_004774 | 146.2 | 158.8 | 0.0300 | 1.22 | | | |
| MED1 | mediator complex subunit 1(MED1) | NM_004774 | 251.7 | 199.4 | 0.0300 | 1.22 | | | |
| | | XM_933621 | 127.6 | 103.3 | 0.0300 | 0.88 | | | |
| SPC25 | SPC25, NDC80 kinetochore complex component(SPC25) | NM_020675 | 90.4 | 97.5 | 0.0300 | 1.16 | | | |
| HOXA6 | homeobox A6(HOXA6) | NM_024014 | 472.9 | 282.8 | 0.0300 | 0.7 | NM_145089 | 67.02 | 12.33 |
| ACBD3 | acyl-CoA binding domain containing 3(ACBD3) | NM_022735 | 286.2 | 184.8 | 0.0300 | 0.72 | NM_013157 | 76.74 | 0.47 |
| MT2A | metallothionein 2A(MT2A) | NM_005953 | 1687.2 | 603.4 | 0.0300 | 0.42 | NM_013157 | 76.74 | 0.47 |

| | | | | | | | | | |
|---|---|---|---|---|---|---|---|---|---|
| BVES | blood vessel epicardial substance(BVES) | NM_147147 | 124.8 | 105 | 0.0300 | 0.92 | NM_013157 | 76.74 | 0.47 |
| FANCG | Fanconi anemia complementation group G(FANCG) | NM_004629 | 133.6 | 164.5 | 0.0300 | 1.38 | | | |
| KRTAP12-3 | keratin associated protein 12-3(KRTAP12-3) | NM_198697 | 104.4 | 106.2 | 0.0300 | 1.11 | | | |
| ZFPM2 | zinc finger protein, FOG family member 2(ZFPM2) | NM_012082 | 94.3 | 105.6 | 0.0300 | 1.22 | | | |
| PTPRB | protein tyrosine phosphatase, receptor type B(PTPRB) | NM_002837 | 151.1 | 197.8 | 0.0300 | 1.48 | | | |
| UCKL1 | uridine-cytidine kinase 1 like 1(UCKL1) | NM_017859 | 413.3 | 491.9 | 0.0300 | 1.36 | | | |
| WHAMM | WAS protein homolog associated with actin, golgi membranes and microtubules(WHAMM) | NM_001080435 | 137.2 | 145.8 | 0.0300 | 1.2 | | | |

| | | | | | | |
|---|---|---|---|---|---|---|
| FAM110B | family with sequence similarity 110 member B(FAM110B) | NM_147189 | 156.1 | 198.6 | 0.0300 | 1.45 |
| | | XM_928195 | 95.5 | 108.2 | 0.0300 | 1.24 |
| BBS2 | Bardet-Biedl syndrome 2(BBS2) | NM_031885 | 468.1 | 314.5 | 0.0300 | 0.74 |
| BBS2 | Bardet-Biedl syndrome 2(BBS2) | NM_031885 | 712.8 | 548.9 | 0.0300 | 0.74 |
| PSAT1 | phosphoserine aminotransferase 1(PSAT1) | NM_021154 | 136.1 | 140.6 | 0.0300 | 1.2 |
| BAZ1A | bromodomain adjacent to zinc finger domain 1A(BAZ1A) | NM_013448 | 95.7 | 113.1 | 0.0300 | 1.3 |
| | | CN304251 | 97.8 | 103.2 | 0.0300 | 1.14 |
| c19orf60 | chromosome 19 open reading frame 60(C19orf60) | NM_001100419 | 682.4 | 483 | 0.0300 | 0.8 |
| SUMO2 | small ubiquitin-like modifier 2(SUMO2) | NM_001005849 | 399.9 | 512.5 | 0.0300 | 1.47 |
| SUMO2 | small ubiquitin-like modifier 2(SUMO2) | NM_001005849 | 697.5 | 875.6 | 0.0300 | 1.47 |

| | | | | | | |
|---|---|---|---|---|---|---|
| qsox1 | quiescin sulfhydryl oxidase 1(QSOX1) | NM_001004128 | 202.9 | 241.6 | 0.0300 | 1.33 |
| | | XR_039017 | 104.9 | 128.7 | 0.0300 | 1.36 |
| HDLBP | high density lipoprotein binding protein(HDLBP) | NM_203346 | 132.5 | 102.2 | 0.0300 | 0.84 |
| RNF138 | ring finger protein 138(RNF138) | NM_016271 | 107.4 | 91.2 | 0.0300 | 0.88 |
| csnk1g3 | casein kinase 1 gamma 3(CSNK1G3) | NM_001031812 | 125.5 | 132.2 | 0.0310 | 1.17 |
| C1orf87 | chromosome 1 open reading frame 87(C1orf87) | NM_152377 | 94.1 | 91.7 | 0.0310 | 1.11 |
| C1orf87 | chromosome 1 open reading frame 87(C1orf87) | NM_152377 | 104.5 | 105.8 | 0.0310 | 1.11 |
| CRISP2 | cysteine rich secretory protein 2(CRISP2) | NM_003296 | 558.3 | 262.7 | 0.0310 | 0.56 |
| | | AW085986 | 224.5 | 145.6 | 0.0310 | 0.73 |
| ESYT2 | extended synaptotagmin 2(ESYT2) | NM_020728 | 135.9 | 99.6 | 0.0310 | 0.79 |

| | | | | | | |
|---|---|---|---|---|---|---|
| | | XM_928722 | 125.3 | 104.5 | 0.0310 | 0.92 |
| LOC284933 | uncharacterized LOC284933(LOC284933) | BC048428 | 110.5 | 94.7 | 0.0310 | 0.9 |
| MOK | MOK protein kinase(MOK) | NM_014226 | 97.2 | 108 | 0.0310 | 1.21 |
| CDK5R1 | cyclin dependent kinase 5 regulatory subunit 1(CDK5R1) | NM_003885 | 81.6 | 95.8 | 0.0310 | 1.26 |
| CDK5R1 | cyclin dependent kinase 5 regulatory subunit 1(CDK5R1) | NM_003885 | 145.7 | 154.1 | 0.0310 | 1.26 |
| ACTRT1 | actin related protein T1(ACTRT1) | NM_138289 | 211.4 | 138.1 | 0.0310 | 0.73 |
| SIRT6 | sirtuin 6(SIRT6) | NM_016539 | 142.2 | 105.3 | 0.0310 | 0.82 |
| C6orf47 | chromosome 6 open reading frame 47(C6orf47) | NM_021184 | 149.3 | 172.3 | 0.0310 | 1.31 |
| TONSL | tonsoku-like, DNA repair protein(TONSL) | NM_013432 | 197.9 | 143.4 | 0.0310 | 0.82 |

| | | | | | | | | | |
|---|---|---|---|---|---|---|---|---|---|
| GDF11 | growth differentiation factor 11(GDF11) | AK091091 | 122.1 | 142.5 | 0.0310 | 1.3 | | | |
| CLUAP1 | clusterin associated protein 1(CLUAP1) | NM_024793 | 111.7 | 106.1 | 0.0310 | 1.13 | | | |
| CLUAP1 | clusterin associated protein 1(CLUAP1) | NM_024793 | 129.3 | 119.1 | 0.0310 | 1.13 | | | |
| CLUAP1 | clusterin associated protein 1(CLUAP1) | NM_024793 | 16683.3 | 17456.7 | 0.0310 | 1.13 | NM_134394 | 3.46 | 15.03 |
| NAV3 | neuron navigator 3(NAV3) | NM_014903 | 99.5 | 128.2 | 0.0310 | 1.43 | | | |
| LPP | LIM domain containing preferred translocation partner in lipoma(LPP) | NM_005578 | 500.5 | 621.5 | 0.0310 | 1.39 | | | |
| C1orf52 | chromosome 1 open reading frame 52(C1orf52) | NM_198077 | 251.3 | 309.2 | 0.0310 | 1.37 | | | |
| TAF6 | TATA-box binding protein associated factor 6(TAF6) | NM_005641 | 119.5 | 99.8 | 0.0310 | 0.91 | | | |

| | | | | | | |
|---|---|---|---|---|---|---|
| TP53BP2 | tumor protein p53 binding protein 2(TP53BP2) | NM_001031685 | 104.8 | 106.5 | 0.0310 | 1.12 |
| TSHZ3 | teashirt zinc finger homeobox 3(TSHZ3) | NM_020856 | 110.8 | 141.1 | 0.0310 | 1.41 |
| PHIP | pleckstrin homology domain interacting protein(PHIP) | BX537762 | 108.4 | 135.4 | 0.0310 | 1.39 |
| | | XR_037871 | 101.6 | 112.9 | 0.0310 | 1.21 |
| | | XM_929738 | 318.1 | 390.1 | 0.0310 | 1.36 |
| | | XM_001721243 | 92.6 | 98 | 0.0310 | 1.14 |
| | | XM_001717548 | 89.2 | 97.8 | 0.0310 | 1.17 |
| | | XR_038911 | 114.5 | 91.2 | 0.0310 | 0.86 |
| KMT2C | lysine methyltransferase 2C(KMT2C) | NM_170606 | 133.5 | 147.2 | 0.0320 | 0.8 |
| KMT2C | lysine methyltransferase 2C(KMT2C) | NM_170606 | 156 | 111.9 | 0.0320 | 0.8 |
| TRAP1 | TNF receptor associated protein 1(TRAP1) | NM_016292 | 127.9 | 104 | 0.0320 | 0.89 |

| | | | | | | |
|---|---|---|---|---|---|---|
| AIM1L | absent in melanoma 1-like(AIM1L) | NM_017977 | 125.2 | 101.7 | 0.0320 | 0.89 |
| NRCAM | neuronal cell adhesion molecule(NRCAM) | NM_005010 | 107.2 | 117.5 | 0.0320 | 1.2 |
| NRCAM | neuronal cell adhesion molecule(NRCAM) | NM_005010 | 145.4 | 187.6 | 0.0320 | 1.2 |
| | | XR_017484 | 292 | 115.3 | 0.0320 | 0.46 |
| TUBB2B | tubulin beta 2B class IIb(TUBB2B) | NM_178012 | 100.2 | 116.1 | 0.0320 | 1.28 |
| SEPT1 | septin 1(SEPT1) | NM_052838 | 82.4 | 99.8 | 0.0320 | 1.3 |
| MED16 | mediator complex subunit 16(MED16) | NM_005481 | 111.7 | 111.1 | 0.0320 | 1.11 |
| MED16 | mediator complex subunit 16(MED16) | NM_005481 | 128.8 | 116.1 | 0.0320 | 1.11 |
| OSBP2 | oxysterol binding protein 2(OSBP2) | NM_030758 | 117.1 | 121.8 | 0.0320 | 1.17 |
| SP140 | SP140 nuclear body protein(SP140) | NM_007237 | 193.9 | 125.1 | 0.0320 | 0.73 |

| | | | | | | |
|---|---|---|---|---|---|---|
| ACADVL | acyl-CoA dehydrogenase, very long chain(ACADVL) | NM_000018 | 443.6 | 256.4 | 0.0320 | 0.67 |
| USP3 | ubiquitin specific peptidase 3(USP3) | NM_006537 | 116.8 | 95.9 | 0.0320 | 0.88 |
| | | XR_038054 | 316.2 | 119.3 | 0.0320 | 0.44 |
| | | XM_941155 | 557.1 | 931 | 0.0320 | 1.94 |
| | | XM_941155 | 4625 | 5065 | 0.0320 | 1.94 |
| USF1 | upstream transcription factor 1(USF1) | NM_207005 | 146.7 | 215.1 | 0.0320 | 1.61 |
| FAM78A | family with sequence similarity 78 member A(FAM78A) | NM_033387 | 124.2 | 97.4 | 0.0320 | 0.84 |
| | | XM_938838 | 191.4 | 135.8 | 0.0320 | 0.81 |
| | | XR_037962 | 112.9 | 120.3 | 0.0320 | 1.17 |
| TLE4 | transducin like enhancer of split 4(TLE4) | NM_007005 | 103.7 | 114.3 | 0.0320 | 1.2 |
| TLE4 | transducin like enhancer of split 4(TLE4) | NM_007005 | 153.9 | 211.2 | 0.0320 | 1.2 |

| | | | | | | |
|---|---|---|---|---|---|---|
| | | AA883754 | 111.4 | 93 | 0.0320 | 0.88 |
| | | XR_019076 | 93.7 | 96 | 0.0320 | 1.1 |
| ENPP6 | ectonucleotide pyrophosphatase/phosphodiesterase 6(ENPP6) | NM_153343 | 106.5 | 87.9 | 0.0320 | 0.87 |
| RPL6 | ribosomal protein L6(RPL6) | NM_001024662 | 114.1 | 131.7 | 0.0320 | 1.32 |
| RPL6 | ribosomal protein L6(RPL6) | NM_001024662 | 1707.2 | 1910.7 | 0.0320 | 1.32 |
| RPL6 | ribosomal protein L6(RPL6) | NM_001024662 | 2873.7 | 3047.8 | 0.0320 | 1.32 |
| | | AW593356 | 116.2 | 96.3 | 0.0320 | 0.88 |
| DNASE1L1 | deoxyribonuclease 1 like 1(DNASE1L1) | NM_006730 | 107.8 | 110.5 | 0.0320 | 1.12 |
| LOC100190986 | uncharacterized LOC100190986(LOC100190986) | NR_024456 | 118.9 | 97.7 | 0.0320 | 0.9 |
| ZNF606 | zinc finger protein 606(ZNF606) | NM_025027 | 121.2 | 125 | 0.0320 | 1.13 |
| | | XM_940806 | 112.5 | 93.1 | 0.0320 | 0.88 |
| ACD | adrenocortical dysplasia homolog(ACD) | NM_022914 | 145.9 | 145.9 | 0.0330 | 1.13 |
| CEBPZ | CCAAT/enhancer binding protein zeta(CEBPZ) | NM_005760 | 424.8 | 463.6 | 0.0330 | 1.23 |

| | | | | | | |
|---|---|---|---|---|---|---|
| CPM | carboxypeptidase M(CPM) | NM_001005502 | 104.8 | 108.6 | 0.0330 | 1.15 |
| CPM | carboxypeptidase M(CPM) | NM_001005502 | 109.2 | 102.9 | 0.0330 | 1.15 |
| HAT1 | histone acetyltransferase 1(HAT1) | NM_003642 | 134.1 | 121.8 | 0.0330 | 1.24 |
| HAT1 | histone acetyltransferase 1(HAT1) | NM_003642 | 172.6 | 188.1 | 0.0330 | 1.24 |
| TINF2 | TERF1 interacting nuclear factor 2(TINF2) | NM_012461 | 152.6 | 157.5 | 0.0330 | 1.18 |
| MFHAS1 | malignant fibrous histiocytoma amplified sequence 1(MFHAS1) | NM_004225 | 137.7 | 174.5 | 0.0330 | 1.38 |
| ETNK2 | ethanolamine kinase 2(ETNK2) | NM_018208 | 218.3 | 134.7 | 0.0330 | 0.69 |
| PDPK1 | 3-phosphoinositide dependent protein kinase 1(PDPK1) | NM_031268 | 105.6 | 89.3 | 0.0330 | 0.9 |
| PDPK1 | 3-phosphoinositide dependent protein kinase 1(PDPK1) | NM_031268 | 472.1 | 387.7 | 0.0330 | 0.9 |

| | | | | | | |
|---|---|---|---|---|---|---|
| slx4ip | SLX4 interacting protein(SLX4IP) | NM_001009608 | 112.8 | 113.6 | 0.0330 | 1.11 |
| pnkd | paroxysmal nonkinesigenic dyskinesia(PNKD) | NM_022572 | 158.3 | 119.9 | 0.0330 | 0.84 |
| pnkd | paroxysmal nonkinesigenic dyskinesia(PNKD) | NM_022572 | 163.9 | 136.1 | 0.0330 | 0.84 |
| TMEM80 | transmembrane protein 80(TMEM80) | NM_174940 | 206.1 | 135.9 | 0.0330 | 0.76 |
| BUB1 | BUB1 mitotic checkpoint serine/threonine kinase(BUB1) | NM_004336 | 80.2 | 101.5 | 0.0330 | 1.38 |
| CACNA2D4 | calcium voltage-gated channel auxiliary subunit alpha2delta 4(CACNA2D4) | NM_001005737 | 123 | 102.4 | 0.0330 | 0.91 |
| CASP10 | caspase 10(CASP10) | NM_001230 | 91.1 | 99.1 | 0.0330 | 1.15 |
| | | BX117604 | 117.2 | 96.6 | 0.0330 | 0.86 |
| RAB27B | RAB27B, member RAS oncogene family(RAB27B) | AF131784 | 127.3 | 147.8 | 0.0330 | 1.31 |
| | | XR_015468 | 108.4 | 89.4 | 0.0330 | 0.85 |

| | | | | | | |
|---|---|---|---|---|---|---|
| TMIGD1 | transmembrane and immunoglobulin domain containing 1(TMIGD1) | NM_206832 | 133.2 | 94.7 | 0.0330 | 0.76 |
| SOCS4 | suppressor of cytokine signaling 4(SOCS4) | NM_199421 | 102.4 | 117.8 | 0.0330 | 1.29 |
| LCA5L | LCA5L, lebercilin like(LCA5L) | NM_152505 | 131.2 | 104.9 | 0.0330 | 0.88 |
| SPTBN4 | spectrin beta, non-erythrocytic 4(SPTBN4) | NM_020971 | 110 | 92.4 | 0.0330 | 0.86 |
| ZBTB16 | zinc finger and BTB domain containing 16(ZBTB16) | NM_006006 | 90.5 | 107.4 | 0.0330 | 1.28 |
| ZBTB16 | zinc finger and BTB domain containing 16(ZBTB16) | NM_006006 | 172.1 | 263 | 0.0330 | 1.28 |
| NRSN2 | neurensin 2(NRSN2) | NM_024958 | 196.8 | 140.5 | 0.0330 | 0.81 |
| VNN2 | vanin 2(VNN2) | NM_004665 | 103.2 | 108 | 0.0330 | 1.17 |
| STRN | striatin(STRN) | NM_003162 | 160 | 118.2 | 0.0330 | 0.83 |
| | | XM_001716173 | 134.1 | 99.2 | 0.0330 | 0.77 |

| | | | | | | | | | |
|---|---|---|---|---|---|---|---|---|---|
| FHIT | fragile histidine triad(FHIT) | NM_002012 | 111.6 | 94.4 | 0.0330 | 0.89 | | | |
| SHISA4 | shisa family member 4(SHISA4) | NM_198149 | 106.5 | 91.8 | 0.0340 | 0.91 | | | |
| TSPYL4 | TSPY like 4(TSPYL4) | NM_021648 | 145.1 | 109.3 | 0.0340 | 0.83 | | | |
| APOOL | apolipoprotein O like(APOOL) | NM_198450 | 189.3 | 206.1 | 0.0340 | 1.24 | | | |
| HDGF | hepatoma-derived growth factor(HDGF) | NM_004494 | 358.3 | 234.1 | 0.0340 | 0.77 | | | |
| CALHM2 | calcium homeostasis modulator 2(CALHM2) | NM_015916 | 100.3 | 105.4 | 0.0340 | 1.13 | | | |
| LRR1 | leucine rich repeat protein 1(LRR1) | NM_203467 | 92.4 | 110.1 | 0.0340 | 1.3 | | | |
| FANCE | Fanconi anemia complementation group E(FANCE) | NM_021922 | 162.8 | 217.7 | 0.0340 | 1.53 | | | |
| | | XR_015615 | 170.5 | 108.4 | 0.0340 | 0.71 | | | |
| TLR10 | toll like receptor 10(TLR10) | NM_030956 | 95.8 | 118.8 | 0.0340 | 1.36 | | | |
| TLR8 | toll like receptor 8(TLR8) | NM_016610 | 91 | 110.7 | 0.0340 | 1.33 | | | |
| WDR73 | WD repeat domain 73(WDR73) | NM_032856 | 114.2 | 130.2 | 0.0340 | 1.23 | NM_001011972 | 5.50 | 125.06 |

| | | | | | | | | | |
|---|---|---|---|---|---|---|---|---|---|
| | | XM_944290 | 333.5 | 163.5 | 0.0340 | 0.57 | | | |
| CKAP2L | cytoskeleton associated protein 2 like(CKAP2L) | NM_152515 | 104.3 | 115 | 0.0340 | 1.21 | | | |
| QRICH1 | glutamine rich 1(QRICH1) | NM_017730 | 156.8 | 180.2 | 0.0340 | 1.3 | | | |
| PCDH17 | protocadherin 17(PCDH17) | NM_014459 | 158.6 | 577.3 | 0.0340 | 3.92 | | | |
| | | BU561024 | 124.9 | 102.1 | 0.0340 | 0.87 | NM_001107867 | 13.22 | 158.63 |
| CLCF1 | cardiotrophin-like cytokine factor 1(CLCF1) | NM_013246 | 212.3 | 141.9 | 0.0340 | 0.75 | | | |
| SYNE1 | spectrin repeat containing nuclear envelope protein 1(SYNE1) | NM_033071 | 123.7 | 86.7 | 0.0340 | 0.75 | | | |
| CLOCK | clock circadian regulator(CLOCK) | NM_004898 | 267.2 | 163.1 | 0.0340 | 0.71 | | | |
| PRKDC | protein kinase, DNA-activated, catalytic polypeptide(PRKDC) | NM_0010811640 | 485.4 | 581.2 | 0.0340 | 1.38 | NM_001014199 | 0.43 | 104.20 |
| TUBB4A | tubulin beta 4A class IVa(TUBB4A) | NM_006087 | 171.7 | 128.1 | 0.0340 | 0.84 | | | |

| | | | | | | | | | |
|---|---|---|---|---|---|---|---|---|---|
| POM121C | POM121 transmembrane nucleoporin C(POM121C) | NM_001099415 | 111.7 | 119.3 | 0.0340 | 1.19 | NM_198745 | 128.13 | 517.66 |
| DUS3L | dihydrouridine synthase 3 like(DUS3L) | NM_020175 | 137 | 153.6 | 0.0340 | 1.26 | NM_198745 | 128.13 | 517.66 |
| | | XM_927468 | 104.4 | 106 | 0.0340 | 1.1 | | | |
| | | XM_496054 | 95.6 | 101.7 | 0.0340 | 1.14 | | | |
| TTPAL | alpha tocopherol transfer protein like(TTPAL) | NM_024331 | 138.7 | 154.6 | 0.0340 | 1.27 | | | |
| | | CK299443 | 149.4 | 117.2 | 0.0340 | 0.88 | NM_001105991 | 1.24 | 549.69 |
| CCT8L1P | chaperonin containing TCP1 subunit 8 like 1, pseudogene(CCT8L1P) | NM_001029866 | 104.4 | 105.4 | 0.0340 | 1.1 | NM_001105991 | 1.24 | 549.69 |
| RBM10 | RNA binding motif protein 10(RBM10) | NM_152856 | 283.8 | 311.5 | 0.0350 | 1.25 | NM_001013929 | 4.59 | 30.85 |
| YIF1B | Yip1 interacting factor homolog B, membrane trafficking protein(YIF1B) | NM_033557 | 106.6 | 107.7 | 0.0350 | 1.11 | NM_001013929 | 4.59 | 30.85 |
| | | XM_001723141 | 695.1 | 854.9 | 0.0350 | 1.35 | | | |

| | | | | | | |
|---|---|---|---|---|---|---|
| | | XR_037908 | 112.4 | 93.7 | 0.0350 | 0.89 |
| PCDHGC4 | protocadherin gamma subfamily C, 4(PCDHGC4) | NM_032406 | 120.7 | 98.4 | 0.0350 | 0.88 |
| KLHL36 | kelch like family member 36(KLHL36) | NM_024731 | 151.1 | 120.3 | 0.0350 | 0.87 |
| | | XM_0011130428 | 130.9 | 105.5 | 0.0350 | 0.89 |
| TAGLN | transgelin(TAGLN) | NM_0010011522 | 126.5 | 104.2 | 0.0350 | 0.89 |
| C15orf56 | chromosome 15 open reading frame 56(C15orf56) | NM_001039905 | 118.4 | 97.5 | 0.0350 | 0.89 |
| RBMXL2 | RNA binding motif protein, X-linked like 2(RBMXL2) | NM_014469 | 106.8 | 88.7 | 0.0350 | 0.87 |
| SEC31A | SEC31 homolog A, COPII coat complex component(SEC31A) | NM_014933 | 115.5 | 110.1 | 0.0350 | 1.37 |
| SEC31A | SEC31 homolog A, COPII coat complex component(SEC31A) | NM_014933 | 207.9 | 230.8 | 0.0350 | 1.37 |

| | | | | | | |
|---|---|---|---|---|---|---|
| SEC31A | SEC31 homolog A, COPII coat complex component(SEC31A) | NM_014933 | 462.7 | 564.9 | 0.0350 | 1.37 |
| MYCBP2 | MYC binding protein 2, E3 ubiquitin protein ligase(MYCBP2) | NM_015057 | 224.3 | 362.9 | 0.0350 | 1.82 |
| GSTA3 | glutathione S-transferase alpha 3(GSTA3) | NM_000847 | 89 | 95.1 | 0.0350 | 1.15 |
| | | AI627272 | 113 | 113.4 | 0.0350 | 1.11 |
| | | BX092585 | 112.7 | 94.3 | 0.0350 | 0.87 |
| SCFD2 | sec1 family domain containing 2(SCFD2) | NM_152540 | 105.5 | 117.9 | 0.0350 | 1.22 |
| CD27-AS1 | CD27 antisense RNA 1(CD27-AS1) | NR_015382 | 107.9 | 116.6 | 0.0350 | 1.18 |
| ATRX | ATRX, chromatin remodeler(ATRX) | NM_000489 | 102.8 | 113 | 0.0350 | 1.2 |
| RWDD1 | RWD domain containing 1(RWDD1) | NM_0010007464 | 605 | 699.3 | 0.0350 | 1.29 |
| | | XM_930558 | 131 | 104.6 | 0.0350 | 0.88 |
| AOAH | acyloxyacyl hydrolase(AOAH) | NM_001637 | 87.8 | 95.9 | 0.0350 | 1.16 |

| | | | | | | |
|---|---|---|---|---|---|---|
| SRPK2 | SRSF protein kinase 2(SRPK2) | NM_182692 | 105.6 | 109.9 | 0.0350 | 1.14 |
| CTDP1 | CTD phosphatase subunit 1(CTDP1) | NM_004715 | 103.6 | 101.4 | 0.0350 | 0.86 |
| CTDP1 | CTD phosphatase subunit 1(CTDP1) | NM_004715 | 122 | 96.1 | 0.0350 | 0.86 |
| | | XM_941100 | 95 | 109.4 | 0.0350 | 1.26 |
| NOTUM | NOTUM, palmitoleoyl-protein carboxylesterase(NOTUM) | NM_178493 | 112.2 | 96.9 | 0.0360 | 0.92 |
| DTD2 | D-tyrosyl-tRNA deacylase 2 (putative)(DTD2) | NM_080664 | 105.6 | 111.8 | 0.0360 | 1.16 |
| USP33 | ubiquitin specific peptidase 33(USP33) | NM_201624 | 95.5 | 100.8 | 0.0360 | 1.13 |
| | | BX641358 | 100.2 | 104 | 0.0360 | 1.13 |
| | | XM_001720809 | 113.1 | 91.1 | 0.0360 | 0.85 |
| DUSP23 | dual specificity phosphatase 23(DUSP23) | NM_017823 | 361.1 | 247.6 | 0.0360 | 0.77 |

| | | | | | | |
|---|---|---|---|---|---|---|
| RITA1 | RBPJ interacting and tubulin associated 1(RITA1) | NM_032848 | 157.2 | 120.9 | 0.0360 | 0.85 |
| LOC613206 | myeloproliferative disease associated tumor antigen 5(LOC613206) | NM_001033016 | 113.9 | 92.9 | 0.0360 | 0.87 |
| DLGAP4 | DLG associated protein 4(DLGAP4) | NM_001042486 | 176.7 | 189.6 | 0.0360 | 1.21 |
| TRIM47 | tripartite motif containing 47(TRIM47) | NM_033452 | 104.5 | 105.6 | 0.0360 | 1.1 |
| SELENOS | selenoprotein S(SELENOS) | NM_018445 | 167.6 | 206.1 | 0.0360 | 1.38 |
| SELENOS | selenoprotein S(SELENOS) | NM_018445 | 911.9 | 899.4 | 0.0360 | 1.38 |
| UGT1A6 | UDP glucuronosyltransferase family 1 member A6(UGT1A6) | NM_001072 | 146.1 | 107.1 | 0.0360 | 0.69 |
| UGT1A6 | UDP glucuronosyltransferase family 1 member A6(UGT1A6) | NM_001072 | 161.1 | 106.8 | 0.0360 | 0.69 |

| | | | | | | | | | |
|---|---|---|---|---|---|---|---|---|---|
| AP1B1 | adaptor related protein complex 1 beta 1 subunit(AP1B1) | NM_145730 | 192.3 | 242.6 | 0.0360 | 1.43 | | | |
| METTL21B | methyltransferase like 21B(METTL21B) | NM_015433 | 698.4 | 358.2 | 0.0360 | 0.57 | | | |
| MIR561 | microRNA 561(MIR561) | NR_030287 | 104 | 108.4 | 0.0360 | 1.15 | | | |
| | | BQ028043 | 119.2 | 98 | 0.0360 | 0.87 | | | |
| | | BQ021435 | 121.2 | 100.1 | 0.0360 | 0.89 | NM_019136 | 6.29 | 354.89 |
| | | XM_944418 | 93.7 | 100 | 0.0360 | 1.15 | | | |
| KIF20A | kinesin family member 20A(KIF20A) | NM_005733 | 95.5 | 103.7 | 0.0360 | 1.17 | | | |
| ASB3 | ankyrin repeat and SOCS box containing 3(ASB3) | NM_145863 | 100.7 | 106.7 | 0.0360 | 1.15 | | | |
| AAED1 | AhpC/TSA antioxidant enzyme domain containing 1(AAED1) | NM_153698 | 124.3 | 134.3 | 0.0360 | 1.21 | | | |
| CLEC2D | C-type lectin domain family 2 member D(CLEC2D) | NM_013269 | 118 | 102 | 0.0360 | 1.3 | | | |

| | | | | | | |
|---|---|---|---|---|---|---|
| CLEC2D | C-type lectin domain family 2 member D(CLEC2D) | NM_013269 | 2236.2 | 2836.9 | 0.0360 | 1.3 |
| | | XM_938002 | 100.5 | 105.8 | 0.0360 | 1.12 |
| NKRF | NFKB repressing factor(NKRF) | NM_017544 | 132.2 | 141 | 0.0360 | 1.2 |
| FAM3D | family with sequence similarity 3 member D(FAM3D) | NM_138805 | 125.4 | 98.5 | 0.0360 | 0.87 |
| FAM162B | family with sequence similarity 162 member B(FAM162B) | NM_001085480 | 107.3 | 114.9 | 0.0360 | 1.19 |
| C8orf82 | chromosome 8 open reading frame 82(C8orf82) | NM_001001795 | 107.7 | 107.7 | 0.0360 | 1.1 |
| OR5P3 | olfactory receptor family 5 subfamily P member 3(OR5P3) | NM_153445 | 127.9 | 106.9 | 0.0360 | 0.91 |
| STOX2 | storkhead box 2(STOX2) | NM_020225 | 141.9 | 197.9 | 0.0360 | 1.58 |
| | | CR748464 | 97.2 | 100 | 0.0370 | 1.11 |
| | | XM_001125808 | 124.7 | 134 | 0.0370 | 1.19 |

| | | | | | | |
|---|---|---|---|---|---|---|
| DDX39B | DExD-box helicase 39B(DDX39B) | NM_080598 | 770.4 | 967 | 0.0370 | 1.42 |
| ITGB4 | integrin subunit beta 4(ITGB4) | NM_001005619 | 103.6 | 97.8 | 0.0370 | 1.12 |
| ITGB4 | integrin subunit beta 4(ITGB4) | NM_001005619 | 105.4 | 108.2 | 0.0370 | 1.12 |
| TBX3 | T-box 3(TBX3) | NM_005996 | 111.7 | 114.8 | 0.0370 | 1.13 |
| KLC4 | kinesin light chain 4(KLC4) | NM_201521 | 107.2 | 87.8 | 0.0370 | 0.87 |
| DNMT3A | DNA methyltransferase 3 alpha(DNMT3A) | NM_175630 | 94.8 | 100.2 | 0.0370 | 1.15 |
| | | AA383317 | 114.3 | 95.5 | 0.0370 | 0.89 |
| | | XM_001717251 | 88.9 | 98.2 | 0.0370 | 1.18 |
| | | XM_928923 | 233.6 | 146.9 | 0.0370 | 0.72 |
| HEATR4 | HEAT repeat containing 4(HEATR4) | NM_203309 | 115.3 | 95.7 | 0.0370 | 0.89 |
| MEF2C | myocyte enhancer factor 2C(MEF2C) | NM_002397 | 107.2 | 199.8 | 0.0370 | 2.08 |
| ZNF75A | zinc finger protein 75a(ZNF75A) | NM_153028 | 97.5 | 103 | 0.0370 | 1.14 |
| ANKRD49 | ankyrin repeat domain 49(ANKRD49) | NM_017704 | 168.1 | 179.1 | 0.0370 | 1.19 |

| | | | | | | |
|---|---|---|---|---|---|---|
| TOX4 | TOX high mobility group box family member 4(TOX4) | NM_014828 | 104.8 | 109.5 | 0.0370 | 1.13 |
| COL11A1 | collagen type XI alpha 1 chain(COL11A1) | NM_080629 | 129.7 | 96.4 | 0.0370 | 0.82 |
| CNST | consortin, connexin sorting protein(CNST) | NM_152609 | 259.4 | 427.8 | 0.0370 | 1.86 |
| | | XM_940361 | 103.5 | 106.4 | 0.0370 | 1.13 |
| PLEC | plectin(PLEC) | NM_201380 | 130.1 | 102.7 | 0.0370 | 0.87 |
| GLIPR2 | GLI pathogenesis related 2(GLIPR2) | NM_022343 | 99.5 | 115.4 | 0.0380 | 1.29 |
| DEFB123 | defensin beta 123(DEFB123) | NM_153324 | 395 | 184.7 | 0.0380 | 0.55 |
| | | XM_938495 | 132.5 | 106.1 | 0.0380 | 0.87 |
| OGDH | oxoglutarate dehydrogenase(OGDH) | NM_001003941 | 98.9 | 103.2 | 0.0380 | 1.12 |
| BPNT1 | 3'(2'), 5'-bisphosphate nucleotidase 1(BPNT1) | NM_006085 | 116.7 | 128.3 | 0.0380 | 1.2 |

| | | | | | | |
|---|---|---|---|---|---|---|
| CTR9 | CTR9 homolog, Paf1/RNA polymerase II complex component(CTR9) | NM_014633 | 118 | 94.8 | 0.0380 | 0.87 |
| RANBP17 | RAN binding protein 17(RANBP17) | NM_022897 | 164.1 | 119.8 | 0.0380 | 0.78 |
| U2AF2 | U2 small nuclear RNA auxiliary factor 2(U2AF2) | NM_007279 | 103.3 | 109.6 | 0.0380 | 1.16 |
| U2AF2 | U2 small nuclear RNA auxiliary factor 2(U2AF2) | NM_007279 | 663.9 | 593.8 | 0.0380 | 1.16 |
| | | XM_941035 | 116.1 | 94 | 0.0380 | 0.88 |
| | | XM_001724530 | 116.3 | 97.3 | 0.0380 | 0.9 |
| SNORD56 | small nucleolar RNA, C/D box 56(SNORD56) | NR_002739 | 105.8 | 120.7 | 0.0380 | 1.26 |
| RPL13 | ribosomal protein L13(RPL13) | NM_033251 | 110.1 | 124.2 | 0.0380 | 1.25 |
| HYDIN | HYDIN, axonemal central pair apparatus protein(HYDIN) | NM_032821 | 103 | 98.1 | 0.0380 | 0.89 |

| | | | | | | |
|---|---|---|---|---|---|---|
| HYDIN | HYDIN, axonemal central pair apparatus protein(HYDIN) | NM_032821 | 112.7 | 94.6 | 0.0380 | 0.89 |
| HYDIN | HYDIN, axonemal central pair apparatus protein(HYDIN) | NM_032821 | 137.8 | 117.2 | 0.0380 | 0.89 |
| HYDIN | HYDIN, axonemal central pair apparatus protein(HYDIN) | NM_032821 | 150.8 | 123.4 | 0.0380 | 0.89 |
| DSP | desmoplakin(DSP) | NM_001008844 | 183.9 | 173.5 | 0.0380 | 0.8 |
| DSP | desmoplakin(DSP) | NM_001008844 | 190.6 | 134.1 | 0.0380 | 0.8 |
| EIF6 | eukaryotic translation initiation factor 6(EIF6) | NM_181469 | 1332.8 | 937.9 | 0.0380 | 0.79 |
| BCAP29 | B-cell receptor associated protein 29(BCAP29) | NM_001008405 | 178.4 | 216.6 | 0.0380 | 1.33 |
| RABGGTA | Rab geranylgeranyltransferase alpha subunit(RABGGTA) | NM_004581 | 99.8 | 104.8 | 0.0380 | 1.14 |
| KIAA1191 | KIAA1191(KIAA1191) | NM_001079685 | 109.4 | 91.3 | 0.0380 | 0.88 |

| | | | | | | | | | |
|---|---|---|---|---|---|---|---|---|---|
| DECR1 | 2,4-dienoyl-CoA reductase 1, mitochondrial(DECR1) | NM_001359 | 1053.4 | 548 | 0.0380 | 0.6 | | | |
| LGALS2 | galectin 2(LGALS2) | NM_006498 | 1166.8 | 439.6 | 0.0380 | 0.41 | | | |
| bag4 | BCL2 associated athanogene 4(BAG4) | NM_004874 | 143.4 | 152.1 | 0.0390 | 1.19 | NM_139082 | 11.11 | 0.48 |
| RACK1 | receptor for activated C kinase 1(RACK1) | NM_006098 | 1619.3 | 1944.8 | 0.0390 | 1.3 | | | |
| ACBD5 | acyl-CoA binding domain containing 5(ACBD5) | NM_145698 | 106.6 | 107.3 | 0.0390 | 1.11 | | | |
| TEDDM1 | transmembrane epididymal protein 1(TEDDM1) | NM_172000 | 188.6 | 132.9 | 0.0390 | 0.81 | | | |
| HAND1 | heart and neural crest derivatives expressed 1(HAND1) | NM_004821 | 92.7 | 98 | 0.0390 | 1.13 | | | |
| ASB16-AS1 | ASB16 antisense RNA 1(ASB16-AS1) | NM_178542 | 153 | 115.6 | 0.0390 | 0.84 | | | |
| CDC14A | cell division cycle 14A(CDC14A) | NM_003672 | 124 | 128 | 0.0390 | 1.14 | | | |

| | | | | | | |
|---|---|---|---|---|---|---|
| GDAP2 | ganglioside induced differentiation associated protein 2(GDAP2) | NM_017686 | 193 | 201 | 0.0390 | 1.19 |
| | | CD516455 | 117 | 97.1 | 0.0390 | 0.89 |
| ZDHHC4 | zinc finger DHHC-type containing 4(ZDHHC4) | NM_018106 | 200.7 | 149 | 0.0390 | 0.83 |
| EEF2 | eukaryotic translation elongation factor 2(EEF2) | NM_001961 | 172.4 | 112.2 | 0.0390 | 0.74 |
| CER1 | cerberus 1, DAN family BMP antagonist(CER1) | NM_005454 | 147.2 | 114.8 | 0.0390 | 0.86 |
| | | BF054947 | 116.6 | 99.1 | 0.0390 | 0.92 |
| | | XR_015400 | 208.9 | 238.3 | 0.0390 | 1.29 |
| TMEM8B | transmembrane protein 8B(TMEM8B) | NM_016446 | 98.6 | 102.4 | 0.0390 | 1.13 |
| | | NM_0010099937 | 171.7 | 127.3 | 0.0390 | 0.82 |
| | | XM_0017155992 | 147.3 | 111.7 | 0.0390 | 0.86 |
| RYK | receptor-like tyrosine kinase(RYK) | NM_0010055861 | 121.1 | 109.5 | 0.0390 | 1.37 |

| | | | | | | |
|---|---|---|---|---|---|---|
| RYK | receptor-like tyrosine kinase(RYK) | NM_001005861 | 217.5 | 269 | 0.0390 | 1.37 |
| ZNF607 | zinc finger protein 607(ZNF607) | NM_032689 | 97.8 | 107 | 0.0390 | 1.19 |
| ZNF607 | zinc finger protein 607(ZNF607) | NM_032689 | 101.9 | 92.3 | 0.0390 | 1.19 |
| | | XM_931025 | 118.9 | 123.8 | 0.0390 | 1.17 |
| RNF181 | ring finger protein 181(RNF181) | NM_016494 | 550.9 | 360.6 | 0.0390 | 0.74 |
| adck5 | aarF domain containing kinase 5(ADCK5) | NM_174922 | 139.7 | 146.4 | 0.0390 | 1.16 |
| | | XM_933681 | 143.3 | 111.4 | 0.0390 | 0.85 |
| | | XR_039005 | 156.7 | 110.3 | 0.0390 | 0.78 |
| DYRK1B | dual specificity tyrosine phosphorylation regulated kinase 1B(DYRK1B) | NM_006484 | 177.2 | 130.8 | 0.0390 | 0.82 |
| HELQ | helicase, POLQ-like(HELQ) | NM_133636 | 164.1 | 191.9 | 0.0390 | 1.3 |
| SYT7 | synaptotagmin 7(SYT7) | NM_004200 | 117.2 | 139.6 | 0.0390 | 1.35 |
| | | XM_942604 | 98.7 | 102.3 | 0.0390 | 1.12 |
| | | XM_001717551 | 101.5 | 103.1 | 0.0390 | 1.09 |

| | | | | | | |
|---|---|---|---|---|---|---|
| NEDD4 | neural precursor cell expressed, developmentally down-regulated 4, E3 ubiquitin protein ligase(NEDD4) | NM_006154 | 95.9 | 106.7 | 0.0390 | 1.21 |
| NEDD4 | neural precursor cell expressed, developmentally down-regulated 4, E3 ubiquitin protein ligase(NEDD4) | NM_006154 | 202.6 | 173.9 | 0.0390 | 1.21 |
| | | XR_017364 | 236.8 | 139.8 | 0.0390 | 0.65 |
| HSD17B7P2 | hydroxysteroid 17-beta dehydrogenase 7 pseudogene 2(HSD17B7P2) | NR_003086 | 98.5 | 99.7 | 0.0390 | 1.09 |
| MEF2A | myocyte enhancer factor 2A(MEF2A) | NM_005587 | 97.5 | 93.3 | 0.0390 | 1.3 |
| MEF2A | myocyte enhancer factor 2A(MEF2A) | NM_005587 | 104 | 122.2 | 0.0390 | 1.3 |
| KATNA1 | katanin catalytic subunit A1(KATNA1) | NM_007044 | 190.6 | 226.5 | 0.0390 | 1.32 |

| | | | | | | | | | |
|---|---|---|---|---|---|---|---|---|---|
| FAM26D | family with sequence similarity 26 member D(FAM26D) | NM_153036 | 106.6 | 92.8 | 0.0400 | 0.92 | | | |
| FAM26D | family with sequence similarity 26 member D(FAM26D) | NM_153036 | 121.7 | 103.4 | 0.0400 | 0.92 | | | |
| | | XM_929427 | 156.5 | 119.7 | 0.0400 | 0.86 | NM_017253 | 1.16 | 16.79 |
| ELP5 | elongator acetyltransferase complex subunit 5(ELP5) | NM_203413 | 132.5 | 137.6 | 0.0400 | 1.19 | | | |
| ELP5 | elongator acetyltransferase complex subunit 5(ELP5) | NM_203413 | 247.8 | 200.5 | 0.0400 | 1.19 | | | |
| HCCS | holocytochrome c synthase(HCCS) | NM_005333 | 109.1 | 100.7 | 0.0400 | 0.66 | | | |
| HCCS | holocytochrome c synthase(HCCS) | NM_005333 | 302 | 166.4 | 0.0400 | 0.66 | | | |
| PTPN23 | protein tyrosine phosphatase, non-receptor type 23(PTPN23) | NM_015466 | 101.1 | 106.7 | 0.0400 | 1.16 | | | |
| RBM39 | RNA binding motif protein 39(RBM39) | NM_184237 | 103.9 | 114.3 | 0.0400 | 1.22 | | | |

| | | | | | | |
|---|---|---|---|---|---|---|
| HMGCS1 | 3-hydroxy-3-methylglutaryl-CoA synthase 1(HMGCS1) | NM_002130 | 599.5 | 385 | 0.0400 | 0.73 |
| SOX21 | SRY-box 21(SOX21) | NM_007084 | 111.9 | 88.8 | 0.0400 | 0.85 |
| SRPK1 | SRSF protein kinase 1(SRPK1) | NM_003137 | 110.4 | 89.9 | 0.0400 | 0.87 |
| SUCLG2 | succinate-CoA ligase GDP-forming beta subunit(SUCLG2) | NM_003848 | 157.2 | 106.9 | 0.0400 | 0.76 |
| | | XM_379270 | 119.7 | 93.6 | 0.0400 | 0.82 |
| | | XM_938817 | 116.7 | 98.7 | 0.0400 | 0.92 |
| STAMBPL1 | STAM binding protein like 1(STAMBPL1) | NM_020799 | 121.7 | 178.1 | 0.0400 | 1.67 |
| H3F3AP4 | H3 histone, family 3A, pseudogene 4(H3F3AP4) | NR_002315 | 468.8 | 162.3 | 0.0400 | 0.41 |
| H3F3AP4 | H3 histone, family 3A, pseudogene 4(H3F3AP4) | NR_002315 | 2589.7 | 2420.9 | 0.0400 | 0.41 |
| LIMS1 | LIM zinc finger domain containing 1(LIMS1) | NM_004987 | 96.9 | 88.8 | 0.0400 | 1.15 |

| | | | | | | |
|---|---|---|---|---|---|---|
| LIMS1 | LIM zinc finger domain containing 1(LIMS1) | NM_004987 | 136 | 138.6 | 0.0400 | 1.15 |
| PRSS3P2 | protease, serine 3 pseudogene 2(PRSS3P2) | NR_001296 | 95.5 | 100.1 | 0.0400 | 1.12 |
| PYGL | phosphorylase, glycogen, liver(PYGL) | NM_002863 | 149.1 | 113.9 | 0.0400 | 0.83 |
| CYP2S1 | cytochrome P450 family 2 subfamily S member 1(CYP2S1) | NM_030622 | 134.6 | 108.5 | 0.0400 | 0.89 |
| S100A10 | S100 calcium binding protein A10(S100A10) | NM_002966 | 128.4 | 98.1 | 0.0400 | 0.82 |
| S100A10 | S100 calcium binding protein A10(S100A10) | NM_002966 | 679.2 | 583.1 | 0.0400 | 0.82 |
| ZNHIT1 | zinc finger HIT-type containing 1(ZNHIT1) | NM_006349 | 1690.2 | 1015 | 0.0400 | 0.68 |
| | | XM_928726 | 147 | 114.2 | 0.0400 | 0.86 |
| DFNA5 | DFNA5, deafness associated tumor suppressor(DFNA5) | NM_004403 | 238.1 | 143.6 | 0.0400 | 0.65 |
| | | XM_942417 | 112.2 | 94.9 | 0.0400 | 0.89 |

| | | | | | | |
|---|---|---|---|---|---|---|
| USP1 | ubiquitin specific peptidase 1(USP1) | NM_003368 | 93.7 | 107.1 | 0.0400 | 1.24 |
| | | BU182441 | 133 | 107.6 | 0.0400 | 0.88 |
| | | XM_936375 | 90.7 | 96.5 | 0.0400 | 1.14 |
| CNOT3 | CCR4-NOT transcription complex subunit 3(CNOT3) | NM_014516 | 102.1 | 107.7 | 0.0400 | 1.15 |
| ABCC3 | ATP binding cassette subfamily C member 3(ABCC3) | NM_020038 | 108.7 | 109.2 | 0.0410 | 1.09 |
| | | XM_931381 | 336.9 | 181.6 | 0.0410 | 0.62 |
| KRTAP13-4 | keratin associated protein 13-4(KRTAP13-4) | NM_181600 | 132.8 | 104.3 | 0.0410 | 0.86 |
| SLC25A48 | solute carrier family 25 member 48(SLC25A48) | NM_145282 | 182.4 | 137.1 | 0.0410 | 0.85 |
| PRAM1 | PML-RARA regulated adaptor molecule 1(PRAM1) | NM_032152 | 117.5 | 101.1 | 0.0410 | 0.92 |
| BIRC2 | baculoviral IAP repeat containing 2(BIRC2) | NM_001166 | 128.7 | 131.1 | 0.0410 | 1.3 |

| | | | | | | |
|---|---|---|---|---|---|---|
| BIRC2 | baculoviral IAP repeat containing 2(BIRC2) | NM_001166 | 171.2 | 202.2 | 0.0410 | 1.3 |
| | | BU734093 | 136.2 | 141.7 | 0.0410 | 1.18 |
| | | XM_001719745 | 104.4 | 123.6 | 0.0410 | 1.29 |
| FKRP | fukutin related protein(FKRP) | NM_024301 | 111.4 | 118.1 | 0.0410 | 1.17 |
| MAP2K2 | mitogen-activated protein kinase kinase 2(MAP2K2) | NM_030662 | 118.9 | 93 | 0.0410 | 0.85 |
| BCOR | BCL6 corepressor(BCOR) | NM_017745 | 215.9 | 250.5 | 0.0410 | 1.3 |
| JADE1 | jade family PHD finger 1(JADE1) | NM_199320 | 202.2 | 150.5 | 0.0410 | 0.83 |
| NFAT5 | nuclear factor of activated T-cells 5(NFAT5) | NM_173215 | 101.4 | 108 | 0.0410 | 1.16 |
| ASUN | asunder, spermatogenesis regulator(ASUN) | NM_018164 | 103.3 | 108.8 | 0.0410 | 1.18 |
| ASUN | asunder, spermatogenesis regulator(ASUN) | NM_018164 | 133.4 | 139.3 | 0.0410 | 1.18 |
| MKNK2 | MAP kinase interacting serine/threonine kinase 2(MKNK2) | NM_199054 | 352.4 | 221.1 | 0.0410 | 0.73 |
| | | CB048233 | 94.5 | 101.6 | 0.0410 | 1.16 |

| | | | | | | |
|---|---|---|---|---|---|---|
| TMED9 | transmembrane p24 trafficking protein 9(TMED9) | NM_017510 | 262.4 | 187.4 | 0.0410 | 0.8 |
| CNIH2 | cornichon family AMPA receptor auxiliary protein 2(CNIH2) | NM_182553 | 111.8 | 103.9 | 0.0410 | 0.83 |
| CNIH2 | cornichon family AMPA receptor auxiliary protein 2(CNIH2) | NM_182553 | 136.5 | 102.3 | 0.0410 | 0.83 |
| CNIH2 | cornichon family AMPA receptor auxiliary protein 2(CNIH2) | NM_182553 | 164.8 | 133.1 | 0.0410 | 0.83 |
| GPR180 | G protein-coupled receptor 180(GPR180) | NM_180989 | 122.3 | 130.2 | 0.0410 | 1.22 |
| | | BU685158 | 98.4 | 101 | 0.0410 | 1.12 |
| PRICKLE1 | prickle planar cell polarity protein 1(PRICKLE1) | NM_153026 | 126.2 | 161.6 | 0.0410 | 1.43 |
| | | XM_943683 | 118.6 | 98.8 | 0.0410 | 0.9 |
| SSR1 | signal sequence receptor subunit 1(SSR1) | NM_003144 | 443 | 498.9 | 0.0410 | 1.26 |
| NFX1 | nuclear transcription factor, X-box binding 1(NFX1) | NM_147133 | 121.5 | 125.1 | 0.0410 | 1.16 |

| | | | | | | |
|---|---|---|---|---|---|---|
| NFX1 | nuclear transcription factor, X-box binding 1(NFX1) | NM_147133 | 181.8 | 136.2 | 0.0410 | 1.16 |
| MT1E | metallothionein 1E(MT1E) | NM_175617 | 130.8 | 103.5 | 0.0410 | 0.41 |
| MT1E | metallothionein 1E(MT1E) | NM_175617 | 5309.7 | 1957.7 | 0.0410 | 0.41 |
| NEMF | nuclear export mediator factor(NEMF) | NM_004713 | 205.5 | 245.1 | 0.0410 | 1.32 |
| POTEH | POTE ankyrin domain family member H(POTEH) | NM_001004053 | 103.1 | 112 | 0.0410 | 1.18 |
| | | XR_040084 | 95.9 | 100 | 0.0410 | 1.13 |
| MAPK8IP2 | mitogen-activated protein kinase 8 interacting protein 2(MAPK8IP2) | NM_016431 | 143 | 124 | 0.0410 | 0.88 |
| MAPK8IP2 | mitogen-activated protein kinase 8 interacting protein 2(MAPK8IP2) | NM_016431 | 146.3 | 115.8 | 0.0410 | 0.88 |
| | | XR_038700 | 115.6 | 89.3 | 0.0420 | 0.84 |

| | | | | | | | | | |
|---|---|---|---|---|---|---|---|---|---|
| RAPGEF1 | Rap guanine nucleotide exchange factor 1(RAPGEF1) | NM_005312 | 144.9 | 115 | 0.0420 | 0.89 | | | |
| STK10 | serine/threonine kinase 10(STK10) | NM_005990 | 211.8 | 147.7 | 0.0420 | 0.8 | NM_019368 | 2.87 | 13.68 |
| | | XM_001725448 | 135 | 109.8 | 0.0420 | 0.88 | | | |
| PSENEN | presenilin enhancer gamma-secretase subunit(PSENEN) | NM_172341 | 269.7 | 198.5 | 0.0420 | 0.85 | | | |
| UIMC1 | ubiquitin interaction motif containing 1(UIMC1) | NM_016290 | 199.8 | 249.3 | 0.0420 | 1.4 | | | |
| | | XM_001128559 | 118.5 | 89.6 | 0.0420 | 0.81 | | | |
| KMT2E | lysine methyltransferase 2E(KMT2E) | NM_182931 | 144.7 | 190.2 | 0.0420 | 1.49 | | | |
| KMT2E | lysine methyltransferase 2E(KMT2E) | NM_182931 | 171 | 135.3 | 0.0420 | 1.49 | | | |
| OGDH | oxoglutarate dehydrogenase(OGDH) | NM_002541 | 333.9 | 208.1 | 0.0420 | 0.71 | NM_001037554 | 15.49 | 101.99 |

| | | | | | | | | | |
|---|---|---|---|---|---|---|---|---|---|
| B3GALNT2 | beta-1,3-N-acetylgalactosaminyltransferase 2(B3GALNT2) | NM_152490 | 118.6 | 137 | 0.0420 | 1.29 | | | |
| DCTD | dCMP deaminase(DCTD) | NM_001921 | 142.3 | 148.9 | 0.0420 | 1.15 | | | |
| HLA-DPB1 | major histocompatibility complex, class II, DP beta 1(HLA-DPB1) | NM_002121 | 116.5 | 95.9 | 0.0420 | 0.89 | NM_017087 | 51.82 | 0.25 |
| | | XM_933749 | 123.3 | 101.2 | 0.0420 | 0.9 | | | |
| NDUFA10 | NADH:ubiquinone oxidoreductase subunit A10(NDUFA10) | NM_004544 | 102.4 | 96.8 | 0.0420 | 0.79 | | | |
| NDUFA10 | NADH:ubiquinone oxidoreductase subunit A10(NDUFA10) | NM_004544 | 118.1 | 88.1 | 0.0420 | 0.79 | | | |
| OR2A5 | olfactory receptor family 2 subfamily A member 5(OR2A5) | NM_012365 | 111.1 | 91.7 | 0.0420 | 0.88 | | | |
| | | XM_001720686 | 194.7 | 126 | 0.0420 | 0.73 | | | |
| GLTSCR1L | GLTSCR1 like(GLTSCR1L) | NM_015349 | 150.6 | 208.6 | 0.0420 | 1.5 | | | |

| | | | | | | |
|---|---|---|---|---|---|---|
| TM7SF3 | transmembrane 7 superfamily member 3(TM7SF3) | NM_016551 | 189.5 | 131.8 | 0.0420 | 0.8 |
| PXMP4 | peroxisomal membrane protein 4(PXMP4) | NM_183397 | 129.5 | 133.2 | 0.0420 | 1.15 |
| | | XR_039263 | 114.9 | 115.4 | 0.0420 | 1.13 |
| SNORD 113-1 | small nucleolar RNA, C/D box 113-1(SNORD113-1) | NR_003229 | 101.4 | 107.8 | 0.0420 | 1.17 |
| CBX5 | chromobox 5(CBX5) | NM_012117 | 110.4 | 144.6 | 0.0420 | 1.48 |
| ALPK3 | alpha kinase 3(ALPK3) | NM_020778 | 179.2 | 129.7 | 0.0420 | 0.83 |
| FARP1 | FERM, ARH/RhoGEF and pleckstrin domain protein 1(FARP1) | NM_005766 | 144 | 112.7 | 0.0420 | 0.86 |
| | | XM_945430 | 110.6 | 120.5 | 0.0420 | 1.22 |
| | | BE673312 | 167.8 | 97.6 | 0.0420 | 0.66 |
| ZEB1-AS1 | ZEB1 antisense RNA 1(ZEB1-AS1) | AK026657 | 112.3 | 91.7 | 0.0420 | 0.87 |
| FBXO28 | F-box protein 28(FBXO28) | NM_015176 | 118.8 | 135.7 | 0.0420 | 1.25 |

| | | | | | | |
|---|---|---|---|---|---|---|
| CAPN5 | calpain 5(CAPN5) | NM_004055 | 399.8 | 209.9 | 0.0430 | 0.61 |
| ARHGAP26 | Rho GTPase activating protein 26(ARHGAP26) | NM_015071 | 116.5 | 95.1 | 0.0430 | 0.88 |
| NSUN4 | NOP2/Sun RNA methyltransferase family member 4(NSUN4) | NM_199044 | 99.7 | 104.7 | 0.0430 | 1.14 |
| ATRAID | all-trans retinoic acid induced differentiation factor(ATRAID) | NM_080592 | 831.4 | 584.6 | 0.0430 | 0.62 |
| ATRAID | all-trans retinoic acid induced differentiation factor(ATRAID) | NM_080592 | 1464.5 | 799.2 | 0.0430 | 0.62 |
| | | XR_017229 | 353.9 | 188 | 0.0430 | 0.61 |
| | | DA453004 | 144 | 110.2 | 0.0430 | 0.86 |
| NPIPB13 | nuclear pore complex interacting protein family, member B13(NPIPB13) | NR_002555 | 209 | 141 | 0.0430 | 0.78 |
| | | XM_0017233520 | 111.9 | 112.7 | 0.0430 | 1.11 |
| | | AI633632 | 156.9 | 120.5 | 0.0430 | 0.86 |
| MT1X | metallothionein 1X(MT1X) | NM_005952 | 6772.5 | 2570 | 0.0430 | 0.44 |

| | | | | | | | | | |
|---|---|---|---|---|---|---|---|---|---|
| | | XR_038314 | 3375.1 | 4770.6 | 0.0430 | 1.35 | NM_001025767 | 5.06 | 85.80 |
| SMC4 | structural maintenance of chromosomes 4(SMC4) | NM_001002800 | 86.2 | 102.3 | 0.0430 | 1.29 | | | |
| RAD54L | RAD54-like (S. cerevisiae)(RAD54L) | NM_003579 | 95.5 | 104.6 | 0.0430 | 1.2 | NM_001037349 | 4.94 | 35.26 |
| ARL5B | ADP ribosylation factor like GTPase 5B(ARL5B) | NM_178815 | 92.4 | 98.2 | 0.0430 | 1.13 | NM_001037349 | 4.94 | 35.26 |
| CYP8B1 | cytochrome P450 family 8 subfamily B member 1(CYP8B1) | NM_004391 | 254.8 | 153.8 | 0.0430 | 0.67 | | | |
| | | XM_926112 | 143.2 | 161.8 | 0.0430 | 1.27 | | | |
| PSME1 | proteasome activator subunit 1(PSME1) | NM_176783 | 113.7 | 112 | 0.0430 | 1.1 | | | |
| MDC1 | mediator of DNA damage checkpoint 1(MDC1) | NM_014641 | 249.2 | 300.8 | 0.0430 | 1.34 | | | |
| | | BF194881 | 131.1 | 107.8 | 0.0430 | 0.87 | | | |
| SCRIB | scribbled planar cell polarity protein(SCRIB) | NM_182706 | 110.8 | 113.3 | 0.0430 | 1.14 | | | |

| | | | | | | | | | |
|---|---|---|---|---|---|---|---|---|---|
| TMEM128 | transmembrane protein 128(TMEM128) | NM_032927 | 119 | 100 | 0.0430 | 0.91 | | | |
| | | XR_038706 | 316.4 | 360.2 | 0.0430 | 1.3 | | | |
| SON | SON DNA binding protein(SON) | NM_138927 | 119.6 | 122.1 | 0.0430 | 1.13 | | | |
| SON | SON DNA binding protein(SON) | NM_138927 | 171.9 | 176 | 0.0430 | 1.13 | NM_001107368 | 10.94 | 1.12 |
| SON | SON DNA binding protein(SON) | NM_138927 | 1267.9 | 1422.7 | 0.0430 | 1.13 | | | |
| ZNF471 | zinc finger protein 471(ZNF471) | NM_020813 | 100 | 113.7 | 0.0430 | 1.23 | | | |
| ZNF471 | zinc finger protein 471(ZNF471) | NM_020813 | 135.5 | 131.7 | 0.0430 | 1.23 | | | |
| IPO11 | importin 11(IPO11) | NM_016338 | 227.6 | 242.9 | 0.0430 | 1.22 | | | |
| | | XM_001132362 | 108.5 | 91.4 | 0.0430 | 0.89 | | | |
| BIRC5 | baculoviral IAP repeat containing 5(BIRC5) | NM_001168 | 94.2 | 113.8 | 0.0440 | 1.31 | | | |
| LSM1 | LSM1 homolog, mRNA degradation associated(LSM1) | NM_014462 | 354.8 | 371 | 0.0440 | 1.17 | | | |

| | | | | | | |
|---|---|---|---|---|---|---|
| CSNK2B | casein kinase 2 beta(CSNK2B) | NM_001320 | 465.5 | 332.4 | 0.0440 | 0.82 |
| | | BF375676 | 104.8 | 114.8 | 0.0440 | 1.2 |
| JAG1 | jagged 1(JAG1) | NM_000214 | 132.6 | 132.6 | 0.0440 | 1.14 |
| DUSP27 | dual specificity phosphatase 27 (putative)(DUSP27) | NM_001080426 | 107.7 | 111.5 | 0.0440 | 1.13 |
| TSPAN17 | tetraspanin 17(TSPAN17) | NM_012171 | 158.7 | 169.5 | 0.0440 | 1.24 |
| SLC30A7 | solute carrier family 30 member 7(SLC30A7) | NM_133496 | 346.9 | 431.7 | 0.0440 | 1.43 |
| SP140 | SP140 nuclear body protein(SP140) | NM_001005176 | 81.4 | 99.1 | 0.0440 | 1.32 |
| | | DB344577 | 136 | 106.1 | 0.0440 | 0.86 |
| | | XM_001126647 | 112.3 | 123.9 | 0.0440 | 1.24 |
| | | XM_001126647 | 118.3 | 108.3 | 0.0440 | 1.24 |
| FER1L5 | fer-1 like family member 5(FER1L5) | NM_001077400 | 167.4 | 127.7 | 0.0440 | 0.87 |
| FER1L5 | fer-1 like family member 5(FER1L5) | NM_001077400 | 249.8 | 184.4 | 0.0440 | 0.87 |
| RWDD2A | RWD domain containing 2A(RWDD2A) | AJ420581 | 228.2 | 154.5 | 0.0440 | 0.78 |

| | | | | | | | | | |
|---|---|---|---|---|---|---|---|---|---|
| ZNF2 | zinc finger protein 2(ZNF2) | NM_021088 | 102.5 | 105.9 | 0.0440 | 1.12 | | | |
| NQO1 | NAD(P)H quinone dehydrogenase 1(NQO1) | NM_000903 | 141.9 | 100.9 | 0.0440 | 0.79 | | | |
| NQO1 | NAD(P)H quinone dehydrogenase 1(NQO1) | NM_000903 | 3501.7 | 2236.7 | 0.0440 | 0.79 | NM_017312 | 61.39 | 265.99 |
| | | BG911604 | 126.8 | 99.5 | 0.0440 | 0.84 | | | |
| LRCH3 | leucine rich repeats and calponin homology domain containing 3(LRCH3) | NM_032773 | 144.3 | 174.9 | 0.0440 | 1.35 | | | |
| PDE11A | phosphodiesterase 11A(PDE11A) | NM_016953 | 110.6 | 91.2 | 0.0440 | 0.87 | | | |
| SRSF5 | serine and arginine rich splicing factor 5(SRSF5) | NM_006925 | 1451.3 | 1528.5 | 0.0440 | 1.28 | | | |
| SRSF5 | serine and arginine rich splicing factor 5(SRSF5) | NM_006925 | 1464.5 | 1688 | 0.0440 | 1.28 | | | |
| MGAM | maltase-glucoamylase(MGAM) | NM_004668 | 117.3 | 91.8 | 0.0440 | 0.84 | | | |

| | | | | | | |
|---|---|---|---|---|---|---|
| IL2RG | interleukin 2 receptor subunit gamma(IL2RG) | NM_000206 | 94.1 | 107.7 | 0.0440 | 1.26 |
| | | XM_001724244 | 116.4 | 117.2 | 0.0440 | 1.14 |
| FAM120A | family with sequence similarity 120A(FAM120A) | NM_014612 | 1682.3 | 1799.4 | 0.0440 | 1.19 |
| ZNF524 | zinc finger protein 524(ZNF524) | NM_153219 | 151.3 | 160.5 | 0.0440 | 1.2 |
| MARS | methionyl-tRNA synthetase(MARS) | NM_004990 | 95 | 96.4 | 0.0440 | 1.08 |
| MT1IP | metallothionein 1I, pseudogene(MT1IP) | NM_175621 | 1855.3 | 667.1 | 0.0440 | 0.42 |
| ZC3H3 | zinc finger CCCH-type containing 3(ZC3H3) | NM_015117 | 177.4 | 182.4 | 0.0450 | 1.18 |
| TTC31 | tetratricopeptide repeat domain 31(TTC31) | NM_022492 | 209.4 | 235.6 | 0.0450 | 1.27 |
| | | XM_001723153 | 123.9 | 88.5 | 0.0450 | 0.78 |
| WDR4 | WD repeat domain 4(WDR4) | NM_033661 | 98 | 100.8 | 0.0450 | 1.1 |
| DERL2 | derlin 2(DERL2) | AK092218 | 142.3 | 112.5 | 0.0450 | 0.86 |
| | | AW136029 | 148.4 | 115.2 | 0.0450 | 0.87 |

| | | | | | | |
|---|---|---|---|---|---|---|
| dhx37 | DEAH-box helicase 37(DHX37) | NM_032656 | 166.2 | 201.6 | 0.0450 | 1.39 |
| st6gal1 | ST6 beta-galactoside alpha-2,6-sialyltransferase 1(ST6GAL1) | NM_003032 | 104.6 | 109.5 | 0.0450 | 0.78 |
| st6gal1 | ST6 beta-galactoside alpha-2,6-sialyltransferase 1(ST6GAL1) | NM_003032 | 167.9 | 116.2 | 0.0450 | 0.78 |
| | | XM_930029 | 348.7 | 229 | 0.0450 | 0.75 |
| | | XM_942439 | 189.3 | 140 | 0.0450 | 0.84 |
| ak5 | adenylate kinase 5(AK5) | NM_012093 | 124.3 | 95.3 | 0.0450 | 0.84 |
| | | NR_002214 | 147.6 | 110.4 | 0.0450 | 0.83 |
| AMPD3 | adenosine monophosphate deaminase 3(AMPD3) | NM_000480 | 137.6 | 111.8 | 0.0450 | 0.89 |
| | | XM_937706 | 375.2 | 242.9 | 0.0450 | 0.74 |
| SF3A3 | splicing factor 3a subunit 3(SF3A3) | NM_006802 | 817.1 | 906.7 | 0.0450 | 1.26 |
| | | XM_935575 | 104.3 | 112 | 0.0450 | 1.2 |

| | | | | | | |
|---|---|---|---|---|---|---|
| EIF2B4 | eukaryotic translation initiation factor 2B subunit delta(EIF2B4) | NM_172195 | 106.5 | 106.4 | 0.0450 | 1.1 |
| YTHDC1 | YTH domain containing 1(YTHDC1) | NM_001031732 | 103.1 | 102.1 | 0.0450 | 1.31 |
| YTHDC1 | YTH domain containing 1(YTHDC1) | NM_001031732 | 104 | 123.2 | 0.0450 | 1.31 |
| YTHDC1 | YTH domain containing 1(YTHDC1) | NM_001031732 | 729.1 | 833.2 | 0.0450 | 1.31 |
| TEX10 | testis expressed 10(TEX10) | NM_017746 | 125.4 | 145.9 | 0.0450 | 1.33 |
| MTNR1B | melatonin receptor 1B(MTNR1B) | NM_005959 | 95.3 | 102.3 | 0.0450 | 1.15 |
| PLIN5 | perilipin 5(PLIN5) | NM_001013706 | 372.8 | 494.9 | 0.0450 | 1.54 |
| SDHAF1 | succinate dehydrogenase complex assembly factor 1(SDHAF1) | NM_001042631 | 114.9 | 119.6 | 0.0460 | 1.16 |
| PI15 | peptidase inhibitor 15(PI15) | NM_015886 | 95.2 | 98.4 | 0.0460 | 1.11 |
| PI15 | peptidase inhibitor 15(PI15) | NM_015886 | 136.8 | 117 | 0.0460 | 1.11 |

| | | | | | | | | | |
|---|---|---|---|---|---|---|---|---|---|
| POU2F2 | POU class 2 homeobox 2(POU2F2) | NM_002698 | 91.4 | 104.4 | 0.0460 | 1.25 | NM_001107453 | 3.00 | 15.94 |
| KLHL7 | kelch like family member 7(KLHL7) | NM_001031710 | 112.5 | 115.2 | 0.0460 | 1.11 | | | |
| ARAP1 | ArfGAP with RhoGAP domain, ankyrin repeat and PH domain 1(ARAP1) | NM_001040118 | 108.1 | 117.7 | 0.0460 | 1.21 | | | |
| ARAP1 | ArfGAP with RhoGAP domain, ankyrin repeat and PH domain 1(ARAP1) | NM_001040118 | 114.3 | 118.6 | 0.0460 | 1.21 | | | |
| | | XM_001721348 | 107 | 115.9 | 0.0460 | 1.2 | | | |
| SOX30 | SRY-box 30(SOX30) | NM_178424 | 203.1 | 134.7 | 0.0460 | 0.76 | | | |
| PDX1 | pancreatic and duodenal homeobox 1(PDX1) | NM_000209 | 150.3 | 121.4 | 0.0460 | 0.9 | | | |
| RWDD2B | RWD domain containing 2B(RWDD2B) | NM_016940 | 299.1 | 185.6 | 0.0460 | 0.72 | | | |
| | | XM_942399 | 98.4 | 103.1 | 0.0460 | 1.15 | | | |
| IL17C | interleukin 17C(IL17C) | NM_013278 | 230.6 | 155.9 | 0.0460 | 0.78 | | | |

| | | | | | | |
|---|---|---|---|---|---|---|
| MAP1LC3A | microtubule associated protein 1 light chain 3 alpha(MAP1LC3A) | NM_181509 | 774.1 | 844.6 | 0.0460 | 1.23 |
| ZNF816 | zinc finger protein 816(ZNF816) | NM_001031665 | 107.7 | 111.6 | 0.0460 | 1.14 |
| | | XM_001717865 | 116.1 | 99.2 | 0.0460 | 0.93 |
| TAGLN3 | transgelin 3(TAGLN3) | NM_001008273 | 116.1 | 97.4 | 0.0460 | 0.91 |
| acap3 | ArfGAP with coiled-coil, ankyrin repeat and PH domains 3(ACAP3) | NM_030649 | 92.1 | 98.2 | 0.0460 | 1.14 |
| SNHG11 | small nucleolar RNA host gene 11(SNHG11) | NR_003239 | 383.3 | 257.4 | 0.0460 | 0.78 |
| MAMSTR | MEF2 activating motif and SAP domain containing transcriptional regulator(MAMSTR) | NM_182574 | 156.7 | 115.6 | 0.0460 | 0.83 |
| ITM2B | integral membrane protein 2B(ITM2B) | NM_021999 | 120.8 | 84.1 | 0.0460 | 0.75 |

| | | | | | | |
|---|---|---|---|---|---|---|
| ITM2B | integral membrane protein 2B(ITM2B) | NM_021999 | 4258 | 3405.8 | 0.0460 | 0.75 |
| ENTPD1 | ectonucleoside triphosphate diphosphohydrolase 1(ENTPD1) | NM_001098175 | 106.3 | 127.1 | 0.0460 | 1.35 |
| GEM | GTP binding protein overexpressed in skeletal muscle(GEM) | NM_181702 | 138.5 | 166.6 | 0.0460 | 1.36 |
| GEM | GTP binding protein overexpressed in skeletal muscle(GEM) | NM_181702 | 154.3 | 177 | 0.0460 | 1.36 |
| KLHL5 | kelch like family member 5(KLHL5) | NM_001007075 | 119.7 | 111.3 | 0.0460 | 0.9 |
| KLHL5 | kelch like family member 5(KLHL5) | NM_001007075 | 133.1 | 108.1 | 0.0460 | 0.9 |
| KLHL5 | kelch like family member 5(KLHL5) | NM_001007075 | 401.5 | 351.6 | 0.0460 | 0.9 |
| C19orf47 | chromosome 19 open reading frame 47(C19orf47) | NM_178830 | 153.4 | 119.6 | 0.0470 | 0.86 |

| | | | | | | | | | |
|---|---|---|---|---|---|---|---|---|---|
| MRPS11 | mitochondrial ribosomal protein S11(MRPS11) | NM_176805 | 249.3 | 172.2 | 0.0470 | 0.77 | | | |
| MRPS11 | mitochondrial ribosomal protein S11(MRPS11) | NM_176805 | 457.1 | 374.9 | 0.0470 | 0.77 | | | |
| | | XM_001132864 | 120.5 | 96 | 0.0470 | 0.86 | | | |
| | | XM_930715 | 127.4 | 101.4 | 0.0470 | 0.87 | | | |
| SEPT2 | septin 2(SEPT2) | NM_004404 | 1066.6 | 1278 | 0.0470 | 1.32 | | | |
| | | XM_001715568 | 126.2 | 122.7 | 0.0470 | 1.1 | | | |
| FAM86JP | family with sequence similarity 86, member A pseudogene(FAM86JP) | NR_024250 | 101.6 | 104.8 | 0.0470 | 1.14 | NM_030848 | 56.65 | 0.04 |
| CYTH3 | cytohesin 3(CYTH3) | NM_004227 | 311 | 125 | 0.0470 | 0.47 | | | |
| ttc32 | tetratricopeptide repeat domain 32(TTC32) | NM_001008237 | 181.9 | 199.3 | 0.0470 | 1.23 | | | |
| ttc32 | tetratricopeptide repeat domain 32(TTC32) | NM_001008237 | 221.6 | 195.1 | 0.0470 | 1.23 | | | |
| CHKA | choline kinase alpha(CHKA) | NM_212469 | 203.1 | 232.7 | 0.0470 | 1.31 | | | |

| | | | | | | |
|---|---|---|---|---|---|---|
| HNRNPM | heterogeneous nuclear ribonucleoprotein M(HNRNPM) | NM_031203 | 242.8 | 324.1 | 0.0470 | 1.49 |
| ABR | active BCR-related(ABR) | NM_001092 | 319.5 | 373.5 | 0.0470 | 1.33 |
| HOXA7 | homeobox A7(HOXA7) | NM_006896 | 175.5 | 129.8 | 0.0470 | 0.83 |
| EXOC3L1 | exocyst complex component 3 like 1(EXOC3L1) | NM_178516 | 102.5 | 113 | 0.0470 | 1.21 |
| NDUFAF3 | NADH:ubiquinone oxidoreductase complex assembly factor 3(NDUFAF3) | NM_199074 | 122.9 | 98.9 | 0.0470 | 0.89 |
| clstn3 | calsyntenin 3(CLSTN3) | NM_014718 | 120.6 | 100.9 | 0.0470 | 0.9 |
| EDNRA | endothelin receptor type A(EDNRA) | NM_001957 | 85.7 | 93.3 | 0.0470 | 1.16 |
| | | XM_001716912 | 90.4 | 96.6 | 0.0470 | 1.14 |
| PHYKPL | 5-phosphohydroxy-L-lysine phospho-lyase(PHYKPL) | NM_153373 | 218.2 | 147.5 | 0.0470 | 0.78 |
| | | XM_929696 | 104.1 | 96.7 | 0.0470 | 0.77 |
| | | XM_929696 | 157.3 | 108 | 0.0470 | 0.77 |

| | | | | | | | | | |
|---|---|---|---|---|---|---|---|---|---|
| | | XM_001128702 | 147.2 | 114.5 | 0.0470 | 0.87 | | | |
| ZMYM6 | zinc finger MYM-type containing 6(ZMYM6) | NM_007167 | 124.4 | 143.4 | 0.0470 | 1.29 | | | |
| | | DB369246 | 111.4 | 112.2 | 0.0470 | 1.11 | | | |
| FBXO38 | F-box protein 38(FBXO38) | NM_030793 | 165.1 | 176.1 | 0.0470 | 1.21 | NM_017258 | 66.46 | 359.19 |
| FBXO38 | F-box protein 38(FBXO38) | NM_030793 | 181.3 | 177.5 | 0.0470 | 1.21 | | | |
| | | XM_001125904 | 126.9 | 135.8 | 0.0470 | 1.22 | | | |
| ZXDB | zinc finger, X-linked, duplicated B(ZXDB) | NM_007157 | 174.1 | 132.8 | 0.0470 | 0.87 | | | |
| zbtb18 | zinc finger and BTB domain containing 18(ZBTB18) | NM_205768 | 289.4 | 185.5 | 0.0470 | 0.75 | | | |
| | | XM_936103 | 141.5 | 158.8 | 0.0470 | 1.28 | | | |
| WDR27 | WD repeat domain 27(WDR27) | NM_182552 | 210.5 | 153.9 | 0.0470 | 0.83 | | | |
| SNX29P2 | sorting nexin 29 pseudogene 2(SNX29P2) | NR_002939 | 401.9 | 263.8 | 0.0470 | 1.89 | | | |
| SNX29P2 | sorting nexin 29 pseudogene 2(SNX29P2) | NR_002939 | 800.3 | 1275 | 0.0470 | 1.89 | | | |

| | | | | | | |
|---|---|---|---|---|---|---|
| nomo1 | NODAL modulator 1(NOMO1) | NM_014287 | 284.9 | 183.8 | 0.0470 | 0.76 |
| ANAPC11 | anaphase promoting complex subunit 11(ANAPC11) | NM_001002244 | 173.5 | 128.4 | 0.0470 | 0.84 |
| RNF135 | ring finger protein 135(RNF135) | NM_197939 | 121.6 | 131.1 | 0.0470 | 1.23 |
| | | XM_001124651 | 129 | 104.1 | 0.0480 | 0.88 |
| BPTF | bromodomain PHD finger transcription factor(BPTF) | NM_182641 | 167.9 | 123.3 | 0.0480 | 0.83 |
| TENM2 | teneurin transmembrane protein 2(TENM2) | NM_001080428 | 109.5 | 108.7 | 0.0480 | 1.09 |
| CTPS1 | CTP synthase 1(CTPS1) | NM_001905 | 110.3 | 148.1 | 0.0480 | 1.52 |
| MEIOB | meiosis specific with OB domains(MEIOB) | NM_152764 | 118.1 | 96 | 0.0480 | 0.85 |
| USP49 | ubiquitin specific peptidase 49(USP49) | NM_018561 | 1581.2 | 2265.6 | 0.0480 | 1.52 |
| PARP8 | poly(ADP-ribose) polymerase family member 8(PARP8) | NM_024615 | 103.5 | 123.3 | 0.0480 | 1.32 |

| | | | | | | |
|---|---|---|---|---|---|---|
| eif2ak2 | eukaryotic translation initiation factor 2 alpha kinase 2(EIF2AK2) | NM_002759 | 110.9 | 121.9 | 0.0480 | 1.22 |
| ZNF394 | zinc finger protein 394(ZNF394) | NM_032164 | 1982.7 | 2445.7 | 0.0480 | 1.31 |
| | | BG201242 | 103.3 | 105.5 | 0.0480 | 1.11 |
| EXTL1 | exostosin like glycosyltransferase 1(EXTL1) | NM_004455 | 116 | 115.9 | 0.0480 | 1.12 |
| RAB3C | RAB3C, member RAS oncogene family(RAB3C) | NM_138453 | 100.1 | 86.4 | 0.0480 | 0.88 |
| RLF | rearranged L-myc fusion(RLF) | NM_012421 | 142.3 | 157.2 | 0.0480 | 1.24 |
| | | XM_001720625 | 3030 | 4299.8 | 0.0480 | 1.35 |
| | | XM_934672 | 106.9 | 104.5 | 0.0480 | 1.06 |
| RPF2 | ribosome production factor 2 homolog(RPF2) | NM_032194 | 158 | 171.6 | 0.0480 | 1.24 |
| | | NR_003253 | 168 | 128.1 | 0.0480 | 0.86 |
| FSIP1 | fibrous sheath interacting protein 1(FSIP1) | NM_152597 | 135.3 | 106.7 | 0.0480 | 0.87 |
| | | XM_940764 | 106.1 | 88.3 | 0.0480 | 0.86 |

| | | | | | | |
|---|---|---|---|---|---|---|
| ZNF622 | zinc finger protein 622(ZNF622) | NM_033414 | 410.6 | 325.4 | 0.0490 | 0.89 |
| CHRNA6 | cholinergic receptor nicotinic alpha 6 subunit(CHRNA6) | NM_004198 | 101.6 | 106.8 | 0.0490 | 1.13 |
| ZBTB32 | zinc finger and BTB domain containing 32(ZBTB32) | NM_014383 | 89.9 | 98.5 | 0.0490 | 1.17 |
| CBFB | core-binding factor beta subunit(CBFB) | NM_001755 | 100.2 | 106.6 | 0.0490 | 1.15 |
| PCDHB16 | protocadherin beta 16(PCDHB16) | NM_020957 | 116 | 93.9 | 0.0490 | 0.85 |
| NLGN4X | neuroligin 4, X-linked(NLGN4X) | NM_020742 | 85.6 | 97.6 | 0.0490 | 1.22 |
| COL20A1 | collagen type XX alpha 1 chain(COL20A1) | NM_020882 | 381.6 | 203.5 | 0.0490 | 0.63 |
| PIAS4 | protein inhibitor of activated STAT 4(PIAS4) | NM_015897 | 467.2 | 510.9 | 0.0490 | 1.24 |
| NOP56 | NOP56 ribonucleoprotein(NOP56) | NM_006392 | 112 | 111.6 | 0.0490 | 1.73 |
| NOP56 | NOP56 ribonucleoprotein(NOP56) | NM_006392 | 343.3 | 532.7 | 0.0490 | 1.73 |

| | | | | | | |
|---|---|---|---|---|---|---|
| | | XR_042317 | 119.9 | 119.6 | 0.0490 | 1.13 |
| FBXO4 | F-box protein 4(FBXO4) | NM_033484 | 90.8 | 99.3 | 0.0490 | 1.18 |
| Krtap5-1 | keratin associated protein 5-1(KRTAP5-1) | NM_001005922 | 118.6 | 100.6 | 0.0490 | 0.93 |
| ITGA2 | integrin subunit alpha 2(ITGA2) | NM_002203 | 99.4 | 104.4 | 0.0490 | 1.13 |
| CPVL | carboxypeptidase, vitellogenic like(CPVL) | NM_019029 | 90.5 | 133.4 | 0.0490 | 1.64 |
| XCL1 | X-C motif chemokine ligand 1(XCL1) | NM_002995 | 106.1 | 90.2 | 0.0490 | 0.9 |
| GUCA1A | guanylate cyclase activator 1A(GUCA1A) | NM_000409 | 129.1 | 105.2 | 0.0490 | 0.87 |
| TSC22D2 | TSC22 domain family member 2(TSC22D2) | NM_014779 | 198.9 | 208 | 0.0490 | 1.35 |
| TSC22D2 | TSC22 domain family member 2(TSC22D2) | NM_014779 | 406.5 | 483.1 | 0.0490 | 1.35 |
| | | XR_039891 | 102 | 101.8 | 0.0490 | 1.08 |
| KRTAP10-8 | keratin associated protein 10-8(KRTAP10-8) | NM_198695 | 100.4 | 100.9 | 0.0490 | 1.1 |

| | | | | | | |
|---|---|---|---|---|---|---|
| CCDC90B | coiled-coil domain containing 90B(CCDC90B) | NM_021825 | 101.4 | 105.8 | 0.0490 | 1.13 |
| DNAJB14 | DnaJ heat shock protein family (Hsp40) member B14(DNAJB14) | NM_001031723 | 98.2 | 117.2 | 0.0490 | 1.2 |
| DNAJB14 | DnaJ heat shock protein family (Hsp40) member B14(DNAJB14) | NM_001031723 | 110.8 | 121.7 | 0.0490 | 1.2 |
| | | XM_943600 | 89.2 | 97.3 | 0.0490 | 1.17 |
| SPRED1 | sprouty related EVH1 domain containing 1(SPRED1) | NM_152594 | 371.3 | 439 | 0.0490 | 1.31 |
| UPK3BL | uroplakin 3B-like(UPK3BL) | NM_001114403 | 116.8 | 91.8 | 0.0490 | 0.85 |
| ZNF830 | zinc finger protein 830(ZNF830) | NM_052857 | 134.4 | 143.2 | 0.0490 | 1.19 |
| STAMBP | STAM binding protein(STAMBP) | NM_006463 | 126.4 | 132 | 0.0490 | 1.19 |
| STAMBP | STAM binding protein(STAMBP) | NM_006463 | 148.9 | 122.4 | 0.0490 | 1.19 |
| ALAD | aminolevulinate dehydratase(ALAD) | NM_001003945 | 114.6 | 103.1 | 0.0500 | 0.69 |

| | | | | | | |
|---|---|---|---|---|---|---|
| ALAD | aminolevulinate dehydratase(ALAD) | NM_001003945 | 370.1 | 223 | 0.0500 | 0.69 |
| | | XR_040201 | 173.3 | 114.2 | 0.0500 | 0.75 |
| CELF1 | CUGBP, Elav-like family member 1(CELF1) | NM_198700 | 141.3 | 151.3 | 0.0500 | 1.19 |
| ZNF621 | zinc finger protein 621(ZNF621) | NM_001098414 | 109.6 | 116.4 | 0.0500 | 1.19 |
| CHM | CHM, Rab escort protein 1(CHM) | NM_000390 | 118 | 122.3 | 0.0500 | 1.17 |
| BTLA | B and T lymphocyte associated(BTLA) | NM_181780 | 86.4 | 103 | 0.0500 | 1.29 |
| BTLA | B and T lymphocyte associated(BTLA) | NM_181780 | 121.9 | 140.4 | 0.0500 | 1.29 |
| | | AA421413 | 154 | 118.7 | 0.0500 | 0.87 |
| | | XM_927848 | 123 | 101.2 | 0.0500 | 0.9 |
| RAB30 | RAB30, member RAS oncogene family(RAB30) | NM_014488 | 115.6 | 252.8 | 0.0500 | 2.42 |
| MT1A | metallothionein 1A(MT1A) | NM_005946 | 11240.4 | 4350.9 | 0.0500 | 0.45 |
| RPL36 | ribosomal protein L36(RPL36) | NM_015414 | 95.5 | 101.3 | 0.0500 | 1.14 |
| FLJ35816 | FLJ35816 protein(FLJ35816) | NM_207489 | 169.5 | 121.2 | 0.0500 | 0.8 |

| | | | | | | |
|---|---|---|---|---|---|---|
| TXLNG | taxilin gamma(TXLNG) | NM_018360 | 211.3 | 232.1 | 0.0500 | 1.23 |
| mocs2 | molybdenum cofactor synthesis 2(MOCS2) | NM_176806 | 148.7 | 104.2 | 0.0500 | 0.75 |
| | | XM_940734 | 160.2 | 110.1 | 0.0500 | 0.77 |
| CYP17A1 | cytochrome P450 family 17 subfamily A member 1(CYP17A1) | NM_000102 | 151.7 | 113.4 | 0.0500 | 0.84 |
| GPKOW | G-patch domain and KOW motifs(GPKOW) | NM_015698 | 122.3 | 130.7 | 0.0500 | 1.21 |
| PURB | purine rich element binding protein B(PURB) | NM_033224 | 536.6 | 636 | 0.0500 | 1.35 |
| BRSK1 | BR serine/threonine kinase 1(BRSK1) | NM_032430 | 146.6 | 154.8 | 0.0500 | 1.22 |
| CCNT2 | cyclin T2(CCNT2) | NM_001241 | 101.4 | 109.2 | 0.0500 | 1.17 |
| MVP | major vault protein(MVP) | NM_005115 | 200.5 | 210.9 | 0.0500 | 1.2 |
| DGKQ | diacylglycerol kinase theta(DGKQ) | NM_001347 | 104.3 | 110.5 | 0.0500 | 1.16 |
| | | XM_940726 | 110.2 | 91.5 | 0.0500 | 0.88 |

| | | | | | | |
|---|---|---|---|---|---|---|
| | | XM_940726 | 136.9 | 109.4 | 0.0500 | 0.88 |
| HM13 | histocompatibility minor 13(HM13) | NM_178580 | 132.4 | 107.8 | 0.0500 | 0.87 |
| HM13 | histocompatibility minor 13(HM13) | NM_178580 | 175 | 143.4 | 0.0500 | 0.87 |
| TNPO3 | transportin 3(TNPO3) | NM_012470 | 279.4 | 297.4 | 0.0500 | 1.2 |
| AGFG1 | ArfGAP with FG repeats 1(AGFG1) | NM_004504 | 160.9 | 183.1 | 0.0500 | 1.28 |
| NRAS | neuroblastoma RAS viral oncogene homolog(NRAS) | NM_002524 | 109 | 114.3 | 0.0500 | 1.15 |

**Supplementary Table 2: Pathways**

| | -log(p-value) | Ratio | End of Pathway | Direction |
|---|---|---|---|---|
| **Female IUGR Chow versus Female CON Chow** | | | | |
| **Oxidative Phosphorylation** | **4.330** | **0.092** | **Y** | **D** |
| **Cell Cycle Control of Chromosomal Replication** | **4.030** | **0.125** | **Y** | **U** |
| **LXR/RXR Activation** | **3.260** | **0.074** | **y** | **D** |
| **Mitochondrial Function** | **2.760** | **0.059** | **Y** | **D** |
| Sirtuin Signaling Pathway | 4.820 | 0.062 | N | |
| FXR/RXR Activation | 4.500 | 0.087 | N | |
| Iron homeostasis signaling pathway | 3.500 | 0.073 | N | |
| EIF2 Signaling | 3.300 | 0.057 | N | |
| Polyamine Regulation in Colon Cancer | 3.130 | 0.182 | N | |
| Extrinsic Prothrombin Activation Pathway | 2.490 | 0.188 | N | |
| Clathrin-mediated Endocytosis Signaling | 2.140 | 0.048 | N | |

**Male IUGR Chow versus Male CON Chow**

| | | | | |
|---|---|---|---|---|
| **IGF-1 Signaling** | **3.540** | **0.133** | **Y** | **D** |
| **PI3K/AKT Signaling** | **3.390** | **0.124** | **Y** | **D** |
| **DNA Double-Strand Break Repair by Non-Homologous End Joining** | **2.460** | **0.286** | **Y** | **U** |
| **Role of BRCA1 in DNA Damage Response** | **2.350** | **0.125** | **Y** | **U** |
| **Toll-like Receptor Signaling** | **2.010** | **0.118** | **Y** | **U** |
| PTEN Signaling | 4.150 | 0.138 | N | |
| Amyloid Processing | 3.920 | 0.196 | N | |
| Phosphatidylethanolamine Biosynthesis II | 3.280 | 0.444 | N | |
| p53 Signaling | 3.030 | 0.124 | N | |
| ERK5 Signaling | 2.830 | 0.145 | N | |
| PDGF Signaling | 2.640 | 0.122 | N | |
| 14-3-3-mediated Signaling | 2.600 | 0.108 | N | |
| Protein Ubiquitination Pathway | 2.590 | 0.089 | N | |
| FAK Signaling | 2.320 | 0.112 | N | |

| | | | |
|---|---|---|---|
| p70S6K Signaling | 2.170 | 0.100 | N |
| Endometrial Cancer Signaling | 2.170 | 0.125 | N |
| IL-2 Signaling | 2.170 | 0.125 | N |
| Integrin Signaling | 2.040 | 0.086 | N |
| Insulin Receptor Signaling | 1.990 | 0.095 | N |

**Female CON HFCS Diet versus Chow Diet**

| | | | | |
|---|---|---|---|---|
| **Hepatic Fibrosis / Hepatic Stellate Cell Activation** | **2.59** | **0.0374** | **Y** | **U** |
| **Acute Phase Response Signaling** | **2.09** | **0.0341** | **Y** | **D** |
| LXR/RXR Activation | 6.47 | 0.0826 | N | |
| FXR/RXR Activation | 5.35 | 0.0714 | N | |
| GP6 Signaling Pathway | 5.13 | 0.0672 | N | |
| Atherosclerosis Signaling | 4.42 | 0.063 | N | |
| Intrinsic Prothrombin Activation Pathway | 4.25 | 0.119 | N | |
| Apelin Liver Signaling Pathway | 2.68 | 0.115 | N | |

**Female IUGR HFCS Diet versus Chow Diet**

| | | | | |
|---|---|---|---|---|
| **EIF2 Signaling** | **13** | **0.115** | **Y** | **D** |
| **mTOR Signaling** | **8.86** | **0.0971** | **Y** | **D** |
| **Regulation of eIF4 and p70S6K Signaling** | **8.09** | **0.104** | **Y** | **D** |
| **PI3K/AKT Signaling** | **3.19** | **0.0692** | **Y** | **D** |
| **p70S6K Signaling** | **2.41** | **0.058** | **Y** | **D** |
| **Gap Junction Signaling** | **2.4** | **0.0498** | **Y** | **D** |
| **PTEN Signaling** | **2.11** | **0.0565** | **Y** | **D** |
| Adrenomedullin signaling pathway | 2.94 | 0.055 | N | |
| D-myo-inositol (1,4,5)-Trisphosphate Biosynthesis | 2.86 | 0.148 | N | |
| ERK5 Signaling | 2.73 | 0.0845 | N | |
| DNA damage-induced 14-3-3σ Signaling | 2.33 | 0.158 | N | |
| AMPK Signaling | 2.19 | 0.0463 | N | |
| RhoA Signaling | 2.11 | 0.0565 | N | |
| Endocannabinoid Developing Neuron Pathway | 2.06 | 0.0551 | N | |
| **Male CON HFCS Diet versus Chow Diet** | | | | |
| **Integrin Signaling** | **3.63** | **0.0731** | **Y** | **U** |
| **Actin Cytoskeleton Signaling** | **3.33** | **0.0687** | **Y** | **U** |
| **PTEN Signaling** | **2.79** | **0.0806** | **Y** | **U** |

| | | | | |
|---|---|---|---|---|
| **Cholesterol Biosynthesis I** | **2.37** | **0.231** | **Y** | **U** |
| **Cholesterol Biosynthesis II (via 24,25-dihydrolanosterol)** | **2.37** | **0.231** | **Y** | **U** |
| **Cholesterol Biosynthesis III (via Desmosterol)** | **2.37** | **0.231** | **Y** | **U** |
| **ILK Signaling** | **2.22** | **0.0609** | **Y** | **U** |
| **FAK Signaling** | **2.19** | **0.0762** | **Y** | **U** |
| Amyloid Processing | 2.68 | 0.118 | N | |
| Cellular Effects of Sildenafil (Viagra) | 2.62 | 0.0763 | N | |
| NRF2-mediated Oxidative Stress Response | 2.6 | 0.065 | N | |
| Adrenomedullin signaling pathway | 2.6 | 0.065 | N | |
| Germ Cell-Sertoli Cell Junction Signaling | 2.55 | 0.067 | N | |
| Corticotropin Releasing Hormone Signaling | 2.43 | 0.0719 | N | |
| RAR Activation | 2.34 | 0.0632 | N | |
| Serotonin Receptor Signaling | 2.29 | 0.116 | N | |
| Melanocyte Development and Pigmentation Signaling | 2.24 | 0.0777 | N | |
| Wnt/β-catenin Signaling | 2.23 | 0.064 | N | |

| | | | | |
|---|---|---|---|---|
| IL-8 Signaling | 2.12 | 0.0591 | N | |
| Molecular Mechanisms of Cancer | 2.07 | 0.0482 | N | |
| p53 Signaling | 2.05 | 0.0721 | N | |
| IGF-1 Signaling | 2.05 | 0.0721 | N | |
| Regulation of Actin-based Motility by Rho | 2.02 | 0.0778 | N | |
| **Male IUGR HFCS Diet versus Chow Diet** | | | | |
| **Mitochondrial Function** | **2.44** | **0.076** | **Y** | **U** |
| **Assembly of RNA Polymerase I Complex** | **2.23** | **0.25** | **Y** | **D** |
| RAR Activation | 2.46 | 0.0737 | N | |
| Protein Ubiquitination Pathway | 2.25 | 0.0642 | N | |
| CDK5 Signaling | 2.17 | 0.0857 | N | |

**Supplementary Table 2: Pathways**

| | Molecules |
|---|---|
| **Female IUGR Chow versus Female CON Chow** | |
| **Oxidative Phosphorylation** | **SDHA,ATP5PF,NDUFB11,ATP5F1D,NDUFA11,UQCRFS1,NDUFB8,COX7A1,NDUFA1,COX4I1** |
| **Cell Cycle Control of Chromosomal Replication** | **CDK12,CDK18,CDC45,CDC6,RPA1,MCM7,ORC1** |
| **LXR/RXR Activation** | **APOE,HPX,TTR,ALB,SREBF1,APOA2,GC,FGA,PON3** |
| **Mitochondrial Function** | **SDHA,ATP5PF,NDUFB11,ATP5F1D,NDUFA11,UQCRFS1,NDUFB8,COX7A1,NDUFA1,COX4I1** |
| Sirtuin Signaling Pathway | ATP5PF,PPARA,SDHA,SLC25A4,ATP5F1D,TIMM9,ATG4B,GLUD1,NDUFB8,TIMM23,NDUFA1,PFKM,NDUFB11,GADD45A,SREBF1,NDUFA11,UQCRFS1,GOT2 |
| FXR/RXR Activation | PPARA,APOE,HPX,TTR,ALB,SREBF1,APOA2,AKT3,GC,FGA,PON3 |
| Iron homeostasis signaling pathway | ATP6V1E1,HAMP,HPX,PDGFA,ATP6V0D1,CIAO2B,CIAO2A,ATP6V0E1,PDGFB,SKP2 |
| EIF2 Signaling | RPS6,RPL32,RPL14,RPS15,EIF1,RPS3A,SREBF1,RPLP2,AKT3,RPS21,PPP1CA,RPL36,EIF3M |
| Polyamine Regulation in Colon Cancer | PSMF1,MXD1,PSMB11,ODC1 |
| Extrinsic Prothrombin Activation Pathway | SERPINC1,FGA,FGG |
| Clathrin-mediated Endocytosis Signaling | APOE,AP2M1,ALB,ACTR3,FGF18,PDGFA,APOA2,MDM2,UBC,PDGFB |

| | |
|---|---|
| **Male IUGR Chow versus Male CON Chow** | |
| **IGF-1 Signaling** | **RAP2B,NEDD4,CSNK2A1,NRAS,CTGF,MAP2K2,YWHAE,YWHAB,PIK3R1,FOXO3,PIK3R6,PDPK1,SOCS4,CSNK2B,TLR9** |
| **PI3K/AKT Signaling** | **RAP2B,TSC1,NRAS,YWHAE,YWHAB,PIK3R1,ITGA2,PDPK1,BCL2,SYNJ1,MAP2K2,LIMS1,FOXO3,INPP5K,GSK3B,ITGA4** |
| **DNA Double-Strand Break Repair by Non-Homologous End Joining** | **PRKDC,WRN,XRCC5,LIG3** |
| **Role of BRCA1 in DNA Damage Response** | **FANCE,PBRM1,BRD7,POU2F1,RFC4,FANCG,BARD1,MDC1,UIMC1,SMARCC2** |
| **Toll-like Receptor Signaling** | **TLR2,TLR10,TICAM1,TLR8,MAP3K7,EIF2AK2,MAPK13,TLR9,UBC** |
| PTEN Signaling | RAP2B,CSNK2A1,NRAS,PIK3R1,ITGA2,FLT4,PDPK1,BCL2,SYNJ1,MAP2K2,SIRT6,NGFR,FOXO3,GSK3B,CSNK2B,INPP5K,ITGA4 |
| Amyloid Processing | CSNK1E,CAPN5,CSNK2A1,PSENEN,APH1A,CSNK1A1,GSK3B,CSNK2B,MAPK13,CDK5R1 |
| Phosphatidylethanolamine Biosynthesis II | ETNK2,CEPT1,PCYT2,CHKA |
| p53 Signaling | PRKDC,MED1,PLAGL1,PIK3R1,TNFRSF10B,TLR9,BIRC5,TP53BP2,BCL2,CASP6,PIK3R6,ADGRB1,GSK3B,PML |
| ERK5 Signaling | RAP2B,NRAS,YWHAE,YWHAB,CREB1,FOXO3,GNAQ,MEF2A,MEF2C,WNK1 |
| PDGF Signaling | RAP2B,CSNK2A1,NRAS,MAP2K2,SYNJ1,PDGFA,PIK3R1,PIK3R6,EIF2AK2,CSNK2B,INPP5K,TLR9 |
| 14-3-3-mediated Signaling | RAP2B,TSC1,TUBB3,NRAS,YWHAE,YWHAB,PIK3R1,NOTUM,TLR9,TUBB2B,SRPK2,MAP2K2,PIK3R6,TUBB4A,GSK3B |
| Protein Ubiquitination Pathway | NEDD4,PSMD9,HSPA9,TRAP1,UBE2Z,FBXW7,USP19,USP1,THOP1,UBE2O,DNAJB14,USP33,SKP2,USP3,PSME1,UBE2D2,USP16,UBE2E3,ANAPC11,UBC,USP49,UBE2D3,UBE2Q2,BIRC2 |
| FAK Signaling | CAPN5,RAP2B,NRAS,ARHGAP26,MAP2K2,ARHGEF7,PIK3R1,ITGA2,PIK3R6,PDPK1,TLR9,ITGA4 |

| | |
|---|---|
| p70S6K Signaling | RAP2B,IL2RG,NRAS,YWHAE,F2R,YWHAB,EEF2,PIK3R1,NOTUM,GNAQ,PDPK1,TLR9,MAP2K2,PIK3R6 |
| Endometrial Cancer Signaling | RAP2B,NRAS,MAP2K2,PIK3R1,FOXO3,PIK3R6,PDPK1,GSK3B,TLR9 |
| IL-2 Signaling | RAP2B,CSNK2A1,IL2RG,NRAS,MAP2K2,PIK3R1,PIK3R6,CSNK2B,TLR9 |
| Integrin Signaling | RAP2B,CAPN5,RAPGEF1,ARHGAP26,NRAS,ARF1,ARHGEF7,PIK3R1,ITGA2,MYLK,TLR9,PPP1R12B,MAP2K2,RND3,LIMS1,PIK3R6,GSK3B,ITGB4,ITGA4 |
| Insulin Receptor Signaling | RAP2B,RAPGEF1,TSC1,NRAS,EIF2B4,PIK3R1,PDPK1,TLR9,SYNJ1,MAP2K2,FOXO3,PIK3R6,INPP5K,GSK3B |
| **Female CON HFCS Diet versus Chow Diet** | |
| **Hepatic Fibrosis / Hepatic Stellate Cell Activation** | **COL1A2,COL5A1,COL1A1,COL6A2,COL6A3,COL6A5,COL3A1** |
| **Acute Phase Response Signaling** | **HAMP,HPX,ALB,ORM1,FGB,FGG** |
| LXR/RXR Activation | SCD,HPX,ALB,ORM1,CD36,FASN,PCYOX1,PLTP,GC,PON3 |
| FXR/RXR Activation | PPARG,HPX,ALB,ORM1,FASN,PCYOX1,PLTP,GC,PON3 |
| GP6 Signaling Pathway | COL1A2,COL5A1,COL1A1,COL6A3,COL6A2,FGB,COL6A5,FGG,COL3A1 |
| Atherosclerosis Signaling | COL1A2,COL1A1,ALB,PLA2G2E,ORM1,CD36,PCYOX1,COL3A1 |
| Intrinsic Prothrombin Activation Pathway | COL1A2,COL1A1,FGB,FGG,COL3A1 |
| Apelin Liver Signaling Pathway | COL1A2,COL1A1,COL3A1 |
| **Female IUGR HFCS Diet versus Chow Diet** | |

| | |
|---|---|
| **EIF2 Signaling** | **EIF2B4,RPS3A,PDPK1,RPL7A,RPS4X,RPL7,BCL2,RPS7,PPP1CC,AKT1,EIF4G2,AKT3,RPL36,RPL36AL,ACTB,RPS8,RPS10,RPL23,RPS21,RPL10A,EIF3M,RPS15,RPS27A,RPS15A,RPL6,RPS14** |
| **mTOR Signaling** | **RHEB,RPS3A,STK11,RPS8,RPS10,PDPK1,RPS21,RPS4X,EIF3M,RPS7,RPS15,AKT1,EIF4G2,PTPA,AKT3,RPS27A,RPS15A,GNB1L,EIF4B,RPS14** |
| **Regulation of eIF4 and p70S6K Signaling** | **EIF2B4,RPS3A,RPS8,RPS10,PDPK1,RPS21,RPS4X,EIF3M,RPS7,RPS15,AKT1,EIF4G2,PTPA,AKT3,RPS27A,RPS15A,RPS14** |
| **PI3K/AKT Signaling** | **RHEB,AKT1,YWHAH,PTPA,AKT3,PDPK1,HSP90AA1,JAK3,BCL2** |
| **p70S6K Signaling** | **PLCD1,AKT1,YWHAH,PTPA,EEF2,PDPK1,AKT3,PLCD4** |
| **Gap Junction Signaling** | **PLCD1,TUBA1A,AKT1,SP3,ACTB,CSNK1G3,AKT3,MAPK7,PLCD4,MAP2K5** |
| **PTEN Signaling** | **AKT1,YWHAH,FLT4,AKT3,PDPK1,FASLG,BCL2** |
| Adrenomedullin signaling pathway | PLCD1,CRCP,IL18,AKT1,MAX,AKT3,MAPK7,PLCD4,RAMP2,MAP2K5,BCL2 |
| D-myo-inositol (1,4,5)-Trisphosphate Biosynthesis | PLCD1,PIP4K2B,PLCD4,PI4KA |
| ERK5 Signaling | IL6ST,AKT1,YWHAH,CREB1,MAPK7,MAP2K5 |
| DNA damage-induced 14-3-3σ Signaling | AKT1,RAD9A,AKT3 |
| AMPK Signaling | AKT1,PTPA,EEF2,ACTB,STK11,CREB1,AKT3,PDPK1,SMARCD1,GNB1L |
| RhoA Signaling | NGEF,ACTR3,ACTB,BAIAP2,PIP4K2B,ARHGAP1,PI4KA |
| Endocannabinoid Developing Neuron Pathway | AKT1,CREB1,AKT3,MAPK7,GNG5,GNB1L,MAP2K5 |
| **Male CON HFCS Diet versus Chow Diet** | |
| **Integrin Signaling** | **ITGB1,ARHGAP26,MPRIP,ILK,MYLK,CRK,RAC3,MYL9,PTK2,CAPNS1,MAP2K2,RHOT1,TSPAN1,FGFR4,PFN2,VCL** |
| **Actin Cytoskeleton Signaling** | **PTK2,MYL9,ITGB1,MYH10,MPRIP,MAP2K2,F2R,PDGFA,FGFR4,FGF12,PFN2,MYLK,CRK,VCL,RAC3,FGF13** |
| **PTEN Signaling** | **ITGB1,PTK2,CSNK2A1,SHARPIN,MAP2K2,FGFR4,FLT4,FOXO3,ILK,RAC3** |

| | |
|---|---|
| **Cholesterol Biosynthesis I** | **SQLE,DHCR7,MSMO1** |
| **Cholesterol Biosynthesis II (via 24,25-dihydrolanosterol)** | **SQLE,DHCR7,MSMO1** |
| **Cholesterol Biosynthesis III (via Desmosterol)** | **SQLE,DHCR7,MSMO1** |
| **ILK Signaling** | **ITGB1,MYL9,PTK2,FOS,MYH10,RHOT1,FGFR4,PPP2R2B,ILK,VCL,PTGS2,HIF1A** |
| **FAK Signaling** | **ITGB1,PTK2,ARHGAP26,CAPNS1,MAP2K2,FGFR4,CRK,VCL** |
| Amyloid Processing | CSNK1E,CSNK2A1,CAPNS1,PRKAR2A,APP,PSEN1 |
| Cellular Effects of Sildenafil (Viagra) | MYL9,CALM1 (includes others),MYH10,ADCY9,MPRIP,ADCY6,PRKAR2A,MYLK,KCNH2,GUCY1B1 |
| NRF2-mediated Oxidative Stress Response | FOS,AKR7A3,MAP2K2,FGFR4,GSTM4,SLC35A2,JUND,DNAJC10,DNAJB6,UBE2E3,JUNB,FKBP5,ABCC4 |
| Adrenomedullin signaling pathway | PTK2,ADCY9,FOS,CALM1 (includes others),MAP2K2,FGFR4,ADCY6,PRKAR2A,MYLK,CFH,HIF1A,KCNH2,GUCY1B1 |
| Germ Cell-Sertoli Cell Junction Signaling | ITGB1,PTK2,TUBB6,MAP2K2,RHOT1,FGFR4,ILK,TUBB4A,VCL,RAC3,MAP3K3,CTNND1 |
| Corticotropin Releasing Hormone Signaling | FOS,CALM1 (includes others),ADCY9,MAP2K2,ADCY6,PRKAR2A,JUND,PTGS2,CACNA2D4,GUCY1B1 |
| RAR Activation | FOS,ADCY9,ADH7,LRAT,CSNK2A1,RARB,ADCY6,PRKAR2A,PML,STAT5B,RBP1,PRMT1 |
| Serotonin Receptor Signaling | ADCY9,HTR2B,SMOX,HTR3A,ADCY6 |
| Melanocyte Development and Pigmentation Signaling | KITLG,ADCY9,MAP2K2,FGFR4,RPS6KA3,ADCY6,PRKAR2A,CRK |
| Wnt/β-catenin Signaling | CSNK1E,SOX4,CSNK2A1,APPL2,DKK3,CSNK1G3,MARK2,PPP2R2B,RARB,ILK,FZD1 |

| | |
|---|---|
| IL-8 Signaling | RAB11FIP2,MYL9,PTK2,FOS,ANGPT2,NAPEPLD,MAP2K2,RHOT1,FGFR4,FLT4,PTGS2,RAC3 |
| Molecular Mechanisms of Cancer | ITGB1,HAT1,PRKAR2A,ADCY6,CRK,HIF1A,FZD1,RAC3,PTK2,ADCY9,FOS,CASP6,CDK8,MAP2K2,RHOT1,FGFR4,CAMK2B,CTNND1,PSEN1 |
| p53 Signaling | CASP6,GADD45B,STAG1,FGFR4,ADGRB1,HIF1A,PML,ST13 |
| IGF-1 Signaling | PTK2,FOS,CSNK2A1,MAP2K2,FGFR4,FOXO3,PRKAR2A,SOCS5 |
| Regulation of Actin-based Motility by Rho | ITGB1,MYL9,MPRIP,RHOT1,PFN2,MYLK,RAC3 |
| **Male IUGR HFCS Diet versus Chow Diet** | |
| **Mitochondrial Function** | **GSR,PDHA1,ATP5PF,NDUFB11,NDUFA9,APH1A,MAPK10,COX5A,NDUFA12,UQCR11,ATP5S,BCL2,UQCRB** |
| **Assembly of RNA Polymerase I Complex** | **TAF1C,TBP,UBTF** |
| RAR Activation | PBRM1,RDH10,ADCY6,PDPK1,SMARCD1,SMARCD3,ADH7,IGFBP3,MAPK10,SORBS3,CSNK2B,ADCY10,PML,SMAD1 |
| Protein Ubiquitination Pathway | PSMD9,HSPA6,UBE2V2,FBXW7,PSMD3,DNAJA1,DNAJB14,UBE2D2,PSME1,UBE2B,PSMC6,PSMB2,USP16,USP47,ANAPC5,UBE2E3,UBC |
| CDK5 Signaling | GNAS,PPP2R3A,PPP2CA,ADCY6,MAPK10,PPP2R2C,MAPK7,ADCY10,GNAL |

**Supplementary Table 3: Causal Networks**

| Upstream Regulator | Expr Fold Change | Molecule Type | Predicted Activation State | Activation z-score | p-value of overlap | Target Molecules in Dataset |
|---|---|---|---|---|---|---|
| **Female IUGR Chow vs Female CON Chow** | | | | | | |
| RB1 | 1.152 | transcription regulator | Inhibited | -2.358 | 3.65E-03 | CDC6,CLDN3,COX4I1,EI24,HSBP1,MCM7,MRPL11,MRPL53,MYL6B,NDUFB8 |
| HOXA10 | | transcription regulator | Activated | 2 | 4.76E-02 | APOE,CYBB,HOXA11,KLF10,NDRG2 |
| **Male IUGR Chow vs Male CON Chow** | | | | | | |
| None | | | | | | |
| **Female CON HFCS Diet versus Chow Diet** | | | | | | |
| MED13 | -1.03 | transcription regulator | Inhibited | -2.236 | 3.35E-09 | CD36,CIDEA,CIDEC,FASN,GPD1,PPARG,SCD,THRSP |
| AHR | 1.166 | ligand-dependent nuclear receptor | Inhibited | -2.284 | 2.33E-07 | CDKN2C,COL1A1,COL1A2,COL3A1,COL5A1,COL6A3,FABP4,FBLN2,GAS1,GPAM |
| FBXW7 | -1.044 | enzyme | Inhibited | -2.216 | 1.00E-06 | CD36,CIDEC,FABP4,GPAM,HSPE1,PPARG |
| HAND2 | | transcription regulator | Inhibited | -2.213 | 6.05E-05 | COL1A1,COL1A2,COL3A1,ECM2,FAP |
| TBX5 | | transcription regulator | Inhibited | -2.213 | 8.48E-05 | COL1A1,COL1A2,COL3A1,ECM2,FAP |
| EOMES | 1.002 | transcription regulator | Inhibited | -2 | 4.54E-04 | ANXA1,COL3A1,COL6A3,PMP22,SOX2 |
| MYOCD | 1.032 | transcription regulator | Inhibited | -2.213 | 6.86E-04 | COL1A1,COL1A2,COL3A1,ECM2,FAP |
| GATA4 | 1.081 | transcription regulator | Inhibited | -2.213 | 2.80E-03 | COL1A1,COL1A2,COL3A1,CTHRC1,ECM2,FAP |
| SPDEF | | transcription regulator | Inhibited | -2 | 3.67E-03 | COL1A1,COL5A1,COL6A2,COL6A3 |

| | | | | | | |
|---|---|---|---|---|---|---|
| PPARA | -1.069 | ligand-dependent nuclear receptor | Activated | 2.453 | 5.43E-09 | CD36,CDKN2C,CEACAM1,CIDEA,CIDEC,ELOVL5,FABP4,FABP5,FASN,FGB |
| IGF2BP1 | -1.232 | translation regulator | Activated | 2.449 | 1.58E-08 | COL1A1,COL1A2,COL5A1,COL6A2,COL6A3,LGALS1 |
| SREBF1 | -1.086 | transcription regulator | Activated | 2.928 | 1.36E-07 | ACLY,ALDOC,CIDEA,FABP5,FASN,GCK,GPAM,NSDHL,PLTP,PMP22 |
| PPARG | 2.294 | ligand-dependent nuclear receptor | Activated | 2.959 | 2.41E-07 | ACLY,AGPAT2,CAV1,CAV2,CD36,CIDEA,CIDEC,COL1A1,COL1A2,FABP4 |
| KLF15 | 1.118 | transcription regulator | Activated | 2.355 | 1.69E-06 | CD36,FABP5,FASN,GPAM,PPARG,SCD |
| SREBF2 | -1.001 | transcription regulator | Activated | 2.599 | 2.82E-06 | ACLY,ALDOC,FABP5,FASN,NSDHL,SCD,THRSP |
| TP53 | 1.075 | transcription regulator | Activated | 2.719 | 1.46E-05 | ACLY,ANXA1,ANXA8/ANXA8L1,CAV1,CAV2,COL1A1,COL1A2,COL3A1,COL6A2,FABP4 |
| NR1I2 | | ligand-dependent nuclear receptor | Activated | 2.104 | 1.20E-02 | CD36,FASN,GSTM5,PPARG,SCD |
| **Female IUGR HFCS Diet versus Chow Diet** | | | | | | |
| SMARCA4 | -1.095 | transcription regulator | Activated | 2.111 | 3.54E-02 | ARL16,CDK20,CLK1,GPX1,HNMT,HYI,ITGA7,KHDRBS3,MAFF,MT1H |
| **Male CON HFCS Diet versus Chow Diet** | | | | | | |
| CREB1 | 1.188 | transcription regulator | Activated | 2.159 | 3.45E-04 | AVP,CH25H,FOS,GADD45B,INHA,JUNB,MSMO1,PSEN1,PTGS2,RHEB |
| AR | -1.297 | ligand-dependent nuclear receptor | Activated | 2.092 | 7.49E-03 | ABCC4,AMD1,CALU,CCDC85B,CCT3,CLDN11,CLSTN1,DHCR7,DSC2,FKBP5 |
| CREM | | transcription regulator | Activated | 2.207 | 1.21E-02 | CH25H,FOS,GADD45B,INHA,JUNB,MSMO1,SERTAD1,SPTY2D1 |

| | | | | | | |
|---|---|---|---|---|---|---|
| SMAD3 | | transcription regulator | Activated | 2.916 | 1.84E-02 | FOS,INHA,ITGB1,JUNB,JUND,mir-21,PTGS2,TBX3,TNC |
| SRF | 1.216 | transcription regulator | Activated | 2.195 | 3.12E-02 | CAPRIN1,EIF6,FHL2,FOS,ITGB1,JUNB,MYL9,MYLK,PTGS2,TAGLN |
| **Male IUGR HFCS Diet versus Chow Diet** | | | | | | |
| PARP1 | -1.867 | enzyme | Inhibited | -2.588 | 3.09E-02 | GBP6,ICAM1,OPRM1,POU2F1,TGFA,TLR9,TNFRSF10B |

**Supplementary Table 4. Normalized and transformed values of identified metabolites F=female, M=male, CON**

| Metabolite | Unique mass | RT_dim1(sec) | RT_dim2(sec) | F1_CON_0 | F2_CON_0 |
|---|---|---|---|---|---|
| Pentanoic acid, trimethylsilyl ester | 75 | 590 | 1.33 | 20340 | 2250 |
| Decane | 57 | 630 | 1.18 | 5930 | 6970 |
| Decanoic acid, trimethylsilyl ester | 147 | 635 | 1.05 | 300 | 380 |
| Benzene, 1,2,3-trimethyl- | 105 | 640 | 1.99 | 900 | 1210 |
| Ethanedioic acid, bis(trimethylsilyl) ester | 147 | 645 | 1.11 | 1200 | 2030 |
| Indane | 117 | 655 | 2.22 | 330 | 440 |
| 3-t-Pentylcyclopentanone | 69 | 670 | 1.25 | #N/A | #N/A |
| 2-Hepten-4-one, 6-methyl- | 69 | 680 | 1.29 | #N/A | #N/A |
| Tridecane | 57 | 685 | 1.18 | 750 | 470 |
| Benzene, 2-ethyl-1,4-dimethyl-/Benzene, 4-ethyl-1,2-dimethyl- | 119 | 700 | 1.88 | 1790 | 2240 |
| 2H-Pyran-2-one, tetrahydro-6-methyl- | 42 | 700 | 4.4 | 1430 | 1680 |
| 2,4,6,8-Tetramethyl-1-undecene | 57 | 720 | 1.2 | 1220 | 1070 |
| N,N-Dimethyloctylamine/3-Buten-1-amine, N,N-dimethyl- | 58 | 790 | 1.39 | 920 | 1280 |
| Propanoic acid, 2-oxo-3-(trimethylsilyl)-, trimethylsilyl ester | 119 | 795 | 2.02 | #N/A | #N/A |
| Heptanoic acid, trimethylsilyl ester | 75 | 890 | 1.49 | 1220 | 980 |
| 2-Methoxyphenol trimethylsilyl ether | 166 | 970 | 2.11 | 1230 | 1010 |
| Urea, N,N'-bis(trimethylsilyl)- | 147 | 990 | 1.98 | 260 | 460 |
| 3-Ethylphenol, trimethylsilyl ether | 179 | 995 | 1.82 | 1000 | 880 |
| Benzeneacetic acid, trimethylsilyl ester | 73 | 1075 | 2.28 | 41110 | 61820 |
| 1,2-Benzenediol bis(trimethylsilyl) ether | 73 | 1135 | 1.64 | 1640 | 4010 |
| 3-Octenoic acid, trimethylsilyl ester | 73 | 1190 | 1.61 | #N/A | 750 |
| 4-Methylcatechol, bis(trimethylsilyl) ether | 73 | 1255 | 1.61 | #N/A | #N/A |
| Benzenepropanoic acid, trimethylsilyl ester | 104 | 1265 | 2.22 | 3300 | 3860 |
| Mercaptoacetic acid, bis(trimethylsilyl)- | 73 | 1330 | 0.97 | #N/A | #N/A |

| | | | | | |
|---|---|---|---|---|---|
| Benzenepropanoic acid, 4-[(trimethylsilyl)oxy]-‡-[[(trimethylsilyl)oxy]imino]-, trimethylsilyl ester | 75 | 1350 | 1.52 | #N/A | #N/A |
| Benzoic acid, 2-[(trimethylsilyl)oxy]-, trimethylsilyl ester | 73 | 1420 | 1.91 | 1700 | 900 |
| Trimethylsilyl 2,4-dichlorobenzoate | 173 | 1460 | 2.28 | 1140 | 1230 |
| Benzenepropanoic acid, ‡-[(trimethylsilyl)oxy]-, trimethylsilyl ester | 73 | 1530 | 1.78 | 1010 | 430 |
| 3-Phenyl-3-trimethylsilyloxypropanoic acid, trimethylsilyl ester | 73 | 1545 | 1.75 | 380 | 710 |
| Benzeneacetic acid, 3-[(trimethylsilyl)oxy]-, trimethylsilyl ester/Benzeneacetic acid, 4-[(trimethylsilyl)oxy]-, trimethylsilyl ester | 73 | 1595 | 1.94 | 640 | 740 |
| 3-Phenyl-3-(trimethylsilyloxy)butan-2-one | 73 | 1605 | 1.84 | 400 | 460 |
| Dodecanoic acid, trimethylsilyl ester | 75 | 1620 | 1.55 | 280 | 130 |
| Benzene, 1,2,3,4-tetramethyl- | 73 | 1685 | 2.18 | #N/A | #N/A |
| 1-Pentadecanamine, N,N-dimethyl- | 58 | 1690 | 1.53 | 5930 | 13880 |
| 1-Octanamine, N-methyl-N-octyl- | 58 | 1725 | 1.5 | 890 | 1450 |
| n-Tridecanoic acid, trimethylsilyl ester | 73 | 1750 | 1.54 | #N/A | 100 |
| Azelaic acid, bis(trimethylsilyl) ester | 75 | 1800 | 1.72 | 1110 | 1710 |
| Glycine, N-benzoyl-, trimethylsilyl ester | 105 | 1815 | 3.26 | 2940 | 4900 |
| Benzenepropanoic acid, 3-methoxy-4[(trimethylsilyl)oxy]-, trimethylsilyl ester | 73 | 1910 | 2.1 | 2130 | 940 |
| n-Pentadecanoic acid, trimethylsilyl ester | 75 | 1945 | 1.53 | 140 | 240 |
| 1H-Indole-3-acetic acid, 1-(trimethylsilyl)-, trimethylsilyl ester | 202 | 1975 | 2.7 | 30360 | 36110 |
| Dodecanedioic acid, bis(trimethylsilyl) ester | 73 | 1980 | 1.61 | 10 | 2730 |
| 1,11-Undecanedioic acid, di(trimethylsilyl) ester | 73 | 2030 | 1.72 | 450 | 660 |
| 1H-Indole-3-propanoic acid, 1-(trimethylsilyl)-, trimethylsilyl ester | 73 | 2115 | 2.52 | 400 | 580 |
| N-Methyl-N-benzyltetradecanamine | 91 | 2155 | 2.04 | 190 | 220 |

| | | | | | |
|---|---|---|---|---|---|
| L-Tryptophan, N,1-bis(trimethylsilyl)-, trimethylsilyl ester | 73 | 2235 | 2.05 | #N/A | 260 |
| 9,12-Octadecadienoic acid (Z,Z)-, trimethylsilyl ester | 75 | 2265 | 1.77 | 3360 | 5130 |
| trans-9-Octadecenoic acid, trimethylsilyl ester | 129 | 2275 | 1.69 | 2530 | 30100 |
| Octadecanoic acid, trimethylsilyl ester | 341 | 2305 | 1.6 | 202050 | 136170 |
| Arachidonic acid, trimethylsilyl ester | 75 | 2415 | 1.94 | #N/A | 320 |
| Sebacic acid, bis(trimethylsilyl) ester | 73 | 2495 | 1.47 | 2250 | 2460 |
| Hexanoic acid, trimethylsilyl ester | 73 | 2510 | 1.47 | 650 | 230 |
| Tetracosanoic acid, trimethylsilyl ester | 117 | 2840 | 1.67 | 190 | 210 |
| Cholesterol trimethylsilyl ether | 129 | 3080 | 2.23 | 1650 | 5820 |

= Control, IUGR = Interuterine Growth Restricted, 0 = Chow Diet, 7 = HFCS diet.

| F3_CON_0 | F1_IUGR_0 | F2_IUGR_0 | F3_IUGR_0 | M1_CON_0 | M2_CON_0 | M3_CON_0 | M1_IUGR_0 | M3_IUGR_0 |
|---|---|---|---|---|---|---|---|---|
| 14080 | 4700 | 12900 | 4710 | 3190 | 1740 | 55450 | #N/A | 2920 |
| 9720 | 12680 | 14330 | 10110 | 11700 | 21150 | 34050 | 58760 | 13930 |
| 950 | 1120 | #N/A | 750 | #N/A | 1160 | 1560 | #N/A | 470 |
| 1720 | 2100 | 2390 | 1660 | 3490 | 3290 | 5520 | 8150 | 2060 |
| 2700 | #N/A | 5210 | #N/A | 6820 | #N/A | #N/A | 9990 | 4400 |
| 640 | 770 | 910 | 620 | 1170 | 1210 | 2070 | 2710 | 740 |
| #N/A | #N/A | #N/A | #N/A | #N/A | #N/A | #N/A | #N/A | #N/A |
| #N/A | #N/A | #N/A | #N/A | #N/A | #N/A | #N/A | #N/A | #N/A |
| 530 | 1520 | 1800 | 750 | 1190 | 1830 | 2820 | 5020 | 1580 |
| 2960 | 3960 | 4650 | 3230 | 1520 | 5580 | 9110 | 14640 | 3740 |
| 4120 | 1180 | 3770 | 840 | 7450 | 1930 | 770 | 10980 | 400 |
| 1980 | 2450 | 3150 | 1030 | #N/A | #N/A | #N/A | #N/A | #N/A |
| 5730 | 2740 | 3130 | 1640 | #N/A | #N/A | #N/A | #N/A | #N/A |
| #N/A | #N/A | #N/A | #N/A | #N/A | #N/A | #N/A | #N/A | #N/A |
| 3310 | 10 | 1160 | 1290 | #N/A | 1230 | 3260 | #N/A | 280 |
| 610 | 690 | 440 | 1060 | #N/A | #N/A | #N/A | #N/A | #N/A |
| 910 | 270 | #N/A | #N/A | 3670 | #N/A | #N/A | 6720 | 2050 |
| 2990 | 2840 | 2790 | 500 | 1470 | #N/A | 4480 | 2610 | 1930 |
| 79030 | 27840 | 46200 | 61650 | 79340 | 13050 | 60420 | 4870 | 45250 |
| 8470 | 6820 | 3790 | 5330 | 2480 | 19070 | 31360 | 1560 | 2410 |
| 3080 | 490 | #N/A | 730 | #N/A | 1330 | 1600 | #N/A | 1100 |
| #N/A | #N/A | #N/A | #N/A | #N/A | #N/A | #N/A | #N/A | #N/A |
| 4070 | 1730 | 3760 | 1690 | 2340 | #N/A | 14320 | 3210 | 1510 |
| #N/A | #N/A | #N/A | #N/A | #N/A | #N/A | #N/A | #N/A | #N/A |

| | | | | | | | | |
|---|---|---|---|---|---|---|---|---|
| #N/A | #N/A | #N/A | #N/A | #N/A | #N/A | #N/A | #N/A | #N/A |
| 2260 | 850 | 1440 | 760 | 4670 | 1250 | 3920 | 20370 | 3740 |
| 5700 | 1440 | 2360 | 1390 | 4020 | 1950 | 10600 | 3620 | 8120 |
| 2500 | 580 | 1280 | 7610 | 2430 | 18500 | 3510 | 10330 | 4330 |
| 1340 | 500 | 830 | #N/A | 2110 | #N/A | 1470 | 2890 | 2250 |
| 1400 | #N/A | #N/A | 960 | 2160 | 1870 | 3640 | 2790 | 2180 |
| 500 | #N/A | #N/A | 290 | #N/A | #N/A | #N/A | #N/A | #N/A |
| 830 | 430 | 380 | 420 | 350 | 620 | 2040 | #N/A | 290 |
| #N/A | #N/A | #N/A | #N/A | #N/A | #N/A | #N/A | #N/A | #N/A |
| 21830 | 21030 | 10400 | 10910 | 4040 | 17870 | #N/A | 4350 | 3900 |
| 7440 | 2890 | 3120 | 1940 | 5150 | 6430 | #N/A | 4460 | 3310 |
| 350 | 260 | #N/A | 100 | #N/A | #N/A | #N/A | #N/A | #N/A |
| 3950 | 850 | 1790 | 1110 | 1750 | 910 | 5110 | #N/A | 3360 |
| 11790 | 3530 | 10200 | 2170 | 15620 | 1640 | 17760 | #N/A | 32690 |
| 2740 | 740 | 700 | 1480 | 7860 | 2760 | 2390 | 4600 | 3540 |
| 280 | 270 | 320 | 490 | #N/A | #N/A | #N/A | #N/A | #N/A |
| 39760 | 11890 | 56080 | 28830 | 90110 | 80290 | 58760 | 68830 | 80450 |
| 3200 | 1970 | 2180 | 220 | 5330 | #N/A | 1120 | 2180 | 4320 |
| 1050 | 510 | 550 | 510 | #N/A | #N/A | #N/A | #N/A | #N/A |
| 1060 | 650 | 790 | 310 | 780 | #N/A | #N/A | #N/A | 1110 |
| 1320 | 280 | 280 | 180 | #N/A | #N/A | #N/A | #N/A | #N/A |

| | | | | | | | | |
|---|---|---|---|---|---|---|---|---|
| 510 | #N/A | #N/A | 280 | 1000 | #N/A | #N/A | 1500 | 900 |
| 20620 | 8100 | 2370 | 5500 | #N/A | #N/A | #N/A | #N/A | #N/A |
| 12770 | 42790 | 5390 | 5050 | 7090 | 22420 | #N/A | 7090 | 17090 |
| 5100 | 152790 | 118920 | 167840 | 40780 | 51910 | #N/A | 63300 | 83360 |
| 600 | 780 | #N/A | #N/A | #N/A | #N/A | #N/A | #N/A | #N/A |
| 9920 | 3310 | 1940 | 4470 | #N/A | #N/A | #N/A | #N/A | #N/A |
| 1240 | 230 | 270 | 440 | #N/A | #N/A | #N/A | #N/A | #N/A |
| 230 | #N/A | #N/A | #N/A | #N/A | #N/A | #N/A | #N/A | #N/A |
| 2620 | 4810 | 1930 | 1440 | 1100 | #N/A | #N/A | #N/A | 620 |

| F1_CON_7 | F2_CON_7 | F3_CON_7 | F1_IUGR_7 | F2_IUGR_7 | F3_IUGR_7 | M1_CON_7 | M2_CON_7 | M3_CON_7 |
|---|---|---|---|---|---|---|---|---|
| 34490 | 4610 | 12340 | 5150 | 51530 | 6850 | #N/A | 3700 | 1690 |
| 10430 | 18050 | 17210 | 17720 | 11050 | 18310 | 37830 | 26510 | 8570 |
| #N/A | 670 | 850 | 810 | 800 | 750 | 860 | 1270 | 870 |
| 1590 | 2300 | 2870 | 2830 | 2790 | 2690 | 5830 | 4150 | 2850 |
| 2790 | 6580 | 10480 | 4740 | 6870 | 3690 | 90 | 21020 | 7250 |
| 610 | 200 | 1080 | 960 | 1140 | 1040 | 2150 | 1590 | 1080 |
| 1760 | 1110 | 2590 | 1170 | 2540 | 1240 | #N/A | 4220 | #N/A |
| #N/A | #N/A | 1320 | #N/A | #N/A | #N/A | 2940 | 2050 | #N/A |
| #N/A | #N/A | #N/A | #N/A | #N/A | #N/A | #N/A | #N/A | #N/A |
| 2980 | 1220 | 4890 | 4930 | 1200 | 4030 | 10040 | 7290 | 1790 |
| #N/A | #N/A | #N/A | #N/A | #N/A | #N/A | #N/A | #N/A | #N/A |
| #N/A | #N/A | #N/A | #N/A | #N/A | #N/A | #N/A | #N/A | #N/A |
| 3010 | 19140 | 14560 | 12340 | 9430 | 16820 | 4130 | 5210 | 4440 |
| 800 | 1750 | 3660 | 1490 | 2150 | 1310 | 2710 | 5280 | 2260 |
| 1520 | 1820 | 4800 | 2830 | 7260 | 2420 | #N/A | 3280 | 2760 |
| 920 | #N/A | 550 | #N/A | #N/A | 990 | #N/A | #N/A | 980 |
| #N/A | #N/A | #N/A | #N/A | #N/A | #N/A | #N/A | #N/A | #N/A |
| 1870 | 270 | 1600 | 1090 | 2100 | 1390 | 700 | 1730 | 1660 |
| 124190 | 8220 | 50380 | 84580 | 33830 | 52880 | 32370 | 20330 | 62060 |
| 5570 | 14900 | 17290 | 12190 | 4080 | 23240 | 1920 | 8070 | 2720 |
| #N/A | 2460 | 1310 | #N/A | 610 | 610 | #N/A | 790 | 11970 |
| #N/A | #N/A | 2020 | 1620 | 870 | 1230 | #N/A | 1040 | #N/A |
| 7920 | 880 | 5930 | 2550 | 5910 | 2690 | 3310 | 1290 | 1510 |
| #N/A | #N/A | 380 | #N/A | #N/A | 570 | 410 | #N/A | #N/A |

| | | | | | | | | |
|---|---|---|---|---|---|---|---|---|
| #N/A | 3540 | 1660 | 1370 | #N/A | 1540 | #N/A | #N/A | 40 |
| 1650 | 1330 | 2070 | 1290 | #N/A | 1910 | 4200 | 2930 | 5090 |
| 740 | 7280 | 4190 | 7770 | 4440 | 2010 | 3270 | 14930 | 7950 |
| #N/A | #N/A | 660 | 500 | #N/A | 890 | 4740 | 4060 | 1770 |
| #N/A | #N/A | #N/A | #N/A | #N/A | #N/A | #N/A | #N/A | #N/A |
| #N/A | 340 | 930 | #N/A | 510 | 670 | 2750 | 2350 | 2200 |
| #N/A | #N/A | #N/A | #N/A | #N/A | #N/A | #N/A | #N/A | #N/A |
| 580 | 930 | 420 | 1270 | 710 | 1320 | 1670 | 840 | 1350 |
| 3530 | 4610 | 5800 | 5650 | 5210 | 4980 | 11920 | 8350 | 6060 |
| 15380 | 27570 | 24490 | 20410 | 22620 | 38450 | 4760 | 16390 | 13610 |
| 890 | 2570 | 12830 | 8160 | 6940 | 3100 | 4340 | 5050 | 4400 |
| #N/A | #N/A | #N/A | #N/A | #N/A | #N/A | #N/A | #N/A | #N/A |
| 630 | 790 | 3190 | 860 | 670 | 1450 | 4260 | 3950 | 6560 |
| 950 | 1430 | 4500 | 2840 | 2560 | 5240 | 17850 | 19770 | 32400 |
| #N/A | #N/A | #N/A | #N/A | #N/A | #N/A | #N/A | #N/A | #N/A |
| #N/A | #N/A | #N/A | #N/A | #N/A | #N/A | #N/A | #N/A | #N/A |
| 33780 | 33670 | 15570 | 22290 | 18990 | 18540 | 163420 | 106450 | 92210 |
| #N/A | #N/A | #N/A | #N/A | #N/A | #N/A | #N/A | #N/A | #N/A |
| #N/A | #N/A | #N/A | #N/A | #N/A | #N/A | #N/A | #N/A | #N/A |
| #N/A | #N/A | #N/A | #N/A | #N/A | #N/A | #N/A | #N/A | #N/A |
| #N/A | #N/A | #N/A | #N/A | #N/A | #N/A | #N/A | #N/A | #N/A |

| | | | | | | | | |
|---|---|---|---|---|---|---|---|---|
| #N/A | #N/A | #N/A | #N/A | #N/A | #N/A | #N/A | #N/A | #N/A |
| 14520 | #N/A | 1780 | 1550 | 6320 | 2000 | #N/A | 1280 | 1810 |
| 12830 | 1760 | 5190 | 3180 | 1750 | 3570 | 1150 | 3770 | 3480 |
| 26260 | 3830 | 7910 | 3180 | 51570 | 5500 | 2100 | 4770 | 4560 |
| #N/A | #N/A | #N/A | #N/A | #N/A | #N/A | #N/A | #N/A | #N/A |
| 930 | #N/A | 900 | #N/A | 4120 | 600 | #N/A | 800 | 330 |
| #N/A | #N/A | #N/A | #N/A | #N/A | #N/A | #N/A | #N/A | #N/A |
| #N/A | #N/A | #N/A | #N/A | #N/A | #N/A | #N/A | #N/A | #N/A |
| 7230 | 710 | 5590 | 2840 | 5020 | 1500 | #N/A | #N/A | 4640 |

| M1_IUGR_7 | M2_IUGR_7 | M3_IUGR_7 |
|---|---|---|
| 2290 | 60 | #N/A |
| 15420 | 29650 | 28400 |
| #N/A | 740 | 740 |
| 2320 | 4190 | 4020 |
| 2130 | 1710 | 3480 |
| 780 | 1580 | 1540 |
| 3300 | #N/A | #N/A |
| #N/A | #N/A | 2020 |
| #N/A | #N/A | #N/A |
| 4020 | 7750 | 6670 |
| #N/A | #N/A | #N/A |
| #N/A | #N/A | #N/A |
| 7980 | 15730 | 12050 |
| 1040 | 2260 | 1870 |
| 1860 | 2560 | 2120 |
| #N/A | 1130 | #N/A |
| #N/A | #N/A | #N/A |
| 600 | 1140 | #N/A |
| 20800 | 11480 | 16320 |
| 4830 | 15600 | 4980 |
| 1980 | 1110 | 910 |
| #N/A | 3610 | 870 |
| #N/A | #N/A | 440 |
| 470 | 680 | 560 |

| | | |
|---|---|---|
| #N/A | 1300 | #N/A |
| 3060 | 2060 | 4420 |
| 151220 | 9160 | 12000 |
| 1480 | #N/A | 3640 |
| #N/A | #N/A | #N/A |
| 1880 | 970 | 3270 |
| #N/A | #N/A | #N/A |
| 270 | 790 | 400 |
| 4800 | 9110 | 8560 |
| 7090 | 31410 | 4860 |
| 9130 | 12690 | 12380 |
| #N/A | #N/A | #N/A |
| 2680 | 1710 | 3880 |
| 7130 | 5230 | 10870 |
| #N/A | #N/A | #N/A |
| #N/A | #N/A | #N/A |
| 38360 | 58870 | 163330 |
| #N/A | #N/A | #N/A |
| #N/A | #N/A | #N/A |
| #N/A | #N/A | #N/A |
| #N/A | #N/A | #N/A |

| | | |
|---|---|---|
| #N/A | #N/A | #N/A |
| 610 | 970 | #N/A |
| 1540 | 2530 | 610 |
| 1920 | 4780 | 830 |
| #N/A | #N/A | #N/A |
| #N/A | #N/A | #N/A |
| #N/A | #N/A | #N/A |
| #N/A | #N/A | #N/A |
| #N/A | 820 | #N/A |

**Supplementary Table 5. Identifiers of metabolites including NIST Identifier, common name, HMDB ID, KEGG ID, PubChem ID, Class**

| NIST Identifier | Common Name | HMDB | KEGG | PubChem | Chemical class | Pathway |
|---|---|---|---|---|---|---|
| Pentanoic acid, trimethylsilyl ester | Valeric acid | HMDB0000892 | C00803 | 7991 | Fatty Acid | NA |
| Decane | Decane | HMDB0031450 | NA | 15600 | Hydrocarbon | NA |
| Benzene, 1,2,3-trimethyl- | Hemimellitene | HMDB0059901 | NA | 10686 | benzenoid | Degradation of Aromatic Compounds |
| Ethanedioic acid, bis(trimethylsilyl) ester | oxalic acid | HMDB0002329 | C00209 | 971 | carboxylic acid | metabolism of glyocylic acid and ascorbic acid excreted in urine |
| Indane | Indane | HMDB0059837 | NA | 10326 | benzenoid | Degradation of Aromatic Compounds |
| 3-Ethylphenol, trimethylsilyl ether | 3-Ethylphenol | HMDB0059873 | C14386 | 6997 | benzenoid | Degradation of Aromatic Compounds |
| Benzene, 2-ethyl-1,4-dimethyl- | 2-ethyl-p-xylene | HMDB0059890 (Predicted) | NA | 15653 | benzenoid | Degradation of Aromatic Compounds |
| Heptanoic acid, trimethylsilyl ester | Heptanoic acid | HMDB0000666 | C17714 | 8094 | carboxylic acid | NA |
| Benzeneacetic acid, trimethylsilyl ester | Phenylacetic acid | HMDB00209 | C07086 | 999 | carboxylic acid | Phenylalanine metabolism |
| 1,2-Benzenediol bis(trimethylsilyl) ether | pyrocatechol | HMDB00957 | C00090 | 289 | benzenoid | Aminobenzoate degradation |
| Benzoic acid, 2-[(trimethylsilyl)oxy]-, trimethylsilyl ester | salicylic acid | HMDB01895 | C00805 | 338 | carboxylic acid | Phenylalanine metabolism |
| Dodecanoic acid, trimethylsilyl ester | Lauric acid | HMDB00638 | C02679 | 3893 | Fatty Acid | fatty acid synthesis |
| 1-Pentadecanamine, N,N-dimethyl- | N,N-dimethyl-1-Pentadecanamine | No result | NA | 87232 | amine | NA |
| 1-Octanamine, N-methyl-N-octyl- | N-methyl-N-octyl-1-Octanamine | No result | NA | 78202 | amine | NA |

| | | | | | | |
|---|---|---|---|---|---|---|
| Azelaic acid, bis(trimethylsilyl) ester | nonanedioic acid | HMDB00784 | C08261 | 2266 | Fatty Acid | NA |
| Glycine, N-benzoyl-, trimethylsilyl ester | hippuric acid | HMDB00714 | C01586 | 464 | carboxylic acid | Phenylalanine metabolism |
| 1H-Indole-3-acetic acid, 1-(trimethylsilyl)-, trimethylsilyl ester | Indoleacetic acid | HMDB00197 | C00954 | 802 | Indoles | tryptophan metabolism |
| trans-9-Octadecenoic acid, trimethylsilyl ester | Elaidic acid | HMDB0000573 | C00712 | 445639 | Fatty acid | Biosynthesis of unsaturated fatty acids |
| Octadecanoic acid, trimethylsilyl ester | Stearic acid | HMDB00827 | C01530 | 5281 | Fatty acid | Biosynthesis of unsaturated fatty acids |
| N,N-Dimethyloctylamine | N,N-Dimethyloctylamine | No result | NA | 16224 | amine | NA |
| Benzenepropanoic acid, trimethylsilyl ester | hydrocinnamic acid | HMDB00764 | C05629 | 107 | phenylpropanoids | phenyalanine metablism |
| Benzeneacetic acid, 3-[(trimethylsilyl)oxy]-, trimethylsilyl ester | 3-hydroxyphenylacetic acid | HMDB0000440 | C05593 | 12122 | phenol | tyrosine metabolism associated with phenylketonuria |
| 3-Octenoic acid, trimethylsilyl ester | (Z)-3-Octenoic acid | No result | NA | 5282716 | Fatty Acid | NA |
| Benzenepropanoic acid, ‡-[(trimethylsilyl)oxy]-, trimethylsilyl ester | Phenyllactic acid | HMDB0000779 | C01479 | 3848 | phenylpropanoids | phenyalanine metablism |
| Cholesterol trimethylsilyl ether | Cholesterol | HMDB0000067 | C00187 | 11025495 | sterol | Steroid biosynthesis |
| N-Methyl-N-benzyltetradecanamine | N-Methyl-N-benzyltetradecanamine | No result | NA | 584166 | amine | NA |
| Trimethylsilyl 2,4-dichlorobenzoate | 2,4-dichlorobenzoate | No result | NA | 5787 | benzenoid | Degradation of Aromatic Compounds |
| Decanoic acid, trimethylsilyl ester | Capric acid | HMDB000511 | NA | 2669 | dicarboxylic acid | normal uninary acid |

| | | | | | | |
|---|---|---|---|---|---|---|
| 9,12-Octadecadienoic acid (Z,Z)-, trimethylsilyl ester | Linoleic acid | HMDB00673 | C01595 | 5280450 | Fatty Acid | Linoleic acid metabolism/Biosynthesis of unsaturated fatty acids |
| 2-Methoxyphenol trimethylsilyl ether | Catechol mono methyl ester | HMDB01398 | C01502 | 460 | benzenoid | Aminobenzoate degradation |
| Sebacic acid, bis(trimethylsilyl) ester | Decanedioic acid | HMDB00792 | C08277 | 5192 | Fatty Acid | NA |
| Propanoic acid, 2-oxo-3-(trimethylsilyl)-, trimethylsilyl ester | (S)3-hydroxyisobutyric acid (3-HIBA) | HMDB0000023 | C06001 | 440873 | hydroxy acid | Valine, leucine and isolucine degradation |
| Benzene, 1,2,3,4-tetramethyl- | prehnitene | HMDB0059823 | NA | 10236 | benzenoid | Degradation of Aromatic Compounds |
| Benzenepropanoic acid, 4-[(trimethylsilyl)oxy]-‡-[[(trimethylsilyl)oxy]imino]-, trimethylsilyl ester | 4-Hydroxyphenylpyruvic acid (4-HPPA) | HMDB0000707 | C01179 | 979 | benzenoid | Phenylalanine, tyrosine and tryptophan biosynthesis/ Ubiquinone and other terpenoid-quinone biosynthesis |
| 3-t-Pentylcyclopentanone | 3-t-Pentylcyclopentanone | No result | NA | 551379 | Ketone | NA |
| 2-Hepten-4-one, 6-methyl- | 6-methyl-2-Hepten-4-one | No result | NA | 12502788 | Ketone | NA |
| 4-Methylcatechol, bis(trimethylsilyl) ether | 4-methylcatechol | HMDB0000873 | C06730 | 9958 | benzenoid | Benzoate degradation/Degradation of Aromatic Compounds |
| Mercaptoacetic acid, bis(trimethylsilyl)- | Mercaptoacetic acid | No result | C02086 | 1133 | carboxylic acid | NA |
| 2H-Pyran-2-one, tetrahydro-6-methyl- | delta Hexalactone | HMDB0000453 | NA | 13204 | lactons | NA |
| Benzenepropanoic acid, 3-methoxy-4[(trimethylsilyl)oxy]-, trimethylsilyl ester | dihydroferulic acid 3-(4-hydroxy-3-methoxyphenyl) propanoic acid | HMDB0062121 | NA | 14340 | phenylpropanoids | NA |

| | | | | | | |
|---|---|---|---|---|---|---|
| 1H-Indole-3-propanoic acid, 1-(trimethylsilyl)-, trimethylsilyl ester | Indole-3-propionic acid | HMDB0002302 | NA | 3744 | Indoles | tryptophan metabolism |
| Tridecane | Tridecane | HMDB0034284 | C13834 | 12388 | hydrocarbons | NA |
| 3-Phenyl-3-trimethylsilyloxypropanoic acid, trimethylsilyl ester | 3-hydroxy-3-phenylpropanoic acid | HMDB0124925 (predicted) | NA | 92959 | phenylpropanoids | related to phenyl pyruvic acid metabolism |
| Urea, N,N'-bis(trimethylsilyl)- | Urea | HMDB00294 | C00086 | 1176 | amide | Arginine biosynthesis |
| 3-Phenyl-3-(trimethylsilyloxy)butan-2-one | 3-hydroxy-3-phenylbutan-2-one | No result | NA | 233220 | ketone | NA |
| L-Tryptophan, N,1-bis(trimethylsilyl)-, trimethylsilyl ester | L-Tryptophan | HMDB00929 | C00078 | 6305 | amino acid | tryptophan metabolism |
| 1,11-Undecanedioic acid, di(trimethylsilyl) ester | Undecanedioic acid | HMDB00888 | NA | 15816 | Fatty Acid | NA |
| Dodecanedioic acid, bis(trimethylsilyl) ester | Dodecanedioic acid | HMDB0000623 | C02678 | 12736 | Fatty Acid | NA |
| n-Pentadecanoic acid, trimethylsilyl ester | Pentadecanoic acid | HMDB0000826 | C16537 | 13849 | Fatty acid | NA |
| Hexanoic acid, trimethylsilyl ester | caproic acid | HMDB0000535 | C01585 | 8892 | Fatty Acid | Beta Oxidation of Very Long Chain Fatty Acids |
| 2,4,6,8-Tetramethyl-1-undecene | 2,4,6,8-Tetramethyl-1-undecene | No result | NA | 536235 | Hydrocarbon | NA |
| n-Tridecanoic acid, trimethylsilyl ester | Tridecanoic acid | HMDB00910 | C17076 | 12530 | Fatty Acid | NA |
| Arachidonic acid, trimethylsilyl ester | Arachidonic acid | HMDB01043 | C00219 | 444899 | Fatty Acid | Biosynthesis of unsaturated fatty acids |
| Tetracosanoic acid, trimethylsilyl ester | lignoceric acid | HMDB0002003 | C08320 | 11197 | Fatty Acid | Biosynthesis of unsaturated fatty acids |

**and Pathway. X = present in all animals, O = not present in all animals.**

| F CON Chow | F IUGR Chow | M CON Chow | M IUGR Chow | F CON HFCS | F IUGR HFCS | M CON HFCS | M IUGR HFCS |
|---|---|---|---|---|---|---|---|
| X | X | X | O | X | X | O | O |
| X | X | X | X | X | X | X | X |
| X | X | X | X | X | X | X | X |
| X | O | O | X | X | X | X | X |
| X | X | X | X | X | X | X | X |
| X | X | O | X | X | X | X | O |
| X | X | X | X | X | X | X | X |
| X | X | X | O | X | X | O | X |
| X | X | X | X | X | X | X | X |
| X | X | X | X | X | X | X | X |
| X | X | X | X | X | O | X | X |
| X | X | X | O | X | X | X | X |
| X | X | O | X | X | X | X | X |
| X | X | O | X | X | X | X | X |

| | | | | | | | |
|---|---|---|---|---|---|---|---|
| X | X | X | O | X | X | X | X |
| X | X | X | O | X | X | X | X |
| X | X | X | X | X | X | X | X |
| X | X | O | X | X | X | X | X |
| X | X | O | X | X | X | X | X |
| X | X | O | O | X | X | X | X |
| X | X | O | X | X | X | X | O |
| X | O | X | X | O | O | X | X |
| O | O | O | O | O | O | O | X |
| X | X | X | X | O | X | O | O |
| X | X | O | O | X | X | O | O |
| X | X | X | X | X | X | O | X |
| X | X | X | X | X | X | X | X |
| O | O | O | O | O | X | X | O |

| | | | | | | | |
|---|---|---|---|---|---|---|---|
| X | X | | O | O | X | O | O |
| X | X | O | O | O | O | O | O |
| X | X | | | O | O | O | O |
| | | | | X | X | X | X |
| | | | | X | X | X | X |
| | | | | O | O | O | O |
| | | | | X | X | O | O |
| | | | | X | O | X | O |
| | | | | O | X | O | O |
| | | | | O | X | O | X |
| X | X | X | X | | | | |
| X | X | X | X | | | | |

| | | | |
|---|---|---|---|
| X | X | O | O |
| X | X | X | X |
| X | O | O | X |
| X | O | O | X |
| X | O | O | O |
| O | O | O | X |
| X | X | O | O |
| X | X | O | O |
| X | X | | |
| X | X | | |
| X | X | | |
| O | O | | |
| O | O | | |
| X | | | |

**Supplementary Table 6: HFCS Diet vs Chow Diet Gene**

| Gene Symbol | Gene Name | Accession No. | CON Chow F Mean | CON HFHS F Mean | p-val CON F HFHS vs. CON F Chow | Ratio CON F HFHS vs. CON F Chow | REFSEQ ID | NEPHRON (MEAN RPKM) | COLLECTING DUCT (MEAN RPKM) |
|---|---|---|---|---|---|---|---|---|---|
| COL6A3 | collagen type VI alpha 3 chain(COL6A3) | NM_004369 | 103.1 | 136.4 | 0.00078 | 1.288 | | | |
| ACLY | ATP citrate lyase(ACLY) | NM_001096 | 317 | 434.6 | 0.00096 | 1.374 | | | |
| | | BQ186959 | 111.6 | 101.8 | 0.00115 | 0.891 | | | |
| NTRK3 | neurotrophic receptor tyrosine kinase 3(NTRK3) | NM_002530 | 102.8 | 119.4 | 0.0013 | 1.13 | | | |
| | | XM_944079 | 122.9 | 159.3 | 0.00131 | 1.257 | | | |
| PPP1CA | protein phosphatase 1 catalytic subunit alpha(PPP1CA) | NM_002708 | 121.5 | 107.1 | 0.00182 | 0.861 | | | |
| | | AW081577 | 104.5 | 123.6 | 0.00203 | 1.156 | | | |
| | | XM_944499 | 109.3 | 100.2 | 0.0026 | 0.897 | | | |
| C1QTNF6 | C1q and tumor necrosis factor related protein 6(C1QTNF6) | NM_031910 | 106.2 | 114.9 | 0.00271 | 1.136 | NM_017193 | 356.13 | 8.18 |
| C1QTNF6 | C1q and tumor necrosis factor related protein 6(C1QTNF6) | NM_031910 | 110.3 | 128.7 | 0.00271 | 1.136 | NM_017193 | 356.13 | 8.18 |

| | | | | | | | | | |
|---|---|---|---|---|---|---|---|---|---|
| SMAGP | small cell adhesion glycoprotein(SMAGP) | NM_001033873 | 190.5 | 156.5 | 0.00286 | 0.813 | NM_017193 | 356.13 | 8.18 |
| SIDT2 | SID1 transmembrane family member 2(SIDT2) | NM_001040455 | 350 | 417.7 | 0.00289 | 1.205 | NM_017193 | 356.13 | 8.18 |
| FAP | fibroblast activation protein alpha(FAP) | NM_004460 | 96.4 | 127.9 | 0.0029 | 1.298 | | | |
| ISPD | isoprenoid synthase domain containing(ISPD) | NM_001101417 | 118.8 | 104 | 0.00308 | 0.854 | | | |
| | | XM_941979 | 99.2 | 125.3 | 0.00341 | 1.217 | | | |
| ABCA9 | ATP binding cassette subfamily A member 9(ABCA9) | NM_080283 | 115.9 | 144.3 | 0.00348 | 1.218 | | | |
| CEP63 | centrosomal protein 63(CEP63) | NM_025180 | 139.3 | 186.9 | 0.00359 | 1.326 | | | |
| SCD | stearoyl-CoA desaturase(SCD) | NM_005063 | 116.4 | 281.9 | 0.00363 | 2.346 | | | |
| CD248 | CD248 molecule(CD248) | NM_020404 | 120.3 | 190.8 | 0.00367 | 1.546 | | | |
| DCUN1D5 | defective in cullin neddylation 1 domain containing 5(DCUN1D5) | AK094744 | 121.4 | 108.2 | 0.00378 | 0.87 | | | |
| ISM1 | isthmin 1(ISM1) | NM_080826 | 93.4 | 114 | 0.00384 | 1.189 | | | |
| GORAB | golgin, RAB6 interacting(GORAB) | NM_152281 | 114.4 | 130.8 | 0.00439 | 1.118 | | | |
| NUAK1 | NUAK family kinase 1(NUAK1) | NM_014840 | 266.5 | 438.6 | 0.00475 | 1.605 | | | |

| | | | | | | |
|---|---|---|---|---|---|---|
| MRPL34 | mitochondrial ribosomal protein L34(MRPL34) | NM_023937 | 399.5 | 330.5 | 0.00481 | 0.823 |
| | | XM_044178 | 107.6 | 90.6 | 0.00495 | 0.832 |
| POSTN | periostin(POSTN) | NM_006475 | 91.7 | 128.2 | 0.00508 | 1.367 |
| NFASC | neurofascin(NFASC) | NM_001005387 | 102.8 | 118.8 | 0.0052 | 1.126 |
| CPXM1 | carboxypeptidase X, M14 family member 1(CPXM1) | NM_019609 | 116.2 | 154.4 | 0.00551 | 1.302 |
| PCSK5 | proprotein convertase subtilisin/kexin type 5(PCSK5) | NM_006200 | 111.9 | 151.8 | 0.00588 | 1.325 |
| CRK | CRK proto-oncogene, adaptor protein(CRK) | NM_016823 | 150.3 | 166.2 | 0.00589 | 1.287 |
| CRK | CRK proto-oncogene, adaptor protein(CRK) | NM_016823 | 153.7 | 144.3 | 0.00589 | 1.287 |
| CRK | CRK proto-oncogene, adaptor protein(CRK) | NM_016823 | 260.4 | 339.6 | 0.00589 | 1.287 |
| COL6A3 | collagen type VI alpha 3 chain(COL6A3) | NM_057165 | 305 | 592.9 | 0.00608 | 1.983 |
| ECM2 | extracellular matrix protein 2(ECM2) | NM_001393 | 105.9 | 139.8 | 0.0061 | 1.292 |
| ECM2 | extracellular matrix protein 2(ECM2) | NM_001393 | 204.9 | 348.7 | 0.0061 | 1.292 |
| NSDHL | NAD(P) dependent steroid dehydrogenase-like(NSDHL) | NM_015922 | 185.9 | 232.8 | 0.00695 | 1.235 |

| | | | | | | |
|---|---|---|---|---|---|---|
| PMEPA1 | prostate transmembrane protein, androgen induced 1(PMEPA1) | NM_199169 | 119.3 | 126.9 | 0.0071 | 1.383 |
| PMEPA1 | prostate transmembrane protein, androgen induced 1(PMEPA1) | NM_199169 | 134 | 189.8 | 0.0071 | 1.383 |
| KCNH2 | potassium voltage-gated channel subfamily H member 2(KCNH2) | NM_172056 | 108.1 | 127.8 | 0.00748 | 1.153 |
| THBS2 | thrombospondin 2(THBS2) | NM_003247 | 143.9 | 291.4 | 0.00795 | 1.976 |
| | | XM_001724468 | 114.1 | 132.4 | 0.00859 | 1.129 |
| OLFML2B | olfactomedin like 2B(OLFML2B) | NM_015441 | 132.8 | 169.1 | 0.00893 | 1.242 |
| SRPX | sushi repeat containing protein, X-linked(SRPX) | NM_006307 | 165.6 | 376.1 | 0.00915 | 2.23 |
| YIF1B | Yip1 interacting factor homolog B, membrane trafficking protein(YIF1B) | NM_033557 | 124 | 110.9 | 0.00937 | 0.876 |
| MPV17L | MPV17 mitochondrial inner membrane protein like(MPV17L) | NM_173803 | 109.6 | 95.4 | 0.00967 | 0.858 |
| | | AA447555 | 118.4 | 99.9 | 0.00973 | 0.827 |
| XRCC5 | X-ray repair cross complementing 5(XRCC5) | NM_021141 | 161.5 | 137.8 | 0.00996 | 0.845 |

| | | | | | | |
|---|---|---|---|---|---|---|
| | | XM_927228 | 109.4 | 124.3 | 0.01005 | 1.111 |
| | | DB302602 | 106.4 | 130.3 | 0.0105 | 1.194 |
| TBC1D8B | TBC1 domain family member 8B(TBC1D8B) | NM_017752 | 113.6 | 108.2 | 0.01075 | 0.82 |
| TBC1D8B | TBC1 domain family member 8B(TBC1D8B) | NM_017752 | 176.8 | 144.8 | 0.01075 | 0.82 |
| ZNF467 | zinc finger protein 467(ZNF467) | NM_207336 | 264.2 | 196.8 | 0.01105 | 0.755 |
| COL5A1 | collagen type V alpha 1 chain(COL5A1) | NM_000093 | 107.3 | 135.9 | 0.01115 | 1.241 |
| PLA2G2E | phospholipase A2 group IIE(PLA2G2E) | NM_014589 | 117.2 | 107.4 | 0.01122 | 0.894 |
| PPM1A | protein phosphatase, Mg2+/Mn2+ dependent 1A(PPM1A) | NM_177952 | 108.1 | 128.4 | 0.01159 | 1.16 |
| PPM1A | protein phosphatase, Mg2+/Mn2+ dependent 1A(PPM1A) | NM_177952 | 216 | 207 | 0.01159 | 1.16 |
| | | XM_0017199616 | 115.5 | 103.7 | 0.0116 | 0.875 |
| SSRP1 | structure specific recognition protein 1(SSRP1) | NM_003146 | 101.1 | 119.6 | 0.01178 | 1.148 |
| hspe1 | heat shock protein family E (Hsp10) member 1(HSPE1) | NM_002157 | 203.9 | 299.6 | 0.01219 | 1.495 |
| hspe1 | heat shock protein family E (Hsp10) member 1(HSPE1) | NM_002157 | 1594.2 | 1583.5 | 0.01219 | 1.495 |

| | | | | | | |
|---|---|---|---|---|---|---|
| MT2A | metallothionein 2A(MT2A) | NM_005953 | 1492.9 | 2670.6 | 0.01243 | 1.87 |
| LINC00560 | long intergenic non-protein coding RNA 560(LINC00560) | BC043278 | 99.1 | 116.9 | 0.01259 | 1.142 |
| | | XM_928029 | 152.5 | 132.6 | 0.0127 | 0.861 |
| MS4A3 | membrane spanning 4-domains A3(MS4A3) | NM_001031666 | 103.5 | 87.4 | 0.0129 | 0.824 |
| ADORA1 | adenosine A1 receptor(ADORA1) | NM_000674 | 112.9 | 113.5 | 0.0132 | 1.264 |
| ADORA1 | adenosine A1 receptor(ADORA1) | NM_000674 | 177.8 | 227.4 | 0.0132 | 1.264 |
| | | XM_209824 | 111 | 125 | 0.0134 | 1.097 |
| HNRNPUL2 | heterogeneous nuclear ribonucleoprotein U like 2(HNRNPUL2) | NM_001079559 | 117.9 | 106.5 | 0.0135 | 0.885 |
| | | XM_929413 | 122.6 | 110.3 | 0.01383 | 0.884 |
| PDGFRL | platelet derived growth factor receptor like(PDGFRL) | NM_006207 | 117.9 | 143.4 | 0.01399 | 1.186 |
| | | CN304166 | 104.5 | 120.1 | 0.01405 | 1.118 |
| CILP | cartilage intermediate layer protein(CILP) | NM_003613 | 115.1 | 117.1 | 0.01464 | 1.504 |
| CILP | cartilage intermediate layer protein(CILP) | NM_003613 | 141 | 218.5 | 0.01464 | 1.504 |

| | | | | | | | | | |
|---|---|---|---|---|---|---|---|---|---|
| NAA60 | N(alpha)-acetyltransferase 60, NatF catalytic subunit(NAA60) | NM_001083601 | 250.3 | 198.7 | 0.01467 | 0.804 | | | |
| EIF2S1 | eukaryotic translation initiation factor 2 subunit alpha(EIF2S1) | NM_004094 | 101.6 | 118.4 | 0.01484 | 1.14 | NM_053502 | 4.02 | 18.01 |
| LARS | leucyl-tRNA synthetase(LARS) | NM_020117 | 112.4 | 100.7 | 0.01492 | 0.884 | NM_053502 | 4.02 | 18.01 |
| TSHR | thyroid stimulating hormone receptor(TSHR) | NM_000369 | 109.8 | 139.1 | 0.01514 | 1.235 | NM_053502 | 4.02 | 18.01 |
| TSHR | thyroid stimulating hormone receptor(TSHR) | NM_000369 | 118.7 | 145.8 | 0.01514 | 1.235 | NM_053502 | 4.02 | 18.01 |
| KCNJ14 | potassium voltage-gated channel subfamily J member 14(KCNJ14) | NM_013348 | 123.1 | 112.7 | 0.01535 | 0.899 | | | |
| COL3A1 | collagen type III alpha 1 chain(COL3A1) | NM_000090 | 114 | 165.3 | 0.01547 | 1.425 | | | |
| TRIM51EP | tripartite motif-containing 51E, pseudogene(TRIM51EP) | NM_001136118 | 133.3 | 153.7 | 0.0157 | 1.125 | | | |
| PMP22 | peripheral myelin protein 22(PMP22) | NM_153321 | 140.7 | 211.1 | 0.01582 | 1.461 | | | |
| PMP22 | peripheral myelin protein 22(PMP22) | NM_153321 | 304.6 | 629.2 | 0.01582 | 1.461 | | | |

| | | | | | | | | | |
|---|---|---|---|---|---|---|---|---|---|
| SRRM2 | serine/arginine repetitive matrix 2(SRRM2) | NM_016333 | 2133.6 | 2612 | 0.01594 | 1.25 | | | |
| hamp | hepcidin antimicrobial peptide(HAMP) | NM_021175 | 136.1 | 114.2 | 0.01607 | 0.82 | NM_001009670 | 33.30 | 2.04 |
| TBC1D7 | TBC1 domain family member 7(TBC1D7) | NM_016495 | 188.3 | 231.6 | 0.01623 | 1.212 | | | |
| EFNA3 | ephrin A3(EFNA3) | NM_004952 | 153.1 | 134.1 | 0.01654 | 0.859 | | | |
| NRDC | nardilysin convertase(NRDC) | NM_002525 | 473.1 | 377.4 | 0.01664 | 0.808 | | | |
| CAPN6 | calpain 6(CAPN6) | NM_014289 | 100.2 | 117.3 | 0.01678 | 1.141 | | | |
| GPAM | glycerol-3-phosphate acyltransferase, mitochondrial(GPAM) | NM_020918 | 95.4 | 123.8 | 0.0168 | 3.075 | | | |
| GPAM | glycerol-3-phosphate acyltransferase, mitochondrial(GPAM) | NM_020918 | 195.7 | 626.4 | 0.0168 | 3.075 | | | |
| ABCA8 | ATP binding cassette subfamily A member 8(ABCA8) | NM_007168 | 202.1 | 315.8 | 0.01685 | 1.509 | | | |
| HAVCR1P1 | hepatitis A virus cellular receptor 1 pseudogene 1(HAVCR1P1) | NR_003603 | 107.9 | 181.6 | 0.01745 | 1.577 | | | |
| HAVCR1P1 | hepatitis A virus cellular receptor 1 pseudogene 1(HAVCR1P1) | NR_003603 | 115.5 | 149.5 | 0.01745 | 1.577 | | | |
| | | XM_927208 | 140.8 | 125.7 | 0.01803 | 0.877 | | | |

| | | | | | | |
|---|---|---|---|---|---|---|
| Hpvc1 | human papillomavirus (type 18) E5 central sequence-like 1(HPVC1) | NR_004422 | 111.2 | 103.6 | 0.01808 | 0.915 |
| HACE1 | HECT domain and ankyrin repeat containing E3 ubiquitin protein ligase 1(HACE1) | NM_020771 | 127.4 | 113.4 | 0.01813 | 0.878 |
| | | XM_939334 | 138 | 123.1 | 0.01831 | 0.878 |
| ECM2 | extracellular matrix protein 2(ECM2) | NM_001393 | 105.9 | 139.8 | 0.01923 | 1.67 |
| COL6A5 | collagen type VI alpha 5 chain(COL6A5) | NM_153264 | 104.1 | 123.1 | 0.01926 | 1.149 |
| COL6A5 | collagen type VI alpha 5 chain(COL6A5) | NM_153264 | 109.8 | 111.8 | 0.01926 | 1.149 |
| SKP2 | S-phase kinase associated protein 2(SKP2) | NM_032637 | 103.7 | 126.8 | 0.01927 | 1.193 |
| TOB1 | transducer of ERBB2, 1(TOB1) | NM_005749 | 1031.1 | 568.8 | 0.01938 | 0.556 |
| ASB8 | ankyrin repeat and SOCS box containing 8(ASB8) | NM_024095 | 118.5 | 110.9 | 0.02023 | 0.913 |
| ASB8 | ankyrin repeat and SOCS box containing 8(ASB8) | NM_024095 | 121.6 | 105.5 | 0.02023 | 0.913 |

| | | | | | | |
|---|---|---|---|---|---|---|
| AGPAT2 | 1-acylglycerol-3-phosphate O-acyltransferase 2(AGPAT2) | NM_006412 | 116 | 160.4 | 0.02024 | 1.347 |
| FASTK | Fas activated serine/threonine kinase(FASTK) | NM_033015 | 120.1 | 113.9 | 0.02038 | 0.86 |
| FASTK | Fas activated serine/threonine kinase(FASTK) | NM_033015 | 212.1 | 183 | 0.02038 | 0.86 |
| ASAH1 | N-acylsphingosine amidohydrolase 1(ASAH1) | NM_177924 | 122.3 | 113.9 | 0.02039 | 0.912 |
| GPD1 | glycerol-3-phosphate dehydrogenase 1(GPD1) | NM_005276 | 101.2 | 125 | 0.02046 | 1.207 |
| MFAP5 | microfibrillar associated protein 5(MFAP5) | NM_003480 | 172.5 | 294.7 | 0.02054 | 1.67 |
| C1orf87 | chromosome 1 open reading frame 87(C1orf87) | NM_152377 | 101.7 | 98.6 | 0.02062 | 0.893 |
| C1orf87 | chromosome 1 open reading frame 87(C1orf87) | NM_152377 | 115.4 | 105.5 | 0.02062 | 0.893 |
| | | XM_944183 | 114.3 | 126.6 | 0.02101 | 1.083 |
| TRIM45 | tripartite motif containing 45(TRIM45) | NM_025188 | 161 | 145.9 | 0.02102 | 0.894 |
| COL1A1 | collagen type I alpha 1 chain(COL1A1) | NM_000088 | 225.5 | 1069.5 | 0.02116 | 4.674 |

| | | | | | | |
|---|---|---|---|---|---|---|
| | | AW135045 | 148.4 | 136.1 | 0.02128 | 0.902 |
| PXK | PX domain containing serine/threonine kinase like(PXK) | NM_017771 | 190.4 | 157.8 | 0.02146 | 0.831 |
| | | XM_938510 | 106.5 | 126 | 0.0215 | 1.151 |
| | | XM_925919 | 112 | 101.5 | 0.02158 | 0.883 |
| | | XM_941821 | 117.9 | 129.5 | 0.02159 | 1.074 |
| MCMDC2 | minichromosome maintenance domain containing 2(MCMDC2) | NM_173518 | 113.7 | 115.6 | 0.02195 | 1.363 |
| MCMDC2 | minichromosome maintenance domain containing 2(MCMDC2) | NM_173518 | 3072.3 | 4141 | 0.02195 | 1.363 |
| MMADHC | methylmalonic aciduria and homocystinuria, cblD type(MMADHC) | NM_015702 | 232.8 | 223.2 | 0.02229 | 0.753 |
| MMADHC | methylmalonic aciduria and homocystinuria, cblD type(MMADHC) | NM_015702 | 637.9 | 568.3 | 0.02229 | 0.753 |
| MMADHC | methylmalonic aciduria and homocystinuria, cblD type(MMADHC) | NM_015702 | 659.1 | 482.7 | 0.02229 | 0.753 |
| COL1A2 | collagen type I alpha 2 chain(COL1A2) | NM_000089 | 184.3 | 825.5 | 0.02232 | 3.163 |
| COL1A2 | collagen type I alpha 2 chain(COL1A2) | NM_000089 | 499.5 | 1554.2 | 0.02232 | 3.163 |

| | | | | | | |
|---|---|---|---|---|---|---|
| | | XM_001726262 | 98.8 | 114.1 | 0.02255 | 1.12 |
| TUBA3FP | tubulin alpha 3f pseudogene(TUBA3FP) | NR_003608 | 120.7 | 112.3 | 0.02259 | 0.91 |
| FBLN2 | fibulin 2(FBLN2) | NM_001998 | 115.3 | 182.4 | 0.02275 | 2.255 |
| FBLN2 | fibulin 2(FBLN2) | NM_001998 | 181 | 416.8 | 0.02275 | 2.255 |
| | | XM_001134012 | 208.6 | 459.5 | 0.02279 | 2.21 |
| FBLN2 | fibulin 2(FBLN2) | NM_001998 | 115.3 | 182.4 | 0.02301 | 1.546 |
| PI15 | peptidase inhibitor 15(PI15) | NM_015886 | 113.6 | 109.2 | 0.02312 | 1.222 |
| PI15 | peptidase inhibitor 15(PI15) | NM_015886 | 121 | 151 | 0.02312 | 1.222 |
| | | XM_927196 | 98.2 | 112.7 | 0.0234 | 1.124 |
| MIR591 | microRNA 591(MIR591) | NR_030322 | 144.2 | 126 | 0.02363 | 0.861 |
| COG4 | component of oligomeric golgi complex 4(COG4) | NM_015386 | 103.1 | 121.6 | 0.02364 | 1.15 |
| | | AK129542 | 131.2 | 120.2 | 0.02374 | 0.902 |
| GSTM5 | glutathione S-transferase mu 5(GSTM5) | NM_000851 | 179.1 | 262.6 | 0.02394 | 1.442 |
| CCDC78 | coiled-coil domain containing 78(CCDC78) | NM_173476 | 141.9 | 122.2 | 0.02415 | 0.843 |

| | | | | | | |
|---|---|---|---|---|---|---|
| ELOVL5 | ELOVL fatty acid elongase 5(ELOVL5) | NM_021814 | 103.7 | 133.2 | 0.02431 | 1.256 |
| PPTC7 | PTC7 protein phosphatase homolog(PPTC7) | NM_139283 | 282 | 207.2 | 0.0247 | 0.726 |
| SLC29A1 | solute carrier family 29 member 1 (Augustine blood group)(SLC29A1) | NM_001078174 | 122.3 | 137.1 | 0.02484 | 1.314 |
| SLC29A1 | solute carrier family 29 member 1 (Augustine blood group)(SLC29A1) | NM_001078174 | 162.9 | 217.2 | 0.02484 | 1.314 |
| CTHRC1 | collagen triple helix repeat containing 1(CTHRC1) | NM_138455 | 104.2 | 140.4 | 0.02514 | 1.375 |
| CTHRC1 | collagen triple helix repeat containing 1(CTHRC1) | NM_138455 | 119 | 165.9 | 0.02514 | 1.375 |
| UBC | ubiquitin C(UBC) | NM_021009 | 10602.5 | 9083.6 | 0.02538 | 0.818 |
| UBC | ubiquitin C(UBC) | NM_021009 | 12020.7 | 12013.2 | 0.02538 | 0.818 |
| | | XM_001718145 | 121.7 | 109.3 | 0.0256 | 0.88 |
| SOX2 | SRY-box 2(SOX2) | NM_003106 | 91.1 | 108 | 0.02573 | 1.154 |
| DACH1 | dachshund family transcription factor 1(DACH1) | NM_080759 | 138.1 | 170.6 | 0.02581 | 1.195 |

| | | | | | | | | | |
|---|---|---|---|---|---|---|---|---|---|
| YBEY | ybeY metallopeptidase (putative)(YBEY) | NM_058181 | 104.5 | 105.4 | 0.02612 | 0.81 | | | |
| YBEY | ybeY metallopeptidase (putative)(YBEY) | NM_058181 | 297.8 | 252.3 | 0.02612 | 0.81 | | | |
| YBEY | ybeY metallopeptidase (putative)(YBEY) | NM_058181 | 301 | 246 | 0.02612 | 0.81 | | | |
| RNASE3 | ribonuclease A family member 3(RNASE3) | NM_002935 | 84.8 | 183.7 | 0.0262 | 2.088 | | | |
| | | XM_001126483 | 127.7 | 115.4 | 0.02629 | 0.886 | | | |
| ANXA1 | annexin A1(ANXA1) | NM_000700 | 270.9 | 944.9 | 0.02645 | 3.411 | | | |
| THRSP | thyroid hormone responsive(THRSP) | NM_003251 | 110.3 | 389 | 0.02685 | 3.374 | NM_001107943 | 0.63 | 112.77 |
| EP400NL | EP400 N-terminal like(EP400NL) | NM_182613 | 96.8 | 110.7 | 0.02693 | 1.118 | | | |
| | | XM_001726438 | 141.4 | 158.5 | 0.02724 | 1.097 | | | |
| POLR2J3 | RNA polymerase II subunit J3(POLR2J3) | NM_001097615 | 525.7 | 409.8 | 0.02754 | 0.774 | | | |
| HPX | hemopexin(HPX) | NM_000613 | 145.1 | 109.6 | 0.0279 | 0.737 | | | |
| TOPBP1 | topoisomerase (DNA) II binding protein 1(TOPBP1) | NM_007027 | 110.4 | 133.2 | 0.02799 | 1.18 | | | |
| FASN | fatty acid synthase(FASN) | NM_004104 | 337 | 1891.8 | 0.02802 | 5.576 | | | |
| COL1A2 | collagen type I alpha 2 chain(COL1A2) | NM_000089 | 184.3 | 825.5 | 0.02838 | 4.39 | | | |

| | | | | | | | | | |
|---|---|---|---|---|---|---|---|---|---|
| PGLYRP2 | peptidoglycan recognition protein 2(PGLYRP2) | NM_052890 | 121.6 | 106.1 | 0.02847 | 0.852 | | | |
| RERE | arginine-glutamic acid dipeptide repeats(RERE) | NM_012102 | 320.9 | 368.4 | 0.02849 | 1.153 | | | |
| TRIM10 | tripartite motif containing 10(TRIM10) | NM_006778 | 123.6 | 115.6 | 0.02856 | 0.917 | | | |
| TVP23B | trans-golgi network vesicle protein 23 homolog B(TVP23B) | NM_016078 | 108.5 | 95.1 | 0.02884 | 0.862 | | | |
| TVP23B | trans-golgi network vesicle protein 23 homolog B(TVP23B) | NM_016078 | 139.9 | 133.8 | 0.02884 | 0.862 | | | |
| | | XM_933591 | 118.5 | 104.7 | 0.0289 | 0.867 | | | |
| | | XR_019367 | 144.5 | 121.6 | 0.02909 | 0.831 | NM_134372 | 22.51 | 0.03 |
| DDX19B | DEAD-box helicase 19B(DDX19B) | NM_001014451 | 97.9 | 112.5 | 0.02916 | 1.12 | | | |
| DDX19B | DEAD-box helicase 19B(DDX19B) | NM_001014451 | 184.1 | 187.7 | 0.02916 | 1.12 | | | |
| PPARG | peroxisome proliferator activated receptor gamma(PPARG) | NM_015869 | 90.1 | 115.6 | 0.02917 | 1.247 | | | |
| PPARG | peroxisome proliferator activated receptor gamma(PPARG) | NM_015869 | 225.9 | 651.8 | 0.02917 | 1.247 | | | |
| CCKAR | cholecystokinin A receptor(CCKAR) | NM_000730 | 109.5 | 125.2 | 0.0293 | 1.116 | | | |

| | | | | | | |
|---|---|---|---|---|---|---|
| FABP5 | fatty acid binding protein 5(FABP5) | NM_001444 | 95 | 136.3 | 0.0293 | 1.398 |
| FABP5 | fatty acid binding protein 5(FABP5) | NM_001444 | 240.5 | 510.9 | 0.0293 | 1.398 |
| SPRY1 | sprouty RTK signaling antagonist 1(SPRY1) | NM_199327 | 500.4 | 756.6 | 0.02948 | 1.547 |
| | | XM_001722644 | 116.6 | 104.8 | 0.02949 | 0.88 |
| NDUFB11 | NADH:ubiquinone oxidoreductase subunit B11(NDUFB11) | NM_019056 | 1498.6 | 1252.4 | 0.02956 | 0.842 |
| | | XM_943964 | 119.2 | 107.1 | 0.02975 | 0.879 |
| | | XM_294093 | 105 | 120.4 | 0.02978 | 1.117 |
| KLF10 | Kruppel like factor 10(KLF10) | NM_005655 | 154.5 | 141.3 | 0.02987 | 0.832 |
| KLF10 | Kruppel like factor 10(KLF10) | NM_005655 | 192.8 | 157.9 | 0.02987 | 0.832 |
| LGALS1 | galectin 1(LGALS1) | NM_002305 | 681.2 | 1942.4 | 0.0299 | 2.938 |
| CAV2 | caveolin 2(CAV2) | NM_001233 | 203.8 | 356.4 | 0.02991 | 1.732 |
| CAV2 | caveolin 2(CAV2) | NM_001233 | 258 | 425.7 | 0.02991 | 1.732 |
| FABP4 | fatty acid binding protein 4(FABP4) | NM_001442 | 282.6 | 6897.1 | 0.03005 | 24.735 |
| DYNAP | dynactin associated protein(DYNAP) | NM_173629 | 116 | 106.5 | 0.03015 | 0.899 |

| | | | | | | | | | |
|---|---|---|---|---|---|---|---|---|---|
| ATP5G2 | ATP synthase, H+ transporting, mitochondrial Fo complex subunit C2 (subunit 9)(ATP5G2) | NM_005176 | 224.2 | 182.4 | 0.03039 | 0.787 | | | |
| ATP5G2 | ATP synthase, H+ transporting, mitochondrial Fo complex subunit C2 (subunit 9)(ATP5G2) | NM_005176 | 2815.4 | 2479.8 | 0.03039 | 0.787 | | | |
| | | DB337880 | 102.2 | 119.2 | 0.03082 | 1.138 | | | |
| PON3 | paraoxonase 3(PON3) | NM_000940 | 111.4 | 97.8 | 0.03097 | 0.857 | NM_001031645 | 1.93 | 10.04 |
| CTHRC1 | collagen triple helix repeat containing 1(CTHRC1) | NM_138455 | 104.2 | 140.4 | 0.03101 | 1.317 | | | |
| AMBN | ameloblastin(AMBN) | NM_016519 | 108.1 | 99.6 | 0.03109 | 0.902 | | | |
| | | AI446524 | 98.8 | 116.4 | 0.03117 | 1.144 | | | |
| | | XM_932043 | 118.7 | 104.5 | 0.03119 | 0.865 | | | |
| VN1R5 | vomeronasal 1 receptor 5 (gene/pseudogene)(VN1R5) | NM_173858 | 139.3 | 123.6 | 0.03126 | 0.871 | | | |
| FBXL8 | F-box and leucine rich repeat protein 8(FBXL8) | NM_018378 | 128.1 | 152.9 | 0.0314 | 1.162 | | | |
| | | AI698256 | 112.9 | 135.6 | 0.03157 | 1.174 | | | |

| | | | | | | | | | |
|---|---|---|---|---|---|---|---|---|---|
| OPN1SW | opsin 1 (cone pigments), short-wave-sensitive(OPN1SW) | NM_001708 | 99.1 | 119.9 | 0.03174 | 1.179 | | | |
| | | XR_037520 | 117.1 | 132.8 | 0.03186 | 1.107 | | | |
| THYN1 | thymocyte nuclear protein 1(THYN1) | NM_199297 | 140.8 | 176.6 | 0.03214 | 1.23 | | | |
| CEACAM1 | carcinoembryonic antigen related cell adhesion molecule 1(CEACAM1) | NM_001712 | 120 | 105 | 0.03226 | 0.858 | | | |
| CPXM2 | carboxypeptidase X, M14 family member 2(CPXM2) | NM_198148 | 172.8 | 232.9 | 0.03227 | 1.31 | | | |
| DSG2 | desmoglein 2(DSG2) | NM_001943 | 121.4 | 110.8 | 0.03236 | 0.891 | | | |
| GID4 | GID complex subunit 4 homolog(GID4) | NM_024052 | 99.4 | 115 | 0.03247 | 1.122 | | | |
| GID4 | GID complex subunit 4 homolog(GID4) | NM_024052 | 131.9 | 129.3 | 0.03247 | 1.122 | NM_144748 | 66.99 | 0.14 |
| | | XM_933382 | 133.4 | 113.7 | 0.03258 | 0.834 | | | |
| HEYL | hes related family bHLH transcription factor with YRPW motif-like(HEYL) | NM_014571 | 168.8 | 269.8 | 0.03269 | 1.587 | | | |
| GSN | gelsolin(GSN) | NM_198252 | 209.4 | 641.6 | 0.03271 | 2.981 | | | |
| SFRP5 | secreted frizzled related protein 5(SFRP5) | NM_003015 | 115.7 | 135.1 | 0.03275 | 1.137 | | | |

| | | | | | | | | | |
|---|---|---|---|---|---|---|---|---|---|
| TIPARP | TCDD inducible poly(ADP-ribose) polymerase(TIPARP) | NM_015508 | 396.4 | 317 | 0.03275 | 0.827 | | | |
| MT1IP | metallothionein 1I, pseudogene(MT1IP) | NM_175621 | 1211.2 | 2412.2 | 0.03281 | 2.159 | | | |
| NBPF11 | neuroblastoma breakpoint family member 11(NBPF11) | NM_183372 | 110 | 121.4 | 0.03288 | 1.078 | | | |
| | | XM_940358 | 99.1 | 113.2 | 0.03291 | 1.106 | | | |
| MID1IP1 | MID1 interacting protein 1(MID1IP1) | NM_021242 | 116 | 116.2 | 0.03294 | 0.815 | NM_001107793 | 77.29 | 6.85 |
| MID1IP1 | MID1 interacting protein 1(MID1IP1) | NM_021242 | 199.5 | 159.5 | 0.03294 | 0.815 | NM_001107793 | 77.29 | 6.85 |
| | | XM_931329 | 118.7 | 106.5 | 0.03309 | 0.879 | NM_001107793 | 77.29 | 6.85 |
| prdm9 | PR/SET domain 9(PRDM9) | NM_020227 | 114 | 126.6 | 0.03339 | 1.088 | NM_001107793 | 77.29 | 6.85 |
| CALR | calreticulin(CALR) | NM_004343 | 203.4 | 172.4 | 0.03356 | 0.848 | | | |
| HSPB2 | heat shock protein family B (small) member 2(HSPB2) | NM_001541 | 116.3 | 101.6 | 0.03388 | 0.858 | | | |
| FABP5 | fatty acid binding protein 5(FABP5) | NM_001444 | 95 | 136.3 | 0.03391 | 2.085 | | | |
| GAS1 | growth arrest specific 1(GAS1) | NM_002048 | 271 | 394.7 | 0.034 | 1.42 | | | |

| | | | | | | |
|---|---|---|---|---|---|---|
| TDGP1 | thymine-DNA glycosylase pseudogene 1(TDGP1) | NR_024382 | 100.4 | 115.6 | 0.03429 | 1.123 |
| IKZF2 | IKAROS family zinc finger 2(IKZF2) | NM_016260 | 114.7 | 131.1 | 0.03431 | 1.12 |
| KCNJ4 | potassium voltage-gated channel subfamily J member 4(KCNJ4) | NM_152868 | 102.1 | 120.4 | 0.0344 | 1.139 |
| KCNJ4 | potassium voltage-gated channel subfamily J member 4(KCNJ4) | NM_152868 | 108.7 | 117.6 | 0.0344 | 1.139 |
| GCK | glucokinase(GCK) | NM_033508 | 86.1 | 106.6 | 0.03459 | 1.21 |
| CIDEC | cell death inducing DFFA like effector c(CIDEC) | NM_022094 | 104.1 | 301.9 | 0.03469 | 2.79 |
| CIDEC | cell death inducing DFFA like effector c(CIDEC) | NM_022094 | 105 | 119.1 | 0.03469 | 2.79 |
| GSTA1 | glutathione S-transferase alpha 1(GSTA1) | NM_145740 | 107.6 | 99.2 | 0.03469 | 0.683 |
| GSTA1 | glutathione S-transferase alpha 1(GSTA1) | NM_145740 | 148.6 | 104.7 | 0.03469 | 0.683 |
| LOC101927635 | uncharacterized LOC101927635(LOC101927635) | BG119374 | 122.9 | 109.5 | 0.03525 | 0.87 |

| | | | | | | |
|---|---|---|---|---|---|---|
| mrap | melanocortin 2 receptor accessory protein(MRAP) | NM_178817 | 114.3 | 198.3 | 0.03526 | 1.676 |
| RNASE2 | ribonuclease A family member 2(RNASE2) | NM_002934 | 92.9 | 130.1 | 0.03553 | 1.361 |
| | | XM_940764 | 98.8 | 117.4 | 0.03563 | 1.164 |
| fgl1 | fibrinogen like 1(FGL1) | NM_201553 | 167.9 | 116 | 0.03579 | 0.671 |
| COL6A2 | collagen type VI alpha 2 chain(COL6A2) | NM_001849 | 167.7 | 217 | 0.03627 | 1.269 |
| COL6A2 | collagen type VI alpha 2 chain(COL6A2) | NM_001849 | 189.9 | 306.3 | 0.03627 | 1.269 |
| CACNG2 | calcium voltage-gated channel auxiliary subunit gamma 2(CACNG2) | NM_006078 | 102.5 | 91.8 | 0.03629 | 0.879 |
| NCALD | neurocalcin delta(NCALD) | NM_032041 | 304.1 | 452 | 0.03661 | 1.512 |
| | | XR_038776 | 132.3 | 116.8 | 0.03678 | 0.866 |
| RNF146 | ring finger protein 146(RNF146) | NM_030963 | 117.2 | 104.1 | 0.0368 | 0.867 |
| RNF146 | ring finger protein 146(RNF146) | NM_030963 | 152.3 | 145.6 | 0.0368 | 0.867 |
| COL6A2 | collagen type VI alpha 2 chain(COL6A2) | NM_001849 | 167.7 | 217 | 0.03704 | 1.571 |
| | | XM_001716052 | 117.9 | 108.2 | 0.03716 | 0.902 |

| | | | | | | | | | |
|---|---|---|---|---|---|---|---|---|---|
| c12orf4 | chromosome 12 open reading frame 4(C12orf4) | NM_020374 | 140.5 | 125.3 | 0.03717 | 0.872 | | | |
| MT1P3 | metallothionein 1 pseudogene 3(MT1P3) | NM_080757 | 1313.1 | 2470.2 | 0.03767 | 1.995 | | | |
| NDUFA10 | NADH:ubiquinone oxidoreductase subunit A10(NDUFA10) | NM_004544 | 103.7 | 111.8 | 0.03787 | 1.25 | | | |
| NDUFA10 | NADH:ubiquinone oxidoreductase subunit A10(NDUFA10) | NM_004544 | 105.1 | 133.8 | 0.03787 | 1.25 | | | |
| GAGE4 | G antigen 4(GAGE4) | NM_001474 | 103 | 124.1 | 0.03808 | 1.171 | | | |
| MIR1185-1 | microRNA 1185-1(MIR1185-1) | NR_031575 | 138.9 | 155.8 | 0.03814 | 1.092 | | | |
| NUGGC | nuclear GTPase, germinal center associated(NUGGC) | NM_001010906 | 117.8 | 104.9 | 0.03852 | 0.872 | | | |
| ZNF585A | zinc finger protein 585A(ZNF585A) | NM_199126 | 137.4 | 117.7 | 0.0386 | 0.834 | | | |
| BPY2B | basic charge, Y-linked, 2B(BPY2B) | NM_001002760 | 111.9 | 101.8 | 0.03865 | 0.89 | NM_001009603 | 28.32 | 0.90 |
| GMFB | glia maturation factor beta(GMFB) | NM_004124 | 114.4 | 134.3 | 0.03872 | 1.145 | | | |
| igsf11 | immunoglobulin superfamily member 11(IGSF11) | NM_152538 | 121.3 | 145.2 | 0.03895 | 1.171 | | | |
| | | XM_938166 | 161.6 | 291.2 | 0.03911 | 1.759 | | | |

| | | | | | | |
|---|---|---|---|---|---|---|
| NLRP1 | NLR family pyrin domain containing 1(NLRP1) | NM_033007 | 113 | 105.6 | 0.03931 | 0.916 |
| | | XM_0011300020 | 130.6 | 118.2 | 0.0394 | 0.898 |
| PHF10 | PHD finger protein 10(PHF10) | NM_133325 | 120.3 | 140.2 | 0.03946 | 1.133 |
| CD36 | CD36 molecule(CD36) | NM_000072 | 183.5 | 1068.4 | 0.03979 | 5.653 |
| WWOX | WW domain containing oxidoreductase(WWOX) | NM_130791 | 123.6 | 141.3 | 0.03993 | 1.113 |
| ALB | albumin(ALB) | NM_000477 | 1072.9 | 336.1 | 0.03999 | 0.286 |
| ALB | albumin(ALB) | NM_000477 | 1474.5 | 661.3 | 0.03999 | 0.286 |
| FGF18 | fibroblast growth factor 18(FGF18) | NM_003862 | 119.7 | 111.1 | 0.04018 | 0.911 |
| TRAF4 | TNF receptor associated factor 4(TRAF4) | NM_004295 | 117.9 | 106.1 | 0.04081 | 0.879 |
| TNN | tenascin N(TNN) | NM_022093 | 113.7 | 137.7 | 0.04092 | 1.184 |
| CDKN2C | cyclin dependent kinase inhibitor 2C(CDKN2C) | NM_078626 | 151.8 | 336.4 | 0.04102 | 2.146 |
| | | XM_0017211172 | 177.4 | 313.4 | 0.04104 | 1.74 |
| PODN | podocan(PODN) | NM_153703 | 104.5 | 134.7 | 0.04122 | 1.26 |

| | | | | | | |
|---|---|---|---|---|---|---|
| FAM19A 4 | family with sequence similarity 19 member A4, C-C motif chemokine like(FAM19A4) | NM_0010 05527 | 103.4 | 115.9 | 0.04132 | 1.097 |
| FAM19A 4 | family with sequence similarity 19 member A4, C-C motif chemokine like(FAM19A4) | NM_0010 05527 | 106.9 | 121.3 | 0.04132 | 1.097 |
| | | XM_29098 5 | 103.8 | 126.5 | 0.04149 | 1.18 |
| GLRB | glycine receptor beta(GLRB) | NM_0008 24 | 146.3 | 255.4 | 0.04167 | 1.678 |
| TRPV4 | transient receptor potential cation channel subfamily V member 4(TRPV4) | NM_0216 25 | 132.3 | 110.5 | 0.04167 | 0.823 |
| | | XM_00172 6471 | 109.8 | 137.9 | 0.04173 | 1.23 |
| ORM1 | orosomucoid 1(ORM1) | NM_0006 07 | 141.1 | 97.6 | 0.04187 | 0.671 |
| TAB3 | TGF-beta activated kinase 1/MAP3K7 binding protein 3(TAB3) | NM_1527 87 | 99.3 | 113.8 | 0.04197 | 1.117 |
| | | XR_01765 9 | 96.9 | 113.8 | 0.042 | 1.133 |
| SLC1A2 | solute carrier family 1 member 2(SLC1A2) | NM_0041 71 | 118.3 | 106.1 | 0.04214 | 0.883 |
| PCBP2 | poly(rC) binding protein 2(PCBP2) | NM_0011 28913 | 115.9 | 101.3 | 0.04224 | 0.853 |

| | | | | | | |
|---|---|---|---|---|---|---|
| ZIC4 | Zic family member 4(ZIC4) | NM_032153 | 122.2 | 112.7 | 0.04233 | 0.905 |
| | | AW044615 | 109.8 | 100.2 | 0.04238 | 0.893 |
| TIGD6 | tigger transposable element derived 6(TIGD6) | NM_030953 | 121.5 | 114.4 | 0.04251 | 0.921 |
| LY9 | lymphocyte antigen 9(LY9) | NM_001033667 | 84.8 | 125.1 | 0.04273 | 1.441 |
| GC | GC, vitamin D binding protein(GC) | NM_000583 | 172 | 110 | 0.04291 | 0.619 |
| abl1 | ABL proto-oncogene 1, non-receptor tyrosine kinase(ABL1) | NM_005157 | 591.2 | 660.4 | 0.043 | 1.125 |
| PCYOX1 | prenylcysteine oxidase 1(PCYOX1) | NM_016297 | 132.5 | 160.6 | 0.04301 | 1.181 |
| PCYOX1 | prenylcysteine oxidase 1(PCYOX1) | NM_016297 | 505.8 | 655.5 | 0.04301 | 1.181 |
| CRCP | CGRP receptor component(CRCP) | NM_014478 | 116.7 | 110.1 | 0.04322 | 1.398 |
| CRCP | CGRP receptor component(CRCP) | NM_014478 | 2497.8 | 3480.4 | 0.04322 | 1.398 |
| FAM124A | family with sequence similarity 124 member A(FAM124A) | NM_145019 | 111.8 | 103.6 | 0.04331 | 0.906 |
| FGB | fibrinogen beta chain(FGB) | NM_005141 | 113.2 | 95.6 | 0.04334 | 0.827 |
| PTAR1 | protein prenyltransferase alpha subunit repeat containing 1(PTAR1) | NM_001099666 | 203.8 | 240 | 0.04337 | 1.149 |

| | | | | | | |
|---|---|---|---|---|---|---|
| FAM196A | family with sequence similarity 196 member A(FAM196A) | NM_001039762 | 85.1 | 111.8 | 0.04343 | 1.271 |
| FGG | fibrinogen gamma chain(FGG) | NM_000509 | 188.8 | 94.8 | 0.04344 | 0.479 |
| | | XM_944654 | 107.9 | 120.1 | 0.04361 | 1.086 |
| FAM3D | family with sequence similarity 3 member D(FAM3D) | NM_138805 | 135.4 | 120.8 | 0.04408 | 0.881 |
| GBGT1 | globoside alpha-1,3-N-acetylgalactosaminyltransferase 1(GBGT1) | NM_021996 | 102.9 | 119.4 | 0.04427 | 1.135 |
| CADM3 | cell adhesion molecule 3(CADM3) | NM_021189 | 102.8 | 120.9 | 0.04434 | 1.143 |
| AGPAT2 | 1-acylglycerol-3-phosphate O-acyltransferase 2(AGPAT2) | NM_001012727 | 138.4 | 213.5 | 0.04444 | 1.507 |
| | | NM_198289 | 121.8 | 596.8 | 0.0445 | 4.655 |
| | | AA906391 | 145.2 | 172.4 | 0.04457 | 1.17 |
| TGM5 | transglutaminase 5(TGM5) | NM_201631 | 104.3 | 116.6 | 0.04462 | 1.094 |
| CASKIN2 | CASK interacting protein 2(CASKIN2) | NM_020753 | 116.9 | 108 | 0.04484 | 0.905 |
| | | AK002088 | 91.1 | 105.7 | 0.045 | 1.129 |
| | | XM_0017225227 | 115 | 135.7 | 0.04521 | 1.147 |

| | | | | | | |
|---|---|---|---|---|---|---|
| PNPLA2 | patatin like phospholipase domain containing 2(PNPLA2) | NM_020376 | 282.3 | 634.6 | 0.04534 | 2.133 |
| NACA | nascent polypeptide-associated complex alpha subunit(NACA) | NM_005594 | 4000.5 | 3255.1 | 0.04555 | 0.822 |
| NACA | nascent polypeptide-associated complex alpha subunit(NACA) | NM_005594 | 5969.1 | 5262.2 | 0.04555 | 0.822 |
| METRN | meteorin, glial cell differentiation regulator(METRN) | NM_024042 | 129.9 | 108 | 0.04557 | 0.821 |
| JADE1 | jade family PHD finger 1(JADE1) | NM_024900 | 215 | 183.4 | 0.04598 | 0.844 |
| CIDEA | cell death-inducing DFFA-like effector a(CIDEA) | NM_001279 | 101.9 | 1110.9 | 0.04601 | 10.315 |
| ADGRB1 | adhesion G protein-coupled receptor B1(ADGRB1) | NM_001702 | 123.6 | 111.7 | 0.04606 | 0.883 |
| PLCL1 | phospholipase C like 1(PLCL1) | NM_006226 | 116.7 | 104.4 | 0.04609 | 0.884 |
| C14orf180 | chromosome 14 open reading frame 180(C14orf180) | NM_001008404 | 93.9 | 124.9 | 0.04611 | 1.3 |
| CAV1 | caveolin 1(CAV1) | NM_001753 | 157.8 | 503.2 | 0.04625 | 3.01 |
| CAV1 | caveolin 1(CAV1) | NM_001753 | 159 | 206.6 | 0.04625 | 3.01 |

| | | | | | | |
|---|---|---|---|---|---|---|
| ALDOC | aldolase, fructose-bisphosphate C(ALDOC) | NM_005165 | 342 | 1042.5 | 0.04629 | 2.928 |
| TDRD1 | tudor domain containing 1(TDRD1) | NM_198795 | 194.2 | 244.2 | 0.04638 | 1.388 |
| TDRD1 | tudor domain containing 1(TDRD1) | NM_198795 | 247.7 | 334.6 | 0.04638 | 1.388 |
| YTHDF2 | YTH N6-methyladenosine RNA binding protein 2(YTHDF2) | NM_016258 | 288.1 | 238.7 | 0.04649 | 0.845 |
| PSMD8 | proteasome 26S subunit, non-ATPase 8(PSMD8) | NM_002812 | 333.6 | 278.8 | 0.0467 | 0.828 |
| UBN2 | ubinuclein 2(UBN2) | NM_173569 | 161.5 | 187.1 | 0.04678 | 1.142 |
| UBN2 | ubinuclein 2(UBN2) | NM_173569 | 234.7 | 243 | 0.04678 | 1.142 |
| ADIRF | adipogenesis regulatory factor(ADIRF) | NM_006829 | 2009.5 | 3414.7 | 0.0469 | 1.753 |
| c2orf42 | chromosome 2 open reading frame 42(C2orf42) | NM_017880 | 125.3 | 110 | 0.04719 | 0.861 |
| MAR2 | membrane associated ring-CH-type finger 2(MARCH2) | NM_001005416 | 114.7 | 132 | 0.04764 | 1.121 |
| MAR2 | membrane associated ring-CH-type finger 2(MARCH2) | NM_001005416 | 154 | 163.8 | 0.04764 | 1.121 |

| | | | | | | |
|---|---|---|---|---|---|---|
| NPSR1-AS1 | NPSR1 antisense RNA 1(NPSR1-AS1) | NM_207285 | 136.8 | 127.8 | 0.04773 | 0.915 |
| | | XM_001130790 | 132.7 | 630.3 | 0.04805 | 4.52 |
| ZNF652 | zinc finger protein 652(ZNF652) | NM_014897 | 4290.7 | 5557.2 | 0.04818 | 1.309 |
| KCNV2 | potassium voltage-gated channel modifier subfamily V member 2(KCNV2) | NM_133497 | 102.7 | 117.1 | 0.04832 | 1.107 |
| | | XR_015597 | 94.3 | 112.9 | 0.04842 | 1.166 |
| HECTD2 | HECT domain E3 ubiquitin protein ligase 2(HECTD2) | NM_182765 | 137.6 | 138 | 0.04849 | 0.862 |
| HECTD2 | HECT domain E3 ubiquitin protein ligase 2(HECTD2) | NM_182765 | 143.7 | 126.4 | 0.04849 | 0.862 |
| TAF6 | TATA-box binding protein associated factor 6(TAF6) | NM_005641 | 113.1 | 126.7 | 0.04867 | 1.089 |
| | | XM_001131201 | 110.9 | 126.7 | 0.04884 | 1.106 |
| c1orf216 | chromosome 1 open reading frame 216(C1orf216) | NM_152374 | 123.1 | 139.7 | 0.04918 | 1.104 |
| | | XM_001732853 | 113.6 | 104.2 | 0.04945 | 0.896 |
| | | CK299443 | 155.9 | 140.5 | 0.04954 | 0.888 |

| | | | | | | | | | |
|---|---|---|---|---|---|---|---|---|---|
| PPARG | peroxisome proliferator activated receptor gamma(PPARG) | NM_138712 | 116.5 | 275.5 | 0.04973 | 2.294 | | | |
| PLTP | phospholipid transfer protein(PLTP) | NM_182676 | 205.3 | 322.8 | 0.04998 | 1.567 | | | |
| APOPT1 | apoptogenic 1, mitochondrial(APOPT1) | NM_032374 | 1407.8 | 1812.9 | 0.05 | 1.328 | | | |
| **Gene Symbol** | **Gene Name** | **Accession No.** | **IUGR Chow F Mean** | **IUGR HFHS F Mean** | **p-val IUGR F HFHS vs. IUGR F Chow** | **Ratio IUGR F HFHS vs. IUGR F Chow** | **REFSEQ ID** | **NEPHRON (MEAN RPKM)** | **COLLECTING DUCT (MEAN RPKM)** |
| | | XR_038951 | 520.6 | 329.4 | 0.00017 | 0.656 | | | |
| FNTB | farnesyltransferase, CAAX box, beta(FNTB) | NM_002028 | 399.5 | 281.9 | 0.00023 | 0.724 | | | |
| CCT7 | chaperonin containing TCP1 subunit 7(CCT7) | NM_006429 | 644.7 | 445.7 | 0.00104 | 0.745 | | | |
| ODC1 | ornithine decarboxylase 1(ODC1) | NM_002539 | 725.5 | 484.4 | 0.0011 | 0.726 | | | |
| | | XM_001719540 | 150.1 | 130.9 | 0.00111 | 0.859 | | | |
| ACTR5 | ARP5 actin-related protein 5 homolog(ACTR5) | NM_024855 | 134 | 122.4 | 0.00143 | 0.885 | | | |
| EIF3M | eukaryotic translation initiation factor 3 subunit M(EIF3M) | NM_006360 | 464.5 | 327.5 | 0.00153 | 0.655 | | | |

| | | | | | | | | | |
|---|---|---|---|---|---|---|---|---|---|
| EIF4A1 | eukaryotic translation initiation factor 4A1(EIF4A1) | NM_001416 | 586.5 | 372.4 | 0.00153 | 0.655 | | | |
| | | XM_934376 | 149.6 | 133.4 | 0.00194 | 0.875 | NM_017193 | 356.13 | 8.18 |
| FLJ41170 | uncharacterized LOC440200(FLJ41170) | NM_001004332 | 183.1 | 236.5 | 0.00197 | 1.391 | NM_017193 | 356.13 | 8.18 |
| | | XM_001713654 | 9306.3 | 6648.7 | 0.00201 | 0.835 | NM_017193 | 356.13 | 8.18 |
| RPS26P11 | ribosomal protein S26 pseudogene 11(RPS26P11) | NR_002309 | 9176.8 | 5893.9 | 0.00255 | 0.745 | NM_017193 | 356.13 | 8.18 |
| SMAGP | small cell adhesion glycoprotein(SMAGP) | NM_001033873 | 235.3 | 188.5 | 0.00284 | 0.826 | | | |
| JAM2 | junctional adhesion molecule 2(JAM2) | NM_021219 | 130 | 158 | 0.003 | 1.202 | | | |
| RPS3A | ribosomal protein S3A(RPS3A) | NM_001006 | 8852.1 | 6542.5 | 0.00312 | 0.75 | | | |
| RPS3A | ribosomal protein S3A(RPS3A) | NM_001006 | 9567.1 | 7835.2 | 0.00312 | 0.75 | | | |
| RPS3A | ribosomal protein S3A(RPS3A) | NM_001006 | 9708 | 6189.4 | 0.00312 | 0.75 | | | |
| IFT52 | intraflagellar transport 52(IFT52) | NM_016004 | 203.8 | 268.1 | 0.00337 | 1.368 | | | |
| | | BU928253 | 128.8 | 153.7 | 0.00349 | 1.144 | | | |
| POMP | proteasome maturation protein(POMP) | NM_015932 | 536.2 | 386.2 | 0.00355 | 0.746 | | | |
| MSLNL | mesothelin-like(MSLNL) | NM_001025190 | 114.5 | 148.6 | 0.00364 | 1.22 | | | |

| | | | | | | |
|---|---|---|---|---|---|---|
| MSLNL | mesothelin-like(MSLNL) | NM_0010 25190 | 123.2 | 139.9 | 0.00364 | 1.22 |
| | | AA759151 | 111 | 135.2 | 0.00375 | 1.166 |
| | | XM_92751 1 | 128.4 | 119.5 | 0.00389 | 0.899 |
| PDPK1 | 3-phosphoinositide dependent protein kinase 1(PDPK1) | NM_0312 68 | 139.4 | 122.7 | 0.00392 | 0.869 |
| PDPK1 | 3-phosphoinositide dependent protein kinase 1(PDPK1) | NM_0312 68 | 574 | 543.4 | 0.00392 | 0.869 |
| | | CA433902 | 187.2 | 222.9 | 0.00423 | 1.229 |
| PPP1CC | protein phosphatase 1 catalytic subunit gamma(PPP1CC) | NM_0027 10 | 2359.4 | 1509.3 | 0.00427 | 0.709 |
| | | XM_94395 6 | 110.5 | 132.6 | 0.00429 | 1.135 |
| TRIM36 | tripartite motif containing 36(TRIM36) | NM_0010 17397 | 133 | 157 | 0.00445 | 1.145 |
| CLUL1 | clusterin like 1(CLUL1) | NM_0144 10 | 325.6 | 432.9 | 0.0045 | 1.482 |
| NETO1 | neuropilin and tolloid like 1(NETO1) | AK123319 | 136.2 | 119.8 | 0.00453 | 0.846 |
| SNORA1 | small nucleolar RNA, H/ACA box 1(SNORA1) | NR_00302 6 | 128.2 | 113.9 | 0.00483 | 0.853 |
| | | XM_94178 5 | 106.4 | 126.8 | 0.00491 | 1.122 |
| NGEF | neuronal guanine nucleotide exchange factor(NGEF) | NM_0198 50 | 164.2 | 201.6 | 0.00498 | 1.231 |

| | | | | | | |
|---|---|---|---|---|---|---|
| MT1H | metallothionein 1H(MT1H) | NM_005951 | 97.7 | 143.8 | 0.00502 | 1.354 |
| MT1H | metallothionein 1H(MT1H) | NM_005951 | 1671.5 | 4509.3 | 0.00502 | 1.354 |
| | | XR_018412 | 132.9 | 120.8 | 0.00508 | 0.874 |
| MAIP1 | matrix AAA peptidase interacting protein 1(MAIP1) | NM_024520 | 232.1 | 168.4 | 0.00533 | 0.726 |
| CREB1 | cAMP responsive element binding protein 1(CREB1) | NM_004379 | 119.2 | 126.4 | 0.00542 | 1.623 |
| CREB1 | cAMP responsive element binding protein 1(CREB1) | NM_004379 | 1238.6 | 1821.8 | 0.00542 | 1.623 |
| CREB1 | cAMP responsive element binding protein 1(CREB1) | NM_004379 | 11307.8 | 13499 | 0.00542 | 1.623 |
| | | XM_938755 | 2625.7 | 1447 | 0.00543 | 0.639 |
| neurl4 | neuralized E3 ubiquitin protein ligase 4(NEURL4) | NM_032442 | 261 | 323.4 | 0.00549 | 1.313 |
| FAAP20 | Fanconi anemia core complex associated protein 20(FAAP20) | NM_182533 | 118.1 | 149.6 | 0.00564 | 1.211 |
| FAAP24 | Fanconi anemia core complex associated protein 24(FAAP24) | NM_152266 | 169.3 | 171.7 | 0.00564 | 1.211 |
| IARS | isoleucyl-tRNA synthetase(IARS) | NM_002161 | 2887.6 | 3967.8 | 0.00567 | 1.516 |

| | | | | | | |
|---|---|---|---|---|---|---|
| ZNF584 | zinc finger protein 584(ZNF584) | NM_173548 | 132.1 | 160.3 | 0.00573 | 1.165 |
| | | XM_001720172 | 722 | 461.2 | 0.00594 | 0.674 |
| USP21 | ubiquitin specific peptidase 21(USP21) | NM_012475 | 142.8 | 137.2 | 0.00597 | 0.822 |
| USP21 | ubiquitin specific peptidase 21(USP21) | NM_012475 | 217.3 | 175.3 | 0.00597 | 0.822 |
| BCL2 | BCL2, apoptosis regulator(BCL2) | NM_000633 | 138.4 | 127.4 | 0.00605 | 0.641 |
| BCL2L1 | BCL2 like 1(BCL2L1) | NM_138578 | 424 | 265.5 | 0.00605 | 0.641 |
| RFX5 | regulatory factor X5(RFX5) | NM_001025603 | 141.5 | 114.3 | 0.00609 | 0.79 |
| | | XR_037398 | 261 | 192.5 | 0.00621 | 0.731 |
| | | XR_037508 | 200.3 | 165.4 | 0.00641 | 0.84 |
| PITPNC1 | phosphatidylinositol transfer protein, cytoplasmic 1(PITPNC1) | NM_181671 | 201.7 | 153.3 | 0.00649 | 0.773 |
| POLE3 | DNA polymerase epsilon 3, accessory subunit(POLE3) | NM_017443 | 402.9 | 289.5 | 0.00651 | 0.744 |
| rbm12 | RNA binding motif protein 12(RBM12) | NM_006047 | 163.5 | 137.4 | 0.00683 | 0.832 |
| rbm12 | RNA binding motif protein 12(RBM12) | NM_006047 | 216.6 | 170.7 | 0.00683 | 0.832 |
| TMEM50B | transmembrane protein 50B(TMEM50B) | NM_006134 | 177.2 | 216.2 | 0.00683 | 1.283 |

| | | | | | | |
|---|---|---|---|---|---|---|
| TMEM50B | transmembrane protein 50B(TMEM50B) | NM_006134 | 568 | 512.4 | 0.00683 | 1.283 |
| MSMO1 | methylsterol monooxygenase 1(MSMO1) | NM_001017369 | 108.6 | 129.1 | 0.00689 | 1.131 |
| MSMO1 | methylsterol monooxygenase 1(MSMO1) | NM_001017369 | 230.5 | 219.5 | 0.00689 | 1.131 |
| | | XM_937648 | 134.9 | 162.9 | 0.0071 | 1.181 |
| SLC35A2 | solute carrier family 35 member A2(SLC35A2) | NM_001032289 | 139.6 | 129.3 | 0.00729 | 0.906 |
| SLC35A2 | solute carrier family 35 member A2(SLC35A2) | NM_001032289 | 409.1 | 388.8 | 0.00729 | 0.906 |
| cyb5r3 | cytochrome b5 reductase 3(CYB5R3) | NM_000398 | 170.6 | 150.8 | 0.00731 | 0.877 |
| | | XM_0011128786 | 505.2 | 1486.6 | 0.00756 | 3.49 |
| MIR1253 | microRNA 1253(MIR1253) | NR_031654 | 191.4 | 221.4 | 0.00782 | 1.2 |
| VAV3 | vav guanine nucleotide exchange factor 3(VAV3) | NM_006113 | 163.6 | 141.1 | 0.00788 | 0.854 |
| VAV3 | vav guanine nucleotide exchange factor 3(VAV3) | NM_006113 | 282.9 | 264.9 | 0.00788 | 0.854 |
| VAV3 | vav guanine nucleotide exchange factor 3(VAV3) | NM_006113 | 495.5 | 559.3 | 0.00788 | 0.854 |

| | | | | | | | | | |
|---|---|---|---|---|---|---|---|---|---|
| OR2G2 | olfactory receptor family 2 subfamily G member 2(OR2G2) | NM_001001915 | 167.2 | 202.2 | 0.00793 | 1.277 | | | |
| RPL10A | ribosomal protein L10a(RPL10A) | NM_007104 | 1974.5 | 1104.7 | 0.00797 | 0.627 | NM_053502 | 4.02 | 18.01 |
| RPL10A | ribosomal protein L10a(RPL10A) | NM_007104 | 6681.9 | 5179.3 | 0.00797 | 0.627 | NM_053502 | 4.02 | 18.01 |
| | | XR_015351 | 119.9 | 140.9 | 0.00801 | 1.154 | NM_053502 | 4.02 | 18.01 |
| DHX40 | DEAH-box helicase 40(DHX40) | NM_024612 | 172.7 | 146.6 | 0.00811 | 0.847 | NM_053502 | 4.02 | 18.01 |
| DHX40 | DEAH-box helicase 40(DHX40) | NM_024612 | 182.5 | 176.2 | 0.00811 | 0.847 | | | |
| DHX40 | DEAH-box helicase 40(DHX40) | NM_024612 | 196.8 | 215.6 | 0.00811 | 0.847 | | | |
| MIR1185-1 | microRNA 1185-1(MIR1185-1) | NR_031575 | 141.9 | 164.5 | 0.00818 | 1.135 | | | |
| | | XM_926231 | 244.2 | 159.5 | 0.00818 | 0.666 | | | |
| SMOC1 | SPARC related modular calcium binding 1(SMOC1) | NM_001034852 | 122.7 | 148.9 | 0.00865 | 1.179 | | | |
| SMOC1 | SPARC related modular calcium binding 1(SMOC1) | NM_001034852 | 132.9 | 136.2 | 0.00865 | 1.179 | | | |
| ANKRD10 | ankyrin repeat domain 10(ANKRD10) | NM_017664 | 229.6 | 204.5 | 0.00871 | 0.87 | NM_001009670 | 33.30 | 2.04 |
| ANKRA2 | ankyrin repeat family A member 2(ANKRA2) | NM_023039 | 150 | 135.5 | 0.00871 | 0.87 | | | |

| | | | | | | |
|---|---|---|---|---|---|---|
| | | XR_018394 | 1706 | 752.9 | 0.00882 | 0.475 |
| prorsd1p | prolyl-tRNA synthetase associated domain containing 1, pseudogene(PRORSD1P) | NM_001080459 | 157.9 | 134.1 | 0.00901 | 0.828 |
| TAOK2 | TAO kinase 2(TAOK2) | NM_016151 | 214.1 | 253.9 | 0.00906 | 1.273 |
| | | XR_038898 | 250 | 168.4 | 0.00906 | 0.659 |
| MCMDC2 | minichromosome maintenance domain containing 2(MCMDC2) | NM_173518 | 119.4 | 134.8 | 0.00946 | 1.382 |
| MCMDC2 | minichromosome maintenance domain containing 2(MCMDC2) | NM_173518 | 4139.8 | 4970.2 | 0.00946 | 1.382 |
| SLC35B2 | solute carrier family 35 member B2(SLC35B2) | NM_178148 | 147.6 | 173.3 | 0.00948 | 1.163 |
| GLE1 | GLE1, RNA export mediator(GLE1) | NM_001003722 | 300.9 | 399.2 | 0.00968 | 1.483 |
| PTPA | protein phosphatase 2 phosphatase activator(PTPA) | NM_178003 | 121.1 | 148.7 | 0.00975 | 1.167 |
| RHBDL2 | rhomboid like 2(RHBDL2) | NM_017821 | 146.7 | 182.8 | 0.00976 | 1.263 |
| RHBDL2 | rhomboid like 2(RHBDL2) | NM_017821 | 518.6 | 594.4 | 0.00976 | 1.263 |
| SEH1L | SEH1 like nucleoporin(SEH1L) | NM_001013437 | 209.3 | 160.1 | 0.00976 | 0.76 |

| | | | | | | |
|---|---|---|---|---|---|---|
| SEH1L | SEH1 like nucleoporin(SEH1L) | NM_001013437 | 273.8 | 261.4 | 0.00976 | 0.76 |
| | | XR_017380 | 139.3 | 114.5 | 0.00979 | 0.812 |
| EEF1B2 | eukaryotic translation elongation factor 1 beta 2(EEF1B2) | NM_001037663 | 7295.5 | 5026 | 0.0098 | 0.797 |
| | | XR_038867 | 179.9 | 142.7 | 0.00981 | 0.799 |
| | | XM_944646 | 156.4 | 177.6 | 0.01007 | 1.163 |
| | | XM_001132307 | 112.5 | 139.4 | 0.01019 | 1.177 |
| MUC4 | mucin 4, cell surface associated(MUC4) | NM_018406 | 115 | 137.6 | 0.01036 | 1.137 |
| DDX19B | DEAD-box helicase 19B(DDX19B) | NM_001014451 | 165 | 133.6 | 0.01038 | 0.799 |
| SP3 | Sp3 transcription factor(SP3) | NM_003111 | 131.7 | 118.4 | 0.01047 | 0.87 |
| | | XR_039278 | 2439.5 | 1596.3 | 0.01057 | 0.724 |
| FUT6 | fucosyltransferase 6(FUT6) | NM_000150 | 114 | 131.2 | 0.01061 | 1.686 |
| FUT7 | fucosyltransferase 7(FUT7) | NM_004479 | 291 | 423.6 | 0.01061 | 1.686 |
| | | XM_929340 | 173.3 | 128.3 | 0.01073 | 0.722 |
| | | XM_929340 | 4436.9 | 3161.7 | 0.01073 | 0.722 |
| | | AW183894 | 1115.9 | 1640.1 | 0.01111 | 1.644 |

| | | | | | | |
|---|---|---|---|---|---|---|
| KCNC1 | potassium voltage-gated channel subfamily C member 1(KCNC1) | AK125480 | 137.2 | 170.3 | 0.01117 | 1.223 |
| SPCS2 | signal peptidase complex subunit 2(SPCS2) | NM_014752 | 1017.4 | 606.7 | 0.01129 | 0.642 |
| | | AI539492 | 211.5 | 290.5 | 0.01131 | 1.474 |
| | | XM_933970 | 507.4 | 329.2 | 0.01136 | 0.674 |
| NDUFAF7 | NADH:ubiquinone oxidoreductase complex assembly factor 7(NDUFAF7) | NM_144736 | 142.2 | 129.7 | 0.01143 | 1.471 |
| NDUFAF7 | NADH:ubiquinone oxidoreductase complex assembly factor 7(NDUFAF7) | NM_144736 | 314.6 | 418.4 | 0.01143 | 1.471 |
| RIPK4 | receptor interacting serine/threonine kinase 4(RIPK4) | NM_020639 | 128.8 | 154.9 | 0.01146 | 1.162 |
| MAPK7 | mitogen-activated protein kinase 7(MAPK7) | NM_139032 | 130.6 | 134.8 | 0.01154 | 0.854 |
| MAPK8IP1 | mitogen-activated protein kinase 8 interacting protein 1(MAPK8IP1) | NM_005456 | 177 | 151.9 | 0.01154 | 0.854 |
| MRPL3 | mitochondrial ribosomal protein L3(MRPL3) | NM_007208 | 124.7 | 148.7 | 0.01165 | 1.137 |

| | | | | | | |
|---|---|---|---|---|---|---|
| EIF4E2 | eukaryotic translation initiation factor 4E family member 2(EIF4E2) | NM_004846 | 1861.8 | 1286.1 | 0.01191 | 0.771 |
| | | XM_001715717 | 160.7 | 131.4 | 0.01204 | 0.809 |
| | | XM_001715717 | 415.3 | 342.5 | 0.01204 | 0.809 |
| BBS1 | Bardet-Biedl syndrome 1(BBS1) | NM_024649 | 179.1 | 201.5 | 0.01237 | 1.141 |
| | | CR605357 | 133 | 158.6 | 0.01242 | 1.17 |
| CYP4A11 | cytochrome P450 family 4 subfamily A member 11(CYP4A11) | NM_000778 | 146.1 | 134.9 | 0.01252 | 0.905 |
| LOC105378397 | uncharacterized LOC105378397(LOC105378397) | BX092644 | 178.7 | 230.7 | 0.01256 | 1.343 |
| CCNK | cyclin K(CCNK) | NM_003858 | 122.3 | 109.6 | 0.01258 | 0.865 |
| TMEM106C | transmembrane protein 106C(TMEM106C) | NM_024056 | 527.8 | 681.4 | 0.01262 | 1.508 |
| cenpv | centromere protein V(CENPV) | NM_181716 | 208.1 | 258.5 | 0.01267 | 1.313 |
| CENPX | centromere protein X(CENPX) | NM_144998 | 416.2 | 502.5 | 0.01267 | 1.313 |
| TMEM267 | transmembrane protein 267(TMEM267) | NM_022483 | 2771.5 | 4189.6 | 0.01278 | 1.703 |
| RPL7A | ribosomal protein L7a(RPL7A) | NM_000972 | 320.5 | 241.5 | 0.01279 | 0.759 |

| | | | | | | |
|---|---|---|---|---|---|---|
| | | XR_036919 | 4757.2 | 2981.9 | 0.01296 | 0.746 |
| EEPD1 | endonuclease/exonuclease/phosphatase family domain containing 1(EEPD1) | NM_030636 | 295.4 | 224.9 | 0.01307 | 0.778 |
| MAR5 | membrane associated ring-CH-type finger 5(MARCH5) | NM_017824 | 101.3 | 113.9 | 0.01308 | 1.137 |
| MAR6 | membrane associated ring-CH-type finger 6(MARCH6) | NM_005885 | 103.4 | 125.5 | 0.01308 | 1.137 |
| | | XR_039439 | 155.4 | 129.1 | 0.0132 | 0.495 |
| | | XR_039439 | 1435 | 675.3 | 0.0132 | 0.495 |
| | | XR_016832 | 224.8 | 168.2 | 0.01326 | 0.752 |
| | | XR_038101 | 843.9 | 504.2 | 0.01336 | 0.628 |
| EPB41L5 | erythrocyte membrane protein band 4.1 like 5(EPB41L5) | NM_020909 | 1038.6 | 725.5 | 0.01341 | 0.752 |
| | | XR_038433 | 297.4 | 177.6 | 0.01345 | 0.569 |
| | | XR_038433 | 2121.1 | 1243.5 | 0.01345 | 0.569 |
| MPZL1 | myelin protein zero like 1(MPZL1) | NM_024569 | 121.6 | 102 | 0.01348 | 0.812 |
| | | XM_001131877 | 117.6 | 139.7 | 0.0135 | 1.136 |

| | | | | | | | | | |
|---|---|---|---|---|---|---|---|---|---|
| RUSC1 | RUN and SH3 domain containing 1(RUSC1) | NM_014328 | 662.7 | 832.5 | 0.01363 | 1.51 | | | |
| THBS3 | thrombospondin 3(THBS3) | NM_007112 | 300.1 | 347.7 | 0.01364 | 1.277 | | | |
| | | NM_001039755 | 5325.7 | 6709 | 0.0137 | 1.431 | | | |
| HBP1 | HMG-box transcription factor 1(HBP1) | NM_012257 | 180.7 | 166.6 | 0.01374 | 0.744 | | | |
| HBQ1 | hemoglobin subunit theta 1(HBQ1) | NM_005331 | 337 | 242.9 | 0.01374 | 0.744 | | | |
| RPL19P9 | ribosomal protein L19 pseudogene 9(RPL19P9) | NM_001080544 | 660.6 | 436.1 | 0.01375 | 0.724 | | | |
| RPL19P9 | ribosomal protein L19 pseudogene 9(RPL19P9) | NM_001080544 | 6597.3 | 4381.6 | 0.01375 | 0.724 | | | |
| | | XR_038693 | 1302 | 690.7 | 0.01381 | 0.563 | | | |
| | | XR_038693 | 1767.9 | 1231.7 | 0.01381 | 0.563 | | | |
| | | XM_001723164 | 151.5 | 167.3 | 0.01411 | 1.098 | | | |
| | | XR_037282 | 263.9 | 176.4 | 0.01417 | 0.673 | NM_001107943 | 0.63 | 112.77 |
| MAFF | MAF bZIP transcription factor F(MAFF) | NM_012323 | 146.7 | 136.4 | 0.01423 | 1.957 | | | |
| MAFG | MAF bZIP transcription factor G(MAFG) | NM_002359 | 487.8 | 800.6 | 0.01423 | 1.957 | | | |
| | | XM_001133677 | 106.4 | 116.7 | 0.01429 | 0.882 | | | |

| | | | | | | | | | |
|---|---|---|---|---|---|---|---|---|---|
| | | XM_001133677 | 138.9 | 123.4 | 0.01429 | 0.882 | | | |
| MAP2K7 | mitogen-activated protein kinase kinase 7(MAP2K7) | NM_145185 | 127.2 | 113.2 | 0.0143 | 0.864 | | | |
| PIPSL | PIP5K1A and PSMD4-like, pseudogene(PIPSL) | NR_002319 | 223.4 | 172.6 | 0.01452 | 0.774 | | | |
| ARL17B | ADP ribosylation factor like GTPase 17B(ARL17B) | NM_001039083 | 4156 | 5459.5 | 0.01458 | 1.473 | | | |
| | | XM_937758 | 805.8 | 480 | 0.01463 | 0.638 | | | |
| c21orf59 | chromosome 21 open reading frame 59(C21orf59) | NM_021254 | 1267.9 | 1675.6 | 0.01472 | 1.473 | | | |
| PI4KA | phosphatidylinositol 4-kinase alpha(PI4KA) | NM_002650 | 252.8 | 190.5 | 0.01475 | 0.761 | | | |
| ZNF280D | zinc finger protein 280D(ZNF280D) | NM_001002844 | 113.1 | 106.4 | 0.0148 | 0.885 | | | |
| | | XM_929608 | 100 | 124.1 | 0.01487 | 1.172 | | | |
| GPRC5A | G protein-coupled receptor class C group 5 member A(GPRC5A) | NM_003979 | 118.4 | 137.2 | 0.01491 | 1.115 | | | |
| | | XR_017653 | 2142.3 | 1463.8 | 0.01502 | 0.77 | NM_134372 | 22.51 | 0.03 |
| | | XM_927468 | 125.8 | 115.6 | 0.01503 | 0.884 | | | |
| TRNT1 | tRNA nucleotidyl transferase 1(TRNT1) | NM_182916 | 117.5 | 105.4 | 0.01514 | 0.851 | | | |

| | | | | | | |
|---|---|---|---|---|---|---|
| TRNT1 | tRNA nucleotidyl transferase 1(TRNT1) | NM_182916 | 126.2 | 120.2 | 0.01514 | 0.851 |
| | | XM_001719035 | 522.4 | 338.7 | 0.01515 | 0.674 |
| ZDHHC24 | zinc finger DHHC-type containing 24(ZDHHC24) | NM_207340 | 262.6 | 290.4 | 0.01519 | 1.181 |
| EWSR1 | EWS RNA binding protein 1(EWSR1) | NM_005243 | 222.5 | 298.9 | 0.01527 | 1.489 |
| MROH7 | maestro heat like repeat family member 7(MROH7) | NM_001039464 | 215.5 | 355.7 | 0.01531 | 1.862 |
| PLCD4 | phospholipase C delta 4(PLCD4) | NM_032726 | 179.8 | 254.8 | 0.01535 | 1.534 |
| FAM63A | family with sequence similarity 63 member A(FAM63A) | NM_018379 | 124.1 | 148.6 | 0.01536 | 1.429 |
| FAM63B | family with sequence similarity 63 member B(FAM63B) | NM_001040450 | 626.5 | 784.2 | 0.01536 | 1.429 |
| | | XR_016872 | 308.9 | 200.2 | 0.01544 | 0.646 |
| SLC35E1 | solute carrier family 35 member E1(SLC35E1) | NM_024881 | 121.9 | 121.4 | 0.01558 | 1.543 |
| SLC35E1 | solute carrier family 35 member E1(SLC35E1) | NM_024881 | 1336.2 | 1830.8 | 0.01558 | 1.543 |
| | | XM_927693 | 142 | 164.3 | 0.01559 | 1.148 |
| | | XM_001721625 | 2052.7 | 2763.4 | 0.0156 | 1.474 |

| | | | | | | | | | |
|---|---|---|---|---|---|---|---|---|---|
| C1orf229 | chromosome 1 open reading frame 229(C1orf229) | NM_207401 | 162 | 144.4 | 0.01563 | 0.886 | | | |
| ZNF394 | zinc finger protein 394(ZNF394) | NM_032164 | 1925.2 | 2622.5 | 0.01565 | 1.525 | | | |
| | | AW450365 | 123.5 | 150.2 | 0.01567 | 1.174 | | | |
| | | AA503583 | 153 | 130 | 0.01584 | 0.811 | | | |
| ACTB | actin beta(ACTB) | NM_001101 | 2599.1 | 1663.7 | 0.01594 | 0.593 | | | |
| ACTB | actin beta(ACTB) | NM_001101 | 7876.7 | 3772.5 | 0.01594 | 0.593 | | | |
| ACTB | actin beta(ACTB) | NM_001101 | 13400.8 | 8681.3 | 0.01594 | 0.593 | | | |
| RAMP2 | receptor activity modifying protein 2(RAMP2) | NM_005854 | 144.1 | 167.6 | 0.01602 | 1.146 | NM_001031645 | 1.93 | 10.04 |
| RAMP2 | receptor activity modifying protein 2(RAMP2) | NM_005854 | 159.1 | 173.6 | 0.01602 | 1.146 | | | |
| | | XM_945223 | 123.9 | 144.5 | 0.01603 | 1.124 | | | |
| | | XM_934920 | 250.7 | 170.2 | 0.01607 | 0.703 | | | |
| ZBTB41 | zinc finger and BTB domain containing 41(ZBTB41) | NM_194314 | 132.7 | 112.5 | 0.0162 | 0.831 | | | |
| | | XR_037881 | 117.9 | 122.3 | 0.01671 | 0.605 | | | |
| | | XR_037881 | 1416.1 | 810.5 | 0.01671 | 0.605 | | | |

| | | | | | | | | | |
|---|---|---|---|---|---|---|---|---|---|
| | | XM_943174 | 115.6 | 147.2 | 0.01677 | 1.249 | | | |
| SLC26A10 | solute carrier family 26 member 10(SLC26A10) | NM_133489 | 99.9 | 105.7 | 0.01684 | 0.882 | | | |
| SLC26A10 | solute carrier family 26 member 10(SLC26A10) | NM_133489 | 144.5 | 130.6 | 0.01684 | 0.882 | | | |
| POTEKP | POTE ankyrin domain family member K, pseudogene(POTEKP) | NM_001017421 | 5570.2 | 2215.8 | 0.01696 | 0.53 | | | |
| ENTPD1 | ectonucleoside triphosphate diphosphohydrolase 1(ENTPD1) | NM_001098175 | 276.2 | 200.2 | 0.01706 | 0.738 | | | |
| | | XM_944499 | 117.6 | 112.3 | 0.01711 | 0.92 | | | |
| PIGL | phosphatidylinositol glycan anchor biosynthesis class L(PIGL) | NM_004278 | 128.1 | 120.8 | 0.01714 | 0.905 | | | |
| | | XM_0011129013 | 153.4 | 158.6 | 0.01715 | 1.741 | | | |
| | | XM_0011129013 | 355.1 | 527.1 | 0.01715 | 1.741 | NM_144748 | 66.99 | 0.14 |
| CDK2AP2 | cyclin dependent kinase 2 associated protein 2(CDK2AP2) | NM_005851 | 166.6 | 209.5 | 0.01716 | 1.28 | | | |
| | | XR_015295 | 5069.4 | 3031.1 | 0.01726 | 0.674 | | | |
| STK11 | serine/threonine kinase 11(STK11) | NM_000455 | 176.8 | 145.8 | 0.0173 | 0.824 | | | |

| | | | | | | | | | |
|---|---|---|---|---|---|---|---|---|---|
| EID2B | EP300 interacting inhibitor of differentiation 2B(EID2B) | NM_152361 | 122.4 | 120.3 | 0.01733 | 1.477 | | | |
| EIF1 | eukaryotic translation initiation factor 1(EIF1) | NM_005801 | 4551.9 | 6130 | 0.01733 | 1.477 | | | |
| | | XM_001720210 | 238 | 184.3 | 0.01735 | 0.765 | | | |
| MUC6 | mucin 6, oligomeric mucus/gel-forming(MUC6) | NM_005961 | 129.1 | 126.6 | 0.01737 | 1.502 | | | |
| MUC6 | mucin 6, oligomeric mucus/gel-forming(MUC6) | NM_005961 | 223.8 | 209.8 | 0.01737 | 1.502 | | | |
| MUC6 | mucin 6, oligomeric mucus/gel-forming(MUC6) | NM_005961 | 307.2 | 393.2 | 0.01737 | 1.502 | NM_001107793 | 77.29 | 6.85 |
| ARF1 | ADP ribosylation factor 1(ARF1) | NM_001024228 | 433.4 | 313.9 | 0.01755 | 0.758 | NM_001107793 | 77.29 | 6.85 |
| | | XR_016027 | 194.3 | 150.1 | 0.01759 | 0.784 | NM_001107793 | 77.29 | 6.85 |
| ARSB | arylsulfatase B(ARSB) | NM_000046 | 149.2 | 133.2 | 0.01764 | 0.874 | NM_001107793 | 77.29 | 6.85 |
| KANK4 | KN motif and ankyrin repeat domains 4(KANK4) | NM_181712 | 105.4 | 142.7 | 0.01773 | 1.263 | | | |
| KANSL1 | KAT8 regulatory NSL complex subunit 1(KANSL1) | NM_015443 | 133.3 | 160.8 | 0.01773 | 1.263 | | | |

| | | | | | | |
|---|---|---|---|---|---|---|
| | | XM_001719452 | 120.8 | 136.8 | 0.01779 | 1.097 |
| STXBP2 | syntaxin binding protein 2(STXBP2) | NM_006949 | 294.2 | 347.2 | 0.01821 | 1.293 |
| | | XR_018937 | 392.6 | 238.8 | 0.01828 | 0.616 |
| | | XR_018937 | 4367.6 | 3029.7 | 0.01828 | 0.616 |
| TRIM13 | tripartite motif containing 13(TRIM13) | NM_001007278 | 126.8 | 127 | 0.01839 | 1.482 |
| TRIM13 | tripartite motif containing 13(TRIM13) | NM_001007278 | 875.2 | 1150.2 | 0.01839 | 1.482 |
| GNG8 | G protein subunit gamma 8(GNG8) | NM_033258 | 211.1 | 169.3 | 0.01839 | 0.797 |
| CTDSPL2 | CTD small phosphatase like 2(CTDSPL2) | NM_016396 | 1542.2 | 1003.5 | 0.01871 | 0.705 |
| CNGA1 | cyclic nucleotide gated channel alpha 1(CNGA1) | NM_000087 | 147.8 | 133.6 | 0.0188 | 0.869 |
| | | XM_001714161 | 171.2 | 192 | 0.01885 | 1.165 |
| C4BPA | complement component 4 binding protein alpha(C4BPA) | NM_000715 | 168.7 | 148.8 | 0.01889 | 0.876 |
| | | XR_019429 | 241.6 | 156.9 | 0.01905 | 0.638 |
| LGALS7B | galectin 7B(LGALS7B) | NM_001042507 | 369 | 836.2 | 0.01938 | 2.701 |
| LGALS8 | galectin 8(LGALS8) | NM_201545 | 983 | 2633.4 | 0.01944 | 2.899 |

| | | | | | | |
|---|---|---|---|---|---|---|
| LGALS7B | galectin 7B(LGALS7B) | NM_001042507 | 628 | 1540.9 | 0.01944 | 2.899 |
| MEX3C | mex-3 RNA binding family member C(MEX3C) | NM_016626 | 285.5 | 194.9 | 0.01952 | 0.694 |
| | | XM_001125808 | 172.8 | 141.4 | 0.01957 | 0.808 |
| lgals9 | galectin 9(LGALS9) | NM_009587 | 983 | 2633.4 | 0.0196 | 3.156 |
| PSMA3 | proteasome subunit alpha 3(PSMA3) | NM_152132 | 701.3 | 478.1 | 0.0198 | 0.733 |
| | | XR_039108 | 184.3 | 148.2 | 0.01993 | 0.793 |
| PTGES3 | prostaglandin E synthase 3(PTGES3) | NM_006601 | 302.1 | 211.8 | 0.01998 | 0.711 |
| | | XR_018710 | 109.6 | 127.1 | 0.02019 | 1.107 |
| brms1 | breast cancer metastasis suppressor 1(BRMS1) | NM_001024958 | 543.3 | 399.1 | 0.02022 | 0.855 |
| HSP90AA1 | heat shock protein 90 alpha family class A member 1(HSP90AA1) | NM_001017963 | 150.3 | 136.6 | 0.02028 | 0.889 |
| HSP90AA1 | heat shock protein 90 alpha family class A member 1(HSP90AA1) | NM_005348 | 170 | 159.3 | 0.02028 | 0.889 |
| LOC606724 | coronin 1A pseudogene(LOC606724) | NR_002454 | 3790 | 5486.9 | 0.02035 | 1.588 |

| | | | | | | | | | |
|---|---|---|---|---|---|---|---|---|---|
| FAM98B | family with sequence similarity 98 member B(FAM98B) | NM_001042429 | 361.1 | 418.5 | 0.02036 | 1.25 | | | |
| YWHAH | tyrosine 3-monooxygenase/tryptophan 5-monooxygenase activation protein eta(YWHAH) | NM_003405 | 3061.8 | 1821.7 | 0.02037 | 0.668 | | | |
| GPSM2 | G-protein signaling modulator 2(GPSM2) | NM_013296 | 117 | 134 | 0.02039 | 1.106 | | | |
| MIR1247 | microRNA 1247(MIR1247) | NR_031649 | 125.6 | 119.4 | 0.02039 | 0.905 | | | |
| SET | SET nuclear proto-oncogene(SET) | NM_003011 | 358.1 | 275.1 | 0.02042 | 0.764 | | | |
| DDX54 | DEAD-box helicase 54(DDX54) | NM_024072 | 4076.4 | 5394.3 | 0.02043 | 1.487 | | | |
| | | BX098521 | 151.7 | 199.5 | 0.02044 | 1.297 | | | |
| | | XR_001416 | 248 | 338.7 | 0.02063 | 1.621 | NM_001009603 | 28.32 | 0.90 |
| | | XR_018705 | 230.8 | 157.9 | 0.02067 | 0.677 | | | |
| CCDC22 | coiled-coil domain containing 22(CCDC22) | NM_014008 | 155.8 | 194.2 | 0.02068 | 1.29 | | | |
| DCUN1D1 | defective in cullin neddylation 1 domain containing 1(DCUN1D1) | NM_020640 | 264.6 | 193.2 | 0.02078 | 0.784 | | | |

| | | | | | | |
|---|---|---|---|---|---|---|
| DCTPP1 | dCTP pyrophosphatase 1(DCTPP1) | NM_024096 | 226.5 | 177.5 | 0.02078 | 0.784 |
| RBBP5 | RB binding protein 5, histone lysine methyltransferase complex subunit(RBBP5) | NM_005057 | 134.7 | 127.2 | 0.02091 | 0.916 |
| YY1 | YY1 transcription factor(YY1) | NM_003403 | 457.6 | 317.6 | 0.02093 | 0.721 |
| YY1 | YY1 transcription factor(YY1) | NM_003403 | 994.4 | 742.5 | 0.02093 | 0.721 |
| DPM1 | dolichyl-phosphate mannosyltransferase subunit 1, catalytic(DPM1) | NM_003859 | 233.9 | 190.1 | 0.02098 | 0.825 |
| DPM2 | dolichyl-phosphate mannosyltransferase subunit 2, regulatory(DPM2) | NM_003863 | 509.3 | 490 | 0.02098 | 0.825 |
| LOC101930164 | uncharacterized LOC101930164(LOC101930164) | AK026751 | 177.1 | 211.8 | 0.02102 | 1.241 |
| MYO3A | myosin IIIA(MYO3A) | NM_017433 | 133.9 | 124.2 | 0.02103 | 0.897 |
| PRRG4 | proline rich and Gla domain 4(PRRG4) | NM_024081 | 188.1 | 246.5 | 0.02121 | 1.372 |
| | | XR_018261 | 1141.9 | 606.6 | 0.02123 | 0.603 |
| NSA2 | NSA2, ribosome biogenesis homolog(NSA2) | NM_014886 | 1514.3 | 1136.2 | 0.02126 | 0.779 |

| | | | | | | |
|---|---|---|---|---|---|---|
| NSA2 | NSA2, ribosome biogenesis homolog(NSA2) | NM_014886 | 2216.3 | 1551.3 | 0.02126 | 0.779 |
| ZC3H3 | zinc finger CCCH-type containing 3(ZC3H3) | NM_015117 | 242.7 | 194.5 | 0.02133 | 0.819 |
| | | XM_0017214 30 | 3671.5 | 5101.1 | 0.02148 | 1.503 |
| ACVR1 | activin A receptor type 1(ACVR1) | NM_001105 | 220.5 | 273.4 | 0.02162 | 1.345 |
| APH1A | aph-1 homolog A, gamma-secretase subunit(APH1A) | NM_001077628 | 841.1 | 566.5 | 0.02167 | 0.724 |
| | | XR_019029 | 175.5 | 155.1 | 0.02168 | 0.88 |
| ACTB | actin beta(ACTB) | NM_001101 | 2599.1 | 1663.7 | 0.0217 | 0.69 |
| ACTB | actin beta(ACTB) | NM_001101 | 7876.7 | 3772.5 | 0.0217 | 0.69 |
| ACTG1 | actin gamma 1(ACTG1) | NM_001614 | 13400.8 | 8681.3 | 0.0217 | 0.69 |
| | | XR_040489 | 125.3 | 143.6 | 0.02172 | 1.111 |
| ZNF468 | zinc finger protein 468(ZNF468) | NM_199132 | 134.6 | 147.4 | 0.02187 | 1.195 |
| ZNF468 | zinc finger protein 468(ZNF468) | NM_199132 | 143 | 171.6 | 0.02187 | 1.195 |
| BNIP3L | BCL2 interacting protein 3 like(BNIP3L) | NM_004331 | 106.5 | 128.3 | 0.02193 | 1.146 |
| BNIPL | BCL2 interacting protein like(BNIPL) | NM_138278 | 136 | 155.3 | 0.02193 | 1.146 |

| | | | | | | |
|---|---|---|---|---|---|---|
| FLYWCH1 | FLYWCH-type zinc finger 1(FLYWCH1) | NM_020912 | 146.8 | 133.3 | 0.02202 | 0.884 |
| | | XM_938988 | 1597.3 | 550.1 | 0.02221 | 0.409 |
| POLRMT | RNA polymerase mitochondrial(POLRMT) | NM_005035 | 252.8 | 290.7 | 0.02229 | 1.226 |
| ETS1 | ETS proto-oncogene 1, transcription factor(ETS1) | NM_005238 | 211 | 290.5 | 0.02231 | 1.423 |
| FASTK | Fas activated serine/threonine kinase(FASTK) | NM_006712 | 116.3 | 97 | 0.02255 | 0.796 |
| KLF10 | Kruppel like factor 10(KLF10) | NM_005655 | 104.3 | 127.1 | 0.02262 | 1.16 |
| | | XM_944246 | 587.6 | 345 | 0.0227 | 0.62 |
| CEP192 | centrosomal protein 192(CEP192) | NM_032142 | 2713.1 | 3587.6 | 0.02299 | 1.456 |
| ART4 | ADP-ribosyltransferase 4 (Dombrock blood group)(ART4) | NM_021071 | 106.9 | 131.7 | 0.02317 | 1.146 |
| DYNC1LI2 | dynein cytoplasmic 1 light intermediate chain 2(DYNC1LI2) | NM_006141 | 132.2 | 123.3 | 0.02332 | 0.897 |
| | | XR_019565 | 2110.2 | 1326.2 | 0.02336 | 0.704 |
| | | XR_019565 | 3209.1 | 2022.7 | 0.02336 | 0.704 |
| DIXDC1 | DIX domain containing 1(DIXDC1) | NM_001037954 | 135.7 | 150 | 0.02338 | 1.077 |

| | | | | | | |
|---|---|---|---|---|---|---|
| | | CD237661 | 289.1 | 400.2 | 0.02363 | 1.601 |
| RPS27A | ribosomal protein S27a(RPS27A) | NM_002954 | 3219 | 2388.2 | 0.02372 | 0.809 |
| RPS27A | ribosomal protein S27a(RPS27A) | NM_002954 | 17436.1 | 14379.2 | 0.02372 | 0.809 |
| | | BF054947 | 125.4 | 141.5 | 0.02391 | 1.09 |
| | | XR_016385 | 386.7 | 274.5 | 0.02404 | 0.74 |
| | | XM_372319 | 138.5 | 160.9 | 0.02422 | 1.165 |
| | | XR_042295 | 210.8 | 303.1 | 0.02437 | 1.58 |
| RPL6 | ribosomal protein L6(RPL6) | NM_001024662 | 165.1 | 148.9 | 0.0244 | 0.726 |
| RPL6 | ribosomal protein L6(RPL6) | NM_001024662 | 2434.3 | 1880.7 | 0.0244 | 0.726 |
| RPL6 | ribosomal protein L6(RPL6) | NM_001024662 | 5324.9 | 3122.2 | 0.0244 | 0.726 |
| TBRG4 | transforming growth factor beta regulator 4(TBRG4) | NM_004749 | 159.3 | 143.5 | 0.02451 | 0.886 |
| | | XR_039678 | 173.8 | 147.4 | 0.02457 | 0.85 |
| ZRANB1 | zinc finger RANBP2-type containing 1(ZRANB1) | NM_017580 | 154.2 | 175.2 | 0.02463 | 1.153 |
| ABHD17B | abhydrolase domain containing 17B(ABHD17B) | NM_016014 | 114.5 | 126.7 | 0.02471 | 0.817 |

| | | | | | | |
|---|---|---|---|---|---|---|
| ABHD17C | abhydrolase domain containing 17C(ABHD17C) | NM_021214 | 154.7 | 129.6 | 0.02471 | 0.817 |
| | | XM_001723141 | 1229.6 | 827.3 | 0.02472 | 0.736 |
| CCT8L1P | chaperonin containing TCP1 subunit 8 like 1, pseudogene(CCT8L1P) | NM_001029866 | 447.8 | 297.4 | 0.0249 | 0.668 |
| | | XR_038984 | 981.9 | 612.8 | 0.02492 | 0.669 |
| BTG2 | BTG anti-proliferation factor 2(BTG2) | NM_006763 | 282.1 | 226.1 | 0.02504 | 0.813 |
| HNMT | histamine N-methyltransferase(HNMT) | NM_006895 | 108.5 | 130.5 | 0.02506 | 1.127 |
| HNRNPA1 | heterogeneous nuclear ribonucleoprotein A1(HNRNPA1) | NM_031157 | 113.3 | 119.9 | 0.02506 | 1.127 |
| | | AW276479 | 1131.4 | 1642 | 0.02509 | 1.62 |
| GSKIP | GSK3B interacting protein(GSKIP) | NM_016472 | 98.9 | 124.5 | 0.02511 | 1.164 |
| PXMP4 | peroxisomal membrane protein 4(PXMP4) | NM_183397 | 150.9 | 177.8 | 0.02518 | 1.171 |
| SHROOM4 | shroom family member 4(SHROOM4) | NM_020717 | 135.1 | 165.1 | 0.0252 | 1.633 |
| SHROOM4 | shroom family member 4(SHROOM4) | NM_020717 | 729 | 1031.1 | 0.0252 | 1.633 |
| ENO1 | enolase 1(ENO1) | NM_001428 | 159.2 | 193.5 | 0.02526 | 1.233 |

| | | | | | | |
|---|---|---|---|---|---|---|
| SEC22C | SEC22 homolog C, vesicle trafficking protein(SEC22C) | NM_032970 | 124.4 | 114.1 | 0.02528 | 0.884 |
| SEC22C | SEC22 homolog C, vesicle trafficking protein(SEC22C) | NM_032970 | 130.9 | 135.7 | 0.02528 | 0.884 |
| EFCAB5 | EF-hand calcium binding domain 5(EFCAB5) | NM_001033562 | 335.3 | 263 | 0.02532 | 0.811 |
| EIF4E3 | eukaryotic translation initiation factor 4E family member 3(EIF4E3) | NM_173359 | 207.5 | 174.1 | 0.02538 | 1.41 |
| EIF4EBP2 | eukaryotic translation initiation factor 4E binding protein 2(EIF4EBP2) | NM_004096 | 232.9 | 286.3 | 0.02538 | 1.41 |
| CSNK2A1 | casein kinase 2 alpha 1(CSNK2A1) | NM_001895 | 190.8 | 163.3 | 0.02543 | 0.856 |
| | | XR_017565 | 605.1 | 385.7 | 0.02553 | 0.652 |
| | | XR_038851 | 505.1 | 672.8 | 0.02554 | 1.552 |
| TDRD1 | tudor domain containing 1(TDRD1) | NM_198795 | 273.5 | 350.9 | 0.02562 | 1.341 |
| TDRD1 | tudor domain containing 1(TDRD1) | NM_198795 | 346 | 425.7 | 0.02562 | 1.341 |
| | | XR_039146 | 344.3 | 230.4 | 0.02562 | 0.673 |
| | | XM_292085 | 1828.7 | 890.3 | 0.02569 | 0.531 |

| | | | | | | | | | |
|---|---|---|---|---|---|---|---|---|---|
| SSPO | SCO-spondin(SSPO) | NM_198455 | 119.1 | 153.3 | 0.02596 | 1.222 | | | |
| ATP6V0E1 | ATPase H+ transporting V0 subunit e1(ATP6V0E1) | NM_003945 | 338.6 | 416.2 | 0.02619 | 1.308 | | | |
| | | W91949 | 129.6 | 107.7 | 0.02624 | 0.791 | | | |
| GDPD3 | glycerophosphodiester phosphodiesterase domain containing 3(GDPD3) | NM_024307 | 340.3 | 412.8 | 0.02631 | 1.35 | | | |
| CCT3 | chaperonin containing TCP1 subunit 3(CCT3) | NM_005998 | 466.1 | 354 | 0.02635 | 0.793 | | | |
| c1orf116 | chromosome 1 open reading frame 116(C1orf116) | NM_023938 | 171.1 | 202.7 | 0.02646 | 1.27 | | | |
| C1orf122 | chromosome 1 open reading frame 122(C1orf122) | NM_198446 | 232.5 | 297 | 0.02646 | 1.27 | | | |
| INO80 | INO80 complex subunit(INO80) | NM_017553 | 7156.1 | 8841.3 | 0.02655 | 1.366 | | | |
| | | XM_371314 | 128.5 | 149 | 0.02668 | 1.129 | | | |
| GPR37 | G protein-coupled receptor 37(GPR37) | NM_005302 | 167.1 | 146.3 | 0.02682 | 0.87 | | | |
| TRMO | tRNA methyltransferase O(TRMO) | NM_016481 | 296 | 226.6 | 0.0269 | 0.775 | | | |
| | | XM_940103 | 171.6 | 196.9 | 0.02703 | 1.197 | NM_144744 | 0.11 | 40.01 |
| AKIRIN1 | akirin 1(AKIRIN1) | NM_024595 | 167.7 | 209.2 | 0.02705 | 1.291 | | | |

| | | | | | | |
|---|---|---|---|---|---|---|
| RPS10 | ribosomal protein S10(RPS10) | NM_0010 14 | 10480.3 | 6847.8 | 0.02706 | 0.756 |
| CLPB | ClpB homolog, mitochondrial AAA ATPase chaperonin(CLPB) | NM_0308 13 | 251.7 | 318.1 | 0.02707 | 1.394 |
| RPL36 | ribosomal protein L36(RPL36) | NM_0336 43 | 1757.7 | 1029 | 0.0272 | 0.689 |
| | | XR_03794 5 | 139.2 | 117.9 | 0.02725 | 0.833 |
| CDCA8 | cell division cycle associated 8(CDCA8) | NM_0181 01 | 214.8 | 163.5 | 0.02732 | 0.769 |
| | | XM_92812 2 | 224.9 | 260.5 | 0.02733 | 1.261 |
| RPL7P9 | ribosomal protein L7 pseudogene 9(RPL7P9) | NM_0010 93763 | 330.3 | 242.5 | 0.02744 | 0.739 |
| | | XM_49718 2 | 114.3 | 104.9 | 0.02747 | 0.866 |
| COPS4 | COP9 signalosome subunit 4(COPS4) | NM_0161 29 | 184.8 | 147.3 | 0.02761 | 0.781 |
| magoh | mago homolog, exon junction complex core component(MAGOH) | NM_0023 70 | 158.5 | 197.2 | 0.0278 | 1.242 |
| DPM1 | dolichyl-phosphate mannosyltransferase subunit 1, catalytic(DPM1) | NM_0038 59 | 135.3 | 118.4 | 0.0278 | 0.854 |
| | | XM_93859 9 | 827.3 | 479.3 | 0.02787 | 0.628 |

| | | | | | | |
|---|---|---|---|---|---|---|
| | | XR_038738 | 359.9 | 244.6 | 0.02806 | 0.701 |
| BRK1 | BRICK1, SCAR/WAVE actin nucleating complex subunit(BRK1) | NM_018462 | 543.3 | 399.1 | 0.02812 | 0.779 |
| | | XM_0011266896 | 137.5 | 127.4 | 0.02812 | 0.9 |
| BRIX1 | BRX1, biogenesis of ribosomes(BRIX1) | NM_018321 | 163.7 | 139.9 | 0.02812 | 0.779 |
| | | DB337588 | 119.6 | 138.6 | 0.02814 | 1.116 |
| HS3ST3A1 | heparan sulfate-glucosamine 3-sulfotransferase 3A1(HS3ST3A1) | NM_006042 | 139.9 | 126.6 | 0.0282 | 0.88 |
| | | XM_932143 | 246.4 | 292.3 | 0.02853 | 1.259 |
| C14orf132 | chromosome 14 open reading frame 132(C14orf132) | NM_020215 | 189.3 | 156.8 | 0.02853 | 0.827 |
| | | XM_938903 | 197.6 | 153.7 | 0.02877 | 0.772 |
| ZNF652 | zinc finger protein 652(ZNF652) | NM_014897 | 6050.7 | 7159.2 | 0.02886 | 1.292 |
| MED20 | mediator complex subunit 20(MED20) | NM_004275 | 542.5 | 381.9 | 0.02887 | 0.75 |
| Tceal7 | transcription elongation factor A like 7(TCEAL7) | NM_152278 | 136.7 | 118.4 | 0.02892 | 0.841 |
| ARMC8 | armadillo repeat containing 8(ARMC8) | NM_015396 | 296.8 | 352.5 | 0.02894 | 1.304 |

| | | | | | | |
|---|---|---|---|---|---|---|
| MSLN | mesothelin(MSLN) | NM_013404 | 120.5 | 139.3 | 0.02896 | 1.115 |
| LRRFIP2 | LRR binding FLII interacting protein 2(LRRFIP2) | NM_017724 | 1687.3 | 2879.9 | 0.02898 | 1.937 |
| SHISA4 | shisa family member 4(SHISA4) | NM_198149 | 103.6 | 122.5 | 0.02903 | 1.112 |
| | | XR_016670 | 139.4 | 125.4 | 0.02906 | 0.87 |
| SIL1 | SIL1 nucleotide exchange factor(SIL1) | NM_001037633 | 167.1 | 180.6 | 0.02911 | 1.372 |
| SIL1 | SIL1 nucleotide exchange factor(SIL1) | NM_001037633 | 332.5 | 405.8 | 0.02911 | 1.372 |
| COLGALT1 | collagen beta(1-O)galactosyltransferase 1(COLGALT1) | NM_024656 | 211.1 | 272.1 | 0.02925 | 1.385 |
| PLCD1 | phospholipase C delta 1(PLCD1) | NM_006225 | 212.9 | 264.2 | 0.02952 | 1.277 |
| FCER1G | Fc fragment of IgE receptor Ig(FCER1G) | NM_004106 | 1717.1 | 2748.9 | 0.02961 | 1.797 |
| | | XM_376420 | 10758.5 | 7086 | 0.02965 | 0.747 |
| ZNF548 | zinc finger protein 548(ZNF548) | NM_152909 | 174.9 | 153.3 | 0.02966 | 0.875 |
| TNPO1 | transportin 1(TNPO1) | NM_153188 | 139.4 | 132.5 | 0.02976 | 0.751 |
| TNPO1 | transportin 1(TNPO1) | NM_153188 | 266.6 | 235.8 | 0.02976 | 0.751 |
| TNPO1 | transportin 1(TNPO1) | NM_153188 | 463.8 | 330.8 | 0.02976 | 0.751 |
| | | BX111413 | 115.3 | 141.8 | 0.02997 | 1.191 |

| | | | | | | |
|---|---|---|---|---|---|---|
| CRCP | CGRP receptor component(CRCP) | NM_014478 | 125.3 | 131.5 | 0.03 | 1.344 |
| CRCT1 | cysteine rich C-terminal 1(CRCT1) | NM_019060 | 3804.2 | 4477.9 | 0.03 | 1.344 |
| TRIM45 | tripartite motif containing 45(TRIM45) | NM_025188 | 179.4 | 192.9 | 0.03024 | 1.101 |
| NEK11 | NIMA related kinase 11(NEK11) | NM_145910 | 118.5 | 140.4 | 0.03027 | 1.14 |
| PSMG1 | proteasome assembly chaperone 1(PSMG1) | NM_203433 | 595 | 441.6 | 0.03029 | 0.784 |
| | | BX091766 | 141.2 | 160.9 | 0.03045 | 1.14 |
| | | AK095707 | 178.6 | 224.2 | 0.03046 | 1.343 |
| BTF3L4 | basic transcription factor 3 like 4(BTF3L4) | NM_152265 | 224.9 | 184.8 | 0.03066 | 0.811 |
| BTG1 | BTG anti-proliferation factor 1(BTG1) | NM_001731 | 282.1 | 226.1 | 0.03066 | 0.811 |
| | | XM_937382 | 120.7 | 138.1 | 0.03067 | 1.094 |
| BICC1 | BicC family RNA binding protein 1(BICC1) | NM_001080512 | 105.2 | 134.3 | 0.03069 | 1.203 |
| | | XM_941107 | 125.4 | 146 | 0.03087 | 1.131 |
| | | XM_926370 | 171.7 | 134.2 | 0.03095 | 0.778 |
| NOXA1 | NADPH oxidase activator 1(NOXA1) | NM_006647 | 269.4 | 350.3 | 0.031 | 1.421 |
| RBM34 | RNA binding motif protein 34(RBM34) | NM_015014 | 123 | 110.9 | 0.03116 | 0.863 |
| ZNF80 | zinc finger protein 80(ZNF80) | NM_007136 | 120.4 | 140.4 | 0.0312 | 1.121 |

| | | | | | | |
|---|---|---|---|---|---|---|
| BAIAP2 | BAI1 associated protein 2(BAIAP2) | NM_006340 | 130.3 | 152.2 | 0.0313 | 1.127 |
| BAIAP2 | BAI1 associated protein 2(BAIAP2) | NM_017450 | 195.3 | 221.3 | 0.0313 | 1.127 |
| | | CF552427 | 186.5 | 227.4 | 0.03139 | 1.315 |
| NFYA | nuclear transcription factor Y subunit alpha(NFYA) | NM_002505 | 138.8 | 126 | 0.03151 | 0.895 |
| CACYBP | calcyclin binding protein(CACYBP) | NM_014412 | 151.6 | 140.4 | 0.03156 | 0.915 |
| | | XR_039373 | 261.1 | 185.7 | 0.03168 | 0.712 |
| | | XM_001720622 | 150.1 | 130.2 | 0.03172 | 0.851 |
| ZNF2 | zinc finger protein 2(ZNF2) | NM_021088 | 117.1 | 138.6 | 0.03172 | 1.125 |
| OSBPL2 | oxysterol binding protein like 2(OSBPL2) | NM_001001691 | 146.9 | 170.7 | 0.03173 | 1.143 |
| PLAC8L1 | PLAC8 like 1(PLAC8L1) | NM_001029869 | 127.1 | 117.6 | 0.03174 | 0.882 |
| AGXT | alanine-glyoxylate aminotransferase(AGXT) | NM_000030 | 117.7 | 140.3 | 0.03175 | 1.134 |
| ALAD | aminolevulinate dehydratase(ALAD) | NM_001003945 | 125.9 | 139.5 | 0.03182 | 1.559 |
| | | XM_929418 | 140.6 | 130.1 | 0.03182 | 0.901 |
| ALAS1 | 5'-aminolevulinate synthase 1(ALAS1) | NM_000688 | 282.5 | 411 | 0.03182 | 1.559 |
| CGN | cingulin(CGN) | NM_020770 | 217.7 | 177.4 | 0.03185 | 0.812 |

| | | | | | | |
|---|---|---|---|---|---|---|
| SAE1 | SUMO1 activating enzyme subunit 1(SAE1) | NM_005500 | 793.7 | 462.1 | 0.03205 | 0.628 |
| PPFIBP1 | PPFIA binding protein 1(PPFIBP1) | NM_003622 | 120.6 | 140.5 | 0.0321 | 0.827 |
| PPFIBP1 | PPFIA binding protein 1(PPFIBP1) | NM_003622 | 128.3 | 110.6 | 0.0321 | 0.827 |
| PPFIBP1 | PPFIA binding protein 1(PPFIBP1) | NM_003622 | 183 | 171.5 | 0.0321 | 0.827 |
| ACTR3B | ARP3 actin related protein 3 homolog B(ACTR3B) | NM_020445 | 310.8 | 169.9 | 0.0321 | 0.555 |
| AKT3 | AKT serine/threonine kinase 3(AKT3) | NM_005465 | 642.2 | 479.4 | 0.03212 | 0.795 |
| | | XR_037673 | 2180.1 | 1427.4 | 0.03212 | 0.712 |
| | | XM_941064 | 124.6 | 143.6 | 0.03213 | 1.104 |
| MOS | v-mos Moloney murine sarcoma viral oncogene homolog(MOS) | NM_005372 | 116.5 | 111.6 | 0.03233 | 0.909 |
| KIAA0513 | KIAA0513(KIAA0513) | NM_014732 | 389.9 | 525.8 | 0.03238 | 1.61 |
| SLK | STE20 like kinase(SLK) | NM_014720 | 153.9 | 138.2 | 0.03257 | 0.872 |
| | | XM_0017166468 | 127.3 | 151 | 0.03262 | 1.168 |

| | | | | | | |
|---|---|---|---|---|---|---|
| SPECC1L | sperm antigen with calponin homology and coiled-coil domains 1 like(SPECC1L) | NM_015330 | 130.9 | 118.1 | 0.03266 | 0.855 |
| SPECC1L | sperm antigen with calponin homology and coiled-coil domains 1 like(SPECC1L) | NM_015330 | 736 | 609.6 | 0.03266 | 0.855 |
| KLHL28 | kelch like family member 28(KLHL28) | NM_017658 | 224.6 | 210.3 | 0.03268 | 1.416 |
| KLHL3 | kelch like family member 3(KLHL3) | NM_017415 | 291.1 | 369.6 | 0.03268 | 1.416 |
| | | XM_001717865 | 124.3 | 138.6 | 0.03272 | 1.082 |
| HINFP | histone H4 transcription factor(HINFP) | NM_015517 | 276.6 | 207.7 | 0.03275 | 0.756 |
| HIKESHI | Hikeshi, heat shock protein nuclear import factor(HIKESHI) | NM_016401 | 136.8 | 132.8 | 0.03275 | 0.756 |
| NUMA1 | nuclear mitotic apparatus protein 1(NUMA1) | NM_006185 | 158 | 149.5 | 0.03284 | 0.849 |
| NUMA1 | nuclear mitotic apparatus protein 1(NUMA1) | NM_006185 | 368.7 | 296 | 0.03284 | 0.849 |
| CNIH2 | cornichon family AMPA receptor auxiliary protein 2(CNIH2) | NM_182553 | 396.4 | 282.3 | 0.03292 | 0.736 |
| CNIH1 | cornichon family AMPA receptor auxiliary protein 1(CNIH1) | NM_005776 | 290.2 | 227.5 | 0.03292 | 0.736 |

| | | | | | | |
|---|---|---|---|---|---|---|
| | | XM_376787 | 323.8 | 214.6 | 0.03295 | 0.663 |
| GNB3 | G protein subunit beta 3(GNB3) | NM_002075 | 150.3 | 133.3 | 0.03302 | 0.871 |
| | | XM_001722111 | 1860.4 | 2514.9 | 0.03305 | 1.48 |
| ZKSCAN8 | zinc finger with KRAB and SCAN domains 8(ZKSCAN8) | NM_006298 | 119.2 | 108.3 | 0.03323 | 0.861 |
| | | CV318379 | 146.9 | 159.7 | 0.03346 | 1.073 |
| | | XM_938181 | 11401.5 | 8116 | 0.03346 | 0.793 |
| RPS4X | ribosomal protein S4, X-linked(RPS4X) | NM_001007 | 4554.2 | 2742.6 | 0.03348 | 0.735 |
| RPS4X | ribosomal protein S4, X-linked(RPS4X) | NM_001007 | 4921 | 3536.8 | 0.03348 | 0.735 |
| FBXW5 | F-box and WD repeat domain containing 5(FBXW5) | NM_018998 | 173.6 | 137.2 | 0.03351 | 0.794 |
| | | XM_927451 | 138.7 | 146.7 | 0.03351 | 0.68 |
| | | XM_927451 | 727.6 | 471.4 | 0.03351 | 0.68 |
| IL9 | interleukin 9(IL9) | NM_000590 | 291.5 | 203.4 | 0.03356 | 0.735 |
| PAXIP1 | PAX interacting protein 1(PAXIP1) | NM_007349 | 237.5 | 195.8 | 0.03361 | 0.832 |
| AMY1B | amylase, alpha 1B (salivary)(AMY1B) | NM_001008218 | 127.8 | 133.2 | 0.03373 | 0.655 |

| | | | | | | |
|---|---|---|---|---|---|---|
| AMY1C | amylase, alpha 1C (salivary)(AMY1C) | NM_0010 08219 | 2362.1 | 1459.6 | 0.03373 | 0.655 |
| SMARC D1 | SWI/SNF related, matrix associated, actin dependent regulator of chromatin, subfamily d, member 1(SMARCD1) | NM_0030 76 | 315.6 | 249.2 | 0.03397 | 0.811 |
| AFTPH | aftiphilin(AFTPH) | NM_0176 57 | 131.3 | 118.2 | 0.03407 | 0.866 |
| IL19 | interleukin 19(IL19) | NM_0133 71 | 11297.9 | 13722.9 | 0.03409 | 1.276 |
| CEP104 | centrosomal protein 104(CEP104) | NM_0147 04 | 176.3 | 167.3 | 0.03415 | 1.172 |
| CEP112 | centrosomal protein 112(CEP112) | NM_1450 36 | 204.2 | 229.3 | 0.03415 | 1.172 |
| RPL36A L | ribosomal protein L36a like(RPL36AL) | NM_0010 01 | 273.2 | 205.1 | 0.03418 | 0.761 |
| RPL36A L | ribosomal protein L36a like(RPL36AL) | NM_0010 01 | 2736.2 | 2005.2 | 0.03418 | 0.761 |
| GPX3 | glutathione peroxidase 3(GPX3) | NM_0020 84 | 3152.3 | 6021.5 | 0.03421 | 2.02 |
| | | XR_01724 9 | 146.2 | 125.6 | 0.03424 | 0.832 |
| EIF4G2 | eukaryotic translation initiation factor 4 gamma 2(EIF4G2) | NM_0014 18 | 557.7 | 376.5 | 0.03426 | 0.71 |
| EIF4G2 | eukaryotic translation initiation factor 4 gamma 2(EIF4G2) | NM_0014 18 | 2383.3 | 1905.2 | 0.03426 | 0.71 |

| | | | | | | | | | |
|---|---|---|---|---|---|---|---|---|---|
| EIF4G2 | eukaryotic translation initiation factor 4 gamma 2(EIF4G2) | NM_001418 | 4924.9 | 3901.3 | 0.03426 | 0.71 | | | |
| CGB5 | chorionic gonadotropin beta subunit 5(CGB5) | NM_033043 | 118.3 | 138.9 | 0.03426 | 1.129 | NM_017196 | 14.73 | 0.69 |
| | | AI198908 | 128.7 | 116.2 | 0.03434 | 0.874 | NM_017196 | 14.73 | 0.69 |
| TUBA1A | tubulin alpha 1a(TUBA1A) | NM_006009 | 777.6 | 439.7 | 0.03443 | 0.595 | | | |
| TUBA1A | tubulin alpha 1a(TUBA1A) | NM_006009 | 6647.5 | 4557.3 | 0.03443 | 0.595 | NM_017196 | 14.73 | 0.69 |
| ARHGAP12 | Rho GTPase activating protein 12(ARHGAP12) | NM_018287 | 907.2 | 943 | 0.03445 | 1.153 | | | |
| | | XM_933679 | 125.4 | 155.5 | 0.03449 | 0.777 | | | |
| | | XM_933679 | 198 | 156.2 | 0.03449 | 0.777 | | | |
| SDR42E1 | short chain dehydrogenase/reductase family 42E, member 1(SDR42E1) | NM_145168 | 119.1 | 135.3 | 0.03451 | 1.091 | NM_031356 | 5.43 | 24.53 |
| | | XM_001128866 | 1655.9 | 2265.5 | 0.03495 | 1.475 | NM_031356 | 5.43 | 24.53 |
| | | XR_017029 | 113.3 | 134.8 | 0.0351 | 1.142 | | | |
| HYLS1 | HYLS1, centriolar and ciliogenesis associated(HYLS1) | NM_145014 | 269.1 | 318.3 | 0.0351 | 1.266 | | | |
| | | BF996074 | 1010.1 | 1588.9 | 0.03514 | 1.785 | | | |
| | | XM_944513 | 116.1 | 140.1 | 0.0352 | 1.144 | | | |

| | | | | | | | | | |
|---|---|---|---|---|---|---|---|---|---|
| | | XR_037873 | 1822.1 | 1195.7 | 0.03524 | 0.738 | | | |
| KIF16B | kinesin family member 16B(KIF16B) | NM_024704 | 649.9 | 787.4 | 0.03528 | 1.355 | | | |
| PPHLN1 | periphilin 1(PPHLN1) | NM_201440 | 269.4 | 184.8 | 0.03533 | 0.684 | | | |
| FBXO38 | F-box protein 38(FBXO38) | NM_030793 | 405.8 | 287 | 0.03534 | 0.724 | | | |
| | | XM_001723702 | 241.1 | 177.9 | 0.0354 | 0.717 | | | |
| ZNF564 | zinc finger protein 564(ZNF564) | NM_144976 | 288.5 | 214.5 | 0.0354 | 0.771 | | | |
| ZNF256 | zinc finger protein 256(ZNF256) | NM_005773 | 190.1 | 141.7 | 0.03546 | 0.74 | | | |
| | | BX093647 | 124.4 | 136 | 0.03547 | 1.056 | | | |
| CDK6 | cyclin dependent kinase 6(CDK6) | NM_001259 | 711.9 | 927.2 | 0.03549 | 1.454 | | | |
| RAD9A | RAD9 checkpoint clamp component A(RAD9A) | NM_004584 | 206.7 | 259.8 | 0.03566 | 1.297 | | | |
| NECTIN4 | nectin cell adhesion molecule 4(NECTIN4) | NM_030916 | 107.5 | 131.7 | 0.03586 | 1.154 | NM_017135 | 41.45 | 2.88 |
| CSN1S1 | casein alpha s1(CSN1S1) | NM_001890 | 114.5 | 135.7 | 0.0359 | 1.118 | NM_017135 | 41.45 | 2.88 |
| | | XM_942516 | 128.7 | 120.3 | 0.03591 | 0.91 | NM_017135 | 41.45 | 2.88 |
| plekhb2 | pleckstrin homology domain containing B2(PLEKHB2) | NM_017958 | 124.6 | 126.5 | 0.0361 | 1.306 | NM_017135 | 41.45 | 2.88 |

| | | | | | | | | | |
|---|---|---|---|---|---|---|---|---|---|
| plekhb2 | pleckstrin homology domain containing B2(PLEKHB2) | NM_017958 | 161.6 | 209.6 | 0.0361 | 1.306 | NM_017135 | 41.45 | 2.88 |
| APP | amyloid beta precursor protein(APP) | NM_201414 | 1835.9 | 2194.8 | 0.03616 | 1.293 | | | |
| TBC1D25 | TBC1 domain family member 25(TBC1D25) | NM_002536 | 130 | 130.3 | 0.03624 | 0.913 | | | |
| TBC1D25 | TBC1 domain family member 25(TBC1D25) | NM_002536 | 161.6 | 147.7 | 0.03624 | 0.913 | | | |
| PIP4K2B | phosphatidylinositol-5-phosphate 4-kinase type 2 beta(PIP4K2B) | NM_138687 | 2108.7 | 2678.9 | 0.03626 | 1.448 | | | |
| DLD | dihydrolipoamide dehydrogenase(DLD) | NM_000108 | 121.3 | 141.5 | 0.03626 | 1.112 | | | |
| RHAG | Rh-associated glycoprotein(RHAG) | NM_000324 | 123.4 | 145.3 | 0.03629 | 1.157 | | | |
| SCRT1 | scratch family transcriptional repressor 1(SCRT1) | NM_031309 | 183.5 | 164.7 | 0.0363 | 0.891 | | | |
| TXNDC5 | thioredoxin domain containing 5(TXNDC5) | NM_022085 | 203.3 | 147 | 0.03632 | 0.729 | NM_001033653 | 43.46 | 1.56 |
| ITPKB | inositol-trisphosphate 3-kinase B(ITPKB) | NM_002221 | 4382.8 | 5647.3 | 0.03635 | 1.399 | | | |
| ITGA9 | integrin subunit alpha 9(ITGA9) | NM_002207 | 178.2 | 204.6 | 0.0364 | 1.182 | | | |
| NISCH | nischarin(NISCH) | NM_007184 | 389.5 | 455.4 | 0.03644 | 1.274 | | | |
| | | XR_019588 | 9256.9 | 6133.3 | 0.03644 | 0.779 | | | |

| | | | | | | |
|---|---|---|---|---|---|---|
| DAZL | deleted in azoospermia like(DAZL) | NM_001351 | 185.8 | 217.9 | 0.03654 | 1.23 |
| | | XM_937091 | 142.2 | 162.4 | 0.03662 | 1.134 |
| | | XR_041286 | 1954 | 1155.3 | 0.03669 | 0.707 |
| | | XR_041286 | 5828 | 3336.9 | 0.03669 | 0.707 |
| CCDC103 | coiled-coil domain containing 103(CCDC103) | NM_213607 | 3129.2 | 3781.1 | 0.03679 | 1.348 |
| KHK | ketohexokinase(KHK) | NM_000221 | 300 | 214.6 | 0.03679 | 0.795 |
| KHDRBS3 | KH RNA binding domain containing, signal transduction associated 3(KHDRBS3) | NM_006558 | 153.4 | 125.1 | 0.03679 | 0.795 |
| HSCB | HscB mitochondrial iron-sulfur cluster cochaperone(HSCB) | NM_172002 | 213.9 | 244.4 | 0.03683 | 1.189 |
| HSD11B1 | hydroxysteroid 11-beta dehydrogenase 1(HSD11B1) | NM_181755 | 411.9 | 490.6 | 0.03683 | 1.189 |
| CMTM6 | CKLF like MARVEL transmembrane domain containing 6(CMTM6) | NM_017801 | 209.9 | 235.3 | 0.03685 | 1.184 |
| EPB41L4B | erythrocyte membrane protein band 4.1 like 4B(EPB41L4B) | NM_019114 | 1100.7 | 755.6 | 0.03692 | 0.736 |

| | | | | | | | | | |
|---|---|---|---|---|---|---|---|---|---|
| | | XR_015954 | 195.4 | 151.5 | 0.03709 | 0.781 | | | |
| | | XR_038568 | 1456.6 | 749.2 | 0.0371 | 0.592 | | | |
| | | XR_038568 | 2160.6 | 1272.1 | 0.0371 | 0.592 | | | |
| CNOT7 | CCR4-NOT transcription complex subunit 7(CNOT7) | NM_013354 | 300.1 | 245.7 | 0.03711 | 0.835 | | | |
| CNOT8 | CCR4-NOT transcription complex subunit 8(CNOT8) | NM_004779 | 472.5 | 373.6 | 0.03711 | 0.835 | | | |
| REXO2 | RNA exonuclease 2(REXO2) | NM_015523 | 1022.6 | 690.5 | 0.03718 | 0.758 | | | |
| CSTF2 | cleavage stimulation factor subunit 2(CSTF2) | NM_001325 | 163.5 | 147 | 0.03721 | 0.889 | | | |
| LYPLA1 | lysophospholipase I(LYPLA1) | NM_006330 | 121.9 | 115.9 | 0.03728 | 0.898 | | | |
| ATP9A | ATPase phospholipid transporting 9A (putative)(ATP9A) | NM_006045 | 275.1 | 167.7 | 0.03729 | 0.599 | NM_134407 | 50.47 | 11.16 |
| MRPS18A | mitochondrial ribosomal protein S18A(MRPS18A) | NM_018135 | 324.5 | 352.4 | 0.03732 | 1.217 | | | |
| SAMD9 | sterile alpha motif domain containing 9(SAMD9) | NM_017654 | 120.6 | 141.6 | 0.03741 | 1.146 | | | |
| CEPT1 | choline/ethanolamine phosphotransferase 1(CEPT1) | NM_006090 | 154.2 | 174.9 | 0.03741 | 1.121 | | | |

| | | | | | | |
|---|---|---|---|---|---|---|
| PCDHGB2 | protocadherin gamma subfamily B, 2(PCDHGB2) | NM_032096 | 143.8 | 130.5 | 0.03748 | 0.894 |
| CREB1 | cAMP responsive element binding protein 1(CREB1) | NM_004379 | 128.9 | 222.5 | 0.03756 | 1.945 |
| UBXN6 | UBX domain protein 6(UBXN6) | NM_025241 | 534 | 616.5 | 0.03757 | 1.293 |
| SNAP23 | synaptosome associated protein 23(SNAP23) | NM_003825 | 130.1 | 127.7 | 0.03759 | 0.848 |
| SNAP23 | synaptosome associated protein 23(SNAP23) | NM_003825 | 188 | 159.5 | 0.03759 | 0.848 |
| CCZ1 | CCZ1 homolog, vacuolar protein trafficking and biogenesis associated(CCZ1) | NM_015622 | 1404.8 | 1039.2 | 0.03761 | 0.811 |
| | | XR_037525 | 501.1 | 353.6 | 0.03764 | 0.722 |
| | | XR_017589 | 227.2 | 170.5 | 0.03766 | 0.738 |
| | | XM_928982 | 96.2 | 117.8 | 0.03772 | 1.145 |
| DYRK1B | dual specificity tyrosine phosphorylation regulated kinase 1B(DYRK1B) | NM_006484 | 145 | 132.5 | 0.03774 | 0.886 |
| | | BU078529 | 120.8 | 137.8 | 0.03804 | 1.097 |

| | | | | | | | | | |
|---|---|---|---|---|---|---|---|---|---|
| DSP | desmoplakin(DSP) | NM_001008844 | 185.5 | 209.3 | 0.03811 | 1.906 | | | |
| DST | dystonin(DST) | NM_001723 | 195.6 | 312.7 | 0.03811 | 1.906 | | | |
| MAX | MYC associated factor X(MAX) | NM_002382 | 150.3 | 127.4 | 0.03814 | 0.839 | | | |
| MBTPS1 | membrane bound transcription factor peptidase, site 1(MBTPS1) | NM_003791 | 210.8 | 197.6 | 0.03838 | 0.84 | | | |
| MBTPS1 | membrane bound transcription factor peptidase, site 1(MBTPS1) | NM_003791 | 264.4 | 221.2 | 0.03838 | 0.84 | | | |
| MRPL47 | mitochondrial ribosomal protein L47(MRPL47) | NM_020409 | 181.6 | 147.8 | 0.03839 | 0.802 | NM_001033706 | 2.42 | 10.86 |
| | | XR_017481 | 146.9 | 129.8 | 0.03843 | 0.876 | | | |
| | | XR_015611 | 213.1 | 161.5 | 0.03843 | 0.754 | | | |
| C1R | complement C1r(C1R) | NM_001733 | 127.4 | 103.8 | 0.03859 | 0.791 | | | |
| | | XM_934846 | 144.2 | 174.7 | 0.03862 | 1.208 | | | |
| | | XM_001720379 | 792.6 | 491.2 | 0.03869 | 0.63 | | | |
| COMMD6 | COMM domain containing 6(COMMD6) | NM_203497 | 700.5 | 520 | 0.03893 | 0.795 | NM_153300 | 0.14 | 61.17 |
| | | XM_001130742 | 575 | 671.1 | 0.0391 | 1.465 | | | |

| | | | | | | | | | |
|---|---|---|---|---|---|---|---|---|---|
| | | XM_001130742 | 1793.8 | 2373.7 | 0.0391 | 1.465 | NM_032416 | 108.51 | 1.87 |
| | | XR_038930 | 7887.1 | 5231.1 | 0.03915 | 0.796 | | | |
| ACTR8 | ARP8 actin-related protein 8 homolog(ACTR8) | NM_022899 | 169.9 | 145.6 | 0.0392 | 0.855 | | | |
| EIF2B5 | eukaryotic translation initiation factor 2B subunit epsilon(EIF2B5) | NM_003907 | 450.7 | 360 | 0.03923 | 0.843 | | | |
| | | XR_017804 | 3711.5 | 2721.1 | 0.03927 | 0.745 | | | |
| | | XR_017804 | 6448.1 | 3895.8 | 0.03927 | 0.745 | | | |
| | | XM_001724539 | 729.2 | 435.7 | 0.03931 | 0.651 | | | |
| | | XR_038097 | 359.1 | 231.6 | 0.03941 | 0.672 | | | |
| KRT2 | keratin 2(KRT2) | NM_000423 | 280.2 | 409.7 | 0.03945 | 1.712 | | | |
| EIF2B4 | eukaryotic translation initiation factor 2B subunit delta(EIF2B4) | NM_172195 | 450.7 | 360 | 0.03947 | 0.842 | | | |
| EIF2B4 | eukaryotic translation initiation factor 2B subunit delta(EIF2B4) | NM_015636 | 398.2 | 321.2 | 0.03947 | 0.842 | | | |
| h2afy | H2A histone family member Y(H2AFY) | NM_138609 | 203.6 | 176 | 0.03952 | 0.79 | | | |
| H2AFY2 | H2A histone family member Y2(H2AFY2) | NM_018649 | 317.1 | 244.3 | 0.03952 | 0.79 | | | |

| | | | | | | | | | |
|---|---|---|---|---|---|---|---|---|---|
| | | XM_001715145 | 4374.5 | 5415.3 | 0.03959 | 1.418 | | | |
| CUTA | cutA divalent cation tolerance homolog(CUTA) | NM_001014840 | 172.7 | 223.9 | 0.03966 | 1.433 | | | |
| NCCRP1 | non-specific cytotoxic cell receptor protein 1 homolog (zebrafish)(NCCRP1) | NM_001001414 | 110.2 | 132.7 | 0.03967 | 1.163 | NM_001191088 | 10.07 | 0.03 |
| NCCRP1 | non-specific cytotoxic cell receptor protein 1 homolog (zebrafish)(NCCRP1) | NM_001001414 | 150.6 | 162.5 | 0.03967 | 1.163 | NM_001191088 | 10.07 | 0.03 |
| | | XM_943583 | 177.1 | 199.1 | 0.03969 | 1.141 | NM_001191088 | 10.07 | 0.03 |
| NIP7 | NIP7, nucleolar pre-rRNA processing protein(NIP7) | NM_016101 | 238 | 183.9 | 0.03981 | 0.791 | | | |
| IMPACT | impact RWD domain protein(IMPACT) | NM_018439 | 191.3 | 163.5 | 0.03986 | 0.853 | | | |
| | | XM_001725448 | 162.6 | 147.2 | 0.03987 | 0.884 | | | |
| | | XM_932890 | 350.2 | 472.2 | 0.0399 | 1.572 | NM_012496 | 1095.39 | 8.76 |
| SPAG9 | sperm associated antigen 9(SPAG9) | NM_003971 | 134.7 | 123.2 | 0.04018 | 0.887 | | | |
| SPAG9 | sperm associated antigen 9(SPAG9) | NM_003971 | 152.5 | 146.2 | 0.04018 | 0.887 | | | |
| RPL7 | ribosomal protein L7(RPL7) | NM_000971 | 456.2 | 334.9 | 0.04021 | 0.774 | | | |

| | | | | | | |
|---|---|---|---|---|---|---|
| TAGAP | T-cell activation RhoGTPase activating protein(TAGAP) | NM_054114 | 137.6 | 116.2 | 0.04031 | 0.825 |
| SIVA1 | SIVA1 apoptosis inducing factor(SIVA1) | NM_006427 | 265.6 | 244 | 0.04034 | 0.765 |
| SIVA1 | SIVA1 apoptosis inducing factor(SIVA1) | NM_006427 | 303.4 | 228.5 | 0.04034 | 0.765 |
| WAPL | WAPL cohesin release factor(WAPL) | NM_015045 | 274.2 | 229.6 | 0.04046 | 0.86 |
| IFRD2 | interferon related developmental regulator 2(IFRD2) | NM_006764 | 105.8 | 128 | 0.04062 | 1.151 |
| SPCS2P4 | signal peptidase complex subunit 2 homolog (S. cerevisiae) pseudogene 4(SPCS2P4) | NM_001080536 | 268.6 | 202 | 0.04067 | 0.741 |
| | | XM_001715568 | 146.4 | 164.2 | 0.04069 | 1.108 |
| SRGAP3 | SLIT-ROBO Rho GTPase activating protein 3(SRGAP3) | NM_001033117 | 124.3 | 157.2 | 0.04079 | 1.232 |
| SRGAP3 | SLIT-ROBO Rho GTPase activating protein 3(SRGAP3) | NM_001033117 | 195.8 | 246.7 | 0.04079 | 1.232 |
| MT1B | metallothionein 1B(MT1B) | NM_005947 | 1509.5 | 5464.9 | 0.04085 | 3.475 |
| ZNF583 | zinc finger protein 583(ZNF583) | NM_152478 | 136.1 | 121.5 | 0.04086 | 0.881 |
| NOP14 | NOP14 nucleolar protein(NOP14) | NM_003703 | 121.1 | 144.3 | 0.04086 | 1.152 |

| | | | | | | |
|---|---|---|---|---|---|---|
| FBXO30 | F-box protein 30(FBXO30) | NM_032145 | 136 | 116.2 | 0.04094 | 0.842 |
| FBXO32 | F-box protein 32(FBXO32) | NM_058229 | 154.8 | 154.4 | 0.04094 | 0.842 |
| OGFOD1 | 2-oxoglutarate and iron dependent oxygenase domain containing 1(OGFOD1) | NM_018233 | 128.5 | 113.4 | 0.04099 | 0.851 |
| OGFOD1 | 2-oxoglutarate and iron dependent oxygenase domain containing 1(OGFOD1) | NM_018233 | 132.2 | 128.5 | 0.04099 | 0.851 |
| DYRK1A | dual specificity tyrosine phosphorylation regulated kinase 1A(DYRK1A) | NM_130438 | 274.5 | 200.8 | 0.04102 | 0.716 |
| DYRK1A | dual specificity tyrosine phosphorylation regulated kinase 1A(DYRK1A) | NM_130437 | 135.2 | 138.6 | 0.04102 | 0.716 |
| RPL17P39 | ribosomal protein L17 pseudogene 39(RPL17P39) | NM_001093733 | 880.1 | 593.3 | 0.04125 | 0.735 |
| MCHR2 | melanin concentrating hormone receptor 2(MCHR2) | NM_032503 | 132.4 | 161.4 | 0.04131 | 1.191 |
| | | XM_001718864 | 238.7 | 158.9 | 0.04134 | 0.653 |
| | | XM_936640 | 120 | 137.9 | 0.0414 | 1.103 |

| | | | | | | |
|---|---|---|---|---|---|---|
| KLK10 | kallikrein related peptidase 10(KLK10) | NM_002776 | 576.6 | 407.6 | 0.04141 | 0.747 |
| FAM131B | family with sequence similarity 131 member B(FAM131B) | NM_014690 | 141.3 | 130 | 0.04142 | 0.894 |
| ZNF100 | zinc finger protein 100(ZNF100) | NM_173531 | 127.9 | 147.3 | 0.04142 | 1.119 |
| MANBAL | mannosidase beta like(MANBAL) | NM_001003897 | 250 | 286.6 | 0.04144 | 1.226 |
| BTBD6 | BTB domain containing 6(BTBD6) | NM_033271 | 187.2 | 158 | 0.04147 | 0.851 |
| NOSTRIN | nitric oxide synthase trafficking(NOSTRIN) | NM_052946 | 223.1 | 288.6 | 0.04147 | 1.434 |
| BTBD3 | BTB domain containing 3(BTBD3) | NM_014962 | 121.8 | 122.7 | 0.04147 | 0.851 |
| HAX1 | HCLS1 associated protein X-1(HAX1) | NM_006118 | 119.9 | 137.9 | 0.04151 | 1.107 |
| | | XM_001132706 | 123 | 113.9 | 0.04153 | 0.888 |
| TOPORS | TOP1 binding arginine/serine rich protein(TOPORS) | NM_005802 | 136.1 | 120.4 | 0.04162 | 0.858 |
| | | AI215519 | 149.9 | 167.1 | 0.04181 | 1.127 |
| | | BG058337 | 238 | 287.3 | 0.04181 | 1.391 |
| PAQR9 | progestin and adipoQ receptor family member 9(PAQR9) | NM_198504 | 128.3 | 264.7 | 0.04194 | 2.447 |
| ZFX | zinc finger protein, X-linked(ZFX) | NM_003410 | 127.2 | 115.6 | 0.04196 | 0.877 |

| | | | | | | | | | |
|---|---|---|---|---|---|---|---|---|---|
| ZSCAN2 | zinc finger and SCAN domain containing 2(ZSCAN2) | NM_017894 | 124.8 | 140.2 | 0.04197 | 1.084 | | | |
| | | XM_944905 | 142.5 | 159.9 | 0.0421 | 1.098 | | | |
| KRTAP4-7 | keratin associated protein 4-7(KRTAP4-7) | NM_033061 | 170.2 | 199.7 | 0.04217 | 1.181 | | | |
| NRG3 | neuregulin 3(NRG3) | NM_001010848 | 199.5 | 227.4 | 0.04225 | 1.213 | NM_013059 | 21.02 | 0.11 |
| | | XR_041205 | 1685.5 | 963.5 | 0.04225 | 0.66 | | | |
| ADAMTSL4 | ADAMTS like 4(ADAMTSL4) | NM_025008 | 172.3 | 150.6 | 0.04228 | 0.865 | | | |
| | | XM_0017157156 | 5234.3 | 3571.6 | 0.04231 | 0.82 | | | |
| CDYL2 | chromodomain Y-like 2(CDYL2) | AK023660 | 96.8 | 149.2 | 0.04232 | 1.591 | | | |
| MED1 | mediator complex subunit 1(MED1) | NM_004774 | 204.6 | 169.2 | 0.04239 | 0.827 | | | |
| MED1 | mediator complex subunit 1(MED1) | NM_004774 | 307.3 | 332 | 0.04239 | 0.827 | | | |
| RTN4IP1 | reticulon 4 interacting protein 1(RTN4IP1) | NM_032730 | 190.4 | 217.2 | 0.04241 | 1.158 | | | |
| FNBP1L | formin binding protein 1 like(FNBP1L) | NM_001024948 | 122.4 | 146.7 | 0.04244 | 1.138 | | | |
| | | XR_037721 | 388.4 | 306.5 | 0.04263 | 0.837 | | | |
| LY86 | lymphocyte antigen 86(LY86) | NM_004271 | 110 | 145.1 | 0.04263 | 1.304 | | | |
| KIAA0319L | KIAA0319 like(KIAA0319L) | NM_024874 | 329 | 264.8 | 0.04271 | 0.82 | | | |

| | | | | | | | | | |
|---|---|---|---|---|---|---|---|---|---|
| | | XM_001133527 | 123.9 | 127.7 | 0.04274 | 1.157 | | | |
| | | XM_001133527 | 142.3 | 165.4 | 0.04274 | 1.157 | | | |
| | | XM_001726456 | 155 | 139.4 | 0.04277 | 0.889 | NM_012816 | 19.78 | 2.28 |
| ZNF283 | zinc finger protein 283(ZNF283) | NM_181845 | 114.2 | 106.6 | 0.0428 | 0.878 | NM_012816 | 19.78 | 2.28 |
| HERPUD1 | homocysteine inducible ER protein with ubiquitin like domain 1(HERPUD1) | NM_001010990 | 278.9 | 211.9 | 0.04286 | 0.775 | | | |
| AATK | apoptosis associated tyrosine kinase(AATK) | NM_001080395 | 323.6 | 368.1 | 0.0429 | 1.24 | | | |
| | | XM_944968 | 107.3 | 125.4 | 0.04293 | 1.095 | | | |
| QRFPR | pyroglutamylated RFamide peptide receptor(QRFPR) | NM_198179 | 1460.7 | 1944.9 | 0.04295 | 1.456 | | | |
| LRRC3DN | LRRC3 downstream neighbor (non-protein coding)(LRRC3DN) | AL117578 | 125.1 | 118.2 | 0.04296 | 0.902 | | | |
| MEIG1 | meiosis/spermiogenesis associated 1(MEIG1) | NM_001080836 | 129.8 | 146.9 | 0.04297 | 1.102 | | | |
| MOCS1 | molybdenum cofactor synthesis 1(MOCS1) | NM_005943 | 234.1 | 268.7 | 0.04301 | 1.237 | | | |
| CLK1 | CDC like kinase 1(CLK1) | NM_004071 | 219.6 | 172.9 | 0.04302 | 0.787 | | | |
| SNRPD2 | small nuclear ribonucleoprotein D2 polypeptide(SNRPD2) | NM_004597 | 221.5 | 182.7 | 0.04304 | 0.818 | | | |

| | | | | | | | | | |
|---|---|---|---|---|---|---|---|---|---|
| SNORA18 | small nucleolar RNA, H/ACA box 18(SNORA18) | NR_002959 | 172 | 154.4 | 0.04316 | 0.892 | | | |
| IL17RE | interleukin 17 receptor E(IL17RE) | NM_153483 | 120.4 | 137.7 | 0.04321 | 1.093 | | | |
| MYO3B | myosin IIIB(MYO3B) | NM_138995 | 696.5 | 949 | 0.04324 | 1.567 | | | |
| | | XM_001721589 | 739.6 | 475.3 | 0.04328 | 0.679 | | | |
| SRSF1 | serine and arginine rich splicing factor 1(SRSF1) | NM_001078166 | 1563.3 | 1172.5 | 0.0433 | 0.803 | | | |
| | | XM_933437 | 161.6 | 189.5 | 0.0433 | 1.176 | | | |
| | | XR_018316 | 715.5 | 385.9 | 0.0434 | 0.559 | | | |
| | | XM_001732883 | 123.4 | 112.3 | 0.04342 | 0.878 | | | |
| HDLBP | high density lipoprotein binding protein(HDLBP) | NM_203346 | 312.5 | 375 | 0.04353 | 1.404 | | | |
| HDLBP | high density lipoprotein binding protein(HDLBP) | NM_005336 | 111.9 | 118.4 | 0.04353 | 1.404 | | | |
| | | XM_936822 | 199.4 | 156.9 | 0.04358 | 0.781 | | | |
| | | XR_015495 | 137.6 | 122.6 | 0.04358 | 0.87 | NM_031544 | 4.79 | 38.59 |
| SERPINB7 | serpin family B member 7(SERPINB7) | NM_003784 | 123.3 | 145.1 | 0.04377 | 1.234 | NM_031544 | 4.79 | 38.59 |

| | | | | | | |
|---|---|---|---|---|---|---|
| SERPINB7 | serpin family B member 7(SERPINB7) | NM_003784 | 171.6 | 205.1 | 0.04377 | 1.234 |
| FKBP14 | FK506 binding protein 14(FKBP14) | NM_017946 | 142.6 | 156 | 0.04379 | 1.371 |
| FKBP1A | FK506 binding protein 1A(FKBP1A) | NM_000801 | 2187.3 | 2702.4 | 0.04379 | 1.371 |
| PPA2 | pyrophosphatase (inorganic) 2(PPA2) | NM_006903 | 4900.5 | 5967 | 0.04389 | 1.367 |
| RPL21P16 | ribosomal protein L21 pseudogene 16(RPL21P16) | NM_001139505 | 117 | 104.5 | 0.04391 | 0.864 |
| | | XR_037860 | 625.2 | 358.5 | 0.04396 | 0.608 |
| CDCA4 | cell division cycle associated 4(CDCA4) | NM_145701 | 134.6 | 101.2 | 0.04402 | 0.741 |
| | | XR_038543 | 5646.2 | 3974 | 0.04404 | 0.775 |
| RAP1GDS1 | Rap1 GTPase-GDP dissociation stimulator 1(RAP1GDS1) | NM_021159 | 215.8 | 175.3 | 0.04412 | 0.831 |
| TRAPPC2 | trafficking protein particle complex 2(TRAPPC2) | NM_001011658 | 140.7 | 131.4 | 0.04412 | 0.918 |
| | | XR_039407 | 620.8 | 465 | 0.04422 | 0.801 |
| NANS | N-acetylneuraminate synthase(NANS) | NM_018946 | 223.9 | 169 | 0.04426 | 0.748 |
| SEC23A | Sec23 homolog A, coat complex II component(SEC23A) | NM_006364 | 115.2 | 133.6 | 0.04427 | 1.109 |

| | | | | | | |
|---|---|---|---|---|---|---|
| SEC23A | Sec23 homolog A, coat complex II component(SEC23A) | NM_006364 | 132.4 | 130.5 | 0.04427 | 1.109 |
| ADAMDEC1 | ADAM like decysin 1(ADAMDEC1) | NM_014479 | 121.7 | 106.7 | 0.04427 | 0.86 |
| RHEB | Ras homolog enriched in brain(RHEB) | NM_005614 | 376 | 314.4 | 0.0443 | 0.861 |
| RNA5S9 | RNA, 5S ribosomal 9(RNA5S9) | NR_023371 | 186.3 | 225.7 | 0.0443 | 1.246 |
| RAX2 | retina and anterior neural fold homeobox 2(RAX2) | NM_032753 | 174.1 | 214.2 | 0.0444 | 1.273 |
| RAX2 | retina and anterior neural fold homeobox 2(RAX2) | NM_032753 | 1050.1 | 1081.8 | 0.0444 | 1.273 |
| XRCC2 | X-ray repair cross complementing 2(XRCC2) | NM_005431 | 118.1 | 123.6 | 0.04448 | 1.462 |
| XRCC2 | X-ray repair cross complementing 2(XRCC2) | NM_005431 | 1119.2 | 1461.7 | 0.04448 | 1.462 |
| | | XR_015615 | 212.3 | 247.8 | 0.04478 | 1.255 |
| | | XR_038081 | 104.2 | 124.1 | 0.04482 | 1.134 |
| LNPEP | leucyl and cystinyl aminopeptidase(LNPEP) | NM_005575 | 1079.3 | 1296.6 | 0.04486 | 1.342 |
| BNIP1 | BCL2 interacting protein 1(BNIP1) | NM_013980 | 2942.2 | 4061.6 | 0.04487 | 1.536 |

| | | | | | | |
|---|---|---|---|---|---|---|
| | | NM_0010 80831 | 206.3 | 168 | 0.0449 | 0.82 |
| AKTIP | AKT interacting protein(AKTIP) | NM_0010 12398 | 117.9 | 111.6 | 0.04493 | 0.881 |
| SUGP1 | SURP and G-patch domain containing 1(SUGP1) | NM_1828 12 | 189.5 | 213.6 | 0.04506 | 1.172 |
| | | XR_03711 7 | 201.4 | 150.5 | 0.0452 | 0.664 |
| | | XR_03711 7 | 1539.6 | 919.6 | 0.0452 | 0.664 |
| | | XM_93746 9 | 111.1 | 133.5 | 0.04522 | 1.142 |
| | | XM_00172 5687 | 1333 | 862.6 | 0.04531 | 0.72 |
| | | BM977869 | 133.1 | 121.8 | 0.04538 | 0.89 |
| ZNF654 | zinc finger protein 654(ZNF654) | NM_0182 93 | 99.8 | 106.1 | 0.04544 | 0.876 |
| ZNF654 | zinc finger protein 654(ZNF654) | NM_0182 93 | 116.2 | 112.9 | 0.04544 | 0.876 |
| ZNF654 | zinc finger protein 654(ZNF654) | NM_0182 93 | 120.1 | 110.9 | 0.04544 | 0.876 |
| ZNF654 | zinc finger protein 654(ZNF654) | NM_0182 93 | 130.5 | 119.2 | 0.04544 | 0.876 |
| | | XM_04529 0 | 611.7 | 424.1 | 0.04551 | 0.745 |
| ZDHHC1 | zinc finger DHHC-type containing 1(ZDHHC1) | NM_0133 04 | 240.9 | 295.7 | 0.04553 | 1.287 |
| DMC1 | DNA meiotic recombinase 1(DMC1) | NM_0070 68 | 321.1 | 355.3 | 0.04558 | 1.21 |

| | | | | | | |
|---|---|---|---|---|---|---|
| RPS21 | ribosomal protein S21(RPS21) | NM_001024 | 165.5 | 149.7 | 0.04564 | 0.898 |
| | | XM_001719081 | 312.4 | 211.5 | 0.04572 | 0.682 |
| | | XM_935907 | 658.5 | 437.2 | 0.04581 | 0.702 |
| SLC10A7 | solute carrier family 10 member 7(SLC10A7) | NM_001029998 | 132.8 | 127.5 | 0.04595 | 0.932 |
| SLC10A7 | solute carrier family 10 member 7(SLC10A7) | NM_001029998 | 150.1 | 141.2 | 0.04595 | 0.932 |
| | | BU622756 | 122.4 | 141.6 | 0.04598 | 1.108 |
| SNHG8 | small nucleolar RNA host gene 8(SNHG8) | NR_003584 | 607.7 | 410.6 | 0.046 | 0.724 |
| ZKSCAN3 | zinc finger with KRAB and SCAN domains 3(ZKSCAN3) | NM_024493 | 132.8 | 124.1 | 0.04603 | 0.896 |
| NECAB2 | N-terminal EF-hand calcium binding protein 2(NECAB2) | NM_019065 | 166.7 | 183.6 | 0.04606 | 1.104 |
| KYAT1 | kynurenine aminotransferase 1(KYAT1) | NM_004059 | 155.2 | 136.1 | 0.04607 | 0.857 |
| MFSD14C | major facilitator superfamily domain containing 14C(MFSD14C) | NR_002894 | 3646.4 | 4446.2 | 0.04622 | 1.378 |
| GLE1 | GLE1, RNA export mediator(GLE1) | NM_001003722 | 468.2 | 977 | 0.04625 | 2.408 |
| | | XM_929297 | 131.5 | 147.9 | 0.04627 | 1.092 |

| | | | | | | |
|---|---|---|---|---|---|---|
| CBX5 | chromobox 5(CBX5) | NM_012117 | 173.6 | 145.5 | 0.04627 | 0.821 |
| TPM1 | tropomyosin 1 (alpha)(TPM1) | NM_001018008 | 130.7 | 155.5 | 0.04645 | 1.158 |
| TPM1 | tropomyosin 1 (alpha)(TPM1) | NM_001018008 | 141.1 | 143.1 | 0.04645 | 1.158 |
| GRK3 | G protein-coupled receptor kinase 3(GRK3) | NM_005160 | 1794 | 2516.4 | 0.04659 | 1.552 |
| RANBP1 | RAN binding protein 1(RANBP1) | NM_002882 | 344.3 | 225 | 0.04659 | 0.677 |
| RANBP1 | RAN binding protein 1(RANBP1) | NM_002882 | 879.4 | 649.2 | 0.04659 | 0.677 |
| ANXA1 | annexin A1(ANXA1) | NM_000700 | 277.1 | 186 | 0.04679 | 0.663 |
| | | XR_040497 | 152.9 | 188.2 | 0.04695 | 1.212 |
| | | XM_939730 | 126.9 | 148.6 | 0.04695 | 1.123 |
| ZNF549 | zinc finger protein 549(ZNF549) | NM_153263 | 178.3 | 196.9 | 0.04698 | 1.137 |
| ZNF549 | zinc finger protein 549(ZNF549) | NM_153263 | 6038.9 | 7142.4 | 0.04698 | 1.137 |
| | | XM_927228 | 137.4 | 131.4 | 0.04711 | 0.927 |
| | | BI827719 | 117.1 | 135.8 | 0.04713 | 1.102 |
| GUSB | glucuronidase beta(GUSB) | NM_000181 | 454.5 | 489.2 | 0.04715 | 1.158 |
| HMGN3 | high mobility group nucleosomal binding domain 3(HMGN3) | NM_004242 | 1643.4 | 877.1 | 0.04716 | 0.603 |

| | | | | | | |
|---|---|---|---|---|---|---|
| HMGN2 | high mobility group nucleosomal binding domain 2(HMGN2) | NM_005517 | 177.1 | 154.4 | 0.04716 | 0.603 |
| AMACR | alpha-methylacyl-CoA racemase(AMACR) | NM_203382 | 168.3 | 125.4 | 0.04725 | 0.726 |
| SPOPL | speckle type BTB/POZ protein like(SPOPL) | NM_001001664 | 148.1 | 127.6 | 0.04729 | 0.849 |
| | | XM_001719439 | 302.5 | 194.1 | 0.04737 | 0.638 |
| | | XR_039723 | 346 | 218.3 | 0.04738 | 0.652 |
| CDC123 | cell division cycle 123(CDC123) | NM_006023 | 126.8 | 148.1 | 0.04739 | 1.139 |
| ZNF600 | zinc finger protein 600(ZNF600) | NM_198457 | 243.7 | 303.9 | 0.04742 | 1.353 |
| PIGO | phosphatidylinositol glycan anchor biosynthesis class O(PIGO) | NM_152850 | 207.2 | 251.3 | 0.04745 | 1.247 |
| WBP11 | WW domain binding protein 11(WBP11) | NM_016312 | 436.2 | 358.1 | 0.04749 | 0.867 |
| PDCD7 | programmed cell death 7(PDCD7) | NM_005707 | 333.5 | 266.3 | 0.04761 | 0.828 |
| PDCD7 | programmed cell death 7(PDCD7) | NM_005707 | 24825.1 | 30661.9 | 0.04761 | 0.828 |
| TRPV5 | transient receptor potential cation channel subfamily V member 5(TRPV5) | NM_019841 | 189.7 | 224.8 | 0.04764 | 1.241 |
| tfcp2 | transcription factor CP2(TFCP2) | NM_005653 | 206.7 | 170.4 | 0.0477 | 0.837 |

| | | | | | | |
|---|---|---|---|---|---|---|
| | | XM_944813 | 121.3 | 138.3 | 0.04772 | 1.084 |
| | | XM_943208 | 145.5 | 159.1 | 0.04772 | 1.081 |
| SH2D1A | SH2 domain containing 1A(SH2D1A) | NM_002351 | 120.3 | 114.5 | 0.04772 | 0.895 |
| YPEL2 | yippee like 2(YPEL2) | NM_001005404 | 160 | 141.3 | 0.04795 | 0.867 |
| GET4 | golgi to ER traffic protein 4(GET4) | NM_015949 | 102.4 | 126.3 | 0.04797 | 1.171 |
| RPS7 | ribosomal protein S7(RPS7) | NM_001011 | 145.4 | 127.6 | 0.04804 | 0.852 |
| RPS7 | ribosomal protein S7(RPS7) | NM_001011 | 194.3 | 202.8 | 0.04804 | 0.852 |
| | | XM_0017222060 | 1000.1 | 1216.9 | 0.04813 | 1.363 |
| CCNG2 | cyclin G2(CCNG2) | NM_004354 | 125 | 116.1 | 0.04814 | 0.678 |
| CCNH | cyclin H(CCNH) | NM_001239 | 226.2 | 154.9 | 0.04814 | 0.678 |
| RPS8 | ribosomal protein S8(RPS8) | NM_001012 | 1470.6 | 725 | 0.04814 | 0.592 |
| | | XM_0017155097 | 129.9 | 149.6 | 0.04821 | 1.136 |
| HNRNPAB | heterogeneous nuclear ribonucleoprotein A/B(HNRNPAB) | NM_004499 | 146.7 | 132 | 0.04826 | 0.877 |
| SNRPF | small nuclear ribonucleoprotein polypeptide F(SNRPF) | NM_003095 | 641.8 | 423.4 | 0.04829 | 0.681 |

| | | | | | | | | | |
|---|---|---|---|---|---|---|---|---|---|
| SNRPF | small nuclear ribonucleoprotein polypeptide F(SNRPF) | NM_003095 | 1328.4 | 906.8 | 0.04829 | 0.681 | | | |
| DSC3 | desmocollin 3(DSC3) | NM_024423 | 111.2 | 182.1 | 0.04845 | 1.811 | | | |
| TMEM54 | transmembrane protein 54(TMEM54) | NM_033504 | 145.8 | 179.7 | 0.04849 | 1.251 | | | |
| CDK7 | cyclin dependent kinase 7(CDK7) | NM_001799 | 719.1 | 859.7 | 0.04854 | 1.352 | | | |
| SGSM2 | small G protein signaling modulator 2(SGSM2) | NM_014853 | 912.5 | 950.8 | 0.04855 | 1.16 | | | |
| | | XM_939709 | 186 | 214.1 | 0.04856 | 1.229 | | | |
| ERAP2 | endoplasmic reticulum aminopeptidase 2(ERAP2) | NM_022350 | 206.1 | 179.1 | 0.0487 | 1.453 | | | |
| ERBB2 | erb-b2 receptor tyrosine kinase 2(ERBB2) | NM_004448 | 5897.6 | 7862.8 | 0.0487 | 1.453 | | | |
| | | DB336496 | 165.9 | 196.3 | 0.04875 | 1.22 | | | |
| NOC4L | nucleolar complex associated 4 homolog(NOC4L) | NM_024078 | 117.2 | 136.9 | 0.04876 | 1.124 | | | |
| NOC4L | nucleolar complex associated 4 homolog(NOC4L) | NM_024078 | 177.9 | 206.1 | 0.04876 | 1.124 | NM_031012 | 19.39 | 0.85 |
| | | XR_039555 | 179.2 | 126.1 | 0.04876 | 0.683 | | | |
| RPL23 | ribosomal protein L23(RPL23) | NM_000978 | 5124.5 | 3196.8 | 0.04884 | 0.747 | | | |

| | | | | | | | | | |
|---|---|---|---|---|---|---|---|---|---|
| ACTL6A | actin like 6A(ACTL6A) | NM_004301 | 1214.3 | 784.4 | 0.04888 | 0.684 | | | |
| OSTC | oligosaccharyltransferase complex non-catalytic subunit(OSTC) | NM_021227 | 199.2 | 178.3 | 0.04891 | 0.751 | NM_012904 | 10.16 | 55.22 |
| OSTC | oligosaccharyltransferase complex non-catalytic subunit(OSTC) | NM_021227 | 757.8 | 544.9 | 0.04891 | 0.751 | | | |
| BAIAP2 | BAI1 associated protein 2(BAIAP2) | NM_006340 | 127.9 | 121.8 | 0.04894 | 0.918 | | | |
| CRHR1 | corticotropin releasing hormone receptor 1(CRHR1) | NM_004382 | 99.8 | 151.5 | 0.04895 | 1.552 | | | |
| | | AI738931 | 137.8 | 172.3 | 0.04898 | 1.258 | | | |
| MBLAC2 | metallo-beta-lactamase domain containing 2(MBLAC2) | NM_203406 | 127.6 | 120.1 | 0.04907 | 0.915 | | | |
| LSM5 | LSM5 homolog, U6 small nuclear RNA and mRNA degradation associated(LSM5) | NM_012322 | 177.6 | 142.4 | 0.04934 | 0.796 | | | |
| LSM7 | LSM7 homolog, U6 small nuclear RNA and mRNA degradation associated(LSM7) | NM_016199 | 566.5 | 466.3 | 0.04934 | 0.796 | | | |
| rps15a | ribosomal protein S15a(RPS15A) | NM_001019 | 182.9 | 136.5 | 0.04938 | 0.741 | | | |
| rps15a | ribosomal protein S15a(RPS15A) | NM_001019 | 1616 | 913.9 | 0.04938 | 0.741 | | | |
| RPS14 | ribosomal protein S14(RPS14) | NM_001025070 | 8805.8 | 5307.3 | 0.04948 | 0.725 | | | |

| | | | | | | | | | |
|---|---|---|---|---|---|---|---|---|---|
| | | XM_001724343 | 839.5 | 430.9 | 0.04948 | 0.571 | | | |
| | | XM_928464 | 435.7 | 340.4 | 0.0495 | 0.831 | NM_012823 | 17.52 | 0.04 |
| | | XR_038937 | 954.2 | 614.2 | 0.04961 | 0.694 | | | |
| DRD3 | dopamine receptor D3(DRD3) | NM_000796 | 113.3 | 125.7 | 0.04964 | 0.895 | | | |
| DRG1 | developmentally regulated GTP binding protein 1(DRG1) | NM_004147 | 146.9 | 133.5 | 0.04964 | 0.895 | | | |
| | | XR_038410 | 135.6 | 120.5 | 0.04965 | 0.866 | | | |
| CCDC53 | coiled-coil domain containing 53(CCDC53) | NM_016053 | 365.4 | 244.6 | 0.04973 | 0.687 | | | |
| TBCE | tubulin folding cofactor E(TBCE) | NM_001079515 | 415.5 | 461.1 | 0.04975 | 1.225 | | | |
| | | XR_017366 | 139 | 124.6 | 0.04983 | 0.886 | | | |
| | | XM_930344 | 500.6 | 383.9 | 0.04997 | 0.799 | | | |
| RHBDD3 | rhomboid domain containing 3(RHBDD3) | NM_012265 | 207.7 | 257.5 | 0.05 | 1.319 | | | |

| Gene Symbol | Gene Name | Accession No. | CON Chow M Mean | CON HFHS M Mean | p-val CON M HFHS vs. CON M Chow | Ratio CON M HFHS vs. CON M Chow | REFSEQ ID | NEPHRON (MEAN RPKM) | COLLECTING DUCT (MEAN RPKM) |
|---|---|---|---|---|---|---|---|---|---|
| | | XM_927511 | 115.2 | 97.2 | 0.00001 | 0.813 | | | |

| | | | | | | | | | |
|---|---|---|---|---|---|---|---|---|---|
| ADCK2 | aarF domain containing kinase 2(ADCK2) | NM_052853 | 107.3 | 133.3 | 0.00016 | 1.171 | | | |
| RPS26 | ribosomal protein S26(RPS26) | NM_001029 | 89.2 | 109.3 | 0.00018 | 1.183 | | | |
| RPS26 | ribosomal protein S26(RPS26) | NM_001029 | 1360.8 | 1666 | 0.00018 | 1.183 | | | |
| C3 | complement C3(C3) | NM_000064 | 116.5 | 99.5 | 0.00024 | 0.805 | | | |
| SLC38A3 | solute carrier family 38 member 3(SLC38A3) | NM_006841 | 210.4 | 129.1 | 0.00044 | 0.564 | | | |
| | | XM_372319 | 144.1 | 116.9 | 0.00046 | 0.762 | | | |
| KLF9 | Kruppel like factor 9(KLF9) | NM_001206 | 332.5 | 774.2 | 0.00046 | 1.981 | | | |
| MTHFD2L | methylenetetrahydrofolate dehydrogenase (NADP+ dependent) 2-like(MTHFD2L) | NM_001004346 | 95.7 | 122.6 | 0.00063 | 1.219 | NM_017193 | 356.13 | 8.18 |
| TSPAN1 | tetraspanin 1(TSPAN1) | NM_005727 | 96 | 132.6 | 0.00076 | 1.291 | NM_017193 | 356.13 | 8.18 |
| | | XM_0017155244 | 115.4 | 105.2 | 0.00082 | 0.864 | NM_017193 | 356.13 | 8.18 |
| SNRPC | small nuclear ribonucleoprotein polypeptide C(SNRPC) | NM_003093 | 237.3 | 363.8 | 0.00102 | 1.336 | NM_017193 | 356.13 | 8.18 |
| SNRPC | small nuclear ribonucleoprotein polypeptide C(SNRPC) | NM_003093 | 355.1 | 429.4 | 0.00102 | 1.336 | | | |
| PHF14 | PHD finger protein 14(PHF14) | NM_014660 | 120.2 | 109.4 | 0.00107 | 0.883 | | | |

| | | | | | | |
|---|---|---|---|---|---|---|
| | | XM_001723369 | 125.5 | 110.8 | 0.00135 | 0.85 |
| ATP6V0E2 | ATPase H+ transporting V0 subunit e2(ATP6V0E2) | NM_145230 | 1451.1 | 1696 | 0.00139 | 1.281 |
| ATP6V0E1 | ATPase H+ transporting V0 subunit e1(ATP6V0E1) | NM_003945 | 625.7 | 926.7 | 0.00139 | 1.281 |
| UGGT2 | UDP-glucose glycoprotein glucosyltransferase 2(UGGT2) | NM_020121 | 135.8 | 193.7 | 0.00139 | 1.276 |
| EFNB2 | ephrin B2(EFNB2) | NM_004093 | 99.5 | 121 | 0.00143 | 1.17 |
| IRF2BP2 | interferon regulatory factor 2 binding protein 2(IRF2BP2) | NM_182972 | 165.2 | 239.2 | 0.00163 | 1.302 |
| IRF2BPL | interferon regulatory factor 2 binding protein like(IRF2BPL) | NM_024496 | 218.4 | 267.1 | 0.00163 | 1.302 |
| KCNH4 | potassium voltage-gated channel subfamily H member 4(KCNH4) | NM_012285 | 108.7 | 135 | 0.002 | 1.181 |
| BZW1 | basic leucine zipper and W2 domains 1(BZW1) | NM_014670 | 124.8 | 114.3 | 0.00206 | 0.878 |
| MYLK | myosin light chain kinase(MYLK) | NM_053026 | 142.6 | 204 | 0.00209 | 1.297 |
| | | XM_929153 | 99.5 | 131.8 | 0.00212 | 1.246 |

| | | | | | | |
|---|---|---|---|---|---|---|
| ADPRHL2 | ADP-ribosylhydrolase like 2(ADPRHL2) | NM_017825 | 145.2 | 133.9 | 0.00222 | 0.856 |
| SHARPIN | SHANK associated RH domain interactor(SHARPIN) | NM_030974 | 114.9 | 152.9 | 0.00224 | 1.261 |
| | | XM_933942 | 144.2 | 131.5 | 0.00226 | 0.86 |
| CDAN1 | codanin 1(CDAN1) | NM_138477 | 200.4 | 163.3 | 0.00254 | 0.732 |
| POTEE | POTE ankyrin domain family member E(POTEE) | NM_001083538 | 102.3 | 133.2 | 0.00259 | 1.239 |
| C8orf59 | chromosome 8 open reading frame 59(C8orf59) | NM_001099672 | 268.8 | 691.5 | 0.00279 | 2.227 |
| PHF7 | PHD finger protein 7(PHF7) | NM_016483 | 116.6 | 108.1 | 0.00286 | 0.891 |
| ARSB | arylsulfatase B(ARSB) | NM_000046 | 124.2 | 109.9 | 0.00294 | 0.848 |
| ARSD | arylsulfatase D(ARSD) | NM_001669 | 149.8 | 144.3 | 0.00294 | 0.848 |
| NEDD8 | neural precursor cell expressed, developmentally down-regulated 8(NEDD8) | NM_006156 | 265.5 | 457.3 | 0.00295 | 1.49 |
| NEDD8 | neural precursor cell expressed, developmentally down-regulated 8(NEDD8) | NM_006156 | 2116.6 | 2390.9 | 0.00295 | 1.49 |
| Amy1a | amylase, alpha 1A (salivary)(AMY1A) | NM_001008221 | 94 | 117 | 0.00296 | 1.191 |

| | | | | | | |
|---|---|---|---|---|---|---|
| KALRN | kalirin, RhoGEF kinase(KALRN) | NM_003947 | 3577.2 | 5517.9 | 0.00297 | 1.382 |
| LRBA | LPS responsive beige-like anchor protein(LRBA) | NM_006726 | 104.3 | 129.4 | 0.00298 | 1.196 |
| | | XM_938867 | 98.8 | 117.9 | 0.003 | 1.168 |
| LYPD6B | LY6/PLAUR domain containing 6B(LYPD6B) | NM_177964 | 135.8 | 123 | 0.00306 | 0.861 |
| CES5A | carboxylesterase 5A(CES5A) | NM_145024 | 120.8 | 105 | 0.00307 | 0.818 |
| | | XM_945223 | 118 | 104.2 | 0.00314 | 0.851 |
| CPNE1 | copine 1(CPNE1) | NM_152931 | 96.3 | 119.8 | 0.00315 | 1.186 |
| | | AI082160 | 151 | 131.1 | 0.00332 | 0.818 |
| CASP6 | caspase 6(CASP6) | NM_001226 | 94.5 | 118 | 0.00333 | 1.193 |
| CASP6 | caspase 6(CASP6) | NM_032992 | 108.7 | 115.2 | 0.00333 | 1.193 |
| RSRC2 | arginine and serine rich coiled-coil 2(RSRC2) | NM_023012 | 127.2 | 139.8 | 0.00359 | 1.329 |
| RSRC2 | arginine and serine rich coiled-coil 2(RSRC2) | NM_023012 | 585.6 | 906.9 | 0.00359 | 1.329 |
| TIGD1 | tigger transposable element derived 1(TIGD1) | NM_145702 | 107.1 | 98.1 | 0.0036 | 0.865 |
| SLCO3A1 | solute carrier organic anion transporter family member 3A1(SLCO3A1) | NM_013272 | 98.3 | 138.1 | 0.00374 | 1.349 |

| | | | | | | |
|---|---|---|---|---|---|---|
| SLCO3A1 | solute carrier organic anion transporter family member 3A1(SLCO3A1) | NM_013272 | 130.5 | 204.6 | 0.00374 | 1.349 |
| | | XM_001717918 | 126.8 | 112.3 | 0.00389 | 0.848 |
| bag4 | BCL2 associated athanogene 4(BAG4) | NM_004874 | 120.2 | 107 | 0.00402 | 0.852 |
| ARMCX6 | armadillo repeat containing, X-linked 6(ARMCX6) | NM_019007 | 99 | 114.4 | 0.00404 | 1.292 |
| ARMT1 | acidic residue methyltransferase 1(ARMT1) | NM_024573 | 115.1 | 160.1 | 0.00404 | 1.292 |
| | | XM_927848 | 123 | 149.3 | 0.00405 | 1.158 |
| CNRIP1 | cannabinoid receptor interacting protein 1(CNRIP1) | NM_015463 | 96.3 | 119.1 | 0.00412 | 1.194 |
| SCARNA12 | small Cajal body-specific RNA 12(SCARNA12) | NR_003010 | 128 | 164 | 0.00432 | 1.201 |
| | | XM_933083 | 143.5 | 127.5 | 0.00439 | 0.848 |
| BPHL | biphenyl hydrolase like(BPHL) | NM_004332 | 106.6 | 95.8 | 0.00458 | 0.852 |
| MIR21 | microRNA 21(MIR21) | NR_029493 | 104 | 177.3 | 0.00469 | 1.562 |

| | | | | | | |
|---|---|---|---|---|---|---|
| NPIPB15 | nuclear pore complex interacting protein family member B15(NPIPB15) | NM_001018059 | 118.4 | 107.9 | 0.00493 | 0.86 |
| RTBDN | retbindin(RTBDN) | NM_001080997 | 130.6 | 122.6 | 0.00515 | 0.897 |
| KHDRBS3 | KH RNA binding domain containing, signal transduction associated 3(KHDRBS3) | NM_006558 | 558.2 | 965.9 | 0.00558 | 1.485 |
| ODF3L2 | outer dense fiber of sperm tails 3 like 2(ODF3L2) | NM_182577 | 137.8 | 124 | 0.00563 | 0.854 |
| MS4A6A | membrane spanning 4-domains A6A(MS4A6A) | NM_022349 | 98.3 | 118.2 | 0.00567 | 1.16 |
| VRK3 | vaccinia related kinase 3(VRK3) | NM_001025778 | 106.9 | 104.7 | 0.00571 | 0.867 |
| VRK3 | vaccinia related kinase 3(VRK3) | NM_001025778 | 167.3 | 155.2 | 0.00571 | 0.867 |
| NFX1 | nuclear transcription factor, X-box binding 1(NFX1) | NM_147133 | 121.5 | 116.9 | 0.00592 | 0.866 |
| NFX1 | nuclear transcription factor, X-box binding 1(NFX1) | NM_147133 | 181.8 | 169 | 0.00592 | 0.866 |
| | | AV649055 | 126.1 | 113.6 | 0.00593 | 0.854 |
| SON | SON DNA binding protein(SON) | NM_138927 | 119.6 | 153.9 | 0.00594 | 1.189 |
| SON | SON DNA binding protein(SON) | NM_138927 | 171.9 | 234 | 0.00594 | 1.189 |

| | | | | | | | | | |
|---|---|---|---|---|---|---|---|---|---|
| SON | SON DNA binding protein(SON) | NM_138927 | 1267.9 | 1431.2 | 0.00594 | 1.189 | NM_053502 | 4.02 | 18.01 |
| CLDN14 | claudin 14(CLDN14) | NM_012130 | 124.8 | 393.1 | 0.00605 | 2.736 | NM_053502 | 4.02 | 18.01 |
| CHRNA6 | cholinergic receptor nicotinic alpha 6 subunit(CHRNA6) | NM_004198 | 116.5 | 103.1 | 0.00626 | 0.82 | NM_053502 | 4.02 | 18.01 |
| BCLAF1 | BCL2 associated transcription factor 1(BCLAF1) | NM_001077440 | 107.8 | 128.6 | 0.00629 | 1.504 | NM_053502 | 4.02 | 18.01 |
| BCO1 | beta-carotene oxygenase 1(BCO1) | NM_017429 | 556.2 | 960.4 | 0.00629 | 1.504 | | | |
| dnase2 | deoxyribonuclease 2, lysosomal(DNASE2) | NM_001375 | 107.8 | 132.5 | 0.00635 | 1.163 | | | |
| ZFP36L1 | ZFP36 ring finger protein like 1(ZFP36L1) | NM_004926 | 802.6 | 1633.2 | 0.00639 | 1.854 | | | |
| | | XM_931775 | 134.4 | 125.6 | 0.00661 | 0.887 | | | |
| TICAM1 | toll like receptor adaptor molecule 1(TICAM1) | NM_182919 | 94.4 | 122.5 | 0.00681 | 1.255 | | | |
| TICAM1 | toll like receptor adaptor molecule 1(TICAM1) | NM_182919 | 114.1 | 118.3 | 0.00681 | 1.255 | | | |
| | | AW274669 | 124.3 | 110.5 | 0.00689 | 0.861 | NM_001009670 | 33.30 | 2.04 |
| RUNDC1 | RUN domain containing 1(RUNDC1) | NM_173079 | 97.3 | 127.8 | 0.00692 | 1.262 | | | |
| MLPH | melanophilin(MLPH) | NM_001042467 | 128.6 | 115.1 | 0.00698 | 0.861 | | | |

| | | | | | | |
|---|---|---|---|---|---|---|
| | | XR_001434 | 123.5 | 114.5 | 0.00698 | 0.88 |
| LOC100190986 | uncharacterized LOC100190986(LOC100190986) | NR_024456 | 123.5 | 109 | 0.00701 | 0.852 |
| BRIX1 | BRX1, biogenesis of ribosomes(BRIX1) | NM_018321 | 121.9 | 109.5 | 0.00741 | 0.871 |
| ALG13 | ALG13, UDP-N-acetylglucosaminyltransferase subunit(ALG13) | NM_018466 | 131 | 121.9 | 0.00742 | 0.874 |
| KLF7 | Kruppel like factor 7(KLF7) | NM_003709 | 159.5 | 275.6 | 0.00745 | 1.504 |
| ankrd40 | ankyrin repeat domain 40(ANKRD40) | NM_052855 | 114.4 | 199.7 | 0.00753 | 1.599 |
| OGFOD1 | 2-oxoglutarate and iron dependent oxygenase domain containing 1(OGFOD1) | NM_018233 | 99.3 | 130 | 0.00761 | 1.228 |
| OGFOD1 | 2-oxoglutarate and iron dependent oxygenase domain containing 1(OGFOD1) | NM_018233 | 108.8 | 120.6 | 0.00761 | 1.228 |
| ACYP1 | acylphosphatase 1(ACYP1) | NM_001107 | 147.2 | 120.3 | 0.00766 | 0.776 |
| | | XM_0017233164 | 130.4 | 153.2 | 0.00769 | 1.108 |
| ATCAY | ATCAY, caytaxin(ATCAY) | NM_033064 | 110.4 | 135.2 | 0.00773 | 1.172 |
| WDFY3 | WD repeat and FYVE domain containing 3(WDFY3) | NM_178583 | 113.6 | 103.2 | 0.00775 | 0.856 |

| | | | | | | |
|---|---|---|---|---|---|---|
| | | XR_038195 | 115.5 | 105.7 | 0.00799 | 0.883 |
| DNMT3L | DNA methyltransferase 3 like(DNMT3L) | NM_013369 | 135.1 | 124.2 | 0.00805 | 0.868 |
| DNMT3B | DNA methyltransferase 3 beta(DNMT3B) | NM_006892 | 100.9 | 100.9 | 0.00805 | 0.868 |
| OPRM1 | opioid receptor mu 1(OPRM1) | NM_001008505 | 109 | 127.7 | 0.00808 | 1.105 |
| OPRM1 | opioid receptor mu 1(OPRM1) | NM_001008505 | 178.1 | 175 | 0.00808 | 1.105 |
| ADIPOQ | adiponectin, C1Q and collagen domain containing(ADIPOQ) | NM_004797 | 91.8 | 113.8 | 0.00808 | 1.184 |
| CACNB1 | calcium voltage-gated channel auxiliary subunit beta 1(CACNB1) | NM_199247 | 123 | 112.6 | 0.00821 | 0.888 |
| ENTPD6 | ectonucleoside triphosphate diphosphohydrolase 6 (putative)(ENTPD6) | NM_001247 | 86.8 | 106.3 | 0.00831 | 1.194 |
| ENTPD7 | ectonucleoside triphosphate diphosphohydrolase 7(ENTPD7) | NM_020354 | 101.7 | 105.3 | 0.00831 | 1.194 |
| rab41 | RAB41, member RAS oncogene family(RAB41) | NM_001032726 | 159.4 | 138.4 | 0.00832 | 0.823 |
| FAM83F | family with sequence similarity 83 member F(FAM83F) | NM_138435 | 124.2 | 103.8 | 0.00838 | 0.795 |

| | | | | | | |
|---|---|---|---|---|---|---|
| PGA3 | pepsinogen 3, group I (pepsinogen A)(PGA3) | NM_0010 79807 | 90.1 | 96.1 | 0.00851 | 0.845 |
| PGA3 | pepsinogen 3, group I (pepsinogen A)(PGA3) | NM_0010 79807 | 113.2 | 103.8 | 0.00851 | 0.845 |
| PGA3 | pepsinogen 3, group I (pepsinogen A)(PGA3) | NM_0010 79807 | 130.2 | 116.3 | 0.00851 | 0.845 |
| SNX13 | sorting nexin 13(SNX13) | NM_0151 32 | 124 | 166.1 | 0.00852 | 1.259 |
| TDGP1 | thymine-DNA glycosylase pseudogene 1(TDGP1) | NR_02438 2 | 94.7 | 117.4 | 0.00859 | 1.164 |
| CCT5 | chaperonin containing TCP1 subunit 5(CCT5) | NM_0120 73 | 453.4 | 761.3 | 0.00873 | 1.48 |
| PRAM1 | PML-RARA regulated adaptor molecule 1(PRAM1) | NM_0321 52 | 117.5 | 111 | 0.00876 | 0.897 |
| SDIM1 | stress responsive DNAJB4 interacting membrane protein 1(SDIM1) | AK129633 | 111.7 | 100.4 | 0.00883 | 0.846 |
| RHBDL1 | rhomboid like 1(RHBDL1) | NM_0039 61 | 125.4 | 110.9 | 0.00884 | 0.858 |
| RBM48 | RNA binding motif protein 48(RBM48) | NM_0321 20 | 111.5 | 147.3 | 0.00888 | 1.228 |
| RBM48 | RNA binding motif protein 48(RBM48) | NM_0321 20 | 113.2 | 123.3 | 0.00888 | 1.228 |
| PBXIP1 | PBX homeobox interacting protein 1(PBXIP1) | NM_0205 24 | 161.6 | 143.6 | 0.00892 | 0.842 |
| | | NM_0010 04302 | 103.3 | 131.4 | 0.00904 | 1.218 |

| | | | | | | |
|---|---|---|---|---|---|---|
| | | XM_001126650 | 93.7 | 111.4 | 0.00908 | 1.155 |
| LMBRD1 | LMBR1 domain containing 1(LMBRD1) | NM_018368 | 147.4 | 118.6 | 0.00911 | 0.755 |
| | | XR_019076 | 93.7 | 108.9 | 0.0092 | 1.124 |
| | | AI351498 | 120.3 | 109.6 | 0.00935 | 0.879 |
| TRIM3 | tripartite motif containing 3(TRIM3) | NM_006458 | 150.2 | 141.6 | 0.0094 | 0.809 |
| TRIM3 | tripartite motif containing 3(TRIM3) | NM_006458 | 207.4 | 183.2 | 0.0094 | 0.809 |
| | | XM_001131286 | 127.8 | 120.4 | 0.00948 | 0.887 |
| | | BG951912 | 120.2 | 108.2 | 0.0095 | 0.873 |
| ALX3 | ALX homeobox 3(ALX3) | NM_006492 | 116.8 | 226.2 | 0.00957 | 1.705 |
| ALX1 | ALX homeobox 1(ALX1) | NM_006982 | 91.2 | 242.9 | 0.00957 | 1.705 |
| TUBA3FP | tubulin alpha 3f pseudogene(TUBA3FP) | NR_003608 | 118.3 | 110.9 | 0.00964 | 0.896 |
| TSNAX | translin associated factor X(TSNAX) | NM_005999 | 167.6 | 278.8 | 0.00969 | 1.43 |
| | | XM_941665 | 377.1 | 707.9 | 0.00979 | 1.679 |
| TAGLN | transgelin(TAGLN) | NM_001001522 | 126.5 | 115 | 0.00986 | 0.86 |
| FAM193A | family with sequence similarity 193 member A(FAM193A) | NM_003704 | 246 | 398.6 | 0.00989 | 1.403 |

| | | | | | | |
|---|---|---|---|---|---|---|
| RNF146 | ring finger protein 146(RNF146) | NM_030963 | 99.4 | 102.1 | 0.00991 | 1.355 |
| RNF146 | ring finger protein 146(RNF146) | NM_030963 | 122.5 | 189.7 | 0.00991 | 1.355 |
| | | BX105500 | 111.7 | 98.4 | 0.00991 | 0.852 |
| PCYOX1L | prenylcysteine oxidase 1 like(PCYOX1L) | NM_024028 | 100.8 | 119.2 | 0.00991 | 1.136 |
| SQLE | squalene epoxidase(SQLE) | NM_003129 | 103 | 141.6 | 0.00998 | 1.296 |
| SQLE | squalene epoxidase(SQLE) | NM_003129 | 105 | 115.6 | 0.00998 | 1.296 |
| | | CR593338 | 122.9 | 114.7 | 0.00999 | 0.873 |
| KDM4B | lysine demethylase 4B(KDM4B) | NM_015015 | 333.1 | 517.7 | 0.01009 | 1.343 |
| UBE2DNL | ubiquitin conjugating enzyme E2 D N-terminal like (pseudogene)(UBE2DNL) | NR_024062 | 105.1 | 95.2 | 0.01014 | 0.861 |
| C11orf58 | chromosome 11 open reading frame 58(C11orf58) | NM_001142705 | 191.7 | 170.1 | 0.01023 | 0.818 |
| PRKAR2A | protein kinase cAMP-dependent type II regulatory subunit alpha(PRKAR2A) | NM_004157 | 98.8 | 121.2 | 0.01026 | 1.171 |
| DNAJC10 | DnaJ heat shock protein family (Hsp40) member C10(DNAJC10) | NM_018981 | 102.7 | 114.7 | 0.01029 | 1.362 |

| | | | | | | | | | |
|---|---|---|---|---|---|---|---|---|---|
| DNAJC11 | DnaJ heat shock protein family (Hsp40) member C11(DNAJC11) | NM_018198 | 514.6 | 812.9 | 0.01029 | 1.362 | | | |
| DNAH6 | dynein axonemal heavy chain 6(DNAH6) | AB051484 | 149.9 | 117.8 | 0.01039 | 0.756 | | | |
| RHEB | Ras homolog enriched in brain(RHEB) | NM_005614 | 256 | 358.2 | 0.01042 | 1.225 | | | |
| DHX15 | DEAH-box helicase 15(DHX15) | NM_001358 | 230 | 328 | 0.01045 | 1.632 | | | |
| DHX16 | DEAH-box helicase 16(DHX16) | NM_003587 | 752.7 | 1388.1 | 0.01045 | 1.632 | | | |
| | | XM_001126824 | 150.6 | 135.5 | 0.01048 | 0.85 | | | |
| | | XM_001716052 | 110.8 | 100.1 | 0.01052 | 0.87 | | | |
| AMY1B | amylase, alpha 1B (salivary)(AMY1B) | NM_001008218 | 112.2 | 120.4 | 0.01056 | 1.624 | | | |
| AMY1C | amylase, alpha 1C (salivary)(AMY1C) | NM_001008219 | 1317.4 | 2434.2 | 0.01056 | 1.624 | | | |
| TMEM123 | transmembrane protein 123(TMEM123) | NM_052932 | 418.7 | 722.5 | 0.01061 | 1.498 | NM_001107943 | 0.63 | 112.77 |
| LEO1 | LEO1 homolog, Paf1/RNA polymerase II complex component(LEO1) | NM_138792 | 107.1 | 125 | 0.01068 | 1.12 | | | |
| QKI | QKI, KH domain containing RNA binding(QKI) | NM_006775 | 96.4 | 119.5 | 0.01078 | 1.165 | | | |

| | | | | | | |
|---|---|---|---|---|---|---|
| QKI | QKI, KH domain containing RNA binding(QKI) | NM_006775 | 105.3 | 119.9 | 0.01078 | 1.165 |
| | | XM_937684 | 132.4 | 125.1 | 0.01083 | 0.894 |
| MIR33B | microRNA 33b(MIR33B) | NR_030361 | 88.7 | 105.8 | 0.01083 | 1.168 |
| HNRNPK | heterogeneous nuclear ribonucleoprotein K(HNRNPK) | NM_031263 | 184.4 | 274.4 | 0.01088 | 1.348 |
| HNRNPK | heterogeneous nuclear ribonucleoprotein K(HNRNPK) | NM_031263 | 825.9 | 1283.7 | 0.01088 | 1.348 |
| HNRNPK | heterogeneous nuclear ribonucleoprotein K(HNRNPK) | NM_031263 | 988.8 | 1505.9 | 0.01088 | 1.348 |
| | | XM_0017163l0 | 107 | 100.3 | 0.01118 | 0.89 |
| | | XM_936314 | 127.8 | 114.9 | 0.01119 | 0.86 |
| FBXW5 | F-box and WD repeat domain containing 5(FBXW5) | NM_178226 | 251.7 | 240.3 | 0.01125 | 0.839 |
| | | XM_496269 | 99.6 | 121.5 | 0.01127 | 1.159 |
| GSTM1 | glutathione S-transferase mu 1(GSTM1) | NM_000561 | 98.9 | 115.8 | 0.01133 | 1.126 |

| | | | | | | | | | |
|---|---|---|---|---|---|---|---|---|---|
| ADAMTS6 | ADAM metallopeptidase with thrombospondin type 1 motif 6(ADAMTS6) | NM_197941 | 170.6 | 291.4 | 0.01137 | 1.484 | NM_134372 | 22.51 | 0.03 |
| FHOD1 | formin homology 2 domain containing 1(FHOD1) | NM_013241 | 296.4 | 997 | 0.01144 | 2.926 | | | |
| | | XM_001715275 | 115 | 106.1 | 0.01146 | 0.877 | | | |
| | | XM_934672 | 106.9 | 102.5 | 0.01155 | 0.918 | | | |
| CYP39A1 | cytochrome P450 family 39 subfamily A member 1(CYP39A1) | NM_016593 | 134.6 | 121 | 0.01157 | 0.858 | | | |
| HSD17B4 | hydroxysteroid 17-beta dehydrogenase 4(HSD17B4) | NM_000414 | 97.4 | 225.8 | 0.01161 | 2.07 | | | |
| MSMO1 | methylsterol monooxygenase 1(MSMO1) | NM_001017369 | 104.2 | 119.7 | 0.01162 | 1.113 | | | |
| MSMO1 | methylsterol monooxygenase 1(MSMO1) | NM_001017369 | 204.8 | 217.2 | 0.01162 | 1.113 | | | |
| RBPMS | RNA binding protein with multiple splicing(RBPMS) | NM_001008710 | 109.3 | 149.7 | 0.01162 | 1.274 | | | |
| PHLDA1 | pleckstrin homology like domain family A member 1(PHLDA1) | NM_007350 | 110.5 | 108.7 | 0.01165 | 1.361 | | | |
| PHLDA1 | pleckstrin homology like domain family A member 1(PHLDA1) | NM_007350 | 186.5 | 293 | 0.01165 | 1.361 | | | |

| | | | | | | |
|---|---|---|---|---|---|---|
| ZFHX4 | zinc finger homeobox 4(ZFHX4) | NM_024721 | 80.5 | 98.4 | 0.01169 | 1.213 |
| rbm12 | RNA binding motif protein 12(RBM12) | NM_006047 | 121.2 | 152.4 | 0.0118 | 1.177 |
| rbm12 | RNA binding motif protein 12(RBM12) | NM_006047 | 138.8 | 179.8 | 0.0118 | 1.177 |
| | | DB336481 | 136.1 | 128.4 | 0.0118 | 0.882 |
| HSD3B2 | hydroxy-delta-5-steroid dehydrogenase, 3 beta- and steroid delta-isomerase 2(HSD3B2) | NM_000198 | 98.5 | 113.7 | 0.01189 | 1.111 |
| CCDC87 | coiled-coil domain containing 87(CCDC87) | NM_018219 | 116.5 | 102.2 | 0.01196 | 0.848 |
| SLC35A2 | solute carrier family 35 member A2(SLC35A2) | NM_001032289 | 109.9 | 126.1 | 0.01206 | 1.094 |
| SLC35A2 | solute carrier family 35 member A2(SLC35A2) | NM_001032289 | 345.8 | 386 | 0.01206 | 1.094 |
| | | AW044615 | 120.4 | 110.6 | 0.01257 | 0.877 |
| | | XM_001720423 | 149.9 | 130 | 0.01262 | 0.833 |
| Snora41 | small nucleolar RNA, H/ACA box 41(SNORA41) | NR_002590 | 109.5 | 143.7 | 0.01267 | 1.264 |
| RBFOX2 | RNA binding protein, fox-1 homolog 2(RBFOX2) | NM_001082576 | 234.9 | 422.2 | 0.0128 | 1.554 |

| | | | | | | | | | |
|---|---|---|---|---|---|---|---|---|---|
| HTR3A | 5-hydroxytryptamine receptor 3A(HTR3A) | NM_000869 | 106.4 | 96.9 | 0.01282 | 0.871 | NM_001031645 | 1.93 | 10.04 |
| HTR3A | 5-hydroxytryptamine receptor 3A(HTR3A) | NM_213621 | 147.9 | 157.9 | 0.01282 | 0.871 | | | |
| MATR3 | matrin 3(MATR3) | NM_199189 | 97.7 | 103.8 | 0.01297 | 1.418 | | | |
| MATR3 | matrin 3(MATR3) | NM_199189 | 100.8 | 111 | 0.01297 | 1.418 | | | |
| MATR3 | matrin 3(MATR3) | NM_199189 | 1659.9 | 2694.5 | 0.01297 | 1.418 | | | |
| MTX1 | metaxin 1(MTX1) | NM_198883 | 171.5 | 158.1 | 0.01297 | 0.84 | | | |
| MOXD1 | monooxygenase DBH like 1(MOXD1) | NM_015529 | 102.9 | 168.3 | 0.01312 | 1.524 | | | |
| TMEM136 | transmembrane protein 136(TMEM136) | NM_174926 | 86.9 | 106.7 | 0.01314 | 1.197 | | | |
| AHCTF1 | AT-hook containing transcription factor 1(AHCTF1) | NM_015446 | 1763.1 | 731.7 | 0.01316 | 0.351 | | | |
| AP4E1 | adaptor related protein complex 4 epsilon 1 subunit(AP4E1) | NM_007347 | 168.8 | 150.5 | 0.01317 | 0.84 | | | |
| MAPKBP1 | mitogen-activated protein kinase binding protein 1(MAPKBP1) | NM_014994 | 125.3 | 135 | 0.01326 | 1.139 | | | |
| MAPKAPK5 | mitogen-activated protein kinase-activated protein kinase 5(MAPKAPK5) | NM_139078 | 91.6 | 107.7 | 0.01326 | 1.139 | | | |

| | | | | | | | | | |
|---|---|---|---|---|---|---|---|---|---|
| SF3A3 | splicing factor 3a subunit 3(SF3A3) | NM_006802 | 817.1 | 1234.5 | 0.01327 | 1.327 | | | |
| IFT172 | intraflagellar transport 172(IFT172) | NM_015662 | 164.7 | 216.9 | 0.01331 | 1.2 | | | |
| B3GAT1 | beta-1,3-glucuronyltransferase 1(B3GAT1) | NM_054025 | 102.7 | 124.5 | 0.01357 | 1.164 | | | |
| RAC3 | ras-related C3 botulinum toxin substrate 3 (rho family, small GTP binding protein Rac3)(RAC3) | NM_005052 | 238.8 | 189.4 | 0.01379 | 0.732 | NM_144748 | 66.99 | 0.14 |
| | | XM_371878 | 94.4 | 114.5 | 0.01381 | 1.173 | | | |
| MSANTD4 | Myb/SANT DNA binding domain containing 4 with coiled-coils(MSANTD4) | NM_032424 | 171.2 | 239.6 | 0.01392 | 1.249 | | | |
| SEBOX | SEBOX homeobox(SEBOX) | NM_001080837 | 114.6 | 106.1 | 0.01403 | 0.876 | | | |
| NAPEPLD | N-acyl phosphatidylethanolamine phospholipase D(NAPEPLD) | NM_198990 | 117.5 | 121.5 | 0.01417 | 1.212 | | | |
| NAPEPLD | N-acyl phosphatidylethanolamine phospholipase D(NAPEPLD) | NM_198990 | 151.5 | 203.1 | 0.01417 | 1.212 | | | |
| CHAD | chondroadherin(CHAD) | NM_001267 | 100 | 122.7 | 0.01418 | 1.187 | | | |

| | | | | | | | | | |
|---|---|---|---|---|---|---|---|---|---|
| PPIE | peptidylprolyl isomerase E(PPIE) | NM_0061 12 | 124.6 | 165.2 | 0.01427 | 1.224 | | | |
| | | XR_03767 2 | 112.5 | 135.6 | 0.01433 | 1.148 | | | |
| HAUS2 | HAUS augmin like complex subunit 2(HAUS2) | NM_0180 97 | 172.6 | 247 | 0.01436 | 1.288 | NM_001 107793 | 77.29 | 6.85 |
| HAT1 | histone acetyltransferase 1(HAT1) | NM_0036 42 | 134.1 | 151 | 0.01436 | 1.288 | NM_001 107793 | 77.29 | 6.85 |
| | | XM_93150 6 | 106.7 | 125.6 | 0.01442 | 1.128 | NM_001 107793 | 77.29 | 6.85 |
| HOXD3 | homeobox D3(HOXD3) | NM_0068 98 | 201.9 | 444.6 | 0.0146 | 1.94 | NM_001 107793 | 77.29 | 6.85 |
| BTLA | B and T lymphocyte associated(BTLA) | NM_1817 80 | 105 | 97.1 | 0.01464 | 0.878 | | | |
| FAM26D | family with sequence similarity 26 member D(FAM26D) | NM_1530 36 | 106.6 | 112.5 | 0.01474 | 0.902 | | | |
| FAM26F | family with sequence similarity 26 member F(FAM26F) | NM_0010 10919 | 121.7 | 114.9 | 0.01474 | 0.902 | | | |
| PI15 | peptidase inhibitor 15(PI15) | NM_0158 86 | 95.2 | 113.2 | 0.01484 | 1.136 | | | |
| PI15 | peptidase inhibitor 15(PI15) | NM_0158 86 | 136.8 | 127.6 | 0.01484 | 1.136 | | | |

| | | | | | | |
|---|---|---|---|---|---|---|
| ELP2 | elongator acetyltransferase complex subunit 2(ELP2) | NM_018255 | 133.4 | 268.5 | 0.01485 | 1.733 |
| ELOVL6 | ELOVL fatty acid elongase 6(ELOVL6) | NM_024090 | 113.7 | 173.4 | 0.01485 | 1.733 |
| FNDC4 | fibronectin type III domain containing 4(FNDC4) | NM_022823 | 338.7 | 599.5 | 0.01487 | 1.536 |
| CAPRIN2 | caprin family member 2(CAPRIN2) | NM_023925 | 160.4 | 234.6 | 0.01487 | 1.311 |
| MPHOSPH6 | M-phase phosphoprotein 6(MPHOSPH6) | NM_005792 | 127.5 | 164.8 | 0.01492 | 1.202 |
| CPSF4 | cleavage and polyadenylation specific factor 4(CPSF4) | NM_001081559 | 152.5 | 191.1 | 0.01498 | 1.18 |
| CPSF3L | cleavage and polyadenylation specific factor 3-like(CPSF3L) | NM_017871 | 97.9 | 122.7 | 0.01498 | 1.18 |
| | | XM_001132711 | 142.6 | 127.2 | 0.01513 | 0.848 |
| CCDC124 | coiled-coil domain containing 124(CCDC124) | NM_138442 | 91.1 | 110.5 | 0.0153 | 1.189 |
| RARB | retinoic acid receptor beta(RARB) | NM_000965 | 122.7 | 195.1 | 0.01545 | 1.398 |
| RARB | retinoic acid receptor beta(RARB) | NM_000965 | 126.3 | 251.4 | 0.01545 | 1.398 |

| | | | | | | |
|---|---|---|---|---|---|---|
| MSLNL | mesothelin-like(MSLNL) | NM_001025190 | 108.4 | 101 | 0.01554 | 0.856 |
| MSLNL | mesothelin-like(MSLNL) | NM_001025190 | 113.5 | 102 | 0.01554 | 0.856 |
| rtf1 | RTF1 homolog, Paf1/RNA polymerase II complex component(RTF1) | NM_015138 | 152.7 | 142.8 | 0.01576 | 0.873 |
| FZD1 | frizzled class receptor 1(FZD1) | NM_003505 | 117.6 | 110.1 | 0.01589 | 0.88 |
| FZD2 | frizzled class receptor 2(FZD2) | NM_001466 | 251.1 | 258.2 | 0.01589 | 0.88 |
| CSNK2A1 | casein kinase 2 alpha 1(CSNK2A1) | NM_001895 | 125.5 | 163.4 | 0.01591 | 1.199 |
| SYTL2 | synaptotagmin like 2(SYTL2) | NM_032943 | 90.5 | 116.1 | 0.01595 | 1.231 |
| tmem132a | transmembrane protein 132A(TMEM132A) | NM_178031 | 98.8 | 130.8 | 0.01611 | 1.252 |
| MORF4L2 | mortality factor 4 like 2(MORF4L2) | NM_012286 | 583.1 | 916.9 | 0.01613 | 1.377 |
| | | NM_001004349 | 124.5 | 112.3 | 0.01613 | 0.847 |
| GDI1 | GDP dissociation inhibitor 1(GDI1) | NM_001493 | 141.8 | 130.3 | 0.01614 | 0.876 |
| CDHR5 | cadherin related family member 5(CDHR5) | NM_021924 | 140.3 | 129.6 | 0.01617 | 0.872 |
| ZCCHC8 | zinc finger CCHC-type containing 8(ZCCHC8) | NM_017612 | 150 | 193.1 | 0.01618 | 1.171 |
| NOXO1 | NADPH oxidase organizer 1(NOXO1) | NM_172167 | 142.3 | 115.5 | 0.01621 | 0.757 |

| | | | | | | | | | |
|---|---|---|---|---|---|---|---|---|---|
| ORAI3 | ORAI calcium release-activated calcium modulator 3(ORAI3) | NM_152288 | 180.3 | 165.3 | 0.01627 | 0.85 | | | |
| mgarp | mitochondria localized glutamic acid rich protein(MGARP) | NM_032623 | 87.5 | 107.8 | 0.01629 | 1.203 | | | |
| mgarp | mitochondria localized glutamic acid rich protein(MGARP) | NM_032623 | 97.1 | 148.2 | 0.01629 | 1.203 | | | |
| MT1X | metallothionein 1X(MT1X) | NM_005952 | 6772.5 | 2593.9 | 0.01632 | 0.32 | | | |
| ZNF174 | zinc finger protein 174(ZNF174) | NM_001032292 | 108.9 | 108.1 | 0.01637 | 1.156 | | | |
| ZNF174 | zinc finger protein 174(ZNF174) | NM_001032292 | 110.7 | 135.4 | 0.01637 | 1.156 | NM_001009603 | 28.32 | 0.90 |
| H2AFJ | H2A histone family member J(H2AFJ) | NM_018267 | 111.7 | 104.7 | 0.01649 | 0.888 | | | |
| DCAF4L1 | DDB1 and CUL4 associated factor 4 like 1(DCAF4L1) | NM_001029955 | 99.8 | 117.6 | 0.01649 | 1.121 | | | |
| FLYWCH1 | FLYWCH-type zinc finger 1(FLYWCH1) | NM_020912 | 124.4 | 112.4 | 0.01661 | 0.874 | | | |
| LRRC46 | leucine rich repeat containing 46(LRRC46) | NM_033413 | 113.2 | 103.7 | 0.01672 | 0.856 | | | |
| LRRC45 | leucine rich repeat containing 45(LRRC45) | NM_144999 | 110.5 | 99.2 | 0.01672 | 0.856 | | | |
| NME3 | NME/NM23 nucleoside diphosphate kinase 3(NME3) | NM_002513 | 1065.4 | 800.5 | 0.01674 | 0.659 | | | |

| | | | | | | |
|---|---|---|---|---|---|---|
| LIPT1 | lipoyltransferase 1(LIPT1) | NM_015929 | 101.6 | 127.9 | 0.01677 | 1.198 |
| LITAF | lipopolysaccharide induced TNF factor(LITAF) | NM_004862 | 109 | 116.6 | 0.01677 | 1.198 |
| HM13 | histocompatibility minor 13(HM13) | NM_178580 | 132.4 | 145.5 | 0.01679 | 1.248 |
| HM13 | histocompatibility minor 13(HM13) | NM_178582 | 175 | 246.4 | 0.01679 | 1.248 |
| CCZ1 | CCZ1 homolog, vacuolar protein trafficking and biogenesis associated(CCZ1) | NM_015622 | 868.3 | 1276 | 0.01691 | 1.288 |
| | | XM_943482 | 115.3 | 107.2 | 0.01716 | 0.878 |
| ITPA | inosine triphosphatase(ITPA) | NM_033453 | 1119.3 | 1967.5 | 0.01725 | 1.573 |
| ITM2C | integral membrane protein 2C(ITM2C) | NM_001012516 | 735.9 | 1104.2 | 0.01725 | 1.573 |
| LOC105378397 | uncharacterized LOC105378397(LOC105378397) | BX092644 | 201.8 | 162.1 | 0.01735 | 0.754 |
| CSNK1G1 | casein kinase 1 gamma 1(CSNK1G1) | NM_022048 | 142.1 | 200.6 | 0.01737 | 1.399 |
| csnk1e | casein kinase 1 epsilon(CSNK1E) | NM_152221 | 127.6 | 197.4 | 0.01737 | 1.399 |
| TSACC | TSSK6 activating cochaperone(TSACC) | NM_144627 | 106.5 | 133.2 | 0.01756 | 1.213 |
| ZNF77 | zinc finger protein 77(ZNF77) | NM_021217 | 108.6 | 137.3 | 0.0176 | 1.194 |

| | | | | | | |
|---|---|---|---|---|---|---|
| TMEM237 | transmembrane protein 237(TMEM237) | NM_001044385 | 145.7 | 225.7 | 0.01761 | 1.367 |
| PCM1 | pericentriolar material 1(PCM1) | NM_006197 | 268.7 | 361.1 | 0.01767 | 1.172 |
| PCM1 | pericentriolar material 1(PCM1) | NM_006197 | 382.6 | 494 | 0.01767 | 1.172 |
| GSTO1 | glutathione S-transferase omega 1(GSTO1) | NM_004832 | 122.2 | 115.2 | 0.0177 | 0.905 |
| | | XM_001714310 | 111.4 | 105.1 | 0.01797 | 0.9 |
| PPIL2 | peptidylprolyl isomerase like 2(PPIL2) | NM_014337 | 132.2 | 117.2 | 0.01812 | 0.844 |
| PPIL2 | peptidylprolyl isomerase like 2(PPIL2) | NM_014337 | 182.9 | 188.4 | 0.01812 | 0.844 |
| PHF12 | PHD finger protein 12(PHF12) | NM_020889 | 114 | 135.7 | 0.01823 | 1.14 |
| TTC19 | tetratricopeptide repeat domain 19(TTC19) | NM_017775 | 97.6 | 113.4 | 0.01824 | 1.114 |
| CAMK1D | calcium/calmodulin dependent protein kinase ID(CAMK1D) | AK097377 | 263.2 | 521.1 | 0.01826 | 1.717 |
| DUSP10 | dual specificity phosphatase 10(DUSP10) | NM_007207 | 103.7 | 94.9 | 0.01855 | 0.862 |
| BRD7 | bromodomain containing 7(BRD7) | NM_013263 | 111.9 | 104.4 | 0.01868 | 0.878 |
| SOX4 | SRY-box 4(SOX4) | NM_003107 | 227.9 | 498.5 | 0.01876 | 1.891 |
| HIF3A | hypoxia inducible factor 3 alpha subunit(HIF3A) | NM_022462 | 114.3 | 139.9 | 0.01887 | 1.166 |

| | | | | | | |
|---|---|---|---|---|---|---|
| | | XM_001715696 | 116.8 | 111 | 0.01896 | 0.91 |
| MAFF | MAF bZIP transcription factor F(MAFF) | NM_012323 | 183.4 | 173.6 | 0.01903 | 0.855 |
| TRIM47 | tripartite motif containing 47(TRIM47) | NM_033452 | 104.5 | 123.2 | 0.01921 | 1.118 |
| ADCY7 | adenylate cyclase 7(ADCY7) | NM_001114 | 260.2 | 480.8 | 0.01934 | 1.625 |
| ADCY6 | adenylate cyclase 6(ADCY6) | NM_020983 | 173 | 239.1 | 0.01934 | 1.625 |
| MAP3K4 | mitogen-activated protein kinase kinase kinase 4(MAP3K4) | NM_005922 | 157 | 140.3 | 0.01935 | 0.822 |
| | | XM_933836 | 122.2 | 115.4 | 0.01944 | 0.907 |
| HLA-DQB2 | major histocompatibility complex, class II, DQ beta 2(HLA-DQB2) | NM_182549 | 147.6 | 137.5 | 0.01945 | 0.88 |
| | | XR_017689 | 128.7 | 124.7 | 0.01946 | 0.914 |
| ASAP2 | ArfGAP with SH3 domain, ankyrin repeat and PH domain 2(ASAP2) | NM_003887 | 112.3 | 100 | 0.01946 | 0.869 |
| ASAP2 | ArfGAP with SH3 domain, ankyrin repeat and PH domain 2(ASAP2) | NM_003887 | 280.6 | 369.1 | 0.01946 | 0.869 |

| | | | | | | |
|---|---|---|---|---|---|---|
| ASAP2 | ArfGAP with SH3 domain, ankyrin repeat and PH domain 2(ASAP2) | NM_003887 | 293 | 451.6 | 0.01946 | 0.869 |
| | | AW088717 | 87.6 | 105.6 | 0.01956 | 1.168 |
| FOXO4 | forkhead box O4(FOXO4) | NM_005938 | 273 | 450.4 | 0.0196 | 1.44 |
| FOXO3 | forkhead box O3(FOXO3) | NM_201559 | 197.9 | 344.8 | 0.0196 | 1.44 |
| ACER3 | alkaline ceramidase 3(ACER3) | NM_018367 | 123.9 | 121.4 | 0.01965 | 0.699 |
| ACHE | acetylcholinesterase (Cartwright blood group)(ACHE) | NM_015831 | 213.5 | 160 | 0.01965 | 0.699 |
| HNRNPH1 | heterogeneous nuclear ribonucleoprotein H1 (H)(HNRNPH1) | NM_005520 | 144.4 | 155.1 | 0.01969 | 1.357 |
| HNRNPH1 | heterogeneous nuclear ribonucleoprotein H1 (H)(HNRNPH1) | NM_005520 | 144.8 | 200.9 | 0.01969 | 1.357 |
| HNRNPH1 | heterogeneous nuclear ribonucleoprotein H1 (H)(HNRNPH1) | NM_005520 | 1152.8 | 1768.4 | 0.01969 | 1.357 |
| CCNK | cyclin K(CCNK) | NM_003858 | 93.5 | 110.5 | 0.01975 | 1.141 |
| DEFA1B | defensin alpha 1B(DEFA1B) | NM_001042500 | 92.5 | 114.2 | 0.01976 | 1.177 |
| OR2B6 | olfactory receptor family 2 subfamily B member 6(OR2B6) | NM_012367 | 113.9 | 108.7 | 0.01984 | 0.919 |

| | | | | | | |
|---|---|---|---|---|---|---|
| BRD9 | bromodomain containing 9(BRD9) | NM_023924 | 387.8 | 306.3 | 0.01987 | 0.71 |
| BRD8 | bromodomain containing 8(BRD8) | NM_183359 | 205.6 | 261.6 | 0.01987 | 0.71 |
| PRPF40B | pre-mRNA processing factor 40 homolog B(PRPF40B) | NM_001031698 | 113.8 | 138.2 | 0.01988 | 1.165 |
| CTAG2 | cancer/testis antigen 2(CTAG2) | NM_020994 | 119.5 | 107 | 0.02006 | 0.879 |
| ATL3 | atlastin GTPase 3(ATL3) | NM_015459 | 88.8 | 106.2 | 0.02007 | 1.169 |
| ZFHX3 | zinc finger homeobox 3(ZFHX3) | NM_006885 | 161.7 | 298.1 | 0.02023 | 1.591 |
| | | XM_926073 | 121.5 | 115.3 | 0.02028 | 0.903 |
| | | XM_001125745 | 102.8 | 105.3 | 0.02028 | 0.861 |
| | | XM_001125745 | 104.3 | 94.6 | 0.02028 | 0.861 |
| TCTE3 | t-complex-associated-testis-expressed 3(TCTE3) | NM_174910 | 95.5 | 117.8 | 0.02032 | 1.186 |
| | | XM_928619 | 177.5 | 168.5 | 0.02033 | 0.882 |
| ACSS2 | acyl-CoA synthetase short-chain family member 2(ACSS2) | NM_001076552 | 116.3 | 101.8 | 0.0204 | 0.822 |
| ACSS2 | acyl-CoA synthetase short-chain family member 2(ACSS2) | NM_018677 | 1898.7 | 1215.1 | 0.0204 | 0.822 |

| | | | | | | |
|---|---|---|---|---|---|---|
| ITGB2 | integrin subunit beta 2(ITGB2) | NM_000211 | 717.3 | 1190.7 | 0.02047 | 1.437 |
| PPP2R2B | protein phosphatase 2 regulatory subunit Bbeta(PPP2R2B) | NM_181676 | 96.8 | 121.6 | 0.0205 | 1.224 |
| PPP2R2B | protein phosphatase 2 regulatory subunit Bbeta(PPP2R2B) | NM_181676 | 165.8 | 336 | 0.0205 | 1.224 |
| ZNF451 | zinc finger protein 451(ZNF451) | NM_001031623 | 101.2 | 112.8 | 0.02055 | 1.245 |
| ZNF451 | zinc finger protein 451(ZNF451) | NM_001031623 | 105.3 | 136.7 | 0.02055 | 1.245 |
| | | XM_940726 | 110.2 | 98.1 | 0.02058 | 0.847 |
| | | XM_940726 | 136.9 | 138.7 | 0.02058 | 0.847 |
| f2rl2 | coagulation factor II thrombin receptor like 2(F2RL2) | NM_004101 | 16746 | 19918.8 | 0.02059 | 1.189 |
| | | XM_927957 | 130.6 | 118.9 | 0.02066 | 0.856 |
| GUF1 | GUF1 homolog, GTPase(GUF1) | NM_021927 | 109 | 151.7 | 0.02068 | 1.295 |
| or8g1 | olfactory receptor family 8 subfamily G member 1 (gene/pseudogene)(OR8G1) | NM_001002905 | 111.3 | 102.1 | 0.02076 | 0.875 |
| ADD1 | adducin 1(ADD1) | NM_001119 | 135.1 | 102.5 | 0.02083 | 0.732 |

| | | | | | | | | | |
|---|---|---|---|---|---|---|---|---|---|
| LOC102724002 | uncharacterized LOC102724002(LOC102724002) | AK025166 | 110.4 | 127.3 | 0.02097 | 1.116 | | | |
| TOR2A | torsin family 2 member A(TOR2A) | NM_130459 | 93 | 122.1 | 0.02099 | 1.263 | | | |
| | | XM_001133423 | 98.7 | 115.5 | 0.02112 | 1.112 | | | |
| LRRN4CL | LRRN4 C-terminal like(LRRN4CL) | NM_203422 | 176.6 | 309.6 | 0.02115 | 1.57 | | | |
| SERTAD1 | SERTA domain containing 1(SERTAD1) | NM_013376 | 288.4 | 483.5 | 0.02118 | 1.464 | | | |
| RSBN1L | round spermatid basic protein 1 like(RSBN1L) | NM_198467 | 107.1 | 132.8 | 0.02119 | 1.186 | | | |
| c14orf166 | chromosome 14 open reading frame 166(C14orf166) | NM_016039 | 185.8 | 441.1 | 0.02125 | 2.099 | | | |
| SAFB2 | scaffold attachment factor B2(SAFB2) | NM_014649 | 125.4 | 127.8 | 0.0213 | 0.788 | | | |
| SAFB2 | scaffold attachment factor B2(SAFB2) | NM_014649 | 758.5 | 683 | 0.0213 | 0.788 | | | |
| | | CK299443 | 149.4 | 135.7 | 0.02132 | 0.864 | | | |
| ELAVL1 | ELAV like RNA binding protein 1(ELAVL1) | NM_001419 | 1332.8 | 1198.1 | 0.02133 | 0.763 | NM_144744 | 0.11 | 40.01 |
| DKKL1 | dickkopf like acrosomal protein 1(DKKL1) | NM_014419 | 847.9 | 2015.2 | 0.02142 | 2.089 | | | |
| DPH3 | diphthamide biosynthesis 3(DPH3) | NM_001047434 | 422.9 | 685.1 | 0.02147 | 1.426 | | | |

| | | | | | | |
|---|---|---|---|---|---|---|
| AIMP2 | aminoacyl tRNA synthetase complex interacting multifunctional protein 2(AIMP2) | NM_006303 | 125.2 | 114.1 | 0.02159 | 0.878 |
| | | XM_379378 | 108.6 | 127.6 | 0.0216 | 1.113 |
| ILK | integrin linked kinase(ILK) | NM_004517 | 200.8 | 295 | 0.02166 | 1.285 |
| MBTPS1 | membrane bound transcription factor peptidase, site 1(MBTPS1) | NM_003791 | 166.4 | 241.1 | 0.02168 | 1.307 |
| MBTPS1 | membrane bound transcription factor peptidase, site 1(MBTPS1) | NM_003791 | 182.9 | 208.1 | 0.02168 | 1.307 |
| C1orf74 | chromosome 1 open reading frame 74(C1orf74) | NM_152485 | 117.9 | 107.9 | 0.02172 | 0.873 |
| | | XM_001721243 | 92.6 | 110.3 | 0.02172 | 1.146 |
| SNORA2C | small nucleolar RNA, H/ACA box 2C(SNORA2C) | NR_002968 | 100.9 | 119.4 | 0.02172 | 1.125 |
| | | XM_001716887 | 115.4 | 154.2 | 0.02184 | 1.268 |
| | | XM_940353 | 114.2 | 106.7 | 0.02191 | 0.894 |
| TAGLN | transgelin(TAGLN) | NM_003186 | 343.4 | 753.9 | 0.02196 | 1.675 |

| | | | | | | |
|---|---|---|---|---|---|---|
| TAGLN | transgelin(TAGLN) | NM_003186 | 1523 | 2874.1 | 0.02196 | 1.675 |
| CEPT1 | choline/ethanolamine phosphotransferase 1(CEPT1) | NM_006090 | 139.7 | 130.3 | 0.02205 | 0.872 |
| MYL9 | myosin light chain 9(MYL9) | NM_006097 | 185.8 | 203.6 | 0.02227 | 1.349 |
| MYL9 | myosin light chain 9(MYL9) | NM_006097 | 222 | 341.8 | 0.02227 | 1.349 |
| | | XM_927196 | 109.4 | 103.7 | 0.02243 | 0.892 |
| S100A13 | S100 calcium binding protein A13(S100A13) | NM_001024211 | 122.2 | 200.3 | 0.02246 | 1.49 |
| S100A13 | S100 calcium binding protein A13(S100A13) | NM_001024211 | 170.6 | 178 | 0.02246 | 1.49 |
| PDGFA | platelet derived growth factor subunit A(PDGFA) | NM_033023 | 123.1 | 166.8 | 0.02255 | 1.263 |
| SIDT2 | SID1 transmembrane family member 2(SIDT2) | NM_001040455 | 350.6 | 349.9 | 0.02287 | 0.87 |
| SEPT4 | septin 4(SEPT4) | NM_080415 | 1066.6 | 1678.5 | 0.02295 | 1.379 |
| TGIF1 | TGFB induced factor homeobox 1(TGIF1) | NM_170695 | 197.5 | 302.5 | 0.02298 | 1.35 |
| TGIF1 | TGFB induced factor homeobox 1(TGIF1) | NM_170695 | 284.3 | 451.6 | 0.02298 | 1.35 |
| | | XM_498969 | 203.5 | 155.4 | 0.02308 | 0.718 |

| | | | | | | |
|---|---|---|---|---|---|---|
| ATG4A | autophagy related 4A cysteine peptidase(ATG4A) | NM_178270 | 100.5 | 93.6 | 0.02313 | 0.877 |
| ORC3 | origin recognition complex subunit 3(ORC3) | NM_181837 | 100.3 | 134.1 | 0.02325 | 1.267 |
| ZFX | zinc finger protein, X-linked(ZFX) | NM_003410 | 95.7 | 116.2 | 0.02342 | 1.16 |
| | | XR_037203 | 378 | 584.7 | 0.02343 | 1.362 |
| | | AI005586 | 101 | 122.5 | 0.02367 | 1.144 |
| NIPAL1 | NIPA like domain containing 1(NIPAL1) | NM_207330 | 127.6 | 148.1 | 0.02372 | 1.113 |
| | | XM_928722 | 125.3 | 118.7 | 0.02373 | 0.911 |
| C1orf87 | chromosome 1 open reading frame 87(C1orf87) | NM_152377 | 94.1 | 109.7 | 0.02376 | 1.141 |
| C1QA | complement C1q A chain(C1QA) | NM_015991 | 104.5 | 115.3 | 0.02376 | 1.141 |
| AMZ2P1 | archaelysin family metallopeptidase 2 pseudogene 1(AMZ2P1) | NM_153032 | 302.1 | 551.9 | 0.02377 | 1.58 |
| PIGV | phosphatidylinositol glycan anchor biosynthesis class V(PIGV) | NM_017837 | 112 | 104.2 | 0.02381 | 0.875 |

| | | | | | | |
|---|---|---|---|---|---|---|
| TMEM158 | transmembrane protein 158 (gene/pseudogene)(TMEM158) | NM_015444 | 224.3 | 1619.8 | 0.02384 | 6.279 |
| scaf8 | SR-related CTD associated factor 8(SCAF8) | NM_014892 | 144.7 | 214 | 0.02396 | 1.296 |
| APCS | amyloid P component, serum(APCS) | NM_001639 | 130.2 | 304.7 | 0.02396 | 2.035 |
| | | XM_001716882 | 119.4 | 187.4 | 0.02399 | 1.452 |
| DCP1A | decapping mRNA 1A(DCP1A) | NM_018403 | 433.8 | 1278.5 | 0.02402 | 2.633 |
| | | BC035064 | 95 | 108.8 | 0.02409 | 1.105 |
| setmar | SET domain and mariner transposase fusion gene(SETMAR) | NM_006515 | 201 | 301.3 | 0.02416 | 1.305 |
| GDF9 | growth differentiation factor 9(GDF9) | NM_005260 | 89.5 | 107.5 | 0.02425 | 1.163 |
| SGCE | sarcoglycan epsilon(SGCE) | NM_001099400 | 391.5 | 695.5 | 0.02436 | 1.536 |
| ZNF789 | zinc finger protein 789(ZNF789) | NM_001013258 | 112.3 | 154.8 | 0.02438 | 1.284 |
| | | XM_941746 | 101.5 | 130.2 | 0.02441 | 1.219 |
| KIAA1429 | KIAA1429(KIAA1429) | NM_183009 | 93.6 | 111.7 | 0.02442 | 1.152 |
| PTK2 | protein tyrosine kinase 2(PTK2) | NM_005607 | 1002.1 | 1702.9 | 0.02444 | 1.484 |

| | | | | | | |
|---|---|---|---|---|---|---|
| PDX1 | pancreatic and duodenal homeobox 1(PDX1) | NM_000209 | 150.3 | 142 | 0.02458 | 0.886 |
| MYLK | myosin light chain kinase(MYLK) | NM_053032 | 239.3 | 559.1 | 0.02464 | 1.993 |
| | | XR_017001 | 118.2 | 108.6 | 0.02469 | 0.868 |
| | | XR_017364 | 236.8 | 160.9 | 0.02471 | 0.62 |
| DNM3 | dynamin 3(DNM3) | NM_015569 | 132 | 123.2 | 0.02486 | 0.878 |
| | | XR_018230 | 90.2 | 109.7 | 0.02487 | 1.179 |
| | | XM_0017133713 | 101.5 | 116.1 | 0.02491 | 1.1 |
| RPS6KA3 | ribosomal protein S6 kinase A3(RPS6KA3) | NM_004586 | 99.5 | 115 | 0.02507 | 1.236 |
| RPS6KA3 | ribosomal protein S6 kinase A3(RPS6KA3) | NM_004586 | 127.2 | 176.7 | 0.02507 | 1.236 |
| HOXC10 | homeobox C10(HOXC10) | NM_017409 | 343.8 | 869.4 | 0.02507 | 2.234 |
| CBL | Cbl proto-oncogene(CBL) | NM_005188 | 171.7 | 142.1 | 0.02513 | 0.799 |
| GPN1 | GPN-loop GTPase 1(GPN1) | NM_007266 | 129.6 | 315.4 | 0.0252 | 1.183 |
| GPM6B | glycoprotein M6B(GPM6B) | NM_001001995 | 96.8 | 119.1 | 0.0252 | 1.183 |
| | | XM_370711 | 108.5 | 100.6 | 0.02535 | 0.885 |

| | | | | | | |
|---|---|---|---|---|---|---|
| ORMDL 1 | ORMDL sphingolipid biosynthesis regulator 1(ORMDL1) | NM_0164 67 | 107.8 | 129.3 | 0.02553 | 1.158 |
| IFNGR2 | interferon gamma receptor 2 (interferon gamma transducer 1)(IFNGR2) | NM_0055 34 | 79.7 | 98.7 | 0.02557 | 1.218 |
| SLC13A 2 | solute carrier family 13 member 2(SLC13A2) | NM_0039 84 | 281.3 | 207.6 | 0.02559 | 0.68 |
| FAM98B | family with sequence similarity 98 member B(FAM98B) | NM_0010 42429 | 347.8 | 311 | 0.02568 | 0.795 |
| CEACA M21 | carcinoembryonic antigen related cell adhesion molecule 21(CEACAM21) | NM_0010 98506 | 106.4 | 130.3 | 0.02569 | 1.163 |
| RUFY3 | RUN and FYVE domain containing 3(RUFY3) | NM_0010 37442 | 104 | 121.5 | 0.02572 | 1.124 |
| RUFY3 | RUN and FYVE domain containing 3(RUFY3) | NM_0010 37442 | 349.3 | 465.4 | 0.02572 | 1.124 |
| OSGEPL 1 | O-sialoglycoprotein endopeptidase like 1(OSGEPL1) | NM_0223 53 | 107.6 | 116.1 | 0.02575 | 0.911 |
| OSGEPL 1 | O-sialoglycoprotein endopeptidase like 1(OSGEPL1) | NM_0223 53 | 111.9 | 105.9 | 0.02575 | 0.911 |
| | | CD242195 | 119.9 | 113.3 | 0.02585 | 0.896 |
| GOLGB1 | golgin B1(GOLGB1) | NM_0044 87 | 767 | 1093.7 | 0.026 | 1.244 |

| | | | | | | |
|---|---|---|---|---|---|---|
| FKBP8 | FK506 binding protein 8(FKBP8) | NM_012181 | 194.9 | 170.1 | 0.02609 | 0.813 |
| HTR3A | 5-hydroxytryptamine receptor 3A(HTR3A) | NM_000869 | 109.4 | 162.8 | 0.02612 | 1.411 |
| PRR4 | proline rich 4 (lacrimal)(PRR4) | NM_001098538 | 116.9 | 108.1 | 0.02613 | 0.858 |
| OGFOD3 | 2-oxoglutarate and iron dependent oxygenase domain containing 3(OGFOD3) | NM_175902 | 108.8 | 99 | 0.02616 | 0.867 |
| | | XM_001725354 | 119.8 | 108.9 | 0.02617 | 0.865 |
| VWA1 | von Willebrand factor A domain containing 1(VWA1) | NM_022834 | 101.9 | 125.3 | 0.02625 | 1.189 |
| fcrl2 | Fc receptor like 2(FCRL2) | NM_030764 | 187.4 | 135.9 | 0.02637 | 0.675 |
| PSEN1 | presenilin 1(PSEN1) | NM_007318 | 111.6 | 130.4 | 0.02646 | 1.108 |
| Dgcr5 | DiGeorge syndrome critical region gene 5 (non-protein coding)(DGCR5) | NR_002733 | 98.8 | 122.5 | 0.02684 | 1.163 |
| TUBGCP4 | tubulin gamma complex associated protein 4(TUBGCP4) | NM_014444 | 96.3 | 114.7 | 0.02686 | 1.148 |
| PCP4 | Purkinje cell protein 4(PCP4) | NM_006198 | 96.1 | 159.9 | 0.02691 | 1.528 |
| PSMA7 | proteasome subunit alpha 7(PSMA7) | NM_152255 | 114.1 | 103 | 0.02694 | 0.857 |

| | | | | | | |
|---|---|---|---|---|---|---|
| TPP1 | tripeptidyl peptidase 1(TPP1) | NM_000391 | 1257 | 865.7 | 0.02701 | 0.592 |
| UTRN | utrophin(UTRN) | NM_007124 | 106.4 | 147.9 | 0.02706 | 1.298 |
| MYO1C | myosin IC(MYO1C) | NM_033375 | 146.8 | 136.1 | 0.02711 | 0.886 |
| CDK4 | cyclin dependent kinase 4(CDK4) | NM_000075 | 1229.8 | 2746.6 | 0.02714 | 2.031 |
| | | AA484800 | 107.6 | 93.7 | 0.02724 | 0.846 |
| TARBP2 | TARBP2, RISC loading complex RNA binding subunit(TARBP2) | NM_134323 | 100 | 119.5 | 0.02734 | 1.137 |
| XRRA1 | X-ray radiation resistance associated 1(XRRA1) | NM_182969 | 134.2 | 125.3 | 0.02735 | 0.889 |
| CRK | CRK proto-oncogene, adaptor protein(CRK) | NM_016823 | 128.2 | 168.4 | 0.02736 | 1.209 |
| CRK | CRK proto-oncogene, adaptor protein(CRK) | NM_016823 | 147.1 | 173 | 0.02736 | 1.209 |
| CRK | CRK proto-oncogene, adaptor protein(CRK) | NM_016823 | 250.1 | 347.5 | 0.02736 | 1.209 |
| | | XM_001713654 | 6200.5 | 6337.7 | 0.02746 | 0.888 |
| VCL | vinculin(VCL) | NM_014000 | 1068 | 1829.8 | 0.02752 | 1.505 |
| PLPP2 | phospholipid phosphatase 2(PLPP2) | NM_177526 | 102.4 | 119.2 | 0.02753 | 1.123 |
| AMOTL2 | angiomotin like 2(AMOTL2) | NM_016201 | 284.1 | 233.1 | 0.02772 | 0.729 |
| LCE1B | late cornified envelope 1B(LCE1B) | NM_178349 | 131.2 | 120 | 0.02772 | 0.875 |

| | | | | | | |
|---|---|---|---|---|---|---|
| | | AW341257 | 126.1 | 117.1 | 0.02774 | 0.848 |
| GPX5 | glutathione peroxidase 5(GPX5) | NM_001509 | 124.2 | 106.9 | 0.02786 | 0.784 |
| GPX8 | glutathione peroxidase 8 (putative)(GPX8) | NM_001008397 | 135.3 | 109.9 | 0.02786 | 0.784 |
| SCN7A | sodium voltage-gated channel alpha subunit 7(SCN7A) | NM_002976 | 143.4 | 125.8 | 0.02787 | 0.838 |
| ZNF691 | zinc finger protein 691(ZNF691) | NM_015911 | 146.1 | 131.7 | 0.02793 | 0.838 |
| | | XM_001126896 | 121.5 | 113.4 | 0.02794 | 0.9 |
| c1orf116 | chromosome 1 open reading frame 116(C1orf116) | NM_023938 | 249.6 | 190.8 | 0.02798 | 0.674 |
| | | CD514751 | 111.1 | 106.3 | 0.028 | 0.917 |
| | | XM_001713613 | 100.5 | 93.9 | 0.02801 | 0.878 |
| ccdc15 | coiled-coil domain containing 15(CCDC15) | NM_025004 | 94.6 | 99.7 | 0.02803 | 1.132 |
| CCDC150 | coiled-coil domain containing 150(CCDC150) | NM_173646 | 107.3 | 128.6 | 0.02803 | 1.132 |
| | | XR_015544 | 218.2 | 193 | 0.02813 | 0.808 |
| MSMO1 | methylsterol monooxygenase 1(MSMO1) | NM_006745 | 144.3 | 246.1 | 0.0282 | 1.452 |

| | | | | | | | | | |
|---|---|---|---|---|---|---|---|---|---|
| POGZ | pogo transposable element with ZNF domain(POGZ) | NM_015100 | 112.2 | 150.6 | 0.02825 | 1.248 | | | |
| SERTAD4-AS1 | SERTAD4 antisense RNA 1(SERTAD4-AS1) | NR_024337 | 112.5 | 153.9 | 0.02827 | 1.291 | | | |
| | | XM_937676 | 143.9 | 135.7 | 0.02827 | 0.89 | | | |
| FERD3L | Fer3 like bHLH transcription factor(FERD3L) | NM_152898 | 249.8 | 267 | 0.02837 | 0.86 | | | |
| FER1L5 | fer-1 like family member 5(FER1L5) | NM_001077400 | 167.4 | 152.7 | 0.02837 | 0.86 | | | |
| FGF13 | fibroblast growth factor 13(FGF13) | NM_033642 | 95.8 | 111 | 0.02843 | 1.118 | | | |
| | | XM_001715647 | 137.4 | 129.6 | 0.02845 | 0.89 | | | |
| NGDN | neuroguidin(NGDN) | NM_015514 | 125 | 127.8 | 0.02871 | 1.31 | NM_017196 | 14.73 | 0.69 |
| NGDN | neuroguidin(NGDN) | NM_015514 | 160.8 | 240 | 0.02871 | 1.31 | NM_017196 | 14.73 | 0.69 |
| ITGA2 | integrin subunit alpha 2(ITGA2) | NM_002203 | 693.2 | 1187.6 | 0.02886 | 1.504 | | | |
| HNRNPU | heterogeneous nuclear ribonucleoprotein U(HNRNPU) | NM_004501 | 475.3 | 1009.4 | 0.02894 | 1.854 | NM_017196 | 14.73 | 0.69 |
| HNRNPR | heterogeneous nuclear ribonucleoprotein R(HNRNPR) | NM_005826 | 402.4 | 867.3 | 0.02894 | 1.854 | | | |
| LOC441666 | zinc finger protein 91 pseudogene(LOC441666) | NR_024380 | 122.9 | 112.8 | 0.02896 | 0.875 | | | |

| | | | | | | | | | |
|---|---|---|---|---|---|---|---|---|---|
| | | XM_001128586 | 95.5 | 121.3 | 0.029 | 1.186 | | | |
| NETO1 | neuropilin and tolloid like 1(NETO1) | AK123319 | 118.2 | 111.3 | 0.02906 | 0.889 | NM_031356 | 5.43 | 24.53 |
| AVP | arginine vasopressin(AVP) | NM_000490 | 133.2 | 124.1 | 0.02907 | 0.894 | NM_031356 | 5.43 | 24.53 |
| AVPI1 | arginine vasopressin induced 1(AVPI1) | NM_021732 | 139.9 | 163.5 | 0.02907 | 0.894 | | | |
| | | XM_001718145 | 94.8 | 113.1 | 0.02919 | 1.145 | | | |
| ZC2HC1A | zinc finger C2HC-type containing 1A(ZC2HC1A) | NM_016010 | 101.2 | 108.9 | 0.0292 | 1.306 | | | |
| ZC2HC1A | zinc finger C2HC-type containing 1A(ZC2HC1A) | NM_016010 | 126.1 | 180 | 0.0292 | 1.306 | | | |
| HVCN1 | hydrogen voltage gated channel 1(HVCN1) | NM_001040107 | 120.9 | 111.6 | 0.02928 | 0.871 | | | |
| DIXDC1 | DIX domain containing 1(DIXDC1) | NM_001037954 | 117 | 163.8 | 0.02932 | 1.305 | | | |
| DKC1 | dyskerin pseudouridine synthase 1(DKC1) | NM_001363 | 154.8 | 168.3 | 0.02932 | 1.305 | | | |
| CD36 | CD36 molecule(CD36) | NM_000072 | 536.7 | 1163.6 | 0.02933 | 1.957 | | | |
| CD34 | CD34 molecule(CD34) | NM_001773 | 157.2 | 158.6 | 0.02933 | 1.957 | | | |
| CD34 | CD34 molecule(CD34) | NM_001773 | 408.5 | 908.2 | 0.02933 | 1.957 | | | |
| LINC01132 | long intergenic non-protein coding RNA 1132(LINC01132) | AK125357 | 112.3 | 105 | 0.02951 | 0.883 | | | |

| | | | | | | | | | |
|---|---|---|---|---|---|---|---|---|---|
| SSBP4 | single stranded DNA binding protein 4(SSBP4) | NM_032627 | 124.6 | 189.1 | 0.02955 | 1.401 | | | |
| RPL36 | ribosomal protein L36(RPL36) | NM_015414 | 95.5 | 115.4 | 0.02965 | 1.161 | | | |
| | | XM_001713717 | 162 | 255.8 | 0.0297 | 1.381 | | | |
| | | XM_001717999 | 174.3 | 148.6 | 0.0297 | 0.797 | NM_017135 | 41.45 | 2.88 |
| HIF1A | hypoxia inducible factor 1 alpha subunit(HIF1A) | NM_001530 | 97.6 | 121.3 | 0.02976 | 1.203 | NM_017135 | 41.45 | 2.88 |
| HIF1A | hypoxia inducible factor 1 alpha subunit(HIF1A) | NM_181054 | 116.8 | 189.3 | 0.02976 | 1.203 | NM_017135 | 41.45 | 2.88 |
| | | XM_001719745 | 104.4 | 149.5 | 0.02976 | 1.313 | NM_017135 | 41.45 | 2.88 |
| AGO3 | argonaute 3, RISC catalytic component(AGO3) | NM_177422 | 159 | 149 | 0.02976 | 0.857 | NM_017135 | 41.45 | 2.88 |
| FKBP1A | FK506 binding protein 1A(FKBP1A) | NM_054014 | 147.4 | 154.1 | 0.02979 | 1.329 | | | |
| FKBP1A | FK506 binding protein 1A(FKBP1A) | NM_054014 | 368.7 | 566.7 | 0.02979 | 1.329 | | | |
| FKBP1A | FK506 binding protein 1A(FKBP1A) | NM_054014 | 412.5 | 668 | 0.02979 | 1.329 | | | |
| ZNF177 | zinc finger protein 177(ZNF177) | NM_003451 | 170.1 | 215.6 | 0.02983 | 1.139 | | | |
| DYNLRB1 | dynein light chain roadblock-type 1(DYNLRB1) | NM_014183 | 112.6 | 121.7 | 0.02987 | 1.216 | | | |

| | | | | | | | | | |
|---|---|---|---|---|---|---|---|---|---|
| DYNLRB1 | dynein light chain roadblock-type 1(DYNLRB1) | NM_014183 | 125.6 | 164.1 | 0.02987 | 1.216 | | | |
| DYNLRB1 | dynein light chain roadblock-type 1(DYNLRB1) | NM_014183 | 214.3 | 285.7 | 0.02987 | 1.216 | | | |
| DYNLRB1 | dynein light chain roadblock-type 1(DYNLRB1) | NM_014183 | 899.2 | 1510.4 | 0.02987 | 1.216 | NM_001033653 | 43.46 | 1.56 |
| | | XM_932276 | 98.7 | 115.2 | 0.0299 | 1.112 | | | |
| ROR2 | receptor tyrosine kinase like orphan receptor 2(ROR2) | NM_004560 | 97.2 | 124.8 | 0.02991 | 1.234 | | | |
| GOLGA6B | golgin A6 family member B(GOLGA6B) | NM_018652 | 193.4 | 265.4 | 0.02996 | 1.204 | | | |
| WSB1 | WD repeat and SOCS box containing 1(WSB1) | NM_134264 | 187.5 | 351 | 0.02996 | 1.625 | | | |
| ECM2 | extracellular matrix protein 2(ECM2) | NM_001393 | 119.7 | 152 | 0.02998 | 1.191 | | | |
| FAM131A | family with sequence similarity 131 member A(FAM131A) | NM_144635 | 200.9 | 487.4 | 0.03022 | 2.11 | | | |
| | | XM_938232 | 106.2 | 97.7 | 0.03028 | 0.871 | | | |
| FAM160B2 | family with sequence similarity 160 member B2(FAM160B2) | NM_022749 | 114.1 | 121 | 0.03036 | 1.137 | | | |

| | | | | | | |
|---|---|---|---|---|---|---|
| FAM161 A | family with sequence similarity 161 member A(FAM161A) | NM_0321 80 | 124.6 | 151.9 | 0.03036 | 1.137 |
| TRIL | TLR4 interactor with leucine rich repeats(TRIL) | NM_0148 17 | 318.2 | 748.9 | 0.03036 | 2.049 |
| BAK1 | BCL2 antagonist/killer 1(BAK1) | NM_0011 88 | 306.1 | 198.8 | 0.03038 | 0.597 |
| NRXN2 | neurexin 2(NRXN2) | NM_1387 34 | 127.5 | 125.9 | 0.03055 | 0.877 |
| NRXN2 | neurexin 2(NRXN2) | NM_1387 34 | 159.3 | 148 | 0.03055 | 0.877 |
| taf7 | TATA-box binding protein associated factor 7(TAF7) | NM_0056 42 | 167.2 | 279.4 | 0.03059 | 1.448 |
| Hist2h2a a3 | histone cluster 2 H2A family member a3(HIST2H2AA3) | NM_0035 16 | 99.5 | 120.8 | 0.03063 | 1.178 |
| P4HA1 | prolyl 4-hydroxylase subunit alpha 1(P4HA1) | NM_0009 17 | 98.8 | 118.6 | 0.03064 | 1.136 |
| | | AA883754 | 111.4 | 102.9 | 0.03065 | 0.876 |
| SELENO M | selenoprotein M(SELENOM) | NM_0804 30 | 137.8 | 236 | 0.03069 | 1.523 |
| arfip1 | ADP ribosylation factor interacting protein 1(ARFIP1) | NM_0010 25595 | 99.4 | 121.9 | 0.03071 | 1.19 |
| B4GALT 3 | beta-1,4-galactosyltransferase 3(B4GALT3) | NM_0037 79 | 118.7 | 106.1 | 0.03077 | 0.874 |

| | | | | | | | | | |
|---|---|---|---|---|---|---|---|---|---|
| SRSF3 | serine and arginine rich splicing factor 3(SRSF3) | NM_003017 | 160.3 | 248.6 | 0.03078 | 1.312 | | | |
| CD34 | CD34 molecule(CD34) | NM_001773 | 157.2 | 158.6 | 0.03083 | 1.913 | | | |
| CD34 | CD34 molecule(CD34) | NM_001773 | 408.5 | 908.2 | 0.03083 | 1.913 | | | |
| CD34 | CD34 molecule(CD34) | NM_001773 | 536.7 | 1163.6 | 0.03083 | 1.913 | NM_134407 | 50.47 | 11.16 |
| TCF12 | transcription factor 12(TCF12) | NM_207038 | 163.4 | 248 | 0.03084 | 1.308 | | | |
| STAT5B | signal transducer and activator of transcription 5B(STAT5B) | NM_012448 | 206.4 | 175.5 | 0.03084 | 0.777 | | | |
| | | XM_373547 | 101.1 | 116.1 | 0.03085 | 1.111 | | | |
| ATP6V1C1 | ATPase H+ transporting V1 subunit C1(ATP6V1C1) | NM_001695 | 101.3 | 102.8 | 0.03087 | 0.858 | | | |
| ATP6V1C2 | ATPase H+ transporting V1 subunit C2(ATP6V1C2) | NM_144583 | 130.8 | 121.1 | 0.03087 | 0.858 | | | |
| ERF | ETS2 repressor factor(ERF) | NM_006494 | 116.7 | 136.3 | 0.03088 | 1.118 | | | |
| MOSPD2 | motile sperm domain containing 2(MOSPD2) | NM_152581 | 122.9 | 116.4 | 0.03089 | 0.903 | | | |
| MOSPD2 | motile sperm domain containing 2(MOSPD2) | NM_152581 | 151.1 | 152.7 | 0.03089 | 0.903 | | | |
| FAM65C | family with sequence similarity 65 member C(FAM65C) | NM_080829 | 453.2 | 344.6 | 0.03096 | 0.669 | | | |

| | | | | | | | | | |
|---|---|---|---|---|---|---|---|---|---|
| | | XM_936382 | 138.4 | 126.5 | 0.03096 | 0.867 | | | |
| | | XM_001716660 | 116 | 103.6 | 0.03107 | 0.85 | | | |
| MBIP | MAP3K12 binding inhibitory protein 1(MBIP) | NM_016586 | 97.4 | 117.6 | 0.03129 | 1.358 | | | |
| MBIP | MAP3K12 binding inhibitory protein 1(MBIP) | NM_016586 | 168.4 | 261.3 | 0.03129 | 1.358 | | | |
| ZNF777 | zinc finger protein 777(ZNF777) | NM_015694 | 122.1 | 125.8 | 0.03145 | 1.213 | | | |
| ZNF777 | zinc finger protein 777(ZNF777) | NM_015694 | 141.1 | 183.9 | 0.03145 | 1.213 | | | |
| ZRANB2 | zinc finger RANBP2-type containing 2(ZRANB2) | NM_005455 | 117.1 | 148.5 | 0.03154 | 1.191 | | | |
| ZRANB2 | zinc finger RANBP2-type containing 2(ZRANB2) | NM_005455 | 155.9 | 197.1 | 0.03154 | 1.191 | | | |
| RCN1 | reticulocalbin 1(RCN1) | NM_002901 | 116.1 | 167.2 | 0.03157 | 1.345 | | | |
| QPRT | quinolinate phosphoribosyltransferase(QPRT) | NM_014298 | 1274.1 | 671.9 | 0.03159 | 0.45 | | | |
| MRO | maestro(MRO) | NM_031939 | 127.7 | 109 | 0.03161 | 0.799 | NM_001033706 | 2.42 | 10.86 |
| | | XM_929740 | 115.6 | 110.6 | 0.03163 | 0.91 | | | |

| | | | | | | | | | |
|---|---|---|---|---|---|---|---|---|---|
| WRNIP1 | Werner helicase interacting protein 1(WRNIP1) | NM_020135 | 148.8 | 203.6 | 0.03164 | 1.226 | | | |
| | | XM_926143 | 107.9 | 100.7 | 0.03167 | 0.879 | | | |
| BEND6 | BEN domain containing 6(BEND6) | NM_152731 | 93.3 | 110.4 | 0.03181 | 1.152 | | | |
| BEND7 | BEN domain containing 7(BEND7) | NM_152751 | 98.5 | 110.8 | 0.03181 | 1.152 | | | |
| MOCS1 | molybdenum cofactor synthesis 1(MOCS1) | NM_005943 | 221.3 | 192.3 | 0.03183 | 0.786 | NM_153300 | 0.14 | 61.17 |
| Miat | myocardial infarction associated transcript (non-protein coding)(MIAT) | NR_003491 | 110.1 | 138 | 0.03188 | 1.197 | | | |
| EFNA4 | ephrin A4(EFNA4) | NM_182689 | 149.9 | 139.8 | 0.03197 | 0.878 | NM_032416 | 108.51 | 1.87 |
| EXT2 | exostosin glycosyltransferase 2(EXT2) | NM_000401 | 242.3 | 383.3 | 0.03198 | 1.364 | | | |
| NSMCE1 | NSE1 homolog, SMC5-SMC6 complex component(NSMCE1) | NM_145080 | 243.5 | 237 | 0.03205 | 0.849 | | | |
| HYDIN | HYDIN, axonemal central pair apparatus protein(HYDIN) | NM_032821 | 103 | 115.2 | 0.03205 | 0.868 | | | |
| HYDIN | HYDIN, axonemal central pair apparatus protein(HYDIN) | NM_032821 | 112.7 | 118.8 | 0.03205 | 0.868 | | | |

| | | | | | | | | |
|---|---|---|---|---|---|---|---|---|
| HYDIN | HYDIN, axonemal central pair apparatus protein(HYDIN) | NM_032821 | 137.8 | 124 | 0.03205 | 0.868 | | |
| HYDIN | HYDIN, axonemal central pair apparatus protein(HYDIN) | NM_032821 | 150.8 | 144.1 | 0.03205 | 0.868 | | |
| N4BP2L2 | NEDD4 binding protein 2 like 2(N4BP2L2) | NM_014887 | 242.5 | 324.2 | 0.0321 | 1.183 | | |
| SMAGP | small cell adhesion glycoprotein(SMAGP) | NM_001031628 | 112.3 | 133.5 | 0.0321 | 1.139 | | |
| RCN3 | reticulocalbin 3(RCN3) | NM_020650 | 151.3 | 142.6 | 0.03212 | 0.886 | | |
| DYNLL2 | dynein light chain LC8-type 2(DYNLL2) | NM_080677 | 265.3 | 487.6 | 0.03212 | 1.593 | | |
| | | XR_038018 | 343.3 | 292.2 | 0.03218 | 0.75 | | |
| C6orf136 | chromosome 6 open reading frame 136(C6orf136) | NM_145029 | 226.6 | 175.9 | 0.03234 | 0.719 | | |
| c6orf106 | chromosome 6 open reading frame 106(C6orf106) | NM_024294 | 102.6 | 107.1 | 0.03234 | 0.719 | | |
| PTGS2 | prostaglandin-endoperoxide synthase 2(PTGS2) | NM_000963 | 114.5 | 293.7 | 0.0324 | 2.188 | | |
| ADAMTSL2 | ADAMTS like 2(ADAMTSL2) | NM_014694 | 100.9 | 116.1 | 0.03243 | 1.104 | NM_001191088 | 10.07 | 0.03 |
| SETBP1 | SET binding protein 1(SETBP1) | NM_015559 | 176 | 271.4 | 0.03246 | 1.362 | NM_001191088 | 10.07 | 0.03 |

| | | | | | | | | | |
|---|---|---|---|---|---|---|---|---|---|
| H3f3c | H3 histone family member 3C(H3F3C) | NM_001013699 | 100.1 | 119.8 | 0.03247 | 1.156 | NM_001191088 | 10.07 | 0.03 |
| H6PD | hexose-6-phosphate dehydrogenase/glucose 1-dehydrogenase(H6PD) | NM_004285 | 103.7 | 114.7 | 0.03247 | 1.156 | | | |
| PCBP3 | poly(rC) binding protein 3(PCBP3) | NM_020528 | 148.1 | 134.5 | 0.0325 | 0.867 | | | |
| SLC10A7 | solute carrier family 10 member 7(SLC10A7) | NM_001029998 | 103.4 | 112.9 | 0.03264 | 1.15 | | | |
| SLC10A7 | solute carrier family 10 member 7(SLC10A7) | NM_001029998 | 125.2 | 152.1 | 0.03264 | 1.15 | NM_012496 | 1095.39 | 8.76 |
| | | XM_001125904 | 126.9 | 168.5 | 0.03274 | 1.239 | | | |
| | | XM_927568 | 118.9 | 112 | 0.03275 | 0.891 | | | |
| SNORA14B | small nucleolar RNA, H/ACA box 14B(SNORA14B) | NR_002956 | 106.5 | 124.6 | 0.03278 | 1.103 | | | |
| | | XM_926372 | 112.8 | 102.5 | 0.03282 | 0.871 | | | |
| APPBP2 | amyloid beta precursor protein binding protein 2(APPBP2) | NM_006380 | 977.2 | 2039.5 | 0.03291 | 1.862 | | | |
| APP | amyloid beta precursor protein(APP) | NM_201414 | 658.1 | 1613.9 | 0.03291 | 1.862 | | | |
| ARMCX3 | armadillo repeat containing, X-linked 3(ARMCX3) | NM_177947 | 278.8 | 350 | 0.03296 | 1.32 | | | |

| | | | | | | |
|---|---|---|---|---|---|---|
| ARMCX3 | armadillo repeat containing, X-linked 3(ARMCX3) | NM_177948 | 343.6 | 527.9 | 0.03296 | 1.32 |
| TUBB4A | tubulin beta 4A class IVa(TUBB4A) | NM_006087 | 171.7 | 154.1 | 0.03321 | 0.84 |
| PASK | PAS domain containing serine/threonine kinase(PASK) | NM_015148 | 114 | 137.2 | 0.03337 | 1.143 |
| PASK | PAS domain containing serine/threonine kinase(PASK) | NM_015148 | 160 | 181.6 | 0.03337 | 1.143 |
| CLSTN1 | calsyntenin 1(CLSTN1) | NM_001009566 | 107.6 | 129.2 | 0.03337 | 1.142 |
| CLSTN1 | calsyntenin 1(CLSTN1) | NM_001009566 | 123.5 | 152.2 | 0.03337 | 1.142 |
| CLSTN1 | calsyntenin 1(CLSTN1) | NM_001009566 | 2058 | 2595.3 | 0.03337 | 1.142 |
| | | XM_930029 | 348.7 | 287.2 | 0.03353 | 0.73 |
| NUMBL | NUMB like, endocytic adaptor protein(NUMBL) | NM_004756 | 94.2 | 109.7 | 0.03374 | 1.124 |
| APRT | adenine phosphoribosyltransferase(APRT) | NM_000485 | 265.9 | 407.6 | 0.03376 | 1.315 |
| NPIPB13 | nuclear pore complex interacting protein family, member B13(NPIPB13) | NR_002555 | 209 | 172.2 | 0.03407 | 0.769 |
| DDX5 | DEAD-box helicase 5(DDX5) | NM_004396 | 217.9 | 303.9 | 0.03409 | 1.215 |

| | | | | | | |
|---|---|---|---|---|---|---|
| SELENOS | selenoprotein S(SELENOS) | NM_018445 | 167.6 | 266.4 | 0.03411 | 1.38 |
| SELENOS | selenoprotein S(SELENOS) | NM_018445 | 911.9 | 1149.3 | 0.03411 | 1.38 |
| CCDC25 | coiled-coil domain containing 25(CCDC25) | NM_018246 | 117 | 176.1 | 0.03412 | 1.34 |
| CCDC28A | coiled-coil domain containing 28A(CCDC28A) | NM_015439 | 280.8 | 296.2 | 0.03412 | 1.34 |
| | | XM_933277 | 112 | 105.3 | 0.03414 | 0.896 |
| | | XM_937706 | 375.2 | 306.2 | 0.03416 | 0.724 |
| | | XM_0011129527 | 218.6 | 388.6 | 0.03418 | 1.568 |
| INHBC | inhibin beta C subunit(INHBC) | NM_005538 | 109.7 | 133.3 | 0.03419 | 1.173 |
| ANO1 | anoctamin 1(ANO1) | NM_018043 | 112.4 | 167.9 | 0.03426 | 1.329 |
| ANO10 | anoctamin 10(ANO10) | NM_018075 | 116.6 | 168.9 | 0.03426 | 1.329 |
| SNHG3 | small nucleolar RNA host gene 3(SNHG3) | NR_002909 | 120.3 | 112.3 | 0.0343 | 0.897 |
| LINC00950 | long intergenic non-protein coding RNA 950(LINC00950) | NR_024006 | 122.7 | 112.3 | 0.03435 | 0.87 |
| SUPT3H | SPT3 homolog, SAGA and STAGA complex component(SUPT3H) | NM_181356 | 105.8 | 132.3 | 0.03445 | 1.191 |

| | | | | | | | | | |
|---|---|---|---|---|---|---|---|---|---|
| PLA2G2D | phospholipase A2 group IID(PLA2G2D) | NM_012400 | 103.8 | 98.5 | 0.03448 | 0.902 | | | |
| PLA2G2D | phospholipase A2 group IID(PLA2G2D) | NM_012400 | 457.9 | 475.7 | 0.03448 | 0.902 | | | |
| AAMDC | adipogenesis associated Mth938 domain containing(AAMDC) | NM_024684 | 122.9 | 125.6 | 0.0346 | 1.123 | | | |
| AAK1 | AP2 associated kinase 1(AAK1) | NM_014911 | 104.7 | 123.6 | 0.0346 | 1.123 | | | |
| SSBP2 | single stranded DNA binding protein 2(SSBP2) | NM_012446 | 312.9 | 513.6 | 0.03464 | 1.437 | | | |
| SPTY2D1 | SPT2 chromatin protein domain containing 1(SPTY2D1) | NM_194285 | 111.3 | 140.2 | 0.03469 | 1.191 | | | |
| C1orf162 | chromosome 1 open reading frame 162(C1orf162) | NM_174896 | 142.2 | 129 | 0.03471 | 0.866 | | | |
| THOC7 | THO complex 7(THOC7) | NM_025075 | 495.7 | 685 | 0.03473 | 1.198 | | | |
| PCDHGA10 | protocadherin gamma subfamily A, 10(PCDHGA10) | NM_032090 | 117.9 | 141.5 | 0.03496 | 1.149 | | | |
| | | NM_058183 | 97.7 | 115.6 | 0.03498 | 1.127 | | | |
| | | XM_942687 | 128.6 | 164.4 | 0.03501 | 1.218 | NM_013059 | 21.02 | 0.11 |
| | | XM_942687 | 160.6 | 189.5 | 0.03501 | 1.218 | | | |

| | | | | | | |
|---|---|---|---|---|---|---|
| SLC7A8 | solute carrier family 7 member 8(SLC7A8) | NM_182728 | 87.1 | 101.3 | 0.03503 | 1.126 |
| SLC7A8 | solute carrier family 7 member 8(SLC7A8) | NM_182728 | 295.1 | 183.7 | 0.03503 | 1.126 |
| MAP2K3 | mitogen-activated protein kinase kinase 3(MAP2K3) | NM_002756 | 118.9 | 101.4 | 0.03505 | 0.846 |
| | | XM_932354 | 142.7 | 130.1 | 0.03512 | 0.874 |
| MLXIPL | MLX interacting protein like(MLXIPL) | NM_032953 | 1590.4 | 709.9 | 0.03513 | 0.394 |
| FBXO11 | F-box protein 11(FBXO11) | NM_025133 | 351.7 | 461.9 | 0.03518 | 1.215 |
| FBXO15 | F-box protein 15(FBXO15) | NM_152676 | 514.1 | 726.3 | 0.03518 | 1.215 |
| BRINP3 | BMP/retinoic acid inducible neural specific 3(BRINP3) | NM_199051 | 102.6 | 173.8 | 0.03519 | 1.557 |
| PLEKHH2 | pleckstrin homology, MyTH4 and FERM domain containing H2(PLEKHH2) | NM_172069 | 146.6 | 195.1 | 0.03523 | 1.261 |
| DNAJC10 | DnaJ heat shock protein family (Hsp40) member C10(DNAJC10) | NM_018981 | 146.3 | 185 | 0.03526 | 1.163 |
| | | NM_001013840 | 122.3 | 145.6 | 0.03533 | 1.127 |
| | | NM_001013840 | 130.5 | 147.3 | 0.03533 | 1.127 |

| | | | | | | | | | |
|---|---|---|---|---|---|---|---|---|---|
| NECAB3 | N-terminal EF-hand calcium binding protein 3(NECAB3) | NM_031231 | 124 | 115 | 0.03539 | 0.868 | NM_012816 | 19.78 | 2.28 |
| FAM30A | family with sequence similarity 30, member A(FAM30A) | NM_014792 | 95 | 112.5 | 0.0354 | 1.134 | NM_012816 | 19.78 | 2.28 |
| | | AW664435 | 110.1 | 103.1 | 0.03543 | 0.864 | | | |
| | | XM_001715935 | 426.2 | 210.4 | 0.03552 | 0.451 | | | |
| MT1IP | metallothionein 1I, pseudogene(MT1IP) | NM_175621 | 1855.3 | 812.8 | 0.03556 | 0.385 | | | |
| ABAT | 4-aminobutyrate aminotransferase(ABAT) | NM_000663 | 131.5 | 137.6 | 0.03556 | 0.888 | | | |
| AATK | apoptosis associated tyrosine kinase(AATK) | NM_001080395 | 121.1 | 112.4 | 0.03556 | 0.888 | | | |
| HOXA5 | homeobox A5(HOXA5) | NM_019102 | 124.7 | 164.9 | 0.03566 | 1.234 | | | |
| MVB12B | multivesicular body subunit 12B(MVB12B) | NM_033446 | 108.1 | 145.9 | 0.03576 | 1.257 | | | |
| TAGLN3 | transgelin 3(TAGLN3) | NM_001008273 | 116.1 | 108.1 | 0.03579 | 0.903 | | | |
| DECR1 | 2,4-dienoyl-CoA reductase 1, mitochondrial(DECR1) | NM_001359 | 121.4 | 155 | 0.03583 | 1.194 | | | |
| SLC5A3 | solute carrier family 5 member 3(SLC5A3) | AK055913 | 98.8 | 128.4 | 0.03587 | 1.247 | | | |
| RSPH10B | radial spoke head 10 homolog B(RSPH10B) | NM_173565 | 101 | 96.7 | 0.03593 | 1.309 | | | |

| | | | | | | | | | |
|---|---|---|---|---|---|---|---|---|---|
| RSPH10B | radial spoke head 10 homolog B(RSPH10B) | NM_173565 | 125.9 | 182.1 | 0.03593 | 1.309 | | | |
| RSPH10B | radial spoke head 10 homolog B(RSPH10B) | NM_173565 | 137.5 | 227.6 | 0.03593 | 1.309 | | | |
| RSPH10B | radial spoke head 10 homolog B(RSPH10B) | NM_173565 | 142.1 | 189.9 | 0.03593 | 1.309 | | | |
| APOD | apolipoprotein D(APOD) | NM_001647 | 105.5 | 125.9 | 0.03605 | 1.134 | | | |
| JUP | junction plakoglobin(JUP) | NM_002230 | 106.2 | 155.1 | 0.03606 | 1.387 | | | |
| ALPPL2 | alkaline phosphatase, placental like 2(ALPPL2) | NM_031313 | 2121.6 | 1017.4 | 0.03611 | 0.405 | | | |
| SNTA1 | syntrophin alpha 1(SNTA1) | NM_003098 | 221.5 | 198 | 0.03626 | 0.821 | | | |
| FAM124A | family with sequence similarity 124 member A(FAM124A) | NM_145019 | 118.4 | 112.8 | 0.03626 | 0.92 | | | |
| | | XR_038380 | 532 | 436 | 0.03629 | 0.714 | | | |
| | | XM_927136 | 104.8 | 123 | 0.03638 | 1.117 | NM_031544 | 4.79 | 38.59 |
| C11ORF49 | chromosome 11 open reading frame 49(C11orf49) | NM_024113 | 105.1 | 98 | 0.0366 | 0.889 | NM_031544 | 4.79 | 38.59 |
| ANGPTL3 | angiopoietin like 3(ANGPTL3) | NM_014495 | 91.2 | 132.8 | 0.03662 | 1.391 | | | |
| RABGGTA | Rab geranylgeranyltransferase alpha subunit(RABGGTA) | NM_182836 | 93.8 | 112.5 | 0.03674 | 1.173 | | | |

| | | | | | | |
|---|---|---|---|---|---|---|
| | | XM_378876 | 127.6 | 120.8 | 0.03679 | 0.892 |
| | | AW051978 | 132.7 | 126.6 | 0.03685 | 0.903 |
| c16orf52 | chromosome 16 open reading frame 52(C16orf52) | NM_173501 | 108.6 | 125.1 | 0.03687 | 1.121 |
| ATP8B4 | ATPase phospholipid transporting 8B4 (putative)(ATP8B4) | NM_024837 | 139 | 130.6 | 0.03691 | 0.877 |
| CSNK2A1 | casein kinase 2 alpha 1(CSNK2A1) | NM_001895 | 133.1 | 194.4 | 0.03697 | 1.316 |
| CSNK2A2 | casein kinase 2 alpha 2(CSNK2A2) | NM_001896 | 270 | 294.8 | 0.03697 | 1.316 |
| NUCKS1 | nuclear casein kinase and cyclin dependent kinase substrate 1(NUCKS1) | AK095855 | 1253.8 | 1798.2 | 0.03699 | 1.26 |
| | | XM_001723239 | 13523.8 | 16849.9 | 0.03706 | 1.176 |
| DUSP22 | dual specificity phosphatase 22(DUSP22) | NM_020185 | 118.8 | 173 | 0.03711 | 1.35 |
| DUSP23 | dual specificity phosphatase 23(DUSP23) | NM_017823 | 177.7 | 290.6 | 0.03711 | 1.35 |
| ILF3 | interleukin enhancer binding factor 3(ILF3) | NM_012218 | 307.4 | 481.7 | 0.03712 | 1.378 |
| PCGF1 | polycomb group ring finger 1(PCGF1) | NM_032673 | 144.5 | 182.6 | 0.03728 | 1.136 |

| | | | | | | |
|---|---|---|---|---|---|---|
| DYNLRB1 | dynein light chain roadblock-type 1(DYNLRB1) | NM_014183 | 1232.5 | 2157.6 | 0.0373 | 1.567 |
| SYTL2 | synaptotagmin like 2(SYTL2) | NM_206928 | 149.7 | 221.5 | 0.0373 | 1.32 |
| STXBP4 | syntaxin binding protein 4(STXBP4) | NM_178509 | 104.3 | 137.7 | 0.03735 | 1.232 |
| SPTLC1 | serine palmitoyltransferase long chain base subunit 1(SPTLC1) | NM_006415 | 120 | 150.5 | 0.03752 | 1.193 |
| TNC | tenascin C(TNC) | NM_002160 | 118.4 | 248.5 | 0.03754 | 1.897 |
| TNC | tenascin C(TNC) | NM_002160 | 128.4 | 274.8 | 0.03754 | 1.897 |
| RAB11FIP2 | RAB11 family interacting protein 2(RAB11FIP2) | NM_014904 | 202.9 | 342.1 | 0.03765 | 1.378 |
| RAB11FIP2 | RAB11 family interacting protein 2(RAB11FIP2) | NM_014904 | 250.6 | 399.4 | 0.03765 | 1.378 |
| LOC105372028 | uncharacterized LOC105372028(LOC105372028) | DA128369 | 111.1 | 102.8 | 0.03765 | 0.893 |
| | | BQ021435 | 121.2 | 113.1 | 0.03773 | 0.89 |
| SIRT6 | sirtuin 6(SIRT6) | NM_016539 | 142.2 | 123.1 | 0.03774 | 0.826 |
| CNIH2 | cornichon family AMPA receptor auxiliary protein 2(CNIH2) | NM_182553 | 260.4 | 346.5 | 0.03775 | 1.302 |

| | | | | | | |
|---|---|---|---|---|---|---|
| CNIH1 | cornichon family AMPA receptor auxiliary protein 1(CNIH1) | NM_005776 | 202.1 | 301.9 | 0.03775 | 1.302 |
| VWA1 | von Willebrand factor A domain containing 1(VWA1) | NM_199121 | 106.7 | 100.5 | 0.03777 | 0.886 |
| TPM1 | tropomyosin 1 (alpha)(TPM1) | NM_001018008 | 108.4 | 139 | 0.0378 | 1.199 |
| TPM1 | tropomyosin 1 (alpha)(TPM1) | NM_001018008 | 109.7 | 138.8 | 0.0378 | 1.199 |
| CD14 | CD14 molecule(CD14) | NM_001040021 | 116.6 | 145.8 | 0.03783 | 1.17 |
| | | XR_017376 | 95.8 | 117.6 | 0.03787 | 1.202 |
| | | XR_017376 | 127.2 | 134.8 | 0.03787 | 1.202 |
| PKM | pyruvate kinase, muscle(PKM) | NM_182470 | 180.2 | 326.4 | 0.03789 | 1.586 |
| DHCR7 | 7-dehydrocholesterol reductase(DHCR7) | NM_001360 | 105.3 | 134 | 0.03791 | 1.205 |
| DHDDS | dehydrodolichyl diphosphate synthase subunit(DHDDS) | NM_024887 | 164 | 196.9 | 0.03791 | 1.205 |
| FGG | fibrinogen gamma chain(FGG) | NM_000509 | 385.4 | 236.6 | 0.03808 | 0.551 |
| ZFAT | zinc finger and AT-hook domain containing(ZFAT) | NM_001029939 | 133.6 | 116.7 | 0.03809 | 0.818 |

| | | | | | | |
|---|---|---|---|---|---|---|
| map4k4 | mitogen-activated protein kinase kinase kinase kinase 4(MAP4K4) | NM_145687 | 212.7 | 330.9 | 0.03818 | 1.324 |
| SMOX | spermine oxidase(SMOX) | NM_175842 | 108.9 | 121.2 | 0.03821 | 0.769 |
| SMOX | spermine oxidase(SMOX) | NM_175842 | 204.4 | 168.1 | 0.03821 | 0.769 |
| ARHGAP28 | Rho GTPase activating protein 28(ARHGAP28) | NM_030672 | 116.5 | 106.3 | 0.03831 | 0.874 |
| CHMP5 | charged multivesicular body protein 5(CHMP5) | NM_016410 | 217.8 | 321.6 | 0.03832 | 1.439 |
| CHMP5 | charged multivesicular body protein 5(CHMP5) | NM_016410 | 369.6 | 623.7 | 0.03832 | 1.439 |
| CHMP5 | charged multivesicular body protein 5(CHMP5) | NM_016410 | 1042.1 | 1140.5 | 0.03832 | 1.439 |
| | | AW294430 | 121.2 | 109.9 | 0.03833 | 0.874 |
| SORCS2 | sortilin related VPS10 domain containing receptor 2(SORCS2) | NM_020777 | 92.2 | 141.6 | 0.0384 | 1.451 |
| KLHL5 | kelch like family member 5(KLHL5) | NM_001007075 | 119.7 | 160.1 | 0.03841 | 1.498 |
| KLHL5 | kelch like family member 5(KLHL5) | NM_001007075 | 133.1 | 132.3 | 0.03841 | 1.498 |
| KLHL5 | kelch like family member 5(KLHL5) | NM_001007075 | 401.5 | 695.6 | 0.03841 | 1.498 |
| UBA3 | ubiquitin like modifier activating enzyme 3(UBA3) | NM_198197 | 147.6 | 189.4 | 0.03843 | 1.192 |

| | | | | | | |
|---|---|---|---|---|---|---|
| PCYT2 | phosphate cytidylyltransferase 2, ethanolamine(PCYT2) | NM_002861 | 161.6 | 142.2 | 0.03852 | 0.821 |
| EFS | embryonal Fyn-associated substrate(EFS) | NM_032459 | 87 | 108.4 | 0.03855 | 1.219 |
| HYLS1 | HYLS1, centriolar and ciliogenesis associated(HYLS1) | NM_145014 | 242.1 | 208.1 | 0.03856 | 0.771 |
| c12orf73 | chromosome 12 open reading frame 73(C12orf73) | NM_001135570 | 99.3 | 115.3 | 0.03857 | 1.112 |
| CFH | complement factor H(CFH) | NM_001014975 | 98.3 | 116.7 | 0.03872 | 1.512 |
| CFH | complement factor H(CFH) | NM_001014975 | 109.6 | 183.6 | 0.03872 | 1.512 |
| CFH | complement factor H(CFH) | NM_001014975 | 111.8 | 131.7 | 0.03872 | 1.512 |
| MAP1A | microtubule associated protein 1A(MAP1A) | NM_002373 | 117.1 | 159.4 | 0.0388 | 1.248 |
| | | BU622756 | 122.1 | 118.7 | 0.03894 | 0.903 |
| MTFMT | mitochondrial methionyl-tRNA formyltransferase(MTFMT) | NM_139242 | 136.7 | 124.5 | 0.03904 | 0.865 |
| MTFMT | mitochondrial methionyl-tRNA formyltransferase(MTFMT) | NM_139242 | 184 | 212.3 | 0.03904 | 0.865 |

| | | | | | | |
|---|---|---|---|---|---|---|
| MOAP1 | modulator of apoptosis 1(MOAP1) | NM_022151 | 116.8 | 149.8 | 0.0391 | 1.191 |
| | | XM_941861 | 259.1 | 350.4 | 0.03912 | 1.18 |
| | | XM_935575 | 104.3 | 129.7 | 0.0393 | 1.202 |
| MKRN2 | makorin ring finger protein 2(MKRN2) | NM_014160 | 227.6 | 334.8 | 0.03931 | 1.298 |
| VCL | vinculin(VCL) | NM_003373 | 139.4 | 198.9 | 0.03935 | 1.298 |
| TH | tyrosine hydroxylase(TH) | NM_199293 | 96.2 | 113.9 | 0.03936 | 1.145 |
| bcam | basal cell adhesion molecule (Lutheran blood group)(BCAM) | NM_005581 | 112 | 119.4 | 0.03939 | 1.197 |
| BCAP29 | B-cell receptor associated protein 29(BCAP29) | NM_001008405 | 120.5 | 152.8 | 0.03939 | 1.197 |
| ORC1 | origin recognition complex subunit 1(ORC1) | NM_004153 | 118.4 | 141.4 | 0.03942 | 1.142 |
| CFAP69 | cilia and flagella associated protein 69(CFAP69) | NM_001039706 | 109.5 | 105.8 | 0.03943 | 0.912 |
| RREB1 | ras responsive element binding protein 1(RREB1) | NM_001003699 | 113.2 | 108 | 0.03947 | 0.898 |
| RREB1 | ras responsive element binding protein 1(RREB1) | NM_001003699 | 116.1 | 114.1 | 0.03947 | 0.898 |

| | | | | | | |
|---|---|---|---|---|---|---|
| RREB1 | ras responsive element binding protein 1(RREB1) | NM_0010 03699 | 118.3 | 138.9 | 0.03947 | 0.898 |
| RREB1 | ras responsive element binding protein 1(RREB1) | NM_0010 03699 | 144.7 | 152.1 | 0.03947 | 0.898 |
| EIF5 | eukaryotic translation initiation factor 5(EIF5) | NM_0019 69 | 200.8 | 318.6 | 0.0395 | 1.387 |
| MMS19 | MMS19 homolog, cytosolic iron-sulfur assembly component(MMS19) | NM_0223 62 | 309.1 | 420.5 | 0.03954 | 1.28 |
| MMS19 | MMS19 homolog, cytosolic iron-sulfur assembly component(MMS19) | NM_0223 62 | 351.4 | 516.4 | 0.03954 | 1.28 |
| FXYD1 | FXYD domain containing ion transport regulator 1(FXYD1) | NM_0219 02 | 148.8 | 122.1 | 0.03956 | 0.786 |
| FXYD1 | FXYD domain containing ion transport regulator 1(FXYD1) | NM_0050 31 | 103.2 | 102.9 | 0.03956 | 0.786 |
| | | XM_00171 4161 | 145.7 | 133.1 | 0.03964 | 0.862 |
| ERCC8 | ERCC excision repair 8, CSA ubiquitin ligase complex subunit(ERCC8) | NM_0000 82 | 127.1 | 151.5 | 0.03993 | 1.138 |

| | | | | | | |
|---|---|---|---|---|---|---|
| ERCC4 | ERCC excision repair 4, endonuclease catalytic subunit(ERCC4) | NM_005236 | 91.9 | 107.8 | 0.03993 | 1.138 |
| | | XM_0011126750 | 93.7 | 107.4 | 0.03996 | 1.116 |
| | | XM_0011126750 | 116.2 | 172 | 0.03996 | 1.116 |
| DZANK1 | double zinc ribbon and ankyrin repeat domains 1(DZANK1) | NM_001099407 | 102.2 | 126.3 | 0.04004 | 1.181 |
| | | BM705347 | 108.7 | 98.5 | 0.04007 | 0.865 |
| AKT1 | AKT serine/threonine kinase 1(AKT1) | NM_001014432 | 7980.7 | 3741.1 | 0.0401 | 0.397 |
| TMEM219 | transmembrane protein 219(TMEM219) | NM_001083613 | 448.4 | 433 | 0.04012 | 0.833 |
| | | CN283269 | 135 | 131 | 0.04023 | 0.925 |
| ACTL6A | actin like 6A(ACTL6A) | NM_004301 | 697.6 | 1184.4 | 0.04033 | 1.478 |
| | | XM_943472 | 96.5 | 112.8 | 0.04063 | 1.104 |
| | | BX441478 | 90.2 | 106.8 | 0.04063 | 1.155 |
| TMEM64 | transmembrane protein 64(TMEM64) | AK095472 | 126.3 | 167.1 | 0.04065 | 1.211 |
| SORL1 | sortilin related receptor 1(SORL1) | NM_003105 | 152 | 340.2 | 0.04066 | 1.952 |
| ODF2 | outer dense fiber of sperm tails 2(ODF2) | NM_002540 | 93.2 | 113.4 | 0.04071 | 1.178 |

| | | | | | | | | | |
|---|---|---|---|---|---|---|---|---|---|
| ODF2 | outer dense fiber of sperm tails 2(ODF2) | NM_002540 | 170.7 | 226.7 | 0.04071 | 1.178 | | | |
| cyp4f3 | cytochrome P450 family 4 subfamily F member 3(CYP4F3) | NM_000896 | 1010.8 | 413.5 | 0.04089 | 0.351 | | | |
| | | XM_001725366 | 122.9 | 156.8 | 0.04093 | 1.188 | | | |
| RBP1 | retinol binding protein 1(RBP1) | NM_002899 | 169.5 | 313.5 | 0.04103 | 1.598 | | | |
| | | XR_017130 | 1504 | 2336.1 | 0.04106 | 1.365 | | | |
| PTK7 | protein tyrosine kinase 7 (inactive)(PTK7) | NM_152880 | 129.3 | 206.9 | 0.04107 | 1.45 | | | |
| UBQLN1 | ubiquilin 1(UBQLN1) | NM_053067 | 140.8 | 183.7 | 0.04107 | 1.189 | | | |
| UBQLN1 | ubiquilin 1(UBQLN1) | NM_053067 | 268.7 | 364.1 | 0.04107 | 1.189 | | | |
| EXOC7 | exocyst complex component 7(EXOC7) | NM_001013839 | 175.7 | 251.2 | 0.0411 | 1.26 | NM_031012 | 19.39 | 0.85 |
| | | XR_001336 | 121.5 | 110.1 | 0.04112 | 0.876 | | | |
| CDKL1 | cyclin dependent kinase like 1(CDKL1) | NM_004196 | 105 | 127.6 | 0.04113 | 1.137 | | | |
| SCRT1 | scratch family transcriptional repressor 1(SCRT1) | NM_031309 | 152 | 142.7 | 0.04125 | 0.891 | | | |
| MAU2 | MAU2 sister chromatid cohesion factor(MAU2) | NM_015329 | 119.3 | 115.2 | 0.04125 | 0.925 | NM_012904 | 10.16 | 55.22 |
| PRMT6 | protein arginine methyltransferase 6(PRMT6) | NM_018137 | 159.9 | 211 | 0.04125 | 1.188 | | | |

| | | | | | | | | | |
|---|---|---|---|---|---|---|---|---|---|
| PRMT6 | protein arginine methyltransferase 6(PRMT6) | NM_018137 | 217.9 | 271.8 | 0.04125 | 1.188 | | | |
| COPRS | coordinator of PRMT5 and differentiation stimulator(COPRS) | NM_018405 | 104.3 | 122 | 0.04128 | 1.129 | | | |
| CUL7 | cullin 7(CUL7) | NM_014780 | 374.7 | 374.2 | 0.04129 | 0.867 | | | |
| ecd | ecdysoneless cell cycle regulator(ECD) | NM_007265 | 106.6 | 133.6 | 0.04145 | 1.156 | | | |
| ECEL1 | endothelin converting enzyme like 1(ECEL1) | NM_004826 | 106.8 | 133.1 | 0.04145 | 1.156 | | | |
| KCNMB2 | potassium calcium-activated channel subfamily M regulatory beta subunit 2(KCNMB2) | NM_181361 | 117.7 | 109.4 | 0.0415 | 0.85 | | | |
| KCNMB2 | potassium calcium-activated channel subfamily M regulatory beta subunit 2(KCNMB2) | NM_005832 | 107.7 | 106.4 | 0.0415 | 0.85 | | | |
| TBC1D8B | TBC1 domain family member 8B(TBC1D8B) | NM_017752 | 111.7 | 114 | 0.04165 | 1.176 | | | |
| TBC1D8B | TBC1 domain family member 8B(TBC1D8B) | NM_017752 | 129.1 | 166.3 | 0.04165 | 1.176 | | | |
| | | XM_927939 | 129.4 | 155 | 0.04183 | 1.142 | | | |
| | | XM_934270 | 103.3 | 121.2 | 0.04184 | 1.132 | NM_012823 | 17.52 | 0.04 |

| | | | | | | |
|---|---|---|---|---|---|---|
| | | NM_178230 | 2370.4 | 3782.8 | 0.04191 | 1.398 |
| VENTXP1 | VENT homeobox pseudogene 1(VENTXP1) | NR_001559 | 112.7 | 119.1 | 0.04199 | 0.889 |
| VENTXP1 | VENT homeobox pseudogene 1(VENTXP1) | NR_001559 | 118.7 | 111.3 | 0.04199 | 0.889 |
| MYADM | myeloid associated differentiation marker(MYADM) | NM_001020820 | 623.5 | 1205.7 | 0.04203 | 1.735 |
| SLC5A10 | solute carrier family 5 member 10(SLC5A10) | NM_152351 | 131.7 | 109.5 | 0.04208 | 0.552 |
| SLC5A10 | solute carrier family 5 member 10(SLC5A10) | NM_152351 | 244.8 | 147.2 | 0.04208 | 0.552 |
| PPT2 | palmitoyl-protein thioesterase 2(PPT2) | NM_138717 | 97.6 | 115 | 0.0421 | 1.156 |
| PPT2 | palmitoyl-protein thioesterase 2(PPT2) | NM_138717 | 172.1 | 220.5 | 0.0421 | 1.156 |
| | | XM_001716173 | 134.1 | 114.2 | 0.0421 | 0.784 |
| | | XM_001722148 | 130.6 | 117.1 | 0.04219 | 0.861 |
| ZNF423 | zinc finger protein 423(ZNF423) | NM_015069 | 105 | 118.2 | 0.04221 | 1.274 |
| ZNF423 | zinc finger protein 423(ZNF423) | NM_015069 | 128.5 | 177.2 | 0.04221 | 1.274 |
| | | XM_113971 | 141.6 | 208.4 | 0.04226 | 1.312 |

| | | | | | | |
|---|---|---|---|---|---|---|
| WIPI2 | WD repeat domain, phosphoinositide interacting 2(WIPI2) | NM_016003 | 258.2 | 199.3 | 0.04226 | 0.689 |
| | | XM_001126128 | 117.8 | 108.1 | 0.04231 | 0.884 |
| EWSR1 | EWS RNA binding protein 1(EWSR1) | NM_013986 | 765.4 | 1037 | 0.04238 | 1.34 |
| EWSR1 | EWS RNA binding protein 1(EWSR1) | NM_005243 | 464.8 | 719.9 | 0.04238 | 1.34 |
| ZMYM5 | zinc finger MYM-type containing 5(ZMYM5) | NM_001039650 | 113.5 | 135 | 0.04239 | 1.143 |
| CALML3-AS1 | CALML3 antisense RNA 1(CALML3-AS1) | AL831877 | 3392.2 | 2742.2 | 0.04246 | 0.697 |
| F7 | coagulation factor VII(F7) | NM_000131 | 92.9 | 107.1 | 0.04254 | 1.124 |
| bola2 | bolA family member 2(BOLA2) | NM_001031827 | 107 | 130.3 | 0.04255 | 1.161 |
| bola2 | bolA family member 2(BOLA2) | NM_001031827 | 108.1 | 112.6 | 0.04255 | 1.161 |
| bola2 | bolA family member 2(BOLA2) | NM_001031827 | 197.4 | 255.1 | 0.04255 | 1.161 |
| bola2 | bolA family member 2(BOLA2) | NM_001031827 | 539.5 | 624.5 | 0.04255 | 1.161 |
| | | XM_927256 | 113.3 | 102 | 0.04261 | 0.876 |
| | | XM_001724530 | 116.3 | 110.4 | 0.04266 | 0.899 |
| CRY1 | cryptochrome circadian clock 1(CRY1) | NM_004075 | 105.8 | 116.6 | 0.04272 | 1.26 |
| cryaa | crystallin alpha A(CRYAA) | NM_000394 | 121.6 | 166.5 | 0.04272 | 1.26 |

| | | | | | | |
|---|---|---|---|---|---|---|
| PRMT1 | protein arginine methyltransferase 1(PRMT1) | NM_198319 | 451.8 | 726.2 | 0.04292 | 1.378 |
| ACOT2 | acyl-CoA thioesterase 2(ACOT2) | NM_006821 | 259.5 | 193.4 | 0.04295 | 0.682 |
| RASEF | RAS and EF-hand domain containing(RASEF) | NM_152573 | 133.6 | 229.4 | 0.04299 | 1.569 |
| ICA1 | islet cell autoantigen 1(ICA1) | NM_004968 | 108.4 | 101.2 | 0.04299 | 0.886 |
| KITLG | KIT ligand(KITLG) | NM_000899 | 136.5 | 148.7 | 0.04309 | 1.289 |
| KLB | klotho beta(KLB) | NM_175737 | 179.8 | 265.7 | 0.04309 | 1.289 |
| MRFAP1L1 | Morf4 family associated protein 1 like 1(MRFAP1L1) | NM_203462 | 454.5 | 751.2 | 0.04315 | 1.416 |
| ANKRD17 | ankyrin repeat domain 17(ANKRD17) | NM_198889 | 98.6 | 125.5 | 0.04317 | 1.202 |
| MAVS | mitochondrial antiviral signaling protein(MAVS) | NM_020746 | 104 | 126.3 | 0.0432 | 1.134 |
| DLX6 | distal-less homeobox 6(DLX6) | NM_005222 | 187.9 | 167 | 0.04327 | 0.831 |
| DTNA | dystrobrevin alpha(DTNA) | NM_032980 | 105.6 | 127.8 | 0.04331 | 1.158 |
| IER2 | immediate early response 2(IER2) | NM_004907 | 99.8 | 111.9 | 0.04342 | 1.088 |
| | | XM_929291 | 103.2 | 96.4 | 0.04361 | 0.897 |

| | | | | | | |
|---|---|---|---|---|---|---|
| TM2D3 | TM2 domain containing 3(TM2D3) | NM_078474 | 144.6 | 210.6 | 0.0437 | 1.311 |
| zswim5 | zinc finger SWIM-type containing 5(ZSWIM5) | NM_020883 | 105.2 | 134.2 | 0.04386 | 1.21 |
| CCDC115 | coiled-coil domain containing 115(CCDC115) | NM_032357 | 104.5 | 123.2 | 0.04389 | 1.125 |
| ZNF548 | zinc finger protein 548(ZNF548) | NM_152909 | 144.8 | 176.6 | 0.04393 | 1.115 |
| RP2 | retinitis pigmentosa 2 (X-linked recessive)(RP2) | NM_006915 | 108.5 | 133.8 | 0.04406 | 1.166 |
| FAM167B | family with sequence similarity 167 member B(FAM167B) | NM_032648 | 100.7 | 117.6 | 0.04406 | 1.118 |
| RASSF4 | Ras association domain family member 4(RASSF4) | NM_032023 | 267.4 | 209.6 | 0.04414 | 0.712 |
| DSCAM | DS cell adhesion molecule(DSCAM) | NM_001389 | 105.3 | 120.2 | 0.04416 | 1.084 |
| | | XR_039278 | 1398 | 2010.8 | 0.04419 | 1.281 |
| LOC284009 | uncharacterized LOC284009(LOC284009) | NM_001025459 | 134.1 | 123.9 | 0.0443 | 0.871 |
| RHOT1 | ras homolog family member T1(RHOT1) | NM_001033566 | 192.4 | 280.1 | 0.04439 | 1.266 |
| RHOT1 | ras homolog family member T1(RHOT1) | NM_001033566 | 316.4 | 379.3 | 0.04439 | 1.266 |
| | | XM_001133534 | 544.4 | 1026.9 | 0.04448 | 1.65 |

| | | | | | | | | | |
|---|---|---|---|---|---|---|---|---|---|
| | | XM_001133534 | 742.4 | 1375.2 | 0.04448 | 1.65 | | | |
| PFN2 | profilin 2(PFN2) | NM_053024 | 631.9 | 1133.6 | 0.0445 | 1.57 | | | |
| PHF6 | PHD finger protein 6(PHF6) | NM_001015877 | 113.8 | 149.1 | 0.04466 | 1.232 | NM_017170 | 13.37 | 0.05 |
| PTK7 | protein tyrosine kinase 7 (inactive)(PTK7) | NM_152882 | 134.6 | 210.4 | 0.04466 | 1.422 | | | |
| | | BF194881 | 131.1 | 124.9 | 0.04474 | 0.876 | | | |
| | | DB337588 | 118.9 | 112.8 | 0.04487 | 0.908 | | | |
| ZHX1 | zinc fingers and homeoboxes 1(ZHX1) | NM_007222 | 98.1 | 121 | 0.04493 | 1.171 | | | |
| | | XM_933543 | 123.3 | 117.8 | 0.04501 | 0.89 | | | |
| GAGE4 | G antigen 4(GAGE4) | NM_001474 | 145.7 | 227 | 0.04506 | 1.382 | | | |
| SORD | sorbitol dehydrogenase(SORD) | NM_003104 | 145.7 | 111.7 | 0.04511 | 0.696 | | | |
| MOBP | myelin-associated oligodendrocyte basic protein(MOBP) | NM_182935 | 115.1 | 106.7 | 0.04511 | 0.89 | | | |
| HCG4B | HLA complex group 4B (non-protein coding)(HCG4B) | NR_001317 | 110.7 | 130.6 | 0.04511 | 1.11 | NM_001106492 | 2.61 | 10.85 |
| SUGP2 | SURP and G-patch domain containing 2(SUGP2) | NM_001017392 | 140.4 | 227.5 | 0.04513 | 1.443 | NM_001106492 | 2.61 | 10.85 |

| | | | | | | | | | |
|---|---|---|---|---|---|---|---|---|---|
| ACTR1B | ARP1 actin-related protein 1 homolog B, centractin beta(ACTR1B) | NM_005735 | 387.6 | 517.3 | 0.04519 | 1.156 | NM_001106492 | 2.61 | 10.85 |
| IRF2BP2 | interferon regulatory factor 2 binding protein 2(IRF2BP2) | NM_182972 | 147.5 | 139.7 | 0.04519 | 0.895 | | | |
| PURB | purine rich element binding protein B(PURB) | NM_033224 | 536.6 | 838.9 | 0.0452 | 1.356 | | | |
| ABCC4 | ATP binding cassette subfamily C member 4(ABCC4) | NM_005845 | 90.7 | 102.9 | 0.04522 | 1.425 | | | |
| ABCC5 | ATP binding cassette subfamily C member 5(ABCC5) | NM_001023587 | 229.1 | 379.6 | 0.04522 | 1.425 | | | |
| DLAT | dihydrolipoamide S-acetyltransferase(DLAT) | NM_001931 | 531.2 | 1140.2 | 0.04524 | 1.862 | | | |
| GPN2 | GPN-loop GTPase 2(GPN2) | NM_018066 | 129.6 | 315.4 | 0.04528 | 2.074 | | | |
| | | XR_015609 | 111.5 | 108.6 | 0.04529 | 0.927 | | | |
| C9orf72 | chromosome 9 open reading frame 72(C9orf72) | NM_018325 | 175.1 | 223.9 | 0.0453 | 1.183 | | | |
| | | BQ185080 | 121.5 | 115.2 | 0.04532 | 0.9 | | | |
| | | XM_940079 | 126.1 | 169.1 | 0.0454 | 1.227 | | | |

| | | | | | | | | | |
|---|---|---|---|---|---|---|---|---|---|
| OR4F21 | olfactory receptor family 4 subfamily F member 21(OR4F21) | NM_001005504 | 144.4 | 131.2 | 0.04541 | 0.851 | | | |
| NOTCH3 | notch 3(NOTCH3) | NM_000435 | 196.5 | 302.7 | 0.04549 | 1.365 | | | |
| LCOR | ligand dependent nuclear receptor corepressor(LCOR) | NM_032440 | 89.9 | 110.8 | 0.04553 | 1.196 | | | |
| UBR2 | ubiquitin protein ligase E3 component n-recognin 2(UBR2) | NM_015255 | 205.4 | 204.3 | 0.04571 | 0.897 | | | |
| | | XR_017963 | 101 | 117.5 | 0.0458 | 1.101 | | | |
| MARK2 | microtubule affinity regulating kinase 2(MARK2) | NM_004954 | 121.8 | 155.8 | 0.04583 | 1.227 | | | |
| MPRIP | myosin phosphatase Rho interacting protein(MPRIP) | NM_015134 | 428.3 | 655.7 | 0.04591 | 1.341 | | | |
| CEACAM1 | carcinoembryonic antigen related cell adhesion molecule 1(CEACAM1) | NM_001712 | 98.8 | 127.4 | 0.04593 | 1.221 | | | |
| ANG | angiogenin(ANG) | NM_001097577 | 131.7 | 176.3 | 0.04596 | 1.203 | NM_001270681 | 26.79 | 0.09 |
| ARMCX3 | armadillo repeat containing, X-linked 3(ARMCX3) | NM_177947 | 275.7 | 414 | 0.04596 | 1.282 | | | |
| PAPOLG | poly(A) polymerase gamma(PAPOLG) | NM_022894 | 122.7 | 113.7 | 0.04608 | 0.904 | | | |

| | | | | | | | | | |
|---|---|---|---|---|---|---|---|---|---|
| HECTD2 | HECT domain E3 ubiquitin protein ligase 2(HECTD2) | NM_182765 | 499.3 | 744.7 | 0.04616 | 1.298 | | | |
| XRCC5 | X-ray repair cross complementing 5(XRCC5) | NM_021141 | 116.8 | 140.3 | 0.04621 | 1.154 | | | |
| STAG1 | stromal antigen 1(STAG1) | NM_005862 | 104 | 130.1 | 0.0463 | 1.302 | | | |
| STAG1 | stromal antigen 1(STAG1) | NM_005862 | 136.5 | 199.9 | 0.0463 | 1.302 | | | |
| TSPYL5 | TSPY like 5(TSPYL5) | NM_033512 | 110.9 | 144.3 | 0.0463 | 1.232 | NM_019373 | 263.23 | 1.01 |
| HPD | 4-hydroxyphenylpyruvate dioxygenase(HPD) | NM_002150 | 705.6 | 634.9 | 0.04632 | 0.773 | | | |
| DUSP27 | dual specificity phosphatase 27 (putative)(DUSP27) | NM_001080426 | 177.7 | 290.6 | 0.04634 | 1.441 | | | |
| PML | promyelocytic leukemia(PML) | NM_033238 | 142.1 | 132.7 | 0.04635 | 0.889 | | | |
| MT2A | metallothionein 2A(MT2A) | NM_005953 | 1687.2 | 886 | 0.04637 | 0.474 | | | |
| POFUT2 | protein O-fucosyltransferase 2(POFUT2) | NM_133635 | 175.3 | 253.2 | 0.04637 | 1.325 | | | |
| POFUT2 | protein O-fucosyltransferase 2(POFUT2) | NM_133635 | 210 | 324 | 0.04637 | 1.325 | | | |
| | | XM_001717507 | 150.8 | 134.5 | 0.04639 | 0.844 | | | |

| | | | | | | | | | |
|---|---|---|---|---|---|---|---|---|---|
| NAT14 | N-acetyltransferase 14 (putative)(NAT14) | NM_020378 | 109.4 | 133.7 | 0.04646 | 1.166 | | | |
| | | XM_001130428 | 130.9 | 122.2 | 0.04651 | 0.897 | | | |
| ZNF112 | zinc finger protein 112(ZNF112) | NM_001083335 | 124 | 121.6 | 0.04658 | 0.921 | | | |
| | | XM_0017184 35 | 1858.9 | 3170.4 | 0.04663 | 1.506 | | | |
| CTNS | cystinosin, lysosomal cystine transporter(CTNS) | NM_004937 | 92.9 | 109.1 | 0.04668 | 1.131 | | | |
| SLC12A1 | solute carrier family 12 member 1(SLC12A1) | NM_000338 | 354.4 | 919.4 | 0.04678 | 2.317 | NM_012778 | 744.78 | 0.71 |
| | | XM_001721828 | 127.3 | 114.6 | 0.0468 | 0.874 | NM_012778 | 744.78 | 0.71 |
| AMDHD2 | amidohydrolase domain containing 2(AMDHD2) | NM_015944 | 208.4 | 387.2 | 0.04693 | 1.585 | NM_012778 | 744.78 | 0.71 |
| ACYP2 | acylphosphatase 2(ACYP2) | NM_138448 | 331 | 583.4 | 0.0471 | 1.561 | | | |
| ACYP1 | acylphosphatase 1(ACYP1) | NM_001107 | 111.4 | 116.1 | 0.0471 | 1.561 | | | |
| | | XM_374200 | 110.6 | 105.2 | 0.0472 | 0.914 | NM_012909 | 6.73 | 5427.66 |
| | | XR_018889 | 161.9 | 252.8 | 0.04754 | 1.368 | NM_022181 | 0.40 | 160.77 |
| | | XM_944079 | 145 | 137.5 | 0.04756 | 0.884 | NM_019157 | 14.48 | 0.05 |
| PLPP5 | phospholipid phosphatase 5(PLPP5) | NM_032483 | 234.8 | 413.3 | 0.04771 | 1.501 | | | |

| | | | | | | |
|---|---|---|---|---|---|---|
| SLC30A5 | solute carrier family 30 member 5(SLC30A5) | NM_022902 | 155.6 | 204.1 | 0.04776 | 1.197 |
| SLC30A5 | solute carrier family 30 member 5(SLC30A5) | NM_022902 | 186.1 | 251.5 | 0.04776 | 1.197 |
| RAI1 | retinoic acid induced 1(RAI1) | NM_030665 | 166.5 | 224.9 | 0.04791 | 1.139 |
| RAI1 | retinoic acid induced 1(RAI1) | NM_030665 | 168.9 | 210.7 | 0.04791 | 1.139 |
| PALLD | palladin, cytoskeletal associated protein(PALLD) | NM_016081 | 289 | 670.3 | 0.04805 | 1.988 |
| HLA-DQB1 | major histocompatibility complex, class II, DQ beta 1(HLA-DQB1) | NM_002123 | 116.5 | 106.6 | 0.04807 | 0.894 |
| BCHE | butyrylcholinesterase(BCHE) | NM_000055 | 194.8 | 271.6 | 0.04814 | 1.219 |
| PRSS12 | protease, serine 12(PRSS12) | NM_003619 | 100.3 | 122.6 | 0.0482 | 1.17 |
| UNC50 | unc-50 inner nuclear membrane RNA binding protein(UNC50) | NM_014044 | 564.2 | 849.1 | 0.04823 | 1.295 |
| CAPRIN1 | cell cycle associated protein 1(CAPRIN1) | NM_005898 | 351.4 | 546.6 | 0.04829 | 1.359 |
| DCP2 | decapping mRNA 2(DCP2) | NM_152624 | 142 | 130.8 | 0.0483 | 0.871 |
| DDX24 | DEAD-box helicase 24(DDX24) | NM_020414 | 165.4 | 253.2 | 0.04831 | 1.363 |
| | | XM_001723486 | 116.7 | 107.4 | 0.04842 | 0.861 |

| | | | | | | | | | |
|---|---|---|---|---|---|---|---|---|---|
| CSK | c-src tyrosine kinase(CSK) | NM_004383 | 93.3 | 112.9 | 0.04845 | 1.16 | | | |
| KAT2B | lysine acetyltransferase 2B(KAT2B) | NM_003884 | 132.2 | 120.9 | 0.04847 | 0.865 | | | |
| URGCP | upregulator of cell proliferation(URGCP) | NM_017920 | 459.2 | 428.6 | 0.0485 | 0.826 | | | |
| UBE2E3 | ubiquitin conjugating enzyme E2 E3(UBE2E3) | NM_006357 | 127.3 | 131.4 | 0.04859 | 1.312 | | | |
| UBE2E3 | ubiquitin conjugating enzyme E2 E3(UBE2E3) | NM_006357 | 157.4 | 234.2 | 0.04859 | 1.312 | | | |
| UBE2E3 | ubiquitin conjugating enzyme E2 E3(UBE2E3) | NM_006357 | 431 | 760.1 | 0.04859 | 1.312 | | | |
| CYP4A11 | cytochrome P450 family 4 subfamily A member 11(CYP4A11) | NM_000778 | 124.5 | 120.8 | 0.0486 | 0.927 | | | |
| SOCS5 | suppressor of cytokine signaling 5(SOCS5) | NM_014011 | 149.1 | 221.4 | 0.04864 | 1.291 | | | |
| LRIG3 | leucine rich repeats and immunoglobulin like domains 3(LRIG3) | NM_153377 | 107.3 | 127.8 | 0.04864 | 1.161 | | | |
| DOK1 | docking protein 1(DOK1) | NM_001381 | 241.9 | 393.2 | 0.04874 | 1.434 | | | |
| | | XM_929565 | 217.9 | 198.6 | 0.04874 | 0.84 | | | |
| | | XR_038768 | 140.2 | 126.6 | 0.04876 | 0.848 | NM_019168 | 8.26 | 61.49 |
| | | AA454665 | 119.9 | 113.6 | 0.04878 | 0.894 | | | |

| | | | | | | |
|---|---|---|---|---|---|---|
| hyal3 | hyaluronoglucosaminidase 3(HYAL3) | NM_003549 | 1365.7 | 1085.5 | 0.04886 | 0.695 |
| MT1A | metallothionein 1A(MT1A) | NM_005946 | 11240.4 | 5630.1 | 0.04894 | 0.441 |
| DPM1 | dolichyl-phosphate mannosyltransferase subunit 1, catalytic(DPM1) | NM_003859 | 167.3 | 214.6 | 0.04898 | 1.208 |
| DPM2 | dolichyl-phosphate mannosyltransferase subunit 2, regulatory(DPM2) | NM_003863 | 463.5 | 648.1 | 0.04898 | 1.208 |
| TGM5 | transglutaminase 5(TGM5) | NM_201631 | 110.8 | 106.2 | 0.04898 | 0.915 |
| ACTR5 | ARP5 actin-related protein 5 homolog(ACTR5) | NM_024855 | 106.5 | 117.3 | 0.04901 | 1.071 |
| ZNF616 | zinc finger protein 616(ZNF616) | NM_178523 | 166.8 | 234 | 0.04903 | 1.24 |
| FGF18 | fibroblast growth factor 18(FGF18) | NM_003862 | 90.5 | 114.3 | 0.04906 | 1.216 |
| FBXO11 | F-box protein 11(FBXO11) | NM_018693 | 113.5 | 102.8 | 0.04911 | 0.876 |
| FBXO11 | F-box protein 11(FBXO11) | NM_025133 | 124.4 | 118.9 | 0.04911 | 0.876 |
| ST13 | suppression of tumorigenicity 13 (colon carcinoma) (Hsp70 interacting protein)(ST13) | NM_003932 | 272.3 | 206.7 | 0.04912 | 0.675 |

| | | | | | | |
|---|---|---|---|---|---|---|
| TTC3 | tetratricopeptide repeat domain 3(TTC3) | NM_003316 | 282.6 | 435.8 | 0.04931 | 1.339 |
| MYH10 | myosin heavy chain 10(MYH10) | NM_005964 | 999.3 | 1734.4 | 0.04936 | 1.546 |
| | | XM_498859 | 118.2 | 110.8 | 0.04938 | 0.898 |
| UGT1A6 | UDP glucuronosyltransferase family 1 member A6(UGT1A6) | NM_001072 | 146.1 | 129 | 0.0496 | 0.714 |
| UGT1A6 | UDP glucuronosyltransferase family 1 member A6(UGT1A6) | NM_001072 | 161.1 | 128.8 | 0.0496 | 0.714 |
| ZNF567 | zinc finger protein 567(ZNF567) | NM_152603 | 100.2 | 117.6 | 0.04961 | 1.128 |
| ATP5A1 | ATP synthase, H+ transporting, mitochondrial F1 complex, alpha subunit 1, cardiac muscle(ATP5A1) | NM_004046 | 164.7 | 291.5 | 0.04966 | 1.527 |
| FAM131C | family with sequence similarity 131 member C(FAM131C) | NM_182623 | 108.1 | 131.4 | 0.04966 | 1.167 |
| FAM132A | family with sequence similarity 132 member A(FAM132A) | NM_001014980 | 173.9 | 170.2 | 0.04966 | 1.167 |
| RAP1GDS1 | Rap1 GTPase-GDP dissociation stimulator 1(RAP1GDS1) | NM_021159 | 152.2 | 196.2 | 0.04973 | 1.197 |

| | | | | | | |
|---|---|---|---|---|---|---|
| sdc1 | syndecan 1(SDC1) | NM_001006946 | 449.1 | 285.7 | 0.04975 | 0.562 |
| ARMC8 | armadillo repeat containing 8(ARMC8) | NM_015396 | 106.4 | 125.8 | 0.04976 | 1.106 |
| ARMC8 | armadillo repeat containing 8(ARMC8) | NM_213654 | 108.6 | 128.3 | 0.04976 | 1.106 |
| DEFB123 | defensin beta 123(DEFB123) | NM_153324 | 102.3 | 126.3 | 0.04976 | 1.152 |
| AGPS | alkylglycerone phosphate synthase(AGPS) | NM_003659 | 153.7 | 145 | 0.04978 | 0.884 |
| SNORD3A | small nucleolar RNA, C/D box 3A(SNORD3A) | NR_006880 | 173.2 | 158.3 | 0.04979 | 0.841 |
| MMGT1 | membrane magnesium transporter 1(MMGT1) | NM_173470 | 182.1 | 208.7 | 0.04979 | 1.271 |
| MMGT1 | membrane magnesium transporter 1(MMGT1) | NM_173470 | 605.3 | 897.2 | 0.04979 | 1.271 |
| OR2B2 | olfactory receptor family 2 subfamily B member 2(OR2B2) | NM_033057 | 129.6 | 118 | 0.04984 | 0.839 |
| STX12 | syntaxin 12(STX12) | NM_177424 | 188.3 | 246.2 | 0.04986 | 1.168 |
| fosl1 | FOS like 1, AP-1 transcription factor subunit(FOSL1) | NM_005438 | 194.5 | 2066.5 | 0.0499 | 10.276 |
| TBX3 | T-box 3(TBX3) | NM_005996 | 111.7 | 132.8 | 0.04995 | 1.121 |
| CHMP2A | charged multivesicular body protein 2A(CHMP2A) | NM_014453 | 130 | 152.2 | 0.05 | 1.116 |

| Gene Symbol | Gene Name | Accession No. | IUGR Chow M Mean | IUGR HFHS M Mean | p-val IUGR M HFHS vs. CON M HFHS | Ratio IUGR M HFHS vs. CON M HFHS | REFSEQ ID | NEPHRON (MEAN RPKM) | COLLECTING DUCT (MEAN RPKM) |
|---|---|---|---|---|---|---|---|---|---|
| | | XM_927511 | 92.1 | 101.9 | 0.00001 | 1.219 | | | |
| VRK3 | vaccinia related kinase 3(VRK3) | NM_001025778 | 96.8 | 95.2 | 0.00003 | 1.28 | | | |
| VRK3 | vaccinia related kinase 3(VRK3) | NM_001025778 | 180.3 | 150.5 | 0.00003 | 1.28 | | | |
| MPHOSPH10 | M-phase phosphoprotein 10(MPHOSPH10) | NM_005791 | 409.1 | 391.6 | 0.00005 | 1.389 | | | |
| | | XM_927848 | 101.2 | 99 | 0.00007 | 0.784 | | | |
| fam134b | family with sequence similarity 134 member B(FAM134B) | NM_001034850 | 368.5 | 320.8 | 0.0001 | 1.602 | | | |
| RPS26 | ribosomal protein S26(RPS26) | NM_001029 | 93 | 82.7 | 0.00011 | 0.838 | | | |
| RPS26 | ribosomal protein S26(RPS26) | NM_001029 | 1438.4 | 1055.6 | 0.00011 | 0.838 | | | |
| TMEM184C | transmembrane protein 184C(TMEM184C) | NM_018241 | 103.1 | 104.8 | 0.00017 | 1.194 | NM_017193 | 356.13 | 8.18 |
| TMEM184C | transmembrane protein 184C(TMEM184C) | NM_018241 | 256.8 | 245.8 | 0.00017 | 1.194 | NM_017193 | 356.13 | 8.18 |
| | | XM_0011133500 | 129.3 | 163.1 | 0.00018 | 1.458 | NM_017193 | 356.13 | 8.18 |
| | | XM_934672 | 104.5 | 101 | 0.00019 | 1.15 | NM_017193 | 356.13 | 8.18 |

| | | | | | | |
|---|---|---|---|---|---|---|
| | | XM_001723164 | 122.5 | 137.8 | 0.0002 | 1.153 |
| SEZ6L2 | seizure related 6 homolog like 2(SEZ6L2) | NM_201575 | 102 | 87.2 | 0.0002 | 0.84 |
| RAI1 | retinoic acid induced 1(RAI1) | NM_030665 | 142.8 | 121.4 | 0.00025 | 0.733 |
| RAI1 | retinoic acid induced 1(RAI1) | NM_030665 | 164.3 | 172 | 0.00025 | 0.733 |
| | | AI350600 | 97.7 | 103.8 | 0.00025 | 1.154 |
| | | XM_001717043 | 129.2 | 145.6 | 0.00032 | 1.42 |
| TTLL9 | tubulin tyrosine ligase like 9(TTLL9) | NM_001008409 | 92.1 | 100.1 | 0.00035 | 1.257 |
| CCND1 | cyclin D1(CCND1) | NM_053056 | 138.6 | 148.6 | 0.00039 | 1.304 |
| CCNC | cyclin C(CCNC) | NM_005190 | 126.3 | 128.5 | 0.00039 | 1.304 |
| RRAGB | Ras related GTP binding B(RRAGB) | NM_006064 | 91.4 | 89 | 0.00041 | 0.653 |
| RRAGB | Ras related GTP binding B(RRAGB) | NM_006064 | 110.8 | 119.2 | 0.00041 | 0.653 |
| EIF6 | eukaryotic translation initiation factor 6(EIF6) | NM_181466 | 457.4 | 426.8 | 0.00046 | 1.382 |
| LOC101927421 | uncharacterized LOC101927421(LOC101927421) | BU517015 | 104.6 | 127.7 | 0.00048 | 1.322 |
| OR52N5 | olfactory receptor family 52 subfamily N member 5(OR52N5) | NM_001001922 | 99.9 | 106 | 0.0005 | 1.172 |

| | | | | | | |
|---|---|---|---|---|---|---|
| | | XM_001714678 | 99.8 | 102.5 | 0.00053 | 1.178 |
| RORA | RAR related orphan receptor A(RORA) | NM_134261 | 96.1 | 108.6 | 0.00054 | 1.188 |
| | | XM_936101 | 168.2 | 178.1 | 0.00061 | 1.303 |
| | | XR_039453 | 109.9 | 102.4 | 0.00064 | 1.167 |
| | | XR_039453 | 203.3 | 176 | 0.00064 | 1.167 |
| TIGD1 | tigger transposable element derived 1(TIGD1) | NM_145702 | 97.5 | 100.2 | 0.0007 | 1.191 |
| | | BX111977 | 101.4 | 93.1 | 0.00072 | 0.846 |
| | | XM_942158 | 98.7 | 101.1 | 0.00074 | 1.201 |
| ATCAY | ATCAY, caytaxin(ATCAY) | NM_033064 | 114.3 | 93.6 | 0.00081 | 0.801 |
| ZNF137P | zinc finger protein 137, pseudogene(ZNF137P) | NM_003438 | 123.9 | 107.3 | 0.00083 | 0.786 |
| RPN2 | ribophorin II(RPN2) | NM_002951 | 330.9 | 310.2 | 0.00087 | 1.341 |
| | | XM_927344 | 101.3 | 113.6 | 0.00091 | 1.175 |
| | | NM_000648 | 93.4 | 90.3 | 0.00092 | 1.291 |
| | | NM_000648 | 96.7 | 108.6 | 0.00092 | 1.291 |
| | | BX103136 | 108.4 | 96.7 | 0.00101 | 0.822 |
| MBP | myelin basic protein(MBP) | NM_001025090 | 252.9 | 264.5 | 0.00114 | 1.672 |

| | | | | | | |
|---|---|---|---|---|---|---|
| PPARA | peroxisome proliferator activated receptor alpha(PPARA) | NM_001001928 | 121.6 | 125.4 | 0.00131 | 1.312 |
| C3orf84 | chromosome 3 open reading frame 84(C3orf84) | NM_001080528 | 110.3 | 112.1 | 0.00132 | 1.229 |
| TRIM51EP | tripartite motif-containing 51E, pseudogene(TRIM51EP) | NM_001136118 | 115.4 | 121.1 | 0.0015 | 1.191 |
| | | XM_936103 | 158.8 | 169.6 | 0.00153 | 1.502 |
| NAA25 | N(alpha)-acetyltransferase 25, NatB auxiliary subunit(NAA25) | NM_024953 | 193.6 | 203.3 | 0.00155 | 1.316 |
| SFTA2 | surfactant associated 2(SFTA2) | NM_205854 | 100.1 | 91.9 | 0.00165 | 0.839 |
| PCDH18 | protocadherin 18(PCDH18) | NM_019035 | 117.6 | 102.5 | 0.00171 | 0.774 |
| PCDH18 | protocadherin 18(PCDH18) | NM_019035 | 223.4 | 162.3 | 0.00171 | 0.774 |
| CYP4X1 | cytochrome P450 family 4 subfamily X member 1(CYP4X1) | NM_178033 | 97.2 | 104.4 | 0.00191 | 1.245 |
| LRBA | LPS responsive beige-like anchor protein(LRBA) | NM_006726 | 99.3 | 92.8 | 0.00204 | 0.828 |
| GPM6B | glycoprotein M6B(GPM6B) | NM_001001995 | 130.7 | 142.1 | 0.00205 | 1.341 |

| | | | | | | |
|---|---|---|---|---|---|---|
| | | XM_944653 | 98.1 | 113.5 | 0.00211 | 1.177 |
| MCTS1 | MCTS1, re-initiation and release factor(MCTS1) | NM_014060 | 184.6 | 192.4 | 0.00215 | 1.407 |
| TBC1D7 | TBC1 domain family member 7(TBC1D7) | NM_016495 | 183.5 | 141.2 | 0.00227 | 0.768 |
| AAMDC | adipogenesis associated Mth938 domain containing(AAMDC) | NM_024684 | 111 | 113.1 | 0.00236 | 0.829 |
| AAK1 | AP2 associated kinase 1(AAK1) | NM_014911 | 100.8 | 89 | 0.00236 | 0.829 |
| | | BM930795 | 105.8 | 117.8 | 0.00236 | 1.192 |
| ARMCX2 | armadillo repeat containing, X-linked 2(ARMCX2) | NM_177949 | 95.7 | 103 | 0.00247 | 1.19 |
| | | BU587445 | 101.9 | 95.2 | 0.0025 | 0.887 |
| RORA | RAR related orphan receptor A(RORA) | AK055969 | 134.3 | 125.8 | 0.0025 | 1.384 |
| MEG8 | maternally expressed 8 (non-protein coding)(MEG8) | NR_024149 | 93.2 | 98.6 | 0.00251 | 1.234 |
| TRIM36 | tripartite motif containing 36(TRIM36) | NM_001017397 | 87.1 | 88.5 | 0.00253 | 0.842 |
| | | XM_0017177551 | 103.1 | 100.8 | 0.00256 | 1.155 |
| | | XM_0017155747 | 91.4 | 102.4 | 0.00263 | 1.195 |

| | | | | | | | | | |
|---|---|---|---|---|---|---|---|---|---|
| RSRC2 | arginine and serine rich coiled-coil 2(RSRC2) | NM_023012 | 112.4 | 102.5 | 0.00269 | 0.743 | | | |
| RSRC2 | arginine and serine rich coiled-coil 2(RSRC2) | NM_023012 | 607.3 | 465.3 | 0.00269 | 0.743 | | | |
| OPRM1 | opioid receptor mu 1(OPRM1) | NM_001008505 | 105.6 | 96.1 | 0.0027 | 0.889 | | | |
| OPRM1 | opioid receptor mu 1(OPRM1) | NM_001008505 | 143.3 | 135.2 | 0.0027 | 0.889 | | | |
| | | XR_017376 | 105.2 | 92.8 | 0.00277 | 0.809 | | | |
| | | XR_017376 | 108.1 | 99.5 | 0.00277 | 0.809 | | | |
| HAGH | hydroxyacylglutathione hydrolase(HAGH) | NM_005326 | 92.2 | 99.8 | 0.00277 | 1.247 | | | |
| SPRN | shadow of prion protein homolog (zebrafish)(SPRN) | NM_001012508 | 91.6 | 90.4 | 0.0028 | 1.152 | | | |
| SPRN | shadow of prion protein homolog (zebrafish)(SPRN) | NM_001012508 | 98.1 | 105.9 | 0.0028 | 1.152 | NM_053502 | 4.02 | 18.01 |
| BTLA | B and T lymphocyte associated(BTLA) | NM_181780 | 92.2 | 98.4 | 0.00283 | 1.179 | NM_053502 | 4.02 | 18.01 |
| PDGFRL | platelet derived growth factor receptor like(PDGFRL) | NM_006207 | 109.9 | 95.1 | 0.00285 | 0.793 | NM_053502 | 4.02 | 18.01 |
| SPAG6 | sperm associated antigen 6(SPAG6) | NM_172242 | 130.5 | 101.6 | 0.00289 | 0.775 | NM_053502 | 4.02 | 18.01 |
| Amy1a | amylase, alpha 1A (salivary)(AMY1A) | NM_001008221 | 90.5 | 87.3 | 0.00292 | 0.839 | | | |

| | | | | | | | | | |
|---|---|---|---|---|---|---|---|---|---|
| TMTC3 | transmembrane and tetratricopeptide repeat containing 3(TMTC3) | NM_181783 | 139.4 | 161.3 | 0.00292 | 1.292 | | | |
| ABCF1 | ATP binding cassette subfamily F member 1(ABCF1) | NM_001025091 | 107.1 | 94.8 | 0.00295 | 0.859 | | | |
| | | XM_001725241 | 102.1 | 85.2 | 0.00296 | 0.862 | | | |
| HEPACAM2 | HEPACAM family member 2(HEPACAM2) | NM_198151 | 123.5 | 128.7 | 0.00299 | 1.295 | | | |
| KALRN | kalirin, RhoGEF kinase(KALRN) | NM_003947 | 3377.9 | 2800.3 | 0.003 | 0.724 | | | |
| | | XM_944999 | 114.1 | 125 | 0.00301 | 1.21 | NM_001009670 | 33.30 | 2.04 |
| DCTN2 | dynactin subunit 2(DCTN2) | NM_006400 | 139.4 | 129 | 0.00302 | 1.179 | | | |
| DCTN1 | dynactin subunit 1(DCTN1) | NM_004082 | 91 | 99 | 0.00302 | 1.179 | | | |
| GORASP2 | golgi reassembly stacking protein 2(GORASP2) | NM_015530 | 103.5 | 98.9 | 0.00303 | 0.888 | | | |
| PPTC7 | PTC7 protein phosphatase homolog(PPTC7) | NM_139283 | 238.9 | 261.2 | 0.00311 | 1.449 | | | |
| ZNF548 | zinc finger protein 548(ZNF548) | NM_152909 | 137.1 | 117.4 | 0.00315 | 0.836 | | | |
| MYO19 | myosin XIX(MYO19) | NM_025109 | 91.7 | 86.7 | 0.00317 | 0.851 | | | |
| | | XM_939706 | 106.9 | 107.8 | 0.00318 | 1.244 | | | |

| | | | | | | |
|---|---|---|---|---|---|---|
| ATG4A | autophagy related 4A cysteine peptidase(ATG4A) | NM_178270 | 91.3 | 96.8 | 0.00324 | 1.193 |
| KCTD10 | potassium channel tetramerization domain containing 10(KCTD10) | NM_031954 | 96.8 | 82.7 | 0.00325 | 0.836 |
| SCN2B | sodium voltage-gated channel beta subunit 2(SCN2B) | NM_004588 | 96.5 | 101.2 | 0.00338 | 1.215 |
| BAG5 | BCL2 associated athanogene 5(BAG5) | NM_004873 | 152.1 | 151.7 | 0.00339 | 1.297 |
| JMJD1C | jumonji domain containing 1C(JMJD1C) | NM_032776 | 103 | 95.9 | 0.00346 | 0.854 |
| JMJD1C | jumonji domain containing 1C(JMJD1C) | NM_032776 | 238.7 | 158.7 | 0.00346 | 0.854 |
| JMJD1C | jumonji domain containing 1C(JMJD1C) | NM_032776 | 396.4 | 291.5 | 0.00346 | 0.854 |
| RRP1B | ribosomal RNA processing 1B(RRP1B) | NM_015056 | 97.9 | 98.3 | 0.0035 | 1.236 |
| CRHBP | corticotropin releasing hormone binding protein(CRHBP) | NM_001882 | 93.8 | 101 | 0.00351 | 1.189 |
| | | BX109970 | 104.8 | 100.8 | 0.00353 | 0.753 |
| | | XR_017532 | 214.7 | 191.6 | 0.00358 | 1.368 |
| | | XR_037047 | 96.6 | 100.5 | 0.00376 | 1.149 |

| | | | | | | |
|---|---|---|---|---|---|---|
| ZNF177 | zinc finger protein 177(ZNF177) | NM_003451 | 153.2 | 134.3 | 0.00385 | 0.827 |
| UBTF | upstream binding transcription factor, RNA polymerase I(UBTF) | NM_014233 | 95.9 | 97.8 | 0.00398 | 0.818 |
| UBTF | upstream binding transcription factor, RNA polymerase I(UBTF) | NM_014233 | 102.9 | 88.1 | 0.00398 | 0.818 |
| ZNF436 | zinc finger protein 436(ZNF436) | NM_030634 | 90.6 | 95.1 | 0.00409 | 0.798 |
| ZNF436 | zinc finger protein 436(ZNF436) | NM_030634 | 108.6 | 93.7 | 0.00409 | 0.798 |
| ABHD12 | abhydrolase domain containing 12(ABHD12) | NM_001042472 | 93.2 | 98.9 | 0.00415 | 1.155 |
| | | AA722034 | 102.7 | 109.3 | 0.00422 | 1.199 |
| FZD1 | frizzled class receptor 1(FZD1) | NM_003505 | 119.8 | 108.1 | 0.00423 | 1.169 |
| FZD2 | frizzled class receptor 2(FZD2) | NM_001466 | 149.7 | 143.3 | 0.00423 | 1.169 |
| PHF12 | PHD finger protein 12(PHF12) | NM_020889 | 103.9 | 97.8 | 0.0043 | 0.845 |
| | | XM_930798 | 105.9 | 90.4 | 0.00433 | 0.825 |
| RBCK1 | RANBP2-type and C3HC4-type zinc finger containing 1(RBCK1) | NM_006462 | 152.4 | 132.4 | 0.00437 | 1.12 |
| | | XM_942469 | 105 | 111.7 | 0.00445 | 1.271 |

| | | | | | | |
|---|---|---|---|---|---|---|
| | | XM_942469 | 150.9 | 128.9 | 0.00445 | 1.271 |
| ZNF138 | zinc finger protein 138(ZNF138) | NM_006524 | 93.5 | 98.2 | 0.00447 | 1.177 |
| GMCL1 | germ cell-less, spermatogenesis associated 1(GMCL1) | NM_178439 | 95.1 | 106.3 | 0.00451 | 1.207 |
| GMDS | GDP-mannose 4,6-dehydratase(GMDS) | NM_001500 | 160.1 | 135.5 | 0.00451 | 1.207 |
| | | XR_038119 | 100.7 | 111.4 | 0.00456 | 1.277 |
| LETMD1 | LETM1 domain containing 1(LETMD1) | NM_015416 | 125.2 | 116.1 | 0.0046 | 1.177 |
| LFNG | LFNG O-fucosylpeptide 3-beta-N-acetylglucosaminyltransferase(LFNG) | NM_001040167 | 130.4 | 136.3 | 0.0046 | 1.177 |
| | | XM_929701 | 102.4 | 108.8 | 0.00463 | 1.149 |
| SEL1L3 | SEL1L family member 3(SEL1L3) | NM_015187 | 116 | 113.6 | 0.00485 | 1.243 |
| | | XR_018424 | 113.2 | 115.5 | 0.00494 | 1.206 |
| c12orf4 | chromosome 12 open reading frame 4(C12orf4) | NM_020374 | 107.1 | 105 | 0.00496 | 1.165 |
| SH3GL1 | SH3 domain containing GRB2 like 1, endophilin A2(SH3GL1) | NM_003025 | 113 | 120.5 | 0.00499 | 1.234 |

| | | | | | | |
|---|---|---|---|---|---|---|
| ZBTB43 | zinc finger and BTB domain containing 43(ZBTB43) | NM_014007 | 139.1 | 120.8 | 0.00505 | 1.24 |
| ST3GAL3 | ST3 beta-galactoside alpha-2,3-sialyltransferase 3(ST3GAL3) | NM_174970 | 113.1 | 99.8 | 0.00511 | 0.864 |
| LST1 | leukocyte specific transcript 1(LST1) | NM_205840 | 100.3 | 103.6 | 0.00513 | 1.151 |
| | | BM681626 | 92.4 | 97.7 | 0.00513 | 1.18 |
| C11ORF49 | chromosome 11 open reading frame 49(C11orf49) | NM_024113 | 90.6 | 99.6 | 0.00514 | 1.177 |
| slc26a6 | solute carrier family 26 member 6(SLC26A6) | NM_001040454 | 88.9 | 84.6 | 0.00515 | 0.834 |
| EEPD1 | endonuclease/exonuclease/phosphatase family domain containing 1(EEPD1) | NM_030636 | 112.2 | 118 | 0.00522 | 0.683 |
| HEATR3 | HEAT repeat containing 3(HEATR3) | NM_182922 | 121.4 | 125.9 | 0.00526 | 1.24 |
| | | XR_040021 | 101.4 | 102.6 | 0.00532 | 1.129 |
| | | XR_015689 | 130.6 | 110.2 | 0.00547 | 0.802 |
| | | XM_001720837 | 102.1 | 103.9 | 0.00549 | 1.146 |
| | | AA705831 | 100.3 | 93.9 | 0.00551 | 0.88 |
| CCNL1 | cyclin L1(CCNL1) | NM_020307 | 105.6 | 109.2 | 0.00556 | 1.195 |

| | | | | | | |
|---|---|---|---|---|---|---|
| ASB8 | ankyrin repeat and SOCS box containing 8(ASB8) | NM_024095 | 104.7 | 99.3 | 0.00562 | 0.891 |
| ASB9 | ankyrin repeat and SOCS box containing 9(ASB9) | NM_024087 | 106.7 | 92.9 | 0.00562 | 0.891 |
| PIGN | phosphatidylinositol glycan anchor biosynthesis class N(PIGN) | NM_176787 | 95.4 | 94 | 0.00566 | 0.839 |
| PIGN | phosphatidylinositol glycan anchor biosynthesis class N(PIGN) | NM_176787 | 97.6 | 94.8 | 0.00566 | 0.839 |
| | | XM_940199 | 96.9 | 103.1 | 0.00573 | 1.14 |
| RORA | RAR related orphan receptor A(RORA) | NM_002943 | 97.5 | 98.3 | 0.00574 | 1.262 |
| RORA | RAR related orphan receptor A(RORA) | NM_002943 | 127.3 | 119.3 | 0.00574 | 1.262 |
| SS18 | SS18, nBAF chromatin remodeling complex subunit(SS18) | NM_005637 | 109 | 123 | 0.00577 | 1.326 |
| SS18 | SS18, nBAF chromatin remodeling complex subunit(SS18) | NM_005637 | 214.4 | 178.5 | 0.00577 | 1.326 |
| SS18 | SS18, nBAF chromatin remodeling complex subunit(SS18) | NM_005637 | 285.2 | 259.8 | 0.00577 | 1.326 |

| | | | | | | | | | |
|---|---|---|---|---|---|---|---|---|---|
| MRGPRD | MAS related GPR family member D(MRGPRD) | NM_198923 | 106.7 | 99.2 | 0.00594 | 0.815 | | | |
| CAPN10 | calpain 10(CAPN10) | NM_023083 | 88.4 | 99.8 | 0.00597 | 1.207 | | | |
| CAPN11 | calpain 11(CAPN11) | NM_007058 | 101 | 104 | 0.00597 | 1.207 | | | |
| CAMK1 | calcium/calmodulin dependent protein kinase I(CAMK1) | NM_003656 | 175.8 | 166 | 0.00599 | 1.243 | | | |
| GCSH | glycine cleavage system protein H(GCSH) | NM_004483 | 110.2 | 89.6 | 0.00601 | 0.785 | | | |
| RAB2B | RAB2B, member RAS oncogene family(RAB2B) | NM_032846 | 348.1 | 310 | 0.00607 | 1.399 | | | |
| E2F4 | E2F transcription factor 4(E2F4) | NM_001950 | 110.4 | 95.9 | 0.00616 | 0.871 | | | |
| STMN3 | stathmin 3(STMN3) | NM_015894 | 378.9 | 410.6 | 0.00632 | 1.407 | NM_001107943 | 0.63 | 112.77 |
| | | XR_017962 | 98.1 | 108.7 | 0.00633 | 1.189 | | | |
| | | XR_017962 | 99.9 | 105.9 | 0.00633 | 1.189 | | | |
| | | XM_001133686 | 91.8 | 104.2 | 0.00634 | 1.215 | | | |
| map4k4 | mitogen-activated protein kinase kinase kinase kinase 4(MAP4K4) | NM_145687 | 185.9 | 156.6 | 0.00636 | 0.66 | | | |

| | | | | | | | | | |
|---|---|---|---|---|---|---|---|---|---|
| | | XM_928495 | 95.6 | 91.9 | 0.00637 | 0.845 | | | |
| TMEM161A | transmembrane protein 161A(TMEM161A) | NM_017814 | 112.1 | 112.7 | 0.00638 | 1.182 | | | |
| KCNMB2 | potassium calcium-activated channel subfamily M regulatory beta subunit 2(KCNMB2) | NM_181361 | 106.3 | 103.7 | 0.00644 | 1.136 | | | |
| KCNMB2 | potassium calcium-activated channel subfamily M regulatory beta subunit 2(KCNMB2) | NM_005832 | 92.5 | 103 | 0.00644 | 1.136 | | | |
| MIR33B | microRNA 33b(MIR33B) | NR_030361 | 84.7 | 81.6 | 0.00664 | 0.845 | | | |
| PLXND1 | plexin D1(PLXND1) | NM_015103 | 104.3 | 84.3 | 0.00664 | 0.768 | | | |
| | | XM_932032 | 103.6 | 94.2 | 0.00665 | 0.839 | | | |
| | | XM_001716372 | 89.9 | 94.9 | 0.00667 | 1.185 | | | |
| MRO | maestro(MRO) | NM_031939 | 106.5 | 115.8 | 0.00679 | 1.332 | | | |
| SPINT1 | serine peptidase inhibitor, Kunitz type 1(SPINT1) | NM_003710 | 114.2 | 110.7 | 0.00698 | 0.604 | NM_134372 | 22.51 | 0.03 |
| RNA5S9 | RNA, 5S ribosomal 9(RNA5S9) | NR_023371 | 218.1 | 182.1 | 0.00698 | 1.316 | | | |

| | | | | | | |
|---|---|---|---|---|---|---|
| c16orf58 | chromosome 16 open reading frame 58(C16orf58) | NM_022744 | 100.8 | 107.7 | 0.00706 | 1.175 |
| RBBP8 | RB binding protein 8, endonuclease(RBBP8) | NM_002894 | 107.6 | 103.9 | 0.00709 | 1.126 |
| NUGGC | nuclear GTPase, germinal center associated(NUGGC) | NM_001010906 | 98.3 | 86.8 | 0.0071 | 0.824 |
| | | XM_001721670 | 98 | 103.6 | 0.0071 | 1.301 |
| NECTIN1 | nectin cell adhesion molecule 1(NECTIN1) | NM_203285 | 113.8 | 120.3 | 0.00713 | 1.223 |
| GSKIP | GSK3B interacting protein(GSKIP) | NM_016472 | 93.2 | 93.3 | 0.00721 | 1.216 |
| CT47A1 | cancer/testis antigen family 47, member A1(CT47A1) | NM_001080146 | 101.6 | 99.8 | 0.00721 | 1.167 |
| EMC4 | ER membrane protein complex subunit 4(EMC4) | NM_016454 | 95.5 | 105.9 | 0.00722 | 1.269 |
| ZNF398 | zinc finger protein 398(ZNF398) | NM_170686 | 99.8 | 106.7 | 0.00723 | 1.208 |
| ARL5B | ADP ribosylation factor like GTPase 5B(ARL5B) | NM_178815 | 128.4 | 140 | 0.0073 | 1.332 |
| DNHD1 | dynein heavy chain domain 1(DNHD1) | NM_173589 | 91.4 | 111 | 0.00735 | 1.379 |
| PRPF40A | pre-mRNA processing factor 40 homolog A(PRPF40A) | NM_017892 | 102.9 | 107 | 0.00735 | 1.175 |

| | | | | | | | | | |
|---|---|---|---|---|---|---|---|---|---|
| UPF3A | UPF3 regulator of nonsense transcripts homolog A (yeast)(UPF3A) | NM_023011 | 347.5 | 270.1 | 0.00755 | 1.418 | | | |
| NCDN | neurochondrin(NCDN) | NM_001014839 | 127.3 | 126.2 | 0.00756 | 1.143 | | | |
| OR8J3 | olfactory receptor family 8 subfamily J member 3(OR8J3) | NM_001004064 | 100.5 | 104.1 | 0.00756 | 1.113 | | | |
| | | XM_928281 | 118.5 | 117.2 | 0.00765 | 1.163 | | | |
| DLGAP4 | DLG associated protein 4(DLGAP4) | NM_001042486 | 116 | 92.9 | 0.00777 | 0.799 | | | |
| SNORA12 | small nucleolar RNA, H/ACA box 12(SNORA12) | NR_002954 | 110.3 | 100.7 | 0.0079 | 1.243 | | | |
| OVGP1 | oviductal glycoprotein 1(OVGP1) | NM_002557 | 102.4 | 101.3 | 0.00794 | 1.153 | | | |
| EIF1B | eukaryotic translation initiation factor 1B(EIF1B) | NM_005875 | 139.4 | 162.7 | 0.00795 | 1.232 | | | |
| EIF1AX | eukaryotic translation initiation factor 1A, X-linked(EIF1AX) | NM_001412 | 122.6 | 127 | 0.00795 | 1.232 | | | |
| TSPAN1 | tetraspanin 1(TSPAN1) | NM_005727 | 100.8 | 94.6 | 0.00802 | 0.835 | NM_001031645 | 1.93 | 10.04 |
| RHOT1 | ras homolog family member T1(RHOT1) | NM_001033566 | 196.7 | 172.1 | 0.00818 | 1.273 | | | |
| RHOT1 | ras homolog family member T1(RHOT1) | NM_001033566 | 316.1 | 331.4 | 0.00818 | 1.273 | | | |

| | | | | | | |
|---|---|---|---|---|---|---|
| IP6K1 | inositol hexakisphosphate kinase 1(IP6K1) | NM_153273 | 100.8 | 89.3 | 0.00819 | 1.358 |
| IP6K2 | inositol hexakisphosphate kinase 2(IP6K2) | NM_001005909 | 320.9 | 350.1 | 0.00819 | 1.358 |
| Snora47 | small nucleolar RNA, H/ACA box 47(SNORA47) | NR_003014 | 102.4 | 101.7 | 0.00826 | 1.14 |
| SCARNA12 | small Cajal body-specific RNA 12(SCARNA12) | NR_003010 | 115.5 | 112.6 | 0.00831 | 0.849 |
| CX3CR1 | C-X3-C motif chemokine receptor 1(CX3CR1) | NM_001337 | 94.9 | 106.3 | 0.00831 | 1.225 |
| CXCL10 | C-X-C motif chemokine ligand 10(CXCL10) | NM_001565 | 108.3 | 103.4 | 0.00831 | 1.225 |
| MIR21 | microRNA 21(MIR21) | NR_029493 | 98.3 | 100.3 | 0.00846 | 0.671 |
| ASCL2 | achaete-scute family bHLH transcription factor 2(ASCL2) | NM_005170 | 97.6 | 89.5 | 0.0085 | 0.858 |
| | | XM_934471 | 108.4 | 115.5 | 0.00856 | 1.181 |
| fxyd7 | FXYD domain containing ion transport regulator 7(FXYD7) | NM_022006 | 180.9 | 96.4 | 0.00857 | 1.136 |
| FXYD5 | FXYD domain containing ion transport regulator 5(FXYD5) | NM_144779 | 113 | 107.7 | 0.00857 | 1.136 |

| | | | | | | | | | |
|---|---|---|---|---|---|---|---|---|---|
| EIF2A | eukaryotic translation initiation factor 2A(EIF2A) | NM_032025 | 139.4 | 162.7 | 0.00872 | 1.352 | | | |
| IL21R | interleukin 21 receptor(IL21R) | NM_181078 | 106.5 | 115.5 | 0.00873 | 1.166 | NM_144748 | 66.99 | 0.14 |
| IL21R | interleukin 21 receptor(IL21R) | NM_181079 | 114.3 | 113.8 | 0.00873 | 1.166 | | | |
| DDX19B | DEAD-box helicase 19B(DDX19B) | NM_001014449 | 219.8 | 201.5 | 0.00874 | 0.834 | | | |
| DDX19A | DEAD-box helicase 19A(DDX19A) | NM_018332 | 109 | 114.6 | 0.00874 | 0.834 | | | |
| ZNF326 | zinc finger protein 326(ZNF326) | NM_182976 | 108.7 | 111.9 | 0.00881 | 1.218 | | | |
| ZNF326 | zinc finger protein 326(ZNF326) | NM_182976 | 120.3 | 125.4 | 0.00881 | 1.218 | | | |
| | | BX441478 | 85.8 | 80.2 | 0.00884 | 0.821 | | | |
| ADGRG3 | adhesion G protein-coupled receptor G3(ADGRG3) | NM_170776 | 103.4 | 102.1 | 0.00884 | 1.145 | | | |
| LINC01559 | long intergenic non-protein coding RNA 1559(LINC01559) | NM_182558 | 103.4 | 108 | 0.00886 | 1.171 | | | |
| bcam | basal cell adhesion molecule (Lutheran blood group)(BCAM) | NM_005581 | 373.6 | 360.4 | 0.00888 | 1.394 | NM_001107793 | 77.29 | 6.85 |
| C8orf59 | chromosome 8 open reading frame 59(C8orf59) | NM_001099672 | 258 | 258.8 | 0.00891 | 0.535 | NM_001107793 | 77.29 | 6.85 |
| | | XM_001715215 | 111.6 | 98.5 | 0.00899 | 0.854 | NM_001107793 | 77.29 | 6.85 |

| | | | | | | | | | |
|---|---|---|---|---|---|---|---|---|---|
| CEP41 | centrosomal protein 41(CEP41) | NM_018718 | 119.8 | 126.7 | 0.00906 | 1.205 | NM_001107793 | 77.29 | 6.85 |
| CEP55 | centrosomal protein 55(CEP55) | NM_018131 | 140.2 | 129.6 | 0.00906 | 1.205 | | | |
| CREBRF | CREB3 regulatory factor(CREBRF) | NM_153607 | 260.6 | 211.6 | 0.00906 | 1.385 | | | |
| CREBZF | CREB/ATF bZIP transcription factor(CREBZF) | NM_001039618 | 332 | 231.8 | 0.00906 | 1.385 | | | |
| | | XM_939026 | 102.8 | 97.1 | 0.00912 | 0.857 | | | |
| TONSL | tonsoku-like, DNA repair protein(TONSL) | NM_013432 | 143.4 | 132.6 | 0.00919 | 0.779 | | | |
| trak1 | trafficking kinesin protein 1(TRAK1) | NM_001042646 | 114.6 | 114.2 | 0.00922 | 1.176 | | | |
| trak1 | trafficking kinesin protein 1(TRAK1) | NM_001042646 | 131.9 | 136.7 | 0.00922 | 1.176 | | | |
| TGM5 | transglutaminase 5(TGM5) | NM_201631 | 99.9 | 102 | 0.00932 | 1.129 | | | |
| CHD4 | chromodomain helicase DNA binding protein 4(CHD4) | NM_001273 | 398.3 | 351 | 0.00936 | 0.761 | | | |
| CHD6 | chromodomain helicase DNA binding protein 6(CHD6) | NM_032221 | 566.9 | 433.9 | 0.00936 | 0.761 | | | |
| ELK3 | ELK3, ETS transcription factor(ELK3) | NM_005230 | 95.7 | 99.3 | 0.00939 | 1.149 | | | |
| CPLX2 | complexin 2(CPLX2) | NM_001008220 | 101.5 | 105.4 | 0.00941 | 1.122 | | | |

| | | | | | | |
|---|---|---|---|---|---|---|
| RRBP1 | ribosome binding protein 1(RRBP1) | NM_0010 42576 | 165.2 | 144.5 | 0.00948 | 1.276 |
| RRBP1 | ribosome binding protein 1(RRBP1) | NM_0010 42576 | 314.1 | 310.3 | 0.00948 | 1.276 |
| FBLN1 | fibulin 1(FBLN1) | NM_0064 86 | 95.7 | 104.7 | 0.00954 | 1.175 |
| ADCK1 | aarF domain containing kinase 1(ADCK1) | NM_0204 21 | 93.2 | 97.5 | 0.00963 | 1.205 |
| | | XM_92793 9 | 121.1 | 106.6 | 0.00977 | 0.835 |
| TENM2 | teneurin transmembrane protein 2(TENM2) | NM_0010 80428 | 108.7 | 106.2 | 0.00979 | 1.127 |
| IL1R2 | interleukin 1 receptor type 2(IL1R2) | NM_1733 43 | 88.6 | 97.1 | 0.00982 | 1.254 |
| TCP10L | t-complex 10-like(TCP10L) | NM_0182 77 | 93.8 | 88.8 | 0.00984 | 0.841 |
| c6orf106 | chromosome 6 open reading frame 106(C6orf106) | NM_0242 94 | 117.8 | 106.5 | 0.00985 | 0.906 |
| PTP4A1 | protein tyrosine phosphatase type IVA, member 1(PTP4A1) | NM_0034 63 | 101.6 | 106.9 | 0.0099 | 1.142 |
| RPL37A | ribosomal protein L37a(RPL37A) | NM_0009 98 | 4299.1 | 3407.7 | 0.00991 | 0.783 |
| SMARC D3 | SWI/SNF related, matrix associated, actin dependent regulator of chromatin, subfamily d, member 3(SMARCD3) | NM_0030 78 | 93.2 | 100.2 | 0.00993 | 1.137 |

| | | | | | | | | | |
|---|---|---|---|---|---|---|---|---|---|
| FREM2 | FRAS1 related extracellular matrix protein 2(FREM2) | NM_207361 | 94.1 | 101.8 | 0.01002 | 1.153 | | | |
| | | XM_930292 | 114.3 | 97.8 | 0.01016 | 0.849 | | | |
| | | DR980429 | 95.6 | 90.7 | 0.01018 | 0.842 | | | |
| | | XM_941821 | 100.1 | 107.3 | 0.01025 | 1.086 | | | |
| | | XR_019567 | 104.9 | 106.3 | 0.01035 | 1.185 | | | |
| TTLL11 | tubulin tyrosine ligase like 11(TTLL11) | NM_194252 | 110.2 | 96.6 | 0.01037 | 0.855 | | | |
| AGAP3 | ArfGAP with GTPase domain, ankyrin repeat and PH domain 3(AGAP3) | NM_001042535 | 106 | 126.3 | 0.01043 | 1.231 | | | |
| L3HYPDH | trans-L-3-hydroxyproline dehydratase(L3HYPDH) | NM_144581 | 116.3 | 116.2 | 0.01044 | 1.189 | | | |
| L3MBTL2 | L3MBTL2 polycomb repressive complex 1 subunit(L3MBTL2) | NM_031488 | 121 | 118.6 | 0.01044 | 1.189 | | | |
| UBE3A | ubiquitin protein ligase E3A(UBE3A) | NM_000462 | 219.6 | 191.4 | 0.01045 | 1.246 | | | |
| UBE3A | ubiquitin protein ligase E3A(UBE3A) | NM_000462 | 504.6 | 507.7 | 0.01045 | 1.246 | | | |
| C1QTNF6 | C1q and tumor necrosis factor related protein 6(C1QTNF6) | NM_031910 | 99.7 | 105 | 0.01046 | 1.112 | NM_001009603 | 28.32 | 0.90 |

| | | | | | | |
|---|---|---|---|---|---|---|
| C1QTNF6 | C1q and tumor necrosis factor related protein 6(C1QTNF6) | NM_182486 | 110.2 | 93.5 | 0.01046 | 1.112 |
| RSPH9 | radial spoke head 9 homolog(RSPH9) | NM_152732 | 98.3 | 93.2 | 0.01049 | 0.828 |
| RDH10 | retinol dehydrogenase 10 (all-trans)(RDH10) | NM_172037 | 197.6 | 344 | 0.01056 | 1.664 |
| SNRPD3 | small nuclear ribonucleoprotein D3 polypeptide(SNRPD3) | NM_004175 | 207.8 | 217.9 | 0.01057 | 1.603 |
| RAB13 | RAB13, member RAS oncogene family(RAB13) | NM_002870 | 113 | 113 | 0.01059 | 1.16 |
| DBT | dihydrolipoamide branched chain transacylase E2(DBT) | NM_001918 | 94.8 | 99.7 | 0.0106 | 1.173 |
| DCAF1 | DDB1 and CUL4 associated factor 1(DCAF1) | NM_014703 | 527 | 473.7 | 0.0106 | 1.173 |
| | | BX114974 | 99.5 | 104 | 0.01064 | 1.196 |
| | | XM_938867 | 97.7 | 91.2 | 0.01069 | 0.881 |
| BZW1 | basic leucine zipper and W2 domains 1(BZW1) | NM_014670 | 97.8 | 104.5 | 0.01076 | 1.195 |
| BZW2 | basic leucine zipper and W2 domains 2(BZW2) | NM_014038 | 109.2 | 104.1 | 0.01076 | 1.195 |
| GPN1 | GPN-loop GTPase 1(GPN1) | NM_007266 | 137.4 | 156.4 | 0.01078 | 0.819 |

| | | | | | | |
|---|---|---|---|---|---|---|
| GPM6B | glycoprotein M6B(GPM6B) | NM_001001995 | 91.1 | 87.2 | 0.01078 | 0.819 |
| MMP19 | matrix metallopeptidase 19(MMP19) | NM_002429 | 98.9 | 88.4 | 0.01079 | 0.881 |
| TUBG1 | tubulin gamma 1(TUBG1) | NM_001070 | 102.4 | 109.8 | 0.01092 | 0.756 |
| TUBG1 | tubulin gamma 1(TUBG1) | NM_001070 | 129.5 | 109.9 | 0.01092 | 0.756 |
| TUBG1 | tubulin gamma 1(TUBG1) | NM_001070 | 164.4 | 186.6 | 0.01092 | 0.756 |
| SIM1 | single-minded family bHLH transcription factor 1(SIM1) | NM_005068 | 104.1 | 95.8 | 0.01096 | 0.791 |
| slc26a6 | solute carrier family 26 member 6(SLC26A6) | NM_134426 | 99.6 | 106.9 | 0.011 | 1.275 |
| slc26a6 | solute carrier family 26 member 6(SLC26A6) | NM_134426 | 112.5 | 122.8 | 0.011 | 1.275 |
| prb1 | proline rich protein BstNI subfamily 1(PRB1) | NM_199354 | 94.5 | 102.6 | 0.01106 | 1.151 |
| CTNNB1 | catenin beta 1(CTNNB1) | NM_001098209 | 96.5 | 96.4 | 0.01106 | 1.239 |
| CTNNB1 | catenin beta 1(CTNNB1) | NM_001098209 | 148.2 | 145.9 | 0.01106 | 1.239 |
| CTNNBIP1 | catenin beta interacting protein 1(CTNNBIP1) | NM_020248 | 154 | 164.5 | 0.01106 | 1.239 |
| C6orf1 | chromosome 6 open reading frame 1(C6orf1) | NM_001008704 | 107.5 | 105.3 | 0.01112 | 1.242 |
| | | AW182385 | 95 | 81.2 | 0.01119 | 0.821 |

| | | | | | | |
|---|---|---|---|---|---|---|
| BAGE2 | B melanoma antigen family member 2(BAGE2) | NM_182482 | 170.4 | 172.4 | 0.01119 | 1.245 |
| TTC16 | tetratricopeptide repeat domain 16(TTC16) | NM_144965 | 109.5 | 105.2 | 0.0112 | 1.156 |
| PIGM | phosphatidylinositol glycan anchor biosynthesis class M(PIGM) | NM_145167 | 112.7 | 92.8 | 0.01129 | 0.797 |
| KDM5D | lysine demethylase 5D(KDM5D) | NM_004653 | 241 | 203.4 | 0.01132 | 1.415 |
| CCPG1 | cell cycle progression 1(CCPG1) | NM_020739 | 272.5 | 284.1 | 0.01138 | 1.444 |
| lrch4 | leucine rich repeats and calponin homology domain containing 4(LRCH4) | NM_002319 | 99.8 | 86.4 | 0.01141 | 0.856 |
| | | XM_938802 | 108.6 | 109.8 | 0.01146 | 1.222 |
| UGGT2 | UDP-glucose glycoprotein glucosyltransferase 2(UGGT2) | NM_020121 | 139.4 | 123.4 | 0.01154 | 0.84 |
| c5orf30 | chromosome 5 open reading frame 30(C5orf30) | NM_033211 | 129.9 | 119.6 | 0.01179 | 0.877 |
| MAPK7 | mitogen-activated protein kinase 7(MAPK7) | NM_139032 | 107.2 | 99.7 | 0.01189 | 0.85 |

| | | | | | | |
|---|---|---|---|---|---|---|
| MAPK8IP1 | mitogen-activated protein kinase 8 interacting protein 1(MAPK8IP1) | NM_005456 | 126.5 | 107.4 | 0.01189 | 0.85 |
| | | XM_927142 | 112.5 | 105.4 | 0.01196 | 1.174 |
| | | CR748464 | 100 | 101.4 | 0.01198 | 1.13 |
| MBP | myelin basic protein(MBP) | NM_001025101 | 158 | 179.2 | 0.01202 | 1.301 |
| CAB39L | calcium binding protein 39 like(CAB39L) | NM_001079670 | 96.5 | 98.9 | 0.01209 | 1.176 |
| | | XM_938374 | 85.8 | 98.5 | 0.01209 | 1.231 |
| TBRG4 | transforming growth factor beta regulator 4(TBRG4) | NM_004749 | 117 | 101 | 0.0121 | 0.866 |
| EGF | epidermal growth factor(EGF) | NM_001963 | 119.3 | 109.3 | 0.01217 | 0.816 |
| SQLE | squalene epoxidase(SQLE) | NM_003129 | 91.9 | 95.3 | 0.01224 | 0.779 |
| SQLE | squalene epoxidase(SQLE) | NM_003129 | 99.5 | 94.2 | 0.01224 | 0.779 |
| | | XM_0017133809 | 664.9 | 830.1 | 0.01224 | 2.787 |
| Hist1h2bk | histone cluster 1 H2B family member k(HIST1H2BK) | NM_080593 | 97.5 | 102 | 0.01226 | 1.157 |

| | | | | | | |
|---|---|---|---|---|---|---|
| UAP1L1 | UDP-N-acetylglucosamine pyrophosphorylase 1 like 1(UAP1L1) | NM_207309 | 109.2 | 112.3 | 0.01228 | 1.172 |
| | | XM_001723218 | 92.1 | 88.6 | 0.01229 | 0.87 |
| NUFIP2 | NUFIP2, FMR1 interacting protein 2(NUFIP2) | NM_020772 | 158.4 | 138.3 | 0.0123 | 1.382 |
| NUFIP2 | NUFIP2, FMR1 interacting protein 2(NUFIP2) | NM_020772 | 341.6 | 290.4 | 0.0123 | 1.382 |
| IVNS1ABP | influenza virus NS1A binding protein(IVNS1ABP) | NM_006469 | 100.2 | 106 | 0.01234 | 1.172 |
| ADGRL2 | adhesion G protein-coupled receptor L2(ADGRL2) | NM_012302 | 96.7 | 104.6 | 0.01237 | 1.182 |
| ALX3 | ALX homeobox 3(ALX3) | NM_006492 | 110.5 | 116.8 | 0.01242 | 0.513 |
| ALX1 | ALX homeobox 1(ALX1) | NM_006982 | 98.4 | 99.8 | 0.01242 | 0.513 |
| MATN4 | matrilin 4(MATN4) | NM_030592 | 85.9 | 85.7 | 0.01245 | 0.817 |
| MATN4 | matrilin 4(MATN4) | NM_030592 | 89 | 88.4 | 0.01245 | 0.817 |
| TNFRSF10B | TNF receptor superfamily member 10b(TNFRSF10B) | NM_003842 | 90 | 103.7 | 0.01247 | 1.21 |

| | | | | | | |
|---|---|---|---|---|---|---|
| TNFRSF 10B | TNF receptor superfamily member 10b(TNFRSF10B) | NM_0038 42 | 98.4 | 95 | 0.01247 | 1.21 |
| ITM2A | integral membrane protein 2A(ITM2A) | NM_0048 67 | 84.9 | 92.9 | 0.01249 | 1.152 |
| MRPL35 | mitochondrial ribosomal protein L35(MRPL35) | NM_1456 44 | 498.9 | 558.5 | 0.01249 | 1.371 |
| GSTM1 | glutathione S-transferase mu 1(GSTM1) | NM_0005 61 | 98.3 | 90.7 | 0.01251 | 0.89 |
| SMIM10 L2B | small integral membrane protein 10 like 2B(SMIM10L2B) | NR_02449 3 | 98.7 | 102.4 | 0.01253 | 1.142 |
| | | XM_94443 8 | 93.3 | 96 | 0.01258 | 1.187 |
| PCYOX1 L | prenylcysteine oxidase 1 like(PCYOX1L) | NM_0240 28 | 96 | 92.3 | 0.01264 | 0.885 |
| Snora2b | small nucleolar RNA, H/ACA box 2B(SNORA2B) | NR_00295 1 | 136 | 122.9 | 0.01265 | 1.285 |
| | | XR_01830 9 | 94.4 | 87.4 | 0.01274 | 0.758 |
| | | NM_0010 04302 | 96.2 | 94.7 | 0.01278 | 0.831 |
| | | XR_01569 5 | 104.4 | 93.5 | 0.01282 | 0.862 |
| | | XR_00149 8 | 97.8 | 105 | 0.01286 | 1.243 |
| | | XM_93114 1 | 96.3 | 100.6 | 0.01286 | 1.149 |

| | | | | | | |
|---|---|---|---|---|---|---|
| DNMT3L | DNA methyltransferase 3 like(DNMT3L) | NM_013369 | 114.9 | 114.2 | 0.01288 | 1.141 |
| DNMT3B | DNA methyltransferase 3 beta(DNMT3B) | NM_006892 | 86.8 | 90.6 | 0.01288 | 1.141 |
| | | XR_039041 | 3488.4 | 2355.4 | 0.01288 | 0.812 |
| | | DA572372 | 92.6 | 101.7 | 0.01293 | 1.163 |
| EIF2AK1 | eukaryotic translation initiation factor 2 alpha kinase 1(EIF2AK1) | NM_014413 | 522.9 | 441.4 | 0.01296 | 0.753 |
| ZSCAN32 | zinc finger and SCAN domain containing 32(ZSCAN32) | NM_017810 | 111.9 | 107.1 | 0.01299 | 1.171 |
| NFASC | neurofascin(NFASC) | NM_001005387 | 98.2 | 92.2 | 0.01303 | 0.902 |
| CSMD1 | CUB and Sushi multiple domains 1(CSMD1) | NM_033225 | 127.1 | 119.8 | 0.01306 | 0.804 |
| | | XR_037583 | 100.9 | 103.8 | 0.01316 | 1.08 |
| MAVS | mitochondrial antiviral signaling protein(MAVS) | NM_020746 | 108.1 | 92.9 | 0.01321 | 0.85 |
| | | AW297854 | 97.9 | 93.4 | 0.01335 | 0.879 |
| SLC35F1 | solute carrier family 35 member F1(SLC35F1) | NM_001029858 | 96.2 | 105.8 | 0.01344 | 1.163 |
| | | AI873192 | 127.3 | 108.1 | 0.01344 | 0.64 |
| MRPL53 | mitochondrial ribosomal protein L53(MRPL53) | NM_053050 | 155.2 | 157.6 | 0.01346 | 1.285 |

| | | | | | | | | | |
|---|---|---|---|---|---|---|---|---|---|
| | | BX116001 | 97.4 | 106.1 | 0.01352 | 1.171 | | | |
| LOC102724002 | uncharacterized LOC102724002(LOC102724002) | AK025166 | 99.7 | 96.9 | 0.01354 | 0.887 | | | |
| | | XM_370758 | 115.6 | 105.4 | 0.01357 | 0.913 | | | |
| DCUN1D4 | defective in cullin neddylation 1 domain containing 4(DCUN1D4) | NM_015115 | 177.1 | 187.7 | 0.01363 | 1.296 | | | |
| | | NM_001015047 | 100.2 | 108.3 | 0.01365 | 1.154 | | | |
| SNORA2C | small nucleolar RNA, H/ACA box 2C(SNORA2C) | NR_002968 | 102.5 | 91.1 | 0.01367 | 0.879 | NM_144744 | 0.11 | 40.01 |
| | | XM_930670 | 122.4 | 128.8 | 0.01372 | 1.364 | | | |
| Mir574 | microRNA 574(MIR574) | NR_030300 | 102.3 | 94.5 | 0.01377 | 0.783 | | | |
| DENR | density regulated re-initiation and release factor(DENR) | NM_003677 | 155.7 | 149.2 | 0.01383 | 1.226 | | | |
| DEPDC5 | DEP domain containing 5(DEPDC5) | NM_001007188 | 843.8 | 567.2 | 0.01383 | 1.226 | | | |
| ARL17A | ADP ribosylation factor like GTPase 17A(ARL17A) | NM_016632 | 163.2 | 163.5 | 0.01388 | 1.274 | | | |
| | | XM_928111 | 92.5 | 99.8 | 0.01393 | 1.106 | | | |
| SLFN11 | schlafen family member 11(SLFN11) | NM_152270 | 102.7 | 95.7 | 0.01394 | 0.876 | | | |

| | | | | | | |
|---|---|---|---|---|---|---|
| SLFN11 | schlafen family member 11(SLFN11) | NM_152270 | 108.1 | 107.6 | 0.01394 | 0.876 |
| SMN1 | survival of motor neuron 1, telomeric(SMN1) | NM_000344 | 98 | 105.1 | 0.01394 | 1.181 |
| PTPN23 | protein tyrosine phosphatase, non-receptor type 23(PTPN23) | NM_015466 | 106.7 | 107.7 | 0.01397 | 1.191 |
| OFD1 | OFD1, centriole and centriolar satellite protein(OFD1) | NM_003611 | 105.5 | 108.9 | 0.01399 | 1.156 |
| HSCB | HscB mitochondrial iron-sulfur cluster cochaperone(HSCB) | NM_172002 | 95.2 | 92.9 | 0.01421 | 0.877 |
| NEDD8 | neural precursor cell expressed, developmentally down-regulated 8(NEDD8) | NM_006156 | 231.6 | 239.3 | 0.01424 | 0.742 |
| NEDD8 | neural precursor cell expressed, developmentally down-regulated 8(NEDD8) | NM_006156 | 1873.4 | 1679.1 | 0.01424 | 0.742 |
| ARID4B | AT-rich interaction domain 4B(ARID4B) | NM_016374 | 127.9 | 108.8 | 0.01428 | 1.674 |
| ARID4B | AT-rich interaction domain 4B(ARID4B) | NM_016374 | 210.3 | 153.7 | 0.01428 | 1.674 |
| ARID4B | AT-rich interaction domain 4B(ARID4B) | NM_016374 | 430.2 | 318.2 | 0.01428 | 1.674 |

| | | | | | | |
|---|---|---|---|---|---|---|
| ERO1B | endoplasmic reticulum oxidoreductase 1 beta(ERO1B) | NM_019891 | 108.7 | 92.1 | 0.01431 | 0.832 |
| RHBDL3 | rhomboid like 3(RHBDL3) | NM_138328 | 141.9 | 115.2 | 0.01436 | 0.808 |
| GON7 | GON7, KEOPS complex subunit homolog(GON7) | NM_032490 | 97.7 | 103.8 | 0.01446 | 1.223 |
| ANGPTL7 | angiopoietin like 7(ANGPTL7) | NM_021146 | 109.6 | 97.3 | 0.01447 | 0.873 |
| PHLDA1 | pleckstrin homology like domain family A member 1(PHLDA1) | NM_007350 | 99.5 | 105.4 | 0.01451 | 0.745 |
| PHLDA1 | pleckstrin homology like domain family A member 1(PHLDA1) | NM_007350 | 157.6 | 155.9 | 0.01451 | 0.745 |
| MOS | v-mos Moloney murine sarcoma viral oncogene homolog(MOS) | NM_005372 | 92.4 | 89.7 | 0.01457 | 0.899 |
| | | XM_001130102 | 122.6 | 117.7 | 0.01466 | 1.189 |
| CSRP2 | cysteine and glycine rich protein 2(CSRP2) | NM_001321 | 102.3 | 91.5 | 0.01466 | 0.903 |
| NECTIN3 | nectin cell adhesion molecule 3(NECTIN3) | NM_015480 | 145.6 | 166.7 | 0.0148 | 1.412 |
| NECTIN3 | nectin cell adhesion molecule 3(NECTIN3) | NM_015480 | 179.6 | 197.3 | 0.0148 | 1.412 |
| | | XM_001720754 | 96.1 | 102.4 | 0.01485 | 1.108 |

| | | | | | | |
|---|---|---|---|---|---|---|
| LOC105372672 | uncharacterized LOC105372672(LOC105372672) | AW661821 | 113.6 | 99.2 | 0.01499 | 0.821 |
| | | BX107471 | 92.6 | 90.4 | 0.01502 | 0.863 |
| KIF5C | kinesin family member 5C(KIF5C) | NM_004522 | 111.7 | 110.1 | 0.01506 | 0.881 |
| mri1 | methylthioribose-1-phosphate isomerase 1(MRI1) | NM_001031727 | 118.3 | 118.3 | 0.01512 | 1.33 |
| mri1 | methylthioribose-1-phosphate isomerase 1(MRI1) | NM_001031727 | 126.3 | 142.2 | 0.01512 | 1.33 |
| | | XR_039704 | 100.4 | 90.1 | 0.01516 | 0.84 |
| LENG8 | leukocyte receptor cluster member 8(LENG8) | NM_052925 | 125.4 | 100.3 | 0.01521 | 0.779 |
| FRS3 | fibroblast growth factor receptor substrate 3(FRS3) | NM_006653 | 187.5 | 189.9 | 0.01526 | 1.243 |
| MIR657 | microRNA 657(MIR657) | NR_030394 | 91.8 | 97.2 | 0.01527 | 1.278 |
| | | XM_001726567 | 139.6 | 137 | 0.01529 | 0.795 |
| C1QTNF1 | C1q and tumor necrosis factor related protein 1(C1QTNF1) | NM_198593 | 101 | 105.7 | 0.01536 | 1.14 |
| C1QTNF1 | C1q and tumor necrosis factor related protein 1(C1QTNF1) | NM_198594 | 139 | 133.4 | 0.01536 | 1.14 |

| | | | | | | |
|---|---|---|---|---|---|---|
| DAPL1 | death associated protein like 1(DAPL1) | NM_001017920 | 159.5 | 127.5 | 0.01541 | 0.71 |
| RBM6 | RNA binding motif protein 6(RBM6) | NM_005777 | 123.5 | 101.2 | 0.01542 | 0.864 |
| ARSB | arylsulfatase B(ARSB) | NM_000046 | 118.4 | 123.1 | 0.01544 | 1.227 |
| UBN2 | ubinuclein 2(UBN2) | NM_173569 | 153.5 | 130.9 | 0.01546 | 0.853 |
| UBN2 | ubinuclein 2(UBN2) | NM_173569 | 242.7 | 199.3 | 0.01546 | 0.853 |
| EPS8L3 | EPS8 like 3(EPS8L3) | NM_139053 | 91 | 83.7 | 0.01554 | 0.862 |
| MAPT | microtubule associated protein tau(MAPT) | NM_016834 | 148.3 | 128.8 | 0.01556 | 0.786 |
| AGFG2 | ArfGAP with FG repeats 2(AGFG2) | NM_006076 | 183.1 | 177.2 | 0.0156 | 1.351 |
| MIR571 | microRNA 571(MIR571) | NR_030297 | 105.3 | 106.7 | 0.01567 | 1.114 |
| SF3B1 | splicing factor 3b subunit 1(SF3B1) | NM_012433 | 148.3 | 148.7 | 0.01586 | 1.302 |
| SF3B1 | splicing factor 3b subunit 1(SF3B1) | NM_012433 | 636.6 | 434.5 | 0.01586 | 1.302 |
| GALE | UDP-galactose-4-epimerase(GALE) | NM_000403 | 98.9 | 102.7 | 0.0159 | 1.194 |
| RANBP3 | RAN binding protein 3(RANBP3) | NM_003624 | 101.5 | 97.4 | 0.01593 | 1.132 |
| RANBP3 | RAN binding protein 3(RANBP3) | NM_003624 | 102.5 | 105.1 | 0.01593 | 1.132 |
| UBE2D3 | ubiquitin conjugating enzyme E2 D3(UBE2D3) | NM_181890 | 1623.6 | 1372.6 | 0.01594 | 1.335 |

| | | | | | | |
|---|---|---|---|---|---|---|
| UBE2D3 | ubiquitin conjugating enzyme E2 D3(UBE2D3) | NM_181890 | 2032.9 | 1807.8 | 0.01594 | 1.335 |
| BPTF | bromodomain PHD finger transcription factor(BPTF) | NM_004459 | 93.9 | 96.4 | 0.01602 | 1.24 |
| FAM220A | family with sequence similarity 220 member A(FAM220A) | NM_001037163 | 104 | 93.3 | 0.01602 | 0.894 |
| LAIR1 | leukocyte associated immunoglobulin like receptor 1(LAIR1) | NM_021706 | 118 | 114.2 | 0.01603 | 1.234 |
| LAMA2 | laminin subunit alpha 2(LAMA2) | NM_000426 | 17263.4 | 17310.7 | 0.01603 | 1.234 |
| | | AI341386 | 101.5 | 119 | 0.01616 | 1.28 |
| TMEM217 | transmembrane protein 217(TMEM217) | NM_145316 | 99.8 | 106.4 | 0.01617 | 1.149 |
| TMEM217 | transmembrane protein 217(TMEM217) | NM_145316 | 142.5 | 140.2 | 0.01617 | 1.149 |
| PARP16 | poly(ADP-ribose) polymerase family member 16(PARP16) | NM_017851 | 104.5 | 99.1 | 0.01623 | 0.851 |
| TOP3A | topoisomerase (DNA) III alpha(TOP3A) | NM_004618 | 93.7 | 91.8 | 0.01623 | 0.799 |
| TOP3A | topoisomerase (DNA) III alpha(TOP3A) | NM_004618 | 108.4 | 96.4 | 0.01623 | 0.799 |
| BCL2L11 | BCL2 like 11(BCL2L11) | NM_006538 | 98.4 | 103.9 | 0.0163 | 1.193 |
| | | BX097731 | 101.5 | 107.1 | 0.01631 | 1.172 |
| | | CR619250 | 95.1 | 99.3 | 0.01637 | 1.159 |

| | | | | | | |
|---|---|---|---|---|---|---|
| CNRIP1 | cannabinoid receptor interacting protein 1(CNRIP1) | NM_015463 | 102.1 | 91.2 | 0.0164 | 0.87 |
| RBM41 | RNA binding motif protein 41(RBM41) | NM_018301 | 154.8 | 157.6 | 0.01644 | 1.28 |
| SAMD9 | sterile alpha motif domain containing 9(SAMD9) | NM_017654 | 91.4 | 87.8 | 0.01646 | 0.84 |
| ZNF467 | zinc finger protein 467(ZNF467) | NM_207336 | 185.3 | 202.1 | 0.01655 | 1.278 |
| DDX10 | DEAD-box helicase 10(DDX10) | NM_004398 | 101.9 | 87.7 | 0.0166 | 0.838 |
| LUC7L | LUC7 like(LUC7L) | NM_018032 | 160.1 | 144.7 | 0.01666 | 0.858 |
| POLR1B | RNA polymerase I subunit B(POLR1B) | NM_019014 | 103.4 | 92.7 | 0.01669 | 0.864 |
| LGALS12 | galectin 12(LGALS12) | NM_033101 | 100.6 | 105.3 | 0.01674 | 1.14 |
| cyb5r3 | cytochrome b5 reductase 3(CYB5R3) | NM_000398 | 113.2 | 106.6 | 0.01675 | 0.887 |
| FAM86JP | family with sequence similarity 86, member A pseudogene(FAM86JP) | NR_024250 | 96.9 | 102.5 | 0.01686 | 1.163 |
| DHX40 | DEAH-box helicase 40(DHX40) | NM_024612 | 117.9 | 110.1 | 0.01687 | 0.864 |
| DHX40 | DEAH-box helicase 40(DHX40) | NM_024612 | 139 | 137.6 | 0.01687 | 0.864 |
| DHX40 | DEAH-box helicase 40(DHX40) | NM_024612 | 169.3 | 171.4 | 0.01687 | 0.864 |

| | | | | | | |
|---|---|---|---|---|---|---|
| EIF2S2 | eukaryotic translation initiation factor 2 subunit beta(EIF2S2) | NM_003908 | 223.4 | 241 | 0.0169 | 1.26 |
| XKR9 | XK related 9(XKR9) | NM_001011720 | 109.2 | 93.9 | 0.01692 | 0.874 |
| PDZD8 | PDZ domain containing 8(PDZD8) | NM_173791 | 285.2 | 267.9 | 0.01699 | 1.213 |
| NFX1 | nuclear transcription factor, X-box binding 1(NFX1) | NM_147134 | 107.6 | 102.5 | 0.017 | 1.131 |
| PHF14 | PHD finger protein 14(PHF14) | NM_014660 | 100.5 | 101.6 | 0.01705 | 1.088 |
| ZNF658B | zinc finger protein 658B (pseudogene)(ZNF658B) | NM_001032297 | 107.5 | 115.7 | 0.01713 | 1.186 |
| | | XM_001721880 | 101.2 | 98.2 | 0.0172 | 1.153 |
| | | XM_001721880 | 177.7 | 157.5 | 0.0172 | 1.153 |
| FBXW4 | F-box and WD repeat domain containing 4(FBXW4) | NM_022039 | 111.1 | 114.8 | 0.01739 | 1.195 |
| Efcab13 | EF-hand calcium binding domain 13(EFCAB13) | NM_152347 | 96.7 | 96.1 | 0.01742 | 1.122 |
| LILRB3 | leukocyte immunoglobulin like receptor B3(LILRB3) | NM_006864 | 129.4 | 93 | 0.01748 | 1.248 |
| LILRB4 | leukocyte immunoglobulin like receptor B4(LILRB4) | NM_001081438 | 14279.9 | 13030.4 | 0.01748 | 1.248 |

| | | | | | | |
|---|---|---|---|---|---|---|
| RNA28S5 | RNA, 28S ribosomal 5(RNA28S5) | NR_003287 | 371.8 | 466.3 | 0.01766 | 2.305 |
| RNA28S5 | RNA, 28S ribosomal 5(RNA28S5) | NR_003287 | 769.5 | 878.2 | 0.01766 | 2.305 |
| STRN | striatin(STRN) | NM_003162 | 118.2 | 110.9 | 0.01769 | 0.808 |
| | | AI239861 | 98.5 | 102.5 | 0.01771 | 1.116 |
| CSNK1D | casein kinase 1 delta(CSNK1D) | NM_139062 | 256 | 242.1 | 0.01774 | 1.437 |
| LINC01018 | long intergenic non-protein coding RNA 1018(LINC01018) | NR_024424 | 106.9 | 97.3 | 0.01782 | 1.181 |
| LINC01011 | long intergenic non-protein coding RNA 1011(LINC01011) | NM_207495 | 99.4 | 104.9 | 0.01782 | 1.181 |
| PSMD3 | proteasome 26S subunit, non-ATPase 3(PSMD3) | NM_002809 | 106.3 | 90.9 | 0.01784 | 0.866 |
| TMEM80 | transmembrane protein 80(TMEM80) | NM_174940 | 135.9 | 123.3 | 0.01799 | 0.722 |
| COG4 | component of oligomeric golgi complex 4(COG4) | NM_015386 | 146.6 | 122 | 0.01803 | 0.77 |
| COG3 | component of oligomeric golgi complex 3(COG3) | NM_031431 | 137.6 | 123.3 | 0.01803 | 0.77 |
| | | XM_939521 | 105.4 | 106.4 | 0.01806 | 1.097 |
| | | DR980387 | 89.4 | 98.9 | 0.01823 | 1.166 |

| | | | | | | |
|---|---|---|---|---|---|---|
| GNB1L | G protein subunit beta 1 like(GNB1L) | NM_053004 | 100.1 | 93.4 | 0.01826 | 1.126 |
| | | XM_930349 | 85.2 | 82 | 0.0183 | 0.865 |
| | | XM_936804 | 109.4 | 104.4 | 0.01837 | 0.78 |
| ALOX12B | arachidonate 12-lipoxygenase, 12R type(ALOX12B) | NM_001139 | 102.3 | 103 | 0.01837 | 1.172 |
| ZNF654 | zinc finger protein 654(ZNF654) | NM_018293 | 94.7 | 96.7 | 0.01839 | 1.158 |
| ZNF654 | zinc finger protein 654(ZNF654) | NM_018293 | 100 | 93.3 | 0.01839 | 1.158 |
| ZNF654 | zinc finger protein 654(ZNF654) | NM_018293 | 101.7 | 99.2 | 0.01839 | 1.158 |
| ZNF654 | zinc finger protein 654(ZNF654) | NM_018293 | 105.3 | 110.3 | 0.01839 | 1.158 |
| | | XM_001724542 | 109.3 | 131.8 | 0.0184 | 1.422 |
| MIR939 | microRNA 939(MIR939) | NR_030635 | 116.6 | 110.5 | 0.01851 | 0.874 |
| | | XR_017667 | 9770 | 7036.8 | 0.01851 | 0.742 |
| E2F1 | E2F transcription factor 1(E2F1) | NM_005225 | 171.1 | 168.6 | 0.01852 | 1.461 |
| GADD45A | growth arrest and DNA damage inducible alpha(GADD45A) | NM_001924 | 104.9 | 99.7 | 0.01853 | 0.763 |
| ITGB2 | integrin subunit beta 2(ITGB2) | NM_000211 | 708.1 | 576.7 | 0.01855 | 0.69 |

| | | | | | | | | | |
|---|---|---|---|---|---|---|---|---|---|
| CCL19 | C-C motif chemokine ligand 19(CCL19) | NM_006274 | 110.3 | 112.8 | 0.01867 | 1.212 | NM_017196 | 14.73 | 0.69 |
| OR1K1 | olfactory receptor family 1 subfamily K member 1(OR1K1) | NM_080859 | 100 | 94.1 | 0.01873 | 0.87 | NM_017196 | 14.73 | 0.69 |
| | | XM_001130104 | 102 | 95.8 | 0.01878 | 0.881 | | | |
| | | XM_001126202 | 194.1 | 183 | 0.01883 | 1.428 | NM_017196 | 14.73 | 0.69 |
| | | XM_929153 | 93.8 | 96.7 | 0.0189 | 0.859 | | | |
| FANCE | Fanconi anemia complementation group E(FANCE) | NM_021922 | 177 | 154.6 | 0.01893 | 1.235 | | | |
| | | XM_944342 | 106.9 | 97.6 | 0.01896 | 0.875 | | | |
| VARS | valyl-tRNA synthetase(VARS) | NM_006295 | 181.3 | 149 | 0.01902 | 0.727 | NM_031356 | 5.43 | 24.53 |
| | | XM_001127768 | 99.9 | 104.9 | 0.01903 | 1.162 | NM_031356 | 5.43 | 24.53 |
| | | XM_001725366 | 110.4 | 104.2 | 0.01904 | 0.815 | | | |
| DTX2 | deltex E3 ubiquitin ligase 2(DTX2) | NM_020892 | 109.1 | 113.2 | 0.01918 | 1.159 | | | |
| DTX3 | deltex E3 ubiquitin ligase 3(DTX3) | NM_178502 | 438 | 382.6 | 0.01918 | 1.159 | | | |
| dhx37 | DEAH-box helicase 37(DHX37) | NM_032656 | 133.3 | 112.5 | 0.0192 | 0.836 | | | |
| | | XM_940292 | 97 | 93.7 | 0.01925 | 0.888 | | | |

| | | | | | | | | | |
|---|---|---|---|---|---|---|---|---|---|
| SLTM | SAFB like transcription modulator(SLTM) | NM_024755 | 132.4 | 131.2 | 0.01929 | 1.268 | | | |
| SLTM | SAFB like transcription modulator(SLTM) | NM_024755 | 342.5 | 294.3 | 0.01929 | 1.268 | | | |
| FIGN | fidgetin, microtubule severing factor(FIGN) | NM_018086 | 93.3 | 99.3 | 0.0193 | 1.132 | | | |
| slx4ip | SLX4 interacting protein(SLX4IP) | NM_001009608 | 113.6 | 112 | 0.01934 | 1.119 | | | |
| | | XM_938977 | 116.5 | 111.7 | 0.01935 | 1.138 | | | |
| SLC25A20 | solute carrier family 25 member 20(SLC25A20) | NM_000387 | 95.2 | 103.1 | 0.0194 | 1.43 | | | |
| SLC25A20 | solute carrier family 25 member 20(SLC25A20) | NM_000387 | 212.9 | 290.9 | 0.0194 | 1.43 | | | |
| TRAF1 | TNF receptor associated factor 1(TRAF1) | NM_005658 | 104.5 | 114.2 | 0.01954 | 1.246 | | | |
| ABTB1 | ankyrin repeat and BTB domain containing 1(ABTB1) | NM_032548 | 114.2 | 97.5 | 0.01957 | 0.843 | | | |
| CHAD | chondroadherin(CHAD) | NM_001267 | 96.4 | 91.9 | 0.01958 | 0.851 | NM_017135 | 41.45 | 2.88 |
| ZFP82 | ZFP82 zinc finger protein(ZFP82) | NM_133466 | 115.8 | 106.4 | 0.01969 | 0.875 | NM_017135 | 41.45 | 2.88 |
| | | XM_931238 | 108.5 | 104.2 | 0.01984 | 0.719 | NM_017135 | 41.45 | 2.88 |
| SHARPIN | SHANK associated RH domain interactor(SHARPIN) | NM_030974 | 116.9 | 108.1 | 0.01985 | 0.852 | NM_017135 | 41.45 | 2.88 |
| CALB2 | calbindin 2(CALB2) | NM_007088 | 103.1 | 113.8 | 0.01986 | 1.279 | NM_017135 | 41.45 | 2.88 |

| | | | | | | | | | |
|---|---|---|---|---|---|---|---|---|---|
| | | XM_944717 | 98.9 | 93.9 | 0.01987 | 0.858 | | | |
| | | AW168583 | 90.7 | 97.5 | 0.02005 | 1.164 | | | |
| | | XM_928447 | 97.7 | 102.2 | 0.02008 | 1.203 | | | |
| | | XM_001719479 | 107.4 | 105.1 | 0.0201 | 1.136 | | | |
| SERPINB7 | serpin family B member 7(SERPINB7) | NM_003784 | 102.5 | 100 | 0.02011 | 0.85 | | | |
| SERPINB7 | serpin family B member 7(SERPINB7) | NM_003784 | 123.2 | 109.9 | 0.02011 | 0.85 | | | |
| | | XM_931877 | 98.6 | 103.2 | 0.02017 | 1.102 | | | |
| | | XR_039881 | 115.4 | 102.6 | 0.02022 | 1.218 | NM_001033653 | 43.46 | 1.56 |
| ARMCX3 | armadillo repeat containing, X-linked 3(ARMCX3) | NM_177947 | 210.1 | 181.4 | 0.02027 | 0.708 | | | |
| ARMCX3 | armadillo repeat containing, X-linked 3(ARMCX3) | NM_177948 | 358.4 | 354.8 | 0.02027 | 0.708 | | | |
| SYNE1 | spectrin repeat containing nuclear envelope protein 1(SYNE1) | NM_033071 | 86.7 | 87.4 | 0.02027 | 0.74 | | | |
| SEC63 | SEC63 homolog, protein translocation regulator(SEC63) | NM_007214 | 103.2 | 112.3 | 0.02028 | 1.168 | | | |

| | | | | | | |
|---|---|---|---|---|---|---|
| TMEM57 | transmembrane protein 57(TMEM57) | NM_018202 | 302.2 | 259 | 0.02049 | 1.361 |
| | | XR_019270 | 99.9 | 97.2 | 0.02052 | 0.878 |
| PA2G4 | proliferation-associated 2G4(PA2G4) | NM_006191 | 118.1 | 124.9 | 0.02056 | 1.19 |
| | | AI056317 | 98.3 | 91.3 | 0.02066 | 0.898 |
| | | XR_041761 | 99 | 104 | 0.02066 | 1.204 |
| NOM1 | nucleolar protein with MIF4G domain 1(NOM1) | NM_138400 | 99.5 | 90.9 | 0.02067 | 0.767 |
| HS6ST2 | heparan sulfate 6-O-sulfotransferase 2(HS6ST2) | NM_147175 | 107.9 | 109.4 | 0.02068 | 1.102 |
| SRSF3 | serine and arginine rich splicing factor 3(SRSF3) | NM_003017 | 140.1 | 138.1 | 0.02078 | 0.742 |
| BTBD6 | BTB domain containing 6(BTBD6) | NM_033271 | 124.3 | 116.8 | 0.02099 | 0.874 |
| BTBD3 | BTB domain containing 3(BTBD3) | NM_014962 | 92.5 | 88.5 | 0.02099 | 0.874 |
| EP300 | E1A binding protein p300(EP300) | NM_001429 | 104.5 | 106.6 | 0.02102 | 1.149 |
| GSTTP2 | glutathione S-transferase theta pseudogene 2(GSTTP2) | NR_003082 | 176.8 | 173.7 | 0.0211 | 1.158 |

| | | | | | | | | | |
|---|---|---|---|---|---|---|---|---|---|
| GSTT2B | glutathione S-transferase theta 2B (gene/pseudogene)(GSTT2B) | NM_001080843 | 92.2 | 106.7 | 0.0211 | 1.158 | | | |
| TM7SF3 | transmembrane 7 superfamily member 3(TM7SF3) | NM_016551 | 131.8 | 120.2 | 0.02121 | 0.767 | | | |
| RBM10 | RNA binding motif protein 10(RBM10) | NM_152856 | 311.5 | 312.6 | 0.02124 | 1.256 | | | |
| | | XM_001713713 | 96.3 | 92.3 | 0.02127 | 0.907 | | | |
| OBSL1 | obscurin like 1(OBSL1) | NM_015311 | 104.6 | 114.2 | 0.02128 | 1.22 | | | |
| cemip | cell migration inducing hyaluronan binding protein(CEMIP) | NM_018689 | 99.3 | 96.4 | 0.02131 | 0.883 | | | |
| CCS | copper chaperone for superoxide dismutase(CCS) | NM_005125 | 104.2 | 111.5 | 0.02143 | 1.276 | | | |
| CCSER2 | coiled-coil serine rich protein 2(CCSER2) | NM_018999 | 117.4 | 111.4 | 0.02143 | 1.276 | NM_134407 | 50.47 | 11.16 |
| CSN1S1 | casein alpha s1(CSN1S1) | NM_001890 | 98.7 | 102.3 | 0.02143 | 1.131 | | | |
| FAHD1 | fumarylacetoacetate hydrolase domain containing 1(FAHD1) | NM_001018104 | 318.4 | 348.8 | 0.02146 | 1.212 | | | |
| ZFYVE28 | zinc finger FYVE-type containing 28(ZFYVE28) | NM_020972 | 92.3 | 96.8 | 0.02152 | 1.133 | | | |
| TNIP2 | TNFAIP3 interacting protein 2(TNIP2) | NM_024309 | 108.2 | 102.8 | 0.02163 | 0.884 | | | |

| | | | | | | |
|---|---|---|---|---|---|---|
| OR11H1 | olfactory receptor family 11 subfamily H member 1(OR11H1) | NM_001005239 | 98.6 | 95.5 | 0.02163 | 0.796 |
| SPTBN5 | spectrin beta, non-erythrocytic 5(SPTBN5) | NM_016642 | 131.5 | 123.8 | 0.02163 | 0.785 |
| | | XR_016400 | 98.2 | 93.9 | 0.02165 | 0.884 |
| | | XR_016218 | 114.4 | 112.6 | 0.0218 | 0.428 |
| SKAP2 | src kinase associated phosphoprotein 2(SKAP2) | NM_003930 | 95 | 93.7 | 0.02188 | 0.803 |
| SKAP2 | src kinase associated phosphoprotein 2(SKAP2) | NM_003930 | 210 | 162.9 | 0.02188 | 0.803 |
| | | XM_930512 | 95.6 | 100.7 | 0.02189 | 1.191 |
| | | XR_016302 | 83.5 | 92.6 | 0.02199 | 1.169 |
| RAMP3 | receptor activity modifying protein 3(RAMP3) | NM_005856 | 135.4 | 126.1 | 0.02208 | 1.338 |
| RAMP3 | receptor activity modifying protein 3(RAMP3) | NM_005856 | 155.8 | 125.8 | 0.02208 | 1.338 |
| adgra3 | adhesion G protein-coupled receptor A3(ADGRA3) | NM_145290 | 132.7 | 137.6 | 0.02213 | 1.561 |
| BBS2 | Bardet-Biedl syndrome 2(BBS2) | NM_031885 | 314.5 | 330.2 | 0.02224 | 0.766 |

| | | | | | | | | | |
|---|---|---|---|---|---|---|---|---|---|
| BBS4 | Bardet-Biedl syndrome 4(BBS4) | NM_033028 | 548.9 | 463.4 | 0.02224 | 0.766 | | | |
| PPIL2 | peptidylprolyl isomerase like 2(PPIL2) | NM_014337 | 106.2 | 112.2 | 0.02228 | 1.178 | | | |
| PPIL2 | peptidylprolyl isomerase like 2(PPIL2) | NM_014337 | 147.3 | 136.1 | 0.02228 | 1.178 | | | |
| | | XM_940079 | 121 | 108.1 | 0.02229 | 0.784 | NM_001033706 | 2.42 | 10.86 |
| C9orf85 | chromosome 9 open reading frame 85(C9orf85) | NM_182505 | 95.9 | 100.4 | 0.02229 | 1.155 | | | |
| TRAPPC4 | trafficking protein particle complex 4(TRAPPC4) | NM_016146 | 351.4 | 304.6 | 0.0223 | 0.842 | | | |
| PTK2 | protein tyrosine kinase 2(PTK2) | NM_153831 | 103.7 | 106.5 | 0.0223 | 1.088 | | | |
| | | XM_937184 | 95.9 | 98.1 | 0.02234 | 1.158 | | | |
| | | XM_0017133911 | 97 | 98.4 | 0.0224 | 1.166 | | | |
| MS4A6A | membrane spanning 4-domains A6A(MS4A6A) | NM_022349 | 104.3 | 92 | 0.02245 | 0.891 | NM_153300 | 0.14 | 61.17 |
| | | XM_0017222445 | 109.3 | 92.3 | 0.02252 | 0.777 | | | |
| setmar | SET domain and mariner transposase fusion gene(SETMAR) | NM_006515 | 184.2 | 161.7 | 0.02255 | 0.763 | NM_032416 | 108.51 | 1.87 |

| | | | | | | |
|---|---|---|---|---|---|---|
| KCNJ14 | potassium voltage-gated channel subfamily J member 14(KCNJ14) | NM_170720 | 97.9 | 101.8 | 0.02263 | 1.11 |
| KRTAP12-1 | keratin associated protein 12-1(KRTAP12-1) | NM_181686 | 101.8 | 102.5 | 0.02264 | 1.162 |
| VPS13A | vacuolar protein sorting 13 homolog A(VPS13A) | NM_001018038 | 95.7 | 102.8 | 0.0227 | 1.183 |
| VPS13A | vacuolar protein sorting 13 homolog A(VPS13A) | NM_001018038 | 100.6 | 97.6 | 0.0227 | 1.183 |
| NUDT22 | nudix hydrolase 22(NUDT22) | NM_032344 | 148.6 | 154.3 | 0.02273 | 1.244 |
| KCNJ4 | potassium voltage-gated channel subfamily J member 4(KCNJ4) | NM_152868 | 96.8 | 100.4 | 0.02275 | 1.124 |
| BICDL2 | BICD family like cargo adaptor 2(BICDL2) | NM_001103175 | 209.9 | 205.9 | 0.0228 | 1.275 |
| | | XM_001723741 | 96.8 | 106.1 | 0.02288 | 1.166 |
| RPLP2 | ribosomal protein lateral stalk subunit P2(RPLP2) | NM_001004 | 9850.2 | 8202.3 | 0.02299 | 1.4 |
| BCORL1 | BCL6 corepressor-like 1(BCORL1) | NM_021946 | 250.5 | 165.9 | 0.02302 | 0.737 |
| PSMD5 | proteasome 26S subunit, non-ATPase 5(PSMD5) | NM_005047 | 156.6 | 143.8 | 0.02303 | 0.834 |
| TTLL3 | tubulin tyrosine ligase like 3(TTLL3) | NM_015644 | 102.6 | 98.6 | 0.02307 | 0.901 |

| | | | | | | | | | |
|---|---|---|---|---|---|---|---|---|---|
| TTLL3 | tubulin tyrosine ligase like 3(TTLL3) | NM_015644 | 152.3 | 116 | 0.02307 | 0.901 | | | |
| TAGLN | transgelin(TAGLN) | NM_003186 | 388.2 | 296.4 | 0.02309 | 0.565 | | | |
| TAGLN | transgelin(TAGLN) | NM_003186 | 1629.7 | 1263 | 0.02309 | 0.565 | NM_001191088 | 10.07 | 0.03 |
| ZNF419 | zinc finger protein 419(ZNF419) | NM_001098495 | 117.2 | 122.7 | 0.02311 | 1.195 | NM_001191088 | 10.07 | 0.03 |
| TBP | TATA-box binding protein(TBP) | NM_003194 | 176.5 | 138.7 | 0.02312 | 0.825 | NM_001191088 | 10.07 | 0.03 |
| | | XM_944734 | 105.6 | 105.9 | 0.02315 | 1.137 | | | |
| SDR16C5 | short chain dehydrogenase/reductase family 16C, member 5(SDR16C5) | NM_138969 | 100.6 | 123 | 0.02318 | 1.399 | | | |
| MANEA | mannosidase endo-alpha(MANEA) | NM_024641 | 284.5 | 258.2 | 0.0232 | 1.18 | | | |
| MANBAL | mannosidase beta like(MANBAL) | NM_001003897 | 93.3 | 98 | 0.0232 | 1.18 | NM_012496 | 1095.39 | 8.76 |
| TMUB2 | transmembrane and ubiquitin like domain containing 2(TMUB2) | NM_001076674 | 96.8 | 102.1 | 0.0232 | 1.093 | | | |
| TAF8 | TATA-box binding protein associated factor 8(TAF8) | NM_138572 | 106 | 117.7 | 0.02321 | 1.203 | | | |

| | | | | | | |
|---|---|---|---|---|---|---|
| TIMMDC 1 | translocase of inner mitochondrial membrane domain containing 1(TIMMDC1) | NM_016589 | 261.6 | 245.6 | 0.0234 | 1.756 |
| OACYLP | O-acyltransferase like, pseudogene(OACYLP) | NR_024021 | 95.9 | 92.1 | 0.02342 | 0.881 |
| ADAMTS 17 | ADAM metallopeptidase with thrombospondin type 1 motif 17(ADAMTS17) | NM_139057 | 100.2 | 101.2 | 0.02354 | 1.182 |
| FOXI3 | forkhead box I3(FOXI3) | NM_001135649 | 125 | 122.7 | 0.02357 | 1.164 |
| FOXI1 | forkhead box I1(FOXI1) | NM_012188 | 95.4 | 100.1 | 0.02357 | 1.164 |
| | | BM674906 | 97.4 | 104.6 | 0.02358 | 1.129 |
| ZNF839 | zinc finger protein 839(ZNF839) | NM_018335 | 115.6 | 122.3 | 0.0236 | 1.15 |
| | | XM_936486 | 85.9 | 93.8 | 0.02362 | 1.19 |
| RAMP2 | receptor activity modifying protein 2(RAMP2) | NM_005854 | 120.2 | 108.4 | 0.02364 | 1.231 |
| RAMP2 | receptor activity modifying protein 2(RAMP2) | NM_005854 | 122.1 | 133.9 | 0.02364 | 1.231 |
| MTMR1 | myotubularin related protein 1(MTMR1) | NM_003828 | 103.1 | 96.4 | 0.02366 | 0.871 |
| ANXA6 | annexin A6(ANXA6) | NM_001155 | 97 | 97.9 | 0.02369 | 0.603 |

| | | | | | | |
|---|---|---|---|---|---|---|
| | | XM_001130122 | 110.9 | 110.4 | 0.02369 | 0.899 |
| | | XM_001130122 | 119 | 109.5 | 0.02369 | 0.899 |
| | | XM_498608 | 95.9 | 94 | 0.0237 | 0.897 |
| UQCR11 | ubiquinol-cytochrome c reductase, complex III subunit XI(UQCR11) | NM_006830 | 102.6 | 110.8 | 0.02379 | 1.137 |
| KRTAP13-3 | keratin associated protein 13-3(KRTAP13-3) | NM_181622 | 106.2 | 101.1 | 0.02383 | 1.117 |
| FAM76B | family with sequence similarity 76 member B(FAM76B) | NM_144664 | 101.8 | 111.2 | 0.02384 | 1.21 |
| NAPEPLD | N-acyl phosphatidylethanolamine phospholipase D(NAPEPLD) | NM_198990 | 109.8 | 105.8 | 0.02406 | 0.842 |
| NAPEPLD | N-acyl phosphatidylethanolamine phospholipase D(NAPEPLD) | NM_198990 | 136.7 | 129.8 | 0.02406 | 0.842 |
| STAR | steroidogenic acute regulatory protein(STAR) | NM_001007243 | 107 | 91.7 | 0.02412 | 0.732 |
| WDR4 | WD repeat domain 4(WDR4) | NM_033661 | 100.8 | 100.2 | 0.02415 | 1.118 |
| BPIFB3 | BPI fold containing family B member 3(BPIFB3) | NM_182658 | 105 | 105.1 | 0.02418 | 1.174 |

| | | | | | | |
|---|---|---|---|---|---|---|
| | | BX102609 | 97.7 | 94.7 | 0.02426 | 0.88 |
| PRUNE2 | prune homolog 2(PRUNE2) | NM_138818 | 101 | 109.8 | 0.02429 | 1.351 |
| PRUNE2 | prune homolog 2(PRUNE2) | NM_138818 | 101.6 | 102.3 | 0.02429 | 1.351 |
| TRAPPC2 | trafficking protein particle complex 2(TRAPPC2) | NM_014563 | 100.3 | 90.3 | 0.02438 | 0.846 |
| TRAPPC2 | trafficking protein particle complex 2(TRAPPC2) | NM_014563 | 204.2 | 161.1 | 0.02438 | 0.846 |
| MLEC | malectin(MLEC) | NM_014730 | 226.4 | 205.3 | 0.02445 | 1.294 |
| MLEC | malectin(MLEC) | NM_014730 | 530.9 | 526.3 | 0.02445 | 1.294 |
| RNF215 | ring finger protein 215(RNF215) | NM_001017981 | 101.8 | 89.7 | 0.0245 | 0.865 |
| | | XM_933836 | 104.9 | 105.5 | 0.02463 | 1.098 |
| GALNT10 | polypeptide N-acetylgalactosaminyltransferase 10(GALNT10) | NM_017540 | 144 | 142.8 | 0.02463 | 1.372 |
| PCGF6 | polycomb group ring finger 6(PCGF6) | NM_001011663 | 94.3 | 91.5 | 0.02464 | 0.859 |
| KLHDC8B | kelch domain containing 8B(KLHDC8B) | NM_173546 | 99 | 101.9 | 0.02474 | 1.127 |
| | | XM_930350 | 121.4 | 116.3 | 0.02484 | 1.183 |

| | | | | | | | | | |
|---|---|---|---|---|---|---|---|---|---|
| KANK3 | KN motif and ankyrin repeat domains 3(KANK3) | NM_198471 | 92 | 96.5 | 0.02484 | 1.131 | | | |
| BACH2 | BTB domain and CNC homolog 2(BACH2) | NM_021813 | 95 | 99.7 | 0.02486 | 1.145 | | | |
| BACH2 | BTB domain and CNC homolog 2(BACH2) | NM_021813 | 110.5 | 88.8 | 0.02486 | 1.145 | | | |
| BACH2 | BTB domain and CNC homolog 2(BACH2) | NM_021813 | 125.9 | 97.7 | 0.02486 | 1.145 | | | |
| HERPUD2 | HERPUD family member 2(HERPUD2) | NM_022373 | 100.6 | 106.9 | 0.02489 | 1.125 | NM_013059 | 21.02 | 0.11 |
| HES1 | hes family bHLH transcription factor 1(HES1) | NM_005524 | 110.3 | 105.6 | 0.02489 | 1.125 | | | |
| MTIF2 | mitochondrial translational initiation factor 2(MTIF2) | NM_001005369 | 724.5 | 656.1 | 0.02491 | 1.3 | | | |
| TUSC3 | tumor suppressor candidate 3(TUSC3) | NM_006765 | 92.7 | 97 | 0.02493 | 1.19 | | | |
| TUSC3 | tumor suppressor candidate 3(TUSC3) | NM_006765 | 162.1 | 190.1 | 0.02493 | 1.19 | | | |
| | | XM_001715936 | 109.1 | 107.4 | 0.02501 | 1.152 | | | |
| DZANK1 | double zinc ribbon and ankyrin repeat domains 1(DZANK1) | NM_001099407 | 99.6 | 91.5 | 0.02504 | 0.83 | | | |
| ZNF471 | zinc finger protein 471(ZNF471) | NM_020813 | 113.7 | 117.6 | 0.02504 | 1.249 | | | |
| ZNF471 | zinc finger protein 471(ZNF471) | NM_020813 | 131.7 | 127.7 | 0.02504 | 1.249 | | | |

| | | | | | | | | | |
|---|---|---|---|---|---|---|---|---|---|
| | | XM_001132307 | 95.6 | 94.6 | 0.02512 | 0.873 | | | |
| mri1 | methylthioribose-1-phosphate isomerase 1(MRI1) | NM_032285 | 138.6 | 130.6 | 0.02515 | 1.226 | | | |
| | | XR_019551 | 118.8 | 121.1 | 0.02526 | 1.225 | | | |
| ABAT | 4-aminobutyrate aminotransferase(ABAT) | NM_000663 | 114.2 | 120.3 | 0.02539 | 1.135 | | | |
| AATK | apoptosis associated tyrosine kinase(AATK) | NM_001080395 | 107.4 | 106.4 | 0.02539 | 1.135 | | | |
| THUMPD1 | THUMP domain containing 1(THUMPD1) | NM_017736 | 141.6 | 149.7 | 0.02554 | 1.191 | NM_012816 | 19.78 | 2.28 |
| LASP1 | LIM and SH3 protein 1(LASP1) | NM_006148 | 94.5 | 102 | 0.02565 | 1.111 | NM_012816 | 19.78 | 2.28 |
| | | XM_0017177976 | 101.7 | 107.2 | 0.0257 | 1.106 | | | |
| WWOX | WW domain containing oxidoreductase(WWOX) | NM_130791 | 117.4 | 101.4 | 0.02601 | 0.884 | | | |
| Mt1f | metallothionein 1F(MT1F) | NM_005949 | 107.2 | 107.1 | 0.02606 | 1.153 | | | |
| POTEE | POTE ankyrin domain family member E(POTEE) | NM_001083538 | 105.2 | 98.8 | 0.02607 | 0.867 | | | |
| FLNC | filamin C(FLNC) | NM_001458 | 102.5 | 88.9 | 0.02607 | 0.81 | | | |
| SNX29 | sorting nexin 29(SNX29) | NM_001009607 | 122.6 | 117.5 | 0.02623 | 1.306 | | | |

| | | | | | | |
|---|---|---|---|---|---|---|
| GGT1 | gamma-glutamyltransferase 1(GGT1) | NM_001032364 | 118.8 | 102.1 | 0.02633 | 0.846 |
| HES5 | hes family bHLH transcription factor 5(HES5) | NM_001010926 | 101.4 | 107.2 | 0.02635 | 1.16 |
| EBAG9 | estrogen receptor binding site associated, antigen, 9(EBAG9) | NM_198120 | 116.1 | 136.2 | 0.02635 | 1.35 |
| | | XR_037520 | 108.7 | 105 | 0.02637 | 0.903 |
| MED7 | mediator complex subunit 7(MED7) | NM_001100816 | 133.3 | 114.7 | 0.02641 | 0.797 |
| MED7 | mediator complex subunit 7(MED7) | NM_001100816 | 138.8 | 130.1 | 0.02641 | 0.797 |
| | | BX093647 | 97.4 | 103.2 | 0.02646 | 1.059 |
| | | BX111229 | 109.1 | 116.9 | 0.02648 | 1.194 |
| | | BX109868 | 124.8 | 118.1 | 0.02649 | 1.126 |
| RINT1 | RAD50 interactor 1(RINT1) | NM_021930 | 124.7 | 125.8 | 0.02653 | 1.177 |
| | | XR_019645 | 115.4 | 101.5 | 0.02655 | 0.864 |
| SLC16A10 | solute carrier family 16 member 10(SLC16A10) | NM_018593 | 238.6 | 234.4 | 0.02665 | 1.749 |
| | | XM_944489 | 251.6 | 275 | 0.02668 | 1.242 |
| CAPZA1 | capping actin protein of muscle Z-line alpha subunit 1(CAPZA1) | NM_006135 | 181.8 | 174.9 | 0.02672 | 1.272 |

| | | | | | | | | | |
|---|---|---|---|---|---|---|---|---|---|
| AASDHPPT | aminoadipate-semialdehyde dehydrogenase-phosphopantetheinyl transferase(AASDHPPT) | NM_015423 | 96.1 | 93.3 | 0.02681 | 0.881 | NM_031544 | 4.79 | 38.59 |
| PRSS3P2 | protease, serine 3 pseudogene 2(PRSS3P2) | NR_001296 | 100.1 | 93.1 | 0.02683 | 1.142 | NM_031544 | 4.79 | 38.59 |
| C15orf56 | chromosome 15 open reading frame 56(C15orf56) | NM_001039905 | 98.3 | 106.9 | 0.02686 | 1.146 | | | |
| ST13 | suppression of tumorigenicity 13 (colon carcinoma) (Hsp70 interacting protein)(ST13) | NM_003932 | 217.8 | 232.6 | 0.02688 | 1.552 | | | |
| Snora26 | small nucleolar RNA, H/ACA box 26(SNORA26) | NR_003016 | 113.5 | 98.3 | 0.02695 | 0.84 | | | |
| PSMD7 | proteasome 26S subunit, non-ATPase 7(PSMD7) | NM_002811 | 152.8 | 133.6 | 0.02697 | 0.822 | | | |
| MFSD14B | major facilitator superfamily domain containing 14B(MFSD14B) | NM_032558 | 95.4 | 95.9 | 0.02706 | 0.864 | | | |
| MFSD14B | major facilitator superfamily domain containing 14B(MFSD14B) | NM_032558 | 101.3 | 95.5 | 0.02706 | 0.864 | | | |

| | | | | | | |
|---|---|---|---|---|---|---|
| MFSD14B | major facilitator superfamily domain containing 14B(MFSD14B) | NM_032558 | 101.7 | 94.1 | 0.02706 | 0.864 |
| MFSD14B | major facilitator superfamily domain containing 14B(MFSD14B) | NM_032558 | 103 | 98.8 | 0.02706 | 0.864 |
| JADE3 | jade family PHD finger 3(JADE3) | NM_001077445 | 150.5 | 135.7 | 0.02712 | 0.817 |
| SRCAP | Snf2 related CREBBP activator protein(SRCAP) | NM_006662 | 95.9 | 88.6 | 0.02716 | 0.886 |
| | | AL602970 | 93.8 | 101.5 | 0.02718 | 1.143 |
| | | XM_001716887 | 121.9 | 102.5 | 0.02723 | 0.798 |
| folr1 | folate receptor 1(FOLR1) | NM_016725 | 124.1 | 103.5 | 0.02737 | 0.781 |
| NAE1 | NEDD8 activating enzyme E1 subunit 1(NAE1) | NM_001018160 | 130.8 | 139.5 | 0.02747 | 0.661 |
| NAE1 | NEDD8 activating enzyme E1 subunit 1(NAE1) | NM_001018160 | 248.3 | 193.4 | 0.02747 | 0.661 |
| LOC284009 | uncharacterized LOC284009(LOC284009) | NM_001025459 | 110.2 | 117 | 0.02751 | 1.164 |
| FAM99A | family with sequence similarity 99 member A (non-protein coding)(FAM99A) | NM_001014374 | 102.9 | 96 | 0.02757 | 1.139 |

| | | | | | | |
|---|---|---|---|---|---|---|
| | | XM_001715669 | 90.3 | 96.4 | 0.02767 | 1.187 |
| CD22 | CD22 molecule(CD22) | NM_001771 | 94.4 | 98.4 | 0.02776 | 1.113 |
| SLC25A26 | solute carrier family 25 member 26(SLC25A26) | NM_001009938 | 101.1 | 98.5 | 0.02781 | 1.124 |
| RASAL2 | RAS protein activator like 2(RASAL2) | NM_170692 | 104.5 | 99.8 | 0.02782 | 1.156 |
| STXBP1 | syntaxin binding protein 1(STXBP1) | NM_001032221 | 100.5 | 104.6 | 0.02784 | 1.143 |
| | | AF009305 | 97.1 | 97.9 | 0.02795 | 1.177 |
| ORC1 | origin recognition complex subunit 1(ORC1) | NM_004153 | 115.9 | 102.3 | 0.02801 | 0.866 |
| KLHL5 | kelch like family member 5(KLHL5) | NM_001007075 | 108.1 | 105.5 | 0.02802 | 0.644 |
| KLHL5 | kelch like family member 5(KLHL5) | NM_001007075 | 111.3 | 104.6 | 0.02802 | 0.644 |
| KLHL5 | kelch like family member 5(KLHL5) | NM_001007075 | 351.6 | 304.8 | 0.02802 | 0.644 |
| | | XM_001131851 | 131.7 | 134.4 | 0.02806 | 1.163 |
| | | XM_929859 | 131.7 | 120.7 | 0.02807 | 0.809 |
| MRPS2 | mitochondrial ribosomal protein S2(MRPS2) | NM_016034 | 158.8 | 151.8 | 0.0281 | 1.263 |
| | | CV318379 | 116.5 | 109.4 | 0.02813 | 0.93 |

| | | | | | | |
|---|---|---|---|---|---|---|
| TIAM2 | T-cell lymphoma invasion and metastasis 2(TIAM2) | NM_012454 | 90.2 | 83 | 0.02818 | 0.869 |
| ZNF544 | zinc finger protein 544(ZNF544) | NM_014480 | 139.9 | 139.6 | 0.02822 | 1.251 |
| MMAA | methylmalonic aciduria (cobalamin deficiency) cblA type(MMAA) | NM_172250 | 104.1 | 107.6 | 0.02831 | 1.107 |
| HAX1 | HCLS1 associated protein X-1(HAX1) | NM_006118 | 106.2 | 95.5 | 0.02836 | 0.896 |
| NFX1 | nuclear transcription factor, X-box binding 1(NFX1) | NM_147133 | 125.1 | 115 | 0.02839 | 1.188 |
| NFX1 | nuclear transcription factor, X-box binding 1(NFX1) | NM_147133 | 136.2 | 123.8 | 0.02839 | 1.188 |
| MINOS1 | mitochondrial inner membrane organizing system 1(MINOS1) | NM_001032363 | 107.5 | 100.1 | 0.0284 | 1.182 |
| MINOS1 | mitochondrial inner membrane organizing system 1(MINOS1) | NM_001032363 | 110.6 | 123.7 | 0.0284 | 1.182 |
| c19orf60 | chromosome 19 open reading frame 60(C19orf60) | NM_001100419 | 97.5 | 105.4 | 0.02846 | 1.138 |
| LINC00161 | long intergenic non-protein coding RNA 161(LINC00161) | NR_026552 | 94.1 | 102.4 | 0.02847 | 1.135 |
| SLC9A8 | solute carrier family 9 member A8(SLC9A8) | NM_015266 | 99 | 96.4 | 0.02853 | 0.878 |

| | | | | | | |
|---|---|---|---|---|---|---|
| POLR2H | RNA polymerase II subunit H(POLR2H) | NM_006232 | 160.7 | 168.4 | 0.0286 | 1.217 |
| | | BU734093 | 141.7 | 135.8 | 0.02871 | 1.192 |
| lemd3 | LEM domain containing 3(LEMD3) | NM_014319 | 213.3 | 209.3 | 0.02873 | 1.179 |
| | | XM_931518 | 88.3 | 82.8 | 0.02884 | 0.835 |
| NFYA | nuclear transcription factor Y subunit alpha(NFYA) | NM_002505 | 105.6 | 91.6 | 0.02884 | 0.886 |
| | | AW150545 | 101.2 | 98.4 | 0.02887 | 1.115 |
| PIN1 | peptidylprolyl cis/trans isomerase, NIMA-interacting 1(PIN1) | NM_006221 | 324 | 338.9 | 0.02887 | 1.19 |
| | | XM_0017223251 | 129.5 | 128.5 | 0.02892 | 1.16 |
| RFESD | Rieske Fe-S domain containing(RFESD) | NM_001131066 | 96.8 | 87.2 | 0.02906 | 0.786 |
| EXO5 | exonuclease 5(EXO5) | NM_022774 | 98 | 94.6 | 0.02922 | 0.808 |
| EXO5 | exonuclease 5(EXO5) | NM_022774 | 99.5 | 94.6 | 0.02922 | 0.808 |
| EXO5 | exonuclease 5(EXO5) | NM_022774 | 1035.5 | 803 | 0.02922 | 0.808 |
| SPDYE8P | speedy/RINGO cell cycle regulator family member E8, pseudogene(SPDYE8P) | NR_003664 | 106.3 | 92.3 | 0.02929 | 0.727 |

| | | | | | | |
|---|---|---|---|---|---|---|
| SPDYE8P | speedy/RINGO cell cycle regulator family member E8, pseudogene(SPDYE8P) | NR_003664 | 13009 | 11409.6 | 0.02929 | 0.727 |
| TNRC6A | trinucleotide repeat containing 6A(TNRC6A) | NM_014494 | 128.5 | 121.8 | 0.0293 | 0.868 |
| TNRC6A | trinucleotide repeat containing 6A(TNRC6A) | NM_014494 | 134.2 | 116.6 | 0.0293 | 0.868 |
| TNRC6A | trinucleotide repeat containing 6A(TNRC6A) | NM_014494 | 310.9 | 258.5 | 0.0293 | 0.868 |
| OR7C2 | olfactory receptor family 7 subfamily C member 2(OR7C2) | NM_012377 | 108.8 | 107.4 | 0.0293 | 1.116 |
| GPS1 | G protein pathway suppressor 1(GPS1) | NM_212492 | 148.3 | 163.4 | 0.02938 | 0.897 |
| GPS1 | G protein pathway suppressor 1(GPS1) | NM_004127 | 103.2 | 98.9 | 0.02938 | 0.897 |
| BRI3BP | BRI3 binding protein(BRI3BP) | NM_080626 | 117 | 118 | 0.02938 | 1.772 |
| BRI3BP | BRI3 binding protein(BRI3BP) | NM_080626 | 124.8 | 116.6 | 0.02938 | 1.772 |
| BRI3BP | BRI3 binding protein(BRI3BP) | NM_080626 | 305.1 | 288.1 | 0.02938 | 1.772 |
| NPR3 | natriuretic peptide receptor 3(NPR3) | NM_000908 | 116.2 | 124.6 | 0.02947 | 1.363 |
| TBC1D9B | TBC1 domain family member 9B(TBC1D9B) | NM_015043 | 112.5 | 98.2 | 0.02948 | 0.814 |

| | | | | | | |
|---|---|---|---|---|---|---|
| | | BQ021435 | 100.1 | 107.3 | 0.02951 | 1.131 |
| | | XM_001726935 | 113 | 104 | 0.02951 | 0.692 |
| TMEM237 | transmembrane protein 237(TMEM237) | NM_001044385 | 136.9 | 130.4 | 0.02976 | 0.758 |
| CCT4 | chaperonin containing TCP1 subunit 4(CCT4) | NM_006430 | 164.5 | 174.9 | 0.02979 | 1.323 |
| KLHL28 | kelch like family member 28(KLHL28) | NM_017658 | 145.5 | 125.2 | 0.02984 | 0.745 |
| | | XM_001132864 | 96 | 93.4 | 0.0299 | 0.844 |
| SLC17A2 | solute carrier family 17 member 2(SLC17A2) | NM_005835 | 94.3 | 100.6 | 0.02994 | 1.194 |
| | | XR_038252 | 103.7 | 111.8 | 0.03001 | 0.752 |
| | | XR_038252 | 610.6 | 508.9 | 0.03001 | 0.752 |
| CD300LG | CD300 molecule like family member g(CD300LG) | NM_145273 | 99.2 | 93.7 | 0.03002 | 0.879 |
| RTKN2 | rhotekin 2(RTKN2) | NM_145307 | 101.3 | 92.8 | 0.03003 | 0.879 |
| PCDHGA10 | protocadherin gamma subfamily A, 10(PCDHGA10) | NM_032090 | 115.6 | 102.7 | 0.03005 | 0.866 |
| | | XM_941706 | 99.8 | 85.9 | 0.03012 | 0.849 |
| SSRP1 | structure specific recognition protein 1(SSRP1) | NM_003146 | 93 | 87.8 | 0.03019 | 0.886 |

| | | | | | | |
|---|---|---|---|---|---|---|
| | | XM_001725293 | 258.1 | 171.5 | 0.03021 | 0.619 |
| KHDRBS3 | KH RNA binding domain containing, signal transduction associated 3(KHDRBS3) | NM_006558 | 536.6 | 500.8 | 0.03022 | 0.757 |
| KDM4B | lysine demethylase 4B(KDM4B) | NM_015015 | 345.9 | 281.4 | 0.03044 | 0.792 |
| HIKESHI | Hikeshi, heat shock protein nuclear import factor(HIKESHI) | NM_016401 | 177.8 | 214.5 | 0.03053 | 1.318 |
| | | XR_038706 | 360.2 | 335.1 | 0.0306 | 1.329 |
| | | AK021616 | 90.9 | 88.8 | 0.03065 | 0.832 |
| ROM1 | retinal outer segment membrane protein 1(ROM1) | NM_000327 | 124 | 142.3 | 0.03065 | 1.154 |
| | | AI263926 | 93.6 | 102.7 | 0.03068 | 1.217 |
| SPO11 | SPO11, initiator of meiotic double stranded breaks(SPO11) | NM_198265 | 103.7 | 99.5 | 0.03071 | 0.834 |
| SPO11 | SPO11, initiator of meiotic double stranded breaks(SPO11) | NM_198265 | 106.4 | 94.2 | 0.03071 | 0.834 |
| JMJD8 | jumonji domain containing 8(JMJD8) | NM_001005920 | 140.9 | 128.2 | 0.03073 | 1.24 |
| ADD3 | adducin 3(ADD3) | NM_001121 | 1598.4 | 1405.8 | 0.03086 | 0.847 |

| | | | | | | |
|---|---|---|---|---|---|---|
| PDE8B | phosphodiesterase 8B(PDE8B) | NM_001029851 | 93.6 | 97.1 | 0.03091 | 1.116 |
| PDE8B | phosphodiesterase 8B(PDE8B) | NM_001029851 | 103.8 | 104.3 | 0.03091 | 1.116 |
| TMED9 | transmembrane p24 trafficking protein 9(TMED9) | NM_017510 | 187.4 | 172.8 | 0.03092 | 0.785 |
| TDP1 | tyrosyl-DNA phosphodiesterase 1(TDP1) | NM_018319 | 94.5 | 101.4 | 0.03101 | 1.125 |
| TDP1 | tyrosyl-DNA phosphodiesterase 1(TDP1) | NM_018319 | 97.8 | 100.2 | 0.03101 | 1.125 |
| TDP1 | tyrosyl-DNA phosphodiesterase 1(TDP1) | NM_018319 | 1550.9 | 1032.5 | 0.03101 | 1.125 |
| FGF21 | fibroblast growth factor 21(FGF21) | NM_019113 | 99.7 | 108 | 0.03104 | 1.098 |
| SUZ12 | SUZ12 polycomb repressive complex 2 subunit(SUZ12) | NM_015355 | 493.9 | 371.7 | 0.03109 | 1.517 |
| | | XR_037524 | 130.6 | 109.2 | 0.0311 | 0.819 |
| CUL5 | cullin 5(CUL5) | NM_003478 | 205.7 | 184.3 | 0.03112 | 0.826 |
| ZNF549 | zinc finger protein 549(ZNF549) | NM_153263 | 131.5 | 120.3 | 0.03118 | 0.862 |
| ZNF549 | zinc finger protein 549(ZNF549) | NM_153263 | 5630.6 | 3969.7 | 0.03118 | 0.862 |

| | | | | | | | | | |
|---|---|---|---|---|---|---|---|---|---|
| ATP1B1 | ATPase Na+/K+ transporting subunit beta 1(ATP1B1) | NM_001677 | 109.6 | 96.6 | 0.0312 | 0.878 | NM_031012 | 19.39 | 0.85 |
| | | XM_936697 | 101.9 | 105.5 | 0.03128 | 1.112 | | | |
| LOC613206 | myeloproliferative disease associated tumor antigen 5(LOC613206) | NM_001033016 | 92.5 | 88 | 0.03131 | 0.876 | | | |
| PUDP | pseudouridine 5'-phosphatase(PUDP) | NM_012080 | 99.4 | 110.4 | 0.03144 | 1.166 | | | |
| PCBP2 | poly(rC) binding protein 2(PCBP2) | NM_001098620 | 1075.5 | 729.4 | 0.03147 | 1.288 | NM_012904 | 10.16 | 55.22 |
| DCUN1D5 | defective in cullin neddylation 1 domain containing 5(DCUN1D5) | NM_032299 | 109 | 105.9 | 0.03148 | 1.098 | | | |
| | | XM_001714155 | 100.1 | 99.8 | 0.03152 | 1.164 | | | |
| GSKIP | GSK3B interacting protein(GSKIP) | NM_016472 | 260.7 | 206.5 | 0.03153 | 1.212 | | | |
| CHIC2 | cysteine rich hydrophobic domain 2(CHIC2) | NM_012110 | 97 | 109.6 | 0.03167 | 1.166 | | | |
| PNMA1 | paraneoplastic Ma antigen 1(PNMA1) | NM_006029 | 186.3 | 186.8 | 0.03178 | 0.762 | | | |
| AKAP11 | A-kinase anchoring protein 11(AKAP11) | NM_016248 | 271.4 | 236.4 | 0.03179 | 0.866 | | | |
| AKAP12 | A-kinase anchoring protein 12(AKAP12) | NM_144497 | 347.5 | 322.1 | 0.03179 | 0.866 | | | |

| | | | | | | | | | |
|---|---|---|---|---|---|---|---|---|---|
| | | DA266249 | 97.2 | 90.4 | 0.03191 | 0.864 | | | |
| ADGRE5 | adhesion G protein-coupled receptor E5(ADGRE5) | NM_078481 | 104.5 | 105.2 | 0.03199 | 1.093 | | | |
| WDR44 | WD repeat domain 44(WDR44) | NM_019045 | 114.7 | 111 | 0.03201 | 1.146 | | | |
| WDR44 | WD repeat domain 44(WDR44) | NM_019045 | 125.3 | 120.4 | 0.03201 | 1.146 | | | |
| POLRMT | RNA polymerase mitochondrial(POLRMT) | NM_005035 | 202.6 | 175 | 0.03202 | 0.834 | NM_012823 | 17.52 | 0.04 |
| | | XM_938965 | 93.5 | 103.4 | 0.03203 | 1.198 | | | |
| SLC10A7 | solute carrier family 10 member 7(SLC10A7) | NM_001029998 | 96.7 | 103 | 0.03204 | 0.869 | | | |
| SLC10A7 | solute carrier family 10 member 7(SLC10A7) | NM_001029998 | 115.6 | 108.9 | 0.03204 | 0.869 | | | |
| | | XR_017484 | 115.3 | 111.7 | 0.03211 | 0.457 | | | |
| INTS6L | integrator complex subunit 6 like(INTS6L) | NM_182540 | 110.7 | 110.5 | 0.03221 | 1.206 | | | |
| SOX18 | SRY-box 18(SOX18) | NM_018419 | 256.5 | 228.1 | 0.0323 | 1.323 | | | |
| | | XM_001713810 | 450.6 | 407.6 | 0.03235 | 1.904 | | | |
| LATS2 | large tumor suppressor kinase 2(LATS2) | NM_014572 | 101.1 | 100.5 | 0.0324 | 1.156 | | | |
| | | XM_929541 | 94.6 | 98.8 | 0.03246 | 1.115 | | | |

| | | | | | | |
|---|---|---|---|---|---|---|
| TMBIM1 | transmembrane BAX inhibitor motif containing 1(TMBIM1) | NM_022152 | 229.2 | 278.1 | 0.03251 | 1.328 |
| | | XM_377725 | 111.5 | 114.1 | 0.03257 | 1.217 |
| SUPV3L1 | Suv3 like RNA helicase(SUPV3L1) | NM_003171 | 129.1 | 116.3 | 0.0326 | 0.894 |
| PIKFYVE | phosphoinositide kinase, FYVE-type zinc finger containing(PIKFYVE) | NM_015040 | 90.8 | 96.9 | 0.03264 | 1.158 |
| DCP2 | decapping mRNA 2(DCP2) | NM_152624 | 111.9 | 122.2 | 0.0327 | 1.162 |
| | | XM_938181 | 7544.4 | 7392.2 | 0.0327 | 1.222 |
| TH | tyrosine hydroxylase(TH) | NM_199293 | 94 | 88.2 | 0.03272 | 0.868 |
| | | AI732113 | 109.2 | 110.2 | 0.03273 | 1.11 |
| TSPAN31 | tetraspanin 31(TSPAN31) | NM_005981 | 280.4 | 249.8 | 0.03275 | 1.214 |
| | | BQ186959 | 100.2 | 97.5 | 0.03276 | 1.069 |
| | | XM_001723047 | 99.5 | 95 | 0.03278 | 0.878 |
| | | XM_937648 | 106.7 | 103.7 | 0.03278 | 0.879 |
| TCEAL9 | transcription elongation factor A like 9(TCEAL9) | NM_001006612 | 167.3 | 169.6 | 0.0328 | 1.264 |
| RHBDD1 | rhomboid domain containing 1(RHBDD1) | NM_032276 | 90.9 | 97 | 0.03282 | 1.129 |

| | | | | | | |
|---|---|---|---|---|---|---|
| RHBDD 1 | rhomboid domain containing 1(RHBDD1) | NM_032276 | 166.5 | 160.2 | 0.03282 | 1.129 |
| POLH | DNA polymerase eta(POLH) | NM_006502 | 108 | 91.9 | 0.03282 | 0.775 |
| | | XM_499363 | 92.3 | 87 | 0.03283 | 0.895 |
| H2AFJ | H2A histone family member J(H2AFJ) | NM_177925 | 129.8 | 116.1 | 0.0329 | 0.622 |
| TAPT1-AS1 | TAPT1 antisense RNA 1 (head to head)(TAPT1-AS1) | CA391675 | 112.2 | 98.4 | 0.03294 | 0.883 |
| CLUAP1 | clusterin associated protein 1(CLUAP1) | NM_015041 | 104.7 | 107.7 | 0.03295 | 0.869 |
| CLUAP1 | clusterin associated protein 1(CLUAP1) | NM_024793 | 110.9 | 104.1 | 0.03295 | 0.869 |
| | | XM_370557 | 98.3 | 90.5 | 0.03298 | 0.864 |
| | | XM_370557 | 111.8 | 105 | 0.03298 | 0.864 |
| | | XM_944469 | 90 | 79.1 | 0.03299 | 0.781 |
| SLC35C2 | solute carrier family 35 member C2(SLC35C2) | NM_173073 | 141.4 | 136.5 | 0.03302 | 1.131 |
| arfip1 | ADP ribosylation factor interacting protein 1(ARFIP1) | NM_001025595 | 95.4 | 90.7 | 0.03309 | 0.843 |
| FHOD1 | formin homology 2 domain containing 1(FHOD1) | NM_013241 | 220.4 | 322.4 | 0.03315 | 0.463 |

| | | | | | | |
|---|---|---|---|---|---|---|
| DISP2 | dispatched RND transporter family member 2(DISP2) | NM_033510 | 112.2 | 113.2 | 0.03316 | 1.122 |
| CSE1L | chromosome segregation 1 like(CSE1L) | NM_177436 | 412.3 | 329 | 0.03336 | 0.743 |
| CSE1L | chromosome segregation 1 like(CSE1L) | NM_001316 | 378.1 | 277.6 | 0.03336 | 0.743 |
| TRAPPC6B | trafficking protein particle complex 6B(TRAPPC6B) | NM_001079537 | 91.8 | 92.4 | 0.03336 | 0.777 |
| TRAPPC6B | trafficking protein particle complex 6B(TRAPPC6B) | NM_001079537 | 109.7 | 102.2 | 0.03336 | 0.777 |
| UFM1 | ubiquitin fold modifier 1(UFM1) | NM_016617 | 209.1 | 185.6 | 0.03344 | 1.324 |
| | | AA454665 | 106.6 | 106.6 | 0.03356 | 1.13 |
| SYNE4 | spectrin repeat containing nuclear envelope family member 4(SYNE4) | NM_001039876 | 98.9 | 91.6 | 0.03363 | 0.903 |
| PSRC1 | proline and serine rich coiled-coil 1(PSRC1) | NM_001032290 | 107.7 | 106.5 | 0.03369 | 0.906 |
| PPM1B | protein phosphatase, Mg2+/Mn2+ dependent 1B(PPM1B) | NM_177968 | 492.6 | 429.7 | 0.03382 | 1.302 |
| PLAC9 | placenta specific 9(PLAC9) | NM_001012973 | 137.1 | 135.9 | 0.03384 | 0.603 |

| | | | | | | | | | |
|---|---|---|---|---|---|---|---|---|---|
| OR2A1 | olfactory receptor family 2 subfamily A member 1(OR2A1) | NM_001005287 | 101.9 | 91.1 | 0.03389 | 0.87 | | | |
| Uqcrh | ubiquinol-cytochrome c reductase hinge protein(UQCRH) | NM_006004 | 97.5 | 94.4 | 0.0339 | 0.839 | | | |
| Uqcrh | ubiquinol-cytochrome c reductase hinge protein(UQCRH) | NM_006004 | 974.9 | 1252.9 | 0.0339 | 0.839 | | | |
| CSNK2A1 | casein kinase 2 alpha 1(CSNK2A1) | NM_001895 | 136.4 | 143.1 | 0.03393 | 0.813 | | | |
| CSNK2A2 | casein kinase 2 alpha 2(CSNK2A2) | NM_001896 | 185.3 | 174 | 0.03393 | 0.813 | | | |
| ADCY3 | adenylate cyclase 3(ADCY3) | NM_004036 | 102.3 | 88.5 | 0.03401 | 0.898 | | | |
| PTPA | protein phosphatase 2 phosphatase activator(PTPA) | NM_178003 | 108.9 | 110.5 | 0.03408 | 1.127 | | | |
| | | XR_001280 | 105.8 | 106.1 | 0.03411 | 1.125 | | | |
| MAP3K4 | mitogen-activated protein kinase kinase kinase 4(MAP3K4) | NM_005922 | 136.5 | 131 | 0.03415 | 1.193 | | | |
| TRMT10A | tRNA methyltransferase 10A(TRMT10A) | NM_152292 | 101 | 96.3 | 0.03425 | 1.133 | NM_017170 | 13.37 | 0.05 |
| EIF4G2 | eukaryotic translation initiation factor 4 gamma 2(EIF4G2) | NM_001418 | 369.1 | 341.9 | 0.03434 | 0.827 | | | |
| EIF4G2 | eukaryotic translation initiation factor 4 gamma 2(EIF4G2) | NM_001418 | 1858.1 | 1378.6 | 0.03434 | 0.827 | | | |

| | | | | | | | | | |
|---|---|---|---|---|---|---|---|---|---|
| EIF4G2 | eukaryotic translation initiation factor 4 gamma 2(EIF4G2) | NM_001418 | 3468.8 | 2977.2 | 0.03434 | 0.827 | | | |
| KDM2A | lysine demethylase 2A(KDM2A) | NM_012308 | 102.2 | 108.3 | 0.03439 | 0.91 | | | |
| spdya | speedy/RINGO cell cycle regulator family member A(SPDYA) | NM_182756 | 94.9 | 100.2 | 0.0345 | 1.15 | | | |
| MSLNL | mesothelin-like(MSLNL) | NM_001025190 | 96.1 | 102 | 0.03452 | 1.157 | | | |
| MSLNL | mesothelin-like(MSLNL) | NM_001025190 | 101.8 | 99.3 | 0.03452 | 1.157 | | | |
| CEP85L | centrosomal protein 85 like(CEP85L) | NM_206921 | 101.6 | 93.4 | 0.03455 | 0.873 | | | |
| | | XM_942374 | 106 | 96.4 | 0.03461 | 0.818 | NM_001106492 | 2.61 | 10.85 |
| NME2 | NME/NM23 nucleoside diphosphate kinase 2(NME2) | NM_001018138 | 96.1 | 99.2 | 0.03465 | 1.129 | NM_001106492 | 2.61 | 10.85 |
| ELOVL3 | ELOVL fatty acid elongase 3(ELOVL3) | NM_152310 | 109.3 | 98.8 | 0.03474 | 0.805 | NM_001106492 | 2.61 | 10.85 |
| LOC100996671 | uncharacterized LOC100996671(LOC100996671) | BX111310 | 113.8 | 106.8 | 0.03476 | 0.834 | | | |
| GNL3L | G protein nucleolar 3 like(GNL3L) | NM_019067 | 147.4 | 139.9 | 0.03477 | 0.771 | | | |
| PUS7L | pseudouridylate synthase 7 like(PUS7L) | NM_001098615 | 107.1 | 97.3 | 0.03477 | 0.858 | | | |

| | | | | | | |
|---|---|---|---|---|---|---|
| | | XM_944547 | 114.3 | 112 | 0.03477 | 1.202 |
| | | XM_929962 | 103.3 | 104.5 | 0.03483 | 1.131 |
| DCAF10 | DDB1 and CUL4 associated factor 10(DCAF10) | NM_024345 | 527 | 473.7 | 0.03486 | 1.6 |
| RABL3 | RAB, member of RAS oncogene family like 3(RABL3) | NM_173825 | 149.1 | 168.3 | 0.0349 | 1.267 |
| RSPH10B | radial spoke head 10 homolog B(RSPH10B) | NM_173565 | 93.6 | 90.3 | 0.03491 | 0.762 |
| RSPH10B | radial spoke head 10 homolog B(RSPH10B) | NM_173565 | 118 | 111.6 | 0.03491 | 0.762 |
| RSPH10B | radial spoke head 10 homolog B(RSPH10B) | NM_173565 | 119.8 | 142.3 | 0.03491 | 0.762 |
| RSPH10B | radial spoke head 10 homolog B(RSPH10B) | NM_173565 | 120.8 | 133.7 | 0.03491 | 0.762 |
| MTFMT | mitochondrial methionyl-tRNA formyltransferase(MTFMT) | NM_139242 | 120.9 | 116.6 | 0.03495 | 1.16 |
| MTFMT | mitochondrial methionyl-tRNA formyltransferase(MTFMT) | NM_139242 | 166.5 | 161.6 | 0.03495 | 1.16 |
| UTRN | utrophin(UTRN) | NM_007124 | 104.4 | 97.4 | 0.03497 | 0.782 |
| NOL6 | nucleolar protein 6(NOL6) | NM_139235 | 136.9 | 115.5 | 0.03505 | 0.772 |

| | | | | | | | | | |
|---|---|---|---|---|---|---|---|---|---|
| CAPN10 | calpain 10(CAPN10) | NM_023083 | 113.5 | 102.7 | 0.03514 | 1.132 | | | |
| TBX19 | T-box 19(TBX19) | NM_005149 | 98.5 | 104.3 | 0.03515 | 1.151 | | | |
| TBX19 | T-box 19(TBX19) | NM_005149 | 120.8 | 125.9 | 0.03515 | 1.151 | | | |
| | | CB858008 | 103.8 | 104.4 | 0.03516 | 1.155 | NM_001270681 | 26.79 | 0.09 |
| CRYL1 | crystallin lambda 1(CRYL1) | NM_015974 | 95.5 | 89.6 | 0.03517 | 0.872 | | | |
| KIAA0430 | KIAA0430(KIAA0430) | NM_014647 | 99 | 110 | 0.03525 | 1.206 | | | |
| CCDC114 | coiled-coil domain containing 114(CCDC114) | NM_144577 | 131.9 | 119.5 | 0.03538 | 0.866 | | | |
| | | XM_937287 | 87.2 | 94.4 | 0.03544 | 1.17 | | | |
| RSPRY1 | ring finger and SPRY domain containing 1(RSPRY1) | NM_133368 | 294.2 | 256.2 | 0.03547 | 1.244 | | | |
| RECQL5 | RecQ like helicase 5(RECQL5) | NM_001003716 | 99.3 | 103 | 0.03559 | 1.086 | | | |
| SLC37A3 | solute carrier family 37 member 3(SLC37A3) | NM_207113 | 91.6 | 87.8 | 0.03569 | 0.91 | NM_019373 | 263.23 | 1.01 |
| ankrd40 | ankyrin repeat domain 40(ANKRD40) | NM_052855 | 107.5 | 112.8 | 0.0357 | 0.719 | | | |
| SRP14 | signal recognition particle 14(SRP14) | NM_003134 | 236.8 | 255.8 | 0.03572 | 0.731 | | | |
| SRP14 | signal recognition particle 14(SRP14) | NM_003134 | 2418.6 | 2337.4 | 0.03572 | 0.731 | | | |

| | | | | | | | | | |
|---|---|---|---|---|---|---|---|---|---|
| AKR1C4 | aldo-keto reductase family 1 member C4(AKR1C4) | NM_001818 | 117.2 | 113.9 | 0.03575 | 1.309 | | | |
| AKR1C8P | aldo-keto reductase family 1 member C8, pseudogene(AKR1C8P) | NM_001007536 | 222.7 | 203.9 | 0.03575 | 1.309 | | | |
| NABP2 | nucleic acid binding protein 2(NABP2) | NM_024068 | 122 | 106.8 | 0.03575 | 0.854 | | | |
| ZBTB20 | zinc finger and BTB domain containing 20(ZBTB20) | AK091904 | 536.2 | 613.9 | 0.03575 | 1.512 | | | |
| EHBP1 | EH domain binding protein 1(EHBP1) | NM_015252 | 117.5 | 122.8 | 0.03579 | 1.258 | | | |
| SLC25A15 | solute carrier family 25 member 15(SLC25A15) | NM_014252 | 109.6 | 106 | 0.03584 | 0.896 | | | |
| | | XR_038676 | 106 | 96.4 | 0.03588 | 0.887 | | | |
| ZNF568 | zinc finger protein 568(ZNF568) | NM_198539 | 101.6 | 101.2 | 0.03606 | 1.096 | | | |
| ZNF568 | zinc finger protein 568(ZNF568) | NM_198539 | 111.7 | 108.7 | 0.03606 | 1.096 | | | |
| MAP3K3 | mitogen-activated protein kinase kinase kinase 3(MAP3K3) | NM_203351 | 119.5 | 116.9 | 0.03609 | 1.197 | NM_012778 | 744.78 | 0.71 |
| PTPRA | protein tyrosine phosphatase, receptor type A(PTPRA) | NM_080840 | 157.4 | 134.9 | 0.03609 | 0.854 | NM_012778 | 744.78 | 0.71 |
| PTPRA | protein tyrosine phosphatase, receptor type A(PTPRA) | NM_080840 | 192.4 | 178.3 | 0.03609 | 0.854 | NM_012778 | 744.78 | 0.71 |

| | | | | | | | | | |
|---|---|---|---|---|---|---|---|---|---|
| | | XM_374200 | 104 | 99.2 | 0.0361 | 1.1 | | | |
| MPG | N-methylpurine DNA glycosylase(MPG) | NM_001015052 | 101.8 | 103.6 | 0.03621 | 1.109 | | | |
| BBS2 | Bardet-Biedl syndrome 2(BBS2) | NM_031885 | 124.6 | 112.4 | 0.03625 | 0.855 | NM_012909 | 6.73 | 5427.66 |
| | | NM_001001700 | 99.8 | 101 | 0.03631 | 1.126 | NM_022181 | 0.40 | 160.77 |
| | | NM_001001700 | 141.1 | 129.8 | 0.03631 | 1.126 | NM_019157 | 14.48 | 0.05 |
| ENTPD1 | ectonucleoside triphosphate diphosphohydrolase 1(ENTPD1) | NM_001098175 | 178.6 | 209.8 | 0.03637 | 1.235 | | | |
| C1orf168 | chromosome 1 open reading frame 168(C1orf168) | NM_001004303 | 90.7 | 94.6 | 0.03637 | 1.191 | | | |
| YY1 | YY1 transcription factor(YY1) | NM_003403 | 282.3 | 279 | 0.03642 | 1.407 | | | |
| YY1 | YY1 transcription factor(YY1) | NM_003403 | 1066.2 | 1084 | 0.03642 | 1.407 | | | |
| PREB | prolactin regulatory element binding(PREB) | NM_013388 | 138.7 | 150.5 | 0.03647 | 1.191 | | | |
| | | BX093763 | 102.8 | 95.7 | 0.03651 | 0.87 | | | |
| | | XM_001134421 | 104.6 | 104.5 | 0.03658 | 1.123 | | | |
| MATR3 | matrin 3(MATR3) | NM_199189 | 91.5 | 98.5 | 0.03663 | 1.115 | | | |
| MATR3 | matrin 3(MATR3) | NM_199189 | 107.2 | 96.5 | 0.03663 | 1.115 | | | |

| | | | | | | |
|---|---|---|---|---|---|---|
| MATR3 | matrin 3(MATR3) | NM_199189 | 2099.5 | 1825.7 | 0.03663 | 1.115 |
| CAPRIN1 | cell cycle associated protein 1(CAPRIN1) | NM_005898 | 244.2 | 279.2 | 0.03664 | 0.718 |
| aire | autoimmune regulator(AIRE) | NM_000658 | 95.7 | 99.7 | 0.03665 | 1.116 |
| SMG6 | SMG6, nonsense mediated mRNA decay factor(SMG6) | NM_017575 | 104.3 | 95.8 | 0.03668 | 0.846 |
| KMT2C | lysine methyltransferase 2C(KMT2C) | NM_170606 | 111.9 | 101.8 | 0.03671 | 0.792 |
| KMT2E | lysine methyltransferase 2E(KMT2E) | NM_018682 | 147.2 | 122 | 0.03671 | 0.792 |
| GALT | galactose-1-phosphate uridylyltransferase(GALT) | NM_000155 | 111 | 102.9 | 0.03675 | 1.239 |
| GAMT | guanidinoacetate N-methyltransferase(GAMT) | NM_000156 | 154 | 156.5 | 0.03675 | 1.239 |
| AGBL5 | ATP/GTP binding protein like 5(AGBL5) | NM_021831 | 109.3 | 97.6 | 0.03679 | 0.846 |
| | | BG216851 | 93.7 | 103.7 | 0.03681 | 1.164 |
| | | XM_933468 | 113.2 | 108.8 | 0.03689 | 1.105 |
| HSF1 | heat shock transcription factor 1(HSF1) | NM_005526 | 98.4 | 99.2 | 0.03691 | 1.128 |
| FBXO48 | F-box protein 48(FBXO48) | NM_001024680 | 103.6 | 105.1 | 0.03699 | 1.099 |

| | | | | | | | | | |
|---|---|---|---|---|---|---|---|---|---|
| IFNA21 | interferon alpha 21(IFNA21) | NM_002175 | 107.8 | 106.8 | 0.03715 | 0.875 | | | |
| CAPS | calcyphosine(CAPS) | NM_004058 | 252.4 | 186.3 | 0.03722 | 0.833 | | | |
| MED16 | mediator complex subunit 16(MED16) | NM_005481 | 111.1 | 98.2 | 0.03728 | 1.192 | | | |
| MED16 | mediator complex subunit 16(MED16) | NM_005481 | 116.1 | 113.6 | 0.03728 | 1.192 | NM_019168 | 8.26 | 61.49 |
| PCBP2 | poly(rC) binding protein 2(PCBP2) | NM_005016 | 2165.2 | 1520.7 | 0.03738 | 1.259 | | | |
| SLC2A4RG | SLC2A4 regulator(SLC2A4RG) | NM_020062 | 150.6 | 159.9 | 0.03738 | 1.339 | | | |
| | | BF054947 | 99.1 | 94.1 | 0.03741 | 0.92 | | | |
| | | XR_037442 | 90 | 102 | 0.03752 | 1.168 | | | |
| BEND3 | BEN domain containing 3(BEND3) | NM_001080450 | 120.3 | 115.2 | 0.03753 | 1.102 | | | |
| RUNX1T1 | RUNX1 translocation partner 1(RUNX1T1) | NM_175636 | 95.3 | 99.8 | 0.03756 | 1.147 | | | |
| RBM45 | RNA binding motif protein 45(RBM45) | NM_152945 | 147.4 | 118.1 | 0.03758 | 0.822 | | | |
| | | DA804594 | 98.1 | 101.9 | 0.03767 | 1.112 | | | |
| | | XR_016054 | 156.9 | 180.8 | 0.03771 | 1.317 | | | |
| CEP41 | centrosomal protein 41(CEP41) | NM_018718 | 199.8 | 178.6 | 0.03777 | 1.191 | | | |
| bag4 | BCL2 associated athanogene 4(BAG4) | NM_004874 | 108.8 | 101.8 | 0.0378 | 1.117 | | | |
| | | AW975165 | 107.5 | 101.8 | 0.03783 | 1.119 | | | |

| | | | | | | |
|---|---|---|---|---|---|---|
| SLC22A7 | solute carrier family 22 member 7(SLC22A7) | NM_153320 | 104.6 | 103 | 0.03789 | 1.119 |
| SLC22A7 | solute carrier family 22 member 7(SLC22A7) | NM_153320 | 350.2 | 461.9 | 0.03789 | 1.119 |
| | | XM_001716323 | 97 | 96.5 | 0.03792 | 0.872 |
| | | XM_001713618 | 119.8 | 134.3 | 0.03797 | 1.2 |
| MARS | methionyl-tRNA synthetase(MARS) | NM_004990 | 96.4 | 98.5 | 0.03802 | 1.079 |
| | | BX098816 | 94.2 | 101.4 | 0.03807 | 1.113 |
| CLCN1 | chloride voltage-gated channel 1(CLCN1) | NM_000083 | 95.9 | 90.6 | 0.03813 | 0.908 |
| C4orf3 | chromosome 4 open reading frame 3(C4orf3) | NM_001001701 | 109.9 | 97.7 | 0.03819 | 0.716 |
| C4orf32 | chromosome 4 open reading frame 32(C4orf32) | NM_152400 | 182.1 | 169.5 | 0.03819 | 0.716 |
| SCN7A | sodium voltage-gated channel alpha subunit 7(SCN7A) | NM_002976 | 127.2 | 117.9 | 0.03821 | 1.18 |
| | | XM_943579 | 104.4 | 92.2 | 0.03823 | 0.876 |
| CTBP2 | C-terminal binding protein 2(CTBP2) | NM_001329 | 137.8 | 117.4 | 0.03851 | 0.834 |
| STX6 | syntaxin 6(STX6) | NM_005819 | 171.4 | 158 | 0.03852 | 0.78 |
| | | XR_041454 | 100.6 | 101.4 | 0.03854 | 1.093 |

| | | | | | | |
|---|---|---|---|---|---|---|
| LRIF1 | ligand dependent nuclear receptor interacting factor 1(LRIF1) | NM_018372 | 95.7 | 102.2 | 0.03856 | 1.112 |
| RAB5C | RAB5C, member RAS oncogene family(RAB5C) | NM_201434 | 267.6 | 255.4 | 0.03861 | 1.26 |
| | | AV700714 | 130 | 115.3 | 0.03868 | 0.803 |
| ABCC10 | ATP binding cassette subfamily C member 10(ABCC10) | NM_033450 | 102.1 | 97.2 | 0.0387 | 0.877 |
| TIMM17B | translocase of inner mitochondrial membrane 17 homolog B (yeast)(TIMM17B) | NM_005834 | 181.2 | 193.7 | 0.03872 | 1.155 |
| | | XM_941677 | 99.8 | 104.6 | 0.03872 | 1.116 |
| SHB | SH2 domain containing adaptor protein B(SHB) | NM_003028 | 125 | 132.2 | 0.03877 | 1.152 |
| | | XM_371901 | 96.9 | 94.7 | 0.03881 | 1.155 |
| RACK1 | receptor for activated C kinase 1(RACK1) | NM_006098 | 1944.8 | 1544.5 | 0.03882 | 1.319 |
| CCDC122 | coiled-coil domain containing 122(CCDC122) | NM_144974 | 112.6 | 119.8 | 0.039 | 1.183 |
| GALNT16 | polypeptide N-acetylgalactosaminyltransferase 16(GALNT16) | NM_020692 | 100.9 | 107 | 0.03902 | 1.171 |
| FGF23 | fibroblast growth factor 23(FGF23) | NM_020638 | 97.4 | 100.3 | 0.03903 | 1.145 |

| | | | | | | |
|---|---|---|---|---|---|---|
| PRNP | prion protein(PRNP) | NM_183079 | 94.5 | 99.9 | 0.03906 | 1.176 |
| | | XM_001726007 | 101.7 | 104.1 | 0.03917 | 1.142 |
| DCP1A | decapping mRNA 1A(DCP1A) | NM_018403 | 442.3 | 405.3 | 0.03928 | 0.442 |
| VPS28 | VPS28, ESCRT-I subunit(VPS28) | NM_016208 | 257.1 | 224.2 | 0.03936 | 0.62 |
| VPS28 | VPS28, ESCRT-I subunit(VPS28) | NM_016208 | 445.5 | 357.9 | 0.03936 | 0.62 |
| LAMTOR5 | late endosomal/lysosomal adaptor, MAPK and MTOR activator 5(LAMTOR5) | NM_006402 | 145.5 | 180.3 | 0.0395 | 1.251 |
| DAZ4 | deleted in azoospermia 4(DAZ4) | NM_020420 | 100.4 | 98.9 | 0.03952 | 0.903 |
| ZNF626 | zinc finger protein 626(ZNF626) | NM_001076675 | 105.2 | 101.2 | 0.03953 | 0.674 |
| ZNF626 | zinc finger protein 626(ZNF626) | NM_001076675 | 121.9 | 106.3 | 0.03953 | 0.674 |
| EDF1 | endothelial differentiation related factor 1(EDF1) | NM_003792 | 122.9 | 109.2 | 0.03956 | 0.661 |
| | | XM_930172 | 95 | 100.7 | 0.03963 | 1.136 |
| | | XM_937832 | 111 | 109.6 | 0.03963 | 1.13 |
| SFXN3 | sideroflexin 3(SFXN3) | NM_030971 | 113.2 | 115 | 0.03979 | 1.162 |

| | | | | | | | | | |
|---|---|---|---|---|---|---|---|---|---|
| | | XR_016984 | 90 | 85.9 | 0.03983 | 0.859 | | | |
| AFTPH | aftiphilin(AFTPH) | NM_203437 | 105.4 | 106.4 | 0.03984 | 1.158 | | | |
| AGAP1 | ArfGAP with GTPase domain, ankyrin repeat and PH domain 1(AGAP1) | NM_014914 | 170.7 | 166.5 | 0.03984 | 1.158 | | | |
| TRIM66 | tripartite motif containing 66(TRIM66) | NM_014818 | 111.5 | 115.9 | 0.03989 | 1.121 | NM_001106061 | 2.22 | 12.22 |
| PPP2R2B | protein phosphatase 2 regulatory subunit Bbeta(PPP2R2B) | NM_181676 | 94.3 | 90 | 0.03992 | 0.841 | | | |
| PPP2R2B | protein phosphatase 2 regulatory subunit Bbeta(PPP2R2B) | NM_181676 | 138.9 | 174.6 | 0.03992 | 0.841 | | | |
| CCNL2 | cyclin L2(CCNL2) | NM_030937 | 94.3 | 94.2 | 0.03993 | 1.207 | | | |
| CCNO | cyclin O(CCNO) | NM_021147 | 148.9 | 146.1 | 0.03993 | 1.207 | | | |
| | | XM_001133393 | 99.7 | 91.2 | 0.03994 | 0.91 | | | |
| | | XM_001133393 | 136.5 | 124.6 | 0.03994 | 0.91 | | | |
| GYS1 | glycogen synthase 1(GYS1) | NM_002103 | 107 | 110 | 0.03994 | 1.226 | | | |
| TRDMT1 | tRNA aspartic acid methyltransferase 1(TRDMT1) | NM_176083 | 98.6 | 105.1 | 0.03995 | 1.126 | | | |

| | | | | | | |
|---|---|---|---|---|---|---|
| JPH4 | junctophilin 4(JPH4) | NM_032452 | 123.1 | 111.4 | 0.03998 | 0.816 |
| | | XM_0017138 29 | 97.8 | 90.1 | 0.04012 | 0.863 |
| LOC606724 | coronin 1A pseudogene(LOC606724) | NR_002454 | 4102.1 | 2649.8 | 0.04015 | 0.694 |
| | | XM_929612 | 102.5 | 108 | 0.0402 | 1.135 |
| | | XM_927238 | 112 | 119.4 | 0.04032 | 1.122 |
| CCT7 | chaperonin containing TCP1 subunit 7(CCT7) | NM_001009570 | 99.2 | 98.7 | 0.04038 | 1.125 |
| ICA1L | islet cell autoantigen 1 like(ICA1L) | NM_178231 | 98.2 | 100.6 | 0.04056 | 1.109 |
| | | AA704464 | 105 | 107.4 | 0.04058 | 1.163 |
| | | NM_001018158 | 116.3 | 120 | 0.04059 | 1.188 |
| | | XM_940033 | 97.7 | 100.7 | 0.04069 | 1.146 |
| HNRNPDL | heterogeneous nuclear ribonucleoprotein D like(HNRNPDL) | NM_031372 | 1754 | 1190.2 | 0.0407 | 0.741 |
| | | XM_936861 | 104.2 | 102.6 | 0.04073 | 1.157 |
| TMEM179B | transmembrane protein 179B(TMEM179B) | NM_199337 | 148.7 | 141.3 | 0.04075 | 1.215 |
| TMEM179B | transmembrane protein 179B(TMEM179B) | NM_199337 | 371.3 | 310.8 | 0.04075 | 1.215 |
| | | XM_001715568 | 122.7 | 117.8 | 0.0408 | 1.106 |

| | | | | | | |
|---|---|---|---|---|---|---|
| RUNDC 1 | RUN domain containing 1(RUNDC1) | NM_1730 79 | 94.9 | 94.6 | 0.04083 | 0.851 |
| | | XM_00112 8220 | 109.4 | 105.1 | 0.04087 | 1.095 |
| CLDN14 | claudin 14(CLDN14) | NM_0121 30 | 116.2 | 161.5 | 0.04089 | 0.554 |
| | | XR_01855 3 | 92.2 | 85.8 | 0.04089 | 0.842 |
| | | XR_01855 3 | 3382.9 | 2599.9 | 0.04089 | 0.842 |
| SUMF1 | sulfatase modifying factor 1(SUMF1) | NM_1827 60 | 184.7 | 201.5 | 0.04092 | 1.158 |
| | | XM_93766 6 | 98.6 | 96.9 | 0.04102 | 1.169 |
| dguok | deoxyguanosine kinase(DGUOK) | NM_0809 16 | 93.7 | 98.1 | 0.04107 | 1.136 |
| MPPE1 | metallophosphoesteras e 1(MPPE1) | NM_0230 75 | 194.9 | 191.8 | 0.04114 | 1.229 |
| TAZ | tafazzin(TAZ) | NM_1813 12 | 130.4 | 114 | 0.04118 | 0.858 |
| | | XM_00173 2883 | 99 | 104.1 | 0.04121 | 1.128 |
| | | XM_00112 6665 | 104.3 | 104.4 | 0.04127 | 1.116 |
| | | AA909564 | 103.4 | 111.6 | 0.0414 | 1.19 |
| osbpl6 | oxysterol binding protein like 6(OSBPL6) | NM_0325 23 | 91.8 | 90.1 | 0.04142 | 1.222 |
| osbpl6 | oxysterol binding protein like 6(OSBPL6) | NM_0325 23 | 118.5 | 118.8 | 0.04142 | 1.222 |

| | | | | | | |
|---|---|---|---|---|---|---|
| PSMC6 | proteasome 26S subunit, ATPase 6(PSMC6) | NM_002806 | 98.3 | 99.4 | 0.04143 | 1.24 |
| PSMC6 | proteasome 26S subunit, ATPase 6(PSMC6) | NM_002806 | 467.8 | 505.9 | 0.04143 | 1.24 |
| HSD17B7 | hydroxysteroid 17-beta dehydrogenase 7(HSD17B7) | NM_016371 | 88.7 | 96.3 | 0.04145 | 1.177 |
| INPP5K | inositol polyphosphate-5-phosphatase K(INPP5K) | NM_130766 | 104 | 98.5 | 0.04166 | 0.914 |
| INPPL1 | inositol polyphosphate phosphatase like 1(INPPL1) | NM_001567 | 128.2 | 112.2 | 0.04166 | 0.914 |
| HRG | histidine rich glycoprotein(HRG) | NM_000412 | 92.7 | 95.2 | 0.04169 | 1.117 |
| | | XM_941665 | 301.3 | 348.4 | 0.04175 | 0.695 |
| GUSBP2 | glucuronidase, beta pseudogene 2(GUSBP2) | NR_003504 | 96.1 | 106 | 0.04179 | 1.119 |
| CSGALNACT2 | chondroitin sulfate N-acetylgalactosaminyltransferase 2(CSGALNACT2) | NM_018590 | 86.5 | 94.4 | 0.04187 | 1.146 |
| NMD3 | NMD3 ribosome export adaptor(NMD3) | NM_015938 | 98.9 | 95.9 | 0.04188 | 0.874 |

| | | | | | | |
|---|---|---|---|---|---|---|
| SLC29A1 | solute carrier family 29 member 1 (Augustine blood group)(SLC29A1) | NM_001078174 | 112.4 | 100 | 0.04192 | 0.854 |
| SLC29A1 | solute carrier family 29 member 1 (Augustine blood group)(SLC29A1) | NM_001078174 | 145.7 | 134.3 | 0.04192 | 0.854 |
| LMBRD1 | LMBR1 domain containing 1(LMBRD1) | NM_018368 | 118.5 | 119.1 | 0.04193 | 1.242 |
| | | XR_041507 | 95.3 | 106.8 | 0.04197 | 1.132 |
| LHX6 | LIM homeobox 6(LHX6) | NM_199160 | 90.8 | 86.1 | 0.04201 | 0.903 |
| GINM1 | glycoprotein integral membrane 1(GINM1) | NM_138785 | 122.3 | 102.7 | 0.04206 | 0.838 |
| GIN1 | gypsy retrotransposon integrase 1(GIN1) | NM_017676 | 101.3 | 100.6 | 0.04206 | 0.838 |
| WDR83 | WD repeat domain 83(WDR83) | NM_032332 | 123.2 | 113 | 0.04207 | 1.161 |
| COL4A5 | collagen type IV alpha 5 chain(COL4A5) | NM_000495 | 411.5 | 346.3 | 0.0421 | 1.547 |
| COL4A3BP | collagen type IV alpha 3 binding protein(COL4A3BP) | NM_005713 | 127.5 | 110.4 | 0.0421 | 1.547 |
| APOBEC3H | apolipoprotein B mRNA editing enzyme catalytic subunit 3H(APOBEC3H) | NM_181773 | 83.4 | 86.7 | 0.04211 | 0.859 |

| | | | | | | | | | |
|---|---|---|---|---|---|---|---|---|---|
| UGGT1 | UDP-glucose glycoprotein glucosyltransferase 1(UGGT1) | NM_020120 | 151.8 | 128.8 | 0.04211 | 1.303 | | | |
| TAOK1 | TAO kinase 1(TAOK1) | NM_020791 | 114 | 104.4 | 0.04226 | 1.121 | | | |
| HIVEP3 | human immunodeficiency virus type I enhancer binding protein 3(HIVEP3) | BC080552 | 86.7 | 97 | 0.04229 | 1.182 | | | |
| BCKDHB | branched chain keto acid dehydrogenase E1 subunit beta(BCKDHB) | NM_000056 | 134.2 | 153.7 | 0.04231 | 1.319 | NM_001014273 | 3.32 | 15.01 |
| BCKDHB | branched chain keto acid dehydrogenase E1 subunit beta(BCKDHB) | NM_183050 | 196 | 178.6 | 0.04231 | 1.319 | NM_001014273 | 3.32 | 15.01 |
| GRIA3 | glutamate ionotropic receptor AMPA type subunit 3(GRIA3) | NM_007325 | 168.1 | 159.5 | 0.04234 | 1.452 | NM_001014273 | 3.32 | 15.01 |
| HSD17B12 | hydroxysteroid 17-beta dehydrogenase 12(HSD17B12) | NM_016142 | 102 | 99.8 | 0.04238 | 0.899 | NM_001014273 | 3.32 | 15.01 |
| HSD17B13 | hydroxysteroid 17-beta dehydrogenase 13(HSD17B13) | NM_178135 | 508.7 | 618.2 | 0.04238 | 0.899 | | | |
| ZNF17 | zinc finger protein 17(ZNF17) | NM_006959 | 103.4 | 94.5 | 0.04251 | 0.914 | | | |
| RCCD1 | RCC1 domain containing 1(RCCD1) | NM_033544 | 92.8 | 101.2 | 0.04252 | 1.135 | | | |
| MYO18A | myosin XVIIIA(MYO18A) | NM_078471 | 162.4 | 161.7 | 0.04255 | 1.205 | | | |

| | | | | | | |
|---|---|---|---|---|---|---|
| LPCAT3 | lysophosphatidylcholine acyltransferase 3(LPCAT3) | NM_005768 | 175.9 | 178.8 | 0.04258 | 1.344 |
| jund | JunD proto-oncogene, AP-1 transcription factor subunit(JUND) | NM_005354 | 401 | 464.7 | 0.04266 | 0.375 |
| ZNF621 | zinc finger protein 621(ZNF621) | NM_001098414 | 116.4 | 117.1 | 0.04272 | 1.186 |
| SGK3 | serum/glucocorticoid regulated kinase family member 3(SGK3) | NM_170709 | 125.1 | 115.9 | 0.04284 | 1.314 |
| SBSPON | somatomedin B and thrombospondin type 1 domain containing(SBSPON) | NM_153225 | 104.3 | 120.9 | 0.04286 | 1.18 |
| RAD17 | RAD17 checkpoint clamp loader component(RAD17) | NM_133340 | 138.7 | 125.1 | 0.04293 | 0.876 |
| RAD17 | RAD17 checkpoint clamp loader component(RAD17) | NM_133340 | 153.6 | 129.1 | 0.04293 | 0.876 |
| SLC4A1 | solute carrier family 4 member 1 (Diego blood group)(SLC4A1) | NM_000342 | 138.3 | 212.5 | 0.04306 | 1.716 |
| | | XR_016385 | 102.5 | 103.5 | 0.0431 | 0.612 |
| | | XM_926345 | 97 | 102.8 | 0.04316 | 1.121 |
| | | XM_496783 | 97.3 | 85.6 | 0.0432 | 0.597 |

| | | | | | | |
|---|---|---|---|---|---|---|
| CEPT1 | choline/ethanolamine phosphotransferase 1(CEPT1) | NM_006090 | 114.9 | 117.1 | 0.04325 | 1.127 |
| | | XM_930777 | 126.2 | 108.9 | 0.04327 | 1.189 |
| ENTPD6 | ectonucleoside triphosphate diphosphohydrolase 6 (putative)(ENTPD6) | NM_001247 | 90.4 | 87.2 | 0.04337 | 1.117 |
| ENTPD7 | ectonucleoside triphosphate diphosphohydrolase 7(ENTPD7) | NM_020354 | 100.4 | 101.5 | 0.04337 | 1.117 |
| | | XM_937442 | 119.2 | 128 | 0.04341 | 1.186 |
| | | XM_929297 | 103.1 | 101.2 | 0.04344 | 0.914 |
| | | AW135045 | 118.2 | 118.4 | 0.0435 | 1.096 |
| VWA8 | von Willebrand factor A domain containing 8(VWA8) | NM_015058 | 97.9 | 93.1 | 0.04351 | 0.874 |
| NAIF1 | nuclear apoptosis inducing factor 1(NAIF1) | NM_197956 | 105.6 | 107.1 | 0.04358 | 1.143 |
| NAIF1 | nuclear apoptosis inducing factor 1(NAIF1) | NM_197956 | 132.5 | 121.1 | 0.04358 | 1.143 |

| | | | | | | |
|---|---|---|---|---|---|---|
| WHAMM | WAS protein homolog associated with actin, golgi membranes and microtubules(WHAMM) | NM_001080435 | 145.8 | 127 | 0.0436 | 1.198 |
| MAGEE1 | MAGE family member E1(MAGEE1) | NM_020932 | 98.4 | 93.2 | 0.04365 | 0.889 |
| MAGEF1 | MAGE family member F1(MAGEF1) | NM_022149 | 108.8 | 104.2 | 0.04365 | 0.889 |
| | | XM_928267 | 103.8 | 109.4 | 0.04378 | 1.162 |
| C11orf58 | chromosome 11 open reading frame 58(C11orf58) | NM_001142705 | 142.5 | 149.8 | 0.04381 | 1.168 |
| | | AW082312 | 95.9 | 100.1 | 0.04392 | 1.109 |
| MSMO1 | methylsterol monooxygenase 1(MSMO1) | NM_001017369 | 96 | 95.8 | 0.04398 | 0.922 |
| MSMO1 | methylsterol monooxygenase 1(MSMO1) | NM_001017369 | 160.9 | 160.5 | 0.04398 | 0.922 |
| YARS | tyrosyl-tRNA synthetase(YARS) | NM_003680 | 118.4 | 121.9 | 0.04401 | 1.132 |
| KIAA0368 | KIAA0368(KIAA0368) | NM_001080398 | 88.2 | 95.4 | 0.04438 | 1.149 |
| | | XM_001128895 | 111.7 | 109.7 | 0.04444 | 1.163 |
| CLYBL | citrate lyase beta like(CLYBL) | NM_206808 | 106.2 | 108.2 | 0.04467 | 1.17 |
| | | XR_040448 | 93.4 | 84 | 0.04468 | 0.868 |

| | | | | | | |
|---|---|---|---|---|---|---|
| CNNM3 | cyclin and CBS domain divalent metal cation transport mediator 3(CNNM3) | NM_017623 | 98.8 | 100.8 | 0.0447 | 1.101 |
| STK40 | serine/threonine kinase 40(STK40) | NM_032017 | 90.8 | 87.6 | 0.04476 | 1.385 |
| STK40 | serine/threonine kinase 40(STK40) | NM_032017 | 274 | 233.2 | 0.04476 | 1.385 |
| | | XM_211040 | 100.9 | 95.8 | 0.04476 | 0.887 |
| SFTPA1 | surfactant protein A1(SFTPA1) | NM_005411 | 86.4 | 94.9 | 0.04479 | 1.342 |
| SOX4 | SRY-box 4(SOX4) | NM_003107 | 180.4 | 215.5 | 0.04487 | 0.608 |
| MBTPS1 | membrane bound transcription factor peptidase, site 1(MBTPS1) | NM_003791 | 156.6 | 150.5 | 0.04489 | 1.18 |
| MBTPS1 | membrane bound transcription factor peptidase, site 1(MBTPS1) | NM_003791 | 195.8 | 179.5 | 0.04489 | 1.18 |
| | | XM_931458 | 97.7 | 95.7 | 0.04495 | 1.154 |
| | | XM_927506 | 90.8 | 94.8 | 0.045 | 1.16 |
| NUTF2 | nuclear transport factor 2(NUTF2) | NM_005796 | 137.9 | 126.8 | 0.04505 | 1.22 |
| ERGIC3 | ERGIC and golgi 3(ERGIC3) | NM_198398 | 196.6 | 157 | 0.04506 | 1.257 |

| | | | | | | |
|---|---|---|---|---|---|---|
| CPT1A | carnitine palmitoyltransferase 1A(CPT1A) | NM_001031847 | 94.2 | 85.4 | 0.04518 | 0.881 |
| PEX7 | peroxisomal biogenesis factor 7(PEX7) | NM_000288 | 201.8 | 164 | 0.04519 | 0.822 |
| CTNNB1 | catenin beta 1(CTNNB1) | NM_001098209 | 96.5 | 96.4 | 0.0452 | 0.697 |
| CTNNB1 | catenin beta 1(CTNNB1) | NM_001098209 | 148.2 | 145.9 | 0.0452 | 0.697 |
| CTNNB1 | catenin beta 1(CTNNB1) | NM_001098209 | 154 | 164.5 | 0.0452 | 0.697 |
| | | XR_039466 | 96.4 | 98.5 | 0.04521 | 1.169 |
| MKRN2 | makorin ring finger protein 2(MKRN2) | NM_014160 | 223.4 | 182.1 | 0.04524 | 0.778 |
| RP1L1 | retinitis pigmentosa 1 like 1(RP1L1) | NM_178857 | 97.9 | 91.9 | 0.04545 | 0.873 |
| | | XR_039516 | 92.6 | 93.2 | 0.04545 | 1.264 |
| | | XR_039516 | 188.3 | 171.5 | 0.04545 | 1.264 |
| KCNJ15 | potassium voltage-gated channel subfamily J member 15(KCNJ15) | NM_170736 | 99.7 | 90.2 | 0.04551 | 0.91 |
| | | XM_931506 | 101 | 97.5 | 0.04557 | 0.91 |
| | | CN283269 | 110.1 | 101.8 | 0.04559 | 0.921 |

| | | | | | | |
|---|---|---|---|---|---|---|
| MAPK3 | mitogen-activated protein kinase 3(MAPK3) | NM_001040056 | 96.9 | 91.8 | 0.04562 | 0.908 |
| TCL1B | T-cell leukemia/lymphoma 1B(TCL1B) | NM_004918 | 138.6 | 108.5 | 0.04562 | 1.148 |
| THG1L | tRNA-histidine guanylyltransferase 1 like(THG1L) | NM_017872 | 103.1 | 99.4 | 0.04565 | 1.105 |
| TPPP3 | tubulin polymerization promoting protein family member 3(TPPP3) | NM_015964 | 98.7 | 89.8 | 0.04577 | 0.89 |
| ANKRD33 | ankyrin repeat domain 33(ANKRD33) | NM_182608 | 99 | 97.6 | 0.04587 | 0.781 |
| MOB4 | MOB family member 4, phocein(MOB4) | NM_001100819 | 105.6 | 104 | 0.04595 | 0.825 |
| WASF3 | WAS protein family member 3(WASF3) | NM_006646 | 108.2 | 124.8 | 0.04602 | 1.202 |
| CTDSP1 | CTD small phosphatase 1(CTDSP1) | NM_021198 | 90.5 | 93.2 | 0.04617 | 1.257 |
| CTDSP2 | CTD small phosphatase 2(CTDSP2) | NM_005730 | 287.9 | 256.9 | 0.04617 | 1.257 |
| SLAMF6 | SLAM family member 6(SLAMF6) | NM_052931 | 92.5 | 86.7 | 0.0462 | 0.841 |
| SLAMF6 | SLAM family member 6(SLAMF6) | NM_052931 | 120.4 | 96.9 | 0.0462 | 0.841 |
| tcf20 | transcription factor 20(TCF20) | NM_181492 | 101 | 100.8 | 0.04623 | 1.105 |

| | | | | | | |
|---|---|---|---|---|---|---|
| tcf20 | transcription factor 20(TCF20) | NM_181492 | 304.4 | 258.7 | 0.04623 | 1.105 |
| | | BM679134 | 114.5 | 109 | 0.04623 | 1.144 |
| TNFRSF11B | TNF receptor superfamily member 11b(TNFRSF11B) | NM_002546 | 176.2 | 176 | 0.04628 | 0.539 |
| CHST4 | carbohydrate sulfotransferase 4(CHST4) | NM_005769 | 95.6 | 92.2 | 0.04639 | 0.895 |
| FAM162B | family with sequence similarity 162 member B(FAM162B) | NM_001085480 | 100.7 | 98.7 | 0.04639 | 1.108 |
| STK25 | serine/threonine kinase 25(STK25) | NM_006374 | 350.5 | 355.1 | 0.04641 | 1.298 |
| | | XM_926412 | 101 | 101 | 0.04651 | 1.127 |
| | | XM_001722392 | 100.1 | 91.8 | 0.04656 | 0.881 |
| RASGRP4 | RAS guanyl releasing protein 4(RASGRP4) | NM_052949 | 108.4 | 114.6 | 0.04659 | 1.118 |
| BCKDHB | branched chain keto acid dehydrogenase E1 subunit beta(BCKDHB) | NM_000056 | 421.2 | 517.8 | 0.04661 | 1.682 |
| PRDM15 | PR/SET domain 15(PRDM15) | NM_001040424 | 98 | 90.2 | 0.04662 | 0.923 |
| ZNF446 | zinc finger protein 446(ZNF446) | NM_017908 | 159.2 | 134.1 | 0.04674 | 0.828 |
| ZFP36L1 | ZFP36 ring finger protein like 1(ZFP36L1) | NM_004926 | 943 | 789.9 | 0.04683 | 0.684 |

| | | | | | | |
|---|---|---|---|---|---|---|
| KIF2C | kinesin family member 2C(KIF2C) | NM_006845 | 104.9 | 104.8 | 0.04683 | 1.123 |
| | | XM_926130 | 141.1 | 123 | 0.04685 | 1.313 |
| | | BQ772469 | 115.1 | 95.4 | 0.04688 | 0.892 |
| MED11 | mediator complex subunit 11(MED11) | NM_001001683 | 92 | 100.3 | 0.04699 | 1.105 |
| TMEM150B | transmembrane protein 150B(TMEM150B) | NM_001085488 | 108.7 | 112.9 | 0.04724 | 1.115 |
| DENND2C | DENN domain containing 2C(DENND2C) | NM_198459 | 218.2 | 181.2 | 0.04726 | 1.251 |
| MAMSTR | MEF2 activating motif and SAP domain containing transcriptional regulator(MAMSTR) | NM_182574 | 116.7 | 109.4 | 0.04729 | 0.833 |
| | | XR_015259 | 98.8 | 93.5 | 0.04738 | 0.697 |
| | | XR_017249 | 93.3 | 88.2 | 0.04739 | 0.829 |
| | | XM_930508 | 91.3 | 84 | 0.0474 | 0.773 |
| MKNK2 | MAP kinase interacting serine/threonine kinase 2(MKNK2) | NM_017572 | 579.1 | 427.2 | 0.04743 | 1.568 |
| PBRM1 | polybromo 1(PBRM1) | NM_181042 | 103.3 | 101 | 0.0475 | 0.877 |
| PBRM1 | polybromo 1(PBRM1) | NM_181042 | 113.7 | 94.4 | 0.0475 | 0.877 |

| | | | | | | | | | |
|---|---|---|---|---|---|---|---|---|---|
| c21orf33 | chromosome 21 open reading frame 33(C21orf33) | NM_004649 | 101.2 | 96.3 | 0.04761 | 0.886 | | | |
| GHSR | growth hormone secretagogue receptor(GHSR) | NM_198407 | 92.6 | 97.5 | 0.04761 | 1.14 | | | |
| | | XM_001721744 | 122 | 128 | 0.04761 | 1.226 | NM_145089 | 67.02 | 12.33 |
| KLHL9 | kelch like family member 9(KLHL9) | NM_018847 | 115.2 | 103 | 0.04767 | 1.121 | | | |
| QKI | QKI, KH domain containing RNA binding(QKI) | NM_006775 | 98.2 | 93 | 0.04774 | 0.895 | NM_013157 | 76.74 | 0.47 |
| QKI | QKI, KH domain containing RNA binding(QKI) | NM_006775 | 102.4 | 106.3 | 0.04774 | 0.895 | NM_013157 | 76.74 | 0.47 |
| | | XR_016047 | 102.3 | 97 | 0.04774 | 1.138 | NM_013157 | 76.74 | 0.47 |
| TRAPPC3 | trafficking protein particle complex 3(TRAPPC3) | NM_014408 | 349 | 308.8 | 0.04783 | 1.167 | NM_013157 | 76.74 | 0.47 |
| IGFBP2 | insulin like growth factor binding protein 2(IGFBP2) | NM_000597 | 88.8 | 101.1 | 0.04793 | 1.21 | | | |
| COL18A1 | collagen type XVIII alpha 1 chain(COL18A1) | NM_130445 | 233.3 | 218 | 0.04794 | 0.769 | | | |
| | | XM_925903 | 100.7 | 98.2 | 0.04795 | 0.875 | | | |
| | | XM_925903 | 103.1 | 90.2 | 0.04795 | 0.875 | | | |

| | | | | | | |
|---|---|---|---|---|---|---|
| LYNX1 | Ly6/neurotoxin 1(LYNX1) | NM_177458 | 100.8 | 90.1 | 0.04813 | 0.905 |
| ZC3H18-AS1 | ZC3H18 antisense RNA 1 (head to head)(ZC3H18-AS1) | NM_001001682 | 101.8 | 94.3 | 0.04814 | 0.915 |
| REV3L | REV3 like, DNA directed polymerase zeta catalytic subunit(REV3L) | NM_002912 | 100.2 | 94.6 | 0.04817 | 0.836 |
| ENPP2 | ectonucleotide pyrophosphatase/phosphodiesterase 2(ENPP2) | NM_001040092 | 116 | 107.4 | 0.04824 | 1.141 |
| | | BQ771772 | 108.4 | 106.7 | 0.04828 | 1.121 |
| | | XR_039891 | 101.8 | 99.2 | 0.04833 | 1.078 |
| ZNF80 | zinc finger protein 80(ZNF80) | NM_007136 | 100.9 | 102.2 | 0.04834 | 1.112 |
| EFEMP1 | EGF containing fibulin like extracellular matrix protein 1(EFEMP1) | NM_001039348 | 98.8 | 108 | 0.0484 | 1.098 |
| MPHOSPH6 | M-phase phosphoprotein 6(MPHOSPH6) | NM_005792 | 120.3 | 140.9 | 0.04844 | 1.132 |
| ZADH2 | zinc binding alcohol dehydrogenase domain containing 2(ZADH2) | NM_175907 | 98.7 | 100.6 | 0.0485 | 1.095 |
| RTN4RL1 | reticulon 4 receptor like 1(RTN4RL1) | NM_178568 | 103.9 | 105 | 0.04862 | 1.097 |

| | | | | | | |
|---|---|---|---|---|---|---|
| SHANK2 | SH3 and multiple ankyrin repeat domains 2(SHANK2) | NM_012309 | 165.9 | 212.4 | 0.04863 | 1.469 |
| | | XM_001726649 | 108.3 | 101.4 | 0.04868 | 0.858 |
| | | XR_015716 | 99.9 | 95.9 | 0.04877 | 0.869 |
| AASDH | aminoadipate-semialdehyde dehydrogenase(AASDH) | NM_181806 | 148.8 | 131.2 | 0.04877 | 0.839 |
| SNX22 | sorting nexin 22(SNX22) | NM_024798 | 101.7 | 97.9 | 0.0488 | 0.83 |
| C4BPA | complement component 4 binding protein alpha(C4BPA) | NM_000715 | 109.4 | 102.1 | 0.04887 | 0.887 |
| RAD21 | RAD21 cohesin complex component(RAD21) | NM_006265 | 296.7 | 266.7 | 0.0489 | 1.333 |
| RAD21 | RAD21 cohesin complex component(RAD21) | NM_006265 | 498 | 390.6 | 0.0489 | 1.333 |
| OPN5 | opsin 5(OPN5) | NM_001030051 | 120.6 | 108.4 | 0.04901 | 0.804 |
| lsm11 | LSM11, U7 small nuclear RNA associated(LSM11) | NM_173491 | 371 | 312.1 | 0.04901 | 1.17 |
| | | XR_041956 | 110.6 | 102.5 | 0.04909 | 0.852 |

| | | | | | | |
|---|---|---|---|---|---|---|
| COX16 | COX16, cytochrome c oxidase assembly homolog(COX16) | NM_016468 | 110 | 105.3 | 0.04912 | 1.149 |
| Cox15 | COX15, cytochrome c oxidase assembly homolog(COX15) | NM_078470 | 100.1 | 103.9 | 0.04912 | 1.149 |
| FRMD1 | FERM domain containing 1(FRMD1) | NM_024919 | 109.2 | 100.4 | 0.04917 | 0.917 |
| | | XM_001725183 | 3889.8 | 2672 | 0.0492 | 0.763 |
| LOC100996549 | uncharacterized LOC100996549(LOC100996549) | AK094576 | 96.1 | 101.7 | 0.04925 | 1.137 |
| SIGLEC7 | sialic acid binding Ig like lectin 7(SIGLEC7) | NM_016543 | 98 | 96 | 0.04931 | 0.896 |
| scaf8 | SR-related CTD associated factor 8(SCAF8) | NM_014892 | 143.9 | 129.2 | 0.04939 | 0.805 |
| | | XM_925876 | 100.2 | 105.2 | 0.0494 | 1.148 |
| KPNA3 | karyopherin subunit alpha 3(KPNA3) | NM_002267 | 104.1 | 104.5 | 0.04953 | 0.871 |
| YPEL2 | yippee like 2(YPEL2) | NM_001005404 | 114.5 | 106.3 | 0.04958 | 0.869 |
| C9orf72 | chromosome 9 open reading frame 72(C9orf72) | NM_018325 | 135.7 | 142 | 0.0496 | 0.848 |
| HERC2 | HECT and RLD domain containing E3 ubiquitin protein ligase 2(HERC2) | NM_004667 | 110.6 | 108.6 | 0.0496 | 1.167 |

| | | | | | | | | | |
|---|---|---|---|---|---|---|---|---|---|
| | | XM_932019 | 98 | 107.3 | 0.04961 | 1.114 | | | |
| | | XM_941746 | 94.7 | 95.7 | 0.04968 | 0.846 | | | |
| c16orf87 | chromosome 16 open reading frame 87(C16orf87) | NM_001001436 | 226.8 | 267.9 | 0.04969 | 1.162 | | | |
| CELSR1 | cadherin EGF LAG seven-pass G-type receptor 1(CELSR1) | NM_014246 | 102.5 | 91.4 | 0.0497 | 0.86 | NM_134394 | 3.46 | 15.03 |
| ZNF607 | zinc finger protein 607(ZNF607) | NM_032689 | 92.3 | 99.2 | 0.04971 | 1.094 | | | |
| ZNF607 | zinc finger protein 607(ZNF607) | NM_032689 | 107 | 98.8 | 0.04971 | 1.094 | | | |
| CPE | carboxypeptidase E(CPE) | NM_001873 | 89.4 | 95.9 | 0.04971 | 1.137 | | | |
| XRN1 | 5'-3' exoribonuclease 1(XRN1) | NM_001042604 | 120.6 | 103.6 | 0.04978 | 0.872 | | | |
| KIAA0907 | KIAA0907(KIAA0907) | NM_014949 | 98.4 | 89.4 | 0.0499 | 0.845 | | | |
| | | XM_931879 | 206.4 | 152.3 | 0.04994 | 0.525 | | | |
| rab11fip5 | RAB11 family interacting protein 5(RAB11FIP5) | NM_015470 | 506.4 | 534.8 | 0.04995 | 1.322 | | | |
| PKNOX1 | PBX/knotted 1 homeobox 1(PKNOX1) | NM_004571 | 91.9 | 86.3 | 0.04996 | 0.908 | | | |
| PKNOX1 | PBX/knotted 1 homeobox 1(PKNOX1) | NM_004571 | 136.7 | 127.3 | 0.04996 | 0.908 | | | |

| | | | | | | |
|---|---|---|---|---|---|---|
| OR2W3 | olfactory receptor family 2 subfamily W member 3(OR2W3) | NM_001001957 | 108.8 | 92.8 | 0.04997 | 0.871 |
| | | NM_001004349 | 103.2 | 106.6 | 0.04999 | 1.142 |
| RAB3A | RAB3A, member RAS oncogene family(RAB3A) | NM_002866 | 93.3 | 99.9 | 0.04999 | 1.127 |
